# The forkhead transcription factor FKH-7/FOXP acts in chemosensory neurons to regulate developmental decision-making

**DOI:** 10.1101/2025.02.17.638733

**Authors:** Cynthia M. Chai, Seth R. Taylor, Carsten H. Tischbirek, Wan-Rong Wong, Long Cai, David M. Miller, Paul W. Sternberg

## Abstract

Autism is a complex neurodevelopmental disorder with many associated genetic factors, including the forkhead transcription factor FOXP1. Although FOXP1’s neuronal role is well-studied, the specific molecular consequences of different FOXP1 pathogenic variants in physiologically-relevant contexts are unknown. Here we ascribe the first function to Caenorhabditis elegans FKH-7/FOXP, which acts in two chemosensory neuron classes to promote the larval decision to enter the alternative, developmentally-arrested dauer life stage. We demonstrate that human FOXP1 can functionally substitute for C. elegans FKH-7 in these neurons and that engineering analogous FOXP1 hypomorphic missense mutations in the endogenous fkh-7 locus also impairs developmental decision-making. In a fkh-7/FOXP1 missense variant, single-cell transcriptomics identifies downregulated expression of autism-associated kcnl-2/KCNN2 calcium-activated potassium channel in a serotonergic sensory neuron. Our findings establish a novel framework linking two evolutionarily-conserved autism-associated genes for deeper characterization of variant-specific molecular pathology at single neuron resolution in the context of a developmental decision-making paradigm.

## INTRODUCTION

The evolutionarily conserved forkhead box (FOX) family of transcription factors share a characteristic, highly conserved winged-helix DNA-binding domain, and have diverse roles in human development and adulthood^1–3^. Dysfunction or dysregulation of the four FOXP paralogs in vertebrates have been implicated in debilitating conditions including immune dysregulation, cancer progression, and cognitive developmental disorders^2,3^. In particular, mutations in human *FOXP1* and *FOXP2* are associated with neurological disorders such as speech and language acquisition deficits, intellectual disability, and autism spectrum disorder (ASD)^4–6^. Whole-brain *Foxp1* conditional knockout mice exhibit reduced striatal area as well as increased frequency of repetitive behaviors and impaired social interactions while heterozygous *Foxp1* mice have altered excitability of striatal neurons and defective ultrasonic volcalization^7,8^. In juvenile zebra finches, *FoxP2* knockdown in Area X of the basal ganglia results in inaccurate imitation of tutor song, prevents context-dependent modulation of singing-related activity in neural components of song control circuitry, and significantly reduced levels of DARPP-32, a key regulator of dopaminergic signaling, in Area X^9,10^. *Drosophila* fruit flies have a sole *FoxP* ortholog, which is expressed in the mushroom bodies of the fly brain, and *FoxP* mutants display prolonged reaction times during difficult perceptual decision-making tasks^11^. Despite the array of neurological functions regulated by FOXP members in disparate organisms, the function of FKH-7, their sole ortholog in the nematode *Caenorhabditis elegans*, has never been explored.

The strong upregulation of *fkh-7* expression during the dauer developmental life stages (diapause) relative to stages along the reproductive growth trajectory suggests that *fkh-7* may mediate dauer formation^12^. When challenged with adverse environmental conditions, *C. elegans* L1 stage larvae can develop into the alternative growth-arrested dauer larval morph by proceeding through a preparatory pre-dauer L2d stage^13,14^ (Figure 1A). Development into the stress-resistant dauer stage entails whole-animal tissue remodeling that confers stage-specific adaptations such as a thickened cuticle, increased density of intestinal storage granules, synaptic rewiring of the nervous system, and a corresponding dauer-specific repertoire of behaviors^13–18^. The larval decision to enter diapause is promoted by low food availability, warmer environmental temperatures, and high secreted pheromone concentrations, which could serve as a proxy for greater conspecific density and competition in the local area^14^. The developmental decision is under sensory control^19^, and the roles of individual sensory neuron classes have been assigned^20–22^. Throughout this study, a pheromone-based dauer formation assay was utilized to induce and quantify outcomes of the binary developmental decision^23^ (see Methods).

**Figure 1.**
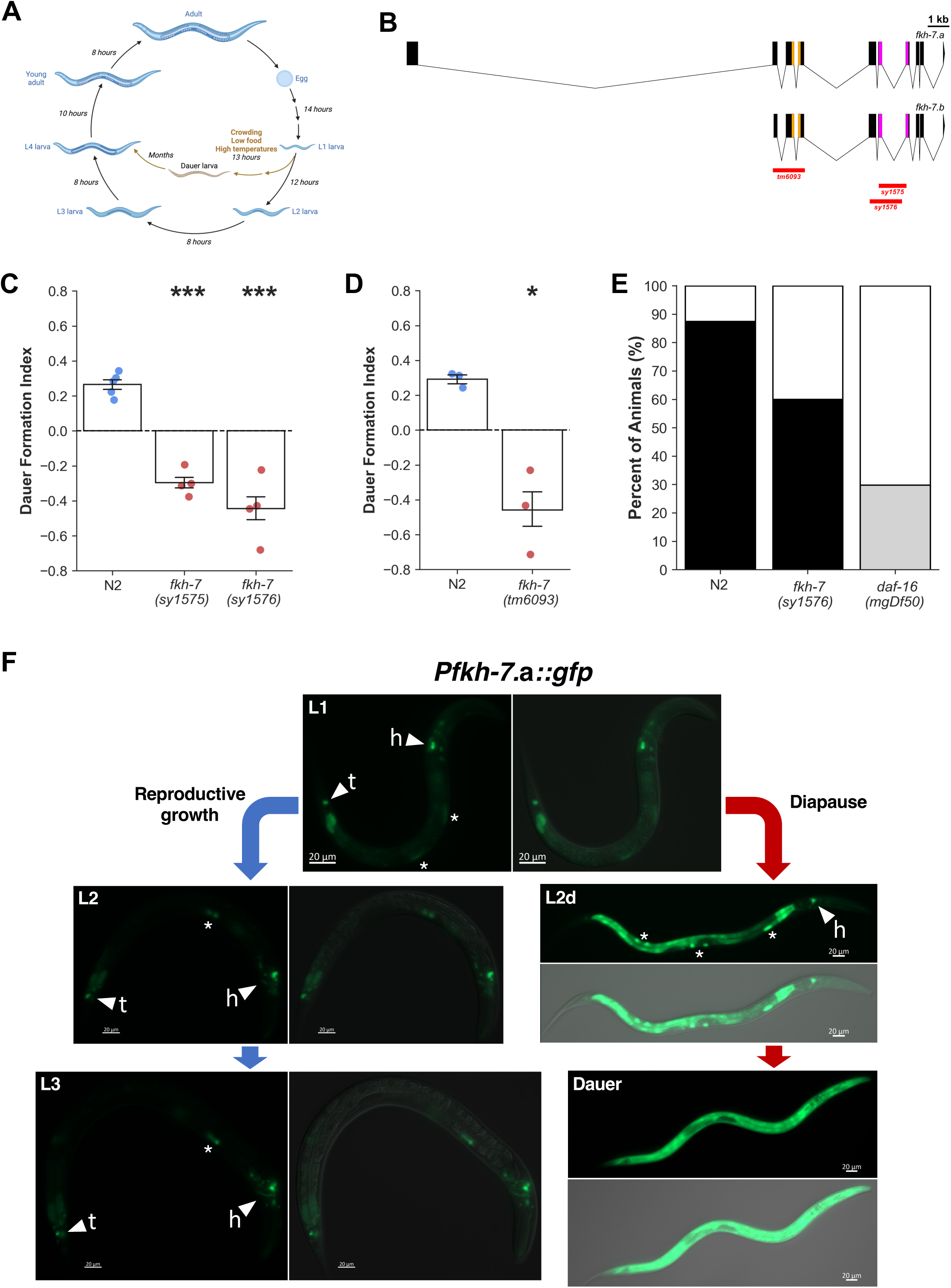
*C. elegans fkh-7* loss-of-function mutants are defective in dauer formation under pheromone dauer-inducing conditions. (A) Entry into the *C. elegans* alternative stress-resistant diapause developmental trajectory is induced by adverse environmental growth conditions. (B) Schematic of *fkh-7* gene model showing overlap in *fkh-7.*a/b transcript sequences, functional domains (coiled-coil domain in yellow and DNA-binding domain in purple), locations of the *sy1575* and *sy1576* CRISPR deletions impacting the DNA-binding domain (red), and location of the *tm6093* deletion impacting the coiled-coil domain (red). (C) Dauer formation assays for 1.26 kb deletion *fkh-7*(*sy1575*) and 1.5 kb deletion *fkh-7(sy1576*) mutants. N=4-5 population assays, 61-72 animals per assay. Data represented as mean ± SEM, one-way ANOVA followed by Dunnett’s post-hoc test, ***p<0.001. (D) Dauer formation assays for *fkh-7*(*tm6093*) mutant. N=3 population assays, 60-66 animals per assay. Data represented as mean ± SEM, Welch’s unpaired t test, *p<0.05. (E) Percentages of reproductive larvae (white), partial dauers (gray), and full dauers (black) for *fkh-7(sy1576)* and *daf-16(mgDf50)* mutants from dauer formation assays using high crude pheromone concentrations that induce >80% N2 wild-type dauer formation. N=4 population assays, 57-73 animals per assay. (F) P*fkh-7*.a::*gfp* expression during different larval stages grown under reproductive (blue arrows) and pheromone-induced (red arrows) dauer formation conditions. White arrowheads labelled ‘h’ indicate head neurons expressing fluorescent reporter. White arrowheads labelled ‘t’ indicate tail neurons expressing fluorescent reporter. White asterisks indicate fluorescence bleed-through in coelomocyte scavenger cells from the P*ofm-1*::*rfp* co-injection marker.

## RESULTS

*C. elegans fkh-7* is predicted to encode twelve protein-coding transcripts with a DNA-binding domain. As there were no available genetic mutants that disrupted all transcripts, we used a CRISPR-mediated non-homologous end joining repair strategy to generate two new *fkh-7* loss-of-function alleles, *sy1575* and *sy1576*. The *sy1575* allele harbors a 1.26 kb in-frame deletion nested within the forkhead domain while the *sy1576* allele harbors a 1.5 kb in-frame deletion that removes the first half of the forkhead domain together with most of the preceding exon (Figure 1B). Animals homozygous for either *fkh-7* deletion allele exhibit a significant reduction in dauer larvae formation under pheromone dauer-inducing conditions, with the *sy1576* allele displaying a stronger phenotype (Figure 1C). The *fkh-7*(*tm6093*) mutant harbors an out-of-frame deletion in the coiled-coil domain-encoding region that introduces a premature stop codon truncating eight of the longest *fkh-7* transcripts (Figure 1B). *fkh-7*(*tm6093*) homozygous mutants also exhibit a significant reduction in dauer larvae formation similar in magnitude to the *sy1576* deletion, suggesting that loss of the longer *fkh-7* transcripts may account for the *fkh-7*(*sy1576*) phenotype (Figure 1D). The genetic and tissue-specific roles of another *C. elegans* forkhead transcription factor, DAF-16/FOXO, in dauer development regulation are well-studied^24–27^. When grown in the presence of high pheromone concentration that induces almost all N2 wild-type larvae to enter dauer development, *fkh-7(sy1576)* mutants formed 27% less dauer larvae compared to the wild-type control while 30% of *daf-16(mgDf50)* null mutants developed into partial dauers with no full dauers formed (Figure 1E). The formation of *daf-16(mgDf50)* mutant partial dauers, which are developmentally arrested larvae that are less radially constricted and have lighter colored bodies relative to full dauers, under pheromone dauer-inducing conditions is consistent with previous studies^25^. During the L1 stage, expression of a *fkh-7*.a transcriptional GFP reporter (P*fkh-7*.a*::gfp*) is localized to a few neurons in the head and tail as well as in the intestine (Figure 1F). Beyond the L1 stage, *fkh-7*.a expression patterns diverge as larvae develop under different growth conditions. Under reproductive growth conditions, P*fkh-7*.a*::gfp* expression in L2 and L3 larvae resembles the L1 stage except with dimmer intestine expression (Figure 1F). Under pheromone dauer-inducing conditions, however, P*fkh-7*.a*::gfp* expression in the intestine increases in intensity during the pre-dauer L2d stage and becomes widespread in several tissues during the dauer stage (Figure 1F).

To determine the *fkh-7*.a longest transcript’s site-of-action, we used tissue-specific promoters to selectively drive *fkh-7*.a cDNA expression in the *fkh-7*(*sy1576*) mutant. *fkh-7*.a cDNA expression in either the nervous system or intestine rescued the *fkh-7(sy1576)* dauer formation defective phenotype, but not expression in the body wall muscle, pharyngeal muscle, or hypodermis (Figure 2A). In the P*fkh-7*.a*::gfp* reporter strain, the processes of GFP-expressing neurons extend to the tip of the larva’s nose suggesting that these are amphid sensory neurons. We leveraged the fact that promoters of the cyclic nucleotide-gated channel subunit-encoding *tax-4* and the vanilloid subfamily of transient receptor potential channel protein-encoding *ocr-2* each drive expression in complementary subsets of *C. elegans* amphid sensory neurons to dissect *fkh-7*.a’s site-of-action in the nervous system^28,29^. Expression of *fkh-7*.a cDNA driven by the *ocr-2* promoter rescued the dauer formation defective phenotype of the *fkh-7(sy1576)* mutant whereas cDNA expression in the sensory neuron subset labelled by the *tax-4* promoter did not (Figure 2B). The 2.5 kb *ocr-2* promoter fragment used to drive *fkh-7*.a cDNA expression is expressed in three amphid sensory neuron classes in the head (ADL, ADF, and ASH), as determined by P*ocr-2::gfp* colocalization with DiI-stained ADL and ASH neurons as well as an ADF-specific P*srh-142::mCherry-H2B* marker^29^ (Figure S1A,B). P*ocr-2::gfp* is also expressed in two neurons in the tail, which are likely to be the PHA and PHB phasmid sensory neurons as previously reported^29^ (Figure S1C).

**Figure 2.**
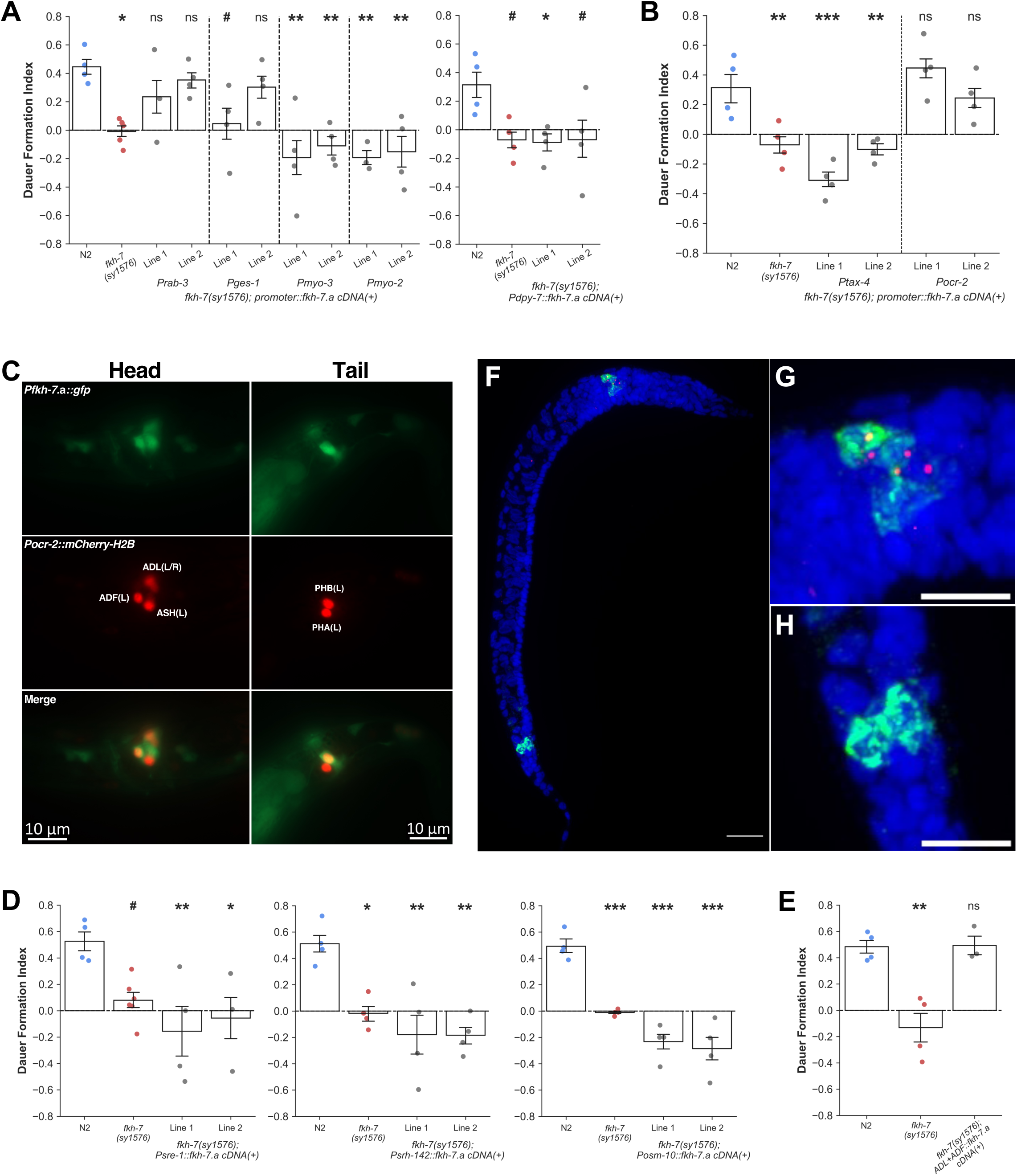
The longest transcript *fkh-7*.a acts in two chemosensory neuron classes to promote larval entry into developmental diapause. (A) Dauer formation assays for tissue-specific *fkh-7*.a cDNA expression in *fkh-7*(*sy1576*) mutants. N=3-5 population assays, 83-130 animals per assay. Data represented as mean ± SEM, one-way ANOVA followed by Dunnett’s post-hoc test, ns=no significance, ^#^p<0.1, *p<0.05, **p<0.01. (B) Dauer formation assays for *fkh-7*.a cDNA expression in *ocr-2* promoter and *tax-4* promoter sensory neuron subsets in *fkh-7*(*sy1576*) mutants. For transgenic rescue lines, only the subset of animals bearing the extrachromosomal array were used to calculate DFI here. For DFIs calculated using all animals, see Data S1. N=4 population assays, 100-120 animals per assay. Data represented as mean ± SEM, one-way ANOVA followed by Dunnett’s post-hoc test, ns=no significance, **p<0.01, ***p<0.001. (C) P*fkh-7.a::gfp* and P*ocr-2*::*mCherry-H2B* colocalization in an L1 stage larva. Head (left) and tail (right) regions shown. (D) Dauer formation assays for ADL (left), ADF (middle), or ASH+PHA+PHB (right) -specific *fkh-7*.a cDNA expression in *fkh-7*(*sy1576*) mutants. N=3-6 population assays, 84-130 animals per assay. Data represented as mean ± SEM, one-way ANOVA followed by Dunnett’s post-hoc test, ^#^p<0.1, *p<0.05, **p<0.01, ***p<0.001. (E) Dauer formation assays for ADL+ADF-specific *fkh-7*.a cDNA expression in *fkh-7*(*sy1576*) mutants. N=3-4 population assays, 93-143 animals per assay. For transgenic rescue line, only the subset of animals bearing the extrachromosomal array were used to calculate DFI here. For DFIs calculated using all animals, see Data S1. Data represented as mean ± SEM, one-way ANOVA followed by Dunnett’s post-hoc test, ns=no significance, **p<0.01. (F to H) Representative images of first exon of *fkh-7*.a mRNA expression (pink) and P*ocr-2::gfp* mRNA expression (green) detected by single-molecule fluorescence *in situ* hybridization (smFISH) in a pheromone-induced pre-dauer L2d stage larva. Nuclei stained with DAPI in blue. (F) Expression in whole animal. Scale bar represents 20 μm. (G) Expression in head region. Scale bar represents 10 μm. (H) Expression in tail region. Scale bar represents 10 μm.

In L1 larvae, expression of *fkh-7*.a and *ocr-2* fluorescent reporter constructs colocalized in the ADL (head), ADF (head), and putative PHB (tail) sensory neuron classes suggesting that *fkh-7*.a functions in either all or a subset of these neurons in the *C. elegans* larval nervous system (Figure 2C). Neither ADL-specific, ADF-specific nor ASH+PHA+PHB-specific cDNA expression was sufficient to rescue the *fkh-7*(*sy1576*) dauer formation defective phenotype (Figure 2D). However, simultaneous expression of *fkh-7*.a cDNA in both the ADL and ADF amphid chemosensory neuron classes was sufficient to rescue the *fkh-7*(*sy1576*) mutant phenotype (Figure 2E). Since only *fkh-7*.a cDNA expression in both the ADL and ADF neurons fully rescues the *fkh-7(sy1576)* dauer formation phenotype, the interactive functions of both chemosensory neurons may be normally required to modulate dauer formation. As a large 16.5 kb-long intron resides between the first and second exons of the *fkh-7*.a transcript, it is possible that regulatory elements in this intronic region that are absent in the P*fkh-7*.a*::gfp* reporter construct examined above may also shape *fkh-7*.a’s expression pattern and dynamics. To directly determine the *fkh-7*.a expression pattern in the larval nervous system, we performed double label single-molecule fluorescence *in situ* hybridization (smFISH) using spectrally-distinct probe sets designed to target the first exon sequence of the *fkh-7*.a mRNA transcript and the *gfp* mRNA sequence in pre-dauer L2d larvae bearing the P*ocr-2*::*gfp* transgene (Figure 2F). *fkh-7*.a*^exon1^* mRNA colocalized with the two most lateral neuronal soma that express *gfp* mRNA in the head ganglia (ADL and ADF; Figure 2G). Although *gfp* mRNA was expressed in two cells in the tail ganglia, *fkh-7*.a*^exon1^* mRNA expression was not observed in the tail region at the detection limit used here (Figure 2H).

Loss of FKH-7 activity could affect sensory neuron cell migration and/or dendritic outgrowth during embryonic development leading to altered sensory neuron function. We examined the expression pattern of a transcriptional GFP reporter construct driven by the *ocr-2* promoter in L1 stage *fkh-7*(*sy1576*) mutants (Figure S2A). The spatial positions of all five neuron classes labelled by P*ocr-2::gfp* in the *fkh-7*(*sy1576*) mutant is comparable with wild-type indicating that loss of FKH-7 activity does not result in aberrant neuronal cell body migration^30^ (Figure S2A). In *C. elegans*, the amphid chemosensory neurons extend their dendrites to the tip of the nose where dendritic endings terminate in ciliated structures that interface with the external environment. The ciliated endings of some chemosensory neuron classes, including ADF and ADL, possess unique morphologies that enable neuron class identification by visual inspection^31^. This aspect of neuron class identity is preserved in the *fkh-7*(*sy1576*) mutants at the L1 stage (Figure S2B,C). Ciliary ultrastructural abnormalities can disrupt dauer formation, and can be diagnosed in a subset of amphid and phasmid sensory neurons, including ADL, by the inability to take up the fluorescent lipophilic dye DiI^32^. *fkh-7(sy1576)* mutant larvae exhibit normal dye-filling of ADL neurons, implying they have functional cilia (Figure S2D).

The forkhead DNA-binding domain of *C. elegans* FKH-7 and *Homo sapiens* FOXP1 share 80.5% amino acid sequence identity (Figure 3A). Large-scale comparative analysis of transcription factor regulatory networks indicate that *in vivo* binding sequence preferences of orthologous transcription factor families are largely conserved^33^. To probe whether this general principle could extend to FKH-7/FOXP, we expressed *H. sapiens FOXP1* cDNA in the subset of sensory neurons labelled by the *ocr-2* promoter, which was sufficient to rescue the *fkh-7(sy1576)* dauer formation defective phenotype (Figure 3B). Therefore, *H. sapiens* FOXP1 can functionally substitute for *C. elegans* FKH-7 in chemosensory neurons, presumably by binding to the same genomic regions and regulating dauer formation-relevant neuronal transcriptional circuits. Advances in clinical exome and whole genome sequencing technology have uncovered many genetic missense variants associated with ASD, a complex neurodevelopmental disorder^34^. A current biomedical challenge is dissecting how these pathogenic missense mutations perturb encoded protein function and/or regulation, as well as identifying variant-specific molecular pathways that are affected and their cell physiological consequences *in vivo*. *FOXP1 de novo* mutations have been recurrently linked to ASD, and the functional conservation of FKH-7/FOXP in *C. elegans* neurons demonstrated above presents a physiologically-relevant system for elucidating the molecular pathology of disease-relevant FKH-7/FOXP partial reduction-of-function alleles^4,5^. To gain basic insights into the molecular mechanism of ASD-associated *FOXP1* variants, we generated analogous *C. elegans fkh-7* hypomorphic missense variants for ASD-associated *H. sapiens FOXP1(R465G)* and *FOXP1(R514C)*, two *de novo* heterozygous *FOXP1* variants identified in patients with intellectual disability^5^ (Figure 3A). When challenged with pheromone dauer-inducing conditions, homozygous *fkh-7(R563G)* and *fkh-7(R612C)* missense variants exhibit dauer formation defective phenotypes that are weaker than the *fkh-7* deletion alleles (Figure 3C). Moreover, *ocr-2* promoter-driven expression of *H. sapiens FOXP1(R514C)* cDNA in *C. elegans* sensory neurons does not rescue the *fkh-7(sy1576)* dauer formation defective phenotype (Figure 3B).

**Figure 3.**
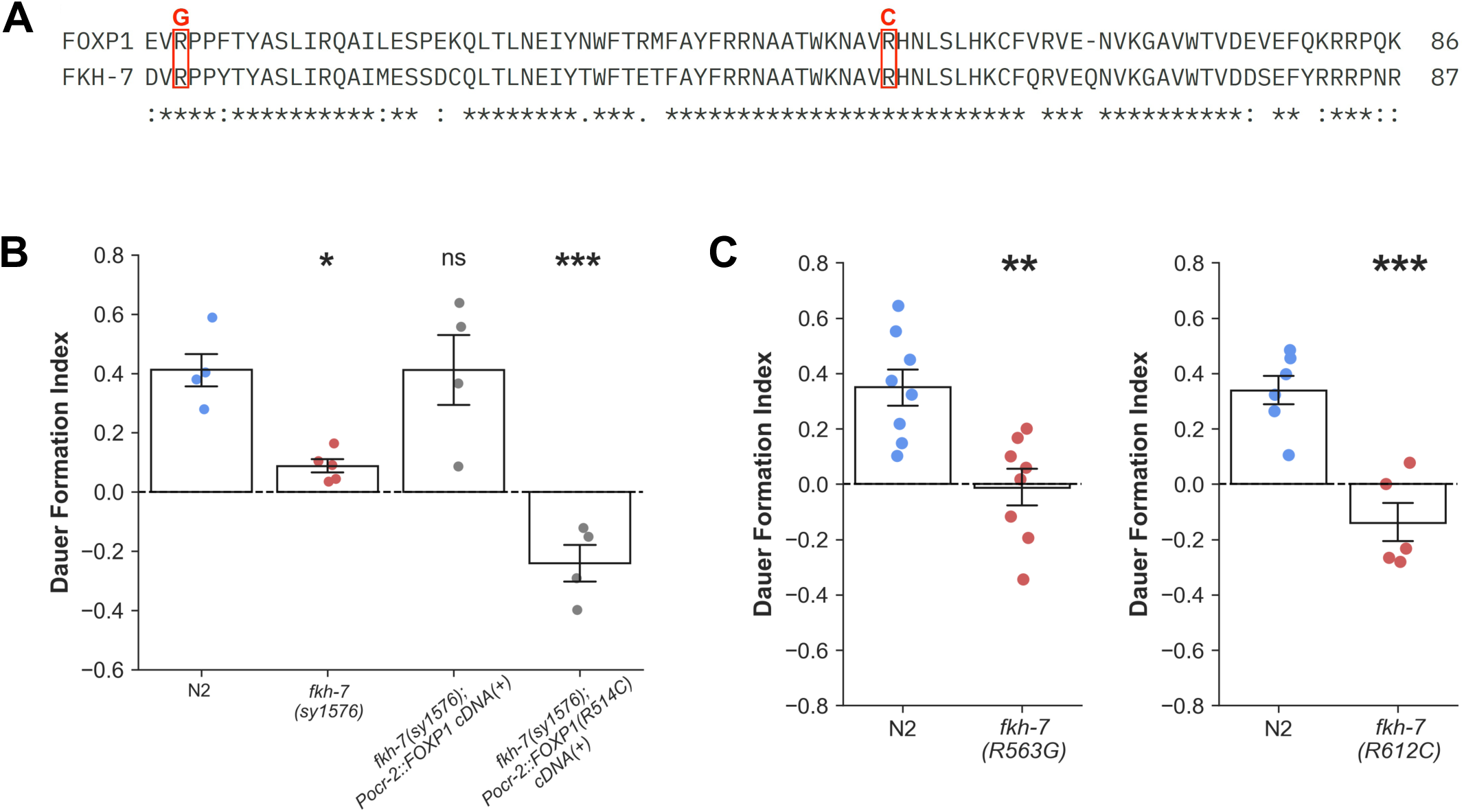
Autism-associated *fkh-7/FOXP1* hypomorphic missense variants are also impaired in developmental decision-making. (A) Alignment of *Homo sapiens* FOXP1 and orthologous *C. elegans* FKH-7 DNA-binding domain protein sequences (InterPro). Red letters and boxes indicate residues with syndromic missense substitutions. (B) Dauer formation assays for *H. sapiens FOXP1* cDNA and *FOXP1(R514C)* cDNA expression in *ocr-2* promoter sensory neuron subset in *fkh-7*(*sy1576*) mutants. N=4-5 population assays, 96-139 animals per assay. Data represented as mean ± SEM, one-way ANOVA followed by Dunnett’s post-hoc test, ns=no significance, *p<0.05, ***p<0.001. (C) Dauer formation assays for *fkh-7(R563G)* and *fkh-7(R612C)* missense variants. N=5-8 population assays, 55-77 animals per assay. Data represented as mean ± SEM, Welch’s unpaired t test, **p<0.01, ***p<0.001.

As a transcription factor, FKH-7 exerts its function by binding to genomic targets and controlling the rate of transcription of associated genes. To identify potential transcriptional targets and molecular pathways that are perturbed in sensory neurons of the autism-associated *fkh-7* missense variant, we compared the single-cell transcriptomes of ADL and ADF neurons between *fkh-7(R612C)* and *fkh-7(wild-type)* L1 stage larvae (Figure 4A,B). ADL transcriptomic profiles did not differ significantly between genotypes, although the singular increase in transmembrane serpentine receptor-encoding *srg-48* expression may alter ADL’s chemosensory function in the *fkh-7(R612C)* variant (Log2FC=1.27, p-adjusted=0.031) (Figure 4C, Data S4). Sequenced ADF cells formed two clusters with divergent gene expression profiles (Cluster 1: 436 cells, Cluster 2: 130 cells; Figure S4A,C, Data S5) and results discussed here refer to the comparison between all *fkh-7(R612C)* ADF cells and all *fkh-7(wild-type)* ADF cells unless stated otherwise. In ADF, the only serotonergic sensory neurons in *C. elegans* hermaphrodites^35,36^, expression of the calcium-activated potassium channel-encoding *kcnl-2/KCNN2* was significantly downregulated in *fkh-7(R612C)* variants relative to the control genotype (Log2FC=-0.92, p-adjusted=2.93E-11) (Figure 4D, Data S5). KCNN2 activation hyperpolarizes the membrane and regulates neuronal excitability^37^, and *KCNN2* is among the common variants that were strongly linked to autism in a landmark large-scale genome-wide association study^38^. Unlike *fkh-7*.a, *kcnl-2* is broadly expressed in the L1 nervous system, including in ADF (Figure 4E).

**Figure 4.**
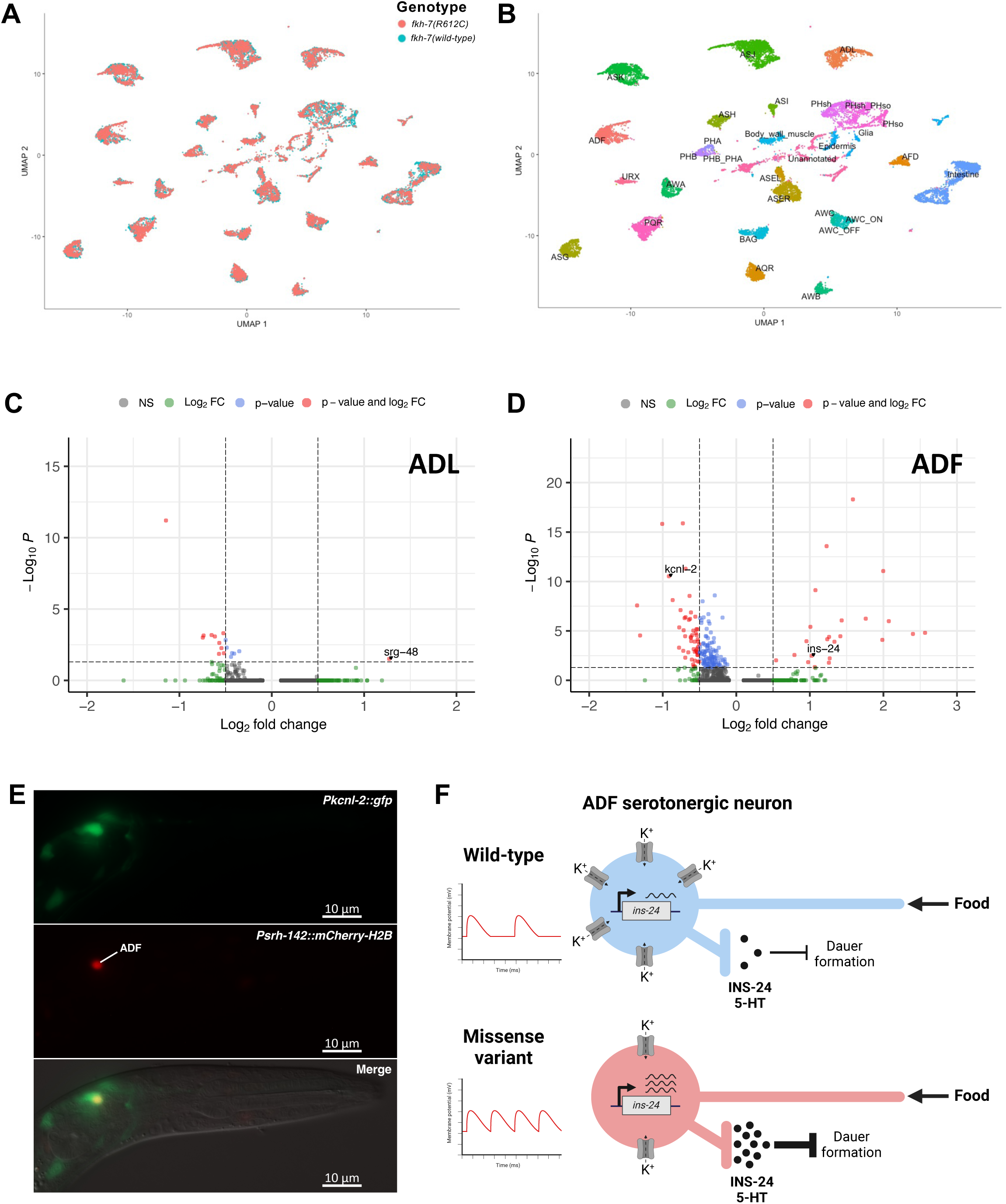
Evolutionarily conserved neuronal function of FKH-7/FOXP enables identification of autism-associated variant-specific molecular pathology at single cell resolution. (A-B) Uniform manifold approximation and projection (UMAP) plots for single-cell RNA-seq of all sensory neurons that express the P*tax-4::gfp* and P*ocr-2::gfp* constructs during the L1 stage. (A) UMAP plot with cells annotated by genotype, *fkh-7(R612C)* in pink and *fkh-7(wild-type)* in teal. (B) UMAP plot with cell clusters annotated by neuron class. (C) Volcano plot showing differentially expressed genes in ADL neuron class for *fkh-7(R612C)* missense mutant compared to *fkh-7(wild-type)* control during the L1 stage (thresholds: Log2FC>|0.5| and adjusted p-value<0.05). *srg-48* gene labeled. Differential gene expression analysis shown here compares all *fkh-7(R612C)* ADL cells with all *fkh-7(wild-type)* ADL cells (see also Figure S3). For the list of genes that are differentially expressed in *fkh-7(R612C)* ADL neurons compared to *fkh-7(wild-type)*, see Data S4. (D) Volcano plot showing differentially expressed genes in ADF neuron class for *fkh-7(R612C)* missense mutant compared to *fkh-7(wild-type)* control during the L1 stage (thresholds: Log2FC>|0.5| and adjusted p-value<0.05). *kcnl-2 and ins-24* genes labeled. Differential gene expression analysis shown here compares all *fkh-7(R612C)* ADF cells with all *fkh-7(wild-type)* ADF cells. *fkh-7(R612C)* ADF cells grouped into two distinct clusters (see Figure S4). For analysis using *fkh-7(R612C)* ADF cells from Cluster 2 only, see Figure S4D. For the list of genes that are differentially expressed in *fkh-7(R612C)* ADF neurons compared to *fkh-7(wild-type)*, see Data S5. (E) P*kcnl-2*::*gfp* and P*srh-142*::*mCherry-H2B* colocalization in an L1 stage larva. Head region shown only. (F) Speculative model: In the *fkh-7(R612C)* missense variant, lower abundance of *kcnl-2/KCNN2* calcium-activated potassium channels may increase the excitability of ADF neurons, and in synergy with higher expression levels of the dauer development-inhibiting *ins-24* insulin-like peptide, could result in increased INS-24 and/or serotonin (5-HT) secretion and stronger inhibition of entry into the dauer developmental trajectory.

One plausible mechanism is that decreased *kcnl-2* expression due to reduction of FKH-7 activity in *fkh-7(R612C)* ADF neurons increases ADF excitability in response to favorable stimuli, such as food, which in turn increases the secretion of dauer development-inhibiting factors (Figure 4F). Although serotonin inhibits dauer formation^35^, expression of *bas-1*, which encodes an amino acid decarboxylase that catalyzes a critical step in serotonin biosynthesis^39^, is downregulated in Cluster 2 ADF *fkh-7(R612C)* cells relative to the control genotype (Log2FC=-1.00, p-adjusted=0.018) (Figure S4D, Data S5). However, the expression of other secreted dauer development-inhibiting factors such as the insulin-like peptide *ins-24* is significantly upregulated in *fkh-7(R612C)* ADF neurons relative to the control genotype^40^ (Log2FC=1.02, p-adjusted=3.26E-03) (Figure 4D, Data S5). Although a subset of *fkh-7(R612C)* ADF cells (Cluster 2) likely produces less serotonin as a result of decreased *bas-1* expression, the general decrease in *kcnl-2* potassium channel and increase in *ins-24* insulin-like peptide expression levels could synergize to inhibit dauer development in *fkh-7(R612C)* variants (Figure 4F).

## DISCUSSION

In summary, we have provided the first functional characterization of clinically significant FKH-7/FOXP in the widely studied model organism *C. elegans*. *fkh-7(sy1576)* mutants exhibit much lower penetrance of the decreased dauer formation phenotype compared with *daf-16(mgDf50)* null mutants (Figure 1E). Dauer formation is almost completely abolished in *ocr-2(ak47)* null mutants under pheromone dauer-inducing conditions, suggesting that loss of FKH-7/FOXP activity in ADL and ADF neurons only partially impairs neuronal function (Figure S1D). Moreover, *fkh-7*.a is expressed and acts in a subset of upstream afferent nodes (sensory neurons) of the physiological network regulating dauer formation^20,21^. In contrast, DAF-16/FOXO likely acts as a downstream integrator of DAF-2-mediated insulin-like signaling in the intestine where it activates cellular stress-response effector pathways that promote dauer formation in response to adverse environmental conditions^27,41,42^. Given the *fkh-7*.a site-of-action in the intestine and the increased intestinal *fkh-7.*a expression post-entry into the dauer developmental trajectory, FKH-7/FOXP intestinal activity may also be regulated by DAF-16/FOXO (Figure 1F, 2A).

In *Foxp1* conditional knockout mice, dopamine receptor 2-expressing indirect pathway striatal spiny projection neurons (D2 SPNs) exhibit decreased expression of *Kcnk2*, *Kcnj2*, and *Kcnj4* potassium channels, which may underlie the increased intrinsic excitability of D2 SPNs with *Foxp1* deletion^43–45^. In *Drosophila* fruit flies, *FoxP* knockdown or knockout in mushroom body αβ core Kenyon cells (αβ_c_ KCs) increases voltage-dependent potassium channel-encoding *Shal* transcript abundance, which impinges upon biophysical properties of αβ_c_ KCs such as lowering input resistance (excitability) and shortening the membrane time constant^46^. Here, our site-of-action experiments coupled with unbiased comparative single-cell transcriptomic analysis of the entire chemosensory apparatus in L1 stage larvae have revealed that a hypomorphic *C. elegans fkh-7* missense variant harbors decreased expression levels of calcium-activated potassium channel-encoding *kcnl-2* in the serotonergic ADF chemosensory neuron (Figure 4D). Collectively, these findings across phylogenetically-distant model organisms suggest that controlling neuronal activity by tuning the stoichiometry of potassium channels in the membrane could be a broadly conserved function of neuronal FOXP1. By combining approaches in molecular genetics and single-cell transcriptomics, we have established a highly tractable *in vivo* system to further probe the neurophysiological link between two autism-associated genes - the forkhead transcription factor *fkh-7/FOXP1* and the potassium channel *kcnl-2/KCNN2* – at the resolution of a single sensory neuron in the context of an adaptive developmental decision.

## DECLARATION OF INTERESTS

The authors declare no competing interests.

## ACKNOWLEDGEMENTS

We thank Stephanie Nava and Wilber Palma for Nematode Growth Media plate preparation, Barbara Perry for strain freezing, Sarah Torres for administrative assistance, the VUMC (Vanderbilt University Medical Center) FCSR (Flow Cytometry Shared Resource) and VANTAGE (Vanderbilt Technologies for Advanced Genomics) for generating single-cell sequencing data. Strains CX4544 and GR1307 were obtained from the *Caenorhabditis* Genetics Center, which is funded by NIH Office of Research Infrastructure Programs (P40 OD010440). The *fkh-7(tm6093)* allele was provided by the lab of Dr. Shohei Mitani as part of the National Bioresource Project. Figure 1A and Figure 4F were generated with BioRender.com. Figure 1B was generated using the Exon-Intron Graphic Marker tool by Dr. Nikhil Bhatla. P.W.S., C.M.C., and W.-R.W. were supported by Simons Foundation SFARI award #367560 and a UF1NS111697 grant to P.W.S., and the Howard Hughes Medical Institute, with which P.W.S. was an investigator. D.M.M. and S.R.T. were supported by grants R01 NS100547, R01 NS113559, and R01 NS106951. L.C. and C.H.T. were supported by an Allen Discovery Grant.

C.M.C. received support from a Neurobiology Graduate Fellowship (Division of Biology and Biological Engineering, California Institute of Technology). The VUMC FCSR is supported by the Vanderbilt Ingram Cancer Center (P30 CA068485) and the Vanderbilt Digestive Disease Research Center (DK058404).

## AUTHOR CONTRIBUTIONS

C.M.C. conceived of the study. C.M.C. performed crude pheromone extraction, dauer formation assays and analysis, CRISPR mutagenesis (deletion alleles), cDNA library generation, molecular cloning, microinjections, transgenesis and genomic integration, DiI live-animal staining, microscopy, and data visualization. C.M.C. and P.W.S. conceived of generating *fkh-7* autism-associated missense alleles, and W.-R.W. performed CRISPR mutagenesis to generate missense alleles. D.M.M., S.R.T., P.W.S., and C.M.C. designed the single-cell RNA-seq experiments. S.R.T. performed single-cell RNA-seq experiments, data analysis, and data visualization. L.C., C.H.T., and C.M.C. designed the smFISH experiments. C.M.C. performed smFISH sample staging, fixation, hybridization, and wash steps. C.H.T. performed smFISH probe design, sample digestion, DAPI staining, and imaging. P.W.S., D.M.M., and L.C. provided resources and acquired funding. C.M.C. wrote the paper with input from all co-authors. All authors edited and approved the final manuscript.

## MATERIALS AND METHODS

### EXPERIMENTAL MODEL AND SUBJECT DETAILS

*Caenorhabditis elegans* strains were cultivated at 21°C on standard Nematode Growth Media (NGM) plates seeded with *Escherichia coli* strain OP50 cultured in Luria-Bertani (LB) broth. Details of all strains used in this study are available in Table S1. Data pertaining to the genomic sequence and gene structures of *C. elegans* were sourced from WormBase^47^.

### METHOD DETAILS

#### Molecular cloning and plasmid construction

All plasmids generated in this study are listed in Table S2. Genomic DNA was extracted and purified from *C. elegans* N2 wild-type strain using the Wizard® Genomic DNA Purification Kit (Promega, Catalog#A1120). Gene promoter fragments were PCR amplified from N2 template genomic DNA using Phusion® High-Fidelity DNA Polymerase (NEB, Catalog#M0530) with the primer sequences listed in Table S3. The promoters of the following genes were amplified: *fkh-7*.a (4.56 kb), *rab-3* (1.21 kb), *ges-1* (2 kb), *myo-2* (962 bp), *myo-3* (2.5 kb), *dpy-7* (343 bp), *ocr-2* (2.5 kb), *tax-4* (3.03 kb), *sre-1(3 kb)* (3.03 kb), *sre-1(1 kb)* (1.02 kb), *srh-142* (3.35 kb), *osm-10* (889 bp), and *kcnl-2* (1.79 kb). To generate transcriptional fluorescent reporter constructs, gene promoters were inserted upstream of the fluorescent protein transgene in *PSM-gfp* (gift from Dr. Cornelia Bargmann) or *PSM-mCherry::H2B* double-digested with FseI (NEB, Catalog#R0588) and AscI (NEB, Catalog#R0558) enzymes, using T4 DNA ligation (NEB, Catalog#M0202) or NEBuilder® HiFi DNA Assembly Master Mix (NEB, Catalog#E2621). *PSM-mCherry::H2B* vector was generated by inserting the *mCherry::H2B* sequence (PCR amplified from pCFJ167, a gift from Dr. Erik Jorgensen) into *PSM-gfp* plasmid double-digested with AgeI-HF (NEB, Catalog#R3552) and EcoRI-HF (NEB, Catalog#3101) to remove the *gfp* sequence, using T4 DNA ligation.

cDNA library was generated from N2 wild-type strain using the Superscript^TM^ III First-Strand Synthesis System (Invitrogen, Catalog#18080051). *fkh-7*.a cDNA was PCR amplified using N2 cDNA library as template with the primers listed in Table S3 (Data S2). The *sl2* bicistronic linker sequence was PCR amplified from plasmid pJL046^48^. The *fkh-7.*a cDNA and *sl2* sequences were assembled upstream of the GFP transgene in the *PSM-gfp* vector backbone double-digested with AscI and KpnI-HF (NEB, Catalog#R3142) restriction enzymes, using NEBuilder® HiFi DNA Assembly Master Mix. The sequence of the resulting *PSM-fkh-7.a cDNA::sl2::gfp* vector was confirmed by Sanger sequencing (Laragen, Culver City, CA) using primers F1-F4 and R1-R4 (see Table S3). *PSM-fkh-7.a cDNA::sl2::gfp* was double-digested with FseI and AscI enzymes, and gene promoters inserted upstream of *fkh-7*.a cDNA sequence using T4 DNA ligation or NEBuilder® HiFi DNA Assembly Master Mix.

*Homo sapiens FOXP1* cDNA (CCDS2914.1) and *FOXP1(R514C)* cDNA sequences were synthesized by Integrated DNA Technologies gBlocks^TM^ (Data S3). The *FOXP1* or *FOXP1(R514C)* cDNA and *sl2* sequences were assembled upstream of the GFP transgene in the *Pocr-2::fkh-7.a cDNA::sl2::gfp* plasmid double-digested with AscI and AgeI-HF to remove the *fkh-7.a cDNA::sl2* sequence, using NEBuilder® HiFi DNA Assembly Master Mix.

#### Transgenesis and genomic integration of extrachromosomal arrays

Transgenic strains were generated by microinjection of plasmid DNA into the syncytial gonad arms of adult *C. elegans* hermaphrodites using standard methods^49,50^. Plasmids were injected at 5-50 ng/µL. *Pofm-1::rfp* (gift from Dr. Cornelia Bargmann) co-injection marker was injected at 40 ng/µL. 1 kb DNA ladder (NEB, Catalog#N3232) was used as a filler to bring final DNA concentration of the injection mix to 200 ng/uL. For strains PS8795: *syIs692[Pocr-2::gfp]* and PS9221: *syIs764[Pocr-2::gfp, Ptax-4::gfp]*, genomic integration of extrachromosomal arrays was performed using X-ray irradiation. Integrants were backcrossed with N2 wild-type strain at least three times prior to experiments.

#### Generation of *fkh-7* DNA-binding domain deletion mutants

Strains PS7375: *fkh-7(sy1575)* and PS7376: *fkh-7(sy1576)* were generated using the CRISPR co-conversion strategy described in Arribere *et al.* 2014^51^, where the *dpy-10* locus (Chromosome II) is co-targeted for repair to introduce a missense mutation that produces a dominant roller phenotype. Forward and reverse oligonucleotides for five sgRNAs targeting *fkh-7*’s DNA-binding domain (Chromosome IV) were annealed at 95°C for 2 min (see Table S3 for sgRNA sequences). Each annealed sgRNA was assembled into plasmid vector pRB1017^51^ digested with BsaI-HF (NEB, Catalog#R3535) using T4 DNA ligation. The injection mix comprised *Peft-3::cas9* plasmid (Addgene#46168^52^), *dpy-10* sgRNA plasmid (pJA58^51^), five *fkh-7* sgRNA plasmids, and *dpy-10(cn64)* repair template single-strand oligonucleotide. F1 progeny exhibiting roller and/or dumpy phenotypes were singled out and propagated. F2 progeny were subjected to PCR-based genotyping of the *fkh-7* locus to identify successful CRISPR mutagenesis events (see Table S3 for genotyping primers). Following a second round of genotyping to isolate mutants homozygous for *fkh-7* deletion alleles, deletion breakpoints were confirmed by Sanger sequencing (Laragen, Culver City, CA) of purified PCR products.

#### Generation of autism-associated *fkh-7/FOXP1* missense strains

Autism-associated missense strains PS8063: *fkh-7(sy906)* and PS8065: *fkh-7(sy1316)* were generated using the CRISPR homology directed repair-based pipeline described in Wong *et al.* 2019^53^. The pipeline utilizes direct injection of *in vitro* assembled Cas9-crRNA-tracrRNA ribonucleoprotein complexes as described in Paix *et al.* 2015^54^ and the *dpy-10* CRISPR co-conversion strategy described in Arribere *et al.* 2014^51^. Additionally, synonymous mutations were engineered during the target gene repair process to introduce a restriction enzyme site to facilitate the PCR-based screening process. See Table S3 for crRNA sequences, universal tracrRNA sequence, and single-strand oligonucleotide repair templates. Purified Cas9 protein was a gift from Dr. Tsui-Fen Chou (Caltech). F1 progeny exhibiting roller and/or dumpy phenotypes were singled out and propagated. F2 progeny were subjected to PCR-based genotyping of the *fkh-7* locus (see Table S3 for genotyping primers) and analysis of enzyme digested PCR product lengths. Following a second round of genotyping to isolate mutants homozygous for *fkh-7* missense alleles, target gene modification was confirmed by Sanger sequencing of purified PCR products.

#### Pheromone-induced dauer formation assays

Crude dauer pheromone extract was prepared from liquid cultures of N2 wild-type strain as previously described in Flatt and Schroeder 2014^55^. To control for differences in pheromone potency between batches, a pheromone dose response curve was generated for each batch and the pheromone volume that yielded approximately 65% dauer formation for the N2 wild-type strain under the experimental conditions described below was selected for dauer formation assays, unless otherwise specified. 8% (weight-to-volume) heat-killed *E.coli* OP50 was prepared by washing bacteria from an overnight culture with virgin S-basal buffer, weighing the bacterial pellet, resuspending with the appropriate volume of virgin S-basal buffer, and killing at 95-100°C on a heat block. 2 µL of the heat-killed bacteria was then streaked on an LB agar plate and incubated overnight at 37°C to confirm heat killing efficacy.

Pheromone-induced dauer formation assays were performed as previously described in Chai *et al.* 2022^23^. The afternoon before experiments, L4 stage larvae were picked from well-fed stock plates onto seeded plates and grown overnight at 21°C. Assay plates were prepared by mixing pheromone of desired volume with 2 mL of NGM agar without peptone in each 35 mm Petri dish. Plates were dried overnight on benchtop at 21°C. On the day of experiments, pheromone plates were seeded with 2 µL heat-killed OP50 and adult animals were transferred to assay plates for egg-laying. Adults were then picked off and 18 µL of heat-killed OP50 added to the plates. Once the food patch had completely dried, plates were sealed with parafilm and placed in an empty pipette tip box, which was placed in a 25.5°C incubator for 72 hours. Control and experimental strains were always assayed simultaneously under the same conditions in the same pipette tip box for statistical comparisons. ‘N’ in the figure legends refers to the number of population assay plates performed per genotype. After the incubation period, the number of dauers and non-dauers per plate were counted and the Dauer Formation Index (DFI) per plate calculated using the following formula:

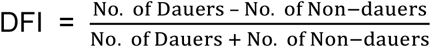

All site-specific rescue experiments were repeated at least twice on two different days, and the results pooled for analysis. The number of dauers, non-dauers, and the total number of animals per population assay plate for all dauer formation assays are recorded in Data S1. Please note that the P*dpy-7*::*fkh-7*.a cDNA rescue (Figure 2A) and the P*tax-4*::*fkh-7*.a/P*ocr-2*::*fkh-7*.a cDNA rescue (Figure 2B) dauer formation experiments utilize the same N2 wild-type and *fkh-7(sy1576)* controls, and are displayed in separate figure panels for organizational clarity.

#### Live-animal DiI staining of sensory neurons

1,1’-Dioctadecyl-3,3,3’,3’-Tetramethylindocarbocyanine Perchlorate (DiI, Invitrogen Catalog#D282) stock solution was prepared at a concentration of 2 mg/mL in dimethyl formamide. DiI working solution (1:200 dilution) was prepared by adding 1 µL of DiI stock solution to 199 µL of M9 buffer in a 1.5 mL microcentrifuge tube and vortexing to mix. Larvae at the appropriate life stage were transferred from NGM plates into the DiI working solution using a worm pick with a small amount of live *E. coli* OP50, and incubated in the dark at room temperature for two hours. Larvae were then transferred to a fresh seeded NGM plate using a P1000 pipette tip, and allowed to roam for one hour at room temperature in the dark to destain prior to microscopy.

#### Microscopy

L1 and L2 stage larvae were identified based on gonad development (see Figure 8 in https://www.wormatlas.org/hermaphrodite/introduction/Introframeset.html) and transferred from well-fed stock plates to glass slides using a worm pick. Larvae were immobilized in 5 µL of 50 mM sodium azide (Sigma, Catalog#S2002) on 2% agarose pads prepared on glass slides. Slides were imaged using a Zeiss Axio Imager.Z2 widefield microscope attached to an AxioCam 506 mono camera capture source. Image in Figure S1C is a maximum intensity projection of a z-stack acquired using the Zeiss ApoTome.2 structured illumination microscopy system. All downstream image processing was performed using Zeiss ZEN Blue microscopy software.

#### Single-molecule fluorescence *in situ* hybridization (smFISH)

##### Sample staging

Pheromone-induced pre-dauer L2d stage larvae were used for smFISH experiments. Golden and Riddle 1984^14^ reported that the presence of pheromone, even at low concentrations that ultimately induce only a few percent dauer larvae, is sufficient to induce all larvae to assume L2d morphology. Dauer formation assay plates were prepared and set up as described above using crude pheromone concentration that induces >95% dauer formation. At 28 hours post egg-laying, larvae were inspected under a dissecting microscope for darkened intestines and cessation of pharyngeal pumping, indicating that larvae are in the L2d developmental stage^14^, before proceeding with the smFISH protocol below.

##### Experimental protocol

smFISH experiments were conducted as previously described with some modifications^56,57^. All solution changes were preceded by centrifugation performed at 4000 RPM for 2 minutes and were performed at room temperature unless specified otherwise. Staged worms were collected in 1.7 mL Eppendorf tubes with water and washed three times in water, or until remaining bacteria were removed. Samples were fixed with freshly prepared, ice - cold 4% PFA in 1x PBS for 45 minutes on a tube rotator. The worms were rinsed with 0.01% 1x PBST (1x PBS with 0.01% Triton X-100) and washed twice in 1x PBST with 15 minutes incubation time. Worms were stored in 100% methanol overnight at -20°C. and rehydrated with 30-minute washes in 75%, 50%, and 25% methanol in 1x PBST. Worms were washed in 1x PBST for another 30 minutes and rinsed with 1x BOT buffer (Borate buffer with 0.01% Triton X-100). Worms were reduced with freshly prepared TCEP solution (1x BOT with TCEP) for 25 minutes, washed with 1x BOT, and oxidized in freshly made H2O2 solution (1X BOT with 0.03% hydrogen peroxide) for 10 minutes. After brief washes with 1x BOT and 1x PBST, worms were cleared in 8% SDS in 1X PBS for 45 minutes at 37°C. Cleared worms were washed twice in 1X PBST for 15 minutes and hybridized with 5 nM *fkh-7^exon1^* and *gfp* primary probes and 1 µM LNAPolyT30-Acryd in pre-warmed hybridization buffer (10% dextran sulfate (Sigma, Catalog#D8906) and 50% formamide in 4x SSC). Primary probes were hybridized for 48 hours at 37°C. See Table S4 for primary probe sequences.

Following hybridization, worms were rinsed with 4x SSCT (0.01% Triton X-100 in 4x SSC) and washed three times in 4x SSCT for 30 minutes on the rotator. Then, samples were washed with 50% wash buffer (50% formamide in 2x SSCT) for 1 hour at 37°C. Samples were again washed three times in 4x SSCT for 30 minutes each. Worms were incubated with gel solution (4% 19:1 acrylamide/bis-acrylamide and VA-044 initiator in 1x PBS) overnight at 4°C. Worms were washed with freshly prepared gel solution (treated with nitrogen gas for 15 minutes on ice) prior to transferring them to a coverslip functionalized with bind-silane and Poly-D-lysin. Worms in gel were gently pressed with a Gel-Slick-coated coverslip. Samples were incubated for 4 hours at 37°C in a humidification chamber filled with nitrogen for gel polymerization. Gel-slick coverslips and excess gel were removed, and a flow cell was attached to the sample coverslip. Worms were digested with proteinase K solution (in mM: 50 Tris-HCl pH 8, 1 EDTA, 0.5% Triton X-100, 500 NaCl, 1% SDS, and 1:100 proteinase K) overnight at 37°C. Samples were washed three times with 4X SSC.

##### Imaging

5 nM readout sequences conjugated to Alexa Fluor 647 for *fkh-7^exon1^* primary probes and Cy3B for *gfp* probes in EC buffer (10% ethylene carbonate and 10% dextran sulfate (Sigma, Catalog#D4911) in 4x SSC) were added to the sample. Primary probes were designed to contain three binding sites for readout probes. The buffer was incubated for 30 minutes at room temperature. After hybridization, the sample was washed with 10% formamide wash buffer (10% formamide and 0.1% Triton X-100 in 4x SSC) to remove excess and non-specifically bound readout probes. Next, the sample was rinsed with 4x SSC before staining with DAPI solution (5 µg/ml DAPI in 4x SSCT) for at least 10 minutes. Anti-bleaching buffer (Trolox, 0.8% glucose, 1:100 diluted catalase, 0.5 mg/ml glucose oxidase, and 50 mM pH 8 Tris-HCl in 2x SSC) was applied to the sample for imaging.

Imaging was performed with a microscope (Leica DMi8) equipped with a confocal scanner unit (Yokogawa CSU-W1), a sCMOS camera (Andor Zyla 4.2 Plus), a 40X oil objective lens (Leica 1.30 NA) and a motorized stage (ASI MS2000). Lasers from CNI and filter sets from Semrock were used. Images were acquired with 1-μm z-steps for z-slices per FOV across 647 nm, 561 nm, and 405 nm fluorescent channels. Imaging was controlled by a custom written script in μManager. Figure 2F is a maximum intensity projection.

#### Single-cell RNA-sequencing of sensory neurons

##### Growth conditions and developmental synchronization

Worms were grown on 8P nutrient agar 150 mm plates seeded with *E. coli* strain NA22. To obtain synchronized cultures of mid-late L1 worms, we scraped worms and eggs off 150 mm plates with a cell scraper and pelleted by centrifugation at 150 rcf for 2.5 minutes. Embryos were isolated by hypochlorite treatment of the pellet and floatation in 30% sucrose. Embryos were washed 2x in M9, and then placed on a nylon mesh grid with 10-micron pores. Embryos will not pass through the pores of the mesh, but hatched L1 larvae are able to crawl through. The mesh was placed on M9 containing NA22 bacteria, and embryos were allowed to hatch at room temperature. Larvae collected in the first 1.5 hours after completion of hypochlorite treatment were discarded, and the mesh grid was placed over a fresh collection media of M9 with NA22 bacteria for two hours. Hatched L1s were pelleted by centrifugation, and then plated onto fresh 8P 150 mm plates with NA22 and incubated at 20°C for 13.5 hours.

##### Dissociation

Single cell suspensions were obtained as previously described with some modifications^58–61^. Worms were collected and separated from bacteria by washing twice with ice-cold M9 and centrifuging at 150 rcf for 2.5 minutes. Worms were transferred to a 1.6 mL centrifuge tube and pelleted at 16,000 rcf for 1 minute. 100 µL pellets of packed worms were treated with 200 µL of SDS-DTT solution (20 mM HEPES, 0.25% SDS, 200 mM DTT, 3% sucrose, pH 8.0) for 2 minutes. Following SDS-DTT treatment, worms were washed five times by diluting with 1 mL egg buffer and pelleting at 16,000 rcf for 30 seconds. Worms were then incubated in pronase (15 mg/mL, Sigma-Aldrich P8811, diluted in egg buffer) for 23 minutes. During the pronase incubation, the solution was triturated by pipetting through a P1000 pipette tip for four sets of 80 repetitions. The status of dissociation was monitored under a fluorescence dissecting microscope at 5-minute intervals. The pronase digestion was stopped by adding 750 µL L-15 media supplemented with 10% fetal bovine serum (L-15-10), and cells were pelleted by centrifuging at 530 rcf for 5 minutes at 4°C. The pellet was resuspended in L-15-10, and single-cells were separated from whole worms and debris by centrifuging at 150 rcf for 2 minutes at 4 °C. The supernatant was then passed through a 35-micron filter into the collection tube. The pellet was resuspended a second time in L-15-10, spun at 150 rcf for 2 minutes at 4°C, and the resulting supernatant was added to the collection tube.

##### Fluorescence-Activated Cell Sorting

Fluorescence Activated Cell Sorting (FACS) was performed on BD FACSAria™ III Cell Sorters equipped with 70-micron diameter nozzles. DAPI was added to the sample (final concentration of 1 µg/mL) to label dead and dying cells. Wild-type and *fkh-7* (R612C) worms (PS9221 and PS9222) were sorted simultaneously on two different machines. N2 worms lacking GFP fluorescence were used to set gates to exclude auto-fluorescent cells. Cells were sorted under the “4-way purity” mask. GFP+ cells were collected into 400 µL of chilled (4°C) L-15 media containing 33% FBS (L-15-33). After ∼300,000 cells we switched to a new collection tube. For PS9221, we collected 300,000 cells (Sample 1), then a subsequent 25,024 cells. For PS9222, we collected 284,706 cells (Sample 2) and 128,859 cells (Sample 3). Cells were stored on ice. We concentrated the collected cells by centrifuging at 500 rcf for 5 minutes at 4°C. We removed all but ∼55 µL of the supernatant, resuspended the cells and counted fluorescent cells on a hemocytometer. Three collection tubes had sufficient concentrations and volumes for 10X Genomics single-cell RNA-sequencing: Sample 1, from PS9221 with 40 µL at 300 cells/µL; Sample 2, from PS9222 with 43 µL at 460 cells/µL; and Sample 3, from PS9222 with 43 µL at 320 cells/µL.

##### Single-cell RNA-Sequencing

Each sample (targeting 10,000 cells per sample) was processed for single cell 3’ RNA sequencing utilizing the 10X Chromium system. Libraries were prepared using P/N 1000121 following the manufacturer’s protocol. The libraries were sequenced using the Illumina NovaSeq 6000 with 150 bp paired end reads. Real-Time Analysis software (RTA, version 2.4.11; Illumina) was used for base calling and analysis was completed using 10X Genomics Cell Ranger software (v6.0.0).

##### Data Analysis

We mapped reads onto a custom WS273 reference genome with extended 3’ UTR sequences^62^. We used the DropletUtils R package (version 1.6.1) function EmptyDrops (with a threshold of 50 UMIs) to distinguish cells from empty droplets. We performed quality control with the scater package (1.14.6), using the percentage of UMIs mapping to the mitochondrial transcripts *nduo-1, nduo-2, nduo-3, nduo-4, nduo-5, nduo-6, ctc-1, ctc-2, ctc-3, ndfl-4, atp-6, ctb-1, MTCE.7,* and *MTCE.33*. The data for the three samples were then combined and further processed using the monocle3 package (version 0.2.2.0). Genes detected in fewer than 5 cells were removed, and data were log-normalized. Initial data analysis revealed that cells from Sample 2 were of much lower quality than cells from Samples 1 and 3; therefore, sample 2 was removed from the analysis.

We performed dimensionality reduction using PCA (using 60 PCs, based on an elbow plot showing the variance explained by each principal component), followed by UMAP (with umap.min_dist = 0.3, umap.n_neighbors = 75). Cells were clustered on UMAP coordinates using the default leiden algorithm (res = 1e-3). Clusters were assigned cell type identities based on the expression of known cell-type specific genes^62,63^.

##### Differential Gene Expression Analysis

To explore differential gene expression between genotypes for ADF and ADL, we generated individual datasets corresponding to only ADF and or ADL neurons. We then repeated dimensionality reduction for ADF (25 PCs for PCA, umap.min_dist = 0.1 and umap.n_neighbors = 50 for UMAP, res = 7e-4 for clustering). This revealed two clusters, one of which almost exclusively contained ADF neurons from *fkh-7(R612C)* worms. The other cluster contained both *fkh-7(wild-type)* and *fkh-7(R612C)* ADF neurons. We converted the data to a Seurat object and performed 3 differential expression analyses: 1) all *fkh-7(R612C)* cells vs. all *fkh-7(wild-type)* cells, 2) cluster 2 (∼97% *fkh-7(R612C)*) vs cluster 1 (mixed genotypes), and 3) cluster 2 *fkh-7(R612C)* ADF neurons vs cluster 1 *fkh-7(wild-type)* ADF neurons. All comparisons used the default Wilcoxon rank-sum test with logFC_threshold = 0.1 and min_pct = 0.05 parameters.

We repeated this for ADL (15 PCs for PCA, umap.min_dist = 0.1 and umap.n_neighbors = 50 for UMAP, res = 1e-3 for clustering). ADL showed two distinct clusters driven not by genotype, but by sequencing depth. For ADL, we performed two differential expression analyses: 1) *fkh-7(R612C)* neurons vs *fkh-7(wild-type)* neurons for all ADL neurons, and 2) *fkh-7(R612C)* neurons vs *fkh-7(wild-type)* neurons from cluster 2, the cluster with the highest quality cells. Differential expression was performed in Seurat, with the same parameters as for ADF. Volcano plots were generated using EnhancedVolcano.

## QUANTIFICATION AND STATISTICAL ANALYSIS

For dauer formation assays, Welch’s unpaired t test or one-way ANOVA followed by Dunnett’s post-hoc test compared to N2 was used to determine statistically significant differences between groups. All statistical analyses were performed using the SciPy package in Python. All p-values are reported in Data S1.

## SUPPLEMENTAL FIGURES

**Figure S1.**
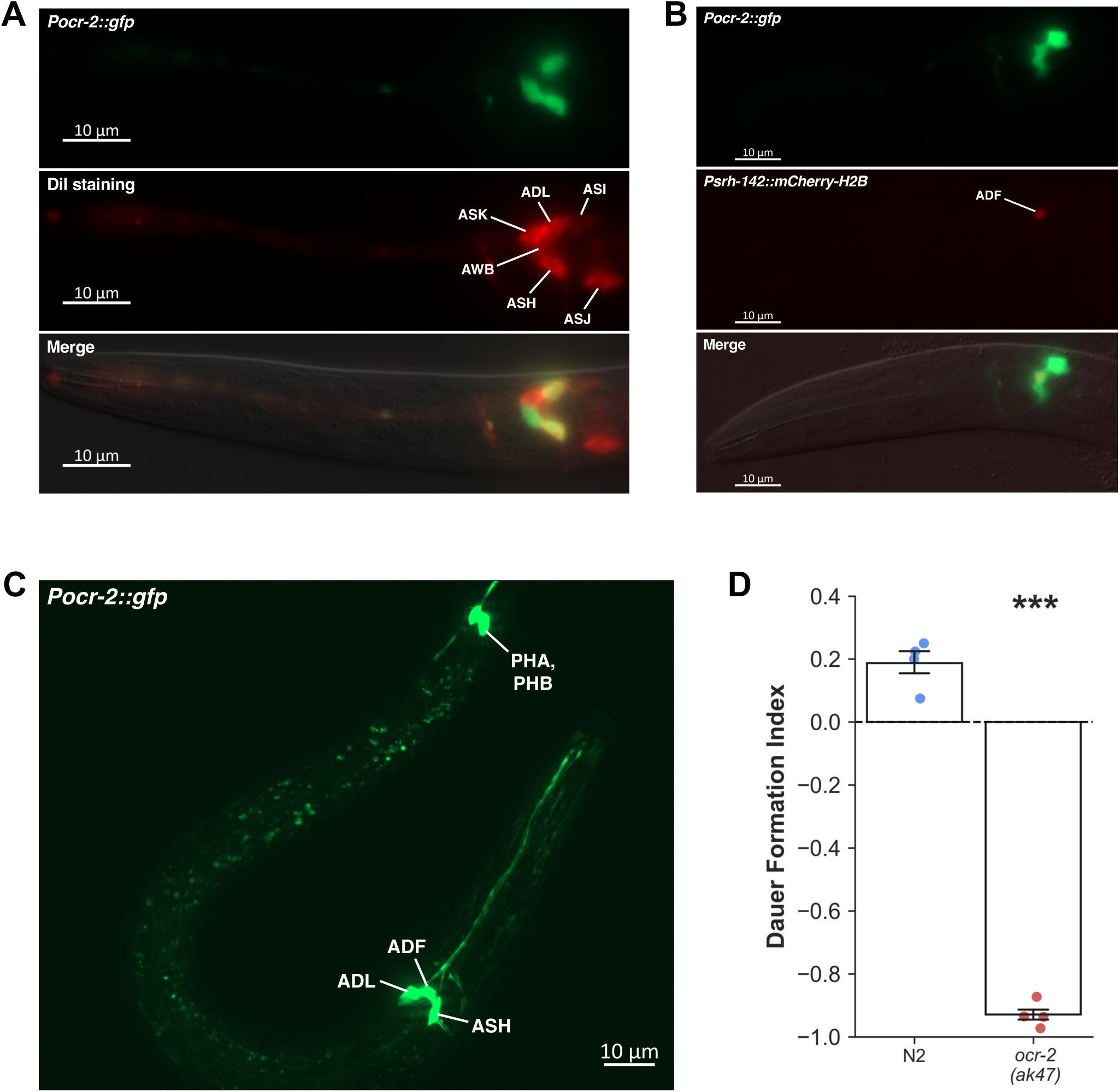
The 2.5 kb upstream regulatory sequence of *ocr-2* drives expression in the ADL, ADF, and ASH sensory neurons in the head as well as in two tail neurons that are likely to be the PHA and PHB sensory neurons (Tobin *et al.* 2002). Related to Figure 2. (A) P*ocr-2::gfp* and DiI staining (ADL and ASH) colocalization in an L1 stage larva. Head region shown only. (B) P*ocr-2::gfp* and P*srh-142*::*mCherry-H2B* (ADF) colocalization in an L1 stage larva. Head region shown only. (C) Whole-animal P*ocr-2*::*gfp* expression in an L1 stage larva. (D) Dauer formation assays for *ocr-2*(*ak47*) mutant. N=4 population assays, 62-74 animals per assay. Data represented as mean ± SEM, Welch’s unpaired t test, ***p<0.001.

**Figure S2.**
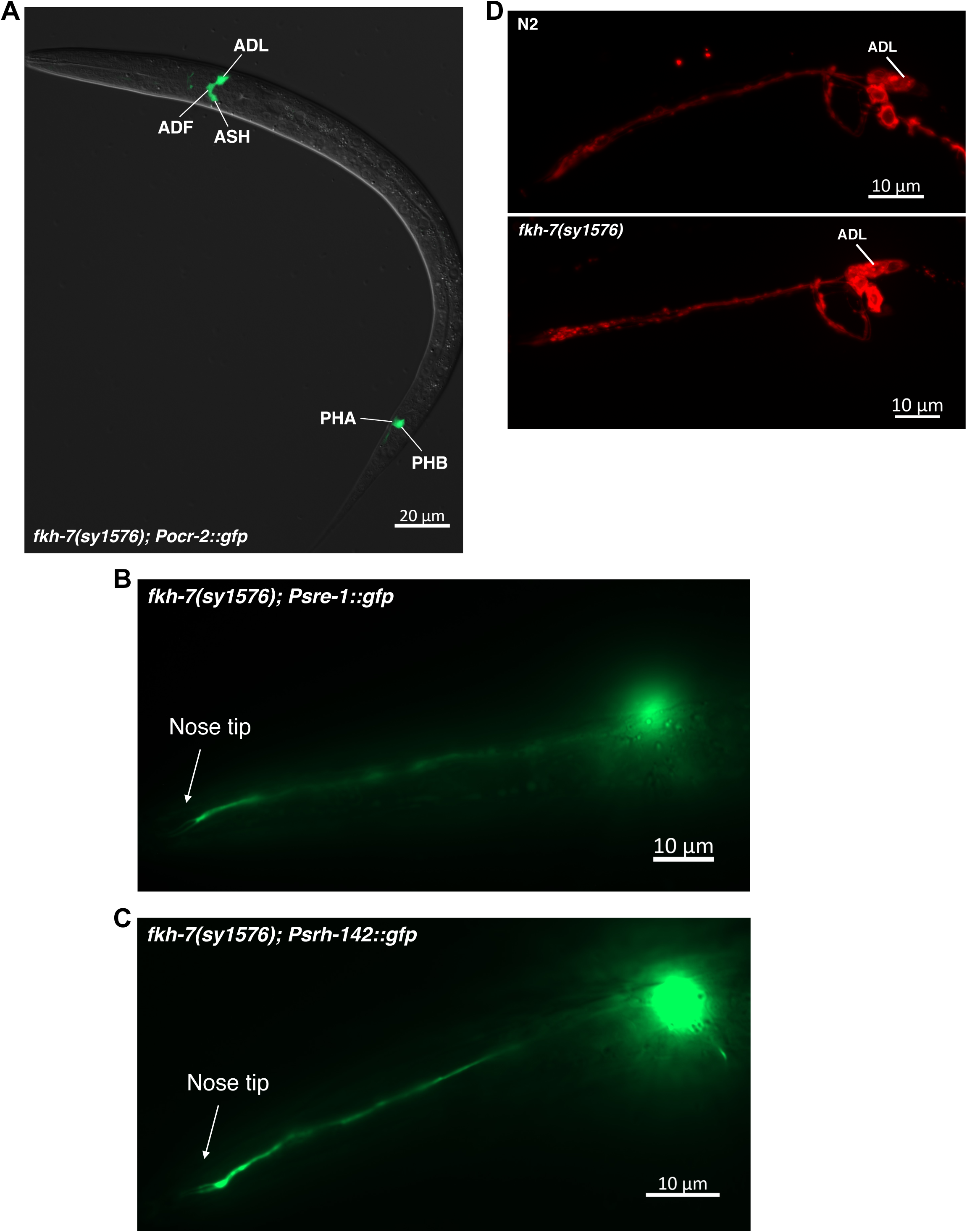
ADL and ADF neurons in *fkh-7(sy1576)* mutants do not exhibit aberrant cell body spatial localization or abnormal dendritic ending anatomy, and are not deficient in DiI fluorescent dye uptake. Related to Figure 2. (A) P*ocr-2*::*gfp* expression in a *fkh-7*(*sy1576*) mutant during the L1 stage. (B) Representative image of P*sre-1::gfp* (ADL) ciliated endings in an L2 stage *fkh-7*(*sy1576*) mutant. (C) Representative image of P*srh-142::gfp* (ADF) ciliated endings in an L2 stage *fkh-7*(*sy1576*) mutant. (D) DiI staining of amphid chemosensory neurons in N2 wild-type and *fkh-7*(*sy1576*) L2 stage animals.

**Figure S3.**
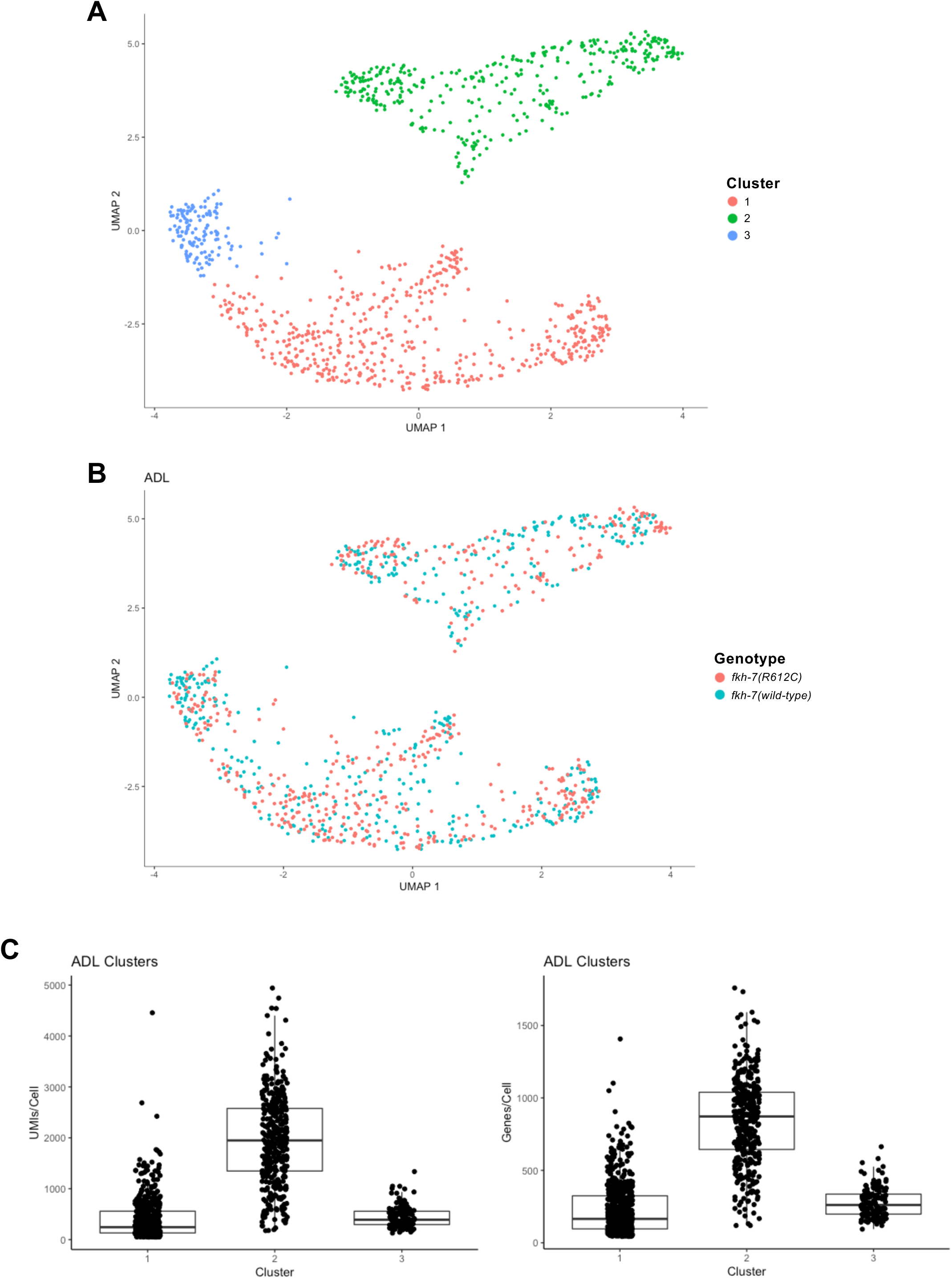
ADL cells form three distinct clusters driven by differences in sequencing depth. Related to Figure 4. (A) ADL cells form three distinct clusters. (B) Distribution of *fkh-7(R612C)* and *fkh-7(wild-type)* ADL cells overlap in all clusters. (C) Median Unique Molecular Identifiers (UMIs) and genes per ADL cell varies between clusters.

**Figure S4.**
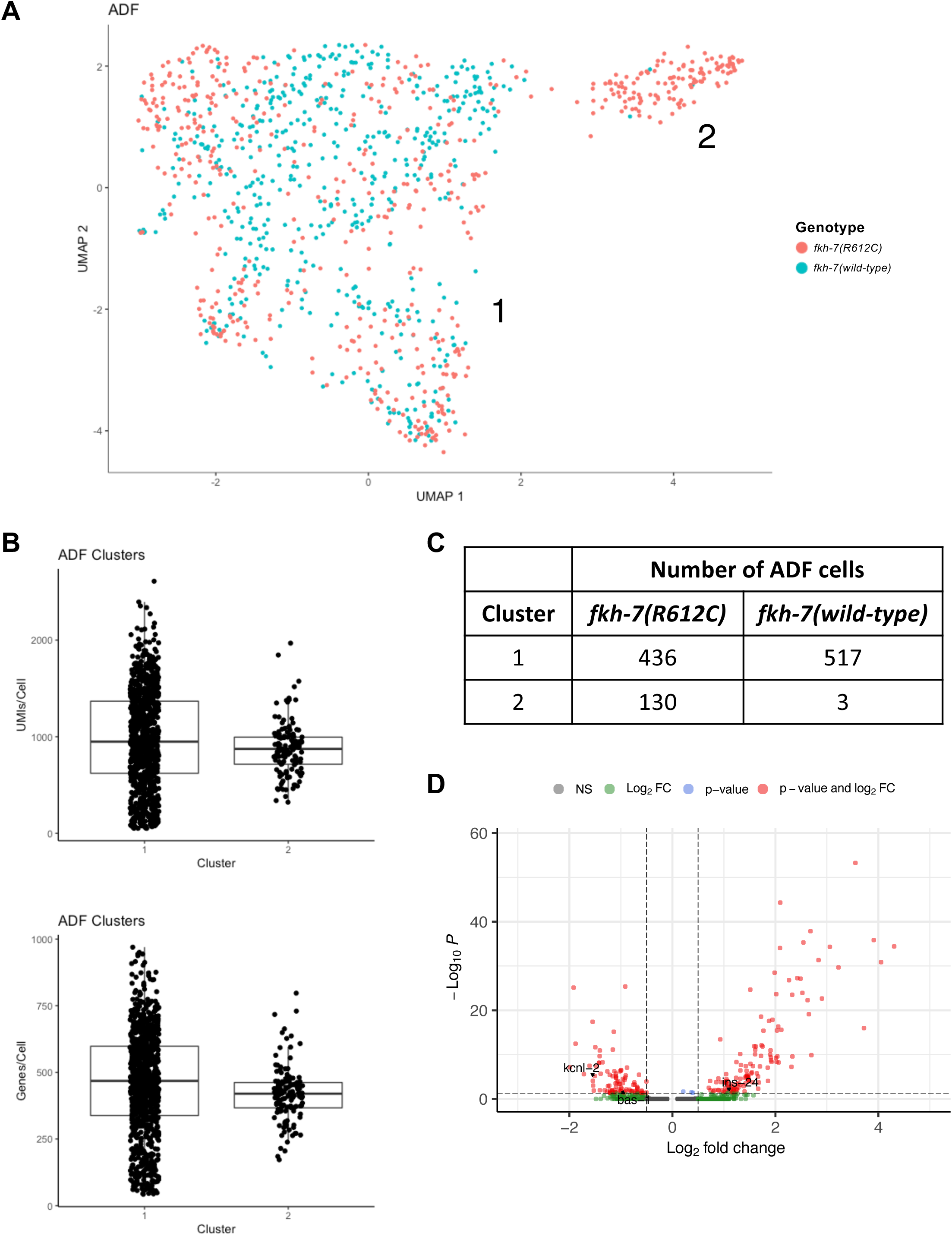
ADF cells form two distinct clusters driven by genotype. Related to Figure 4. (A) ADF cells form two distinct clusters. Cluster 1 (left) is comprised of ADF cells from both genotypes and Cluster 2 (right) is comprised mostly of *fkh-7(R612C)* ADF cells. (B) Median Unique Molecular Identifiers (UMIs) and genes per ADF cell do not vary between clusters. (C) Number of *fkh-7(R612C)* and *fkh-7(wild-type)* ADF cells in each cluster. (D) Volcano plot for differential gene expression analysis between Cluster 2 *fkh-7(R612C)* cells and Cluster 1 *fkh-7(wild-type)* cells (thresholds: Log2FC>|0.5| and adjusted p-value<0.05). *kcnl-*2 (Log2FC=-1.51, p-adjusted=6.57E-06), *ins-24* (Log2FC=1.06, p-adjusted=0.012), and *bas-1* (Log2FC=-1.00, p-adjusted=0.018) genes labeled.

## SUPPLEMENTAL TABLES AND DATA

**Table S1.**
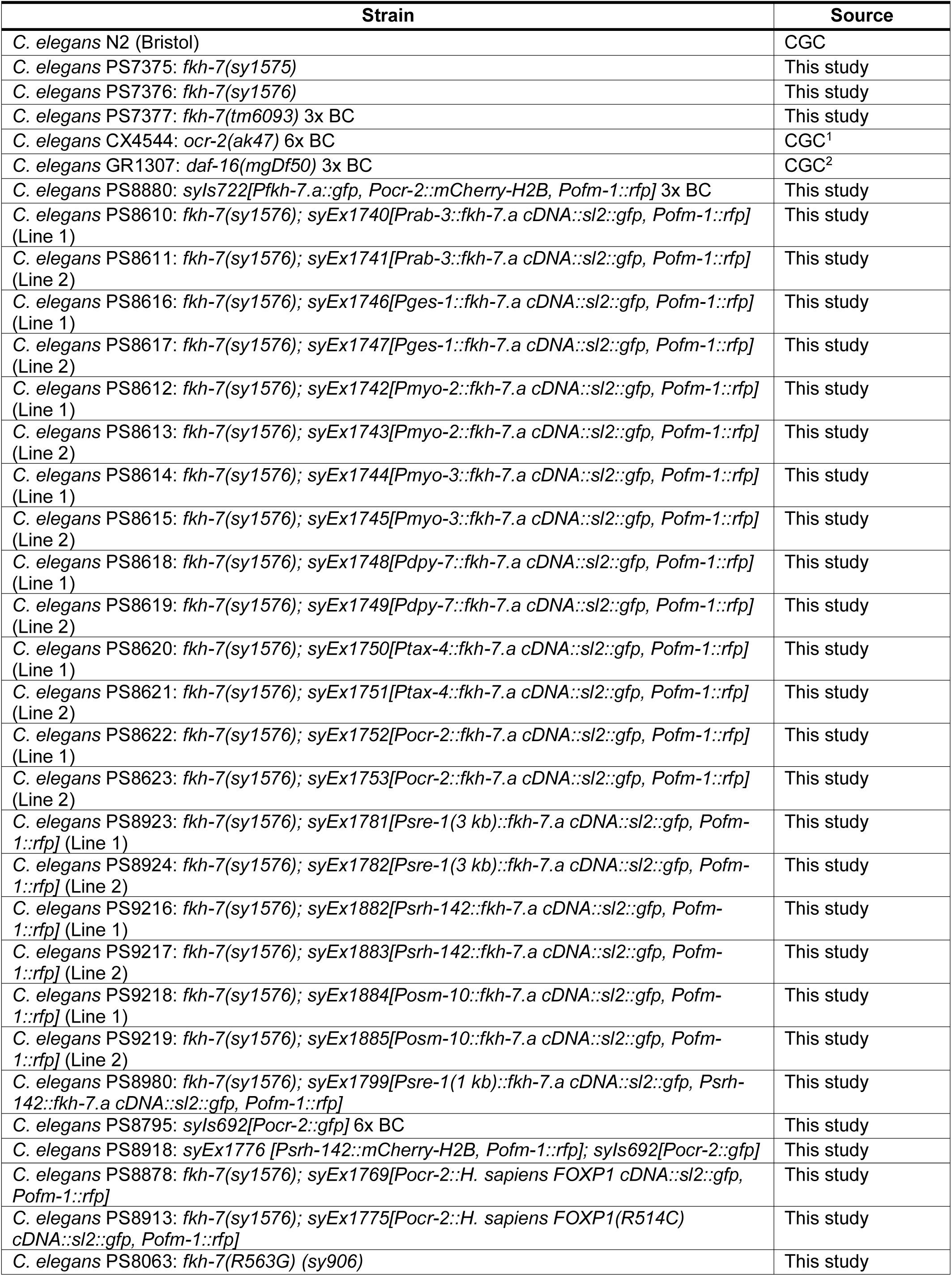
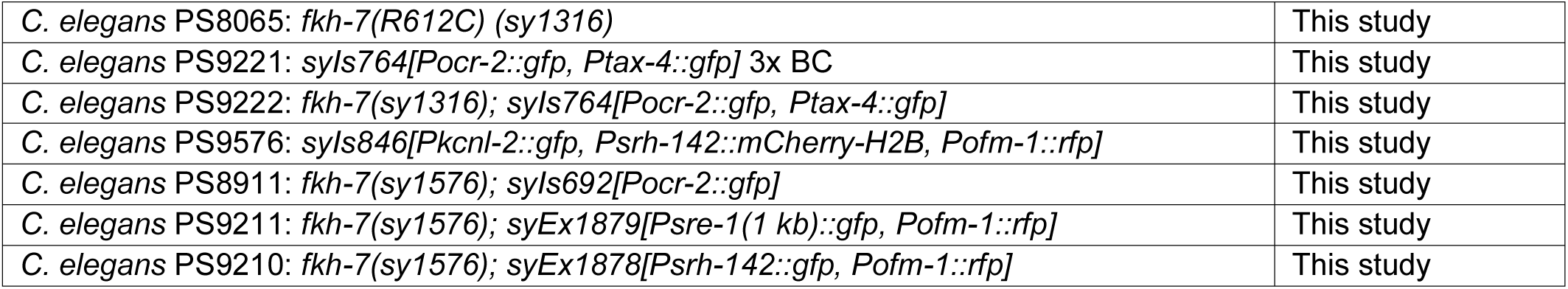
Details of all *Caenorhabditis elegans* strains used in this study.

**Table S2.** All plasmids generated in this study.

| Recombinant DNA |
| --- |
| PSM_Pfkh-7.a::gfp |
| PSM_Prab-3::fkh-7.a cDNA::sl2::gfp |
| PSM_Pges-1::fkh-7.a cDNA::sl2::gfp |
| PSM_Pmyo-2::fkh-7.a cDNA::sl2::gfp |
| PSM_Pmyo-3::fkh-7.a cDNA::sl2::gfp |
| PSM_Pdpy-7::fkh-7.a cDNA::sl2::gfp |
| PSM_Ptax-4::fkh-7.a cDNA::sl2::gfp |
| PSM_Pocr-2::fkh-7.a cDNA::sl2::gfp |
| PSM_Psre-1(3 kb)::fkh-7.a cDNA::sl2::gfp |
| PSM_Psrh-142::fkh-7.a cDNA::sl2::gfp |
| PSM_Posm-10::fkh-7.a cDNA::sl2::gfp |
| PSM_Psre-1(1 kb)::fkh-7.a cDNA::sl2::gfp |
| PSM_Pocr-2::gfp |
| PSM_Ptax-4::gfp |
| PSM_Pocr-2::mCherry-H2B |
| PSM_Psrh-142::mCherry-H2B |
| PSM_Pocr-2::H. sapiens FOXP1 cDNA::sl2::gfp |
| PSM_Pocr-2::H. sapiens FOXP1(R514C) cDNA::sl2::gfp |
| PSM_Pkcnl-2::gfp |
| PSM_Psre-1(1 kb)::gfp |
| PSM_Psrh-142::gfp |
| pRB1017_fkh-7(deletion)_sgRNA1 |
| pRB1017_fkh-7(deletion)_sgRNA2 |
| pRB1017_fkh-7(deletion)_sgRNA3 |
| pRB1017_fkh-7(deletion)_sgRNA4 |
| pRB1017_fkh-7(deletion)_sgRNA5 |

**Table S3.**
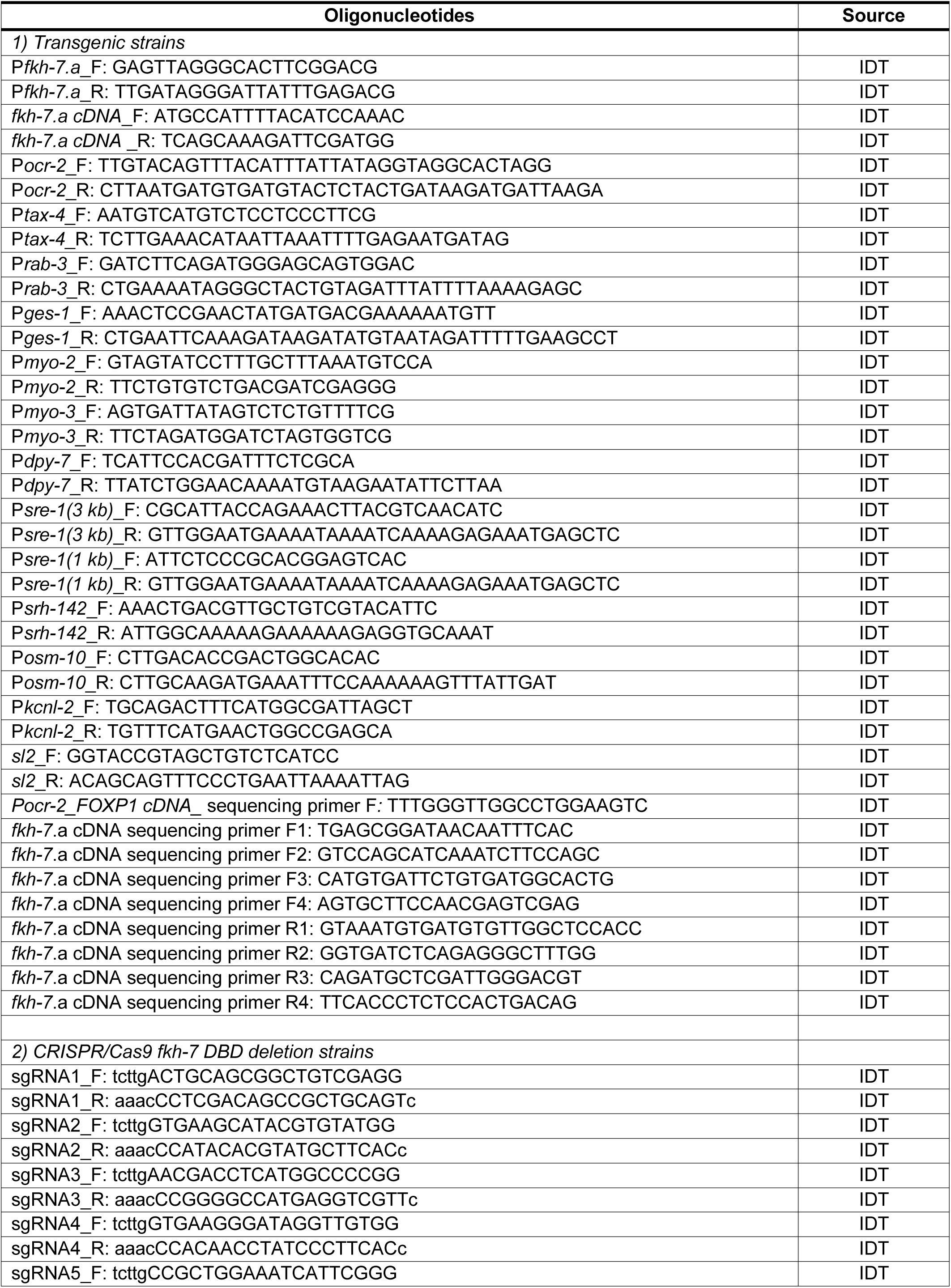
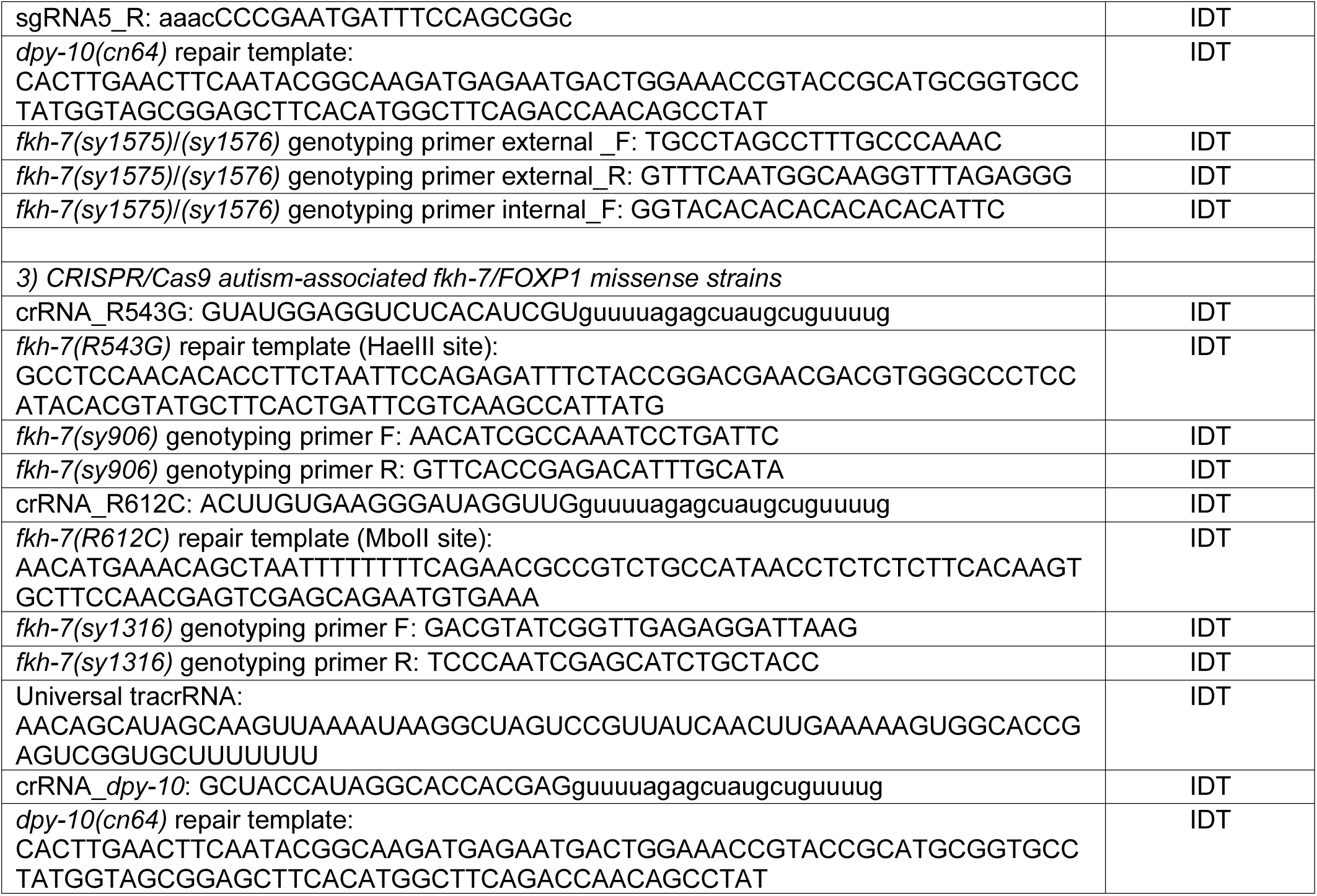
All oligonucleotides used in this study.

**Table S4.**
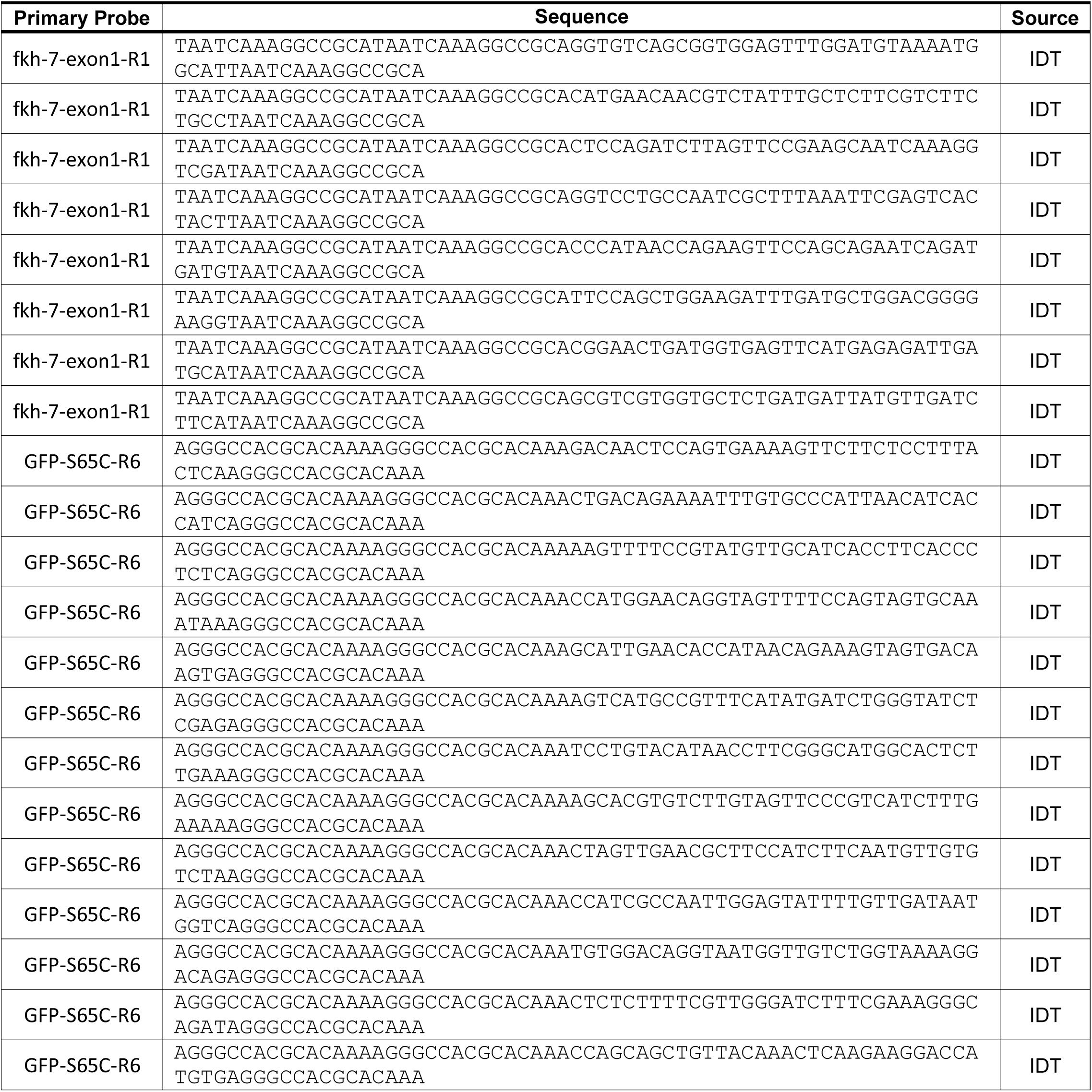
smFISH primary probe sequences.

**Data S1. Table of dauer formation assay raw data and p-values.**

**Data S2. Caenorhabditis elegans fkh-7.a cDNA sequence.**

**Data S3. Homo sapiens FOXP1 and FOXP1(R514C) cDNA sequences.**

**Data S4. Single-cell RNA-seq ADL gene list table.**

**Data S5. Single-cell RNA-seq ADF gene list table.**

## Notes

### Competing Interest Statement

The authors have declared no competing interest.

