## Supplemental Data S1 for "The forkhead transcription factor FKH-7/FOXP acts in chemosensory neurons to regulate developmental decision-making"

**Data S1. Table of dauer formation assay raw data and p-values.**

**Figure 1C**

| Strain Genotype | # of Dauers | # of non-Dauers | Total | %Dauers | DFI |
|---|---|---|---|---|---|
| N2 | 40 | 28 | 68 | 0.58823529 | 0.17647059 |
| N2 | 43 | 23 | 66 | 0.65151515 | 0.30303030 |
| N2 | 47 | 23 | 70 | 0.67142857 | 0.34285714 |
| N2 | 43 | 24 | 67 | 0.64179104 | 0.28358209 |
| N2 | 44 | 28 | 72 | 0.61111111 | 0.22222222 |
| fkh-7(sy1575) | 27 | 40 | 67 | 0.40298507 | -0.19402985 |
| fkh-7(sy1575) | 22 | 41 | 63 | 0.34920635 | -0.30158730 |
| fkh-7(sy1575) | 23 | 44 | 67 | 0.34328358 | -0.31343284 |
| fkh-7(sy1575) | 19 | 42 | 61 | 0.31147541 | -0.37704918 |
| fkh-7(sy1576) | 18 | 45 | 63 | 0.28571429 | -0.42857143 |
| fkh-7(sy1576) | 26 | 41 | 67 | 0.3880597 | -0.22388060 |
| fkh-7(sy1576) | 18 | 47 | 65 | 0.27692308 | -0.44615385 |
| fkh-7(sy1576) | 11 | 58 | 69 | 0.15942029 | -0.68115942 |

| | one-way ANOVA followed by Dunnett's post-hoc compared to N2 |
|---|---|
| N2 | 1 |
| fkh-7(sy1575) | 3.5593829813551636e-05 |
| fkh-7(sy1576) | 4.180175477652703e-06 |

**Figure 1D**

| Strain Genotype | # of Dauers | # of non-Dauers | Total | %Dauers | DFI |
|---|---|---|---|---|---|
| N2 | 41 | 25 | 66 | 0.621212121 | 0.24242424 |
| N2 | 42 | 22 | 64 | 0.65625 | 0.3125 |
| N2 | 41 | 21 | 62 | 0.661290323 | 0.32258065 |
| fkh-7(tm6093) | 9 | 54 | 63 | 0.142857143 | -0.71428571 |
| fkh-7(tm6093) | 17 | 43 | 60 | 0.283333333 | -0.43333333 |
| fkh-7(tm6093) | 25 | 40 | 65 | 0.384615385 | -0.23076923 |

| | Welch's unpaired t-test |
|---|---|
| fkh-7(tm6093) | 0.02974673 |

**Figure 1E**

| Strain Genotype | # of Dauers | # of Partial Dauers | # of Reproductive Larvae | Total | %Dauers | %PartialDauers |
|---|---|---|---|---|---|---|
| N2 | 57 | | 11 | 68 | 0.83823529 | |
| N2 | 59 | | 11 | 70 | 0.84285714 | |
| N2 | 70 | | 3 | 73 | 0.95890411 | |
| N2 | 53 | | 9 | 62 | 0.85483871 | |
| fkh-7(sy1576) | 34 | | 26 | 60 | 0.56666667 | |
| fkh-7(sy1576) | 44 | | 19 | 63 | 0.69841270 | |
| fkh-7(sy1576) | 32 | | 31 | 63 | 0.50793651 | |
| fkh-7(sy1576) | 42 | | 25 | 67 | 0.62686567 | |
| daf-16(mgDf50) | | 15 | 52 | 67 | | 0.22388060 |
| daf-16(mgDf50) | | 23 | 40 | 63 | | 0.36507937 |
| daf-16(mgDf50) | | 22 | 46 | 68 | | 0.32352941 |
| daf-16(mgDf50) | | 16 | 41 | 57 | | 0.28070175 |

**Figure 2A**

| Strain Genotype | # of Dauers | # of non-Dauers | Total | %Dauers | DFI |
|---|---|---|---|---|---|
| N2 | 86 | 32 | 118 | 0.72881356 | 0.45762712 |
| N2 | 97 | 24 | 121 | 0.80165289 | 0.60330579 |
| N2 | 85 | 37 | 122 | 0.69672131 | 0.39344262 |
| N2 | 77 | 39 | 116 | 0.66379310 | 0.32758621 |
| fkh-7(sy1576) | 67 | 63 | 130 | 0.51538462 | 0.03076923 |
| fkh-7(sy1576) | 60 | 51 | 111 | 0.54054054 | 0.08108108 |
| fkh-7(sy1576) | 60 | 54 | 114 | 0.52631579 | 0.05263158 |
| fkh-7(sy1576) | 47 | 54 | 101 | 0.46534653 | -0.06930693 |
| fkh-7(sy1576) | 48 | 64 | 112 | 0.42857143 | -0.14285714 |
| Prab-3 rescue Line 1 | 65 | 18 | 83 | 0.78313253 | 0.56626506 |
| Prab-3 rescue Line 1 | 66 | 42 | 108 | 0.61111111 | 0.22222222 |
| Prab-3 rescue Line 1 | 48 | 57 | 105 | 0.45714286 | -0.08571429 |
| Prab-3 rescue Line 2 | 66 | 34 | 100 | 0.66000000 | 0.32000000 |
| Prab-3 rescue Line 2 | 84 | 28 | 112 | 0.75000000 | 0.50000000 |
| Prab-3 rescue Line 2 | 55 | 35 | 90 | 0.61111111 | 0.22222222 |
| Prab-3 rescue Line 2 | 61 | 28 | 89 | 0.68539326 | 0.37078652 |
| Pges-1 rescue Line 1 | 73 | 38 | 111 | 0.65765766 | 0.31531532 |
| Pges-1 rescue Line 1 | 56 | 43 | 99 | 0.56565657 | 0.13131313 |
| Pges-1 rescue Line 1 | 53 | 49 | 102 | 0.51960784 | 0.03921569 |
| Pges-1 rescue Line 1 | 32 | 60 | 92 | 0.34782609 | -0.30434783 |
| Pges-1 rescue Line 2 | 72 | 38 | 110 | 0.65454545 | 0.30909091 |
| Pges-1 rescue Line 2 | 78 | 37 | 115 | 0.67826087 | 0.35652174 |
| Pges-1 rescue Line 2 | 83 | 28 | 111 | 0.74774775 | 0.49549550 |
| Pges-1 rescue Line 2 | 55 | 50 | 105 | 0.52380952 | 0.04761905 |
| Pmyo-3 rescue Line 1 | 68 | 43 | 111 | 0.61261261 | 0.22522523 |
| Pmyo-3 rescue Line 1 | 48 | 64 | 112 | 0.42857143 | -0.14285714 |
| Pmyo-3 rescue Line 1 | 40 | 67 | 107 | 0.37383178 | -0.25233645 |
| Pmyo-3 rescue Line 1 | 19 | 77 | 96 | 0.19791667 | -0.60416667 |
| Pmyo-3 rescue Line 2 | 54 | 59 | 113 | 0.47787611 | -0.04424779 |
| Pmyo-3 rescue Line 2 | 59 | 53 | 112 | 0.52678571 | 0.05357143 |
| Pmyo-3 rescue Line 2 | 41 | 68 | 109 | 0.37614679 | -0.24770642 |
| Pmyo-3 rescue Line 2 | 43 | 65 | 108 | 0.39814815 | -0.20370370 |
| Pmyo-2 rescue Line 1 | 35 | 61 | 96 | 0.36458333 | -0.27083333 |
| Pmyo-2 rescue Line 1 | 51 | 60 | 111 | 0.45945946 | -0.08108108 |
| Pmyo-2 rescue Line 1 | 44 | 70 | 114 | 0.38596491 | -0.22807018 |
| Pmyo-2 rescue Line 2 | 55 | 54 | 109 | 0.50458716 | 0.00917431 |
| Pmyo-2 rescue Line 2 | 57 | 47 | 104 | 0.54807692 | 0.09615385 |
| Pmyo-2 rescue Line 2 | 36 | 66 | 102 | 0.35294118 | -0.29411765 |
| Pmyo-2 rescue Line 2 | 31 | 76 | 107 | 0.28971963 | -0.42056075 |
| N2 | 77 | 30 | 107 | 0.71962617 | 0.43925234 |
| N2 | 85 | 26 | 111 | 0.76576577 | 0.53153153 |
| N2 | 66 | 46 | 112 | 0.58928571 | 0.17857143 |
| N2 | 63 | 51 | 114 | 0.55263158 | 0.10526316 |

| | | | | | |
|---|---|---|---|---|---|
| fkh-7(sy1576) | 44 | 71 | 115 | 0.38260870 | -0.23478261 |
| fkh-7(sy1576) | 60 | 50 | 110 | 0.54545455 | 0.09090909 |
| fkh-7(sy1576) | 53 | 55 | 108 | 0.49074074 | -0.01851852 |
| fkh-7(sy1576) | 50 | 64 | 114 | 0.43859649 | -0.12280702 |
| Pdpy-7 rescue Line 1 | 49 | 58 | 107 | 0.45794393 | -0.08411215 |
| Pdpy-7 rescue Line 1 | 54 | 57 | 111 | 0.48648649 | -0.02702703 |
| Pdpy-7 rescue Line 1 | 36 | 62 | 98 | 0.36734694 | -0.26530612 |
| Pdpy-7 rescue Line 1 | 49 | 47 | 96 | 0.51041667 | 0.02083333 |
| Pdpy-7 rescue Line 2 | 68 | 37 | 105 | 0.64761905 | 0.29523810 |
| Pdpy-7 rescue Line 2 | 51 | 52 | 103 | 0.49514563 | -0.00970874 |
| Pdpy-7 rescue Line 2 | 25 | 68 | 93 | 0.26881720 | -0.46236559 |
| Pdpy-7 rescue Line 2 | 48 | 59 | 107 | 0.44859813 | -0.10280374 |

one-way ANOVA followed by Dunnett's post-hoc compared to N2

| | |
|---|---|
| N2 | 1 |
| fkh-7(sy1576) | 0.019432758348377543 |
| rab-3_Line1 | 0.7048435967693896 |
| rab-3_Line2 | 0.9934748247280665 |
| ges-1_Line1 | 0.06753166 |
| ges-1_Line2 | 0.9143818057220862 |
| myo-3_Line1 | 0.00113487 |
| myo-3_Line2 | 0.00519267 |
| myo-2_Line1 | 0.00273582 |
| myo-2_Line2 | 0.00245213 |

one-way ANOVA followed by Dunnett's post-hoc compared to N2

| | |
|---|---|
| N2 | 1 |
| fkh-7(sy1576) | 0.056984386765555506 |
| dpy-7_Line1 | 0.046170844901738284 |
| dpy-7_Line2 | 0.0579122 |

**Figure 2B**

| Strain Genotype | # of Dauers | # of non-Dauers | Total | Transgenic Dauers | Transgenic Non-Dauers | Transgenic Total | %Dauers | DFI |
|---|---|---|---|---|---|---|---|---|
| N2 | 77 | 30 | 107 | | | | 0.71962617 | 0.43925234 |
| N2 | 85 | 26 | 111 | | | | 0.76576577 | 0.53153153 |
| N2 | 66 | 46 | 112 | | | | 0.58928571 | 0.17857143 |
| N2 | 63 | 51 | 114 | | | | 0.55263158 | 0.10526316 |
| fkh-7(sy1576) | 44 | 71 | 115 | | | | 0.38260870 | -0.23478261 |
| fkh-7(sy1576) | 60 | 50 | 110 | | | | 0.54545455 | 0.09090909 |
| fkh-7(sy1576) | 53 | 55 | 108 | | | | 0.49074074 | -0.01851852 |
| fkh-7(sy1576) | 50 | 64 | 114 | | | | 0.43859649 | -0.12280702 |
| Ptax-4 rescue Line 1 | 49 | 61 | 110 | 42 | 59 | 101 | 0.41584158 | -0.16831683 |
| Ptax-4 rescue Line 1 | 40 | 70 | 110 | 33 | 67 | 100 | 0.33000000 | -0.34000000 |
| Ptax-4 rescue Line 1 | 41 | 71 | 112 | 38 | 68 | 106 | 0.35849057 | -0.28301887 |
| Ptax-4 rescue Line 1 | 29 | 71 | 100 | 27 | 71 | 98 | 0.27551020 | -0.44897959 |
| Ptax-4 rescue Line 2 | 47 | 54 | 101 | 28 | 31 | 59 | 0.47457627 | -0.05084746 |
| Ptax-4 rescue Line 2 | 50 | 50 | 100 | 32 | 41 | 73 | 0.43835616 | -0.12328767 |
| Ptax-4 rescue Line 2 | 52 | 54 | 106 | 30 | 32 | 62 | 0.48387097 | -0.03225807 |
| Ptax-4 rescue Line 2 | 52 | 55 | 107 | 26 | 39 | 65 | 0.40000000 | -0.20000000 |
| Pocr-2 rescue Line 1 | 85 | 35 | 120 | 52 | 10 | 62 | 0.83870968 | 0.67741936 |
| Pocr-2 rescue Line 1 | 65 | 48 | 113 | 37 | 14 | 51 | 0.72549020 | 0.45098039 |
| Pocr-2 rescue Line 1 | 63 | 50 | 113 | 41 | 26 | 67 | 0.61194030 | 0.22388060 |
| Pocr-2 rescue Line 1 | 81 | 36 | 117 | 58 | 23 | 81 | 0.71604938 | 0.43209877 |
| Pocr-2 rescue Line 2 | 58 | 46 | 104 | 38 | 26 | 64 | 0.59375000 | 0.18750000 |
| Pocr-2 rescue Line 2 | 71 | 37 | 108 | 47 | 18 | 65 | 0.72307692 | 0.44615385 |
| Pocr-2 rescue Line 2 | 58 | 59 | 117 | 40 | 35 | 75 | 0.53333333 | 0.06666670 |
| Pocr-2 rescue Line 2 | 65 | 44 | 109 | 53 | 30 | 83 | 0.63855422 | 0.27710840 |

one-way ANOVA followed by Dunnett's post-hoc compared to N2

| | |
|---|---|
| Transgenic | |
| N2 | 1 |
| fkh-7(sy1576) | 0.00954084 |
| tax-4_Line1 | 0.00012502 |
| tax-4_Line2 | 0.00519388 |
| ocr-2_Line1 | 0.6261690404456011 |
| ocr-2_Line2 | 0.94912165 |
| All | |
| N2 | 1 |
| fkh-7 | 0.0051726 |
| tax-4_1a | 7.205943279742222e-05 |
| tax-4_1b | 0.012941809697378348 |
| ocr-2_3a | 0.9858718417600434 |
| ocr-2_4a | 0.39150892072311205 |

**Figure 2D_ADL**

| Strain Genotype | # of Dauers | # of non-Dauers | Total | %Dauers | DFI |
|---|---|---|---|---|---|
| N2 | 92 | 17 | 109 | 0.84403670 | 0.68807339 |
| N2 | 93 | 21 | 114 | 0.81578947 | 0.63157895 |
| N2 | 80 | 34 | 114 | 0.70175439 | 0.40350877 |
| N2 | 89 | 40 | 129 | 0.68992248 | 0.37984496 |
| fkh-7(sy1576) | 69 | 36 | 105 | 0.65714286 | 0.31428571 |
| fkh-7(sy1576) | 42 | 60 | 102 | 0.41176471 | -0.17647059 |
| fkh-7(sy1576) | 59 | 55 | 114 | 0.51754386 | 0.03508772 |
| fkh-7(sy1576) | 71 | 51 | 122 | 0.58196721 | 0.16393443 |
| fkh-7(sy1576) | 54 | 45 | 99 | 0.54545455 | 0.09090909 |
| fkh-7(sy1576) | 59 | 54 | 113 | 0.52212389 | 0.04424779 |
| Psre-1(3kb) rescue Line 1 | 27 | 66 | 93 | 0.29032258 | -0.41935484 |
| Psre-1(3kb) rescue Line 1 | 33 | 92 | 110 | 0.30000000 | -0.53636364 |
| Psre-1(3kb) rescue Line 1 | 70 | 35 | 105 | 0.66666667 | 0.33333333 |
| Psre-1(3kb) rescue Line 1 | 52 | 52 | 104 | 0.50000000 | 0.00000000 |
| Psre-1(3kb) rescue Line 2 | 51 | 50 | 101 | 0.50495050 | 0.00990099 |
| Psre-1(3kb) rescue Line 2 | 27 | 73 | 100 | 0.27000000 | -0.46000000 |
| Psre-1(3kb) rescue Line 2 | 75 | 42 | 117 | 0.64102564 | 0.28205128 |

| | one-way ANOVA followed by Dunnett's post-hoc compared to N2 |
|---|---|
| N2 | 1 |
| fkh-7(sy1576) | 0.06109049 |
| sre-1(3kb)_Line1 | 0.00945892 |
| sre-1(3kb)_Line2 | 0.03832476 |

**Figure 2D_ADF**

| Strain Genotype | # of Dauers | # of non-Dauers | Total | %Dauers | DFI |
|---|---|---|---|---|---|
| N2 | 72 | 26 | 98 | 0.73469388 | 0.46938776 |
| N2 | 99 | 16 | 115 | 0.86086957 | 0.72173913 |
| N2 | 67 | 33 | 100 | 0.67000000 | 0.34000000 |
| N2 | 81 | 26 | 107 | 0.75700935 | 0.51401869 |
| fkh-7(sy1576) | 66 | 49 | 115 | 0.57391304 | 0.14782609 |
| fkh-7(sy1576) | 51 | 57 | 108 | 0.47222222 | -0.05555556 |
| fkh-7(sy1576) | 58 | 60 | 118 | 0.49152542 | -0.01694915 |
| fkh-7(sy1576) | 48 | 64 | 112 | 0.42857143 | -0.14285714 |
| Psrh-142 rescue Line 1 | 70 | 46 | 116 | 0.60344828 | 0.20689655 |
| Psrh-142 rescue Line 1 | 61 | 62 | 123 | 0.49593496 | -0.00813008 |
| Psrh-142 rescue Line 1 | 36 | 70 | 106 | 0.33962264 | -0.32075472 |
| Psrh-142 rescue Line 1 | 23 | 91 | 114 | 0.20175439 | -0.59649123 |
| Psrh-142 rescue Line 2 | 32 | 52 | 84 | 0.38095238 | -0.23809524 |
| Psrh-142 rescue Line 2 | 41 | 56 | 97 | 0.42268041 | -0.15463918 |
| Psrh-142 rescue Line 2 | 57 | 57 | 114 | 0.50000000 | 0.00000000 |
| Psrh-142 rescue Line 2 | 37 | 76 | 113 | 0.32743363 | -0.34513274 |

one-way ANOVA followed by Dunnett's post-hoc compared to N2

| | |
|---|---|
| N2 | 1 |
| fkh-7(sy1576) | 0.012143309446961292 |
| srh-142_Line1 | 0.00180998 |
| srh-142_Line2 | 0.00179335 |

**Figure 2D_ASH+PHA+PHB**

| Strain Genotype | # of Dauers | # of non-Dauers | Total | %Dauers | DFI |
|---|---|---|---|---|---|
| N2 | 83 | 29 | 112 | 0.74107143 | 0.48214286 |
| N2 | 80 | 30 | 110 | 0.72727273 | 0.45454546 |
| N2 | 84 | 37 | 121 | 0.69421488 | 0.38842975 |
| N2 | 91 | 20 | 111 | 0.81981982 | 0.63963964 |
| fkh-7(sy1576) | 45 | 46 | 91 | 0.49450549 | -0.01098901 |
| fkh-7(sy1576) | 47 | 51 | 98 | 0.47959184 | -0.04081633 |
| fkh-7(sy1576) | 52 | 53 | 105 | 0.49523810 | -0.00952381 |
| fkh-7(sy1576) | 64 | 62 | 126 | 0.50793651 | 0.01587302 |
| Posm-10 rescue Line 1 | 42 | 63 | 105 | 0.40000000 | -0.20000000 |
| Posm-10 rescue Line 1 | 54 | 67 | 121 | 0.44628099 | -0.10743802 |
| Posm-10 rescue Line 1 | 52 | 78 | 130 | 0.40000000 | -0.20000000 |
| Posm-10 rescue Line 1 | 32 | 79 | 111 | 0.28828829 | -0.42342342 |
| Posm-10 rescue Line 2 | 56 | 62 | 118 | 0.47457627 | -0.05084746 |
| Posm-10 rescue Line 2 | 37 | 56 | 93 | 0.39784946 | -0.20430108 |
| Posm-10 rescue Line 2 | 34 | 69 | 103 | 0.33009709 | -0.33980583 |
| Posm-10 rescue Line 2 | 22 | 75 | 97 | 0.22680412 | -0.54639175 |

one-way ANOVA followed by Dunnett's post-hoc compared to N2

| | |
|---|---|
| N2 | 1 |
| fkh-7(sy1576) | 0.00054605 |
| osm-10_Line1 | 2.1408082660490102e-05 |
| osm-10_Line2 | 5.121226092419384e-06 |

**Figure 2E**

| Strain Genotype | # of Dauers | # of non-Dauers | Total | Transgenic Dauers | Transgenic Non-Dauers | Transgenic Total | %Dauers | DFI |
|---|---|---|---|---|---|---|---|---|
| N2 | 80 | 34 | 114 | | | | 0.70175439 | 0.40350877 |
| N2 | 89 | 40 | 129 | | | | 0.68992248 | 0.37984496 |
| N2 | 111 | 32 | 143 | | | | 0.77622378 | 0.55244755 |
| N2 | 91 | 23 | 114 | | | | 0.79824561 | 0.59649123 |
| fkh-7(sy1576) | 54 | 45 | 99 | | | | 0.54545455 | 0.09090909 |
| fkh-7(sy1576) | 59 | 54 | 113 | | | | 0.52212389 | 0.04424779 |
| fkh-7(sy1576) | 34 | 78 | 112 | | | | 0.30357143 | -0.39285714 |
| fkh-7(sy1576) | 43 | 75 | 118 | | | | 0.36440678 | -0.27118644 |
| ADL+ADF rescue | 65 | 28 | 93 | 41 | 9 | 50 | 0.82000000 | 0.64000000 |
| ADL+ADF rescue | 78 | 38 | 116 | 67 | 28 | 95 | 0.70526316 | 0.41052632 |
| ADL+ADF rescue | 71 | 45 | 116 | 55 | 22 | 77 | 0.71428571 | 0.42857143 |

one-way ANOVA followed by Dunnett's post-hoc compared to N2

| Transgenic | |
|---|---|
| N2 | 1 |
| fkh-7(sy1576) | 0.00193053 |
| ADL+ADF rescue | 0.99586255 |

| All | |
|---|---|
| N2 | 1 |
| fkh-7(71) | 0.00146962 |
| ADL+ADF | 0.38515619 |

**Figure 3B**

| Strain Genotype | # of Dauers | # of non-Dauers | Total | %Dauers | DFI |
|---|---|---|---|---|---|
| N2 | 85 | 22 | 107 | 0.79439252 | 0.58878505 |
| N2 | 71 | 40 | 111 | 0.63963964 | 0.27927928 |
| N2 | 80 | 34 | 114 | 0.70175439 | 0.40350877 |
| N2 | 89 | 40 | 129 | 0.68992248 | 0.37984496 |
| fkh-7(sy1576) | 53 | 43 | 96 | 0.55208333 | 0.10416667 |
| fkh-7(sy1576) | 59 | 55 | 114 | 0.51754386 | 0.03508772 |
| fkh-7(sy1576) | 71 | 51 | 122 | 0.58196721 | 0.16393443 |
| fkh-7(sy1576) | 54 | 45 | 99 | 0.54545455 | 0.09090909 |
| fkh-7(sy1576) | 59 | 54 | 113 | 0.52212389 | 0.04424779 |
| Pocr-2::FOXP1 rescue | 86 | 19 | 105 | 0.81904762 | 0.63809524 |
| Pocr-2::FOXP1 rescue | 69 | 32 | 101 | 0.68316832 | 0.36633663 |
| Pocr-2::FOXP1 rescue | 88 | 25 | 113 | 0.77876106 | 0.55752212 |
| Pocr-2::FOXP1 rescue | 63 | 53 | 116 | 0.54310345 | 0.08620690 |
| Pocr-2::FOXP1(R514C) rescue | 59 | 80 | 139 | 0.42446043 | -0.15107914 |
| Pocr-2::FOXP1(R514C) rescue | 39 | 71 | 110 | 0.35454545 | -0.29090909 |
| Pocr-2::FOXP1(R514C) rescue | 31 | 72 | 103 | 0.30097087 | -0.39805825 |
| Pocr-2::FOXP1(R514C) rescue | 47 | 60 | 107 | 0.43925234 | -0.12149533 |

one-way ANOVA followed by Dunnett's post-hoc compared to N2

| | |
|---|---|
| N2 | 1 |
| fkh-7(sy1576) | 0.0167612 |
| Pocr-2::FOXP1 rescue | 0.9999996434486182 |
| Pocr-2::FOXP1(R514C) rescue | 8.924100792095935e-05 |

**Figure 3C**

| Strain Genotype | # of Dauers | # of non-Dauers | Total | %Dauers | DFI |
|---|---|---|---|---|---|
| N2 | 46 | 21 | 67 | 0.68656716 | 0.37313433 |
| N2 | 45 | 13 | 58 | 0.77586207 | 0.55172414 |
| N2 | 60 | 13 | 73 | 0.82191781 | 0.64383562 |
| N2 | 50 | 19 | 69 | 0.72463768 | 0.44927536 |
| N2 | 42 | 27 | 69 | 0.60869565 | 0.21739130 |
| N2 | 35 | 26 | 61 | 0.57377049 | 0.14754098 |
| N2 | 38 | 31 | 69 | 0.55072464 | 0.10144928 |
| N2 | 43 | 22 | 65 | 0.66153846 | 0.32307692 |
| fkh-7(R563G) | 33 | 27 | 60 | 0.55000000 | 0.10000000 |
| fkh-7(R563G) | 30 | 38 | 68 | 0.44117647 | -0.11764706 |
| fkh-7(R563G) | 35 | 25 | 60 | 0.58333333 | 0.16666667 |
| fkh-7(R563G) | 36 | 32 | 68 | 0.52941176 | 0.05882353 |
| fkh-7(R563G) | 31 | 46 | 77 | 0.40259740 | -0.19480520 |
| fkh-7(R563G) | 33 | 22 | 55 | 0.60000000 | 0.20000000 |
| fkh-7(R563G) | 19 | 39 | 58 | 0.32758621 | -0.34482759 |
| fkh-7(R563G) | 30 | 29 | 59 | 0.50847458 | 0.01694915 |

| | | | | | |
|---|---|---|---|---|---|
| N2 | 41 | 21 | 62 | 0.66129032 | 0.32258065 |
| N2 | 36 | 21 | 57 | 0.63157895 | 0.26315790 |
| N2 | 44 | 19 | 63 | 0.69841270 | 0.39682540 |
| N2 | 46 | 16 | 62 | 0.74193548 | 0.48387097 |
| N2 | 48 | 18 | 66 | 0.72727273 | 0.45454546 |
| N2 | 37 | 30 | 67 | 0.55223881 | 0.10447761 |
| fkh-7(R612C) | 23 | 37 | 60 | 0.38333333 | -0.23333333 |
| fkh-7(R612C) | 22 | 38 | 60 | 0.36666667 | -0.26666667 |
| fkh-7(R612C) | 30 | 30 | 60 | 0.50000000 | 0.00000000 |
| fkh-7(R612C) | 35 | 30 | 65 | 0.53846154 | 0.07692308 |
| fkh-7(R612C) | 23 | 41 | 64 | 0.35937500 | -0.28125000 |

| | Welch's unpaired t-test |
|---|---|
| fkh-7(R563G) | 0.001834977 |

| | Welch's unpaired t-test |
|---|---|
| fkh-7(R612C) | 0.000981263 |

**Figure S1D**

| Strain Genotype | # of Dauers | # of non-Dauers | Total | %Dauers | DFI |
|---|---|---|---|---|---|
| N2 | 36 | 31 | 67 | 0.53731343 | 0.07462687 |
| N2 | 41 | 26 | 67 | 0.61194030 | 0.22388060 |
| N2 | 42 | 28 | 70 | 0.60000000 | 0.20000000 |
| N2 | 45 | 27 | 72 | 0.62500000 | 0.25000000 |
| ocr-2(ak47) | 4 | 59 | 63 | 0.06349206 | -0.87301587 |
| ocr-2(ak47) | 1 | 73 | 74 | 0.01351351 | -0.97297297 |
| ocr-2(ak47) | 2 | 61 | 63 | 0.03174603 | -0.93650794 |
| ocr-2(ak47) | 2 | 60 | 62 | 0.03225806 | -0.93548387 |

| | Welch's unpaired t-test |
|---|---|
| ocr-2(ak47) | 4.2097E-06 |
