## Supplemental Data S2 for "The forkhead transcription factor FKH-7/FOXP acts in chemosensory neurons to regulate developmental decision-making"

**Data S2. *Caenorhabditis elegans fkh-7*.a cDNA sequence.**

> *Caenorhabditis elegans fkh-7*.a WB_CDS_F26D12.1a (2337 bp)
ATGCCATTTTACATCCAAACTCCACCGCTGACACCAGCACAACATTTATCCTCAACATCTACAGTTACATCTC
CAACTAGTCAAAATAATCATTTATTGGCAGAAGACGAAGAGCAAATAGACGTTGTTCATGAAGAAGAGGAAG
AAGATATGGAAGAAATGAAAATGGAAATTGAAATGCAATCAAAGAGTTTACATCAGAAACCATCAGGATTACG
AAGAAGTTTTCGACCTTTGATTGCTTCGGAACTAAGATCTGGAGGAATTCATAAAACTAGTAGTGACTCGAA
TTTAAAGCGATTGGCAGGACCATCATCATCTGATTCTGCTGGAACTTCTGGTTATGGGTACCTTCCCCGTCC
AGCATCAAATCTTCCAGCTGGAATAAATCATGAATTCAAAATTGATCCACTTCAAACAGTTCAACAATTTCTT
CAAATGCATCAATCTCTCATGAACTCACCATCAGTTCCGACATGCCTTTCGAAACTTTTGATCGATGAAGATC
AACATAATCATCAGAGCACCACGACGCTGAACAACGTCCTGGCGCCGCCGATGATCACCTCTGAATCATTG
CCACGACCTACACAGGCTCAATCATTTGCCAACTCGACAATTGGGGGTGGAGCCAACACATCACATTTACC
AGTCACACTGGACACTGGAATCAACGATAATTTGATATTATTAGCGTTACAGGAAAGACTTATGCGTCCGCA
GCAACCAACTCTAACCCAATCTGCCACCACACCATCTCTGTCCCATTTGGCAGCTGCCACAAATTCCTCCA
CAAATGCCACGTCACTGGGCTCGGCGGACACAACTCGTCTCATTCAGCAAATGTTTCTGGGGCAGATGCT
GATTCCACAACTTCCAGCCGCGTTGGATTTGGCATCGACTATTCAGTCGGCTTCAATGTCACTTCCATTACA
AACGACTATACCTCAACCGATGAGTGCACCACAACATGCTCTCTGGCAGCATGGGATGTGTGCGTGGCCA
AGTTGTGATCAACCATGTGATTCTGTGATGGCACTGATCACACACTTGCAACATGAACATCCGCCGTCTGAT
AAATCCAACGAGGAAATGCGAAATCAAATCGAAAAAGTCGAGAGCATAGAGCACAAGTTGTCTGTCGAACG
GAGTAGGCTTCAAGGAATGATGCAACATCTTCGGATGAAGCCATCTCCTGACACAACTACCCCGAATCTTG
TGAAAATGGAGGCCCAAAGCCCTCTGAGATCACCTAAAATTGAGGGTGCTGCGTTTTCAATTCAACCAGCC
CAGCAGTTTCAACAACAGACGACTAGTCAGCCACAGGTTTCACCAACATCCGAAGCAGCATCTTCACTGCT
AAGCATCGCAGCTACCGTTGCCGCCTCGACAGCCGCTGCAGTTACCTCCCCGATCAATCAAATATCTACAG
TACCCTCTGTTAGTTCAATGCCTTCATTTCTGAATCATCAACTATCGACATCATCCCAACCGTCTTCTCAACA
AGCCAGTGGATCATCTGGACCACTTCTTCAACGGGCAGCTAGTTCGGCATCAACTGAAACATCGCCAAATC
CTGATTCTAAATCATTTGTGCCAAGAAGAAGTCGGATATCAGACAAGACAGTTCAACCAATTGCCACAGATA
TTGCAAAGAATCGAGATTTCTACCGGACCAACGATGTGAGACCTCCATACACGTATGCTTCACTGATTCGTC
AAGCCATTATGGAATCGTCAGATTGCCAACTAACACTGAATGAGATCTACACATGGTTCACCGAGACATTTG
CATACTTCAGAAGGAACGCTGCCACGTGGAAGAACGCCGTCCGCCACAACCTATCCCTTCACAAGTGCTT
CCAACGAGTCGAGCAGAATGTGAAAGGAGCTGTATGGACTGTAGACGATTCTGAATTTTATCGAAGACGTC
CCAATCGAGCATCTGCTACCAGAAGTCAGCCACAGACACCTTTGCCCGACGATATTTCGCAGCAAAAATTG
TTTGACACGAGCGCTTTGAGCTCATTTTTGGAAATGCAGAACTTCGACCCCGCATCGCTGACTGGTGATCA
ATTCCAGCTGAATGGCAATTTAGACAGTGTTTTGTCGCTTCTTGCAAACGCCGATGTGAACAATCCGTTGCA
AATGCTATCAGCTGCTGCGGCCGCTGGAAATCATTCGGGTGGTATTCTAGAAGGACAGTTGCTGAATAATG
TGAAAGAAGAGATGATGGATGTTAGCGAGCCACACAATGGACATTTATTGAGAGTGGCTCGGCACATTCAA
AAAGCTTCAAAGCGGCCTGCGTCAGCAAATCCATCGAATCTTTGCTGA
