## Supplemental Data S3 for "The forkhead transcription factor FKH-7/FOXP acts in chemosensory neurons to regulate developmental decision-making"

**Data S3. *Homo sapiens FOXP1* and *FOXP1(R514C)* cDNA sequences.**

> *Homo sapiens FOXP1* CCDS2914.1 (2034 bp)
ATGATGCAAGAATCTGGGACTGAGACAAAAAGTAACGGTTCAGCCATCCAGAATGGGTCGGGCGGCAGCA
ACCACTTACTAGAGTGCGGCGGTCTTCGGGAGGGGCGGTCCAACGGAGAGACGCCGGCCGTGGACATC
GGGGCAGCTGACCTCGCCCACGCCCAGCAGCAGCAGCAACAGGCACTTCAGGTGGCAAGACAGCTCCT
TCTTCAGCAGCAACAGCAGCAGCAAGTTAGTGGATTAAAATCTCCCAAGAGGAATGACAAACAACCAGCTC
TTCAGGTTCCCGTGTCAGTGGCTATGATGACACCTCAAGTTATCACTCCCCAGCAAATGCAGCAGATCCTC
CAGCAACAAGTGCTGAGCCCTCAGCAGCTCCAGGTTCTCCTCCAGCAGCAGCAGGCCCTCATGCTTCAA
CAGCAGCAGCTTCAAGAGTTTTATAAAAAACAACAGGAACAGTTGCAGCTTCAACTTTTACAACAACAACAT
GCTGGAAAACAGCCTAAAGAGCAACAGCAGGTGGCTACCCAGCAGTTGGCTTTTCAGCAGCAGCTTTTAC
AGATGCAGCAGTTACAGCAGCAGCACCTCCTGTCTTTGCAGCGCCAAGGCCTTCTGACAATTCAGCCCGG
GCAGCCTGCCCTTCCCCTTCAACCTCTTGCTCAAGGCATGATTCCAACAGAACTGCAGCAGCTCTGGAAA
GAAGTGACAAGTGCTCATACTGCAGAAGAAACCACAGGCAACAATCACAGCAGTTTGGATCTGACCACGA
CATGTGTCTCCTCCTCTGCACCTTCCAAGACCTCCTTAATAATGAACCCACATGCCTCTACCAATGGACAGC
TCTCAGTCCACACTCCCAAAAGGGAAAGTTTGTCCCATGAGGAGCACCCCCATAGCCATCCTCTCTATGGA
CATGGTGTATGCAAGTGGCCAGGCTGTGAAGCAGTGTGCGAAGATTTCCAATCATTTCTAAAACATCTCAAC
AGTGAGCATGCGCTGGACGATAGAAGTACAGCCCAATGTAGAGTACAAATGCAGGTTGTACAGCAGTTAGA
GCTACAGCTTGCAAAAGACAAAGAACGCCTGCAAGCCATGATGACCCACCTGCATGTGAAGTCTACAGAA
CCCAAAGCCGCCCCTCAGCCCTTGAATCTGGTATCAAGTGTCACTCTCTCCAAGTCCGCATCGGAGGCTT
CTCCACAGAGCTTACCTCATACTCCAACGACCCCAACCGCCCCCCTGACTCCCGTCACCCAAGGCCCCTC
TGTCATCACAACCACCAGCATGCACACGGTGGGACCCATCCGCAGGCGGTACTCAGACAAATACAACGTG
CCCATTTCGTCAGCAGATATTGCGCAGAACCAAGAATTTTATAAGAACGCAGAAGTTAGACCACCATTTACAT
ATGCATCTTTAATTAGGCAGGCCATTCTCGAATCTCCAGAAAAGCAGCTAACACTAAATGAGATCTATAACTG
GTTCACACGAATGTTTGCTTACTTCCGACGCAACGCGGCCACGTGGAAGAATGCAGTGCGTCATAATCTTA
GTCTTCACAAGTGTTTTGTGCGAGTAGAAAACGTTAAAGGGGCAGTATGGACAGTGGATGAAGTAGAATTC
CAAAAACGAAGGCCACAAAAGATCAGTGGTAACCCTTCCCTTATTAAAAACATGCAGAGCAGCCACGCCTA
CTGCACACCTCTCAATGCAGCTTTACAGGCTTCAATGGCTGAGAATAGTATACCTCTATACACTACCGCTTC
CATGGGAAATCCCACTCTGGGCAACTTAGCCAGCGCAATACGGGAAGAGCTGAACGGGGCAATGGAGCAT
ACCAACAGCAACGAGAGTGACAGCAGTCCAGGCAGATCTCCTATGCAAGCCGTGCATCCTGTACACGTCA
AAGAAGAGCCCCTCGATCCAGAGGAAGCTGAAGGGCCCCTGTCCTTAGTGACAACAGCCAACCACAGTC
CAGATTTTGACCATGACAGAGATTACGAAGATGAACCAGTAAACGAGGACATGGAGTGA

> *Homo sapiens FOXP1* CCDS2914.1 [1540C>T] (2034 bp)
ATGATGCAAGAATCTGGGACTGAGACAAAAAGTAACGGTTCAGCCATCCAGAATGGGTCGGGCGGCAGCA
ACCACTTACTAGAGTGCGGCGGTCTTCGGGAGGGGCGGTCCAACGGAGAGACGCCGGCCGTGGACATC
GGGGCAGCTGACCTCGCCCACGCCCAGCAGCAGCAGCAACAGGCACTTCAGGTGGCAAGACAGCTCCT
TCTTCAGCAGCAACAGCAGCAGCAAGTTAGTGGATTAAAATCTCCCAAGAGGAATGACAAACAACCAGCTC
TTCAGGTTCCCGTGTCAGTGGCTATGATGACACCTCAAGTTATCACTCCCCAGCAAATGCAGCAGATCCTC
CAGCAACAAGTGCTGAGCCCTCAGCAGCTCCAGGTTCTCCTCCAGCAGCAGCAGGCCCTCATGCTTCAA
CAGCAGCAGCTTCAAGAGTTTTATAAAAAACAACAGGAACAGTTGCAGCTTCAACTTTTACAACAACAACAT
GCTGGAAAACAGCCTAAAGAGCAACAGCAGGTGGCTACCCAGCAGTTGGCTTTTCAGCAGCAGCTTTTAC
AGATGCAGCAGTTACAGCAGCAGCACCTCCTGTCTTTGCAGCGCCAAGGCCTTCTGACAATTCAGCCCGG
GCAGCCTGCCCTTCCCCTTCAACCTCTTGCTCAAGGCATGATTCCAACAGAACTGCAGCAGCTCTGGAAA
GAAGTGACAAGTGCTCATACTGCAGAAGAAACCACAGGCAACAATCACAGCAGTTTGGATCTGACCACGA
CATGTGTCTCCTCCTCTGCACCTTCCAAGACCTCCTTAATAATGAACCCACATGCCTCTACCAATGGACAGC
TCTCAGTCCACACTCCCAAAAGGGAAAGTTTGTCCCATGAGGAGCACCCCCATAGCCATCCTCTCTATGGA
CATGGTGTATGCAAGTGGCCAGGCTGTGAAGCAGTGTGCGAAGATTTCCAATCATTTCTAAAACATCTCAAC
AGTGAGCATGCGCTGGACGATAGAAGTACAGCCCAATGTAGAGTACAAATGCAGGTTGTACAGCAGTTAGA
GCTACAGCTTGCAAAAGACAAAGAACGCCTGCAAGCCATGATGACCCACCTGCATGTGAAGTCTACAGAA
CCCAAAGCCGCCCCTCAGCCCTTGAATCTGGTATCAAGTGTCACTCTCTCCAAGTCCGCATCGGAGGCTT
CTCCACAGAGCTTACCTCATACTCCAACGACCCCAACCGCCCCCCTGACTCCCGTCACCCAAGGCCCCTC
TGTCATCACAACCACCAGCATGCACACGGTGGGACCCATCCGCAGGCGGTACTCAGACAAATACAACGTG
CCCATTTCGTCAGCAGATATTGCGCAGAACCAAGAATTTTATAAGAACGCAGAAGTTAGACCACCATTTACAT
ATGCATCTTTAATTAGGCAGGCCATTCTCGAATCTCCAGAAAAGCAGCTAACACTAAATGAGATCTATAACTG
GTTCACACGAATGTTTGCTTACTTCCGACGCAACGCGGCCACGTGGAAGAATGCAGTGTGTCATAATCTTA
GTCTTCACAAGTGTTTTGTGCGAGTAGAAAACGTTAAAGGGGCAGTATGGACAGTGGATGAAGTAGAATTC
CAAAAACGAAGGCCACAAAAGATCAGTGGTAACCCTTCCCTTATTAAAAACATGCAGAGCAGCCACGCCTA
CTGCACACCTCTCAATGCAGCTTTACAGGCTTCAATGGCTGAGAATAGTATACCTCTATACACTACCGCTTC
CATGGGAAATCCCACTCTGGGCAACTTAGCCAGCGCAATACGGGAAGAGCTGAACGGGGCAATGGAGCAT
ACCAACAGCAACGAGAGTGACAGCAGTCCAGGCAGATCTCCTATGCAAGCCGTGCATCCTGTACACGTCA
AAGAAGAGCCCCTCGATCCAGAGGAAGCTGAAGGGCCCCTGTCCTTAGTGACAACAGCCAACCACAGTC
CAGATTTTGACCATGACAGAGATTACGAAGATGAACCAGTAAACGAGGACATGGAGTGA
