## Supplemental Data S4 for "The forkhead transcription factor FKH-7/FOXP acts in chemosensory neurons to regulate developmental decision-making"

**Data S4. Single-cell RNA-seq ADL gene list table.**

This comparison is between all ADL neurons in *fkh-7(R612C)* missense variants and all ADL neurons in *fkh-7(wild-type)* control animals.

| id | gene_name | fkh-7.avg | wt.avg | avg_log2FC | fkh7.pc | wt.pct | p_val | p_val_adj |
|---|---|---|---|---|---|---|---|---|
| WBGene00010516 | K02E11.7 | 0.132829739 | 1.244791545 | -1.604221782 | 0.023 | 0.079 | 0.019119622 | 1 |
| WBGene00002113 | ins-30 | 0.421127812 | 1.218163973 | -1.149880116 | 0.04 | 0.079 | 0.136721436 | 1 |
| WBGene00004451 | rpl-37 | 1.585195649 | 2.379318864 | -1.145677625 | 0.434 | 0.832 | 9.98E-16 | 6.19E-12 |
| WBGene00017613 | F20A1.1 | 0.388843542 | 1.112538812 | -1.044071578 | 0.058 | 0.074 | 0.582398012 | 1 |
| WBGene00016114 | flp-27 | 0.123832607 | 0.775017561 | -0.939461303 | 0.023 | 0.053 | 0.154330178 | 1 |
| WBGene00000962 | dhc-1 | 0.467371866 | 1.008831586 | -0.781161254 | 0.092 | 0.226 | 0.001078294 | 1 |
| WBGene00009122 | tct-1 | 0.664055571 | 1.181641999 | -0.746719373 | 0.139 | 0.416 | 1.59E-07 | 0.000984573 |
| WBGene00015117 | B0294.1 | 0.832306039 | 1.348394473 | -0.744558225 | 0.04 | 0.1 | 0.028056114 | 1 |
| WBGene00006980 | zig-3 | 1.217187802 | 1.728713964 | -0.737976258 | 0.173 | 0.458 | 1.15E-07 | 0.000711276 |
| WBGene00022297 | Y76B12C.3 | 0.164330496 | 0.663871272 | -0.720685 | 0.023 | 0.1 | 0.003148578 | 1 |
| WBGene00002086 | ins-3 | 0.240845069 | 0.72453893 | -0.697822735 | 0.023 | 0.068 | 0.041546132 | 1 |
| WBGene00015514 | nlp-77 | 0.405308302 | 0.885533274 | -0.692818186 | 0.029 | 0.147 | 0.00012067 | 0.748030917 |
| WBGene00012088 | T27E7.3 | 0.145245481 | 0.614964498 | -0.677661297 | 0.023 | 0.068 | 0.040732284 | 1 |
| WBGene00000984 | dhs-21 | 0.390891224 | 0.858460078 | -0.674559267 | 0.075 | 0.153 | 0.022930851 | 1 |
| WBGene00015339 | C02E7.6 | 0.492605266 | 0.95300394 | -0.664214884 | 0.064 | 0.095 | 0.284437909 | 1 |
| WBGene00006479 | tmbi-4 | 0.945328681 | 1.396786426 | -0.65131585 | 0.191 | 0.521 | 1.06E-07 | 0.000659048 |
| WBGene00008601 | mob-2 | 0.223294658 | 0.674641164 | -0.651155366 | 0.029 | 0.179 | 8.15E-06 | 0.050500286 |
| WBGene00019994 | cdh-1 | 0.063574601 | 0.512706761 | -0.64796074 | 0.012 | 0.1 | 0.000358076 | 1 |
| WBGene00000567 | cnx-1 | 0.631411746 | 1.0796813 | -0.646716263 | 0.133 | 0.321 | 0.000154206 | 0.955923484 |
| WBGene00000221 | atf-4 | 1.205798373 | 1.653887378 | -0.646455785 | 0.306 | 0.589 | 1.15E-05 | 0.071555696 |
| WBGene00013317 | Y57G11C.22 | 0.62041777 | 1.06847509 | -0.646410073 | 0.116 | 0.311 | 6.07E-05 | 0.376354561 |
| WBGene00014021 | svop-1 | 0.344199749 | 0.787347482 | -0.639327037 | 0.052 | 0.174 | 0.000513464 | 1 |
| WBGene00000840 | cul-5 | 0.342762369 | 0.781376546 | -0.632786497 | 0.058 | 0.211 | 5.63E-05 | 0.349010075 |
| WBGene00004442 | rpl-28 | 1.497367121 | 1.924287179 | -0.61591545 | 0.358 | 0.695 | 1.34E-07 | 0.000832054 |
| WBGene00009069 | F23A7.4 | 1.597816805 | 2.022001219 | -0.61196875 | 0.133 | 0.168 | 0.407002285 | 1 |
| WBGene00003759 | nlp-21 | 0.067355935 | 0.489891267 | -0.609589628 | 0.012 | 0.063 | 0.010768752 | 1 |
| WBGene00011428 | nlp-80 | 0.753699483 | 1.172984377 | -0.604900237 | 0.127 | 0.116 | 0.644091777 | 1 |
| WBGene00007554 | pptr-2 | 1.178399844 | 1.592504014 | -0.597426032 | 0.254 | 0.505 | 7.59E-05 | 0.47046985 |
| WBGene00005432 | srh-222 | 0.483848349 | 0.891808366 | -0.588561893 | 0.081 | 0.242 | 0.000111385 | 0.690476995 |
| WBGene00001480 | fmo-5 | 0.354924881 | 0.761953328 | -0.587217922 | 0.052 | 0.137 | 0.00807774 | 1 |
| WBGene00000959 | dgk-2 | 0.041149942 | 0.44412646 | -0.581372224 | 0.006 | 0.121 | 1.18E-05 | 0.07301606 |
| WBGene00013071 | Y51A2B.8 | 0.402187464 | 0.801309333 | -0.575811141 | 0.069 | 0.158 | 0.013951922 | 1 |
| WBGene00001030 | dnj-12 | 0.4153523 | 0.811430584 | -0.571420175 | 0.075 | 0.284 | 2.22E-06 | 0.013733869 |
| WBGene00012136 | T28F4.1 | 0.444975151 | 0.840149916 | -0.570116674 | 0.069 | 0.189 | 0.001462133 | 1 |
| WBGene00002017 | hsp-16.11 | 3.382695895 | 3.776609046 | -0.56829655 | 0.798 | 0.9 | 3.77E-07 | 0.002335563 |
| WBGene00200808 | B0034.16 | 0.134949024 | 0.52820715 | -0.567351549 | 0.023 | 0.137 | 0.000116788 | 0.723969716 |
| WBGene00014215 | obr-3 | 0.798581464 | 1.191643961 | -0.567069316 | 0.156 | 0.379 | 3.54E-05 | 0.219228449 |
| WBGene00017858 | F27C1.11 | 0.559755082 | 0.952592495 | -0.566744589 | 0.116 | 0.289 | 0.000279612 | 1 |
| WBGene00020735 | T23F2.2 | 0.191617438 | 0.582011917 | -0.563220179 | 0.029 | 0.089 | 0.017575581 | 1 |
| WBGene00016463 | C35E7.11 | 0.999102844 | 1.388692293 | -0.562058767 | 0.225 | 0.484 | 2.18E-05 | 0.135293446 |
| WBGene00020727 | npr-21 | 1.355871589 | 1.740238242 | -0.554523864 | 0.277 | 0.537 | 0.000115767 | 0.717640406 |
| WBGene00007880 | C33A12.1 | 0.258931792 | 0.642401014 | -0.553229145 | 0.04 | 0.132 | 0.003221544 | 1 |
| WBGene00018604 | uggt-1 | 0.274367963 | 0.652033782 | -0.544856604 | 0.035 | 0.153 | 0.000227604 | 1 |
| WBGene00004477 | rps-8 | 2.448873024 | 2.826473473 | -0.544762296 | 0.688 | 0.895 | 8.72E-07 | 0.005403694 |
| WBGene00001368 | exc-7 | 0.919634095 | 1.29627654 | -0.543380188 | 0.179 | 0.337 | 0.004316952 | 1 |
| WBGene00006537 | tbb-2 | 0.886124197 | 1.261995006 | -0.542266952 | 0.168 | 0.405 | 1.33E-05 | 0.082379365 |
| WBGene00006410 | nck-1 | 0.443013514 | 0.812109621 | -0.532493122 | 0.092 | 0.237 | 0.000633652 | 1 |
| WBGene00000509 | cka-1 | 0 | 0.367186424 | -0.529738034 | 0 | 0.058 | 0.001340226 | 1 |
| WBGene00020225 | sre-40 | 0.541149211 | 0.906089103 | -0.526496973 | 0.087 | 0.237 | 0.00030295 | 1 |

| Legend | |
|---|---|
| id | WormBase GeneID |
| gene_name | Gene symbol (WS273) |
| fkh-7.avg | Log of averaged normalized expression across all fkh-7 ADL neurons |
| wt.avg | Log of averaged normalized expression across all wt ADL neurons |
| avg_log2FC | log fold-change (fkh-7 - wt) |
| fkh7.pct | Percent of fkh-7 ADL neurons where gene is detected (≥ 1 UMI) |
| wt.pct | Percent of wt ADL neurons where gene is detected (≥ 1 UMI) |
| p_val | p-value |
| p_val_adj | Adjusted p-value, based on bonferroni correction |

| | | | | | | | | |
|---|---|---|---|---|---|---|---|---|
| WBGene00016977 | akap-1 | 0.905309668 | 1.268873972 | -0.524512418 | 0.185 | 0.4 | 0.000185003 | 1 |
| WBGene00004487 | rps-18 | 2.353935311 | 2.717035394 | -0.523842688 | 0.671 | 0.921 | 7.88E-08 | 0.000488447 |
| WBGene00011578 | npr-20 | 0.382326757 | 0.743773188 | -0.521456974 | 0.064 | 0.205 | 0.000207103 | 1 |
| WBGene00005383 | srh-167 | 0.62714669 | 0.986600397 | -0.51858208 | 0.098 | 0.3 | 1.46E-05 | 0.090411498 |
| WBGene00004489 | rps-20 | 2.282927431 | 2.64224942 | -0.518392053 | 0.63 | 0.879 | 1.98E-06 | 0.012252833 |
| WBGene00006201 | str-155 | 0.443845453 | 0.801761872 | -0.516364243 | 0.069 | 0.189 | 0.001346348 | 1 |
| WBGene00021497 | Y40C5A.4 | 0.234999675 | 0.591572876 | -0.514426389 | 0.046 | 0.142 | 0.00266686 | 1 |
| WBGene00018849 | spt-16 | 0.357753431 | 0.714304339 | -0.514394228 | 0.075 | 0.232 | 0.000146762 | 0.90977551 |
| WBGene00008641 | pch-2 | 0.063224645 | 0.417131772 | -0.510580057 | 0.012 | 0.053 | 0.029428911 | 1 |
| WBGene00016611 | bicd-1 | 0.035468106 | 0.388975882 | -0.510003915 | 0.006 | 0.079 | 0.000723655 | 1 |
| WBGene00012386 | agef-1 | 0.136559133 | 0.489386404 | -0.509022155 | 0.023 | 0.121 | 0.000500606 | 1 |
| WBGene00009945 | bath-38 | 0.284138937 | 0.636822121 | -0.508814281 | 0.052 | 0.147 | 0.004337646 | 1 |
| WBGene00219300 | W10G6.6 | 0.070797599 | 0.423386505 | -0.508678267 | 0.012 | 0.126 | 2.96E-05 | 0.183247207 |
| WBGene00004267 | rab-3 | 0.3648706 | 0.715273195 | -0.505524086 | 0.075 | 0.216 | 0.000421759 | 1 |
| WBGene00003019 | lin-33 | 0.939578983 | 1.289318428 | -0.504567363 | 0.237 | 0.432 | 0.000922691 | 1 |
| WBGene00009995 | F53F4.13 | 0.913698247 | 1.26180474 | -0.502211511 | 0.04 | 0.089 | 0.075098635 | 1 |
| WBGene00008149 | pyp-1 | 0.550723365 | 0.898680543 | -0.501996094 | 0.116 | 0.289 | 0.000502903 | 1 |
| WBGene00005180 | srg-23 | 0.137832041 | 0.485274793 | -0.501253936 | 0.023 | 0.111 | 0.001216738 | 1 |
| WBGene00004478 | rps-9 | 2.587301595 | 2.933392384 | -0.499303466 | 0.717 | 0.932 | 2.36E-07 | 0.001462968 |
| WBGene00006791 | unc-57 | 0.167099539 | 0.510974092 | -0.496106113 | 0.035 | 0.111 | 0.006466338 | 1 |
| WBGene00013378 | emc-3 | 0.355692317 | 0.69904259 | -0.495349737 | 0.069 | 0.205 | 0.000419873 | 1 |
| WBGene00021936 | zipt-15 | 0.663856695 | 1.004187736 | -0.490993904 | 0.104 | 0.226 | 0.004509302 | 1 |
| WBGene00001178 | egl-9 | 1.568025726 | 1.907951194 | -0.490408787 | 0.382 | 0.653 | 0.000108441 | 0.672225097 |
| WBGene00006772 | unc-36 | 0.404991063 | 0.743541689 | -0.488425309 | 0.064 | 0.221 | 6.14E-05 | 0.380593945 |
| WBGene00021369 | siah-1 | 0.432525688 | 0.770538499 | -0.487649407 | 0.087 | 0.226 | 0.000757407 | 1 |
| WBGene00001448 | flp-5 | 2.180458439 | 2.518294863 | -0.487394934 | 0.462 | 0.753 | 8.09E-07 | 0.005016159 |
| WBGene00000209 | asg-1 | 0.151612262 | 0.489174343 | -0.48699914 | 0.017 | 0.116 | 0.00028765 | 1 |
| WBGene00045048 | D1007.19 | 0.370542497 | 0.707976431 | -0.486814264 | 0.058 | 0.179 | 0.000751868 | 1 |
| WBGene00017156 | F01E11.3 | 0.437817536 | 0.774455005 | -0.485665207 | 0.081 | 0.211 | 0.001408978 | 1 |
| WBGene00004416 | rpl-5 | 2.230601948 | 2.566076548 | -0.483987541 | 0.607 | 0.868 | 1.51E-05 | 0.093334814 |
| WBGene00013208 | Y54G9A.5 | 0.787183444 | 1.122510961 | -0.483775346 | 0.127 | 0.337 | 2.62E-05 | 0.162225396 |
| WBGene00015767 | hex-2 | 0.508207054 | 0.843506984 | -0.483735546 | 0.098 | 0.2 | 0.015722272 | 1 |
| WBGene00005270 | srh-47 | 0.03273604 | 0.367617887 | -0.483132379 | 0.006 | 0.068 | 0.002059944 | 1 |
| WBGene00195164 | K12B6.11 | 0.082660857 | 0.41709319 | -0.482483868 | 0.012 | 0.084 | 0.001628262 | 1 |
| WBGene00010471 | cdr-6 | 0.911642938 | 1.244088142 | -0.479617047 | 0.179 | 0.384 | 0.000239919 | 1 |
| WBGene00002045 | icd-1 | 1.429544315 | 1.758044591 | -0.473925719 | 0.335 | 0.642 | 4.23E-05 | 0.26236811 |
| WBGene00018078 | F35H12.6 | 0.350358735 | 0.678154996 | -0.47291004 | 0.058 | 0.216 | 4.75E-05 | 0.294414843 |
| WBGene00006655 | tub-1 | 0.705915016 | 1.032739188 | -0.471507613 | 0.133 | 0.305 | 0.000795803 | 1 |
| WBGene00020931 | cytb-5.2 | 0.578096449 | 0.903476202 | -0.469423756 | 0.104 | 0.226 | 0.006083073 | 1 |
| WBGene00003044 | lir-1 | 0.385298792 | 0.708876618 | -0.466824126 | 0.081 | 0.153 | 0.051756865 | 1 |
| WBGene00010306 | golg-4 | 0.079978416 | 0.401908119 | -0.464446385 | 0.012 | 0.079 | 0.002691151 | 1 |
| WBGene00004414 | rpl-3 | 2.17366885 | 2.49533555 | -0.464066952 | 0.584 | 0.832 | 6.14E-05 | 0.380839482 |
| WBGene00014091 | ZK822.4 | 0.404821286 | 0.722796505 | -0.458741271 | 0.04 | 0.137 | 0.002114925 | 1 |
| WBGene00003632 | nhr-42 | 1.466606609 | 1.784410008 | -0.458493388 | 0.301 | 0.516 | 0.000925304 | 1 |
| WBGene00006814 | unc-82 | 0.155558003 | 0.471266647 | -0.455471295 | 0.029 | 0.111 | 0.003279559 | 1 |
| WBGene00044638 | F23A7.8 | 1.263195506 | 1.578192874 | -0.454445141 | 0.116 | 0.137 | 0.60983366 | 1 |
| WBGene00010898 | nphp-1 | 0.660791094 | 0.973119834 | -0.450595124 | 0.116 | 0.268 | 0.001092033 | 1 |
| WBGene00009920 | abts-1 | 1.281360679 | 1.59287239 | -0.449416401 | 0.289 | 0.553 | 0.00022272 | 1 |
| WBGene00002196 | kin-10 | 0.805322381 | 1.11668949 | -0.449207784 | 0.162 | 0.363 | 0.000297668 | 1 |
| WBGene00003123 | mag-1 | 0.358911536 | 0.670157063 | -0.449032377 | 0.058 | 0.216 | 4.96E-05 | 0.307234755 |
| WBGene00001431 | fkb-6 | 1.0162748 | 1.326823617 | -0.448027238 | 0.202 | 0.426 | 0.000154787 | 0.959524998 |
| WBGene00010648 | K08C7.6 | 0.138296626 | 0.448257828 | -0.447179489 | 0.017 | 0.068 | 0.020098499 | 1 |
| WBGene00014249 | ZK1307.8 | 0.029435099 | 0.339271908 | -0.447000029 | 0.006 | 0.079 | 0.000731105 | 1 |

| | | | | | | | | |
|---|---|---|---|---|---|---|---|---|
| WBGene00008402 | wdfy-2 | 0.071470089 | 0.379301098 | -0.444106269 | 0.012 | 0.111 | 0.000138167 | 0.85649983 |
| WBGene00022668 | ZK154.6 | 1.410245006 | 1.717926741 | -0.443890912 | 0.283 | 0.489 | 0.000730602 | 1 |
| WBGene00004055 | pmk-1 | 0.062746277 | 0.369822855 | -0.443017857 | 0.012 | 0.084 | 0.001541424 | 1 |
| WBGene00003616 | nhr-17 | 2.313301643 | 2.618953538 | -0.440962472 | 0.578 | 0.858 | 3.54E-06 | 0.02191552 |
| WBGene00016138 | flh-2 | 0.358317398 | 0.663589349 | -0.44041433 | 0.075 | 0.174 | 0.007159925 | 1 |
| WBGene00004101 | hgrs-1 | 0.766133431 | 1.070554652 | -0.439186986 | 0.162 | 0.316 | 0.00230505 | 1 |
| WBGene00012847 | srxa-15 | 1.174722774 | 1.478797051 | -0.438686451 | 0.26 | 0.489 | 0.000700683 | 1 |
| WBGene00009953 | F53A2.9 | 0.027333216 | 0.330096439 | -0.436794999 | 0.006 | 0.068 | 0.002039269 | 1 |
| WBGene00304791 | Y62H9A.16 | 0.13712256 | 0.439528028 | -0.436278869 | 0.023 | 0.105 | 0.002009052 | 1 |
| WBGene00003904 | pabp-2 | 0.28365991 | 0.584419574 | -0.433904475 | 0.052 | 0.163 | 0.001235601 | 1 |
| WBGene00044061 | tbc-12 | 0.548079224 | 0.848203043 | -0.432987145 | 0.092 | 0.232 | 0.000956151 | 1 |
| WBGene00007180 | B0457.6 | 1.610327587 | 1.909527075 | -0.431653618 | 0.358 | 0.668 | 1.29E-05 | 0.080177319 |
| WBGene00013034 | tat-1 | 0 | 0.29911588 | -0.431532997 | 0 | 0.079 | 0.000165421 | 1 |
| WBGene00009992 | F53F4.10 | 0.239311276 | 0.537535929 | -0.430247227 | 0.046 | 0.163 | 0.00057511 | 1 |
| WBGene00004436 | rpl-24.1 | 2.872525841 | 3.170401781 | -0.429744141 | 0.803 | 0.953 | 1.96E-06 | 0.012153014 |
| WBGene00012419 | srz-73 | 0 | 0.297524554 | -0.429237198 | 0 | 0.053 | 0.002261451 | 1 |
| WBGene00015484 | atgl-1 | 0 | 0.297331327 | -0.428958431 | 0 | 0.058 | 0.001340226 | 1 |
| WBGene00004367 | ric-8 | 0.228437931 | 0.525062914 | -0.427939392 | 0.035 | 0.163 | 0.000101381 | 0.628462899 |
| WBGene00007029 | mys-1 | 0.085838176 | 0.382188088 | -0.427542548 | 0.017 | 0.089 | 0.003006188 | 1 |
| WBGene00015025 | mrpl-9 | 0.250158806 | 0.54623414 | -0.427146416 | 0.046 | 0.132 | 0.00662496 | 1 |
| WBGene00017240 | calu-2 | 0.440487994 | 0.736302483 | -0.426770096 | 0.069 | 0.147 | 0.027629312 | 1 |
| WBGene00006713 | ubc-18 | 0.925560388 | 1.221192033 | -0.426506308 | 0.191 | 0.432 | 6.07E-05 | 0.37653197 |
| WBGene00016643 | vps-45 | 0.338146615 | 0.632523337 | -0.424695838 | 0.069 | 0.189 | 0.001624484 | 1 |
| WBGene00016366 | nhr-140 | 0.684461921 | 0.978129623 | -0.423672937 | 0.127 | 0.279 | 0.001733732 | 1 |
| WBGene00022063 | nlp-81 | 0.0817173 | 0.374922063 | -0.423005057 | 0.017 | 0.063 | 0.029505231 | 1 |
| WBGene00004435 | rpl-23 | 1.982320833 | 2.275406176 | -0.422832771 | 0.526 | 0.779 | 0.000108382 | 0.671859908 |
| WBGene00044238 | C30H6.12 | 0.280469282 | 0.572377222 | -0.421134138 | 0.052 | 0.158 | 0.001951768 | 1 |
| WBGene00017735 | did-2 | 0.391486423 | 0.683267723 | -0.420951434 | 0.081 | 0.158 | 0.035138395 | 1 |
| WBGene00021831 | Y54E10A.11 | 0.128788468 | 0.418942389 | -0.418603623 | 0.023 | 0.1 | 0.003322109 | 1 |
| WBGene00016162 | crh-2 | 0.465719864 | 0.755717329 | -0.418377905 | 0.087 | 0.195 | 0.006276224 | 1 |
| WBGene00015652 | srz-79 | 0 | 0.286652864 | -0.413552666 | 0 | 0.063 | 0.000794656 | 1 |
| WBGene00004836 | sls-2.4 | 0.502087442 | 0.788182256 | -0.41274757 | 0.052 | 0.163 | 0.001292519 | 1 |
| WBGene00011827 | T19A6.1 | 0.137326676 | 0.422901736 | -0.411997723 | 0.023 | 0.116 | 0.000834066 | 1 |
| WBGene00004567 | rrn-2.1 | 1.678499061 | 1.963473974 | -0.411131894 | 0.347 | 0.611 | 0.00024194 | 1 |
| WBGene00019902 | R05G6.10 | 0.218552067 | 0.502604286 | -0.409800727 | 0.04 | 0.142 | 0.001438037 | 1 |
| WBGene00011975 | T24B1.1 | 0.076438956 | 0.360256466 | -0.409462114 | 0.012 | 0.089 | 0.000991519 | 1 |
| WBGene00004419 | rpl-7A | 1.891446177 | 2.17460327 | -0.408509334 | 0.526 | 0.763 | 0.002850878 | 1 |
| WBGene00008840 | F15A4.5 | 2.884816951 | 3.166802313 | -0.406818883 | 0.786 | 0.937 | 2.13E-06 | 0.013228362 |
| WBGene00013496 | Y70D2A.1 | 0.715581971 | 0.9974903 | -0.406707747 | 0.145 | 0.268 | 0.014415937 | 1 |
| WBGene00005175 | srg-18 | 0.525650776 | 0.806886418 | -0.405737267 | 0.11 | 0.237 | 0.003856013 | 1 |
| WBGene00013276 | sre-42 | 0.427632239 | 0.707360243 | -0.403562204 | 0.075 | 0.2 | 0.001489636 | 1 |
| WBGene00003634 | nhr-44 | 0.359537127 | 0.638790639 | -0.402877658 | 0.069 | 0.142 | 0.038983373 | 1 |
| WBGene00004449 | rpl-35 | 2.546892678 | 2.825820331 | -0.402407542 | 0.723 | 0.889 | 1.32E-05 | 0.081595128 |
| WBGene00019284 | K01A2.10 | 0.562294367 | 0.840677016 | -0.401621267 | 0.087 | 0.226 | 0.0006775 | 1 |
| WBGene00008288 | C54C6.6 | 0.322928758 | 0.599746109 | -0.39936302 | 0.058 | 0.163 | 0.002836022 | 1 |
| WBGene00304813 | F07G6.11 | 2.110661058 | 2.386535983 | -0.398003386 | 0.064 | 0.084 | 0.437750354 | 1 |
| WBGene00005562 | sri-50 | 0.028992159 | 0.304609582 | -0.397631889 | 0.006 | 0.063 | 0.003435925 | 1 |
| WBGene00001945 | his-71 | 1.395904011 | 1.671424016 | -0.397491345 | 0.358 | 0.584 | 0.001257053 | 1 |
| WBGene00009716 | F44G4.7 | 1.745648924 | 2.020441694 | -0.396442166 | 0.468 | 0.679 | 0.001073751 | 1 |
| WBGene00004434 | rpl-22 | 2.466074269 | 2.740832938 | -0.396392969 | 0.717 | 0.889 | 3.97E-05 | 0.246025333 |
| WBGene00004470 | rps-1 | 2.742778544 | 3.017221674 | -0.395937742 | 0.775 | 0.932 | 1.71E-05 | 0.106286135 |
| WBGene00005389 | srh-173 | 0.202152594 | 0.476173118 | -0.39532805 | 0.04 | 0.116 | 0.010550289 | 1 |
| WBGene00021350 | Y37E3.8 | 2.821420842 | 3.095093999 | -0.394826906 | 0.786 | 0.942 | 3.92E-05 | 0.242831129 |

| | | | | | | | | |
|---|---|---|---|---|---|---|---|---|
| WBGene00004484 | rps-15 | 2.105814914 | 2.378186327 | -0.392948886 | 0.572 | 0.853 | 2.82E-05 | 0.174988964 |
| WBGene00000834 | cua-1 | 0.541348168 | 0.813072521 | -0.392015376 | 0.098 | 0.226 | 0.00290873 | 1 |
| WBGene00006519 | cox-6A | 0.747586488 | 1.018138586 | -0.39032417 | 0.145 | 0.332 | 0.000365235 | 1 |
| WBGene00005395 | srh-180 | 0.810541375 | 1.079799156 | -0.388456865 | 0.116 | 0.237 | 0.005546805 | 1 |
| WBGene00006982 | zig-5 | 0.706802506 | 0.97578427 | -0.388058657 | 0.133 | 0.258 | 0.01363716 | 1 |
| WBGene00004430 | rpl-18 | 2.516644807 | 2.784730767 | -0.386766284 | 0.705 | 0.926 | 3.62E-05 | 0.224448576 |
| WBGene00021510 | frpr-19 | 1.313681072 | 1.581510248 | -0.386395825 | 0.289 | 0.442 | 0.017131131 | 1 |
| WBGene00014943 | Y68A4A.8 | 0.804327813 | 1.071932498 | -0.386071952 | 0.116 | 0.316 | 5.00E-05 | 0.309906023 |
| WBGene00000192 | arl-8 | 0.34260017 | 0.609747887 | -0.385412687 | 0.064 | 0.189 | 0.00095365 | 1 |
| WBGene00004680 | rars-2 | 0.273377969 | 0.540236015 | -0.38499478 | 0.046 | 0.168 | 0.000431907 | 1 |
| WBGene00004781 | set-1 | 0.136972947 | 0.403496368 | -0.384512018 | 0.029 | 0.116 | 0.002173311 | 1 |
| WBGene00018803 | fbxa-24 | 1.711465289 | 1.977735324 | -0.384146459 | 0.37 | 0.626 | 0.000157408 | 0.975774728 |
| WBGene00004677 | rrn-3.56 | 0.782800267 | 1.04887538 | -0.383865246 | 0.121 | 0.226 | 0.023949598 | 1 |
| WBGene00017901 | igeg-1 | 0.308089845 | 0.573957477 | -0.383565914 | 0.035 | 0.132 | 0.001422173 | 1 |
| WBGene00017642 | F20D12.2 | 0.09830204 | 0.364159594 | -0.383551375 | 0.017 | 0.068 | 0.019357007 | 1 |
| WBGene00011956 | T23F11.4 | 0 | 0.264264297 | -0.381252791 | 0 | 0.089 | 5.79E-05 | 0.358687848 |
| WBGene00004441 | rpl-27 | 1.940675825 | 2.202592075 | -0.377865275 | 0.572 | 0.779 | 0.001662062 | 1 |
| WBGene00010043 | F54C9.7 | 0.479191092 | 0.740022918 | -0.376300782 | 0.075 | 0.168 | 0.014281997 | 1 |
| WBGene00003002 | lin-13 | 0.444003051 | 0.704308924 | -0.375541993 | 0.075 | 0.174 | 0.008726177 | 1 |
| WBGene00001352 | evl-14 | 0.075324056 | 0.335543089 | -0.375416709 | 0.012 | 0.095 | 0.000607904 | 1 |
| WBGene00019365 | rei-2 | 0.346993744 | 0.606978713 | -0.375079026 | 0.058 | 0.184 | 0.000663069 | 1 |
| WBGene00004471 | rps-2 | 1.950362332 | 2.209562986 | -0.373947497 | 0.514 | 0.805 | 0.000168727 | 1 |
| WBGene00016981 | rpn-13 | 0.452157891 | 0.711066103 | -0.373525594 | 0.092 | 0.163 | 0.073917947 | 1 |
| WBGene00000953 | del-2 | 0.803905826 | 1.062358344 | -0.372868166 | 0.168 | 0.289 | 0.016808299 | 1 |
| WBGene00004493 | rps-24 | 2.202148562 | 2.460251989 | -0.372364535 | 0.607 | 0.847 | 0.000251299 | 1 |
| WBGene00022847 | ZK1055.6 | 0.175351503 | 0.433356289 | -0.372222225 | 0.023 | 0.116 | 0.000847627 | 1 |
| WBGene00001329 | epn-1 | 0.680987491 | 0.938668343 | -0.371754888 | 0.15 | 0.279 | 0.014067225 | 1 |
| WBGene00000092 | ags-3 | 1.326532398 | 1.583666985 | -0.370966794 | 0.26 | 0.542 | 2.97E-05 | 0.183878355 |
| WBGene00015494 | C05E11.3 | 0.282335291 | 0.53869111 | -0.369843269 | 0.029 | 0.089 | 0.019131504 | 1 |
| WBGene00201458 | VF15C11L.2 | 0.108627621 | 0.364969489 | -0.369823142 | 0.017 | 0.105 | 0.000754163 | 1 |
| WBGene00023480 | srz-68 | 1.119981475 | 1.376236947 | -0.369698499 | 0.243 | 0.426 | 0.002849632 | 1 |
| WBGene00000288 | cal-4 | 1.198274576 | 1.454511518 | -0.369671766 | 0.277 | 0.484 | 0.002689361 | 1 |
| WBGene00017311 | F09G2.2 | 0.265395387 | 0.521507088 | -0.369491081 | 0.052 | 0.158 | 0.002225601 | 1 |
| WBGene00012031 | gpdh-3 | 0.41080025 | 0.6668594 | -0.369415265 | 0.064 | 0.147 | 0.015630629 | 1 |
| WBGene00023470 | srz-87 | 0.350574997 | 0.606574743 | -0.369329564 | 0.046 | 0.147 | 0.002070936 | 1 |
| WBGene00016610 | paqr-1 | 0.064531746 | 0.320278055 | -0.368963931 | 0.012 | 0.084 | 0.001499623 | 1 |
| WBGene00002223 | klp-12 | 0.196119057 | 0.451119565 | -0.367887968 | 0.04 | 0.105 | 0.021608887 | 1 |
| WBGene00004428 | rpl-16 | 2.274085734 | 2.52861989 | -0.367215163 | 0.642 | 0.868 | 0.000210963 | 1 |
| WBGene00004485 | rps-16 | 1.53324396 | 1.787510561 | -0.366829164 | 0.387 | 0.632 | 0.001844746 | 1 |
| WBGene00019205 | kcc-2 | 1.199071264 | 1.453150231 | -0.366558466 | 0.272 | 0.458 | 0.003829173 | 1 |
| WBGene00004474 | rps-5 | 2.516179395 | 2.769823777 | -0.365931493 | 0.757 | 0.9 | 7.50E-05 | 0.465149965 |
| WBGene00001497 | fars-1 | 0.174945622 | 0.428552404 | -0.365877247 | 0.023 | 0.095 | 0.005334264 | 1 |
| WBGene00004475 | rps-6 | 1.616265652 | 1.869504008 | -0.36534572 | 0.399 | 0.695 | 0.000106549 | 0.660499753 |
| WBGene00012140 | T28F4.5 | 2.292277114 | 2.5442925 | -0.363581346 | 0.642 | 0.816 | 0.000627911 | 1 |
| WBGene00015478 | mapk-15 | 0.382587368 | 0.6343629 | -0.36323531 | 0.081 | 0.179 | 0.011278852 | 1 |
| WBGene00017711 | F22E5.13 | 0.443739997 | 0.695453756 | -0.363146192 | 0.069 | 0.189 | 0.001834779 | 1 |
| WBGene00018413 | cyp-33C4 | 0.084861835 | 0.336473173 | -0.36299843 | 0.017 | 0.084 | 0.004890429 | 1 |
| WBGene00011111 | snfc-5 | 0.174760621 | 0.426323942 | -0.362929156 | 0.029 | 0.116 | 0.002048719 | 1 |
| WBGene00010642 | mks-6 | 0.555335341 | 0.805725065 | -0.361236014 | 0.11 | 0.221 | 0.011396597 | 1 |
| WBGene00008547 | F07A11.4 | 0.036065675 | 0.286390212 | -0.361141967 | 0.006 | 0.068 | 0.002080811 | 1 |
| WBGene00010735 | frpr-15 | 0.330902344 | 0.580800064 | -0.360526201 | 0.04 | 0.142 | 0.001218745 | 1 |
| WBGene00003674 | nhr-84 | 0.915746463 | 1.164543104 | -0.35893768 | 0.191 | 0.358 | 0.002642276 | 1 |
| WBGene00006919 | vha-10 | 0.62417882 | 0.872578183 | -0.358364529 | 0.116 | 0.263 | 0.001774598 | 1 |

| | | | | | | | | |
|---|---|---|---|---|---|---|---|---|
| WBGene00004440 | rpl-26 | 2.154835312 | 2.402924525 | -0.357917077 | 0.636 | 0.821 | 0.001334595 | 1 |
| WBGene00016514 | C38C3.6 | 0.267127192 | 0.514737061 | -0.357225531 | 0.046 | 0.137 | 0.004453116 | 1 |
| WBGene00004421 | rpl-10 | 2.591960973 | 2.838739466 | -0.356026108 | 0.751 | 0.916 | 0.001226273 | 1 |
| WBGene00018076 | pifk-1 | 0 | 0.246703781 | -0.355918322 | 0 | 0.058 | 0.001340226 | 1 |
| WBGene00018672 | sorf-2 | 0.116521754 | 0.363045584 | -0.355658706 | 0.023 | 0.095 | 0.005138332 | 1 |
| WBGene00013415 | ilcr-1 | 0.522521277 | 0.76894414 | -0.355513043 | 0.104 | 0.237 | 0.00332062 | 1 |
| WBGene00016617 | C43H6.4 | 0.08117427 | 0.327589139 | -0.35550151 | 0.012 | 0.079 | 0.002667073 | 1 |
| WBGene00202501 | Y57E12B.10 | 0.546650935 | 0.792832119 | -0.355164373 | 0.087 | 0.221 | 0.001088206 | 1 |
| WBGene00017035 | D1065.3 | 0.131397225 | 0.377429384 | -0.354949375 | 0.017 | 0.084 | 0.004930051 | 1 |
| WBGene00007168 | B0393.3 | 0.417412533 | 0.66294408 | -0.354227145 | 0.075 | 0.174 | 0.008082794 | 1 |
| WBGene00002215 | klc-2 | 0.348335346 | 0.593262356 | -0.353354982 | 0.058 | 0.163 | 0.003155467 | 1 |
| WBGene00004450 | rpl-36 | 2.576907409 | 2.821759206 | -0.353246472 | 0.734 | 0.921 | 2.07E-05 | 0.128512353 |
| WBGene00004431 | rpl-19 | 2.149422224 | 2.394076533 | -0.352961559 | 0.584 | 0.832 | 6.40E-05 | 0.396659129 |
| WBGene00007042 | pbrm-1 | 0.41979811 | 0.664247225 | -0.352665526 | 0.087 | 0.189 | 0.008380728 | 1 |
| WBGene00001995 | hpl-1 | 0.242379503 | 0.486371635 | -0.352006238 | 0.046 | 0.132 | 0.00670575 | 1 |
| WBGene00000371 | cox-5B | 1.282426378 | 1.526383187 | -0.351955279 | 0.249 | 0.526 | 1.06E-05 | 0.065802046 |
| WBGene00004483 | rps-14 | 3.143596019 | 3.387510535 | -0.351894262 | 0.89 | 0.958 | 1.45E-06 | 0.008984882 |
| WBGene00016250 | hsp-110 | 1.919391839 | 2.163151108 | -0.351670288 | 0.52 | 0.721 | 0.014252951 | 1 |
| WBGene00011528 | fndc-1 | 0.084480265 | 0.32820071 | -0.351614278 | 0.017 | 0.089 | 0.003031046 | 1 |
| WBGene00011527 | cchl-1 | 0.031590918 | 0.275100905 | -0.351310651 | 0.006 | 0.074 | 0.001259409 | 1 |
| WBGene00022401 | Y97E10AR.6 | 0.428157766 | 0.671132943 | -0.350539082 | 0.069 | 0.189 | 0.001662745 | 1 |
| WBGene00000837 | cul-2 | 0.066472739 | 0.309384333 | -0.350447352 | 0.006 | 0.095 | 0.000163847 | 1 |
| WBGene00017993 | cec-5 | 0.701489438 | 0.944325872 | -0.350338919 | 0.145 | 0.311 | 0.001599591 | 1 |
| WBGene00007707 | fath-1 | 0.123481223 | 0.365827156 | -0.349631276 | 0.023 | 0.095 | 0.005138332 | 1 |
| WBGene00017811 | F26A1.13 | 0.739958024 | 0.982233714 | -0.349529937 | 0.133 | 0.289 | 0.001609418 | 1 |
| WBGene00004447 | rpl-33 | 2.205014701 | 2.447114752 | -0.349276543 | 0.624 | 0.847 | 0.000354454 | 1 |
| WBGene00014015 | panl-3 | 0.17060513 | 0.412522945 | -0.349013631 | 0.029 | 0.105 | 0.005251168 | 1 |
| WBGene00004032 | pkc-1 | 0.970001709 | 1.211815403 | -0.348863417 | 0.191 | 0.395 | 0.000588269 | 1 |
| WBGene00011481 | imp-2 | 0.532241632 | 0.773700062 | -0.34835088 | 0.087 | 0.216 | 0.002071204 | 1 |
| WBGene00008720 | F12F6.1 | 0 | 0.241116717 | -0.347857892 | 0 | 0.063 | 0.000794656 | 1 |
| WBGene00019488 | K07D4.9 | 2.86367428 | 3.104430499 | -0.347337804 | 0.798 | 0.953 | 0.000267799 | 1 |
| WBGene00004432 | rpl-20 | 2.359567267 | 2.600134848 | -0.347065656 | 0.642 | 0.863 | 0.00028369 | 1 |
| WBGene00016706 | C46C11.3 | 1.516595874 | 1.756865911 | -0.346636391 | 0.353 | 0.637 | 0.000142176 | 0.881350017 |
| WBGene00016343 | cnnm-3 | 0.02960092 | 0.269629511 | -0.346288057 | 0.006 | 0.053 | 0.009548988 | 1 |
| WBGene00010272 | amph-1 | 0.09212324 | 0.332070244 | -0.346170353 | 0.006 | 0.074 | 0.001285202 | 1 |
| WBGene00044324 | ufm-1 | 0.423515091 | 0.663390294 | -0.346066765 | 0.087 | 0.205 | 0.004602026 | 1 |
| WBGene00016871 | inso-1 | 0.104837762 | 0.344309868 | -0.34548522 | 0.023 | 0.089 | 0.007795301 | 1 |
| WBGene00003565 | ncs-3 | 0.356145239 | 0.595383969 | -0.34514853 | 0.058 | 0.137 | 0.017499843 | 1 |
| WBGene00000814 | csn-2 | 0.175036409 | 0.413993384 | -0.344742042 | 0.029 | 0.089 | 0.017691201 | 1 |
| WBGene00011892 | erh-1 | 0.168212559 | 0.407026629 | -0.344535874 | 0.029 | 0.111 | 0.003140062 | 1 |
| WBGene00021826 | txl-1 | 0.303208427 | 0.540949653 | -0.342988087 | 0.052 | 0.116 | 0.044067391 | 1 |
| WBGene00022306 | hxk-3 | 0.159515825 | 0.397126594 | -0.342799877 | 0.029 | 0.105 | 0.005325669 | 1 |
| WBGene00020562 | T19C3.4 | 0.245355704 | 0.482332235 | -0.341884867 | 0.04 | 0.147 | 0.00103642 | 1 |
| WBGene00012825 | Y43F8C.3 | 0.093680084 | 0.330134617 | -0.341131782 | 0.012 | 0.089 | 0.001009978 | 1 |
| WBGene00005584 | sri-72 | 0.799113561 | 1.03550607 | -0.341042301 | 0.121 | 0.3 | 0.000230898 | 1 |
| WBGene00013376 | Y62E10A.6 | 0.028449883 | 0.264602764 | -0.34069659 | 0.006 | 0.053 | 0.009548988 | 1 |
| WBGene00021264 | fbxa-13 | 0.194957126 | 0.430902408 | -0.340397088 | 0.035 | 0.074 | 0.116051488 | 1 |
| WBGene00004417 | rpl-6 | 1.80363832 | 2.039550529 | -0.340349374 | 0.486 | 0.758 | 0.00014836 | 0.91968551 |
| WBGene00000490 | che-11 | 1.456363338 | 1.692141827 | -0.340156457 | 0.312 | 0.558 | 0.000420392 | 1 |
| WBGene00018266 | nhr-183 | 0.574804155 | 0.810541919 | -0.340097703 | 0.11 | 0.242 | 0.0051874 | 1 |
| WBGene00001459 | flp-16 | 2.688770305 | 2.924450009 | -0.340013941 | 0.746 | 0.958 | 0.001105922 | 1 |
| WBGene00007955 | selt-1.1 | 0.190528236 | 0.42595633 | -0.339650943 | 0.035 | 0.105 | 0.012615414 | 1 |
| WBGene00009546 | fbxb-3 | 0.464730995 | 0.699804647 | -0.339139592 | 0.069 | 0.142 | 0.036577723 | 1 |

| | | | | | | | | |
|---|---|---|---|---|---|---|---|---|
| WBGene00006839 | unc-115 | 0.176276641 | 0.411085175 | -0.338757108 | 0.035 | 0.111 | 0.007381632 | 1 |
| WBGene00018481 | F45F2.9 | 0.138035849 | 0.372800572 | -0.338693901 | 0.017 | 0.105 | 0.000679914 | 1 |
| WBGene00011156 | rbm-3.2 | 2.372885597 | 2.60760786 | -0.338632645 | 0.676 | 0.842 | 0.003141696 | 1 |
| WBGene00016365 | nhr-139 | 0.449104687 | 0.683603443 | -0.338310193 | 0.075 | 0.179 | 0.007124219 | 1 |
| WBGene00044418 | C08G5.7 | 0.892964102 | 1.127410779 | -0.338235059 | 0.191 | 0.379 | 0.001575525 | 1 |
| WBGene00015373 | C03B1.2 | 0.49111189 | 0.725255297 | -0.337797532 | 0.069 | 0.189 | 0.001479401 | 1 |
| WBGene00012468 | Y17G7B.17 | 1.019998123 | 1.254101976 | -0.337740468 | 0.231 | 0.447 | 0.001171574 | 1 |
| WBGene00000938 | dcp-66 | 0.800898479 | 1.034749677 | -0.337375964 | 0.156 | 0.337 | 0.000907261 | 1 |
| WBGene00019385 | irld-41 | 0.164125864 | 0.397578741 | -0.336801308 | 0.029 | 0.1 | 0.008117972 | 1 |
| WBGene00004364 | ric-4 | 2.213939083 | 2.447225111 | -0.336560595 | 0.59 | 0.858 | 0.000171877 | 1 |
| WBGene00015096 | B0261.8 | 0.146691333 | 0.379960076 | -0.33653566 | 0.029 | 0.1 | 0.007550476 | 1 |
| WBGene00003952 | pbs-6 | 0.631162391 | 0.863430821 | -0.335092512 | 0.139 | 0.289 | 0.005375236 | 1 |
| WBGene00018846 | eef-1B.1 | 2.203436979 | 2.435655915 | -0.335021107 | 0.63 | 0.795 | 0.002374961 | 1 |
| WBGene00004492 | rps-23 | 2.648210846 | 2.880213412 | -0.334708952 | 0.763 | 0.932 | 0.000177763 | 1 |
| WBGene00015555 | ufl-1 | 0.079199819 | 0.311165357 | -0.334655532 | 0.017 | 0.074 | 0.012075417 | 1 |
| WBGene00016570 | gnrr-4 | 0.733151714 | 0.964456456 | -0.333702205 | 0.11 | 0.289 | 0.000170872 | 1 |
| WBGene00004919 | snr-6 | 0.654396439 | 0.885619094 | -0.333583778 | 0.139 | 0.274 | 0.00684222 | 1 |
| WBGene00001972 | hmg-1.2 | 0.74969824 | 0.980460209 | -0.332919148 | 0.162 | 0.316 | 0.004712888 | 1 |
| WBGene00011507 | T05H10.1 | 0.188069808 | 0.417924202 | -0.331609794 | 0.029 | 0.121 | 0.001522569 | 1 |
| WBGene00005295 | srh-74 | 0.527610643 | 0.757160764 | -0.331170823 | 0.11 | 0.205 | 0.025207569 | 1 |
| WBGene00009672 | F43G9.12 | 0.066884966 | 0.296388763 | -0.33110399 | 0.012 | 0.079 | 0.002667073 | 1 |
| WBGene00008480 | eif-2A | 0.289662212 | 0.518218169 | -0.329736546 | 0.052 | 0.132 | 0.013598532 | 1 |
| WBGene00018608 | F48E3.9 | 0.994300963 | 1.222388899 | -0.329061334 | 0.22 | 0.353 | 0.023033044 | 1 |
| WBGene00015063 | B0228.6 | 0.516819011 | 0.744785253 | -0.328885768 | 0.104 | 0.226 | 0.005536757 | 1 |
| WBGene00001915 | his-41 | 2.853661254 | 3.081386327 | -0.328537832 | 0.798 | 0.895 | 0.001738837 | 1 |
| WBGene00004946 | sop-3 | 0.198748696 | 0.426436508 | -0.328484077 | 0.035 | 0.079 | 0.081850986 | 1 |
| WBGene00012717 | Y39E4B.6 | 0.640672772 | 0.867721686 | -0.327562342 | 0.127 | 0.242 | 0.017736062 | 1 |
| WBGene00004433 | rpl-21 | 2.418974135 | 2.645537789 | -0.326862259 | 0.699 | 0.9 | 0.000838028 | 1 |
| WBGene00185067 | Y54G11A.17 | 0.071338274 | 0.297521558 | -0.326313503 | 0.012 | 0.058 | 0.018996363 | 1 |
| WBGene00013298 | Y57G11B.6 | 1.146951914 | 1.372665834 | -0.325636353 | 0.243 | 0.495 | 0.000410441 | 1 |
| WBGene00006704 | ubc-7 | 0.507940609 | 0.733291521 | -0.325112644 | 0.11 | 0.253 | 0.002678287 | 1 |
| WBGene00004491 | rps-22 | 2.436576765 | 2.661905209 | -0.325080228 | 0.688 | 0.863 | 0.00096324 | 1 |
| WBGene00020481 | T13C2.6 | 0.590681326 | 0.815931172 | -0.324966836 | 0.11 | 0.211 | 0.024669168 | 1 |
| WBGene00016931 | C54G6.2 | 0.231690129 | 0.456831058 | -0.324809703 | 0.04 | 0.132 | 0.003672085 | 1 |
| WBGene00010785 | top-2 | 0.135749407 | 0.360590755 | -0.324377498 | 0.029 | 0.079 | 0.041338202 | 1 |
| WBGene00000911 | daf-15 | 0.085842399 | 0.310549772 | -0.324184213 | 0.012 | 0.089 | 0.001038264 | 1 |
| WBGene00011282 | R74.8 | 0.199518952 | 0.423920312 | -0.323742729 | 0.04 | 0.105 | 0.022116121 | 1 |
| WBGene00003073 | lars-1 | 0.239475251 | 0.463477267 | -0.323166599 | 0.04 | 0.116 | 0.011214274 | 1 |
| WBGene00019157 | H05C05.1 | 0.088308394 | 0.311955801 | -0.322655005 | 0.017 | 0.074 | 0.012358958 | 1 |
| WBGene00016237 | C29H12.6 | 0.566952726 | 0.790080159 | -0.321904841 | 0.098 | 0.242 | 0.001057404 | 1 |
| WBGene00021511 | Y41D4B.1 | 0.05996387 | 0.2830399 | -0.321830683 | 0.012 | 0.068 | 0.007099123 | 1 |
| WBGene00020824 | T26A8.1 | 0.159486164 | 0.38215192 | -0.321238782 | 0.029 | 0.079 | 0.040093391 | 1 |
| WBGene00011980 | T24B8.7 | 0.19775425 | 0.420213199 | -0.320940423 | 0.029 | 0.084 | 0.028937341 | 1 |
| WBGene00002060 | ife-2 | 0.120400259 | 0.342758096 | -0.320794549 | 0.023 | 0.074 | 0.028544789 | 1 |
| WBGene00017001 | D1007.3 | 0.414829294 | 0.637126718 | -0.320707391 | 0.075 | 0.179 | 0.0084731 | 1 |
| WBGene00014016 | ZK632.9 | 1.190836261 | 1.412559298 | -0.319878726 | 0.283 | 0.421 | 0.033156355 | 1 |
| WBGene00003802 | npp-16 | 0.116811351 | 0.338406107 | -0.319693655 | 0.023 | 0.084 | 0.012205527 | 1 |
| WBGene00008672 | F11A5.4 | 0.454857534 | 0.676396999 | -0.319613887 | 0.069 | 0.179 | 0.003901891 | 1 |
| WBGene00006331 | sup-26 | 0.43144666 | 0.652827401 | -0.319384898 | 0.069 | 0.153 | 0.019290455 | 1 |
| WBGene00009966 | isy-1 | 0.244507547 | 0.465553228 | -0.318901508 | 0.052 | 0.137 | 0.009648435 | 1 |
| WBGene00020185 | pgk-1 | 0.396831711 | 0.617831783 | -0.318835708 | 0.069 | 0.163 | 0.009867239 | 1 |
| WBGene00006697 | uaf-1 | 0.25720631 | 0.477833575 | -0.318297861 | 0.046 | 0.132 | 0.006466 | 1 |
| WBGene00003519 | nac-3 | 0.900561844 | 1.120717449 | -0.3176174 | 0.179 | 0.3 | 0.04378755 | 1 |

| | | | | | | | | |
|---|---|---|---|---|---|---|---|---|
| WBGene00003162 | mdh-2 | 0.626824248 | 0.846646354 | -0.317136263 | 0.098 | 0.253 | 0.000687546 | 1 |
| WBGene00002169 | isw-1 | 0.094161565 | 0.313955138 | -0.317095097 | 0.012 | 0.068 | 0.007223737 | 1 |
| WBGene00010470 | cdr-4 | 0.897087496 | 1.116845242 | -0.317043411 | 0.156 | 0.247 | 0.050323988 | 1 |
| WBGene00016655 | acbp-1 | 0.093067422 | 0.312472611 | -0.316534779 | 0.017 | 0.084 | 0.005050668 | 1 |
| WBGene00000210 | asg-2 | 0.679272837 | 0.898558418 | -0.31636222 | 0.139 | 0.3 | 0.00204549 | 1 |
| WBGene00003511 | mxl-3 | 1.71407636 | 1.933186015 | -0.316108412 | 0.434 | 0.632 | 0.006219695 | 1 |
| WBGene00002101 | ins-18 | 0.233462066 | 0.452371505 | -0.315819562 | 0.017 | 0.053 | 0.077492615 | 1 |
| WBGene00004413 | rpl-2 | 2.521274721 | 2.740155101 | -0.315777639 | 0.717 | 0.889 | 3.13E-05 | 0.194225567 |
| WBGene00001769 | gst-21 | 1.589743109 | 1.808566235 | -0.315695038 | 0.266 | 0.389 | 0.044778408 | 1 |
| WBGene00023464 | srz-80 | 0.156326044 | 0.375094556 | -0.315616247 | 0.029 | 0.105 | 0.005477543 | 1 |
| WBGene00016937 | tag-294 | 0.165080192 | 0.383618307 | -0.315283855 | 0.029 | 0.089 | 0.019381267 | 1 |
| WBGene00002980 | lgg-1 | 2.921346454 | 3.139806498 | -0.315171223 | 0.855 | 0.958 | 0.000338212 | 1 |
| WBGene00020402 | T10B11.6 | 0.325191518 | 0.54349462 | -0.314944803 | 0.058 | 0.142 | 0.012003774 | 1 |
| WBGene00015975 | cas-2 | 0.443936525 | 0.661876971 | -0.3144216 | 0.075 | 0.205 | 0.001283761 | 1 |
| WBGene00008428 | eif-2Bepsilon | 0.053452726 | 0.270828787 | -0.313607365 | 0.012 | 0.063 | 0.011145391 | 1 |
| WBGene00000164 | apm-3 | 0.248528961 | 0.465764165 | -0.313404152 | 0.046 | 0.126 | 0.010138912 | 1 |
| WBGene00044602 | C31B8.16 | 0.099716816 | 0.316927468 | -0.313368731 | 0.017 | 0.068 | 0.019503364 | 1 |
| WBGene00000761 | coq-1 | 0.436561826 | 0.653694565 | -0.313256325 | 0.064 | 0.111 | 0.140097138 | 1 |
| WBGene00000472 | cey-1 | 1.591638618 | 1.80872586 | -0.313190688 | 0.387 | 0.647 | 0.002345271 | 1 |
| WBGene00044443 | ZC21.10 | 0.333736993 | 0.550756823 | -0.313093432 | 0.04 | 0.121 | 0.007285231 | 1 |
| WBGene00003061 | lpd-5 | 0.226232391 | 0.44316614 | -0.312969244 | 0.035 | 0.111 | 0.008521243 | 1 |
| WBGene00001955 | hlh-11 | 0.337411368 | 0.554315066 | -0.31292589 | 0.064 | 0.153 | 0.010922002 | 1 |
| WBGene00045366 | Y17D7C.3 | 0.183145457 | 0.399502343 | -0.312137006 | 0.035 | 0.116 | 0.00553395 | 1 |
| WBGene00012055 | srsx-36 | 0.435176488 | 0.651355839 | -0.311880877 | 0.087 | 0.174 | 0.024730577 | 1 |
| WBGene00009455 | phip-1 | 0.197630242 | 0.413295303 | -0.311138914 | 0.035 | 0.111 | 0.008249249 | 1 |
| WBGene00020683 | ribo-1 | 0.566454269 | 0.78204281 | -0.311028518 | 0.087 | 0.221 | 0.001243487 | 1 |
| WBGene00001803 | lite-1 | 0.223701026 | 0.439112189 | -0.310772617 | 0.035 | 0.1 | 0.017883503 | 1 |
| WBGene00004486 | rps-17 | 2.528524602 | 2.743909813 | -0.310735176 | 0.757 | 0.879 | 0.001732621 | 1 |
| WBGene00014938 | mdt-9 | 0.319404102 | 0.534783522 | -0.310726822 | 0.064 | 0.158 | 0.00812256 | 1 |
| WBGene00021347 | rpb-10 | 0.143687833 | 0.3589459 | -0.310551745 | 0.029 | 0.105 | 0.005251168 | 1 |
| WBGene00004719 | sad-1 | 0.162878815 | 0.377983871 | -0.310330998 | 0.029 | 0.074 | 0.063361313 | 1 |
| WBGene00017751 | F23F12.3 | 0.067990399 | 0.282542619 | -0.309533424 | 0.012 | 0.053 | 0.030159245 | 1 |
| WBGene00045331 | F14D7.12 | 1.591603581 | 1.805941925 | -0.309224865 | 0.312 | 0.521 | 0.002364668 | 1 |
| WBGene00016292 | tbc-7 | 0.134631865 | 0.348752793 | -0.308911201 | 0.029 | 0.095 | 0.012300723 | 1 |
| WBGene00001194 | egl-27 | 0.851590761 | 1.065387493 | -0.308443485 | 0.179 | 0.342 | 0.006275813 | 1 |
| WBGene00003808 | npr-2 | 0.316990412 | 0.530740868 | -0.308376722 | 0.046 | 0.121 | 0.014268322 | 1 |
| WBGene00006741 | unc-1 | 0.240076608 | 0.453812424 | -0.308355602 | 0.046 | 0.126 | 0.01081395 | 1 |
| WBGene00001215 | ego-2 | 0.099109646 | 0.312342397 | -0.307629833 | 0.017 | 0.084 | 0.005010169 | 1 |
| WBGene00006608 | tre-2 | 0.352678812 | 0.565867991 | -0.307566971 | 0.069 | 0.158 | 0.015226524 | 1 |
| WBGene00003949 | pbs-3 | 0.352295938 | 0.56546565 | -0.307538887 | 0.058 | 0.163 | 0.00271998 | 1 |
| WBGene00005353 | srh-136 | 0.145899179 | 0.358982199 | -0.307413816 | 0.023 | 0.084 | 0.012740227 | 1 |
| WBGene00018988 | F56F11.1 | 1.051127268 | 1.264163697 | -0.307346601 | 0.197 | 0.453 | 5.32E-05 | 0.329946197 |
| WBGene00002010 | hsp-6 | 0.14041427 | 0.353379125 | -0.307243342 | 0.023 | 0.068 | 0.043073865 | 1 |
| WBGene00001005 | dlc-1 | 1.152727606 | 1.365298554 | -0.306675052 | 0.266 | 0.484 | 0.001998713 | 1 |
| WBGene00013994 | ZK524.4 | 0.698256229 | 0.910643215 | -0.306409651 | 0.116 | 0.268 | 0.001392241 | 1 |
| WBGene00001898 | his-24 | 0.553211139 | 0.764866247 | -0.305353775 | 0.098 | 0.189 | 0.026022938 | 1 |
| WBGene00015251 | B0546.4 | 0.698922233 | 0.909411225 | -0.303671426 | 0.145 | 0.3 | 0.004119722 | 1 |
| WBGene00015938 | anat-1 | 0.362955157 | 0.573266134 | -0.303414603 | 0.04 | 0.116 | 0.011843322 | 1 |
| WBGene00007885 | ugt-21 | 0.332691283 | 0.542788551 | -0.303106287 | 0.017 | 0.084 | 0.004930051 | 1 |
| WBGene00003724 | nhr-134 | 0.447657274 | 0.657667093 | -0.302980124 | 0.058 | 0.179 | 0.000927534 | 1 |
| WBGene00014254 | cyp-13A10 | 0.139152903 | 0.348781871 | -0.302430672 | 0.017 | 0.079 | 0.008042287 | 1 |
| WBGene00015249 | otub-4 | 0.099144582 | 0.308766037 | -0.302419833 | 0.017 | 0.068 | 0.020098499 | 1 |
| WBGene00003710 | nhr-120 | 0.646985681 | 0.856290005 | -0.301962312 | 0.116 | 0.289 | 0.00040002 | 1 |

| | | | | | | | | |
|---|---|---|---|---|---|---|---|---|
| WBGene00022732 | ZK418.2 | 0.867160623 | 1.076452849 | -0.301944856 | 0.145 | 0.326 | 0.00058897 | 1 |
| WBGene00004438 | rpl-25.1 | 1.15553129 | 1.364806513 | -0.301920325 | 0.249 | 0.511 | 0.000190037 | 1 |
| WBGene00019679 | spcs-3 | 0.147265426 | 0.356263399 | -0.301520339 | 0.029 | 0.084 | 0.028756087 | 1 |
| WBGene00001677 | gpa-15 | 0.905002751 | 1.113699598 | -0.301085906 | 0.15 | 0.347 | 0.000261583 | 1 |
| WBGene00000001 | aap-1 | 0.138563757 | 0.34721054 | -0.301013679 | 0.023 | 0.058 | 0.105744875 | 1 |
| WBGene00015667 | C10A4.6 | 0.032010825 | 0.239895204 | -0.299913763 | 0.006 | 0.063 | 0.003505266 | 1 |
| WBGene00007514 | catp-8 | 0.531819802 | 0.739603736 | -0.299768851 | 0.081 | 0.179 | 0.013667311 | 1 |
| WBGene00004444 | rpl-30 | 1.804257196 | 2.011619872 | -0.299161105 | 0.445 | 0.721 | 0.001348872 | 1 |
| WBGene00018519 | F46H5.3 | 1.029653099 | 1.236628048 | -0.298601731 | 0.185 | 0.384 | 0.001050154 | 1 |
| WBGene00077491 | K08H10.10 | 0.839123639 | 1.046071473 | -0.298562614 | 0.173 | 0.311 | 0.011371548 | 1 |
| WBGene00008208 | nhr-167 | 0.708169876 | 0.915029977 | -0.298436041 | 0.098 | 0.295 | 3.36E-05 | 0.208122692 |
| WBGene00021346 | Y37E3.1 | 0.306586741 | 0.513340615 | -0.298282788 | 0.052 | 0.153 | 0.003537078 | 1 |
| WBGene00021466 | eif-2gamma | 0.120676901 | 0.327140563 | -0.2978641 | 0.023 | 0.1 | 0.003504332 | 1 |
| WBGene00012644 | rabx-5 | 0.640802272 | 0.847232218 | -0.29781546 | 0.121 | 0.247 | 0.004983877 | 1 |
| WBGene00014233 | ZK1128.7 | 0.481852341 | 0.688178164 | -0.297665242 | 0.081 | 0.211 | 0.001557856 | 1 |
| WBGene00020181 | T02H6.11 | 0.829103879 | 1.035405135 | -0.297629798 | 0.179 | 0.358 | 0.001957423 | 1 |
| WBGene00015048 | faah-2 | 0.542180556 | 0.74814583 | -0.29714508 | 0.087 | 0.211 | 0.003212043 | 1 |
| WBGene00007023 | mdt-27 | 0 | 0.205919328 | -0.297078793 | 0 | 0.058 | 0.001340226 | 1 |
| WBGene00015644 | C09E8.1 | 0.097887748 | 0.303731216 | -0.29696935 | 0.017 | 0.068 | 0.019650686 | 1 |
| WBGene00001007 | dli-1 | 0.538282942 | 0.7432631 | -0.295723858 | 0.116 | 0.237 | 0.011359134 | 1 |
| WBGene00004472 | rps-3 | 2.084363318 | 2.289329406 | -0.295703559 | 0.561 | 0.805 | 0.000105292 | 0.652705595 |
| WBGene00012372 | W09H1.1 | 1.258381208 | 1.462927815 | -0.295098375 | 0.289 | 0.511 | 0.003961418 | 1 |
| WBGene00004497 | rps-28 | 2.719512731 | 2.924044524 | -0.295077004 | 0.809 | 0.921 | 0.001443187 | 1 |
| WBGene00009912 | srz-94 | 0.891486627 | 1.095585971 | -0.294453111 | 0.162 | 0.337 | 0.001462062 | 1 |
| WBGene00002222 | klp-11 | 0.177139003 | 0.381041841 | -0.294169614 | 0.035 | 0.095 | 0.024626872 | 1 |
| WBGene00000137 | amx-1 | 0.092595241 | 0.296478165 | -0.294140883 | 0.017 | 0.068 | 0.019650686 | 1 |
| WBGene00005667 | sru-4 | 0.033073219 | 0.23651393 | -0.293502905 | 0.006 | 0.063 | 0.003540411 | 1 |
| WBGene00008237 | nstp-10 | 0.261202154 | 0.463941053 | -0.292490404 | 0.046 | 0.084 | 0.172671711 | 1 |
| WBGene00000236 | bag-1 | 0.319486416 | 0.521851759 | -0.291951478 | 0.052 | 0.142 | 0.006844935 | 1 |
| WBGene00045514 | ZK1037.12 | 1.316484274 | 1.518709113 | -0.291748773 | 0.283 | 0.468 | 0.004646872 | 1 |
| WBGene00011326 | T01D3.1 | 1.567673496 | 1.769279756 | -0.290856351 | 0.335 | 0.589 | 0.000625947 | 1 |
| WBGene00021610 | nhr-237 | 0.761923849 | 0.963514758 | -0.290834205 | 0.168 | 0.295 | 0.01874279 | 1 |
| WBGene00012907 | cpt-1 | 0.80981622 | 1.011336841 | -0.290732801 | 0.156 | 0.305 | 0.006145769 | 1 |
| WBGene00002070 | ile-1 | 0.154351972 | 0.355843358 | -0.290690622 | 0.035 | 0.089 | 0.039071283 | 1 |
| WBGene00004423 | rpl-11.2 | 2.664803303 | 2.866139353 | -0.290466521 | 0.746 | 0.926 | 0.006245591 | 1 |
| WBGene00020100 | mks-1 | 0.699527695 | 0.900679776 | -0.29020111 | 0.15 | 0.253 | 0.061034029 | 1 |
| WBGene00019993 | rbx-2 | 0.089171656 | 0.290066605 | -0.289830147 | 0.012 | 0.079 | 0.002715427 | 1 |
| WBGene00016501 | C37C3.9 | 0.099537568 | 0.30030512 | -0.289646352 | 0.017 | 0.053 | 0.072905566 | 1 |
| WBGene00013606 | cand-1 | 0.104976713 | 0.305274282 | -0.288968309 | 0.017 | 0.063 | 0.030606713 | 1 |
| WBGene00006888 | vbh-1 | 0.112776477 | 0.312782267 | -0.288547362 | 0.023 | 0.089 | 0.007738189 | 1 |
| WBGene00003403 | mps-1 | 0.493674233 | 0.693531094 | -0.288332502 | 0.098 | 0.179 | 0.056304793 | 1 |
| WBGene00004463 | rpn-7 | 0.67904094 | 0.87882417 | -0.288226275 | 0.15 | 0.232 | 0.10502547 | 1 |
| WBGene00006910 | vha-1 | 1.025938377 | 1.225490264 | -0.287892518 | 0.214 | 0.432 | 0.000959083 | 1 |
| WBGene00015511 | C06A6.7 | 0.031940067 | 0.231293823 | -0.287606675 | 0.006 | 0.058 | 0.005900907 | 1 |
| WBGene00001748 | gsp-2 | 0.527610833 | 0.726895439 | -0.287506913 | 0.104 | 0.2 | 0.030549681 | 1 |
| WBGene00004488 | rps-19 | 2.700489938 | 2.899157377 | -0.286616529 | 0.757 | 0.926 | 0.004895881 | 1 |
| WBGene00013202 | Y54E5A.7 | 0.261354053 | 0.459787749 | -0.286279309 | 0.046 | 0.084 | 0.16832072 | 1 |
| WBGene00004925 | snt-5 | 0.371700809 | 0.570055218 | -0.286164922 | 0.035 | 0.121 | 0.003331672 | 1 |
| WBGene00008419 | wdr-23 | 0.501269084 | 0.699436073 | -0.285894531 | 0.092 | 0.205 | 0.006443102 | 1 |
| WBGene00219816 | C14E2.12 | 0.249246137 | 0.447412884 | -0.285894184 | 0.046 | 0.063 | 0.471177555 | 1 |
| WBGene00019200 | H14E04.3 | 0.423057472 | 0.620417349 | -0.284730116 | 0.029 | 0.121 | 0.001340577 | 1 |
| WBGene00019980 | chil-14 | 0.153126867 | 0.350312315 | -0.284478467 | 0.017 | 0.068 | 0.019948248 | 1 |
| WBGene00003627 | nhr-34 | 0.378570177 | 0.575662906 | -0.284344703 | 0.064 | 0.168 | 0.003895573 | 1 |

| | | | | | | | | |
|---|---|---|---|---|---|---|---|---|
| WBGene00004476 | rps-7 | 1.97216798 | 2.169190804 | -0.284243851 | 0.509 | 0.795 | 0.000145302 | 0.900729308 |
| WBGene00000554 | cnb-1 | 2.259630927 | 2.456578171 | -0.284134813 | 0.601 | 0.821 | 0.000801165 | 1 |
| WBGene00004284 | rab-35 | 0.197913833 | 0.394733213 | -0.283950344 | 0.035 | 0.1 | 0.018105245 | 1 |
| WBGene00010405 | erfa-3 | 0.18997749 | 0.386593935 | -0.283657569 | 0.035 | 0.1 | 0.017232493 | 1 |
| WBGene00020994 | W03F8.4 | 0.133487158 | 0.330092679 | -0.28364181 | 0.017 | 0.089 | 0.00315817 | 1 |
| WBGene00017698 | F22B7.9 | 0.783270418 | 0.979763747 | -0.283479951 | 0.116 | 0.147 | 0.477548306 | 1 |
| WBGene00004800 | sir-2.1 | 0.176128446 | 0.372198121 | -0.282868747 | 0.029 | 0.111 | 0.003291425 | 1 |
| WBGene00011682 | snt-7 | 0.401598899 | 0.597626647 | -0.28280826 | 0.058 | 0.137 | 0.017317523 | 1 |
| WBGene00015507 | C06A6.2 | 0.254847849 | 0.450640347 | -0.282468866 | 0.04 | 0.132 | 0.003329192 | 1 |
| WBGene00002191 | kin-3 | 0.642716482 | 0.838417093 | -0.282336301 | 0.133 | 0.268 | 0.00622806 | 1 |
| WBGene00013322 | Y57G11C.32 | 0.062587716 | 0.257939212 | -0.281832634 | 0.012 | 0.068 | 0.007223737 | 1 |
| WBGene00022370 | Y92H12BR.4 | 0.145902794 | 0.341217597 | -0.281779697 | 0.017 | 0.063 | 0.030606713 | 1 |
| WBGene00016519 | srz-56 | 0.49018062 | 0.685379191 | -0.281612011 | 0.092 | 0.142 | 0.173322461 | 1 |
| WBGene00019764 | M03F8.5 | 0.089889659 | 0.284858708 | -0.28128088 | 0.017 | 0.084 | 0.005010169 | 1 |
| WBGene00022465 | ift-20 | 1.709737547 | 1.904673885 | -0.281233689 | 0.434 | 0.695 | 0.000186504 | 1 |
| WBGene00007144 | B0334.4 | 0.09828619 | 0.293026833 | -0.28095136 | 0.017 | 0.084 | 0.005173946 | 1 |
| WBGene00018582 | F48A9.2 | 0.309993634 | 0.504499333 | -0.280612408 | 0.046 | 0.121 | 0.016357083 | 1 |
| WBGene00004480 | rps-11 | 2.743047531 | 2.937293107 | -0.28023713 | 0.763 | 0.884 | 0.006159024 | 1 |
| WBGene00016833 | sre-51 | 0.121504109 | 0.315721478 | -0.280196434 | 0.023 | 0.084 | 0.013015271 | 1 |
| WBGene00023411 | srz-3 | 0.952585346 | 1.146424233 | -0.279650401 | 0.191 | 0.384 | 0.001614278 | 1 |
| WBGene00003858 | ogt-1 | 0.711613603 | 0.905265745 | -0.279380985 | 0.116 | 0.274 | 0.001148896 | 1 |
| WBGene00009078 | rpb-12 | 0.382894992 | 0.57654197 | -0.279373535 | 0.035 | 0.137 | 0.000905204 | 1 |
| WBGene00021597 | spsb-1 | 0.335662613 | 0.529272705 | -0.27932032 | 0.052 | 0.1 | 0.107974812 | 1 |
| WBGene00018892 | cir-1 | 0.148438191 | 0.341853818 | -0.279039766 | 0.023 | 0.058 | 0.10449823 | 1 |
| WBGene00004068 | pnk-1 | 0.035381258 | 0.228653016 | -0.278832208 | 0.006 | 0.053 | 0.009737833 | 1 |
| WBGene00009976 | swan-2 | 0.15417237 | 0.347341131 | -0.278683612 | 0.029 | 0.084 | 0.029487017 | 1 |
| WBGene00011060 | R06C1.6 | 0.713603295 | 0.906665431 | -0.278529786 | 0.15 | 0.258 | 0.037200208 | 1 |
| WBGene00016002 | gyf-1 | 1.176979637 | 1.36968976 | -0.278021939 | 0.272 | 0.442 | 0.015907967 | 1 |
| WBGene00000197 | aars-2 | 0.09036904 | 0.283027203 | -0.277946977 | 0.017 | 0.068 | 0.019948248 | 1 |
| WBGene00000262 | bra-1 | 0.313688082 | 0.506006656 | -0.277457052 | 0.052 | 0.121 | 0.027134267 | 1 |
| WBGene00004420 | rpl-9 | 2.123789507 | 2.315780224 | -0.276984055 | 0.595 | 0.8 | 0.010462509 | 1 |
| WBGene00004481 | rps-12 | 2.177303359 | 2.369147844 | -0.276773087 | 0.601 | 0.795 | 0.00461618 | 1 |
| WBGene00002182 | kap-1 | 0.345643823 | 0.537412442 | -0.276663636 | 0.064 | 0.147 | 0.015392191 | 1 |
| WBGene00017635 | F20D6.5 | 0.059159909 | 0.250672338 | -0.276294032 | 0.012 | 0.068 | 0.007099123 | 1 |
| WBGene00005428 | srh-218 | 0.089240396 | 0.280458372 | -0.275869225 | 0.012 | 0.079 | 0.002739891 | 1 |
| WBGene00019117 | nhr-144 | 0.636456322 | 0.827531002 | -0.275662493 | 0.104 | 0.232 | 0.005100348 | 1 |
| WBGene00008204 | C49C3.15 | 0.118285491 | 0.309282378 | -0.275550261 | 0.023 | 0.079 | 0.019658024 | 1 |
| WBGene00017068 | D2092.8 | 0.117506304 | 0.308455354 | -0.275481248 | 0.012 | 0.063 | 0.011933937 | 1 |
| WBGene00004087 | ppk-1 | 0.416870997 | 0.607755894 | -0.275388695 | 0.087 | 0.174 | 0.03001579 | 1 |
| WBGene00006579 | tlk-1 | 0.402859576 | 0.593485829 | -0.27501555 | 0.064 | 0.147 | 0.014848247 | 1 |
| WBGene00006770 | unc-34 | 0.027646995 | 0.217921926 | -0.274508698 | 0.006 | 0.063 | 0.003540411 | 1 |
| WBGene00001225 | eif-3.B | 0.370482695 | 0.5605461 | -0.274203532 | 0.069 | 0.147 | 0.02882081 | 1 |
| WBGene00009180 | nurf-1 | 1.580863506 | 1.770812081 | -0.274037867 | 0.329 | 0.526 | 0.010313714 | 1 |
| WBGene00002238 | kars-1 | 0.073493172 | 0.263230983 | -0.2737338 | 0.012 | 0.063 | 0.011832736 | 1 |
| WBGene00012951 | Y47H9C.8 | 0 | 0.189609073 | -0.27354807 | 0 | 0.053 | 0.002261451 | 1 |
| WBGene00011218 | R10E11.6 | 0.021822539 | 0.211050913 | -0.272998837 | 0.006 | 0.058 | 0.005560396 | 1 |
| WBGene00001086 | dpy-27 | 0.470415177 | 0.65962313 | -0.272969375 | 0.087 | 0.153 | 0.084658249 | 1 |
| WBGene00007246 | nspb-10 | 0.636642028 | 0.825837064 | -0.27295074 | 0.081 | 0.184 | 0.008154388 | 1 |
| WBGene00018639 | tbc-16 | 0.669656712 | 0.858673693 | -0.272693861 | 0.139 | 0.268 | 0.009051245 | 1 |
| WBGene00008920 | eef-1G | 2.101543308 | 2.290535675 | -0.27265835 | 0.618 | 0.842 | 0.000620962 | 1 |
| WBGene00008528 | F02E9.1 | 0.210375178 | 0.399357815 | -0.272644314 | 0.023 | 0.111 | 0.001426176 | 1 |
| WBGene00009974 | F53C11.4 | 0.546685518 | 0.735638225 | -0.272601133 | 0.092 | 0.216 | 0.003417761 | 1 |
| WBGene00020184 | uba-5 | 0.098674238 | 0.287515446 | -0.272440274 | 0.017 | 0.074 | 0.012746181 | 1 |

| | | | | | | | | |
|---|---|---|---|---|---|---|---|---|
| WBGene00018995 | ccpp-1 | 0.262897689 | 0.451299474 | -0.271806321 | 0.046 | 0.095 | 0.085145311 | 1 |
| WBGene00002233 | kqt-1 | 0.602895919 | 0.791212649 | -0.271683612 | 0.092 | 0.179 | 0.033826098 | 1 |
| WBGene00021304 | nphp-2 | 0.53327605 | 0.721474555 | -0.271513049 | 0.116 | 0.2 | 0.060292442 | 1 |
| WBGene00001992 | hot-7 | 0.351411753 | 0.539480518 | -0.271325875 | 0.052 | 0.137 | 0.008606747 | 1 |
| WBGene00006515 | txdc-9 | 0.346150296 | 0.533942717 | -0.270927193 | 0.058 | 0.179 | 0.001147717 | 1 |
| WBGene00017150 | EGAP5.1 | 2.510876154 | 2.69824168 | -0.270311316 | 0.728 | 0.847 | 0.01051197 | 1 |
| WBGene00016405 | hecd-1 | 0.597815791 | 0.784955555 | -0.269985608 | 0.116 | 0.232 | 0.014129608 | 1 |
| WBGene00000158 | apg-1 | 0.223844398 | 0.41071589 | -0.269598575 | 0.029 | 0.089 | 0.019007681 | 1 |
| WBGene00009940 | strl-1 | 0.764790974 | 0.951367748 | -0.269173386 | 0.145 | 0.237 | 0.055523429 | 1 |
| WBGene00021973 | emc-2 | 0.150484143 | 0.337052339 | -0.26916101 | 0.023 | 0.074 | 0.029925064 | 1 |
| WBGene00021402 | Y38C1AA.12 | 0.087903108 | 0.274195831 | -0.268763588 | 0.017 | 0.074 | 0.013043606 | 1 |
| WBGene00019005 | F57B10.8 | 0.055496294 | 0.240872644 | -0.267441541 | 0.012 | 0.058 | 0.018681607 | 1 |
| WBGene00017568 | F18E9.1 | 0.17535282 | 0.360533161 | -0.267158759 | 0.029 | 0.089 | 0.019007681 | 1 |
| WBGene00017926 | cox-6C | 1.110859233 | 1.295677895 | -0.266636966 | 0.254 | 0.479 | 0.00112661 | 1 |
| WBGene00003656 | nhr-66 | 0.672543573 | 0.857315108 | -0.266568977 | 0.133 | 0.216 | 0.072168048 | 1 |
| WBGene00008447 | E01G4.5 | 0.180787508 | 0.365373219 | -0.26630089 | 0.035 | 0.084 | 0.05906811 | 1 |
| WBGene00014862 | Y7A9C.2 | 0.25386787 | 0.438296076 | -0.266073658 | 0.046 | 0.074 | 0.289382408 | 1 |
| WBGene00004885 | smg-7 | 0.108120717 | 0.292429167 | -0.265900887 | 0.017 | 0.053 | 0.073905225 | 1 |
| WBGene00019053 | F58F6.5 | 0.202239456 | 0.386465265 | -0.265781662 | 0.023 | 0.095 | 0.005414509 | 1 |
| WBGene00002025 | hsp-60 | 0.029646469 | 0.213615797 | -0.265411637 | 0.006 | 0.058 | 0.005900907 | 1 |
| WBGene00012548 | Y37D8A.6 | 2.138079343 | 2.321947341 | -0.265265449 | 0.549 | 0.784 | 0.002506213 | 1 |
| WBGene00003645 | nhr-55 | 0.102274781 | 0.286108092 | -0.265215407 | 0.012 | 0.068 | 0.007414341 | 1 |
| WBGene00007918 | sphk-1 | 0.348001142 | 0.531795696 | -0.265159491 | 0.046 | 0.126 | 0.010258782 | 1 |
| WBGene00005327 | srh-108 | 0.516859445 | 0.700620705 | -0.265111458 | 0.069 | 0.211 | 0.000420304 | 1 |
| WBGene00004926 | snt-6 | 0.454395518 | 0.637735458 | -0.264503622 | 0.087 | 0.189 | 0.010844962 | 1 |
| WBGene00004215 | ptp-3 | 0.650111069 | 0.833370479 | -0.264387442 | 0.116 | 0.237 | 0.008362189 | 1 |
| WBGene00015810 | C16A3.5 | 0.226089334 | 0.408955884 | -0.263820664 | 0.04 | 0.105 | 0.024111591 | 1 |
| WBGene00001837 | hda-4 | 0.133631438 | 0.316377127 | -0.263646299 | 0.023 | 0.074 | 0.03094576 | 1 |
| WBGene00006572 | tin-9.1 | 0.452651525 | 0.635127287 | -0.263256877 | 0.087 | 0.168 | 0.037384456 | 1 |
| WBGene00009216 | F28D1.6 | 0.563869531 | 0.746327941 | -0.263231844 | 0.075 | 0.168 | 0.012208009 | 1 |
| WBGene00008144 | gasr-8 | 0.490715067 | 0.672992345 | -0.262970525 | 0.092 | 0.174 | 0.040822233 | 1 |
| WBGene00021643 | ccep-290 | 0.241464825 | 0.423693651 | -0.262900624 | 0.046 | 0.089 | 0.123029953 | 1 |
| WBGene00004495 | rps-26 | 3.000661978 | 3.182582334 | -0.262455595 | 0.85 | 0.947 | 0.002258213 | 1 |
| WBGene00017157 | tyra-2 | 0.192423674 | 0.374162111 | -0.262193142 | 0.029 | 0.105 | 0.005477543 | 1 |
| WBGene00019807 | jamp-1 | 0.079094065 | 0.260716885 | -0.262026341 | 0.017 | 0.053 | 0.074916061 | 1 |
| WBGene00018466 | F45E1.5 | 0.02790056 | 0.209449375 | -0.261919576 | 0.006 | 0.053 | 0.009737833 | 1 |
| WBGene00001817 | haf-7 | 0 | 0.18152392 | -0.261883659 | 0 | 0.053 | 0.002261451 | 1 |
| WBGene00019896 | R05G6.1 | 1.830958075 | 2.012340962 | -0.261680192 | 0.422 | 0.632 | 0.013225327 | 1 |
| WBGene00021736 | wrb-1 | 0 | 0.181367602 | -0.26165814 | 0 | 0.053 | 0.002261451 | 1 |
| WBGene00007646 | nkb-1 | 0.663133809 | 0.844497717 | -0.261652811 | 0.11 | 0.221 | 0.015803103 | 1 |
| WBGene00012615 | dct-16 | 0.39568974 | 0.577026387 | -0.261613482 | 0.075 | 0.074 | 0.925458682 | 1 |
| WBGene00004212 | ptl-1 | 0.227728234 | 0.408829669 | -0.261274143 | 0.04 | 0.116 | 0.011630247 | 1 |
| WBGene00003948 | pbs-2 | 0.740026134 | 0.921066001 | -0.26118532 | 0.145 | 0.284 | 0.005329442 | 1 |
| WBGene00010564 | ath-1 | 0.068951593 | 0.249923951 | -0.261087923 | 0.012 | 0.058 | 0.018681536 | 1 |
| WBGene00009740 | F45H10.3 | 0.295131573 | 0.476076695 | -0.261048631 | 0.046 | 0.132 | 0.007563699 | 1 |
| WBGene00015527 | C06E2.1 | 0.29681561 | 0.477754098 | -0.261039059 | 0.052 | 0.116 | 0.041319614 | 1 |
| WBGene00022092 | Y69A2AR.21 | 0.05533364 | 0.23624577 | -0.261001033 | 0.006 | 0.058 | 0.005959452 | 1 |
| WBGene00002076 | imb-2 | 0.265146943 | 0.446024974 | -0.260951839 | 0.046 | 0.111 | 0.030939696 | 1 |
| WBGene00011551 | T06G6.12 | 0.542838539 | 0.723537021 | -0.260692804 | 0.081 | 0.179 | 0.011115644 | 1 |
| WBGene00001168 | eef-1A.1 | 2.239039527 | 2.419700745 | -0.260639043 | 0.595 | 0.826 | 0.004757065 | 1 |
| WBGene00008430 | hgap-2 | 0.167217265 | 0.347849372 | -0.260597046 | 0.029 | 0.095 | 0.012804276 | 1 |
| WBGene00012528 | pap-1 | 0.623092801 | 0.803031248 | -0.259596306 | 0.121 | 0.237 | 0.012754403 | 1 |
| WBGene00019245 | sacy-1 | 0.101601273 | 0.281483552 | -0.259515272 | 0.017 | 0.058 | 0.048752734 | 1 |

| | | | | | | | | |
|---|---|---|---|---|---|---|---|---|
| WBGene00009915 | F52A8.1 | 0 | 0.179851657 | -0.259471094 | 0 | 0.053 | 0.002261451 | 1 |
| WBGene00004439 | rpl-25.2 | 1.119475543 | 1.299215963 | -0.259310613 | 0.249 | 0.411 | 0.017288817 | 1 |
| WBGene00009191 | F27D4.7 | 0.037541523 | 0.217246057 | -0.25925884 | 0.006 | 0.063 | 0.003540411 | 1 |
| WBGene00013365 | Y60A3A.19 | 0.280614517 | 0.460313456 | -0.259250768 | 0.052 | 0.132 | 0.01405677 | 1 |
| WBGene00000881 | cyn-5 | 0.665933237 | 0.84498496 | -0.258317033 | 0.104 | 0.268 | 0.000400205 | 1 |
| WBGene00302980 | F53E2.2 | 0.529262849 | 0.707769548 | -0.257530729 | 0.098 | 0.211 | 0.009342497 | 1 |
| WBGene00000833 | cts-1 | 0.670637186 | 0.848900872 | -0.257180137 | 0.127 | 0.247 | 0.010693247 | 1 |
| WBGene00017571 | jmjd-3.1 | 1.099118441 | 1.276940767 | -0.256543388 | 0.254 | 0.484 | 0.002640882 | 1 |
| WBGene00013075 | ints-5 | 0.046712372 | 0.224518159 | -0.256519527 | 0.012 | 0.058 | 0.017914602 | 1 |
| WBGene00016096 | C25E10.7 | 3.39899351 | 3.57669094 | -0.2563632 | 0.867 | 0.958 | 0.005336776 | 1 |
| WBGene00004187 | prp-8 | 0.773879908 | 0.951387566 | -0.256089417 | 0.133 | 0.274 | 0.005764964 | 1 |
| WBGene00016792 | C49H3.6 | 0.624168729 | 0.801574861 | -0.255942947 | 0.116 | 0.216 | 0.027341334 | 1 |
| WBGene00020275 | atp-4 | 0.758161723 | 0.935551045 | -0.255918696 | 0.156 | 0.295 | 0.011176083 | 1 |
| WBGene00004385 | rnp-2 | 0.219389808 | 0.396530474 | -0.255559961 | 0.04 | 0.1 | 0.034644049 | 1 |
| WBGene00004426 | rpl-14 | 2.437008729 | 2.614049239 | -0.255415467 | 0.723 | 0.889 | 0.002038004 | 1 |
| WBGene00021039 | W05F2.7 | 1.98574662 | 2.162740769 | -0.255348582 | 0.532 | 0.763 | 0.032619477 | 1 |
| WBGene00016194 | C28H8.3 | 0.375266453 | 0.552157426 | -0.25519973 | 0.064 | 0.121 | 0.077674912 | 1 |
| WBGene00021558 | Y45G5AM.6 | 0.295798573 | 0.472447399 | -0.254850385 | 0.035 | 0.116 | 0.005141982 | 1 |
| WBGene00044367 | T05A8.8 | 0.937638465 | 1.114282367 | -0.254843282 | 0.197 | 0.405 | 0.001012158 | 1 |
| WBGene00001393 | fat-1 | 0.370897443 | 0.547330667 | -0.254539337 | 0.069 | 0.153 | 0.020648128 | 1 |
| WBGene00004425 | rpl-13 | 2.191110564 | 2.367220412 | -0.254072805 | 0.613 | 0.816 | 0.005384255 | 1 |
| WBGene00015776 | dct-1 | 1.744306591 | 1.920262657 | -0.253850945 | 0.462 | 0.7 | 0.008212475 | 1 |
| WBGene00000882 | cyn-6 | 0.287886662 | 0.463586608 | -0.253481442 | 0.035 | 0.132 | 0.00145298 | 1 |
| WBGene00015115 | B0286.1 | 0.192216657 | 0.36785643 | -0.253394629 | 0.035 | 0.095 | 0.02584019 | 1 |
| WBGene00003790 | npp-4 | 0.027333216 | 0.202793369 | -0.253135491 | 0.006 | 0.053 | 0.009737833 | 1 |
| WBGene00005965 | srx-74 | 0.643546911 | 0.818975327 | -0.253089706 | 0.121 | 0.247 | 0.008518993 | 1 |
| WBGene00018213 | F39H12.2 | 1.899856741 | 2.075271165 | -0.253069521 | 0.514 | 0.726 | 0.011864164 | 1 |
| WBGene00012097 | abcf-2 | 0.546168233 | 0.721549675 | -0.253021938 | 0.098 | 0.226 | 0.003273253 | 1 |
| WBGene00006468 | rhgf-1 | 0.8033571 | 0.978724917 | -0.253002281 | 0.173 | 0.342 | 0.005116431 | 1 |
| WBGene00016118 | C25H3.9 | 0.227425246 | 0.402476104 | -0.252545005 | 0.046 | 0.111 | 0.032118211 | 1 |
| WBGene00003372 | mlc-4 | 0.357376976 | 0.532263025 | -0.252307237 | 0.064 | 0.142 | 0.022738972 | 1 |
| WBGene00021033 | srz-32 | 0.070370042 | 0.24517807 | -0.252194676 | 0.012 | 0.053 | 0.030406126 | 1 |
| WBGene00017999 | emc-6 | 0.270459008 | 0.445266432 | -0.252193804 | 0.052 | 0.111 | 0.058940921 | 1 |
| WBGene00023513 | srz-71 | 0.122040779 | 0.296816818 | -0.252148525 | 0.017 | 0.063 | 0.030606713 | 1 |
| WBGene00017789 | F25E5.8 | 0.237941429 | 0.412680638 | -0.25209539 | 0.046 | 0.111 | 0.035699538 | 1 |
| WBGene00006066 | sto-4 | 0.05488738 | 0.22953654 | -0.251965477 | 0.012 | 0.058 | 0.018996363 | 1 |
| WBGene00010631 | cash-1 | 0.29978129 | 0.474426676 | -0.251960032 | 0.035 | 0.116 | 0.0053884 | 1 |
| WBGene00009627 | cup-15 | 0 | 0.174531885 | -0.251796286 | 0 | 0.053 | 0.002261451 | 1 |
| WBGene00008932 | F18A11.5 | 0.370489391 | 0.544992905 | -0.251755355 | 0.075 | 0.126 | 0.133572271 | 1 |
| WBGene00013379 | Y62E10A.13 | 0.089128082 | 0.263239404 | -0.25118954 | 0.017 | 0.058 | 0.047731237 | 1 |
| WBGene00001208 | egl-44 | 1.140986729 | 1.314956538 | -0.250985382 | 0.243 | 0.374 | 0.040150086 | 1 |
| WBGene00008185 | best-10 | 0.281144623 | 0.45483061 | -0.250575913 | 0.046 | 0.1 | 0.061929905 | 1 |
| WBGene00004412 | rpl-1 | 2.272769345 | 2.446439824 | -0.250553538 | 0.694 | 0.884 | 0.010944192 | 1 |
| WBGene00020475 | sut-1 | 0.244025856 | 0.417669018 | -0.25051413 | 0.046 | 0.095 | 0.097144695 | 1 |
| WBGene00012461 | glb-29 | 0.507429954 | 0.680954942 | -0.25034364 | 0.081 | 0.174 | 0.016014783 | 1 |
| WBGene00008643 | tmem-138 | 0.121460161 | 0.294956784 | -0.250302718 | 0.023 | 0.074 | 0.030126856 | 1 |
| WBGene00017359 | F10E9.10 | 0.309668953 | 0.483083289 | -0.250184003 | 0.04 | 0.1 | 0.036509048 | 1 |
| WBGene00006434 | prdx-2 | 1.741740839 | 1.9151167 | -0.250128495 | 0.457 | 0.689 | 0.004723116 | 1 |
| WBGene00012440 | nlp-53 | 1.584059878 | 1.757106791 | -0.249653922 | 0.341 | 0.621 | 0.00031063 | 1 |
| WBGene00022733 | tmem-17 | 0.674775929 | 0.847400047 | -0.249043959 | 0.127 | 0.174 | 0.332220142 | 1 |
| WBGene00206380 | C42D4.18 | 0.578503148 | 0.751124865 | -0.249040495 | 0.116 | 0.211 | 0.033463632 | 1 |
| WBGene00013195 | Y54E2A.10 | 0.560822115 | 0.732799759 | -0.248111293 | 0.081 | 0.179 | 0.011333726 | 1 |
| WBGene00007197 | prdh-1 | 0.297933483 | 0.469879807 | -0.248066109 | 0.058 | 0.105 | 0.129163533 | 1 |

| | | | | | | | | |
|---|---|---|---|---|---|---|---|---|
| WBGene00011449 | T04H1.2 | 0.145997303 | 0.317689684 | -0.247699746 | 0.029 | 0.1 | 0.008873331 | 1 |
| WBGene00003598 | nhl-2 | 0.227034878 | 0.398680011 | -0.247631583 | 0.046 | 0.116 | 0.023570827 | 1 |
| WBGene00001464 | flp-21 | 2.1399125 | 2.3111297 | -0.247014205 | 0.624 | 0.811 | 0.010762096 | 1 |
| WBGene00008856 | F15D3.6 | 0.067145984 | 0.238171201 | -0.246737232 | 0.012 | 0.063 | 0.011933937 | 1 |
| WBGene00011720 | T11G6.5 | 0.179875109 | 0.35060125 | -0.246305756 | 0.029 | 0.089 | 0.018884559 | 1 |
| WBGene00000835 | cuc-1 | 0.202494879 | 0.373177166 | -0.246242489 | 0.04 | 0.1 | 0.034548247 | 1 |
| WBGene00010910 | M106.2 | 0.893565896 | 1.064111847 | -0.246045798 | 0.133 | 0.284 | 0.002441258 | 1 |
| WBGene00009164 | hrdl-1 | 0.559574441 | 0.728968707 | -0.244384268 | 0.098 | 0.179 | 0.047338523 | 1 |
| WBGene00009161 | cox-7C | 0.983125948 | 1.152426417 | -0.244248946 | 0.191 | 0.426 | 0.000275382 | 1 |
| WBGene00000105 | alg-1 | 0.299511025 | 0.468727484 | -0.244127747 | 0.046 | 0.116 | 0.021824073 | 1 |
| WBGene00002016 | hsp-16.2 | 3.754549041 | 3.923518467 | -0.243771353 | 0.52 | 0.663 | 0.013572629 | 1 |
| WBGene00001029 | dnj-11 | 0.077870601 | 0.246580453 | -0.243396866 | 0.017 | 0.053 | 0.071917 | 1 |
| WBGene00016331 | obr-4 | 0.085749026 | 0.254389733 | -0.243297112 | 0.017 | 0.068 | 0.019503364 | 1 |
| WBGene00019156 | H04M03.12 | 0.930091204 | 1.098713172 | -0.243270078 | 0.179 | 0.326 | 0.006872228 | 1 |
| WBGene00018075 | tbc-11 | 0.146153679 | 0.314774874 | -0.243268962 | 0.023 | 0.084 | 0.012877063 | 1 |
| WBGene00008034 | prcc-1 | 0.68952504 | 0.858113888 | -0.243222294 | 0.116 | 0.216 | 0.020993662 | 1 |
| WBGene00019580 | oac-58 | 0.420982703 | 0.58900059 | -0.242398572 | 0.081 | 0.163 | 0.027750462 | 1 |
| WBGene00018281 | tbc-18 | 0.212020789 | 0.379822382 | -0.242086526 | 0.035 | 0.1 | 0.019133141 | 1 |
| WBGene00303368 | Y49F6B.19 | 0.147364252 | 0.315040533 | -0.241905739 | 0.023 | 0.058 | 0.102039969 | 1 |
| WBGene00020137 | T01B10.5 | 0.096150186 | 0.263820549 | -0.241897202 | 0.017 | 0.068 | 0.020098499 | 1 |
| WBGene00012158 | ucr-2.1 | 0.221572804 | 0.388799561 | -0.241257214 | 0.046 | 0.105 | 0.043633703 | 1 |
| WBGene00020511 | immt-1 | 0.098308113 | 0.265432945 | -0.241110166 | 0.017 | 0.063 | 0.03151315 | 1 |
| WBGene00044354 | ZK418.11 | 1.025875703 | 1.192781728 | -0.240794494 | 0.208 | 0.326 | 0.044806431 | 1 |
| WBGene00019433 | acdh-3 | 0.144016913 | 0.310909778 | -0.240775509 | 0.029 | 0.079 | 0.041086654 | 1 |
| WBGene00016629 | C44B7.7 | 0.39230943 | 0.558911592 | -0.240356113 | 0.075 | 0.105 | 0.401785198 | 1 |
| WBGene00011344 | T01G9.2 | 0.274446323 | 0.440816098 | -0.240020849 | 0.052 | 0.105 | 0.08198695 | 1 |
| WBGene00003030 | lin-45 | 0.38179297 | 0.548065128 | -0.239880018 | 0.069 | 0.137 | 0.055617112 | 1 |
| WBGene00012768 | eef-1B.2 | 0.718778859 | 0.885039014 | -0.239862702 | 0.139 | 0.279 | 0.007113825 | 1 |
| WBGene00012343 | W08E3.2 | 0.146038167 | 0.311859887 | -0.239230173 | 0.029 | 0.089 | 0.019633875 | 1 |
| WBGene00003090 | lys-1 | 1.075619278 | 1.241343671 | -0.239089759 | 0.22 | 0.368 | 0.015690511 | 1 |
| WBGene00003922 | pas-1 | 0.342273966 | 0.507899095 | -0.238946552 | 0.064 | 0.121 | 0.078343995 | 1 |
| WBGene00000883 | cyn-7 | 2.233044901 | 2.398658918 | -0.238930522 | 0.671 | 0.832 | 0.003944915 | 1 |
| WBGene00044764 | W01A8.7 | 0.570779285 | 0.736343386 | -0.238858507 | 0.104 | 0.2 | 0.026618903 | 1 |
| WBGene00043052 | srz-104 | 0.232455906 | 0.397972782 | -0.238790376 | 0.04 | 0.111 | 0.01602533 | 1 |
| WBGene00004947 | sos-1 | 0.189193187 | 0.354530229 | -0.238530931 | 0.035 | 0.084 | 0.056512902 | 1 |
| WBGene00005390 | srh-174 | 0.514195192 | 0.678885065 | -0.237597263 | 0.098 | 0.211 | 0.009428945 | 1 |
| WBGene00021374 | Y37E11B.1 | 0.227548909 | 0.392054987 | -0.237332102 | 0.04 | 0.095 | 0.05290333 | 1 |
| WBGene00001909 | his-35 | 2.039842815 | 2.204311853 | -0.237278665 | 0.555 | 0.758 | 0.017447976 | 1 |
| WBGene00007200 | vamp-8 | 0.700660495 | 0.864415572 | -0.236248638 | 0.133 | 0.279 | 0.00410615 | 1 |
| WBGene00021000 | W03F9.2 | 0.042512023 | 0.206078967 | -0.235977219 | 0.006 | 0.058 | 0.005959452 | 1 |
| WBGene00008956 | nekl-3 | 0.169913206 | 0.333435644 | -0.235913011 | 0.035 | 0.079 | 0.081850986 | 1 |
| WBGene00015336 | C02D5.2 | 0.245980849 | 0.409398646 | -0.235762046 | 0.052 | 0.105 | 0.078130307 | 1 |
| WBGene00022250 | Y73B6BL.29 | 0.555960135 | 0.71880814 | -0.234940009 | 0.092 | 0.253 | 0.000454214 | 1 |
| WBGene00000041 | aco-2 | 0.223324638 | 0.386103063 | -0.234839627 | 0.035 | 0.116 | 0.005874491 | 1 |
| WBGene00017011 | eaf-1 | 0.078668835 | 0.241356436 | -0.234708596 | 0.012 | 0.063 | 0.011933937 | 1 |
| WBGene00004445 | rpl-31 | 2.201648547 | 2.364324877 | -0.234692333 | 0.624 | 0.858 | 0.001001176 | 1 |
| WBGene00284844 | H38K22.9 | 0.293168412 | 0.455746319 | -0.23455034 | 0.058 | 0.126 | 0.040983095 | 1 |
| WBGene00007235 | ahsa-1 | 0.762097964 | 0.924608634 | -0.234453337 | 0.168 | 0.321 | 0.009743827 | 1 |
| WBGene00019321 | K02E10.7 | 0.209374931 | 0.371849767 | -0.234401641 | 0.035 | 0.079 | 0.084939254 | 1 |
| WBGene00014017 | ZK632.10 | 1.096512873 | 1.25896682 | -0.234371505 | 0.237 | 0.421 | 0.00446745 | 1 |
| WBGene00009448 | zfp-2 | 0.143219469 | 0.305632429 | -0.234312372 | 0.029 | 0.084 | 0.028937341 | 1 |
| WBGene00018193 | F39C12.1 | 0.305319181 | 0.467679335 | -0.234236189 | 0.046 | 0.1 | 0.066360985 | 1 |
| WBGene00001991 | hot-6 | 0.198504941 | 0.360607225 | -0.233864163 | 0.035 | 0.1 | 0.019016465 | 1 |

| | | | | | | | | |
|---|---|---|---|---|---|---|---|---|
| WBGene00021097 | cdc-37 | 0.297939726 | 0.459946228 | -0.233725978 | 0.058 | 0.126 | 0.036683365 | 1 |
| WBGene00001025 | dnj-7 | 0.11418523 | 0.276189695 | -0.233723038 | 0.017 | 0.058 | 0.048410214 | 1 |
| WBGene00016703 | rdl-1 | 0.274730799 | 0.436666321 | -0.233623575 | 0.035 | 0.084 | 0.056200069 | 1 |
| WBGene00007969 | acs-19 | 0.434785874 | 0.596706648 | -0.233602297 | 0.046 | 0.121 | 0.015368515 | 1 |
| WBGene00001563 | gei-6 | 0.185249754 | 0.347028858 | -0.233397911 | 0.029 | 0.074 | 0.06262602 | 1 |
| WBGene00010097 | srpa-68 | 0.184515775 | 0.346240524 | -0.233319494 | 0.029 | 0.084 | 0.028575811 | 1 |
| WBGene00004498 | rps-29 | 2.769354404 | 2.931061007 | -0.233293315 | 0.798 | 0.926 | 0.001516183 | 1 |
| WBGene00008990 | smgl-1 | 0.192260745 | 0.353864265 | -0.233144598 | 0.023 | 0.074 | 0.03053395 | 1 |
| WBGene00006709 | ubc-14 | 0.17975767 | 0.341156197 | -0.232848855 | 0.029 | 0.084 | 0.028937341 | 1 |
| WBGene00017792 | F25E5.16 | 0.237822751 | 0.399170091 | -0.232775007 | 0.029 | 0.084 | 0.029858437 | 1 |
| WBGene00007752 | wdfy-3 | 0.486028863 | 0.64734017 | -0.232723023 | 0.092 | 0.163 | 0.069851626 | 1 |
| WBGene00007044 | cpna-5 | 0.875260087 | 1.036564646 | -0.232713288 | 0.185 | 0.305 | 0.028349561 | 1 |
| WBGene00001673 | gpa-11 | 1.698227206 | 1.859192603 | -0.23222398 | 0.399 | 0.695 | 0.00092538 | 1 |
| WBGene00001189 | egl-21 | 2.627812053 | 2.788775581 | -0.232221285 | 0.728 | 0.953 | 0.002974866 | 1 |
| WBGene00001324 | eor-1 | 0.031556422 | 0.192465848 | -0.23214323 | 0.006 | 0.058 | 0.005959452 | 1 |
| WBGene00016406 | C34D10.1 | 0.065276926 | 0.226077485 | -0.231986169 | 0.012 | 0.053 | 0.030406126 | 1 |
| WBGene00017699 | flcn-1 | 1.360494308 | 1.521097774 | -0.231701823 | 0.26 | 0.484 | 0.000863897 | 1 |
| WBGene00009969 | F53B7.7 | 1.509210014 | 1.669695826 | -0.231532085 | 0.347 | 0.526 | 0.017040229 | 1 |
| WBGene00004479 | rps-10 | 2.786041787 | 2.946515837 | -0.231515117 | 0.815 | 0.921 | 0.001839667 | 1 |
| WBGene00003695 | nhr-105 | 1.357169856 | 1.517563372 | -0.23139893 | 0.277 | 0.458 | 0.011092633 | 1 |
| WBGene00022252 | Y73B6BL.31 | 0.035886529 | 0.196120341 | -0.231168526 | 0.006 | 0.053 | 0.009833498 | 1 |
| WBGene00018373 | F43B10.1 | 0.032643596 | 0.192703842 | -0.230918123 | 0.006 | 0.058 | 0.005900907 | 1 |
| WBGene00003132 | mat-1 | 0.027089815 | 0.186960215 | -0.230644234 | 0.006 | 0.053 | 0.009642998 | 1 |
| WBGene00008386 | cdc-5L | 0.069161231 | 0.229014606 | -0.230619672 | 0.012 | 0.058 | 0.019155473 | 1 |
| WBGene00021959 | Y57E12AL.6 | 0.672009717 | 0.831412026 | -0.229968921 | 0.156 | 0.253 | 0.096420562 | 1 |
| WBGene00044077 | pacs-1 | 0.161509441 | 0.320840985 | -0.229866828 | 0.023 | 0.095 | 0.00553692 | 1 |
| WBGene00013766 | prmt-1 | 0.38502044 | 0.544260592 | -0.229734979 | 0.069 | 0.174 | 0.006216733 | 1 |
| WBGene00008805 | git-1 | 0.061264858 | 0.220401165 | -0.229585161 | 0.012 | 0.053 | 0.030654741 | 1 |
| WBGene00001974 | hmg-4 | 0.286582109 | 0.445709138 | -0.229571775 | 0.058 | 0.116 | 0.065774224 | 1 |
| WBGene00007189 | pigm-1 | 0.100579385 | 0.259676792 | -0.229529039 | 0.017 | 0.074 | 0.013043606 | 1 |
| WBGene00022776 | perm-3 | 0.282847247 | 0.441904079 | -0.229470502 | 0.052 | 0.126 | 0.02064095 | 1 |
| WBGene00021807 | Y53G8AM.7 | 0.221494234 | 0.380492154 | -0.22938551 | 0.04 | 0.105 | 0.024669389 | 1 |
| WBGene00022410 | Y97E10C.1 | 1.335415959 | 1.494272737 | -0.229181887 | 0.329 | 0.521 | 0.024782758 | 1 |
| WBGene00000405 | cdk-1 | 0.295245852 | 0.454046122 | -0.229100362 | 0.058 | 0.1 | 0.152297154 | 1 |
| WBGene00013456 | Y67A10A.7 | 0.183969677 | 0.342681267 | -0.228972424 | 0.029 | 0.089 | 0.019381267 | 1 |
| WBGene00050941 | C33F10.14 | 0.097576585 | 0.256116064 | -0.228724121 | 0.017 | 0.074 | 0.012844649 | 1 |
| WBGene00269424 | K09D9.14 | 1.657255792 | 1.815739744 | -0.228644012 | 0.364 | 0.595 | 0.003882161 | 1 |
| WBGene00022697 | cyy-1 | 0.156006888 | 0.314235215 | -0.228275223 | 0.023 | 0.084 | 0.012740227 | 1 |
| WBGene00001089 | dre-1 | 0.191530213 | 0.349495517 | -0.227895761 | 0.029 | 0.068 | 0.090670326 | 1 |
| WBGene00001505 | fut-1 | 1.189518554 | 1.347466128 | -0.227870181 | 0.254 | 0.463 | 0.002771764 | 1 |
| WBGene00004277 | rab-18 | 0.192710641 | 0.350273325 | -0.227314904 | 0.035 | 0.095 | 0.027429253 | 1 |
| WBGene00015567 | bath-44 | 0.104977801 | 0.262340997 | -0.227027103 | 0.023 | 0.068 | 0.046422382 | 1 |
| WBGene00010476 | rnf-113 | 0.199843857 | 0.357109605 | -0.226886514 | 0.023 | 0.1 | 0.003531092 | 1 |
| WBGene00045056 | F13H10.9 | 1.068195944 | 1.22545308 | -0.226874091 | 0.202 | 0.363 | 0.006749869 | 1 |
| WBGene00022242 | sfrp-1 | 0.051246061 | 0.208318178 | -0.226607164 | 0.012 | 0.053 | 0.029670647 | 1 |
| WBGene00000204 | arx-6 | 0.168809659 | 0.325862395 | -0.226579204 | 0.029 | 0.095 | 0.01323775 | 1 |
| WBGene00004166 | pqn-85 | 0.085691641 | 0.242369402 | -0.226038228 | 0.017 | 0.063 | 0.031974939 | 1 |
| WBGene00004452 | rpl-38 | 3.35520687 | 3.511823571 | -0.225950137 | 0.867 | 0.942 | 0.002797745 | 1 |
| WBGene00012072 | T26H8.4 | 0.134545984 | 0.291116924 | -0.225884119 | 0.023 | 0.084 | 0.013295554 | 1 |
| WBGene00012326 | W07E11.1 | 0.313402093 | 0.469916662 | -0.225802792 | 0.052 | 0.095 | 0.144115149 | 1 |
| WBGene00018521 | F46H5.5 | 0.11608422 | 0.27224463 | -0.22529185 | 0.023 | 0.079 | 0.020774836 | 1 |
| WBGene00009661 | patr-1 | 0.418289092 | 0.57427667 | -0.225042504 | 0.064 | 0.158 | 0.008756438 | 1 |
| WBGene00015179 | B0416.3 | 0.059082241 | 0.214940584 | -0.224856058 | 0.012 | 0.053 | 0.030406126 | 1 |

| | | | | | | | | |
|---|---|---|---|---|---|---|---|---|
| WBGene00012329 | sre-44 | 0.244892186 | 0.400702932 | -0.224787391 | 0.035 | 0.1 | 0.019016465 | 1 |
| WBGene00022677 | rfth-1 | 0.178200564 | 0.333917677 | -0.224652308 | 0.023 | 0.095 | 0.00553692 | 1 |
| WBGene00013238 | trap-4 | 0.384727992 | 0.540297112 | -0.224438797 | 0.064 | 0.147 | 0.014848247 | 1 |
| WBGene00012641 | epg-6 | 0.395927255 | 0.551489457 | -0.224428818 | 0.064 | 0.147 | 0.016872206 | 1 |
| WBGene00015746 | C13F10.7 | 0.054303118 | 0.209839171 | -0.224391092 | 0.012 | 0.053 | 0.030159245 | 1 |
| WBGene00020932 | dhod-1 | 0.166316908 | 0.321716813 | -0.224194673 | 0.029 | 0.068 | 0.089659311 | 1 |
| WBGene00019872 | R04E5.9 | 0.300109724 | 0.455404159 | -0.224042511 | 0.052 | 0.121 | 0.029103055 | 1 |
| WBGene00015745 | C13F10.6 | 0.02790056 | 0.182970372 | -0.223718449 | 0.006 | 0.053 | 0.009833498 | 1 |
| WBGene00004418 | rpl-7 | 2.575395097 | 2.730382044 | -0.2235989 | 0.746 | 0.921 | 0.009256914 | 1 |
| WBGene00000184 | arf-6 | 0.173052349 | 0.328020044 | -0.223571124 | 0.035 | 0.084 | 0.054963118 | 1 |
| WBGene00009126 | pyk-1 | 0.780339478 | 0.93507689 | -0.223238897 | 0.15 | 0.279 | 0.016392847 | 1 |
| WBGene00013803 | snx-13 | 0.286898271 | 0.441597937 | -0.223184441 | 0.052 | 0.1 | 0.107028899 | 1 |
| WBGene00013277 | sre-41 | 0.212311494 | 0.366804542 | -0.222886354 | 0.035 | 0.095 | 0.027592701 | 1 |
| WBGene00002981 | lgg-2 | 1.743537165 | 1.897943797 | -0.222761682 | 0.439 | 0.668 | 0.010446269 | 1 |
| WBGene00010697 | uda-1 | 0.131566053 | 0.285966373 | -0.222752576 | 0.017 | 0.063 | 0.03151315 | 1 |
| WBGene00000392 | cdd-2 | 0.412779186 | 0.566910751 | -0.222364844 | 0.069 | 0.142 | 0.039515997 | 1 |
| WBGene00003407 | mrp-1 | 0.2827532 | 0.43687995 | -0.222357898 | 0.052 | 0.121 | 0.031113854 | 1 |
| WBGene00005265 | srh-42 | 0.692620652 | 0.846507104 | -0.222011221 | 0.121 | 0.189 | 0.130747456 | 1 |
| WBGene00004468 | rpn-12 | 0.325449937 | 0.479142634 | -0.221731691 | 0.052 | 0.142 | 0.007379977 | 1 |
| WBGene00018328 | F42A6.6 | 0.091449496 | 0.244987865 | -0.221509045 | 0.012 | 0.063 | 0.011933937 | 1 |
| WBGene00004349 | rgs-6 | 0.293761225 | 0.447136489 | -0.221273733 | 0.04 | 0.089 | 0.072103787 | 1 |
| WBGene00003638 | nhr-48 | 0.070651553 | 0.223939758 | -0.221148133 | 0.012 | 0.053 | 0.030654741 | 1 |
| WBGene00019180 | H10D18.5 | 0.27634052 | 0.429620051 | -0.22113562 | 0.035 | 0.063 | 0.231427344 | 1 |
| WBGene00219675 | linc-44 | 0.063122844 | 0.21602136 | -0.220585931 | 0.012 | 0.063 | 0.012035905 | 1 |
| WBGene00021312 | szy-2 | 0.538952959 | 0.691658773 | -0.220307919 | 0.098 | 0.226 | 0.004686637 | 1 |
| WBGene00004681 | rsd-2 | 0.070935666 | 0.223411924 | -0.219976742 | 0.012 | 0.063 | 0.011933937 | 1 |
| WBGene00017984 | F32D1.5 | 0.24163813 | 0.394010543 | -0.219826924 | 0.04 | 0.121 | 0.008252264 | 1 |
| WBGene00004701 | rsp-4 | 0.354327067 | 0.506411134 | -0.219410929 | 0.075 | 0.126 | 0.142175669 | 1 |
| WBGene00013695 | epg-8 | 0.112573049 | 0.264652973 | -0.219404951 | 0.023 | 0.068 | 0.045528421 | 1 |
| WBGene00000884 | cyn-8 | 0.280150758 | 0.432024273 | -0.219107167 | 0.058 | 0.121 | 0.052875484 | 1 |
| WBGene00019305 | snap-29 | 0.354593296 | 0.505880873 | -0.218261837 | 0.052 | 0.147 | 0.005094404 | 1 |
| WBGene00014096 | ZK829.7 | 0.588623517 | 0.739750482 | -0.218030122 | 0.116 | 0.2 | 0.071097598 | 1 |
| WBGene00011944 | T23D8.3 | 0.134277712 | 0.285250935 | -0.217808321 | 0.023 | 0.063 | 0.069983892 | 1 |
| WBGene00014202 | mmcm-1 | 0.306333593 | 0.456892132 | -0.217210057 | 0.052 | 0.105 | 0.083496137 | 1 |
| WBGene00043061 | Y54G2A.38 | 0.134200128 | 0.284679363 | -0.217095645 | 0.017 | 0.053 | 0.075425698 | 1 |
| WBGene00016025 | xbx-4 | 1.071840582 | 1.222110783 | -0.216794073 | 0.208 | 0.389 | 0.005176018 | 1 |
| WBGene00005337 | srh-118 | 0.85665406 | 1.006867542 | -0.216712245 | 0.133 | 0.3 | 0.001091059 | 1 |
| WBGene00016964 | C56C10.7 | 0.255037903 | 0.405177772 | -0.216606045 | 0.046 | 0.095 | 0.094465634 | 1 |
| WBGene00017289 | F09E5.11 | 0.148252564 | 0.298222845 | -0.216361381 | 0.035 | 0.079 | 0.080555626 | 1 |
| WBGene00008256 | glb-12 | 1.264324839 | 1.41414969 | -0.216151569 | 0.254 | 0.5 | 0.000456923 | 1 |
| WBGene00009738 | hecw-1 | 0.306765849 | 0.456536498 | -0.216073373 | 0.058 | 0.095 | 0.20575498 | 1 |
| WBGene00022114 | Y71F9AL.9 | 0.592328436 | 0.742030539 | -0.215974482 | 0.092 | 0.216 | 0.003917521 | 1 |
| WBGene00009558 | F39B2.8 | 0.215566046 | 0.365212194 | -0.215893756 | 0.029 | 0.084 | 0.028396509 | 1 |
| WBGene00007574 | C14B1.3 | 0.892113717 | 1.041649475 | -0.215734496 | 0.168 | 0.353 | 0.001360897 | 1 |
| WBGene00002099 | ins-16 | 0.158190084 | 0.307515753 | -0.215431402 | 0.035 | 0.053 | 0.416573822 | 1 |
| WBGene00012553 | cox-5A | 1.01844436 | 1.167740752 | -0.215389164 | 0.208 | 0.426 | 0.000965633 | 1 |
| WBGene00007104 | B0024.15 | 0.357496086 | 0.506715639 | -0.215278309 | 0.058 | 0.137 | 0.020130906 | 1 |
| WBGene00003129 | map-1 | 0.31248669 | 0.461622697 | -0.215157779 | 0.064 | 0.1 | 0.249987985 | 1 |
| WBGene00043308 | tmem-107 | 0.342234526 | 0.491144672 | -0.214831929 | 0.058 | 0.1 | 0.17890698 | 1 |
| WBGene00005380 | srh-164 | 1.283874261 | 1.432751149 | -0.214783948 | 0.295 | 0.505 | 0.010536684 | 1 |
| WBGene00002180 | jtr-1 | 0.065334246 | 0.213857098 | -0.214273182 | 0.012 | 0.053 | 0.0309051 | 1 |
| WBGene00007048 | nfx-1 | 0.198439043 | 0.346947259 | -0.214252067 | 0.017 | 0.074 | 0.013043606 | 1 |
| WBGene00016120 | C25H3.11 | 0.243274071 | 0.391747574 | -0.214201987 | 0.04 | 0.105 | 0.02370045 | 1 |

| | | | | | | | | |
|---|---|---|---|---|---|---|---|---|
| WBGene00019355 | K03B4.4 | 0.323538568 | 0.471945262 | -0.214105602 | 0.012 | 0.058 | 0.01947719 | 1 |
| WBGene00001262 | emb-8 | 1.43232795 | 1.580658574 | -0.213995857 | 0.347 | 0.495 | 0.054303369 | 1 |
| WBGene00012674 | bed-1 | 0.225046205 | 0.373281803 | -0.213858763 | 0.04 | 0.111 | 0.018133668 | 1 |
| WBGene00020629 | T20F5.7 | 0.191183127 | 0.339375875 | -0.213796942 | 0.035 | 0.084 | 0.055270208 | 1 |
| WBGene00016060 | mms-19 | 0.191221397 | 0.339402547 | -0.213780211 | 0.035 | 0.068 | 0.158060991 | 1 |
| WBGene00000430 | ceh-5 | 0.263653824 | 0.411532492 | -0.213343821 | 0.046 | 0.132 | 0.008468588 | 1 |
| WBGene00012290 | W05H12.2 | 0.366446202 | 0.514316398 | -0.213331598 | 0.069 | 0.132 | 0.073593506 | 1 |
| WBGene00007106 | B0035.3 | 0.046254655 | 0.193988299 | -0.213134595 | 0.006 | 0.053 | 0.009929999 | 1 |
| WBGene00018111 | F36H9.2 | 0.242888476 | 0.390532886 | -0.213005859 | 0.035 | 0.079 | 0.084492388 | 1 |
| WBGene00023281 | F39E9.8 | 1.451162813 | 1.598797949 | -0.212992479 | 0.358 | 0.511 | 0.143128642 | 1 |
| WBGene00017513 | F16F9.1 | 0.503241061 | 0.650770351 | -0.212839774 | 0.092 | 0.205 | 0.008269686 | 1 |
| WBGene00010199 | bet-2 | 0.33813195 | 0.485391354 | -0.212450411 | 0.052 | 0.116 | 0.039701856 | 1 |
| WBGene00019947 | htz-1 | 0.348418285 | 0.495517159 | -0.212218816 | 0.069 | 0.142 | 0.042652239 | 1 |
| WBGene00003036 | lin-53 | 0.205754628 | 0.352772953 | -0.212102608 | 0.029 | 0.1 | 0.008401416 | 1 |
| WBGene00013530 | ntl-11 | 0.061683046 | 0.208661954 | -0.212045741 | 0.006 | 0.053 | 0.009833498 | 1 |
| WBGene00010986 | sprr-1 | 0.698598306 | 0.845490688 | -0.21192091 | 0.11 | 0.237 | 0.005569625 | 1 |
| WBGene00271636 | W05G11.8 | 0.711627085 | 0.857985742 | -0.211150908 | 0.156 | 0.274 | 0.034080389 | 1 |
| WBGene00022812 | uvs-1 | 0.163093682 | 0.309226053 | -0.210824447 | 0.029 | 0.068 | 0.091690345 | 1 |
| WBGene00043983 | C39D10.11 | 0.723385182 | 0.869406653 | -0.210664452 | 0.104 | 0.253 | 0.000962102 | 1 |
| WBGene00020942 | W02D7.6 | 0.65697219 | 0.802873808 | -0.210491541 | 0.133 | 0.253 | 0.016521293 | 1 |
| WBGene00017236 | nlp-60 | 2.929827891 | 3.075680912 | -0.210421429 | 0.746 | 0.874 | 0.019250649 | 1 |
| WBGene00015186 | misc-1 | 0.091100193 | 0.236655798 | -0.209992351 | 0.017 | 0.053 | 0.074916061 | 1 |
| WBGene00012089 | T27E7.4 | 0.511422054 | 0.656797413 | -0.20973231 | 0.075 | 0.168 | 0.013669378 | 1 |
| WBGene00016428 | dmsr-7 | 0.098409177 | 0.243730843 | -0.209654847 | 0.017 | 0.058 | 0.049097285 | 1 |
| WBGene00008454 | E02A10.4 | 3.98815342 | 4.133465898 | -0.20964159 | 0.96 | 0.974 | 0.001404317 | 1 |
| WBGene00018509 | xrep-4 | 0.355570316 | 0.500846388 | -0.209589069 | 0.064 | 0.153 | 0.013629402 | 1 |
| WBGene00014036 | ZK643.5 | 0.484515763 | 0.629772901 | -0.209561753 | 0.087 | 0.189 | 0.012060805 | 1 |
| WBGene00022175 | Y71H2AM.10 | 0.397770706 | 0.542961119 | -0.209465489 | 0.069 | 0.126 | 0.089093012 | 1 |
| WBGene00015501 | C06A5.3 | 0.349972255 | 0.4951079 | -0.209386475 | 0.046 | 0.132 | 0.007318442 | 1 |
| WBGene00011940 | T23B5.3 | 0.136397483 | 0.281529893 | -0.209381807 | 0.017 | 0.058 | 0.049443878 | 1 |
| WBGene00010427 | hpo-11 | 0.520359091 | 0.665362676 | -0.209195953 | 0.052 | 0.126 | 0.022587034 | 1 |
| WBGene00002845 | let-711 | 0.094258189 | 0.239224949 | -0.209142827 | 0.017 | 0.063 | 0.031974939 | 1 |
| WBGene00015972 | C18E3.3 | 1.748667502 | 1.893589525 | -0.209078284 | 0.41 | 0.621 | 0.018299758 | 1 |
| WBGene00018330 | elks-1 | 0.155204847 | 0.299880844 | -0.208723343 | 0.029 | 0.074 | 0.065610102 | 1 |
| WBGene00004274 | rab-11.1 | 1.097548713 | 1.242171048 | -0.208645926 | 0.249 | 0.453 | 0.006025015 | 1 |
| WBGene00000377 | cct-1 | 0.364457319 | 0.509049617 | -0.208602591 | 0.069 | 0.163 | 0.012794604 | 1 |
| WBGene00016407 | unk-1 | 0.400456618 | 0.544191887 | -0.207366159 | 0.075 | 0.132 | 0.107685298 | 1 |
| WBGene00195086 | F21C10.13 | 1.140985261 | 1.284481743 | -0.207021663 | 0.272 | 0.442 | 0.021794246 | 1 |
| WBGene00219936 | F37A8.7 | 0.134071735 | 0.277422445 | -0.206811359 | 0.023 | 0.079 | 0.019658076 | 1 |
| WBGene00000492 | che-13 | 0.897895844 | 1.040900249 | -0.206311746 | 0.197 | 0.332 | 0.038540292 | 1 |
| WBGene00022268 | Y73E7A.1 | 0.129006219 | 0.271765125 | -0.205957566 | 0.023 | 0.074 | 0.03094576 | 1 |
| WBGene00012989 | Y48C3A.5 | 0.597356845 | 0.740034217 | -0.205839937 | 0.098 | 0.184 | 0.034021743 | 1 |
| WBGene00001582 | gfi-2 | 0.451480694 | 0.594077847 | -0.205724206 | 0.064 | 0.179 | 0.002247622 | 1 |
| WBGene00021960 | tmem-258 | 0.582340533 | 0.724772766 | -0.205486276 | 0.104 | 0.211 | 0.016825058 | 1 |
| WBGene00010428 | dcn-1 | 0.061978586 | 0.204406006 | -0.205479332 | 0.012 | 0.053 | 0.030654741 | 1 |
| WBGene00020901 | T28F2.2 | 0.258970619 | 0.401192006 | -0.205182089 | 0.046 | 0.058 | 0.671372158 | 1 |
| WBGene00013638 | Y105C5A.14 | 0.379547969 | 0.521684103 | -0.205059096 | 0.069 | 0.158 | 0.015340449 | 1 |
| WBGene00010221 | F57G12.1 | 0.266276132 | 0.408405631 | -0.205049523 | 0.029 | 0.068 | 0.087664937 | 1 |
| WBGene00007635 | npr-4 | 0.798306979 | 0.940148537 | -0.204634112 | 0.121 | 0.205 | 0.05136339 | 1 |
| WBGene00001520 | gas-1 | 0.035080609 | 0.176846042 | -0.204524288 | 0.006 | 0.053 | 0.009833498 | 1 |
| WBGene00010262 | droe-4 | 1.414622488 | 1.556089499 | -0.204093756 | 0.092 | 0.126 | 0.340383561 | 1 |
| WBGene00000771 | cpb-2 | 0.389390031 | 0.530759806 | -0.203953473 | 0.069 | 0.105 | 0.244485603 | 1 |
| WBGene00003883 | osm-1 | 0.827872106 | 0.968914455 | -0.203481098 | 0.15 | 0.268 | 0.018935253 | 1 |

| | | | | | | | | |
|---|---|---|---|---|---|---|---|---|
| WBGene00000266 | bre-1 | 0.833190937 | 0.974226062 | -0.203470675 | 0.191 | 0.305 | 0.046428616 | 1 |
| WBGene00008688 | rbm-34 | 0.051712818 | 0.192665081 | -0.203351132 | 0.012 | 0.058 | 0.018681607 | 1 |
| WBGene00008504 | pigq-1 | 0.092842403 | 0.233720229 | -0.20324374 | 0.017 | 0.053 | 0.075938161 | 1 |
| WBGene00012004 | dyrb-1 | 0.222452254 | 0.363272029 | -0.203159992 | 0.035 | 0.105 | 0.012300431 | 1 |
| WBGene00016827 | sre-53 | 0.406860846 | 0.547635754 | -0.203095262 | 0.052 | 0.137 | 0.008857283 | 1 |
| WBGene00045249 | T07A9.15 | 0.134856474 | 0.275610694 | -0.203065415 | 0.023 | 0.074 | 0.031153447 | 1 |
| WBGene00020496 | spat-3 | 0.330381731 | 0.471000476 | -0.202869965 | 0.035 | 0.116 | 0.005316929 | 1 |
| WBGene00000802 | crt-1 | 1.461447661 | 1.601851329 | -0.202559676 | 0.301 | 0.568 | 0.000309911 | 1 |
| WBGene00002242 | kvs-1 | 0.887612616 | 1.027475078 | -0.201778881 | 0.179 | 0.326 | 0.01573014 | 1 |
| WBGene00019116 | nhr-143 | 0.881001794 | 1.020747845 | -0.201610935 | 0.179 | 0.337 | 0.010586687 | 1 |
| WBGene00007547 | nhr-154 | 0.07641296 | 0.216149137 | -0.20159669 | 0.012 | 0.063 | 0.011933937 | 1 |
| WBGene00006920 | vha-11 | 0.372312692 | 0.511787177 | -0.201219148 | 0.069 | 0.142 | 0.038983373 | 1 |
| WBGene00012651 | orc-4 | 0.306572884 | 0.445703853 | -0.200723559 | 0.058 | 0.111 | 0.087125446 | 1 |
| WBGene00077698 | lurp-4 | 2.441267168 | 2.5803687 | -0.20068109 | 0.682 | 0.879 | 0.018117188 | 1 |
| WBGene00010813 | pitp-1 | 0.438993703 | 0.578020316 | -0.200573006 | 0.075 | 0.147 | 0.044139676 | 1 |
| WBGene00000112 | alh-6 | 0.245499591 | 0.384512735 | -0.200553573 | 0.052 | 0.111 | 0.057548656 | 1 |
| WBGene00009082 | dlat-1 | 0.051384284 | 0.190205727 | -0.200277007 | 0.012 | 0.053 | 0.030406126 | 1 |
| WBGene00020507 | vha-15 | 0.104492416 | 0.243302813 | -0.200261072 | 0.012 | 0.068 | 0.007350311 | 1 |
| WBGene00020324 | T07F12.4 | 0.12134359 | 0.259602442 | -0.199465359 | 0.023 | 0.068 | 0.046422382 | 1 |
| WBGene00004348 | rgs-5 | 0.171766305 | 0.310003117 | -0.199433563 | 0.029 | 0.084 | 0.028756087 | 1 |
| WBGene00020801 | T25D10.4 | 0.720606395 | 0.858738186 | -0.199282051 | 0.127 | 0.221 | 0.0524052 | 1 |
| WBGene00001779 | gst-31 | 0.278619273 | 0.416572386 | -0.199024272 | 0.04 | 0.116 | 0.011771915 | 1 |
| WBGene00219421 | Y66D12A.30 | 0.10292203 | 0.240748071 | -0.198840947 | 0.017 | 0.053 | 0.073905225 | 1 |
| WBGene00022527 | ZC132.9 | 2.283020994 | 2.420572604 | -0.198445026 | 0.572 | 0.753 | 0.005662428 | 1 |
| WBGene00020069 | R13H7.2 | 0.542173999 | 0.67968462 | -0.19838589 | 0.087 | 0.142 | 0.132873411 | 1 |
| WBGene00006577 | tlf-1 | 0.131046686 | 0.268510188 | -0.198317913 | 0.023 | 0.068 | 0.047026425 | 1 |
| WBGene00006658 | twk-3 | 0.097395481 | 0.234614224 | -0.197964799 | 0.017 | 0.058 | 0.048410214 | 1 |
| WBGene00012532 | crp-1 | 0.032442046 | 0.169626847 | -0.197915832 | 0.006 | 0.053 | 0.009929999 | 1 |
| WBGene00016289 | lntl-1 | 0.395537154 | 0.532689781 | -0.197869415 | 0.04 | 0.084 | 0.098547972 | 1 |
| WBGene00023488 | srz-84 | 0.779536949 | 0.916672092 | -0.197844191 | 0.173 | 0.289 | 0.04882875 | 1 |
| WBGene00014701 | C43D7.1 | 0.684239821 | 0.821183384 | -0.1975678 | 0.139 | 0.232 | 0.057593517 | 1 |
| WBGene00010913 | M110.3 | 0.362702516 | 0.499468124 | -0.197311064 | 0.075 | 0.1 | 0.477882065 | 1 |
| WBGene00012289 | W05H12.1 | 1.268040012 | 1.404425094 | -0.19676208 | 0.266 | 0.468 | 0.003420626 | 1 |
| WBGene00000367 | cca-1 | 0.240632472 | 0.376972171 | -0.196696608 | 0.04 | 0.095 | 0.05152085 | 1 |
| WBGene00011144 | R08D7.4 | 0.6831201 | 0.819328202 | -0.196506754 | 0.145 | 0.268 | 0.022325333 | 1 |
| WBGene00018042 | mks-3 | 0.584989325 | 0.721140802 | -0.19642506 | 0.081 | 0.163 | 0.032960188 | 1 |
| WBGene00003053 | lmp-1 | 0.468880672 | 0.604880766 | -0.19620666 | 0.069 | 0.147 | 0.029916798 | 1 |
| WBGene00013281 | sre-27 | 0.524887397 | 0.660873183 | -0.196186018 | 0.075 | 0.179 | 0.007272223 | 1 |
| WBGene00018285 | smk-1 | 1.038736073 | 1.174457892 | -0.195805195 | 0.185 | 0.337 | 0.008841022 | 1 |
| WBGene00007025 | mdt-29 | 0.10534456 | 0.240946548 | -0.195632316 | 0.023 | 0.058 | 0.105744875 | 1 |
| WBGene00018565 | F47E1.1 | 0.112316827 | 0.24789583 | -0.195599156 | 0.017 | 0.068 | 0.020864658 | 1 |
| WBGene00000566 | cnt-2 | 0.13004958 | 0.265594518 | -0.19555001 | 0.023 | 0.068 | 0.046122767 | 1 |
| WBGene00020397 | pcbd-1 | 0.67872233 | 0.814248896 | -0.195523505 | 0.133 | 0.263 | 0.012526439 | 1 |
| WBGene00020399 | ztf-4 | 0.167648333 | 0.303041771 | -0.195331442 | 0.035 | 0.084 | 0.056512902 | 1 |
| WBGene00018018 | F33G12.6 | 0.169886627 | 0.304874861 | -0.194746856 | 0.029 | 0.063 | 0.141454385 | 1 |
| WBGene00019741 | nhr-201 | 0.10667529 | 0.241512189 | -0.194528526 | 0.023 | 0.068 | 0.047026528 | 1 |
| WBGene00004014 | phb-1 | 0.232555054 | 0.367357817 | -0.194479277 | 0.035 | 0.1 | 0.018784979 | 1 |
| WBGene00012203 | rga-1 | 0.322630635 | 0.457401507 | -0.194433268 | 0.058 | 0.126 | 0.039252256 | 1 |
| WBGene00004408 | rla-0 | 1.761356755 | 1.895930589 | -0.194149003 | 0.445 | 0.632 | 0.026317039 | 1 |
| WBGene00002202 | kin-19 | 0.810053996 | 0.944505917 | -0.19397312 | 0.133 | 0.284 | 0.002765216 | 1 |
| WBGene00022400 | rpb-9 | 0.209097038 | 0.34319932 | -0.193468697 | 0.04 | 0.068 | 0.273337111 | 1 |
| WBGene00000157 | aps-2 | 0.276317188 | 0.410345074 | -0.193361366 | 0.052 | 0.105 | 0.076523423 | 1 |
| WBGene00009337 | uig-1 | 0.446002414 | 0.579920838 | -0.193203446 | 0.087 | 0.153 | 0.07892498 | 1 |

| | | | | | | | | |
|---|---|---|---|---|---|---|---|---|
| WBGene00009242 | sre-6 | 1.450560374 | 1.584420644 | -0.193119548 | 0.295 | 0.505 | 0.002413683 | 1 |
| WBGene00020709 | T23B3.1 | 0.145165574 | 0.278895576 | -0.192931611 | 0.012 | 0.079 | 0.002814546 | 1 |
| WBGene00009188 | lsy-22 | 0.13261476 | 0.266057284 | -0.192516867 | 0.023 | 0.053 | 0.150885864 | 1 |
| WBGene00018654 | F49H12.3 | 0.814676618 | 0.948106706 | -0.192498927 | 0.145 | 0.232 | 0.082505431 | 1 |
| WBGene00021799 | Y53G8AL.1 | 0.357274637 | 0.490680593 | -0.192464111 | 0.04 | 0.142 | 0.001642393 | 1 |
| WBGene00021682 | Y48G1C.9 | 0.089458519 | 0.222697782 | -0.192223624 | 0.017 | 0.058 | 0.048069718 | 1 |
| WBGene00017158 | hpo-10 | 0.290246678 | 0.42347601 | -0.192209296 | 0.058 | 0.111 | 0.092601033 | 1 |
| WBGene00016313 | set-4 | 0.285137892 | 0.418359963 | -0.192198822 | 0.052 | 0.126 | 0.022706319 | 1 |
| WBGene00015748 | C14A11.2 | 0.851576871 | 0.98455089 | -0.191840957 | 0.173 | 0.295 | 0.026962165 | 1 |
| WBGene00006789 | unc-54 | 4.63189874 | 4.764865227 | -0.191830093 | 0.977 | 0.995 | 0.006799869 | 1 |
| WBGene00011512 | T05H10.8 | 2.003382621 | 2.136115818 | -0.191493525 | 0.491 | 0.763 | 0.003359834 | 1 |
| WBGene00003996 | pgp-2 | 0.104299206 | 0.236926476 | -0.191340705 | 0.012 | 0.058 | 0.019155473 | 1 |
| WBGene00000518 | ckk-1 | 1.069737218 | 1.202347349 | -0.191315978 | 0.231 | 0.332 | 0.123562016 | 1 |
| WBGene00003926 | pas-5 | 0.604549463 | 0.73706496 | -0.19117945 | 0.133 | 0.242 | 0.031705401 | 1 |
| WBGene00018023 | set-11 | 0.07394111 | 0.206432357 | -0.191144466 | 0.012 | 0.053 | 0.0309051 | 1 |
| WBGene00017663 | F21D12.3 | 1.466547458 | 1.599014439 | -0.191109456 | 0.324 | 0.574 | 0.000974194 | 1 |
| WBGene00011273 | R53.4 | 1.383261678 | 1.515672183 | -0.191027978 | 0.347 | 0.579 | 0.005496462 | 1 |
| WBGene00017217 | F07F6.4 | 0.342802932 | 0.47509208 | -0.190852897 | 0.046 | 0.053 | 0.803420961 | 1 |
| WBGene00011912 | T22C1.1 | 0.154275677 | 0.286235551 | -0.190377857 | 0.023 | 0.079 | 0.020918247 | 1 |
| WBGene00019166 | tat-2 | 0.221076743 | 0.352987917 | -0.190307596 | 0.029 | 0.074 | 0.060462371 | 1 |
| WBGene00014022 | ZK637.2 | 0.119161326 | 0.250976774 | -0.190169493 | 0.023 | 0.074 | 0.030739263 | 1 |
| WBGene00001449 | flp-6 | 0.562642488 | 0.69412905 | -0.18969501 | 0.098 | 0.095 | 0.877995091 | 1 |
| WBGene00004738 | scc-3 | 0.369957316 | 0.501240863 | -0.189402121 | 0.052 | 0.111 | 0.054581752 | 1 |
| WBGene00008041 | ufc-1 | 0.304659948 | 0.435873957 | -0.1893018 | 0.058 | 0.121 | 0.05641433 | 1 |
| WBGene00000099 | air-2 | 0.527614439 | 0.658740766 | -0.189175302 | 0.081 | 0.163 | 0.029794993 | 1 |
| WBGene00016316 | C32D5.8 | 1.429029618 | 1.55967663 | -0.188483795 | 0.37 | 0.484 | 0.130145291 | 1 |
| WBGene00016624 | hrpl-1 | 0.117070146 | 0.24725719 | -0.187820204 | 0.017 | 0.068 | 0.020401965 | 1 |
| WBGene00002140 | inx-18 | 0.750652195 | 0.880519127 | -0.187358378 | 0.121 | 0.284 | 0.000951813 | 1 |
| WBGene00005165 | srg-7 | 0.212947332 | 0.342795021 | -0.187330616 | 0.035 | 0.074 | 0.11782539 | 1 |
| WBGene00001337 | ears-1 | 0.096214616 | 0.225909333 | -0.187109924 | 0.017 | 0.053 | 0.075425698 | 1 |
| WBGene00015250 | B0546.3 | 0.142268626 | 0.27186041 | -0.186961424 | 0.012 | 0.053 | 0.031157212 | 1 |
| WBGene00018391 | rpb-4 | 0.189252771 | 0.318740847 | -0.186811806 | 0.035 | 0.089 | 0.03952377 | 1 |
| WBGene00007718 | otub-1 | 0.1638301 | 0.292856177 | -0.186145281 | 0.029 | 0.084 | 0.030045651 | 1 |
| WBGene00009141 | ncbp-2 | 1.19043969 | 1.319308244 | -0.185918023 | 0.249 | 0.447 | 0.002519547 | 1 |
| WBGene00012889 | icap-1 | 0.074236316 | 0.20304319 | -0.185829038 | 0.012 | 0.053 | 0.031157212 | 1 |
| WBGene00021547 | Y44E3A.1 | 0.105789468 | 0.234380379 | -0.18551747 | 0.023 | 0.063 | 0.068261235 | 1 |
| WBGene00001155 | ech-6 | 0.104885674 | 0.233439164 | -0.185463482 | 0.017 | 0.058 | 0.050143224 | 1 |
| WBGene00017597 | F19C7.8 | 1.284507715 | 1.413008444 | -0.185387365 | 0.312 | 0.447 | 0.059553871 | 1 |
| WBGene00013375 | srsx-25 | 0.149093676 | 0.27755809 | -0.185334973 | 0.029 | 0.079 | 0.045270682 | 1 |
| WBGene00010756 | K10G9.2 | 0.655941627 | 0.78411892 | -0.184920745 | 0.127 | 0.226 | 0.042178558 | 1 |
| WBGene00004501 | rpt-1 | 0.231285952 | 0.359407803 | -0.184840759 | 0.046 | 0.089 | 0.119197948 | 1 |
| WBGene00005416 | srh-206 | 0.352334054 | 0.480260525 | -0.184558885 | 0.069 | 0.095 | 0.443337647 | 1 |
| WBGene00020679 | ogdh-1 | 0.328748429 | 0.456539446 | -0.184363467 | 0.058 | 0.142 | 0.01402123 | 1 |
| WBGene00008586 | snrp-40.1 | 0.436249111 | 0.563960693 | -0.184248867 | 0.075 | 0.137 | 0.084311027 | 1 |
| WBGene00021503 | ctsa-4.2 | 1.518662942 | 1.646353594 | -0.18421867 | 0.301 | 0.532 | 0.000744936 | 1 |
| WBGene00044084 | sre-46 | 0.428118333 | 0.555620704 | -0.183947038 | 0.069 | 0.137 | 0.053251978 | 1 |
| WBGene00000102 | akt-1 | 0.995423053 | 1.12281499 | -0.183787717 | 0.173 | 0.353 | 0.001013193 | 1 |
| WBGene00015418 | pac-1 | 0.414518243 | 0.541785963 | -0.183608508 | 0.081 | 0.158 | 0.044008538 | 1 |
| WBGene00020446 | T12B3.3 | 0.111235427 | 0.238496558 | -0.183599002 | 0.017 | 0.068 | 0.020555192 | 1 |
| WBGene00000232 | avr-14 | 0.336977151 | 0.464032509 | -0.183302136 | 0.058 | 0.1 | 0.166174502 | 1 |
| WBGene00004494 | rps-25 | 3.292071919 | 3.418935243 | -0.183025089 | 0.902 | 0.947 | 0.011661225 | 1 |
| WBGene00015551 | C06G3.5 | 0.415681055 | 0.541972137 | -0.182199517 | 0.064 | 0.158 | 0.008593942 | 1 |
| WBGene00017494 | irld-31 | 0.409204557 | 0.535446859 | -0.182129144 | 0.058 | 0.1 | 0.186602631 | 1 |

| | | | | | | | | |
|---|---|---|---|---|---|---|---|---|
| WBGene00008422 | D2045.2 | 0.157024874 | 0.283115149 | -0.181909814 | 0.023 | 0.079 | 0.020490582 | 1 |
| WBGene00194910 | ZC404.15 | 1.040959653 | 1.167049063 | -0.181908567 | 0.214 | 0.374 | 0.013702226 | 1 |
| WBGene00004448 | rpl-34 | 2.239934245 | 2.365878216 | -0.181698743 | 0.613 | 0.8 | 0.020034698 | 1 |
| WBGene00001229 | eif-3.F | 0.078711245 | 0.204462949 | -0.18142136 | 0.012 | 0.053 | 0.0309051 | 1 |
| WBGene00018606 | F48E3.6 | 1.04446214 | 1.170189699 | -0.181386526 | 0.231 | 0.4 | 0.025220348 | 1 |
| WBGene00007892 | C33B4.4 | 2.930165801 | 3.055825805 | -0.181289065 | 0.769 | 0.9 | 0.117779957 | 1 |
| WBGene00017925 | F29B9.11 | 0.885855439 | 1.011345661 | -0.18104412 | 0.191 | 0.342 | 0.015235848 | 1 |
| WBGene00017085 | E01A2.2 | 0.268943729 | 0.394379813 | -0.180966016 | 0.052 | 0.095 | 0.148394166 | 1 |
| WBGene00013598 | vps-28 | 0.153492526 | 0.27892229 | -0.180956899 | 0.029 | 0.068 | 0.094280766 | 1 |
| WBGene00009812 | suca-1 | 0.31109995 | 0.436487767 | -0.180896382 | 0.058 | 0.116 | 0.071500606 | 1 |
| WBGene00000985 | dhs-22 | 0.195183941 | 0.320474379 | -0.180755894 | 0.029 | 0.074 | 0.064103725 | 1 |
| WBGene00021554 | Y45G5AL.1 | 0.18605863 | 0.311274673 | -0.180648565 | 0.035 | 0.084 | 0.058098888 | 1 |
| WBGene00006802 | unc-69 | 0.89735424 | 1.022525702 | -0.180584247 | 0.162 | 0.358 | 0.000700656 | 1 |
| WBGene00001494 | frm-8 | 0.119712168 | 0.244818823 | -0.180490752 | 0.023 | 0.058 | 0.10449823 | 1 |
| WBGene00021849 | Y54F10AM.5 | 0.252497699 | 0.377384303 | -0.180173285 | 0.04 | 0.095 | 0.052624381 | 1 |
| WBGene00004152 | pqn-70 | 0.383213948 | 0.508043126 | -0.180090436 | 0.069 | 0.153 | 0.024528182 | 1 |
| WBGene00004895 | smu-1 | 0.090911229 | 0.21546656 | -0.17969536 | 0.012 | 0.063 | 0.012138644 | 1 |
| WBGene00022019 | sut-2 | 0.31362025 | 0.438099614 | -0.179585761 | 0.046 | 0.111 | 0.032118211 | 1 |
| WBGene00003160 | mdf-1 | 0.072024323 | 0.196449132 | -0.179507055 | 0.006 | 0.058 | 0.005959452 | 1 |
| WBGene00000122 | aly-3 | 0.820778441 | 0.945195512 | -0.179495891 | 0.156 | 0.295 | 0.009036649 | 1 |
| WBGene00012673 | dyf-17 | 0.172281491 | 0.296564605 | -0.179302633 | 0.029 | 0.074 | 0.066374164 | 1 |
| WBGene00010217 | sre-31 | 0.131296154 | 0.255396622 | -0.179039129 | 0.017 | 0.053 | 0.075938161 | 1 |
| WBGene00019121 | npl-4.2 | 0.254613467 | 0.378710225 | -0.179033777 | 0.046 | 0.053 | 0.846947579 | 1 |
| WBGene00007813 | jmjd-3.3 | 0.124638912 | 0.248424349 | -0.178584636 | 0.023 | 0.058 | 0.102650177 | 1 |
| WBGene00022696 | ift-139 | 0.933006724 | 1.056672131 | -0.178411468 | 0.179 | 0.321 | 0.01456825 | 1 |
| WBGene00000914 | daf-19 | 0.533409484 | 0.656871908 | -0.178118628 | 0.104 | 0.184 | 0.058329737 | 1 |
| WBGene00006720 | ubc-25 | 0.62192372 | 0.745331371 | -0.178039606 | 0.116 | 0.232 | 0.015359246 | 1 |
| WBGene00006481 | plrg-1 | 0.286031814 | 0.409060424 | -0.177492766 | 0.035 | 0.1 | 0.019133141 | 1 |
| WBGene00011250 | R11H6.2 | 0.470521961 | 0.593547765 | -0.177488718 | 0.058 | 0.147 | 0.008532176 | 1 |
| WBGene00006648 | ttb-1 | 0.213126521 | 0.336102189 | -0.177416386 | 0.04 | 0.079 | 0.136721436 | 1 |
| WBGene00004424 | rpl-12 | 3.471985735 | 3.594875042 | -0.177291794 | 0.873 | 0.953 | 0.03273578 | 1 |
| WBGene00004326 | rde-4 | 0.062079884 | 0.184890508 | -0.177178278 | 0.012 | 0.058 | 0.01947719 | 1 |
| WBGene00004095 | pqe-1 | 0.327857179 | 0.450504625 | -0.176942863 | 0.046 | 0.105 | 0.045354031 | 1 |
| WBGene00001078 | dpy-19 | 0.408266945 | 0.530760397 | -0.176720697 | 0.081 | 0.137 | 0.1463126 | 1 |
| WBGene00003789 | npp-3 | 0.075837658 | 0.198220007 | -0.176560408 | 0.012 | 0.053 | 0.031283818 | 1 |
| WBGene00003630 | nhr-40 | 0.548449783 | 0.670294963 | -0.175785437 | 0.087 | 0.179 | 0.019481011 | 1 |
| WBGene00022199 | pfk-1.1 | 0.132523898 | 0.254330121 | -0.175729234 | 0.023 | 0.058 | 0.103263284 | 1 |
| WBGene00001854 | hil-3 | 0.183484376 | 0.305228705 | -0.175639939 | 0.029 | 0.068 | 0.093758041 | 1 |
| WBGene00002027 | hsr-9 | 0.310405224 | 0.431595615 | -0.174840777 | 0.046 | 0.063 | 0.520841053 | 1 |
| WBGene00044386 | C27A2.7 | 0.358706691 | 0.479879836 | -0.174815894 | 0.075 | 0.142 | 0.068444614 | 1 |
| WBGene00005335 | srh-116 | 0.794655994 | 0.915787818 | -0.174756283 | 0.156 | 0.274 | 0.025648587 | 1 |
| WBGene00004482 | rps-13 | 2.159798903 | 2.280787835 | -0.174550132 | 0.555 | 0.842 | 0.001828879 | 1 |
| WBGene00010943 | M176.4 | 0.089603469 | 0.210225653 | -0.174021026 | 0.017 | 0.053 | 0.076453461 | 1 |
| WBGene00021800 | nduf-9 | 0.223048549 | 0.343507823 | -0.173785997 | 0.035 | 0.095 | 0.027266641 | 1 |
| WBGene00012996 | pinn-4 | 0.12345335 | 0.243687055 | -0.173460571 | 0.023 | 0.063 | 0.068688597 | 1 |
| WBGene00012178 | glb-27 | 2.582190305 | 2.702305121 | -0.173289049 | 0.711 | 0.821 | 0.090168289 | 1 |
| WBGene00000195 | arr-1 | 1.518737672 | 1.638649469 | -0.172996155 | 0.312 | 0.484 | 0.029326837 | 1 |
| WBGene00016374 | swd-2.2 | 0.145090006 | 0.264881673 | -0.172822844 | 0.029 | 0.079 | 0.0455437 | 1 |
| WBGene00004281 | rab-28 | 1.716239752 | 1.835783429 | -0.172465069 | 0.457 | 0.663 | 0.047330727 | 1 |
| WBGene00019403 | K05B2.2 | 0.238470221 | 0.358000507 | -0.172445751 | 0.04 | 0.1 | 0.037319508 | 1 |
| WBGene00003799 | npp-13 | 0.162906469 | 0.282375949 | -0.172358027 | 0.023 | 0.084 | 0.013677559 | 1 |
| WBGene00000479 | cgh-1 | 0.388343851 | 0.50759274 | -0.172039781 | 0.069 | 0.174 | 0.006118297 | 1 |
| WBGene00003649 | nhr-59 | 0.348370828 | 0.467566491 | -0.171962993 | 0.052 | 0.121 | 0.028507293 | 1 |

| | | | | | | | | |
|---|---|---|---|---|---|---|---|---|
| WBGene00018804 | lido-15 | 0.104394262 | 0.223541905 | -0.171893714 | 0.023 | 0.063 | 0.069118155 | 1 |
| WBGene00005581 | sri-69 | 0.231286823 | 0.350287405 | -0.17168155 | 0.035 | 0.084 | 0.057460111 | 1 |
| WBGene00019259 | H34C03.2 | 0.141399365 | 0.260267004 | -0.171489754 | 0.029 | 0.058 | 0.191839191 | 1 |
| WBGene00001127 | dyf-11 | 1.165100474 | 1.28395669 | -0.171473273 | 0.254 | 0.442 | 0.007358466 | 1 |
| WBGene00003130 | map-2 | 0.088991854 | 0.207771263 | -0.171362464 | 0.017 | 0.058 | 0.049097285 | 1 |
| WBGene00008221 | nhr-168 | 0.272455396 | 0.390973493 | -0.17098547 | 0.04 | 0.089 | 0.073215187 | 1 |
| WBGene00021710 | cyp-33C11 | 0.226364569 | 0.344836331 | -0.170918623 | 0.035 | 0.079 | 0.088583383 | 1 |
| WBGene00012354 | cox-4 | 0.922529718 | 1.040833992 | -0.170676989 | 0.202 | 0.3 | 0.15389122 | 1 |
| WBGene00006352 | sur-6 | 0.581556475 | 0.699837623 | -0.170643627 | 0.11 | 0.205 | 0.0353134 | 1 |
| WBGene00006715 | ubc-20 | 0.447173923 | 0.565305157 | -0.170427345 | 0.092 | 0.179 | 0.040830788 | 1 |
| WBGene00004387 | rnp-4 | 0.626147664 | 0.744099012 | -0.170167824 | 0.121 | 0.253 | 0.008018152 | 1 |
| WBGene00004273 | rab-10 | 0.594815225 | 0.712728598 | -0.170113039 | 0.104 | 0.221 | 0.008863854 | 1 |
| WBGene00001561 | gei-4 | 0.844988041 | 0.962888997 | -0.170095126 | 0.168 | 0.295 | 0.023065762 | 1 |
| WBGene00006829 | unc-101 | 0.12863313 | 0.24648054 | -0.170017873 | 0.017 | 0.063 | 0.032208001 | 1 |
| WBGene00022805 | nhr-256 | 0.259195241 | 0.376641903 | -0.169439717 | 0.035 | 0.105 | 0.013266874 | 1 |
| WBGene00011520 | nhr-213 | 0.437586209 | 0.554948696 | -0.169318278 | 0.075 | 0.174 | 0.011437887 | 1 |
| WBGene00012540 | Y37A1B.7 | 1.447926374 | 1.565137923 | -0.169100522 | 0.347 | 0.532 | 0.03849902 | 1 |
| WBGene00015285 | gmeb-1 | 0.405383958 | 0.522459592 | -0.168904436 | 0.069 | 0.142 | 0.041522104 | 1 |
| WBGene00020442 | gen-1 | 0.179162805 | 0.29610829 | -0.168716671 | 0.029 | 0.089 | 0.020278012 | 1 |
| WBGene00001787 | gst-39 | 1.181316752 | 1.298177383 | -0.168594253 | 0.266 | 0.426 | 0.027660021 | 1 |
| WBGene00014454 | MTCE.7 | 2.602288737 | 2.719063069 | -0.16846975 | 0.688 | 0.874 | 0.007752984 | 1 |
| WBGene00016619 | C43H6.6 | 0.77090197 | 0.887582795 | -0.168334847 | 0.145 | 0.226 | 0.10619294 | 1 |
| WBGene00004410 | rla-2 | 2.311730754 | 2.428281999 | -0.168147903 | 0.694 | 0.853 | 0.023301221 | 1 |
| WBGene00010556 | rack-1 | 2.197371093 | 2.313796706 | -0.167966655 | 0.618 | 0.837 | 0.017561908 | 1 |
| WBGene00019469 | srz-89 | 0.134148666 | 0.250451586 | -0.167789646 | 0.029 | 0.068 | 0.095862969 | 1 |
| WBGene00003902 | pab-1 | 1.433196835 | 1.549290846 | -0.167488254 | 0.329 | 0.553 | 0.020839631 | 1 |
| WBGene00009583 | aagr-3 | 0.460943068 | 0.57681877 | -0.167173301 | 0.075 | 0.121 | 0.215454946 | 1 |
| WBGene00019937 | R07E4.3 | 0.367791674 | 0.483549326 | -0.167002991 | 0.052 | 0.142 | 0.007727595 | 1 |
| WBGene00022382 | Y94H6A.10 | 0.458696828 | 0.574448996 | -0.16699508 | 0.058 | 0.116 | 0.06789488 | 1 |
| WBGene00022783 | tomm-7 | 0.060829416 | 0.176470638 | -0.166835019 | 0.012 | 0.053 | 0.031666734 | 1 |
| WBGene00018317 | F41H10.4 | 0.06655278 | 0.181932746 | -0.166458106 | 0.012 | 0.058 | 0.019639811 | 1 |
| WBGene00022489 | Y119D3B.12 | 0.503898141 | 0.618978509 | -0.166025877 | 0.081 | 0.158 | 0.041036679 | 1 |
| WBGene00012930 | obr-1 | 0.124696434 | 0.239716078 | -0.16593827 | 0.023 | 0.053 | 0.152600626 | 1 |
| WBGene00020112 | pfd-5 | 0.159153138 | 0.274164431 | -0.165926221 | 0.023 | 0.089 | 0.008954632 | 1 |
| WBGene00016837 | syf-1 | 0.134738689 | 0.249502909 | -0.165569771 | 0.023 | 0.074 | 0.031572411 | 1 |
| WBGene00018344 | npr-27 | 1.990239545 | 2.104968823 | -0.16551936 | 0.486 | 0.732 | 0.003997046 | 1 |
| WBGene00007505 | pezo-1 | 0.318665951 | 0.433098785 | -0.165091683 | 0.069 | 0.116 | 0.158631805 | 1 |
| WBGene00008750 | F13E9.1 | 0.113716122 | 0.228126148 | -0.165058777 | 0.017 | 0.058 | 0.050495994 | 1 |
| WBGene00006059 | stc-1 | 0.369414132 | 0.48366552 | -0.164829911 | 0.069 | 0.126 | 0.100533341 | 1 |
| WBGene00006965 | xtr-1 | 0.424567778 | 0.53881823 | -0.164828561 | 0.064 | 0.142 | 0.022404084 | 1 |
| WBGene00011517 | cgt-1 | 0.914747322 | 1.028695224 | -0.164392073 | 0.197 | 0.316 | 0.037999466 | 1 |
| WBGene00008919 | vps-36 | 0.248949218 | 0.362879811 | -0.164367102 | 0.04 | 0.079 | 0.150645041 | 1 |
| WBGene00005559 | sri-47 | 0.083055388 | 0.196926951 | -0.164281939 | 0.017 | 0.053 | 0.075938161 | 1 |
| WBGene00017023 | D1022.4 | 0.363890045 | 0.47770325 | -0.164197747 | 0.052 | 0.147 | 0.004975239 | 1 |
| WBGene00119203 | T04A8.18 | 0.625379464 | 0.739150299 | -0.164136619 | 0.133 | 0.195 | 0.260026741 | 1 |
| WBGene00019967 | cyp-33C8 | 1.739908672 | 1.85364028 | -0.164080027 | 0.353 | 0.521 | 0.02920035 | 1 |
| WBGene00015128 | B0303.7 | 0.198503744 | 0.312106029 | -0.163893453 | 0.04 | 0.095 | 0.055179705 | 1 |
| WBGene00014023 | lnkn-1 | 0.248462346 | 0.361977315 | -0.163767482 | 0.046 | 0.1 | 0.06867261 | 1 |
| WBGene00021785 | Y51H7C.7 | 0.139113685 | 0.252590806 | -0.16371288 | 0.017 | 0.053 | 0.075938161 | 1 |
| WBGene00000380 | cct-5 | 1.327645908 | 1.440925122 | -0.16342736 | 0.277 | 0.5 | 0.000740742 | 1 |
| WBGene00003000 | lin-11 | 1.729414013 | 1.842420905 | -0.163034484 | 0.41 | 0.705 | 0.000696114 | 1 |
| WBGene00006728 | ubq-2 | 2.706313083 | 2.81922637 | -0.16289944 | 0.763 | 0.911 | 0.0077136 | 1 |
| WBGene00016506 | abhd-5.1 | 0.343485049 | 0.456375472 | -0.162866453 | 0.069 | 0.116 | 0.175668767 | 1 |

| | | | | | | | | |
|---|---|---|---|---|---|---|---|---|
| WBGene00016393 | C34B2.8 | 0.440793104 | 0.553636951 | -0.162799259 | 0.081 | 0.168 | 0.024889376 | 1 |
| WBGene00002074 | ima-3 | 0.27751953 | 0.389978315 | -0.16224373 | 0.035 | 0.095 | 0.028254922 | 1 |
| WBGene00012094 | T27E9.2 | 0.789484074 | 0.901908294 | -0.162193865 | 0.168 | 0.263 | 0.084006685 | 1 |
| WBGene00003034 | lin-49 | 0.480828787 | 0.593142771 | -0.162034827 | 0.081 | 0.121 | 0.265315081 | 1 |
| WBGene00004702 | rsp-5 | 0.37324591 | 0.485548491 | -0.162018376 | 0.058 | 0.142 | 0.015098175 | 1 |
| WBGene00044027 | F27E5.8 | 0.130803745 | 0.24267974 | -0.161402944 | 0.023 | 0.058 | 0.103263284 | 1 |
| WBGene00173615 | 21ur-12311 | 0.298925607 | 0.41075848 | -0.161340732 | 0.04 | 0.095 | 0.047553629 | 1 |
| WBGene00012466 | ippk-1 | 0.074171417 | 0.185938654 | -0.161246039 | 0.017 | 0.053 | 0.075938161 | 1 |
| WBGene00009507 | mus-101 | 0.548006356 | 0.659595356 | -0.160988897 | 0.104 | 0.184 | 0.061430724 | 1 |
| WBGene00003563 | ncs-1 | 0.26190317 | 0.373478002 | -0.160968457 | 0.052 | 0.111 | 0.061809396 | 1 |
| WBGene00000923 | daf-31 | 0.139836688 | 0.251407663 | -0.160962892 | 0.029 | 0.063 | 0.133487939 | 1 |
| WBGene00008018 | tbc-8 | 0.635951119 | 0.747389023 | -0.160770912 | 0.11 | 0.195 | 0.057642942 | 1 |
| WBGene00019762 | M03F8.3 | 0.341679036 | 0.453097841 | -0.160743358 | 0.058 | 0.111 | 0.100043336 | 1 |
| WBGene00013242 | Y56A3A.30 | 0.123292206 | 0.234435052 | -0.160345232 | 0.023 | 0.053 | 0.154330178 | 1 |
| WBGene00007943 | C34F6.9 | 0.353661304 | 0.464756739 | -0.160276833 | 0.058 | 0.132 | 0.027668271 | 1 |
| WBGene00018847 | 6-Mar | 0.669209132 | 0.780240174 | -0.160183934 | 0.121 | 0.247 | 0.010330501 | 1 |
| WBGene00020163 | T02G5.3 | 1.302134869 | 1.413070173 | -0.160045812 | 0.289 | 0.537 | 0.000627029 | 1 |
| WBGene00050906 | F20E11.17 | 0.334372415 | 0.44528935 | -0.160019313 | 0.046 | 0.105 | 0.046651526 | 1 |
| WBGene00022169 | Y71H2AM.4 | 0.122815402 | 0.233679153 | -0.159942584 | 0.017 | 0.063 | 0.031974939 | 1 |
| WBGene00017715 | F22F1.3 | 0.302427104 | 0.413072114 | -0.159627007 | 0.046 | 0.084 | 0.169761811 | 1 |
| WBGene00020216 | trap-2 | 0.39223139 | 0.502876047 | -0.159626498 | 0.069 | 0.126 | 0.096596496 | 1 |
| WBGene00017778 | nono-1 | 0.619830867 | 0.730290535 | -0.159359615 | 0.127 | 0.195 | 0.125780519 | 1 |
| WBGene00013042 | cccp-1 | 0.761307596 | 0.871685163 | -0.159241169 | 0.121 | 0.195 | 0.112958052 | 1 |
| WBGene00015974 | snrp-40.2 | 0.627101801 | 0.737474474 | -0.159234109 | 0.104 | 0.147 | 0.297281012 | 1 |
| WBGene00008768 | F13G3.10 | 0.238364494 | 0.348654668 | -0.159115086 | 0.04 | 0.095 | 0.055469909 | 1 |
| WBGene00003357 | mir-264 | 0.630460855 | 0.740597818 | -0.15889405 | 0.11 | 0.205 | 0.033247355 | 1 |
| WBGene00011367 | snpc-3.4 | 0.079275369 | 0.189347783 | -0.158800925 | 0.012 | 0.053 | 0.031666734 | 1 |
| WBGene00002230 | klp-20 | 0.126545662 | 0.236502791 | -0.158634605 | 0.023 | 0.063 | 0.07174207 | 1 |
| WBGene00018160 | odr-8 | 1.245339712 | 1.355290197 | -0.158625018 | 0.289 | 0.453 | 0.032709043 | 1 |
| WBGene00009498 | tat-5 | 0.110191375 | 0.219907155 | -0.158286412 | 0.017 | 0.068 | 0.020555192 | 1 |
| WBGene00012056 | nhr-285 | 0.334642256 | 0.444329814 | -0.158245695 | 0.052 | 0.089 | 0.193454636 | 1 |
| WBGene00011693 | T10G3.2 | 1.230561054 | 1.340104934 | -0.158038412 | 0.266 | 0.447 | 0.020301402 | 1 |
| WBGene00010616 | K07A1.10 | 0.181475938 | 0.290980742 | -0.157982038 | 0.029 | 0.079 | 0.044728784 | 1 |
| WBGene00004276 | rab-14 | 0.75792494 | 0.867110038 | -0.157520799 | 0.145 | 0.268 | 0.015885989 | 1 |
| WBGene00019054 | F58F6.6 | 0.530541437 | 0.639689881 | -0.157467918 | 0.092 | 0.121 | 0.465175036 | 1 |
| WBGene00001303 | sec-61.G | 1.582736118 | 1.69185905 | -0.157431112 | 0.451 | 0.574 | 0.248564707 | 1 |
| WBGene00020719 | natc-1 | 0.194221014 | 0.303299256 | -0.157366638 | 0.035 | 0.068 | 0.167437587 | 1 |
| WBGene00022855 | tcer-1 | 0.181039185 | 0.290058164 | -0.15728114 | 0.035 | 0.074 | 0.121436957 | 1 |
| WBGene00004759 | sel-1 | 0.184122875 | 0.292698111 | -0.156640955 | 0.035 | 0.079 | 0.084939254 | 1 |
| WBGene00015011 | B0041.8 | 0.27582014 | 0.384326023 | -0.156540899 | 0.052 | 0.089 | 0.193454636 | 1 |
| WBGene00002244 | laf-1 | 0.302978926 | 0.411186572 | -0.156110635 | 0.04 | 0.105 | 0.025527037 | 1 |
| WBGene00003076 | lsm-1 | 0.327787 | 0.435984462 | -0.156095942 | 0.046 | 0.121 | 0.01626496 | 1 |
| WBGene00000229 | atp-2 | 1.106792356 | 1.214883642 | -0.155942762 | 0.249 | 0.437 | 0.011469123 | 1 |
| WBGene00012561 | Y37D8A.26 | 0.247807771 | 0.355853254 | -0.155876682 | 0.035 | 0.089 | 0.040326254 | 1 |
| WBGene00018727 | jbts-14 | 0.856355214 | 0.964221474 | -0.155618119 | 0.139 | 0.295 | 0.003384931 | 1 |
| WBGene00012113 | T28B8.1 | 0.863664592 | 0.971510687 | -0.155589026 | 0.139 | 0.305 | 0.001166169 | 1 |
| WBGene00003745 | nlp-7 | 2.583971631 | 2.691755977 | -0.155499941 | 0.717 | 0.921 | 0.002761519 | 1 |
| WBGene00000910 | daf-14 | 1.211626257 | 1.319360688 | -0.15542793 | 0.283 | 0.4 | 0.053629323 | 1 |
| WBGene00014019 | ZK632.12 | 0.229649919 | 0.337339287 | -0.155362918 | 0.023 | 0.1 | 0.003640001 | 1 |
| WBGene00004437 | rpl-24.2 | 0.308799545 | 0.416282725 | -0.155065451 | 0.058 | 0.121 | 0.053121953 | 1 |
| WBGene00018491 | mdh-1 | 0.449493286 | 0.556956975 | -0.155037332 | 0.081 | 0.163 | 0.035251982 | 1 |
| WBGene00010677 | gtbp-1 | 0.591531971 | 0.698953258 | -0.154976159 | 0.11 | 0.216 | 0.022530827 | 1 |
| WBGene00001489 | frm-2 | 0.264563736 | 0.371931809 | -0.154899386 | 0.04 | 0.089 | 0.074718813 | 1 |

| | | | | | | | | |
|---|---|---|---|---|---|---|---|---|
| WBGene00007416 | ceh-57 | 0.833976803 | 0.941314215 | -0.154855152 | 0.185 | 0.295 | 0.082968509 | 1 |
| WBGene00017121 | cyc-2.1 | 1.66679791 | 1.774120236 | -0.154833387 | 0.416 | 0.653 | 0.013821553 | 1 |
| WBGene00008747 | phf-31 | 0.105551457 | 0.212630747 | -0.15448276 | 0.023 | 0.058 | 0.102650177 | 1 |
| WBGene00043994 | C07A9.12 | 0.249079182 | 0.356091711 | -0.154386445 | 0.04 | 0.095 | 0.053183514 | 1 |
| WBGene00022790 | ZK682.7 | 0.789419139 | 0.896391052 | -0.154327848 | 0.173 | 0.295 | 0.046548143 | 1 |
| WBGene00022534 | syx-16 | 0.23052455 | 0.337433115 | -0.154236457 | 0.04 | 0.089 | 0.072472709 | 1 |
| WBGene00018739 | F53B1.8 | 0.960478655 | 1.067321094 | -0.154141056 | 0.168 | 0.316 | 0.006294301 | 1 |
| WBGene00012558 | rbm-7 | 0.091844485 | 0.198556513 | -0.153952913 | 0.012 | 0.058 | 0.019639811 | 1 |
| WBGene00022300 | Y76B12C.6 | 0.097761511 | 0.204283116 | -0.153678191 | 0.017 | 0.053 | 0.077492615 | 1 |
| WBGene00001112 | duo-3 | 0.110891125 | 0.217344078 | -0.153579148 | 0.017 | 0.058 | 0.049792522 | 1 |
| WBGene00012197 | W02B8.1 | 0.309104187 | 0.415479965 | -0.153467807 | 0.058 | 0.111 | 0.100470392 | 1 |
| WBGene00008570 | kcnl-2 | 0.363834947 | 0.470066349 | -0.153259517 | 0.064 | 0.105 | 0.196946033 | 1 |
| WBGene00007617 | rrbs-1 | 0.243199868 | 0.349318024 | -0.153096136 | 0.04 | 0.074 | 0.196772319 | 1 |
| WBGene00004891 | smr-1 | 0.334358211 | 0.440450939 | -0.153059452 | 0.04 | 0.137 | 0.002276936 | 1 |
| WBGene00019106 | irld-38 | 0.093183503 | 0.199250512 | -0.153022347 | 0.012 | 0.053 | 0.031157212 | 1 |
| WBGene00006529 | tba-2 | 1.038380963 | 1.144226566 | -0.152702927 | 0.214 | 0.374 | 0.010796163 | 1 |
| WBGene00004446 | rpl-32 | 2.828730178 | 2.934490424 | -0.152579782 | 0.792 | 0.926 | 0.051545585 | 1 |
| WBGene00016630 | acer-1 | 0.400840495 | 0.506452488 | -0.152365897 | 0.075 | 0.132 | 0.106862953 | 1 |
| WBGene00006836 | unc-112 | 0.134332614 | 0.239708699 | -0.152025554 | 0.017 | 0.058 | 0.048752734 | 1 |
| WBGene00022664 | ZK121.2 | 1.499249412 | 1.604587021 | -0.151970047 | 0.312 | 0.474 | 0.050854254 | 1 |
| WBGene00016902 | C53D5.1 | 0.183851515 | 0.288760691 | -0.151351948 | 0.029 | 0.058 | 0.186150557 | 1 |
| WBGene00009829 | tmed-10 | 0.406956631 | 0.511806309 | -0.15126611 | 0.064 | 0.132 | 0.048117701 | 1 |
| WBGene00001088 | dpy-30 | 0.728616612 | 0.833381179 | -0.151143321 | 0.133 | 0.253 | 0.015692383 | 1 |
| WBGene00021132 | nfyb-1 | 0.210438082 | 0.315150196 | -0.151067648 | 0.029 | 0.068 | 0.093237643 | 1 |
| WBGene00016990 | vps-37 | 0.238715419 | 0.343225463 | -0.150776123 | 0.052 | 0.1 | 0.116284608 | 1 |
| WBGene00010809 | M01F1.3 | 0.102256171 | 0.206677074 | -0.150647518 | 0.017 | 0.053 | 0.076971609 | 1 |
| WBGene00008444 | tbc-14 | 0.463055759 | 0.567464833 | -0.150630454 | 0.069 | 0.142 | 0.041522059 | 1 |
| WBGene00005362 | srh-146 | 1.708101762 | 1.812484804 | -0.150592898 | 0.416 | 0.621 | 0.03365225 | 1 |
| WBGene00018696 | pho-6 | 1.119411515 | 1.223705426 | -0.150464307 | 0.214 | 0.416 | 0.002675344 | 1 |
| WBGene00017380 | F11D5.1 | 0.527226957 | 0.631440308 | -0.150348085 | 0.087 | 0.137 | 0.206307241 | 1 |
| WBGene00003619 | nhr-20 | 0.993872951 | 1.098038576 | -0.150279231 | 0.208 | 0.289 | 0.27884821 | 1 |
| WBGene00001081 | dpy-22 | 0.386846653 | 0.490919292 | -0.150145081 | 0.052 | 0.084 | 0.273999666 | 1 |
| WBGene00005587 | sri-75 | 0.324264901 | 0.428275845 | -0.150056074 | 0.058 | 0.121 | 0.050462941 | 1 |
| WBGene00011737 | sqst-1 | 1.515895156 | 1.61968983 | -0.14974406 | 0.376 | 0.537 | 0.105079753 | 1 |
| WBGene00011679 | ucr-2.2 | 0.170300578 | 0.274042523 | -0.14966799 | 0.023 | 0.063 | 0.07174207 | 1 |
| WBGene00012794 | Y43E12A.2 | 0.309701947 | 0.413309893 | -0.14947467 | 0.046 | 0.111 | 0.032634946 | 1 |
| WBGene00016747 | nmur-1 | 0.161404395 | 0.264957406 | -0.149395417 | 0.023 | 0.079 | 0.020918247 | 1 |
| WBGene00001427 | fkb-2 | 1.673428421 | 1.776781771 | -0.149107366 | 0.445 | 0.616 | 0.112174202 | 1 |
| WBGene00006721 | ubh-1 | 0.16162151 | 0.264774137 | -0.148817782 | 0.029 | 0.068 | 0.095333223 | 1 |
| WBGene00005218 | srg-61 | 0.281095286 | 0.384243879 | -0.148811964 | 0.052 | 0.105 | 0.082738755 | 1 |
| WBGene00003806 | npp-20 | 0.199296205 | 0.302203595 | -0.148463982 | 0.035 | 0.053 | 0.427970971 | 1 |
| WBGene00003078 | lsm-4 | 0.208306619 | 0.311195722 | -0.148437599 | 0.04 | 0.079 | 0.140594242 | 1 |
| WBGene00020950 | dlst-1 | 0.256250312 | 0.35912653 | -0.14841901 | 0.035 | 0.068 | 0.169845756 | 1 |
| WBGene00017238 | F08B4.7 | 0.406475742 | 0.509194696 | -0.148192125 | 0.075 | 0.126 | 0.161220545 | 1 |
| WBGene00003700 | nhr-110 | 0.274835551 | 0.377518562 | -0.14814027 | 0.052 | 0.095 | 0.143511662 | 1 |
| WBGene00011488 | nra-2 | 0.212057799 | 0.314665025 | -0.148030936 | 0.029 | 0.074 | 0.065610102 | 1 |
| WBGene00009925 | F52B11.2 | 0.18837933 | 0.290923797 | -0.147940394 | 0.029 | 0.068 | 0.092719566 | 1 |
| WBGene00003635 | nhr-45 | 0.528189332 | 0.630725077 | -0.147927811 | 0.092 | 0.158 | 0.089643366 | 1 |
| WBGene00017653 | F21C10.3 | 2.576619231 | 2.679122743 | -0.147881307 | 0.705 | 0.853 | 0.017705072 | 1 |
| WBGene00002190 | kin-2 | 0.915699065 | 1.018078768 | -0.14770269 | 0.173 | 0.279 | 0.070596559 | 1 |
| WBGene00004469 | rps-0 | 2.33340733 | 2.435723641 | -0.147611235 | 0.665 | 0.826 | 0.026125563 | 1 |
| WBGene00007129 | B0272.3 | 0.232718641 | 0.335032854 | -0.147608207 | 0.04 | 0.079 | 0.149958391 | 1 |
| WBGene00013690 | hprt-1 | 0.172257757 | 0.27448383 | -0.14748105 | 0.029 | 0.063 | 0.135625358 | 1 |

| | | | | | | | | |
|---|---|---|---|---|---|---|---|---|
| WBGene00001749 | gst-1 | 0.501279144 | 0.603449787 | -0.14740108 | 0.098 | 0.137 | 0.317196476 | 1 |
| WBGene00009578 | F40F8.5 | 1.347175641 | 1.449013367 | -0.146920782 | 0.318 | 0.463 | 0.112948974 | 1 |
| WBGene00003990 | pfn-2 | 0.965937278 | 1.067630423 | -0.146712196 | 0.185 | 0.253 | 0.348166377 | 1 |
| WBGene00000199 | arx-1 | 0.563634541 | 0.66527104 | -0.146630473 | 0.092 | 0.153 | 0.120721101 | 1 |
| WBGene00016246 | daf-37 | 1.520390044 | 1.621946556 | -0.146515076 | 0.335 | 0.521 | 0.02982455 | 1 |
| WBGene00009664 | idha-1 | 0.284438602 | 0.385811716 | -0.146250488 | 0.04 | 0.116 | 0.012654514 | 1 |
| WBGene00020246 | clic-1 | 0.899229385 | 1.000538617 | -0.146158326 | 0.179 | 0.326 | 0.01501203 | 1 |
| WBGene00004918 | snr-5 | 0.440871039 | 0.541864731 | -0.145703099 | 0.087 | 0.163 | 0.05857683 | 1 |
| WBGene00002324 | let-49 | 0.227320192 | 0.328281822 | -0.145656842 | 0.035 | 0.1 | 0.018442371 | 1 |
| WBGene00020334 | epg-7 | 0.280632603 | 0.38131207 | -0.145249769 | 0.04 | 0.116 | 0.012503465 | 1 |
| WBGene00009691 | F44E5.4 | 4.093066997 | 4.193558783 | -0.144979001 | 0.942 | 0.974 | 0.047930547 | 1 |
| WBGene00001232 | eif-3.I | 0.179112086 | 0.279523334 | -0.14486281 | 0.029 | 0.068 | 0.096395069 | 1 |
| WBGene00020031 | suco-1 | 0.29118368 | 0.391332957 | -0.144484866 | 0.052 | 0.1 | 0.106558411 | 1 |
| WBGene00003924 | pas-3 | 0.866295456 | 0.966418966 | -0.144447692 | 0.191 | 0.295 | 0.087156076 | 1 |
| WBGene00020346 | rbm-5 | 0.19632573 | 0.296386597 | -0.144357317 | 0.029 | 0.079 | 0.043135813 | 1 |
| WBGene00003239 | mig-2 | 0.276712269 | 0.376644323 | -0.144171478 | 0.052 | 0.084 | 0.249760465 | 1 |
| WBGene00007256 | swsn-9 | 0.280026046 | 0.379865423 | -0.144037774 | 0.052 | 0.105 | 0.085027746 | 1 |
| WBGene00005696 | sru-33 | 0.490276119 | 0.589954953 | -0.14380616 | 0.092 | 0.142 | 0.205785268 | 1 |
| WBGene00005400 | srh-185 | 0.15348734 | 0.253119443 | -0.143738741 | 0.029 | 0.053 | 0.265388085 | 1 |
| WBGene00007676 | srz-57 | 0.339651971 | 0.439171184 | -0.143575875 | 0.035 | 0.105 | 0.012222782 | 1 |
| WBGene00020018 | R12B2.2 | 0.137051201 | 0.236542125 | -0.143535062 | 0.012 | 0.068 | 0.00747887 | 1 |
| WBGene00000149 | apl-1 | 1.473121495 | 1.572499023 | -0.143371466 | 0.341 | 0.553 | 0.020114866 | 1 |
| WBGene00004927 | snx-1 | 0.199585305 | 0.298905748 | -0.143289112 | 0.035 | 0.063 | 0.223228413 | 1 |
| WBGene00015755 | C14B9.10 | 1.028145193 | 1.127211402 | -0.142922328 | 0.197 | 0.405 | 0.000996823 | 1 |
| WBGene00021011 | W03G1.5 | 0.150533749 | 0.24935954 | -0.142575479 | 0.023 | 0.053 | 0.154330178 | 1 |
| WBGene00001708 | grk-1 | 0.30956333 | 0.408331228 | -0.142491956 | 0.058 | 0.079 | 0.459815204 | 1 |
| WBGene00000247 | bec-1 | 0.692887554 | 0.791607808 | -0.142423221 | 0.156 | 0.237 | 0.136015079 | 1 |
| WBGene00022835 | ZK973.9 | 0.122536667 | 0.221143758 | -0.142259962 | 0.017 | 0.058 | 0.049097285 | 1 |
| WBGene00020838 | trak-1 | 0.435258076 | 0.533752088 | -0.142096822 | 0.087 | 0.147 | 0.120743823 | 1 |
| WBGene00017162 | ddx-23 | 0.099413748 | 0.197893779 | -0.142076652 | 0.017 | 0.058 | 0.051207776 | 1 |
| WBGene00001340 | etr-1 | 1.373799457 | 1.471935498 | -0.141580379 | 0.283 | 0.442 | 0.035900708 | 1 |
| WBGene00000931 | dao-5 | 0.139882699 | 0.237869489 | -0.141365056 | 0.006 | 0.053 | 0.009929999 | 1 |
| WBGene00007848 | cytb-5.1 | 0.995331533 | 1.093279869 | -0.141309578 | 0.191 | 0.358 | 0.006102532 | 1 |
| WBGene00016158 | ari-1.1 | 0.408257343 | 0.505891467 | -0.140856267 | 0.064 | 0.147 | 0.018568561 | 1 |
| WBGene00003953 | pbs-7 | 0.782993275 | 0.880601032 | -0.140818226 | 0.156 | 0.279 | 0.026648113 | 1 |
| WBGene00001503 | fum-1 | 0.356488025 | 0.454059804 | -0.140766321 | 0.064 | 0.137 | 0.0380526 | 1 |
| WBGene00022599 | daf-41 | 1.746503668 | 1.843775462 | -0.140333535 | 0.491 | 0.674 | 0.148031825 | 1 |
| WBGene00010800 | srsx-37 | 0.259861027 | 0.357108548 | -0.140298516 | 0.04 | 0.1 | 0.039627642 | 1 |
| WBGene00001669 | gpa-7 | 0.425307247 | 0.522371876 | -0.140034659 | 0.075 | 0.158 | 0.030896492 | 1 |
| WBGene00006602 | tps-1 | 0.277160016 | 0.374180972 | -0.139971653 | 0.052 | 0.095 | 0.155954432 | 1 |
| WBGene00020647 | pqbp-1.1 | 0.102701638 | 0.199222892 | -0.139250735 | 0.017 | 0.063 | 0.03291594 | 1 |
| WBGene00000235 | baf-1 | 0.205556574 | 0.301834515 | -0.138899708 | 0.029 | 0.079 | 0.043926202 | 1 |
| WBGene00011223 | R10H10.4 | 0.689190674 | 0.785245052 | -0.138577174 | 0.133 | 0.221 | 0.063710502 | 1 |
| WBGene00007136 | B0285.3 | 0.129628938 | 0.225560945 | -0.138400631 | 0.023 | 0.063 | 0.072634636 | 1 |
| WBGene00005640 | srn-1 | 0.129858716 | 0.225644207 | -0.138189252 | 0.017 | 0.058 | 0.050143224 | 1 |
| WBGene00002210 | kin-29 | 0.290317457 | 0.386080543 | -0.13815693 | 0.046 | 0.105 | 0.046532354 | 1 |
| WBGene00021628 | Y47D9A.1 | 0.18515571 | 0.280633469 | -0.137745289 | 0.029 | 0.079 | 0.044728784 | 1 |
| WBGene00021522 | nhr-274 | 0.159774583 | 0.255009276 | -0.13739462 | 0.017 | 0.058 | 0.049792522 | 1 |
| WBGene00012766 | Y41E3.8 | 0.650876866 | 0.745956713 | -0.137171224 | 0.121 | 0.226 | 0.029970185 | 1 |
| WBGene00023264 | D1065.2 | 1.001722948 | 1.096760933 | -0.137110829 | 0.214 | 0.358 | 0.0432974 | 1 |
| WBGene00017894 | F28B12.1 | 1.766265376 | 1.861269415 | -0.137061856 | 0.474 | 0.689 | 0.057235329 | 1 |
| WBGene00009676 | F44A6.4 | 0.361361771 | 0.456331933 | -0.137012982 | 0.069 | 0.132 | 0.077035657 | 1 |
| WBGene00005329 | srh-110 | 0.425444647 | 0.520364187 | -0.136939949 | 0.075 | 0.126 | 0.151198384 | 1 |

| | | | | | | | | |
|---|---|---|---|---|---|---|---|---|
| WBGene00000200 | arx-2 | 0.220572218 | 0.315469916 | -0.136908438 | 0.029 | 0.074 | 0.064853305 | 1 |
| WBGene00011996 | T24F1.4 | 0.618996516 | 0.713763339 | -0.136719624 | 0.127 | 0.2 | 0.120271305 | 1 |
| WBGene00008205 | sams-1 | 0.233741102 | 0.328505317 | -0.136715862 | 0.023 | 0.1 | 0.003296807 | 1 |
| WBGene00016928 | srb-19 | 0.609646325 | 0.704396011 | -0.136694903 | 0.087 | 0.174 | 0.021017007 | 1 |
| WBGene00009194 | F27E5.5 | 0.55313148 | 0.647756588 | -0.136515174 | 0.064 | 0.168 | 0.004360864 | 1 |
| WBGene00007456 | srz-29 | 1.331221821 | 1.425825039 | -0.136483594 | 0.312 | 0.453 | 0.097665974 | 1 |
| WBGene00004116 | pqn-27 | 0.238890577 | 0.333224582 | -0.136095201 | 0.046 | 0.1 | 0.0707063 | 1 |
| WBGene00010484 | dph-3 | 0.319478915 | 0.413520731 | -0.135673661 | 0.069 | 0.111 | 0.212356216 | 1 |
| WBGene00010882 | atg-7 | 0.384208207 | 0.478223908 | -0.135635987 | 0.069 | 0.089 | 0.495460462 | 1 |
| WBGene00044780 | F18E9.8 | 0.511585448 | 0.605540862 | -0.13554901 | 0.104 | 0.179 | 0.086873495 | 1 |
| WBGene00008506 | tkt-1 | 0.121022067 | 0.214955863 | -0.135517821 | 0.023 | 0.053 | 0.15520052 | 1 |
| WBGene00003646 | nhr-56 | 0.52971635 | 0.623603876 | -0.135451068 | 0.11 | 0.168 | 0.198641993 | 1 |
| WBGene00018164 | sup-36 | 0.683017818 | 0.776592378 | -0.134999553 | 0.127 | 0.247 | 0.018014046 | 1 |
| WBGene00012547 | Y37D8A.5 | 1.050226425 | 1.143709839 | -0.134868057 | 0.214 | 0.358 | 0.028095112 | 1 |
| WBGene00007350 | sucl-1 | 0.442201265 | 0.535650918 | -0.13481935 | 0.098 | 0.158 | 0.15997752 | 1 |
| WBGene00011216 | usp-46 | 0.10430667 | 0.197651947 | -0.134668767 | 0.017 | 0.058 | 0.050495994 | 1 |
| WBGene00002083 | inf-1 | 0.713777928 | 0.807101794 | -0.134637877 | 0.139 | 0.253 | 0.029064005 | 1 |
| WBGene00011858 | T20D3.5 | 0.126814707 | 0.220105501 | -0.134590166 | 0.029 | 0.058 | 0.191839191 | 1 |
| WBGene00020696 | pygl-1 | 0.479930577 | 0.573161047 | -0.134503138 | 0.075 | 0.158 | 0.029536671 | 1 |
| WBGene00018219 | F40A3.4 | 0.508034129 | 0.601175098 | -0.134374014 | 0.069 | 0.121 | 0.134653212 | 1 |
| WBGene00006523 | tam-1 | 0.416708441 | 0.509810223 | -0.134317479 | 0.069 | 0.111 | 0.207247524 | 1 |
| WBGene00014165 | puf-12 | 0.109913428 | 0.202276204 | -0.133251318 | 0.012 | 0.058 | 0.019639811 | 1 |
| WBGene00007980 | C36E8.1 | 0.373686773 | 0.465989632 | -0.133164878 | 0.075 | 0.142 | 0.070138629 | 1 |
| WBGene00004456 | rpl-43 | 2.699403901 | 2.791541975 | -0.132927142 | 0.775 | 0.916 | 0.195438762 | 1 |
| WBGene00020235 | T05A12.4 | 0.14028381 | 0.232376626 | -0.132861849 | 0.023 | 0.063 | 0.072634636 | 1 |
| WBGene00008000 | slc-17.2 | 0.953206561 | 1.045160789 | -0.132661909 | 0.173 | 0.295 | 0.029873732 | 1 |
| WBGene00021331 | glrx-10 | 0.457945674 | 0.549834656 | -0.13256778 | 0.087 | 0.137 | 0.177197494 | 1 |
| WBGene00005554 | sri-42 | 0.199123024 | 0.2906002 | -0.131973669 | 0.029 | 0.074 | 0.063361313 | 1 |
| WBGene00006055 | ssr-2 | 0.620386923 | 0.711862713 | -0.131971669 | 0.104 | 0.179 | 0.078721128 | 1 |
| WBGene00001334 | ero-1 | 0.271773519 | 0.363160999 | -0.131844265 | 0.046 | 0.095 | 0.092278607 | 1 |
| WBGene00009100 | rgef-1 | 0.334576623 | 0.425938426 | -0.131807219 | 0.04 | 0.126 | 0.005401656 | 1 |
| WBGene00015352 | C02F5.12 | 0.523263511 | 0.61460829 | -0.131782661 | 0.098 | 0.174 | 0.068569681 | 1 |
| WBGene00007053 | chd-7 | 1.289367229 | 1.380701274 | -0.131767174 | 0.277 | 0.463 | 0.019695954 | 1 |
| WBGene00004766 | sel-9 | 0.252695287 | 0.343981606 | -0.13169832 | 0.035 | 0.089 | 0.040908003 | 1 |
| WBGene00011015 | R04F11.2 | 1.062837712 | 1.154040524 | -0.131577844 | 0.254 | 0.453 | 0.015946059 | 1 |
| WBGene00009975 | F53C11.5 | 0.34750479 | 0.438544047 | -0.131341885 | 0.052 | 0.121 | 0.031918391 | 1 |
| WBGene00010961 | nduo-2 | 0.95102139 | 1.042013583 | -0.131273985 | 0.191 | 0.253 | 0.294109938 | 1 |
| WBGene00000403 | casy-1 | 2.352495956 | 2.4433356 | -0.131053903 | 0.613 | 0.8 | 0.076369099 | 1 |
| WBGene00014294 | C53C7.5 | 1.889197088 | 1.98001902 | -0.131028351 | 0.439 | 0.658 | 0.024650666 | 1 |
| WBGene00010425 | lpin-1 | 0.174389649 | 0.265159077 | -0.130952604 | 0.035 | 0.063 | 0.226278314 | 1 |
| WBGene00016925 | srb-18 | 1.083832687 | 1.174401687 | -0.130663447 | 0.191 | 0.321 | 0.03205019 | 1 |
| WBGene00219748 | linc-46 | 0.15025909 | 0.240653394 | -0.130411414 | 0.029 | 0.068 | 0.097466357 | 1 |
| WBGene00005040 | sra-14 | 1.33269207 | 1.423012654 | -0.130305059 | 0.341 | 0.4 | 0.70102412 | 1 |
| WBGene00018156 | ncbp-1 | 0.285684196 | 0.375919882 | -0.130182578 | 0.058 | 0.105 | 0.135087246 | 1 |
| WBGene00044134 | D1025.10 | 0.258996374 | 0.34920787 | -0.130147678 | 0.035 | 0.084 | 0.057142918 | 1 |
| WBGene00001980 | hmr-1 | 0.514722394 | 0.604896379 | -0.130093562 | 0.087 | 0.158 | 0.059191062 | 1 |
| WBGene00006615 | trp-2 | 0.213686539 | 0.303824349 | -0.130041372 | 0.029 | 0.063 | 0.131376679 | 1 |
| WBGene00017766 | peli-1 | 1.652795379 | 1.742920077 | -0.130022454 | 0.387 | 0.611 | 0.019172794 | 1 |
| WBGene00008852 | ubql-1 | 0.812729598 | 0.902821312 | -0.12997487 | 0.162 | 0.305 | 0.011993453 | 1 |
| WBGene00004305 | ran-4 | 0.292485204 | 0.382336649 | -0.129628234 | 0.046 | 0.116 | 0.02409167 | 1 |
| WBGene00011033 | R05D11.5 | 0.115867838 | 0.205716147 | -0.12962371 | 0.023 | 0.058 | 0.109877914 | 1 |
| WBGene00006576 | tkr-1 | 0.815036393 | 0.904881492 | -0.129619079 | 0.133 | 0.184 | 0.338573145 | 1 |
| WBGene00003980 | pes-7 | 0.314389363 | 0.404190405 | -0.129555518 | 0.052 | 0.1 | 0.115782277 | 1 |

| | | | | | | | | |
|---|---|---|---|---|---|---|---|---|
| WBGene00000414 | cec-1 | 0.993537383 | 1.083317286 | -0.129525021 | 0.162 | 0.347 | 0.001236574 | 1 |
| WBGene00016005 | ift-74 | 1.008621262 | 1.09823427 | -0.129284241 | 0.191 | 0.389 | 0.002217943 | 1 |
| WBGene00000488 | che-7 | 0.470947619 | 0.560488037 | -0.129179517 | 0.092 | 0.142 | 0.191455714 | 1 |
| WBGene00017084 | E01A2.1 | 0.401420706 | 0.490789452 | -0.128931847 | 0.075 | 0.163 | 0.024683083 | 1 |
| WBGene00012769 | hrpu-1 | 0.272665766 | 0.361921666 | -0.128769045 | 0.046 | 0.095 | 0.097144695 | 1 |
| WBGene00001056 | dpf-3 | 0.198000179 | 0.287221193 | -0.128718714 | 0.017 | 0.068 | 0.020864658 | 1 |
| WBGene00007107 | pfd-4 | 0.556420572 | 0.645584993 | -0.128637069 | 0.116 | 0.211 | 0.041993643 | 1 |
| WBGene00020636 | T20H4.5 | 0.317718664 | 0.406882239 | -0.128635848 | 0.058 | 0.116 | 0.072628741 | 1 |
| WBGene00021854 | Y54F10AR.1 | 0.474329317 | 0.563474779 | -0.128609715 | 0.087 | 0.163 | 0.057203468 | 1 |
| WBGene00022188 | Y71H2AR.1 | 0.147646668 | 0.236752351 | -0.128552327 | 0.017 | 0.063 | 0.032442517 | 1 |
| WBGene00015801 | C15H9.5 | 1.029350302 | 1.118360772 | -0.128414964 | 0.173 | 0.226 | 0.313896579 | 1 |
| WBGene00008136 | C47D12.2 | 0.208976247 | 0.297963405 | -0.128381331 | 0.023 | 0.068 | 0.047945141 | 1 |
| WBGene00021170 | arx-4 | 0.225623697 | 0.31452006 | -0.128250342 | 0.035 | 0.079 | 0.088583383 | 1 |
| WBGene00012344 | ola-1 | 0.286259458 | 0.375142968 | -0.128231798 | 0.046 | 0.116 | 0.023960533 | 1 |
| WBGene00008398 | D2005.3 | 0.816207169 | 0.905059267 | -0.128186481 | 0.145 | 0.237 | 0.079174219 | 1 |
| WBGene00019426 | cutl-16 | 0.737832734 | 0.826628586 | -0.128105335 | 0.139 | 0.242 | 0.044198368 | 1 |
| WBGene00012221 | W03C9.5 | 0.75295208 | 0.841707131 | -0.128046472 | 0.145 | 0.253 | 0.036299856 | 1 |
| WBGene00016126 | nhr-258 | 2.184935984 | 2.273526443 | -0.127809017 | 0.601 | 0.768 | 0.08945627 | 1 |
| WBGene00010603 | nhr-198 | 1.276942131 | 1.365530863 | -0.127806525 | 0.283 | 0.395 | 0.172507024 | 1 |
| WBGene00004882 | smg-4 | 0.11955922 | 0.208044316 | -0.12765701 | 0.017 | 0.058 | 0.050495994 | 1 |
| WBGene00004888 | smo-1 | 1.187470129 | 1.275875466 | -0.127541941 | 0.295 | 0.437 | 0.110401756 | 1 |
| WBGene00012908 | Y46G5A.18 | 0.326196088 | 0.414506562 | -0.127405082 | 0.058 | 0.116 | 0.075261848 | 1 |
| WBGene00011303 | R166.3 | 0.446586928 | 0.534894108 | -0.12740033 | 0.058 | 0.111 | 0.105709196 | 1 |
| WBGene00004105 | szy-20 | 0.349787061 | 0.43804464 | -0.127328771 | 0.058 | 0.105 | 0.133451249 | 1 |
| WBGene00019840 | R02F11.2 | 3.268832711 | 3.356958344 | -0.127138413 | 0.902 | 0.979 | 0.18233722 | 1 |
| WBGene00003603 | nhr-4 | 0.774049309 | 0.861925324 | -0.126778291 | 0.133 | 0.242 | 0.025940874 | 1 |
| WBGene00021963 | lipl-6 | 0.097715721 | 0.185515281 | -0.12666799 | 0.017 | 0.053 | 0.077492615 | 1 |
| WBGene00016397 | maph-9 | 1.354854357 | 1.442571275 | -0.126548763 | 0.324 | 0.484 | 0.033974899 | 1 |
| WBGene00009973 | F53C11.3 | 1.26758852 | 1.355163622 | -0.126344165 | 0.231 | 0.379 | 0.019359758 | 1 |
| WBGene00010507 | K02E2.6 | 0.793606755 | 0.881145263 | -0.126291371 | 0.179 | 0.311 | 0.045765634 | 1 |
| WBGene00001017 | dnc-1 | 0.19543136 | 0.282886589 | -0.126171225 | 0.023 | 0.079 | 0.021062521 | 1 |
| WBGene00018009 | dhhc-3 | 0.255672583 | 0.343048632 | -0.126056992 | 0.04 | 0.1 | 0.038563827 | 1 |
| WBGene00044268 | T27E7.10 | 0.678388576 | 0.765592042 | -0.125808007 | 0.11 | 0.205 | 0.027547993 | 1 |
| WBGene00016496 | C37C3.2 | 0.706528976 | 0.793700531 | -0.12576197 | 0.145 | 0.247 | 0.057907702 | 1 |
| WBGene00019537 | K08D12.3 | 1.221557153 | 1.30861201 | -0.125593612 | 0.272 | 0.453 | 0.019561917 | 1 |
| WBGene00016140 | rpb-2 | 0.23091171 | 0.317920382 | -0.125526981 | 0.04 | 0.074 | 0.196772319 | 1 |
| WBGene00006072 | str-4 | 1.333606836 | 1.420570807 | -0.125462489 | 0.312 | 0.463 | 0.066145813 | 1 |
| WBGene00004754 | sec-23 | 0.174816983 | 0.26169423 | -0.125337374 | 0.029 | 0.068 | 0.092719566 | 1 |
| WBGene00016318 | C32D5.11 | 0.472683449 | 0.559397344 | -0.125101706 | 0.092 | 0.168 | 0.070415745 | 1 |
| WBGene00000906 | daf-10 | 1.429005196 | 1.515433579 | -0.124689801 | 0.353 | 0.521 | 0.103437556 | 1 |
| WBGene00009451 | cids-2 | 0.207110118 | 0.293311856 | -0.12436282 | 0.035 | 0.089 | 0.043552551 | 1 |
| WBGene00001130 | dyn-1 | 0.13520821 | 0.221307338 | -0.124214785 | 0.023 | 0.063 | 0.072634636 | 1 |
| WBGene00013518 | Y73F8A.5 | 0.356679335 | 0.442735399 | -0.124152657 | 0.064 | 0.121 | 0.090453068 | 1 |
| WBGene00016744 | bbs-9 | 1.227369668 | 1.313250052 | -0.123899205 | 0.237 | 0.495 | 0.000148721 | 0.921919345 |
| WBGene00003621 | nhr-22 | 0.20802868 | 0.293898082 | -0.12388336 | 0.035 | 0.074 | 0.120829088 | 1 |
| WBGene00001440 | fkh-8 | 2.547472682 | 2.633186096 | -0.123658317 | 0.613 | 0.768 | 0.204037447 | 1 |
| WBGene00003748 | nlp-10 | 2.401054752 | 2.486696274 | -0.123554599 | 0.653 | 0.853 | 0.05838467 | 1 |
| WBGene00009764 | srz-17 | 0.546822756 | 0.632399739 | -0.12346149 | 0.121 | 0.163 | 0.376026031 | 1 |
| WBGene00004415 | rpl-4 | 1.937586658 | 2.023159044 | -0.123454857 | 0.462 | 0.774 | 0.000863615 | 1 |
| WBGene00044650 | Y71G10AR.4 | 0.27933839 | 0.364894095 | -0.123430791 | 0.052 | 0.079 | 0.331683824 | 1 |
| WBGene00021539 | Y42H9AR.4 | 0.410025237 | 0.495523627 | -0.123348103 | 0.069 | 0.137 | 0.060094824 | 1 |
| WBGene00011575 | madf-4 | 0.339367544 | 0.424856158 | -0.123333999 | 0.058 | 0.116 | 0.077287539 | 1 |
| WBGene00003196 | mel-11 | 0.202634569 | 0.288008019 | -0.123167853 | 0.035 | 0.074 | 0.120223603 | 1 |

| | | | | | | | | |
|---|---|---|---|---|---|---|---|---|
| WBGene00011382 | dop-5 | 0.524140546 | 0.60935383 | -0.122936783 | 0.069 | 0.147 | 0.029916798 | 1 |
| WBGene00021844 | sec-11 | 0.307286333 | 0.392445402 | -0.122858567 | 0.052 | 0.116 | 0.045386799 | 1 |
| WBGene00006889 | pfd-3 | 0.55217363 | 0.63726687 | -0.122763595 | 0.092 | 0.153 | 0.137934845 | 1 |
| WBGene00018212 | F39H12.1 | 0.131037683 | 0.216126925 | -0.122757828 | 0.023 | 0.053 | 0.154330178 | 1 |
| WBGene00005405 | srh-192 | 0.402501625 | 0.487556914 | -0.122708844 | 0.046 | 0.095 | 0.090558424 | 1 |
| WBGene00017676 | F21F3.7 | 0.872457462 | 0.957401994 | -0.122549054 | 0.162 | 0.305 | 0.010527132 | 1 |
| WBGene00009367 | F33H2.3 | 0.533488229 | 0.618137895 | -0.122123653 | 0.069 | 0.179 | 0.004178973 | 1 |
| WBGene00019554 | K09C4.10 | 1.039958411 | 1.124472993 | -0.121928769 | 0.185 | 0.342 | 0.01105365 | 1 |
| WBGene00001663 | gpa-1 | 2.040002596 | 2.124474132 | -0.121866667 | 0.486 | 0.747 | 0.008514382 | 1 |
| WBGene00000193 | arl-6 | 0.259342949 | 0.343710929 | -0.121717267 | 0.046 | 0.105 | 0.052027455 | 1 |
| WBGene00004270 | rab-6.2 | 0.307118417 | 0.391288928 | -0.121432378 | 0.035 | 0.105 | 0.012937517 | 1 |
| WBGene00010990 | tceb-3 | 0.442883162 | 0.527003882 | -0.121360546 | 0.075 | 0.174 | 0.012495342 | 1 |
| WBGene00001047 | dnj-29 | 0.172758247 | 0.256848878 | -0.121317137 | 0.035 | 0.079 | 0.089982002 | 1 |
| WBGene00006450 | tag-77 | 0.113757035 | 0.197677398 | -0.121071491 | 0.023 | 0.053 | 0.15346355 | 1 |
| WBGene00009692 | F44E5.5 | 3.977286821 | 4.061062491 | -0.120862745 | 0.913 | 0.963 | 0.109702126 | 1 |
| WBGene00017830 | rpb-8 | 0.195879546 | 0.279627605 | -0.120822909 | 0.017 | 0.063 | 0.032678494 | 1 |
| WBGene00004148 | pqn-65 | 0.719605693 | 0.803255023 | -0.120680474 | 0.139 | 0.247 | 0.036371352 | 1 |
| WBGene00021966 | Y57G7A.2 | 0.160364307 | 0.243996795 | -0.120656176 | 0.029 | 0.068 | 0.098005558 | 1 |
| WBGene00015303 | rga-6 | 2.004972382 | 2.088373096 | -0.120321796 | 0.486 | 0.705 | 0.009917214 | 1 |
| WBGene00008404 | D2013.6 | 0.218336445 | 0.301606614 | -0.120133459 | 0.035 | 0.058 | 0.32605644 | 1 |
| WBGene00000381 | cct-6 | 0.729564915 | 0.812799833 | -0.120082603 | 0.133 | 0.2 | 0.146828123 | 1 |
| WBGene00009103 | rtfo-1 | 0.240044634 | 0.323187951 | -0.119950451 | 0.035 | 0.089 | 0.042333027 | 1 |
| WBGene00010890 | ddb-1 | 0.198673633 | 0.281815387 | -0.119948196 | 0.029 | 0.058 | 0.194730997 | 1 |
| WBGene00005251 | srh-28 | 0.794266762 | 0.87728081 | -0.119763954 | 0.179 | 0.247 | 0.226777511 | 1 |
| WBGene00005354 | srh-137 | 0.241583205 | 0.32436371 | -0.119427024 | 0.04 | 0.074 | 0.199360183 | 1 |
| WBGene00016342 | C33C12.10 | 0.450771818 | 0.533402875 | -0.119211417 | 0.069 | 0.142 | 0.040054785 | 1 |
| WBGene00022053 | cisd-3.2 | 0.337989059 | 0.420453472 | -0.118970999 | 0.069 | 0.132 | 0.085694724 | 1 |
| WBGene00011423 | ipla-7 | 0.612655343 | 0.695024732 | -0.118833909 | 0.116 | 0.2 | 0.068674604 | 1 |
| WBGene00004454 | rpl-36.A | 2.242845587 | 2.325166796 | -0.118764401 | 0.647 | 0.837 | 0.124960659 | 1 |
| WBGene00050944 | C14A11.9 | 0.436539046 | 0.518582113 | -0.118363126 | 0.075 | 0.153 | 0.038797338 | 1 |
| WBGene00018735 | F53B1.2 | 0.151873314 | 0.233490524 | -0.117748744 | 0.029 | 0.053 | 0.274210726 | 1 |
| WBGene00011283 | ttbk-7 | 0.229962418 | 0.3114247 | -0.117525231 | 0.029 | 0.074 | 0.065610102 | 1 |
| WBGene00009900 | F49E12.7 | 1.910830871 | 1.992229683 | -0.117433662 | 0.445 | 0.653 | 0.018293615 | 1 |
| WBGene00005655 | srr-4 | 1.516881 | 1.598082183 | -0.117148545 | 0.341 | 0.542 | 0.030368863 | 1 |
| WBGene00007344 | C05E7.2 | 0.149586399 | 0.230582779 | -0.116853075 | 0.023 | 0.063 | 0.073084316 | 1 |
| WBGene00014055 | ZK669.5 | 1.733582295 | 1.814570784 | -0.11684169 | 0.416 | 0.658 | 0.016044647 | 1 |
| WBGene00004744 | scp-1 | 0.82802102 | 0.908943667 | -0.116746703 | 0.058 | 0.116 | 0.062552293 | 1 |
| WBGene00012002 | T24H10.4 | 1.380951547 | 1.461852419 | -0.116715287 | 0.324 | 0.474 | 0.084718524 | 1 |
| WBGene00021059 | atat-2 | 0.174965496 | 0.255720033 | -0.116504169 | 0.023 | 0.074 | 0.032641017 | 1 |
| WBGene00012245 | rbm-25 | 0.562520373 | 0.643238582 | -0.11645176 | 0.098 | 0.189 | 0.035061698 | 1 |
| WBGene00010732 | K10C3.5 | 0.152757036 | 0.233458993 | -0.116428314 | 0.029 | 0.063 | 0.139246339 | 1 |
| WBGene00002065 | iff-2 | 1.767323332 | 1.847892211 | -0.116236322 | 0.439 | 0.642 | 0.021455059 | 1 |
| WBGene00013200 | Y54E5A.5 | 0.17021236 | 0.250593179 | -0.115965009 | 0.029 | 0.058 | 0.191839191 | 1 |
| WBGene00018837 | F54G2.1 | 0.571299785 | 0.651490277 | -0.115690425 | 0.116 | 0.142 | 0.600553988 | 1 |
| WBGene00008446 | E01G4.3 | 1.007958563 | 1.087932967 | -0.115378675 | 0.173 | 0.305 | 0.01854915 | 1 |
| WBGene00015716 | sox-4 | 0.361855241 | 0.441799997 | -0.115335903 | 0.046 | 0.1 | 0.063483468 | 1 |
| WBGene00007971 | rpb-3 | 0.189133196 | 0.268999594 | -0.115222856 | 0.029 | 0.063 | 0.137789115 | 1 |
| WBGene00016202 | kle-2 | 0.081734777 | 0.161587419 | -0.115203011 | 0.012 | 0.053 | 0.031666734 | 1 |
| WBGene00016311 | efr-3 | 0.196042665 | 0.275883988 | -0.11518668 | 0.035 | 0.074 | 0.11782539 | 1 |
| WBGene00020131 | gcy-28 | 0.950706369 | 1.03034135 | -0.114888992 | 0.15 | 0.3 | 0.004157665 | 1 |
| WBGene00008255 | C51E3.10 | 0.585246415 | 0.66480978 | -0.114785672 | 0.116 | 0.153 | 0.428067709 | 1 |
| WBGene00235102 | Y54G2A.75 | 0.736994643 | 0.816466173 | -0.114653181 | 0.15 | 0.242 | 0.108304722 | 1 |
| WBGene00005431 | srh-221 | 1.268349158 | 1.34752261 | -0.114223147 | 0.272 | 0.432 | 0.032391289 | 1 |

| | | | | | | | | |
|---|---|---|---|---|---|---|---|---|
| WBGene00006833 | unc-108 | 2.337918934 | 2.417024683 | -0.114125472 | 0.694 | 0.821 | 0.146318655 | 1 |
| WBGene00020779 | T24G10.2 | 0.15101198 | 0.229770067 | -0.113623901 | 0.029 | 0.058 | 0.196676554 | 1 |
| WBGene00012208 | W02D9.2 | 0.297278964 | 0.376003003 | -0.113574781 | 0.046 | 0.111 | 0.036265079 | 1 |
| WBGene00010054 | gtf-2E2 | 0.563489931 | 0.642172826 | -0.113515422 | 0.127 | 0.205 | 0.136214184 | 1 |
| WBGene00206532 | F11A6.15 | 0.727929455 | 0.806486444 | -0.113333778 | 0.139 | 0.226 | 0.080844176 | 1 |
| WBGene00021691 | Y48G8AL.13 | 0.137102021 | 0.215617715 | -0.113274203 | 0.023 | 0.063 | 0.072187223 | 1 |
| WBGene00003600 | nhr-1 | 0.156009611 | 0.234438 | -0.113148248 | 0.029 | 0.063 | 0.135625358 | 1 |
| WBGene00007555 | dohh-1 | 0.174616777 | 0.253027277 | -0.11312244 | 0.023 | 0.063 | 0.072634636 | 1 |
| WBGene00002018 | hsp-16.41 | 3.51467829 | 3.592881592 | -0.112823517 | 0.538 | 0.695 | 0.009966742 | 1 |
| WBGene00022240 | shl-1 | 1.030799558 | 1.108983699 | -0.112795872 | 0.202 | 0.337 | 0.040515653 | 1 |
| WBGene00005648 | srp-7 | 0.397136112 | 0.475296908 | -0.112762193 | 0.058 | 0.105 | 0.129693571 | 1 |
| WBGene00010006 | F53H2.3 | 0.604901761 | 0.682728377 | -0.112280072 | 0.127 | 0.195 | 0.170083605 | 1 |
| WBGene00008576 | F08G2.4 | 0.830830477 | 0.908536647 | -0.112106307 | 0.145 | 0.263 | 0.023485271 | 1 |
| WBGene00005118 | srd-40 | 0.228480509 | 0.306118479 | -0.112007915 | 0.04 | 0.079 | 0.152025526 | 1 |
| WBGene00020866 | vps-24 | 0.292506835 | 0.370103457 | -0.111948262 | 0.046 | 0.089 | 0.1241423 | 1 |
| WBGene00021938 | Y55F3BR.1 | 0.11022385 | 0.187790583 | -0.111905141 | 0.017 | 0.053 | 0.078016491 | 1 |
| WBGene00003950 | pbs-4 | 0.695142696 | 0.772568616 | -0.111701991 | 0.133 | 0.258 | 0.018156816 | 1 |
| WBGene00018349 | F42C5.9 | 0.76583781 | 0.843221771 | -0.111641457 | 0.133 | 0.2 | 0.18070937 | 1 |
| WBGene00019322 | ahcy-1 | 0.962431005 | 1.039579249 | -0.111301389 | 0.173 | 0.305 | 0.021044711 | 1 |
| WBGene00011270 | R31.2 | 0.421194886 | 0.498252489 | -0.111170622 | 0.058 | 0.158 | 0.005512103 | 1 |
| WBGene00021820 | nipa-1 | 0.58902378 | 0.665857025 | -0.110846942 | 0.11 | 0.189 | 0.078356477 | 1 |
| WBGene00010138 | anoh-1 | 0.723446658 | 0.800160609 | -0.110674837 | 0.121 | 0.211 | 0.053724206 | 1 |
| WBGene00003087 | lsy-2 | 0.222256361 | 0.298746775 | -0.110352341 | 0.035 | 0.084 | 0.060050715 | 1 |
| WBGene00003081 | lsm-7 | 0.309220057 | 0.385617466 | -0.110218163 | 0.046 | 0.116 | 0.025300177 | 1 |
| WBGene00009144 | del-3 | 0.934914172 | 1.01077358 | -0.109441991 | 0.202 | 0.326 | 0.046452007 | 1 |
| WBGene00007193 | B0491.6 | 0.573948721 | 0.649748831 | -0.109356443 | 0.11 | 0.179 | 0.108048129 | 1 |
| WBGene00014218 | ZK1098.1 | 0.300359131 | 0.376104955 | -0.109278125 | 0.052 | 0.105 | 0.086581995 | 1 |
| WBGene00018700 | lmd-3 | 0.335554794 | 0.411234734 | -0.109183074 | 0.058 | 0.1 | 0.169457132 | 1 |
| WBGene00012779 | cdkl-1 | 0.717260748 | 0.79293951 | -0.109181375 | 0.11 | 0.242 | 0.00442577 | 1 |
| WBGene00019240 | H24K24.3 | 0.155861931 | 0.23148109 | -0.109095386 | 0.023 | 0.058 | 0.107003289 | 1 |
| WBGene00010143 | F56A12.2 | 0.31694155 | 0.392367804 | -0.108817083 | 0.064 | 0.074 | 0.760703071 | 1 |
| WBGene00023067 | rpl-41.1 | 0.419496712 | 0.494725211 | -0.108531782 | 0.081 | 0.137 | 0.134420299 | 1 |
| WBGene00021461 | nekl-1 | 0.680911285 | 0.75605179 | -0.108404834 | 0.104 | 0.247 | 0.002045183 | 1 |
| WBGene00001665 | gpa-3 | 2.246920306 | 2.32202635 | -0.108355118 | 0.642 | 0.821 | 0.082445555 | 1 |
| WBGene00004344 | rgs-1 | 1.072154334 | 1.147218001 | -0.108293981 | 0.168 | 0.374 | 0.000331303 | 1 |
| WBGene00021685 | herc-1 | 0.766665993 | 0.841667807 | -0.108204745 | 0.15 | 0.237 | 0.11442716 | 1 |
| WBGene00013732 | aakg-1 | 0.254234992 | 0.328835021 | -0.107625093 | 0.035 | 0.068 | 0.16348111 | 1 |
| WBGene00003172 | mec-8 | 1.268955588 | 1.343505024 | -0.107552102 | 0.272 | 0.4 | 0.1070245 | 1 |
| WBGene00006439 | ant-1.1 | 2.864113706 | 2.938636136 | -0.10751314 | 0.798 | 0.926 | 0.033860261 | 1 |
| WBGene00000263 | F23H11.5 | 1.351384593 | 1.425868354 | -0.107457352 | 0.329 | 0.511 | 0.036491345 | 1 |
| WBGene00009580 | xbx-6 | 2.565806837 | 2.640246507 | -0.107393742 | 0.717 | 0.874 | 0.098503045 | 1 |
| WBGene00008224 | slrp-1 | 0.433530185 | 0.50793542 | -0.107344064 | 0.092 | 0.158 | 0.113440414 | 1 |
| WBGene00020854 | T27C4.1 | 2.031417617 | 2.10572827 | -0.107207611 | 0.601 | 0.737 | 0.188245227 | 1 |
| WBGene00019680 | K12H4.5 | 0.448176672 | 0.522349245 | -0.107008404 | 0.081 | 0.158 | 0.052295636 | 1 |
| WBGene00008860 | romo-1 | 0.199800362 | 0.27389431 | -0.10689497 | 0.04 | 0.058 | 0.456813356 | 1 |
| WBGene00022836 | ZK973.11 | 0.530535315 | 0.604608687 | -0.106865286 | 0.098 | 0.168 | 0.099304114 | 1 |
| WBGene00017283 | F09E5.3 | 0.169304638 | 0.243308922 | -0.106765614 | 0.023 | 0.058 | 0.107003289 | 1 |
| WBGene00003060 | lpd-3 | 0.440502003 | 0.514472737 | -0.10671721 | 0.081 | 0.132 | 0.165124417 | 1 |
| WBGene00006514 | tdp-1 | 0.190854808 | 0.264635695 | -0.10644332 | 0.035 | 0.084 | 0.060381247 | 1 |
| WBGene00202280 | Y108G3AL.9 | 0.178871172 | 0.25256341 | -0.106315427 | 0.029 | 0.079 | 0.045818104 | 1 |
| WBGene00012037 | T26E3.4 | 0.237597374 | 0.311268357 | -0.106284761 | 0.035 | 0.095 | 0.029446786 | 1 |
| WBGene00022144 | pghm-1 | 2.976618029 | 3.050184342 | -0.106133756 | 0.844 | 0.953 | 0.188315659 | 1 |
| WBGene00009952 | acaa-2 | 0.275544581 | 0.349073595 | -0.106079943 | 0.046 | 0.1 | 0.071050004 | 1 |

| | | | | | | | | |
|---|---|---|---|---|---|---|---|---|
| WBGene00014219 | ZK1098.2 | 0.286467259 | 0.359541964 | -0.105424513 | 0.04 | 0.1 | 0.038563888 | 1 |
| WBGene00008765 | ttx-7 | 0.696973044 | 0.769953949 | -0.10528919 | 0.133 | 0.226 | 0.064926334 | 1 |
| WBGene00003026 | lin-41 | 0.169853101 | 0.242735318 | -0.105146814 | 0.023 | 0.058 | 0.102039969 | 1 |
| WBGene00019946 | chat-1 | 0.600405037 | 0.673168277 | -0.104975167 | 0.104 | 0.179 | 0.090257146 | 1 |
| WBGene00004914 | snr-1 | 0.750802255 | 0.823446148 | -0.104802985 | 0.145 | 0.247 | 0.049197901 | 1 |
| WBGene00004462 | rpn-6.1 | 0.394184516 | 0.466747838 | -0.104686744 | 0.075 | 0.158 | 0.033927872 | 1 |
| WBGene00271642 | R106.8 | 0.987602069 | 1.060119738 | -0.104620881 | 0.197 | 0.342 | 0.021851482 | 1 |
| WBGene00005726 | srv-15 | 1.266201896 | 1.338701986 | -0.104595521 | 0.22 | 0.389 | 0.006993066 | 1 |
| WBGene00000160 | apb-1 | 0.377318627 | 0.449803426 | -0.104573459 | 0.064 | 0.1 | 0.249125705 | 1 |
| WBGene00020762 | zer-1 | 0.525762484 | 0.59809124 | -0.104348338 | 0.104 | 0.179 | 0.095530574 | 1 |
| WBGene00013308 | nuo-3 | 1.273753083 | 1.345870067 | -0.104042815 | 0.306 | 0.463 | 0.087802535 | 1 |
| WBGene00019781 | M60.6 | 0.479228025 | 0.551138328 | -0.103744637 | 0.075 | 0.163 | 0.023411982 | 1 |
| WBGene00010061 | F54E12.2 | 0.293132784 | 0.364927186 | -0.103577429 | 0.052 | 0.111 | 0.064792457 | 1 |
| WBGene00005269 | srh-46 | 0.400528471 | 0.472163146 | -0.10334699 | 0.046 | 0.111 | 0.033868419 | 1 |
| WBGene00003962 | pdi-1 | 0.308722764 | 0.380267366 | -0.103217043 | 0.058 | 0.105 | 0.13729263 | 1 |
| WBGene00005386 | srh-170 | 0.202169489 | 0.273683 | -0.103172188 | 0.035 | 0.058 | 0.320695042 | 1 |
| WBGene00044095 | srh-121 | 0.645872293 | 0.717128787 | -0.10280139 | 0.139 | 0.189 | 0.358165227 | 1 |
| WBGene00001946 | his-72 | 2.038379167 | 2.109624129 | -0.102784752 | 0.561 | 0.7 | 0.182495335 | 1 |
| WBGene00002098 | ins-15 | 0.532289331 | 0.603404378 | -0.102597325 | 0.081 | 0.111 | 0.396041525 | 1 |
| WBGene00001145 | eat-16 | 1.4362017 | 1.507315909 | -0.102596117 | 0.324 | 0.474 | 0.113216045 | 1 |
| WBGene00006918 | vha-9 | 0.948596388 | 1.019683204 | -0.102556596 | 0.22 | 0.316 | 0.210409441 | 1 |
| WBGene00016238 | mob-4 | 0.121375825 | 0.192418805 | -0.102493355 | 0.023 | 0.053 | 0.161397566 | 1 |
| WBGene00022122 | trap-1 | 0.326426167 | 0.397402655 | -0.102397427 | 0.064 | 0.105 | 0.197664081 | 1 |
| WBGene00013577 | Y76A2B.5 | 0.904503168 | 0.975454636 | -0.102361331 | 0.162 | 0.289 | 0.02124243 | 1 |
| WBGene00045024 | C43F9.11 | 0.151778393 | 0.222660117 | -0.102260712 | 0.017 | 0.053 | 0.078016491 | 1 |
| WBGene00019915 | tbc-10 | 0.344255222 | 0.415013671 | -0.102082865 | 0.046 | 0.095 | 0.092278607 | 1 |
| WBGene00006386 | taf-5 | 0.238313886 | 0.308883439 | -0.101810344 | 0.029 | 0.079 | 0.041845212 | 1 |
| WBGene00005518 | sri-6 | 0.427978081 | 0.498450916 | -0.101670809 | 0.035 | 0.111 | 0.007933129 | 1 |
| WBGene00013168 | arp-1 | 0.348022714 | 0.418001482 | -0.100958023 | 0.069 | 0.116 | 0.178210044 | 1 |
| WBGene00006812 | unc-80 | 0.302853928 | 0.37275348 | -0.100843736 | 0.04 | 0.084 | 0.103478983 | 1 |
| WBGene00011935 | scrm-1 | 2.131642072 | 2.201499692 | -0.100783242 | 0.607 | 0.789 | 0.108246703 | 1 |
| WBGene00012811 | Y43F8A.5 | 0.279601049 | 0.349378377 | -0.100667405 | 0.052 | 0.089 | 0.191162579 | 1 |
| WBGene00219734 | linc-128 | 0.151824417 | 0.221581484 | -0.100638175 | 0.023 | 0.053 | 0.159608215 | 1 |
| WBGene00014031 | ZK637.14 | 0.778515698 | 0.848216392 | -0.100556846 | 0.156 | 0.242 | 0.117587779 | 1 |
| WBGene00017733 | ubxn-1 | 0.132663578 | 0.202335282 | -0.100515023 | 0.023 | 0.053 | 0.160501006 | 1 |
| WBGene00009200 | F28C1.1 | 0.140641331 | 0.210216834 | -0.100376233 | 0.023 | 0.058 | 0.107636931 | 1 |
| WBGene00235283 | Y55F3BL.6 | 2.333071142 | 2.263264743 | 0.100709345 | 0.613 | 0.737 | 0.847726524 | 1 |
| WBGene00007027 | ssl-1 | 0.471132929 | 0.401261354 | 0.100803375 | 0.064 | 0.105 | 0.21624235 | 1 |
| WBGene00004504 | rpt-4 | 0.857735486 | 0.787829093 | 0.100853606 | 0.15 | 0.226 | 0.170946077 | 1 |
| WBGene00008434 | D2089.3 | 1.760144389 | 1.69023536 | 0.10085741 | 0.364 | 0.505 | 0.080841651 | 1 |
| WBGene00000368 | ccb-1 | 0.415947612 | 0.346034735 | 0.10086296 | 0.087 | 0.095 | 0.939969075 | 1 |
| WBGene00003791 | npp-5 | 0.384218611 | 0.314252258 | 0.100940112 | 0.046 | 0.068 | 0.414038914 | 1 |
| WBGene00004459 | rpn-2 | 0.613108134 | 0.542930444 | 0.101245005 | 0.139 | 0.137 | 0.76923808 | 1 |
| WBGene00004389 | rnp-6 | 0.376741087 | 0.306548524 | 0.101266461 | 0.075 | 0.089 | 0.763441217 | 1 |
| WBGene00268194 | F36H9.10 | 0.24880978 | 0.1785561 | 0.101354636 | 0.035 | 0.053 | 0.442887915 | 1 |
| WBGene00018376 | F43C9.2 | 0.396159654 | 0.325887863 | 0.101380765 | 0.058 | 0.047 | 0.642299408 | 1 |
| WBGene00019971 | ergo-1 | 0.25857793 | 0.188295734 | 0.101395775 | 0.04 | 0.053 | 0.646796835 | 1 |
| WBGene00001668 | gpa-6 | 0.818502172 | 0.748146351 | 0.101501995 | 0.156 | 0.247 | 0.12628758 | 1 |
| WBGene00010075 | F55A11.1 | 0.462715182 | 0.392324045 | 0.101552945 | 0.087 | 0.111 | 0.594742089 | 1 |
| WBGene00010110 | F55D12.1 | 0.494476409 | 0.424084761 | 0.101553683 | 0.092 | 0.121 | 0.536203987 | 1 |
| WBGene00005558 | sri-46 | 0.395001565 | 0.324585605 | 0.101588756 | 0.058 | 0.089 | 0.319717704 | 1 |
| WBGene00005368 | srh-152 | 0.297880542 | 0.227423822 | 0.101647561 | 0.058 | 0.058 | 0.950658957 | 1 |
| WBGene00012193 | vps-25 | 1.522178509 | 1.451649596 | 0.101751712 | 0.382 | 0.526 | 0.338787504 | 1 |

| | | | | | | | | |
|---|---|---|---|---|---|---|---|---|
| WBGene00018782 | cct-3 | 0.294766361 | 0.224019482 | 0.102066171 | 0.052 | 0.063 | 0.709510674 | 1 |
| WBGene00019955 | R08C7.12 | 0.447011218 | 0.376127032 | 0.102264264 | 0.058 | 0.089 | 0.308082542 | 1 |
| WBGene00007675 | C18D4.6 | 1.438391054 | 1.367426686 | 0.102379942 | 0.266 | 0.395 | 0.115747106 | 1 |
| WBGene00006757 | unc-18 | 0.297332857 | 0.226211256 | 0.102606781 | 0.046 | 0.053 | 0.842779777 | 1 |
| WBGene00007091 | eri-12 | 1.032514122 | 0.961322013 | 0.102708502 | 0.197 | 0.337 | 0.04412794 | 1 |
| WBGene00015920 | eif-3.L | 0.312484506 | 0.241254939 | 0.102762544 | 0.052 | 0.068 | 0.600076873 | 1 |
| WBGene00004884 | smg-6 | 0.327306892 | 0.256011618 | 0.102857339 | 0.052 | 0.074 | 0.470170063 | 1 |
| WBGene00003009 | lin-23 | 0.385024286 | 0.313728637 | 0.102857878 | 0.064 | 0.084 | 0.552993448 | 1 |
| WBGene00020144 | T01C8.3 | 2.395217354 | 2.323859179 | 0.102948085 | 0.671 | 0.774 | 0.872550712 | 1 |
| WBGene00005557 | sri-45 | 1.991053888 | 1.919462709 | 0.103284238 | 0.491 | 0.642 | 0.364948595 | 1 |
| WBGene00021722 | srz-5 | 0.675454825 | 0.603590407 | 0.10367844 | 0.092 | 0.153 | 0.149432486 | 1 |
| WBGene00000971 | dhs-7 | 0.417742054 | 0.345790904 | 0.103803569 | 0.064 | 0.089 | 0.435149565 | 1 |
| WBGene00012132 | ifta-2 | 1.738626679 | 1.666602906 | 0.10390834 | 0.462 | 0.605 | 0.318970274 | 1 |
| WBGene00018795 | F54C4.4 | 0.247513115 | 0.175475893 | 0.103927742 | 0.035 | 0.053 | 0.456393439 | 1 |
| WBGene00020936 | hrpf-1 | 0.304988135 | 0.232896793 | 0.104005822 | 0.058 | 0.068 | 0.779578592 | 1 |
| WBGene00009671 | mfap-1 | 0.291100711 | 0.218968215 | 0.104065195 | 0.035 | 0.068 | 0.181428021 | 1 |
| WBGene00019287 | K01A12.3 | 0.297537239 | 0.225220215 | 0.104331412 | 0.04 | 0.063 | 0.383979501 | 1 |
| WBGene00003599 | nhl-3 | 0.67092907 | 0.598597139 | 0.104352918 | 0.092 | 0.168 | 0.070151967 | 1 |
| WBGene00004117 | sin-3 | 0.40172816 | 0.328709681 | 0.105343398 | 0.052 | 0.068 | 0.57588584 | 1 |
| WBGene00004760 | sel-2 | 0.65438176 | 0.581362312 | 0.105344795 | 0.081 | 0.137 | 0.145291816 | 1 |
| WBGene00007531 | C11H1.7 | 0.393644116 | 0.320585914 | 0.105400706 | 0.052 | 0.079 | 0.341032543 | 1 |
| WBGene00022391 | prhg-1 | 0.934577915 | 0.861498704 | 0.105431016 | 0.185 | 0.263 | 0.282659432 | 1 |
| WBGene00023282 | F39E9.9 | 1.108250279 | 1.035073542 | 0.105571715 | 0.208 | 0.3 | 0.206809021 | 1 |
| WBGene00000241 | bbs-1 | 0.545138797 | 0.47175379 | 0.105872185 | 0.092 | 0.137 | 0.30026309 | 1 |
| WBGene00001226 | eif-3.C | 0.478132294 | 0.404488743 | 0.106245185 | 0.081 | 0.105 | 0.524134714 | 1 |
| WBGene00019007 | ucr-11 | 0.266298418 | 0.192413033 | 0.106594078 | 0.046 | 0.053 | 0.844863135 | 1 |
| WBGene00009880 | F49C12.11 | 0.589732737 | 0.515776207 | 0.106696719 | 0.116 | 0.147 | 0.56012136 | 1 |
| WBGene00018161 | F38A5.2 | 0.339798382 | 0.265832298 | 0.106710503 | 0.064 | 0.068 | 0.932367901 | 1 |
| WBGene00015996 | C18H7.6 | 2.900270701 | 2.826132561 | 0.106958726 | 0.85 | 0.905 | 0.640067327 | 1 |
| WBGene00271778 | T01A4.9 | 3.322063372 | 3.247901548 | 0.106992897 | 0.908 | 0.937 | 0.371578386 | 1 |
| WBGene00016121 | ift-43 | 1.336171668 | 1.262006434 | 0.106997816 | 0.289 | 0.437 | 0.135834752 | 1 |
| WBGene00021078 | unc-132 | 0.681750691 | 0.607552336 | 0.107045599 | 0.098 | 0.174 | 0.08586899 | 1 |
| WBGene00011605 | sftb-1 | 0.346853326 | 0.272550248 | 0.107196682 | 0.046 | 0.084 | 0.186228952 | 1 |
| WBGene00006843 | unc-119 | 1.374962295 | 1.300571794 | 0.107322807 | 0.341 | 0.432 | 0.617445773 | 1 |
| WBGene00014214 | ZK1073.2 | 0.735620794 | 0.660865299 | 0.107849382 | 0.145 | 0.168 | 0.738005619 | 1 |
| WBGene00009134 | lurp-2 | 1.18606823 | 1.111188622 | 0.108028439 | 0.249 | 0.368 | 0.151039475 | 1 |
| WBGene00007191 | lgc-20 | 1.492356521 | 1.417197266 | 0.108431885 | 0.324 | 0.495 | 0.049639693 | 1 |
| WBGene00016917 | C54E4.1 | 0.515652232 | 0.440367811 | 0.10861246 | 0.064 | 0.1 | 0.267675198 | 1 |
| WBGene00012606 | Y38F1A.2 | 0.31411559 | 0.238724752 | 0.108765988 | 0.046 | 0.058 | 0.683715288 | 1 |
| WBGene00003668 | nhr-78 | 0.656504438 | 0.580909074 | 0.109061056 | 0.116 | 0.174 | 0.247620912 | 1 |
| WBGene00009804 | F47B8.3 | 1.445380435 | 1.369779558 | 0.10906901 | 0.318 | 0.511 | 0.029788767 | 1 |
| WBGene00015817 | C16A11.4 | 1.559806177 | 1.484155494 | 0.109140866 | 0.393 | 0.511 | 0.597537803 | 1 |
| WBGene00010465 | K01D12.6 | 0.348463955 | 0.272731898 | 0.109258262 | 0.046 | 0.074 | 0.323727192 | 1 |
| WBGene00010700 | nipi-3 | 1.322148923 | 1.246388607 | 0.109299033 | 0.266 | 0.384 | 0.157950562 | 1 |
| WBGene00005578 | sri-66 | 0.626325716 | 0.550445419 | 0.109472129 | 0.092 | 0.147 | 0.206844802 | 1 |
| WBGene00014261 | shk-1 | 1.719120158 | 1.642896879 | 0.109966947 | 0.382 | 0.511 | 0.395646602 | 1 |
| WBGene00000961 | dgn-1 | 0.570558603 | 0.494280927 | 0.110045425 | 0.092 | 0.121 | 0.496474597 | 1 |
| WBGene00010724 | K09E9.1 | 0.301826302 | 0.225481156 | 0.110142763 | 0.04 | 0.058 | 0.498058581 | 1 |
| WBGene00044728 | Y53F4B.45 | 1.829417774 | 1.752977051 | 0.110280652 | 0.358 | 0.463 | 0.324402255 | 1 |
| WBGene00020424 | T10H9.1 | 0.442452686 | 0.365986461 | 0.110317444 | 0.058 | 0.079 | 0.4865504 | 1 |
| WBGene00011684 | T10B10.8 | 0.281412853 | 0.204919871 | 0.110356046 | 0.052 | 0.037 | 0.444563506 | 1 |
| WBGene00008368 | D1053.3 | 0.977826689 | 0.901299406 | 0.110405532 | 0.191 | 0.274 | 0.256167132 | 1 |
| WBGene00011375 | T02E1.2 | 0.435237149 | 0.358683156 | 0.110444065 | 0.069 | 0.105 | 0.327813054 | 1 |

| | | | | | | | | |
|---|---|---|---|---|---|---|---|---|
| WBGene00004740 | scd-2 | 1.873251969 | 1.796691878 | 0.110452864 | 0.399 | 0.584 | 0.105910791 | 1 |
| WBGene00021075 | sms-5 | 0.329898426 | 0.253190155 | 0.110666642 | 0.046 | 0.053 | 0.846947579 | 1 |
| WBGene00009803 | F47B8.2 | 2.777609933 | 2.700524948 | 0.111210126 | 0.728 | 0.816 | 0.84861744 | 1 |
| WBGene00015074 | natc-2 | 0.565080945 | 0.487946956 | 0.111280823 | 0.092 | 0.126 | 0.456136245 | 1 |
| WBGene00002235 | kqt-3 | 0.510340355 | 0.433087048 | 0.111452963 | 0.087 | 0.121 | 0.40852683 | 1 |
| WBGene00002879 | let-754 | 0.289528184 | 0.212174913 | 0.111597181 | 0.046 | 0.053 | 0.842779777 | 1 |
| WBGene00004728 | sax-2 | 0.446847982 | 0.369412865 | 0.111715259 | 0.064 | 0.1 | 0.264070791 | 1 |
| WBGene00001093 | drp-1 | 0.381809473 | 0.304323975 | 0.111787943 | 0.058 | 0.063 | 0.89691538 | 1 |
| WBGene00012625 | Y38H6C.13 | 0.366060249 | 0.28856059 | 0.111808374 | 0.058 | 0.074 | 0.616330077 | 1 |
| WBGene00001122 | dyf-6 | 1.868349349 | 1.790842282 | 0.111819061 | 0.451 | 0.616 | 0.17904015 | 1 |
| WBGene00011629 | T08G11.1 | 0.892664263 | 0.815076879 | 0.111934933 | 0.145 | 0.232 | 0.107761576 | 1 |
| WBGene00015495 | C05E11.6 | 2.32791861 | 2.250021597 | 0.112381635 | 0.665 | 0.753 | 0.590344336 | 1 |
| WBGene00194710 | mks-2 | 2.148844359 | 2.070771956 | 0.112634668 | 0.613 | 0.737 | 0.850919484 | 1 |
| WBGene00006143 | str-84 | 1.082673645 | 1.004560216 | 0.112693856 | 0.214 | 0.321 | 0.133022753 | 1 |
| WBGene00018615 | F48G7.4 | 0.435478179 | 0.357324543 | 0.112751864 | 0.075 | 0.079 | 0.964602806 | 1 |
| WBGene00008664 | nubp-1 | 0.44823847 | 0.370038507 | 0.112818699 | 0.075 | 0.1 | 0.511332266 | 1 |
| WBGene00011983 | fbxa-81 | 0.33068269 | 0.252219175 | 0.113198924 | 0.064 | 0.068 | 0.965690145 | 1 |
| WBGene00018635 | nprl-2 | 0.322456743 | 0.243986821 | 0.113208167 | 0.04 | 0.058 | 0.504836767 | 1 |
| WBGene00018265 | nhr-182 | 0.287848237 | 0.209313839 | 0.113301188 | 0.046 | 0.058 | 0.683715288 | 1 |
| WBGene00194985 | Y25C1A.14 | 0.394564818 | 0.315917862 | 0.113463574 | 0.058 | 0.089 | 0.313336075 | 1 |
| WBGene00021922 | atg-3 | 0.321983136 | 0.243047353 | 0.113880263 | 0.046 | 0.063 | 0.532417249 | 1 |
| WBGene00019230 | ttll-11 | 0.9255592 | 0.846566875 | 0.113961836 | 0.173 | 0.253 | 0.195273029 | 1 |
| WBGene00020146 | got-1.2 | 0.437797264 | 0.358488896 | 0.11441779 | 0.075 | 0.089 | 0.728511742 | 1 |
| WBGene00010959 | nduo-1 | 1.746336381 | 1.666661719 | 0.11494624 | 0.382 | 0.579 | 0.045971662 | 1 |
| WBGene00009112 | tag-353 | 0.400194248 | 0.320272951 | 0.115302058 | 0.064 | 0.084 | 0.551521388 | 1 |
| WBGene00010419 | atp-1 | 1.552436324 | 1.472505735 | 0.115315464 | 0.387 | 0.553 | 0.263032809 | 1 |
| WBGene00005372 | srh-156 | 0.657571884 | 0.577584729 | 0.115397072 | 0.092 | 0.153 | 0.144075063 | 1 |
| WBGene00007433 | swsn-7 | 0.707062971 | 0.62670075 | 0.115938178 | 0.133 | 0.184 | 0.38309828 | 1 |
| WBGene00008624 | crld-1 | 0.295631921 | 0.215198498 | 0.116040901 | 0.046 | 0.063 | 0.545804821 | 1 |
| WBGene00001082 | dpy-23 | 0.641065131 | 0.560565561 | 0.11613633 | 0.11 | 0.147 | 0.475658173 | 1 |
| WBGene00012156 | ebp-2 | 0.256956986 | 0.176456339 | 0.116137884 | 0.04 | 0.053 | 0.650732115 | 1 |
| WBGene00018898 | F55F10.1 | 0.416335015 | 0.33578465 | 0.116209612 | 0.058 | 0.089 | 0.313336075 | 1 |
| WBGene00007534 | fubl-1 | 1.239007699 | 1.158372636 | 0.116331804 | 0.272 | 0.347 | 0.51773353 | 1 |
| WBGene00015734 | copd-1 | 0.452898767 | 0.37221683 | 0.11639943 | 0.081 | 0.111 | 0.460378697 | 1 |
| WBGene00022635 | ZC581.9 | 1.758863472 | 1.678129669 | 0.116474258 | 0.422 | 0.542 | 0.57919711 | 1 |
| WBGene00016400 | C34D4.4 | 0.313008925 | 0.232236323 | 0.116530233 | 0.052 | 0.074 | 0.481913512 | 1 |
| WBGene00018545 | nhr-189 | 0.482520374 | 0.401633031 | 0.116695769 | 0.092 | 0.121 | 0.555401946 | 1 |
| WBGene00008066 | fbxb-65 | 0.725848889 | 0.644933414 | 0.116736355 | 0.15 | 0.147 | 0.714036358 | 1 |
| WBGene00005411 | srh-200 | 0.939739912 | 0.858661005 | 0.116972137 | 0.139 | 0.237 | 0.067476667 | 1 |
| WBGene00001169 | eef-1A.2 | 2.460700189 | 2.379608052 | 0.116991224 | 0.728 | 0.842 | 0.919845101 | 1 |
| WBGene00018159 | F37F2.2 | 0.414289006 | 0.333167026 | 0.117034278 | 0.064 | 0.084 | 0.542729684 | 1 |
| WBGene00008438 | DH11.5 | 1.53680167 | 1.45524085 | 0.11766739 | 0.37 | 0.495 | 0.500939907 | 1 |
| WBGene00003225 | mev-1 | 0.645611836 | 0.563816616 | 0.118005558 | 0.127 | 0.158 | 0.610900683 | 1 |
| WBGene00023465 | srz-81 | 1.074689296 | 0.992745906 | 0.118219322 | 0.243 | 0.305 | 0.536804478 | 1 |
| WBGene00011474 | aldo-1 | 1.779321035 | 1.697275783 | 0.118366278 | 0.474 | 0.595 | 0.890481243 | 1 |
| WBGene00022354 | Y82E9BR.22 | 0.315046601 | 0.232770841 | 0.118698832 | 0.029 | 0.053 | 0.275487275 | 1 |
| WBGene00020363 | fbxa-3 | 0.300274091 | 0.217712276 | 0.119111521 | 0.04 | 0.058 | 0.503137627 | 1 |
| WBGene00021883 | Y54G2A.18 | 0.853702545 | 0.771102939 | 0.119166042 | 0.139 | 0.2 | 0.226077632 | 1 |
| WBGene00014058 | ZK673.2 | 0.280501734 | 0.197691132 | 0.119470446 | 0.046 | 0.053 | 0.832379764 | 1 |
| WBGene00013039 | scav-3 | 0.484800988 | 0.401705895 | 0.119880879 | 0.081 | 0.111 | 0.437538558 | 1 |
| WBGene00022774 | swsn-6 | 0.290748547 | 0.20762451 | 0.119922637 | 0.052 | 0.058 | 0.890213303 | 1 |
| WBGene00018972 | srz-19 | 2.043159438 | 1.960001409 | 0.119971676 | 0.462 | 0.568 | 0.685927025 | 1 |
| WBGene00022842 | ZK994.6 | 0.687575458 | 0.604362289 | 0.120051226 | 0.15 | 0.174 | 0.873653302 | 1 |

| | | | | | | | | |
|---|---|---|---|---|---|---|---|---|
| WBGene00005401 | srh-186 | 0.334572877 | 0.251343587 | 0.120074484 | 0.064 | 0.063 | 0.919709224 | 1 |
| WBGene00006000 | srx-109 | 0.419813042 | 0.336570332 | 0.120093845 | 0.058 | 0.068 | 0.732701517 | 1 |
| WBGene00015162 | algn-11 | 0.44044338 | 0.357190397 | 0.120108665 | 0.064 | 0.095 | 0.354678576 | 1 |
| WBGene00022583 | mrps-18A | 0.405226705 | 0.321969636 | 0.12011456 | 0.058 | 0.1 | 0.190901999 | 1 |
| WBGene00019465 | acl-14 | 0.707078596 | 0.623678245 | 0.120321272 | 0.11 | 0.168 | 0.205677962 | 1 |
| WBGene00005331 | srh-112 | 0.768551969 | 0.685019691 | 0.120511604 | 0.145 | 0.195 | 0.367070398 | 1 |
| WBGene00005379 | srh-163 | 0.263098328 | 0.179541009 | 0.12054773 | 0.04 | 0.058 | 0.509952465 | 1 |
| WBGene00018974 | fcho-1 | 1.115683442 | 1.03204639 | 0.120662759 | 0.208 | 0.295 | 0.233268893 | 1 |
| WBGene00012903 | vps-2 | 0.307513014 | 0.223629974 | 0.121017647 | 0.052 | 0.047 | 0.786990731 | 1 |
| WBGene00269427 | D1065.7 | 1.138052277 | 1.053956504 | 0.121324554 | 0.26 | 0.305 | 0.95180419 | 1 |
| WBGene00022059 | Y67D8A.2 | 0.404178441 | 0.319954886 | 0.121508906 | 0.046 | 0.095 | 0.09988416 | 1 |
| WBGene00022745 | ZK470.2 | 1.905967537 | 1.821727274 | 0.121533009 | 0.468 | 0.611 | 0.532309496 | 1 |
| WBGene00023515 | srz-97 | 0.682170998 | 0.597922868 | 0.12154436 | 0.098 | 0.158 | 0.172584706 | 1 |
| WBGene00012005 | 1-Jun | 1.089335729 | 1.004944699 | 0.12175052 | 0.202 | 0.363 | 0.026326025 | 1 |
| WBGene00003746 | nlp-8 | 2.874033324 | 2.789620761 | 0.121781586 | 0.861 | 0.932 | 0.534769028 | 1 |
| WBGene00001491 | frm-4 | 0.747261507 | 0.662817706 | 0.121826653 | 0.133 | 0.2 | 0.212650003 | 1 |
| WBGene00005299 | srh-78 | 1.553212793 | 1.468647492 | 0.122001941 | 0.341 | 0.468 | 0.229272002 | 1 |
| WBGene00018750 | F53C3.6 | 0.455091713 | 0.370162625 | 0.122526773 | 0.058 | 0.089 | 0.298773856 | 1 |
| WBGene00011644 | T09B9.3 | 4.585643441 | 4.50060929 | 0.122678347 | 1 | 0.995 | 0.167730744 | 1 |
| WBGene00016653 | ssb-1 | 0.272307616 | 0.187244419 | 0.122720252 | 0.046 | 0.058 | 0.69902319 | 1 |
| WBGene00019185 | H10E21.5 | 1.707471621 | 1.622408091 | 0.122720732 | 0.399 | 0.547 | 0.378429642 | 1 |
| WBGene00001309 | emr-1 | 0.317863305 | 0.232696607 | 0.122869572 | 0.064 | 0.063 | 0.89152367 | 1 |
| WBGene00017769 | hmgs-1 | 0.305198698 | 0.21996948 | 0.122959769 | 0.052 | 0.053 | 0.96171181 | 1 |
| WBGene00195073 | F02E11.7 | 1.370217192 | 1.284954405 | 0.123008201 | 0.347 | 0.432 | 0.788685109 | 1 |
| WBGene00003733 | nhx-5 | 0.689586795 | 0.604258769 | 0.12310232 | 0.121 | 0.189 | 0.191164376 | 1 |
| WBGene00015237 | B0511.12 | 0.43822201 | 0.352675953 | 0.123416873 | 0.052 | 0.089 | 0.215826611 | 1 |
| WBGene00004756 | sec-24.2 | 0.272063014 | 0.186488949 | 0.123457278 | 0.012 | 0.053 | 0.031666734 | 1 |
| WBGene00017385 | F11G11.5 | 0.62070415 | 0.535066478 | 0.123549044 | 0.104 | 0.153 | 0.295795872 | 1 |
| WBGene00015206 | B0495.7 | 0.727289336 | 0.641576285 | 0.123657794 | 0.104 | 0.205 | 0.029422435 | 1 |
| WBGene00000166 | apt-9 | 0.298407472 | 0.212402733 | 0.124078612 | 0.058 | 0.053 | 0.766010839 | 1 |
| WBGene00015355 | C02F12.5 | 0.430636904 | 0.344424143 | 0.124378723 | 0.052 | 0.068 | 0.573409982 | 1 |
| WBGene00009851 | F48F7.6 | 0.281048357 | 0.194822488 | 0.124397633 | 0.04 | 0.053 | 0.648763249 | 1 |
| WBGene00016067 | C24H10.1 | 0.617664481 | 0.531408955 | 0.12444042 | 0.098 | 0.137 | 0.392170504 | 1 |
| WBGene00006669 | twk-14 | 1.844418583 | 1.758104127 | 0.124525437 | 0.399 | 0.568 | 0.150376659 | 1 |
| WBGene00003725 | nhr-135 | 0.319452186 | 0.233135045 | 0.124529311 | 0.058 | 0.074 | 0.651102366 | 1 |
| WBGene00022648 | ZK54.3 | 1.950628894 | 1.86428419 | 0.124569076 | 0.289 | 0.368 | 0.25523508 | 1 |
| WBGene00000832 | ctn-1 | 0.412847157 | 0.325777936 | 0.125614334 | 0.058 | 0.105 | 0.145230558 | 1 |
| WBGene00011966 | T23G7.3 | 0.355506774 | 0.268202664 | 0.125953206 | 0.069 | 0.079 | 0.827476036 | 1 |
| WBGene00010896 | M28.5 | 0.320568943 | 0.233224695 | 0.126011113 | 0.064 | 0.053 | 0.60753298 | 1 |
| WBGene00022717 | hipr-1 | 0.297855468 | 0.210502415 | 0.126023816 | 0.017 | 0.063 | 0.031284409 | 1 |
| WBGene00007055 | tag-196 | 0.322452831 | 0.235081196 | 0.126050625 | 0.052 | 0.047 | 0.815801331 | 1 |
| WBGene00012359 | W09D10.1 | 0.713682465 | 0.626234788 | 0.12616033 | 0.121 | 0.184 | 0.227464406 | 1 |
| WBGene00012833 | Y43F8C.11 | 2.390113827 | 2.302522739 | 0.126367229 | 0.659 | 0.784 | 0.633933473 | 1 |
| WBGene00005811 | srw-64 | 0.310560214 | 0.222965981 | 0.126371765 | 0.035 | 0.068 | 0.179738428 | 1 |
| WBGene00020628 | T20F5.6 | 0.422838144 | 0.334977139 | 0.126756637 | 0.064 | 0.1 | 0.28905652 | 1 |
| WBGene00005437 | srh-229 | 0.693398778 | 0.605226221 | 0.127206111 | 0.15 | 0.168 | 0.928063189 | 1 |
| WBGene00005234 | srh-8 | 0.498908255 | 0.410577209 | 0.127434762 | 0.092 | 0.111 | 0.732067435 | 1 |
| WBGene00008729 | F13B12.1 | 0.345803564 | 0.25743192 | 0.127493333 | 0.052 | 0.068 | 0.591683319 | 1 |
| WBGene00003555 | nas-39 | 0.846876904 | 0.758462523 | 0.12755499 | 0.179 | 0.221 | 0.684022438 | 1 |
| WBGene00000369 | ccf-1 | 0.385170533 | 0.296677392 | 0.127668615 | 0.058 | 0.074 | 0.622890476 | 1 |
| WBGene00016235 | C29H12.2 | 1.83823574 | 1.749703031 | 0.1277257 | 0.428 | 0.611 | 0.230355396 | 1 |
| WBGene00019686 | K12H6.6 | 0.373237629 | 0.283807195 | 0.129020844 | 0.069 | 0.068 | 0.889322785 | 1 |
| WBGene00004381 | rnf-5 | 0.39057556 | 0.30114366 | 0.129022959 | 0.052 | 0.068 | 0.566832112 | 1 |

| | | | | | | | | |
|---|---|---|---|---|---|---|---|---|
| WBGene00021348 | moag-4 | 0.290686927 | 0.201205006 | 0.129095123 | 0.046 | 0.063 | 0.55935725 | 1 |
| WBGene00006607 | tre-1 | 0.433873882 | 0.344041879 | 0.129600184 | 0.081 | 0.089 | 0.863839109 | 1 |
| WBGene00010647 | K08C7.4 | 0.472930785 | 0.383090772 | 0.12961174 | 0.087 | 0.089 | 0.942367126 | 1 |
| WBGene00008728 | sre-36 | 0.767678716 | 0.677708393 | 0.129799738 | 0.127 | 0.2 | 0.170874896 | 1 |
| WBGene00013751 | Y113G7A.14 | 1.863346775 | 1.773300373 | 0.129909499 | 0.457 | 0.626 | 0.323966835 | 1 |
| WBGene00017120 | tctn-1 | 0.787166333 | 0.697013742 | 0.130062696 | 0.145 | 0.2 | 0.368585762 | 1 |
| WBGene00005964 | srx-73 | 1.789177661 | 1.698983184 | 0.130123125 | 0.422 | 0.584 | 0.307975782 | 1 |
| WBGene00016260 | rege-1 | 0.526489161 | 0.436072878 | 0.130443122 | 0.092 | 0.084 | 0.73608874 | 1 |
| WBGene00019170 | H06I04.6 | 0.52479934 | 0.434370584 | 0.130461118 | 0.081 | 0.111 | 0.458598617 | 1 |
| WBGene00004151 | pqn-68 | 0.289697157 | 0.199063064 | 0.130757358 | 0.04 | 0.053 | 0.640912419 | 1 |
| WBGene00017744 | fbxc-54 | 0.313700318 | 0.223031949 | 0.130806807 | 0.052 | 0.047 | 0.786990731 | 1 |
| WBGene00012028 | srz-99 | 0.622105236 | 0.531299695 | 0.131004704 | 0.133 | 0.147 | 0.92933706 | 1 |
| WBGene00019759 | M03F4.6 | 0.342998086 | 0.25206797 | 0.131184427 | 0.046 | 0.063 | 0.530755566 | 1 |
| WBGene00044742 | Y55F3BR.10 | 0.452098537 | 0.360929109 | 0.131529681 | 0.081 | 0.095 | 0.7701071 | 1 |
| WBGene00012216 | W02D9.10 | 2.755689455 | 2.664422936 | 0.131669754 | 0.705 | 0.784 | 0.709078418 | 1 |
| WBGene00017290 | F09E5.12 | 0.560928773 | 0.469661268 | 0.131671177 | 0.104 | 0.147 | 0.384846347 | 1 |
| WBGene00017372 | F10G7.9 | 0.486957183 | 0.395507285 | 0.131934315 | 0.087 | 0.105 | 0.687781819 | 1 |
| WBGene00017349 | farl-11 | 0.398187731 | 0.306582314 | 0.132158681 | 0.058 | 0.074 | 0.619606491 | 1 |
| WBGene00021921 | rbm-39 | 0.603924734 | 0.512219294 | 0.132302984 | 0.104 | 0.126 | 0.676376422 | 1 |
| WBGene00001686 | gpd-4 | 0.522199905 | 0.430374511 | 0.13247604 | 0.058 | 0.1 | 0.185185526 | 1 |
| WBGene00012441 | glb-28 | 0.885246451 | 0.793031442 | 0.133038136 | 0.173 | 0.237 | 0.38440884 | 1 |
| WBGene00005007 | spr-2 | 0.997811867 | 0.905412854 | 0.133303597 | 0.179 | 0.295 | 0.076200746 | 1 |
| WBGene00045507 | F10A3.17 | 0.283660801 | 0.191178921 | 0.13342315 | 0.046 | 0.053 | 0.851119668 | 1 |
| WBGene00015219 | B0507.2 | 0.326211153 | 0.233664208 | 0.133517017 | 0.035 | 0.058 | 0.330115504 | 1 |
| WBGene00003927 | pas-6 | 0.682763819 | 0.590209336 | 0.133527894 | 0.133 | 0.163 | 0.633751241 | 1 |
| WBGene00006438 | nrfl-1 | 0.636414796 | 0.543780162 | 0.133643527 | 0.121 | 0.147 | 0.703772759 | 1 |
| WBGene00003182 | mef-2 | 0.641159629 | 0.548417079 | 0.133799217 | 0.127 | 0.153 | 0.72218165 | 1 |
| WBGene00019128 | F59G1.4 | 1.352976714 | 1.260201409 | 0.133846473 | 0.312 | 0.405 | 0.498112748 | 1 |
| WBGene00249815 | F20D12.12 | 0.668409041 | 0.575324655 | 0.134292383 | 0.11 | 0.105 | 0.712895062 | 1 |
| WBGene00011093 | R07B7.9 | 0.591767048 | 0.498573327 | 0.134450119 | 0.104 | 0.147 | 0.370790563 | 1 |
| WBGene00010408 | mboa-2 | 0.609493418 | 0.516139396 | 0.134681385 | 0.121 | 0.147 | 0.736575208 | 1 |
| WBGene00004831 | slo-2 | 1.108494102 | 1.014949981 | 0.134955641 | 0.249 | 0.305 | 0.660453089 | 1 |
| WBGene00011642 | T09B9.1 | 0.415926654 | 0.322173286 | 0.13525752 | 0.058 | 0.084 | 0.414187221 | 1 |
| WBGene00005461 | srh-255 | 1.786156159 | 1.692246687 | 0.13548273 | 0.445 | 0.553 | 0.779629066 | 1 |
| WBGene00001664 | gpa-2 | 1.6540047 | 1.559962784 | 0.135673805 | 0.416 | 0.542 | 0.426382106 | 1 |
| WBGene00006702 | ubc-3 | 2.263391678 | 2.169208747 | 0.135877247 | 0.613 | 0.763 | 0.834928106 | 1 |
| WBGene00001041 | dnj-23 | 0.953931617 | 0.859021645 | 0.136926146 | 0.191 | 0.247 | 0.453855538 | 1 |
| WBGene00021487 | comt-3 | 0.435183542 | 0.339996679 | 0.137325615 | 0.087 | 0.1 | 0.845269963 | 1 |
| WBGene00000514 | ckb-4 | 0.52626875 | 0.431060089 | 0.137357062 | 0.069 | 0.111 | 0.227452195 | 1 |
| WBGene00019334 | K02F3.12 | 0.328852316 | 0.233296418 | 0.137858021 | 0.046 | 0.063 | 0.535748496 | 1 |
| WBGene00008686 | F11A10.5 | 0.504245778 | 0.408605642 | 0.137979551 | 0.075 | 0.111 | 0.327210488 | 1 |
| WBGene00010261 | srh-308 | 0.250832202 | 0.155017715 | 0.138231085 | 0.052 | 0.037 | 0.451301926 | 1 |
| WBGene00044176 | C30G7.2 | 1.327150821 | 1.231318834 | 0.138256332 | 0.295 | 0.405 | 0.366956469 | 1 |
| WBGene00003625 | nhr-31 | 0.446436201 | 0.350384196 | 0.138573752 | 0.04 | 0.074 | 0.207272817 | 1 |
| WBGene00016831 | sre-56 | 0.474781924 | 0.378530459 | 0.138861512 | 0.052 | 0.079 | 0.356580316 | 1 |
| WBGene00001842 | her-1 | 0.826340957 | 0.730076351 | 0.13888047 | 0.104 | 0.158 | 0.222233136 | 1 |
| WBGene00000170 | aqp-2 | 0.735144888 | 0.638773982 | 0.139033829 | 0.127 | 0.168 | 0.471064987 | 1 |
| WBGene00010480 | tasp-1 | 1.393275183 | 1.296841092 | 0.139124985 | 0.358 | 0.453 | 0.701215165 | 1 |
| WBGene00007882 | C33A12.3 | 0.848008609 | 0.751574205 | 0.139125437 | 0.191 | 0.237 | 0.635996702 | 1 |
| WBGene00006936 | glp-4 | 0.277236916 | 0.180759266 | 0.139187826 | 0.029 | 0.058 | 0.206618117 | 1 |
| WBGene00009050 | F22D6.2 | 2.106970195 | 2.010109617 | 0.139740275 | 0.584 | 0.726 | 0.782976141 | 1 |
| WBGene00013189 | Y54E2A.4 | 0.282366809 | 0.185446902 | 0.13982587 | 0.046 | 0.058 | 0.69518443 | 1 |
| WBGene00015468 | madf-11 | 0.310646016 | 0.213542708 | 0.140090462 | 0.035 | 0.053 | 0.427970971 | 1 |

| | | | | | | | | |
|---|---|---|---|---|---|---|---|---|
| WBGene00004980 | spk-1 | 0.661270499 | 0.564128965 | 0.140145608 | 0.121 | 0.142 | 0.748455678 | 1 |
| WBGene00000466 | cel-1 | 0.377049621 | 0.279692988 | 0.140455932 | 0.075 | 0.079 | 0.992227457 | 1 |
| WBGene00007253 | C01H6.3 | 1.558675221 | 1.460743404 | 0.141285748 | 0.364 | 0.511 | 0.249722416 | 1 |
| WBGene00021845 | rpb-7 | 0.404586639 | 0.306132517 | 0.142039274 | 0.064 | 0.089 | 0.455804534 | 1 |
| WBGene00004490 | rps-21 | 2.602259711 | 2.503601253 | 0.142334069 | 0.746 | 0.853 | 0.95710313 | 1 |
| WBGene00000534 | cpi-2 | 1.582363674 | 1.483152385 | 0.143131635 | 0.399 | 0.521 | 0.65156733 | 1 |
| WBGene00001447 | flp-4 | 3.765072422 | 3.665810746 | 0.143204329 | 0.936 | 0.953 | 0.369507347 | 1 |
| WBGene00001241 | elo-3 | 1.136299359 | 1.037035681 | 0.143207215 | 0.197 | 0.321 | 0.062105318 | 1 |
| WBGene00018636 | oef-1 | 1.446102742 | 1.346807839 | 0.143252264 | 0.289 | 0.432 | 0.077452548 | 1 |
| WBGene00004506 | rpt-6 | 0.713845512 | 0.6144834 | 0.143349227 | 0.133 | 0.184 | 0.37290588 | 1 |
| WBGene00044045 | R11G10.3 | 0.319268372 | 0.219798397 | 0.143504839 | 0.035 | 0.053 | 0.449611924 | 1 |
| WBGene00007009 | wwp-1 | 0.766734094 | 0.667128003 | 0.143701214 | 0.145 | 0.179 | 0.596606808 | 1 |
| WBGene00011824 | T18D3.7 | 0.406702034 | 0.307057457 | 0.143756737 | 0.069 | 0.1 | 0.403876168 | 1 |
| WBGene00017467 | F14F9.4 | 0.597217007 | 0.497570754 | 0.143759156 | 0.092 | 0.116 | 0.604890735 | 1 |
| WBGene00012765 | Y41E3.7 | 0.93035193 | 0.830671189 | 0.143808912 | 0.191 | 0.253 | 0.493341666 | 1 |
| WBGene00008287 | C54C6.5 | 0.388291224 | 0.288599179 | 0.143825219 | 0.052 | 0.053 | 0.978263198 | 1 |
| WBGene00006531 | tba-5 | 1.255629789 | 1.155648697 | 0.144242225 | 0.26 | 0.353 | 0.240207857 | 1 |
| WBGene00021088 | W08E12.7 | 0.415658749 | 0.31563082 | 0.144309796 | 0.075 | 0.095 | 0.644238231 | 1 |
| WBGene00015508 | mvb-12 | 0.476146128 | 0.376083218 | 0.144360264 | 0.092 | 0.111 | 0.780644668 | 1 |
| WBGene00016539 | madd-2 | 0.365979581 | 0.265767473 | 0.144575511 | 0.04 | 0.068 | 0.286868238 | 1 |
| WBGene00045419 | lsy-12 | 0.8289996 | 0.728464308 | 0.145041768 | 0.162 | 0.211 | 0.474107176 | 1 |
| WBGene00000508 | cit-1.2 | 0.567313575 | 0.466395209 | 0.145594426 | 0.092 | 0.147 | 0.203057405 | 1 |
| WBGene00009650 | ccnk-1 | 0.292661901 | 0.191517435 | 0.145920619 | 0.046 | 0.058 | 0.70286966 | 1 |
| WBGene00019495 | sdz-24 | 2.684283453 | 2.583110838 | 0.14596123 | 0.694 | 0.811 | 0.87374169 | 1 |
| WBGene00018327 | F42A6.5 | 1.259903925 | 1.15846783 | 0.146341351 | 0.289 | 0.395 | 0.397143915 | 1 |
| WBGene00011722 | rbm-22 | 0.376465513 | 0.275028011 | 0.146343381 | 0.064 | 0.084 | 0.572303588 | 1 |
| WBGene00019579 | oac-57 | 1.034784066 | 0.933151569 | 0.146624699 | 0.191 | 0.253 | 0.371758482 | 1 |
| WBGene00009722 | F45D3.2 | 0.961131994 | 0.859475222 | 0.146659722 | 0.139 | 0.216 | 0.102270598 | 1 |
| WBGene00009826 | F47G4.5 | 0.680588558 | 0.578743428 | 0.146931464 | 0.121 | 0.168 | 0.367183081 | 1 |
| WBGene00016562 | C41D11.3 | 0.306862856 | 0.204979544 | 0.146986549 | 0.017 | 0.053 | 0.076971609 | 1 |
| WBGene00019774 | M04G7.3 | 1.422243612 | 1.320334414 | 0.147023894 | 0.329 | 0.421 | 0.541892058 | 1 |
| WBGene00004320 | rbx-1 | 0.423265764 | 0.321163005 | 0.147303143 | 0.064 | 0.079 | 0.672824472 | 1 |
| WBGene00007247 | C01G12.7 | 3.116318598 | 3.014098822 | 0.147471963 | 0.861 | 0.937 | 0.281430473 | 1 |
| WBGene00014122 | ZK863.1 | 1.785878241 | 1.683615559 | 0.147533865 | 0.434 | 0.632 | 0.204511868 | 1 |
| WBGene00009201 | F28C1.3 | 0.302257792 | 0.199958307 | 0.14758696 | 0.052 | 0.053 | 0.947242617 | 1 |
| WBGene00002001 | hars-1 | 0.317432709 | 0.214955217 | 0.147843769 | 0.058 | 0.063 | 0.931491041 | 1 |
| WBGene00017929 | grld-1 | 0.524781649 | 0.422101059 | 0.148136779 | 0.075 | 0.1 | 0.479146097 | 1 |
| WBGene00003404 | mpz-1 | 0.679855593 | 0.577163756 | 0.148153004 | 0.133 | 0.153 | 0.80077887 | 1 |
| WBGene00007226 | C01G6.4 | 1.249141162 | 1.14626029 | 0.148425723 | 0.277 | 0.384 | 0.325840609 | 1 |
| WBGene00022712 | ZK355.2 | 0.415819241 | 0.312580787 | 0.148941606 | 0.046 | 0.068 | 0.401422608 | 1 |
| WBGene00003640 | nhr-50 | 1.0908487 | 0.987460596 | 0.149157505 | 0.202 | 0.274 | 0.315715199 | 1 |
| WBGene00010560 | eif-2beta | 0.83231102 | 0.728803161 | 0.149330274 | 0.173 | 0.221 | 0.597599617 | 1 |
| WBGene00001851 | hif-1 | 1.829465326 | 1.725881871 | 0.149439337 | 0.434 | 0.616 | 0.235475062 | 1 |
| WBGene00022251 | blos-2 | 0.449378487 | 0.345600407 | 0.149720121 | 0.058 | 0.105 | 0.1511124 | 1 |
| WBGene00007953 | hda-11 | 0.350420041 | 0.246092942 | 0.150512189 | 0.064 | 0.053 | 0.617957552 | 1 |
| WBGene00021857 | iffb-1 | 0.271390539 | 0.166666334 | 0.151085091 | 0.058 | 0.047 | 0.598357668 | 1 |
| WBGene00011263 | R13H4.5 | 1.69506591 | 1.590160689 | 0.151346243 | 0.41 | 0.505 | 0.914419712 | 1 |
| WBGene00022347 | Y82E9BR.14 | 0.423553897 | 0.318327347 | 0.151809821 | 0.064 | 0.089 | 0.435149565 | 1 |
| WBGene00013477 | Y69E1A.5 | 0.455579801 | 0.350295431 | 0.151893238 | 0.075 | 0.105 | 0.442398632 | 1 |
| WBGene00000230 | atp-3 | 0.942414727 | 0.837121877 | 0.151905472 | 0.185 | 0.258 | 0.289393126 | 1 |
| WBGene00012019 | dkf-2 | 1.650245748 | 1.544832109 | 0.152079734 | 0.364 | 0.484 | 0.559542326 | 1 |
| WBGene00008338 | pafo-1 | 0.328078581 | 0.222656028 | 0.152092594 | 0.035 | 0.053 | 0.446242709 | 1 |
| WBGene00003637 | nhr-47 | 1.219741444 | 1.113982009 | 0.152578612 | 0.289 | 0.395 | 0.482699661 | 1 |

| | | | | | | | | |
|---|---|---|---|---|---|---|---|---|
| WBGene00009322 | tbc-3 | 0.483947244 | 0.378023191 | 0.152816106 | 0.081 | 0.105 | 0.547625792 | 1 |
| WBGene00001745 | gsa-1 | 1.130959095 | 1.024982367 | 0.152892101 | 0.231 | 0.316 | 0.317951714 | 1 |
| WBGene00019158 | H05C05.2 | 0.321289891 | 0.215262401 | 0.152965334 | 0.046 | 0.063 | 0.557654302 | 1 |
| WBGene00007633 | C16D2.1 | 0.404851831 | 0.298698459 | 0.153146944 | 0.069 | 0.074 | 0.970514473 | 1 |
| WBGene00004040 | pld-1 | 0.369727337 | 0.263299342 | 0.153543141 | 0.069 | 0.079 | 0.858476101 | 1 |
| WBGene00016661 | C45B2.6 | 0.510775435 | 0.404315064 | 0.153589849 | 0.087 | 0.068 | 0.450589728 | 1 |
| WBGene00011974 | xbx-5 | 1.003353199 | 0.896766502 | 0.1537721 | 0.168 | 0.221 | 0.465608574 | 1 |
| WBGene00022127 | yop-1 | 0.657873889 | 0.551144949 | 0.153977313 | 0.133 | 0.168 | 0.581325801 | 1 |
| WBGene00015819 | fbxc-44 | 0.310777978 | 0.203760276 | 0.154393908 | 0.046 | 0.053 | 0.849033095 | 1 |
| WBGene00021734 | Y50C1A.2 | 0.377659228 | 0.270422518 | 0.154709869 | 0.058 | 0.089 | 0.328357867 | 1 |
| WBGene00017080 | lec-12 | 0.533665475 | 0.426416066 | 0.154728191 | 0.104 | 0.105 | 0.880889006 | 1 |
| WBGene00001196 | egl-30 | 1.725801724 | 1.618221541 | 0.155205395 | 0.491 | 0.553 | 0.647611724 | 1 |
| WBGene00010559 | K04D7.6 | 3.877675346 | 3.769816603 | 0.155607274 | 0.919 | 0.958 | 0.084035478 | 1 |
| WBGene00019467 | K07B1.7 | 0.480402712 | 0.372150922 | 0.156174321 | 0.092 | 0.095 | 0.942349558 | 1 |
| WBGene00014166 | ZK945.4 | 0.274630371 | 0.16595338 | 0.156787755 | 0.052 | 0.037 | 0.452995517 | 1 |
| WBGene00018544 | nhr-188 | 1.345043961 | 1.236314612 | 0.156863294 | 0.26 | 0.389 | 0.122875388 | 1 |
| WBGene00008744 | F13D12.10 | 0.511179619 | 0.402449442 | 0.156864487 | 0.087 | 0.1 | 0.802863716 | 1 |
| WBGene00004057 | pmk-3 | 1.115885042 | 1.00681598 | 0.157353394 | 0.231 | 0.316 | 0.353041996 | 1 |
| WBGene00011920 | T22C1.11 | 0.686735114 | 0.577394769 | 0.157744774 | 0.121 | 0.153 | 0.610743548 | 1 |
| WBGene00011511 | gpcp-2 | 0.389194157 | 0.279659359 | 0.158025311 | 0.081 | 0.079 | 0.813085635 | 1 |
| WBGene00005530 | sri-18 | 0.535211368 | 0.425168323 | 0.158758556 | 0.081 | 0.105 | 0.543674735 | 1 |
| WBGene00009813 | haly-1 | 0.904469526 | 0.794393993 | 0.158805425 | 0.191 | 0.242 | 0.550348627 | 1 |
| WBGene00020391 | cct-7 | 0.796880195 | 0.686727227 | 0.15891714 | 0.156 | 0.195 | 0.648852467 | 1 |
| WBGene00001718 | grl-9 | 0.421162301 | 0.310532007 | 0.159605777 | 0.052 | 0.079 | 0.366366676 | 1 |
| WBGene00006807 | unc-75 | 0.324311226 | 0.213494002 | 0.159875461 | 0.035 | 0.068 | 0.181428021 | 1 |
| WBGene00001117 | dyf-1 | 1.30385677 | 1.193030532 | 0.159888465 | 0.266 | 0.379 | 0.284694098 | 1 |
| WBGene00003653 | nhr-63 | 0.454717697 | 0.343802607 | 0.160016649 | 0.081 | 0.095 | 0.781028133 | 1 |
| WBGene00013538 | ari-1.4 | 0.410078248 | 0.29892108 | 0.160365896 | 0.069 | 0.079 | 0.844668356 | 1 |
| WBGene00022139 | tub-2 | 0.416772642 | 0.305364493 | 0.160727985 | 0.069 | 0.084 | 0.716900929 | 1 |
| WBGene00003697 | nhr-107 | 0.712852673 | 0.601384449 | 0.160814654 | 0.121 | 0.174 | 0.304647049 | 1 |
| WBGene00006433 | sdhb-1 | 0.728649368 | 0.617177885 | 0.160819357 | 0.116 | 0.179 | 0.18651282 | 1 |
| WBGene00017286 | F09E5.8 | 0.810819455 | 0.69931594 | 0.160865568 | 0.133 | 0.168 | 0.492017085 | 1 |
| WBGene00000101 | aka-1 | 0.554277277 | 0.44250142 | 0.161258474 | 0.087 | 0.132 | 0.284855832 | 1 |
| WBGene00001233 | eif-3.K | 0.395468733 | 0.283634987 | 0.16134199 | 0.052 | 0.079 | 0.376318852 | 1 |
| WBGene00021043 | pck-1 | 0.354828693 | 0.242836021 | 0.161571272 | 0.064 | 0.058 | 0.750144647 | 1 |
| WBGene00004825 | skr-19 | 0.660186802 | 0.547773725 | 0.162177789 | 0.092 | 0.158 | 0.123658677 | 1 |
| WBGene00008370 | D1054.1 | 0.402490384 | 0.28995952 | 0.16234772 | 0.075 | 0.095 | 0.66812479 | 1 |
| WBGene00014241 | ZK1251.3 | 0.990355067 | 0.8776881 | 0.162544075 | 0.243 | 0.263 | 0.826242182 | 1 |
| WBGene00021967 | Y57G7A.3 | 0.343183634 | 0.23032513 | 0.162820403 | 0.064 | 0.042 | 0.352231333 | 1 |
| WBGene00008820 | nlp-55 | 2.638499027 | 2.525606564 | 0.162869396 | 0.763 | 0.832 | 0.364392962 | 1 |
| WBGene00005149 | sre-1 | 2.410888862 | 2.297582362 | 0.163466725 | 0.647 | 0.784 | 0.906478464 | 1 |
| WBGene00003777 | nmy-2 | 0.421592354 | 0.308084655 | 0.163756995 | 0.069 | 0.1 | 0.409719176 | 1 |
| WBGene00021316 | Y32H12A.8 | 0.291246736 | 0.177729808 | 0.163770309 | 0.046 | 0.053 | 0.855295927 | 1 |
| WBGene00011729 | set-16 | 0.400502852 | 0.286619537 | 0.164298894 | 0.069 | 0.089 | 0.609184905 | 1 |
| WBGene00001129 | dyf-13 | 1.424940091 | 1.310840019 | 0.164611608 | 0.347 | 0.426 | 0.990901514 | 1 |
| WBGene00012130 | slc-17.4 | 0.570824938 | 0.456624396 | 0.164756556 | 0.104 | 0.121 | 0.783184387 | 1 |
| WBGene00001137 | eat-6 | 1.317441254 | 1.20319403 | 0.164823904 | 0.26 | 0.421 | 0.071981733 | 1 |
| WBGene00005588 | sri-76 | 2.006200699 | 1.891867449 | 0.164948013 | 0.434 | 0.616 | 0.082534085 | 1 |
| WBGene00000268 | bre-3 | 0.618622228 | 0.503932713 | 0.165461994 | 0.121 | 0.137 | 0.886015879 | 1 |
| WBGene00000920 | daf-28 | 2.382674512 | 2.267978743 | 0.165471017 | 0.486 | 0.532 | 0.857828107 | 1 |
| WBGene00022579 | ZC262.2 | 0.455146318 | 0.340446617 | 0.165476689 | 0.058 | 0.074 | 0.614694727 | 1 |
| WBGene00018468 | cla-1 | 1.70262539 | 1.587913918 | 0.165493673 | 0.382 | 0.479 | 0.611113989 | 1 |
| WBGene00010742 | K10D6.2 | 1.21522458 | 1.100453931 | 0.165579046 | 0.254 | 0.358 | 0.262975687 | 1 |

| | | | | | | | | |
|---|---|---|---|---|---|---|---|---|
| WBGene00009274 | F30F8.5 | 0.300572311 | 0.185784309 | 0.165604081 | 0.04 | 0.053 | 0.646796835 | 1 |
| WBGene00007855 | C31H5.4 | 0.845880905 | 0.731087185 | 0.16561233 | 0.162 | 0.195 | 0.807632888 | 1 |
| WBGene00000484 | che-2 | 0.708556963 | 0.593641128 | 0.165788505 | 0.121 | 0.147 | 0.711296396 | 1 |
| WBGene00003072 | lrp-2 | 0.474877325 | 0.359920694 | 0.165847361 | 0.081 | 0.111 | 0.483871297 | 1 |
| WBGene00010260 | ddx-17 | 0.959653465 | 0.844317993 | 0.166393914 | 0.225 | 0.268 | 0.951014747 | 1 |
| WBGene00003587 | ned-8 | 0.616549018 | 0.500977087 | 0.166735052 | 0.092 | 0.158 | 0.130126131 | 1 |
| WBGene00020894 | T28D9.1 | 2.235000237 | 2.119305701 | 0.166911933 | 0.59 | 0.747 | 0.535985819 | 1 |
| WBGene00001670 | gpa-8 | 1.05048266 | 0.934773085 | 0.166933631 | 0.197 | 0.284 | 0.223655738 | 1 |
| WBGene00019127 | cgt-3 | 0.314981169 | 0.199110253 | 0.167166396 | 0.04 | 0.053 | 0.635050327 | 1 |
| WBGene00000089 | aex-6 | 1.680303545 | 1.564393066 | 0.167223473 | 0.434 | 0.547 | 0.608496349 | 1 |
| WBGene00022402 | lmtr-2 | 0.353915374 | 0.23797878 | 0.167261149 | 0.058 | 0.063 | 0.929566163 | 1 |
| WBGene00012121 | T28C6.7 | 0.558639703 | 0.442694584 | 0.167273449 | 0.052 | 0.074 | 0.434510537 | 1 |
| WBGene00019207 | fgt-1 | 0.361771195 | 0.245720819 | 0.167425301 | 0.052 | 0.074 | 0.484874106 | 1 |
| WBGene00018609 | F48E8.2 | 0.297087534 | 0.180343686 | 0.16842577 | 0.04 | 0.053 | 0.652703421 | 1 |
| WBGene00043992 | bbs-4 | 1.214594534 | 1.097850087 | 0.168426634 | 0.283 | 0.379 | 0.490922138 | 1 |
| WBGene00012546 | Y37D8A.4 | 0.303196083 | 0.186270861 | 0.168687438 | 0.064 | 0.037 | 0.228328031 | 1 |
| WBGene00017064 | ppfr-2 | 0.441413424 | 0.324372895 | 0.168853791 | 0.04 | 0.084 | 0.110180231 | 1 |
| WBGene00003826 | ntl-3 | 0.507451273 | 0.390349678 | 0.16894189 | 0.087 | 0.105 | 0.684905549 | 1 |
| WBGene00000068 | acy-1 | 0.422930027 | 0.305504904 | 0.169408642 | 0.058 | 0.084 | 0.400241916 | 1 |
| WBGene00044206 | T26H5.9 | 0.414229606 | 0.296673889 | 0.16959705 | 0.058 | 0.079 | 0.499501754 | 1 |
| WBGene00017952 | F31E3.6 | 0.246405617 | 0.128839187 | 0.169612506 | 0.052 | 0.042 | 0.587115436 | 1 |
| WBGene00000542 | clp-1 | 0.589895503 | 0.472008626 | 0.170074814 | 0.081 | 0.132 | 0.180375503 | 1 |
| WBGene00021296 | Y25C1A.13 | 0.647066104 | 0.529074192 | 0.170226346 | 0.092 | 0.132 | 0.310088169 | 1 |
| WBGene00006750 | unc-10 | 0.38532217 | 0.267033448 | 0.170654553 | 0.035 | 0.053 | 0.436221768 | 1 |
| WBGene00014115 | gld-4 | 0.369160385 | 0.250859627 | 0.170671916 | 0.052 | 0.047 | 0.799304942 | 1 |
| WBGene00019162 | eif-1.A | 0.493095208 | 0.374550513 | 0.171023844 | 0.087 | 0.105 | 0.709495483 | 1 |
| WBGene00019641 | K10G6.4 | 0.397544689 | 0.27888572 | 0.171188706 | 0.052 | 0.074 | 0.448583388 | 1 |
| WBGene00016754 | cebp-2 | 0.37020652 | 0.251444039 | 0.171338043 | 0.04 | 0.068 | 0.290320505 | 1 |
| WBGene00012337 | scyl-1 | 0.924058634 | 0.805221511 | 0.171445729 | 0.145 | 0.253 | 0.055708638 | 1 |
| WBGene00020026 | R12C12.6 | 0.403476617 | 0.284623845 | 0.171468305 | 0.058 | 0.068 | 0.774124925 | 1 |
| WBGene00003707 | nhr-117 | 0.866802185 | 0.747754096 | 0.171750087 | 0.168 | 0.221 | 0.454192173 | 1 |
| WBGene00004830 | slo-1 | 0.509766514 | 0.390560605 | 0.171977774 | 0.052 | 0.111 | 0.065710199 | 1 |
| WBGene00003602 | nhr-3 | 0.631256333 | 0.512033244 | 0.172002558 | 0.098 | 0.142 | 0.331653907 | 1 |
| WBGene00022614 | ZC449.4 | 1.301080339 | 1.181660335 | 0.172286647 | 0.318 | 0.368 | 0.977373182 | 1 |
| WBGene00020482 | T13C2.7 | 0.733012448 | 0.613522221 | 0.172387958 | 0.116 | 0.174 | 0.236465024 | 1 |
| WBGene00011312 | trcs-2 | 1.445401816 | 1.325571655 | 0.172878379 | 0.364 | 0.442 | 0.782848052 | 1 |
| WBGene00007167 | rbg-3 | 0.333025218 | 0.213102204 | 0.173012338 | 0.064 | 0.068 | 0.980522768 | 1 |
| WBGene00018395 | mtch-1 | 1.295226788 | 1.175184671 | 0.173184167 | 0.283 | 0.358 | 0.634470599 | 1 |
| WBGene00019479 | K07C11.7 | 1.583528161 | 1.463341518 | 0.173392674 | 0.405 | 0.484 | 0.631006353 | 1 |
| WBGene00012626 | famh-161 | 0.644332866 | 0.523789563 | 0.173907225 | 0.116 | 0.137 | 0.725668697 | 1 |
| WBGene00005326 | srh-107 | 0.534660726 | 0.413889304 | 0.174236331 | 0.064 | 0.095 | 0.338312498 | 1 |
| WBGene00007184 | ctr-9 | 0.626012949 | 0.505213654 | 0.174276543 | 0.087 | 0.142 | 0.172558243 | 1 |
| WBGene00010967 | nduo-5 | 1.499009847 | 1.37811072 | 0.174420572 | 0.329 | 0.432 | 0.383257392 | 1 |
| WBGene00001425 | fis-2 | 0.651077226 | 0.530148486 | 0.174463293 | 0.127 | 0.158 | 0.648258179 | 1 |
| WBGene00006705 | ubc-8 | 0.593804966 | 0.472872336 | 0.174468907 | 0.104 | 0.137 | 0.496956412 | 1 |
| WBGene00002239 | ksr-1 | 0.6480604 | 0.526454042 | 0.17544089 | 0.098 | 0.142 | 0.314357268 | 1 |
| WBGene00017174 | tat-6 | 0.368375721 | 0.246557043 | 0.175747202 | 0.046 | 0.068 | 0.429773214 | 1 |
| WBGene00001193 | egl-26 | 0.341440894 | 0.219313985 | 0.176191886 | 0.069 | 0.058 | 0.570814886 | 1 |
| WBGene00005321 | srh-102 | 0.345371946 | 0.222886028 | 0.176709826 | 0.064 | 0.063 | 0.882157075 | 1 |
| WBGene00009254 | capg-1 | 0.620427762 | 0.497940069 | 0.176712387 | 0.087 | 0.137 | 0.196662128 | 1 |
| WBGene00017184 | ncam-1 | 0.424459629 | 0.301851662 | 0.176885906 | 0.052 | 0.089 | 0.224251667 | 1 |
| WBGene00020910 | dgtr-1 | 0.483149576 | 0.360413876 | 0.177070187 | 0.081 | 0.089 | 0.911163331 | 1 |
| WBGene00017961 | nhr-180 | 0.359469704 | 0.236397319 | 0.177555919 | 0.058 | 0.063 | 0.941121218 | 1 |

| | | | | | | | | |
|---|---|---|---|---|---|---|---|---|
| WBGene00020437 | stt-3 | 0.653241545 | 0.530110751 | 0.177640185 | 0.081 | 0.116 | 0.343572008 | 1 |
| WBGene00010731 | K10C3.4 | 0.505297827 | 0.381933211 | 0.177977519 | 0.069 | 0.105 | 0.319823704 | 1 |
| WBGene00011815 | bath-43 | 0.518458731 | 0.394863036 | 0.178310896 | 0.092 | 0.095 | 0.948658151 | 1 |
| WBGene00017692 | F22B7.1 | 1.185089918 | 1.061456129 | 0.178365853 | 0.225 | 0.353 | 0.107191542 | 1 |
| WBGene00018784 | F54A3.5 | 0.4573383 | 0.333577558 | 0.178549008 | 0.069 | 0.079 | 0.822334026 | 1 |
| WBGene00019711 | M01E11.2 | 0.335972748 | 0.212176191 | 0.178600678 | 0.023 | 0.063 | 0.073536271 | 1 |
| WBGene00005391 | srh-175 | 0.516099535 | 0.392276838 | 0.178638391 | 0.081 | 0.105 | 0.525426095 | 1 |
| WBGene00004917 | snr-4 | 0.874744561 | 0.750696972 | 0.178962841 | 0.173 | 0.232 | 0.48166675 | 1 |
| WBGene00020888 | T28C12.1 | 0.411965709 | 0.287784241 | 0.179155988 | 0.052 | 0.058 | 0.844385287 | 1 |
| WBGene00021697 | gcn-1 | 0.47669096 | 0.352468833 | 0.179214646 | 0.081 | 0.089 | 0.87929659 | 1 |
| WBGene00004063 | pmr-1 | 1.43072523 | 1.306285926 | 0.179527968 | 0.353 | 0.437 | 0.861853808 | 1 |
| WBGene00015861 | C16D9.6 | 1.636701885 | 1.511869937 | 0.180094432 | 0.353 | 0.484 | 0.321194999 | 1 |
| WBGene00001681 | gpc-1 | 2.76052135 | 2.635672294 | 0.180119114 | 0.769 | 0.874 | 0.439072985 | 1 |
| WBGene00014145 | ZK899.7 | 0.347047246 | 0.222163268 | 0.180169495 | 0.052 | 0.037 | 0.468398321 | 1 |
| WBGene00011763 | T14B1.1 | 0.521878268 | 0.396866351 | 0.180354073 | 0.098 | 0.074 | 0.371534101 | 1 |
| WBGene00009625 | F41E7.9 | 0.560650703 | 0.435598181 | 0.180412653 | 0.098 | 0.126 | 0.597801889 | 1 |
| WBGene00000899 | daf-3 | 0.972124431 | 0.847016974 | 0.180491907 | 0.156 | 0.237 | 0.177066437 | 1 |
| WBGene00011372 | npr-26 | 1.716500169 | 1.591141889 | 0.180853768 | 0.41 | 0.537 | 0.550671727 | 1 |
| WBGene00235291 | C01G5.25 | 1.084572192 | 0.959087287 | 0.181036449 | 0.214 | 0.3 | 0.297269407 | 1 |
| WBGene00006823 | unc-94 | 0.720955455 | 0.595395932 | 0.181144101 | 0.139 | 0.195 | 0.376183277 | 1 |
| WBGene00008405 | ttll-12 | 0.598991233 | 0.473424206 | 0.181154927 | 0.058 | 0.105 | 0.141786578 | 1 |
| WBGene00003124 | mai-1 | 0.406481571 | 0.280594419 | 0.181616769 | 0.069 | 0.068 | 0.894733415 | 1 |
| WBGene00018270 | eas-1 | 0.321472448 | 0.195537722 | 0.181685404 | 0.046 | 0.053 | 0.849033095 | 1 |
| WBGene00020747 | T24A6.7 | 0.367959215 | 0.241856721 | 0.181927443 | 0.058 | 0.074 | 0.637760663 | 1 |
| WBGene00015790 | C15C7.4 | 2.241025427 | 2.114584519 | 0.182415671 | 0.399 | 0.395 | 0.598092276 | 1 |
| WBGene00010013 | F54B3.1 | 0.424917989 | 0.298476351 | 0.182416724 | 0.075 | 0.079 | 0.990500299 | 1 |
| WBGene00010239 | F58B4.6 | 2.664826668 | 2.538329014 | 0.182497539 | 0.775 | 0.826 | 0.409694176 | 1 |
| WBGene00008380 | prp-38 | 0.279757215 | 0.15316258 | 0.182637451 | 0.023 | 0.053 | 0.163202025 | 1 |
| WBGene00005519 | sri-7 | 0.800076068 | 0.6733605 | 0.182811921 | 0.116 | 0.174 | 0.233252576 | 1 |
| WBGene00013266 | Y57A10A.26 | 0.982179922 | 0.8554407 | 0.182846046 | 0.168 | 0.274 | 0.098735138 | 1 |
| WBGene00016601 | acin-1 | 0.31796113 | 0.191101065 | 0.183020386 | 0.029 | 0.058 | 0.197654655 | 1 |
| WBGene00004357 | rho-1 | 1.130479549 | 1.003477711 | 0.183224922 | 0.225 | 0.342 | 0.148770167 | 1 |
| WBGene00018081 | F36A4.2 | 1.164106697 | 1.036982287 | 0.183401756 | 0.266 | 0.289 | 0.873584517 | 1 |
| WBGene00007877 | nfki-1 | 2.083291733 | 1.955832079 | 0.183885411 | 0.555 | 0.679 | 0.844415858 | 1 |
| WBGene00018017 | golg-2 | 0.302444148 | 0.17452428 | 0.184549359 | 0.046 | 0.053 | 0.863660503 | 1 |
| WBGene00004143 | pqn-59 | 0.869264824 | 0.741344753 | 0.184549653 | 0.173 | 0.195 | 0.878246014 | 1 |
| WBGene00022794 | snu-23 | 0.665059398 | 0.537013985 | 0.184730482 | 0.116 | 0.158 | 0.392991508 | 1 |
| WBGene00020930 | hlh-30 | 1.131909383 | 1.003742199 | 0.184906161 | 0.254 | 0.311 | 0.7178091 | 1 |
| WBGene00000912 | daf-16 | 1.49007038 | 1.361833258 | 0.18500706 | 0.37 | 0.495 | 0.66460869 | 1 |
| WBGene00011756 | ctg-2 | 0.770876541 | 0.642268512 | 0.185542166 | 0.162 | 0.163 | 0.752821262 | 1 |
| WBGene00022453 | ncap-1 | 0.614951586 | 0.486328091 | 0.185564479 | 0.11 | 0.137 | 0.624863922 | 1 |
| WBGene00001008 | dlk-1 | 0.686955515 | 0.558320727 | 0.185580771 | 0.127 | 0.158 | 0.596001722 | 1 |
| WBGene00012444 | Y15E3A.5 | 0.803859559 | 0.675085201 | 0.185782127 | 0.104 | 0.189 | 0.052898961 | 1 |
| WBGene00016509 | adss-1 | 0.612333177 | 0.483491323 | 0.185879503 | 0.081 | 0.142 | 0.12051299 | 1 |
| WBGene00003393 | mog-5 | 0.542305475 | 0.413230907 | 0.186215239 | 0.087 | 0.105 | 0.712409017 | 1 |
| WBGene00002994 | lin-5 | 0.420533668 | 0.291076167 | 0.186767696 | 0.075 | 0.074 | 0.867128219 | 1 |
| WBGene00010179 | F57A8.4 | 0.973888022 | 0.844256303 | 0.187019038 | 0.156 | 0.184 | 0.629863785 | 1 |
| WBGene00011480 | enpl-1 | 1.37824353 | 1.248510191 | 0.187165645 | 0.329 | 0.368 | 0.989209664 | 1 |
| WBGene00003715 | nhr-125 | 0.492224352 | 0.362378452 | 0.187328035 | 0.052 | 0.089 | 0.208441101 | 1 |
| WBGene00007761 | rad-26 | 0.741286612 | 0.611403569 | 0.187381622 | 0.139 | 0.132 | 0.703147084 | 1 |
| WBGene00015064 | B0228.7 | 0.837039656 | 0.707119482 | 0.187435191 | 0.156 | 0.216 | 0.405381182 | 1 |
| WBGene00016315 | C32D5.7 | 2.214586096 | 2.084559702 | 0.187588435 | 0.578 | 0.737 | 0.678327693 | 1 |
| WBGene00003401 | mpk-1 | 1.531078089 | 1.401009306 | 0.187649588 | 0.382 | 0.479 | 0.944359265 | 1 |

| | | | | | | | | |
|---|---|---|---|---|---|---|---|---|
| WBGene00015207 | B0495.8 | 0.84723123 | 0.717010884 | 0.187868246 | 0.168 | 0.242 | 0.310497588 | 1 |
| WBGene00011311 | T01B7.5 | 0.769481235 | 0.638990031 | 0.188259013 | 0.15 | 0.179 | 0.760367058 | 1 |
| WBGene00011352 | rskn-1 | 1.760919022 | 1.630234443 | 0.188537994 | 0.439 | 0.568 | 0.609112502 | 1 |
| WBGene00013219 | Y54G11A.11 | 0.68540665 | 0.554611424 | 0.188697624 | 0.139 | 0.168 | 0.770004829 | 1 |
| WBGene00002162 | isp-1 | 0.917705447 | 0.786810902 | 0.18884091 | 0.185 | 0.247 | 0.474939285 | 1 |
| WBGene00005523 | sri-11 | 0.441531854 | 0.310609023 | 0.188881719 | 0.087 | 0.079 | 0.675643944 | 1 |
| WBGene00008439 | mfb-1 | 1.097483012 | 0.966518294 | 0.18894215 | 0.237 | 0.289 | 0.713952688 | 1 |
| WBGene00021068 | W06H8.6 | 0.448555965 | 0.31733847 | 0.189306829 | 0.064 | 0.095 | 0.358010814 | 1 |
| WBGene00018361 | F42G8.10 | 0.671597176 | 0.540279885 | 0.189450804 | 0.133 | 0.158 | 0.736317418 | 1 |
| WBGene00022750 | tres-1 | 0.656095189 | 0.524575772 | 0.189742412 | 0.116 | 0.168 | 0.322757036 | 1 |
| WBGene00017193 | F07C3.2 | 0.549006641 | 0.41721745 | 0.190131612 | 0.104 | 0.132 | 0.62486687 | 1 |
| WBGene00003831 | nuo-1 | 0.697793195 | 0.566000883 | 0.190136115 | 0.121 | 0.158 | 0.531495123 | 1 |
| WBGene00013761 | Y113G7B.11 | 0.387080214 | 0.254986865 | 0.190570419 | 0.052 | 0.068 | 0.578367014 | 1 |
| WBGene00010962 | ctc-3 | 2.646801819 | 2.514676223 | 0.190616941 | 0.763 | 0.842 | 0.640546255 | 1 |
| WBGene00011566 | nhr-214 | 0.495408403 | 0.362846102 | 0.191246975 | 0.058 | 0.095 | 0.238504631 | 1 |
| WBGene00022475 | Y119C1B.10 | 0.416233012 | 0.283366796 | 0.191685431 | 0.069 | 0.084 | 0.721758692 | 1 |
| WBGene00010833 | M03B6.1 | 0.855225622 | 0.722273794 | 0.191808943 | 0.156 | 0.2 | 0.564032349 | 1 |
| WBGene00005591 | srj-1 | 0.470013796 | 0.33706171 | 0.191809316 | 0.087 | 0.1 | 0.828246303 | 1 |
| WBGene00009566 | srsx-1 | 0.416310045 | 0.283225815 | 0.191999958 | 0.058 | 0.079 | 0.521487631 | 1 |
| WBGene00010529 | srz-21 | 0.400405375 | 0.267187617 | 0.192192598 | 0.069 | 0.058 | 0.59194553 | 1 |
| WBGene00013735 | Y111B2A.12 | 0.409536617 | 0.276124795 | 0.192472574 | 0.064 | 0.074 | 0.804502418 | 1 |
| WBGene00018802 | F54D8.6 | 0.34644179 | 0.212893048 | 0.192670107 | 0.046 | 0.058 | 0.69231035 | 1 |
| WBGene00004405 | rop-1 | 0.48427491 | 0.350574543 | 0.192888856 | 0.087 | 0.105 | 0.712409017 | 1 |
| WBGene00019006 | spg-20 | 0.983646118 | 0.849041558 | 0.194193331 | 0.197 | 0.263 | 0.455587836 | 1 |
| WBGene00003423 | msi-1 | 0.416809191 | 0.282183302 | 0.194224102 | 0.081 | 0.089 | 0.955512568 | 1 |
| WBGene00012896 | snrp-200 | 0.792332612 | 0.657453557 | 0.194589345 | 0.133 | 0.168 | 0.553054517 | 1 |
| WBGene00010200 | F57C7.4 | 1.120274307 | 0.985328824 | 0.194685178 | 0.197 | 0.305 | 0.108534312 | 1 |
| WBGene00008275 | nstp-1 | 0.786450785 | 0.651472507 | 0.194732491 | 0.156 | 0.211 | 0.448113015 | 1 |
| WBGene00006975 | zfp-1 | 1.230988415 | 1.095708865 | 0.195167135 | 0.272 | 0.311 | 0.941852725 | 1 |
| WBGene00012351 | W09C5.1 | 0.573496707 | 0.438059041 | 0.195395249 | 0.104 | 0.132 | 0.629929362 | 1 |
| WBGene00001954 | hlh-10 | 0.831400225 | 0.695794526 | 0.19563767 | 0.127 | 0.205 | 0.126775135 | 1 |
| WBGene00008973 | F20D1.1 | 0.621554394 | 0.485575932 | 0.196175453 | 0.092 | 0.121 | 0.515539223 | 1 |
| WBGene00019109 | F59E11.2 | 0.568468272 | 0.432431494 | 0.196259585 | 0.092 | 0.1 | 0.96568378 | 1 |
| WBGene00016200 | dpff-1 | 0.482941764 | 0.346720731 | 0.196525409 | 0.104 | 0.079 | 0.340128898 | 1 |
| WBGene00004138 | pqn-53 | 0.660395117 | 0.524091462 | 0.196644607 | 0.139 | 0.168 | 0.767569895 | 1 |
| WBGene00011288 | R102.1 | 2.848145595 | 2.711728414 | 0.196808391 | 0.491 | 0.532 | 0.72642113 | 1 |
| WBGene00004388 | rnp-5 | 0.384060547 | 0.247445016 | 0.197094549 | 0.04 | 0.053 | 0.635050485 | 1 |
| WBGene00006734 | ufd-2 | 0.378660542 | 0.241952648 | 0.1972278 | 0.064 | 0.042 | 0.341589598 | 1 |
| WBGene00008407 | D2023.3 | 0.312286306 | 0.175541476 | 0.197281088 | 0.058 | 0.047 | 0.591168621 | 1 |
| WBGene00019399 | K04F10.7 | 0.716000738 | 0.579192626 | 0.197372385 | 0.116 | 0.137 | 0.712708493 | 1 |
| WBGene00019747 | ipla-1 | 0.353028253 | 0.215706333 | 0.198113652 | 0.035 | 0.063 | 0.245202704 | 1 |
| WBGene00018152 | acs-4 | 0.608235229 | 0.470882477 | 0.198158134 | 0.104 | 0.153 | 0.313587394 | 1 |
| WBGene00000482 | chd-3 | 1.0030294 | 0.865584803 | 0.198290638 | 0.145 | 0.263 | 0.043397447 | 1 |
| WBGene00004363 | ric-3 | 0.452079733 | 0.314058663 | 0.199122314 | 0.069 | 0.079 | 0.825761199 | 1 |
| WBGene00009208 | F28C6.9 | 1.2339369 | 1.09564 | 0.199520251 | 0.289 | 0.363 | 0.772397342 | 1 |
| WBGene00012434 | Y11D7A.10 | 0.409304979 | 0.270953285 | 0.199599302 | 0.069 | 0.084 | 0.736398746 | 1 |
| WBGene00012062 | T26G10.5 | 0.595796822 | 0.457393203 | 0.199674215 | 0.075 | 0.105 | 0.415451108 | 1 |
| WBGene00220183 | Y55F3BR.19 | 0.580573112 | 0.441901452 | 0.200060916 | 0.087 | 0.1 | 0.780727353 | 1 |
| WBGene00010137 | ztf-26 | 0.457005916 | 0.318220894 | 0.200224463 | 0.069 | 0.095 | 0.506817841 | 1 |
| WBGene00022749 | ZK484.3 | 0.438285763 | 0.299234411 | 0.200608696 | 0.075 | 0.074 | 0.864530891 | 1 |
| WBGene00003954 | pcm-1 | 0.993554832 | 0.854299487 | 0.200902995 | 0.191 | 0.237 | 0.667133224 | 1 |
| WBGene00021474 | dot-1.1 | 0.483452301 | 0.344130726 | 0.200998546 | 0.075 | 0.095 | 0.625095909 | 1 |
| WBGene00012648 | Y39A1A.9 | 0.758541634 | 0.619218986 | 0.201000095 | 0.156 | 0.189 | 0.767874713 | 1 |

| | | | | | | | | |
|---|---|---|---|---|---|---|---|---|
| WBGene00006708 | ubc-13 | 0.540704628 | 0.401312693 | 0.201100053 | 0.11 | 0.1 | 0.620102594 | 1 |
| WBGene00013468 | srz-96 | 0.35284881 | 0.21335117 | 0.201252554 | 0.069 | 0.053 | 0.456476518 | 1 |
| WBGene00006776 | unc-40 | 0.376117902 | 0.236220514 | 0.201829268 | 0.04 | 0.074 | 0.214495448 | 1 |
| WBGene00001258 | emb-4 | 0.637828055 | 0.497911766 | 0.201856536 | 0.081 | 0.142 | 0.117707361 | 1 |
| WBGene00004220 | ptr-5 | 1.653583501 | 1.51319641 | 0.20253576 | 0.358 | 0.5 | 0.307327731 | 1 |
| WBGene00011709 | T11F9.1 | 0.613837316 | 0.473409051 | 0.202595162 | 0.081 | 0.142 | 0.116854504 | 1 |
| WBGene00001118 | dyf-2 | 0.950781648 | 0.810317469 | 0.202646973 | 0.168 | 0.237 | 0.315104941 | 1 |
| WBGene00003368 | mkk-4 | 0.708281665 | 0.567732513 | 0.202769564 | 0.121 | 0.132 | 0.979954729 | 1 |
| WBGene00020447 | mob-3 | 0.638803908 | 0.49825377 | 0.202770987 | 0.127 | 0.142 | 0.950253454 | 1 |
| WBGene00007058 | dmd-6 | 2.570856716 | 2.430213176 | 0.202905737 | 0.688 | 0.816 | 0.750625198 | 1 |
| WBGene00000374 | cyp-31A1 | 0.41336827 | 0.272032536 | 0.203904362 | 0.064 | 0.079 | 0.694206945 | 1 |
| WBGene00000585 | cogc-2 | 0.3130644 | 0.171667713 | 0.203992299 | 0.052 | 0.053 | 0.930727955 | 1 |
| WBGene00002265 | lec-2 | 1.443212332 | 1.301607713 | 0.204292282 | 0.301 | 0.405 | 0.422361779 | 1 |
| WBGene00021281 | ell-1 | 0.247787561 | 0.105749738 | 0.204917263 | 0.052 | 0.032 | 0.303232807 | 1 |
| WBGene00013645 | Y105C5B.1 | 0.389802656 | 0.247518919 | 0.205272042 | 0.081 | 0.058 | 0.351965845 | 1 |
| WBGene00018117 | nlp-47 | 1.977281413 | 1.834759952 | 0.205615006 | 0.474 | 0.668 | 0.266645045 | 1 |
| WBGene00009323 | best-13 | 1.131397042 | 0.988584554 | 0.206034868 | 0.208 | 0.321 | 0.105727251 | 1 |
| WBGene00005938 | srx-47 | 0.275218147 | 0.132386753 | 0.206062143 | 0.052 | 0.026 | 0.191101431 | 1 |
| WBGene00019401 | nuo-4 | 0.43952871 | 0.296552655 | 0.206270846 | 0.092 | 0.074 | 0.438364029 | 1 |
| WBGene00020420 | T10E10.3 | 1.187314523 | 1.04415274 | 0.206538795 | 0.185 | 0.321 | 0.034109407 | 1 |
| WBGene00001325 | eor-2 | 0.380429752 | 0.236754408 | 0.207279706 | 0.046 | 0.058 | 0.679908241 | 1 |
| WBGene00007967 | cyp-25A4 | 0.382013701 | 0.237987366 | 0.207786079 | 0.035 | 0.074 | 0.12891929 | 1 |
| WBGene00007939 | C34F6.5 | 0.470676631 | 0.326649924 | 0.207786615 | 0.064 | 0.084 | 0.53400838 | 1 |
| WBGene00008343 | C56A3.4 | 0.547602283 | 0.403056576 | 0.208535375 | 0.075 | 0.111 | 0.357411116 | 1 |
| WBGene00001853 | hil-2 | 0.554354443 | 0.409533448 | 0.208932531 | 0.075 | 0.084 | 0.850470016 | 1 |
| WBGene00008143 | C47E8.4 | 0.357393141 | 0.21221336 | 0.20945015 | 0.058 | 0.053 | 0.750592032 | 1 |
| WBGene00004502 | rpt-2 | 0.666393414 | 0.52105277 | 0.209682227 | 0.127 | 0.153 | 0.752811353 | 1 |
| WBGene00003868 | ooc-3 | 0.505907157 | 0.360454907 | 0.209843241 | 0.069 | 0.084 | 0.680050598 | 1 |
| WBGene00020839 | usp-5 | 0.283928177 | 0.138289075 | 0.21011281 | 0.052 | 0.042 | 0.590888017 | 1 |
| WBGene00010349 | dyf-18 | 0.559253062 | 0.413117395 | 0.210829202 | 0.11 | 0.126 | 0.825566726 | 1 |
| WBGene00022374 | nhr-277 | 2.291473679 | 2.145193785 | 0.211037279 | 0.52 | 0.684 | 0.478308407 | 1 |
| WBGene00020831 | T26C12.1 | 0.457030615 | 0.310711472 | 0.211093902 | 0.058 | 0.079 | 0.508236628 | 1 |
| WBGene00012000 | T24H10.1 | 0.319534175 | 0.173096156 | 0.211265403 | 0.058 | 0.047 | 0.60016111 | 1 |
| WBGene00017852 | F27C1.2 | 1.08789624 | 0.941266909 | 0.211541408 | 0.179 | 0.305 | 0.042057859 | 1 |
| WBGene00000074 | adm-2 | 0.405133779 | 0.25834026 | 0.211778282 | 0.058 | 0.074 | 0.649428415 | 1 |
| WBGene00022580 | iglr-2 | 0.469451275 | 0.322100151 | 0.212582737 | 0.092 | 0.079 | 0.552502742 | 1 |
| WBGene00021810 | phf-15 | 0.43545455 | 0.288039335 | 0.2126752 | 0.087 | 0.084 | 0.789034559 | 1 |
| WBGene00011471 | T05C12.8 | 1.769150196 | 1.62160446 | 0.212863502 | 0.457 | 0.605 | 0.781995618 | 1 |
| WBGene00019733 | M02E1.1 | 0.986958069 | 0.839335694 | 0.212974068 | 0.185 | 0.2 | 0.998839533 | 1 |
| WBGene00018797 | ept-1 | 0.318989656 | 0.171231936 | 0.21316933 | 0.052 | 0.042 | 0.608000966 | 1 |
| WBGene00007016 | mdt-15 | 1.012514148 | 0.864381029 | 0.213710917 | 0.179 | 0.253 | 0.280441164 | 1 |
| WBGene00014569 | bkip-1 | 0.679790339 | 0.531635878 | 0.213741705 | 0.133 | 0.158 | 0.735702772 | 1 |
| WBGene00005315 | srh-95 | 0.377814725 | 0.229582427 | 0.213854001 | 0.04 | 0.068 | 0.288015901 | 1 |
| WBGene00001951 | hlh-4 | 2.510392929 | 2.362026355 | 0.214047721 | 0.665 | 0.758 | 0.494880768 | 1 |
| WBGene00005263 | srh-40 | 0.806785641 | 0.657917736 | 0.214770987 | 0.133 | 0.2 | 0.254839854 | 1 |
| WBGene00018508 | F46F11.1 | 0.723434471 | 0.574500931 | 0.214865679 | 0.092 | 0.174 | 0.056950272 | 1 |
| WBGene00045381 | F28B1.9 | 0.378416655 | 0.229295584 | 0.21513623 | 0.052 | 0.037 | 0.459805644 | 1 |
| WBGene00008366 | tpra-1 | 0.467997021 | 0.318745126 | 0.215324968 | 0.069 | 0.089 | 0.603232313 | 1 |
| WBGene00015678 | mcp-1 | 0.855680904 | 0.7063141 | 0.215490747 | 0.162 | 0.216 | 0.483505317 | 1 |
| WBGene00000897 | daf-1 | 0.343179937 | 0.193589908 | 0.215812794 | 0.04 | 0.053 | 0.646796835 | 1 |
| WBGene00015043 | cyp-34A8 | 0.497960478 | 0.347871791 | 0.216532204 | 0.052 | 0.074 | 0.454284177 | 1 |
| WBGene00045295 | fbxa-222 | 0.348802574 | 0.198674939 | 0.216588395 | 0.052 | 0.058 | 0.900230155 | 1 |
| WBGene00005328 | srh-109 | 0.372072935 | 0.221701676 | 0.216939869 | 0.046 | 0.058 | 0.691353469 | 1 |

| | | | | | | | | |
|---|---|---|---|---|---|---|---|---|
| WBGene00004920 | snr-7 | 1.135818861 | 0.98522188 | 0.217265518 | 0.237 | 0.337 | 0.356247503 | 1 |
| WBGene00008649 | F10D11.3 | 0.943309497 | 0.792703054 | 0.217279169 | 0.162 | 0.232 | 0.270889188 | 1 |
| WBGene00011638 | ostb-1 | 0.750903899 | 0.599941308 | 0.217792982 | 0.121 | 0.168 | 0.359009726 | 1 |
| WBGene00020347 | ech-1.2 | 0.610139539 | 0.459060555 | 0.217960901 | 0.098 | 0.137 | 0.403122378 | 1 |
| WBGene00005111 | srd-33 | 1.144139791 | 0.992980824 | 0.218076291 | 0.243 | 0.274 | 0.99377443 | 1 |
| WBGene00016093 | srsx-34 | 0.796834116 | 0.645591801 | 0.218196539 | 0.173 | 0.174 | 0.6931728 | 1 |
| WBGene00003690 | nhr-100 | 0.785352519 | 0.634066521 | 0.218259559 | 0.127 | 0.195 | 0.205540994 | 1 |
| WBGene00023491 | tsp-21 | 1.553520016 | 1.402067041 | 0.218500455 | 0.318 | 0.479 | 0.160555025 | 1 |
| WBGene00004189 | pars-1 | 0.377338346 | 0.225446122 | 0.219134158 | 0.058 | 0.068 | 0.796003794 | 1 |
| WBGene00021929 | dcap-1 | 0.337321892 | 0.185394331 | 0.219185138 | 0.04 | 0.053 | 0.648763249 | 1 |
| WBGene00006483 | dgk-4 | 0.936755201 | 0.784756609 | 0.219287615 | 0.133 | 0.189 | 0.244186574 | 1 |
| WBGene00010279 | letm-1 | 0.255377232 | 0.10333433 | 0.21935154 | 0.052 | 0.026 | 0.192122513 | 1 |
| WBGene00011240 | mask-1 | 0.945200629 | 0.793054333 | 0.219500706 | 0.179 | 0.247 | 0.35933481 | 1 |
| WBGene00022748 | oaz-1 | 2.020641672 | 1.868412399 | 0.219620417 | 0.543 | 0.653 | 0.634740613 | 1 |
| WBGene00195143 | nlp-78 | 1.41731486 | 1.264634801 | 0.220270764 | 0.353 | 0.411 | 0.98173019 | 1 |
| WBGene00001227 | eif-3.D | 0.531474417 | 0.378645742 | 0.220485172 | 0.104 | 0.111 | 0.952457832 | 1 |
| WBGene00020051 | mfsd-6 | 0.631782793 | 0.478770419 | 0.220750194 | 0.11 | 0.137 | 0.636053631 | 1 |
| WBGene00044995 | Y105C5A.28 | 0.356258277 | 0.203004576 | 0.221098355 | 0.046 | 0.053 | 0.842779777 | 1 |
| WBGene00008777 | F13H10.6 | 3.145231903 | 2.991509519 | 0.221774522 | 0.85 | 0.916 | 0.072888424 | 1 |
| WBGene00000140 | anc-1 | 1.550301373 | 1.396433572 | 0.221984312 | 0.301 | 0.389 | 0.482671232 | 1 |
| WBGene00013703 | rbm-12 | 0.36924573 | 0.215376347 | 0.221986596 | 0.058 | 0.063 | 0.923794045 | 1 |
| WBGene00001207 | egl-43 | 0.881715988 | 0.72781575 | 0.222031111 | 0.145 | 0.205 | 0.310583474 | 1 |
| WBGene00008281 | C53D6.4 | 0.814735919 | 0.660699643 | 0.222227372 | 0.11 | 0.147 | 0.412363849 | 1 |
| WBGene00015022 | B0205.8 | 0.388378763 | 0.233974858 | 0.222757747 | 0.046 | 0.068 | 0.432670859 | 1 |
| WBGene00018184 | F38E9.1 | 0.662743732 | 0.508014077 | 0.223227705 | 0.104 | 0.137 | 0.510482676 | 1 |
| WBGene00005668 | sru-5 | 0.371079322 | 0.215954039 | 0.223798477 | 0.04 | 0.063 | 0.386820615 | 1 |
| WBGene00006149 | str-90 | 0.95444575 | 0.799311584 | 0.223811291 | 0.145 | 0.2 | 0.355005806 | 1 |
| WBGene00268206 | F54E2.9 | 1.906142086 | 1.750947864 | 0.223897934 | 0.457 | 0.626 | 0.309725731 | 1 |
| WBGene00017197 | F07C3.9 | 0.49693774 | 0.341584194 | 0.224127791 | 0.081 | 0.089 | 0.920999732 | 1 |
| WBGene00005031 | sra-5 | 0.600309713 | 0.444199904 | 0.225218848 | 0.087 | 0.116 | 0.504397358 | 1 |
| WBGene00020549 | nmt-1 | 0.356289945 | 0.200156602 | 0.2252528 | 0.052 | 0.058 | 0.89822546 | 1 |
| WBGene00009189 | F27D4.4 | 0.377460859 | 0.22128871 | 0.225308785 | 0.064 | 0.053 | 0.586923126 | 1 |
| WBGene00010794 | dld-1 | 0.371674377 | 0.215133769 | 0.225840358 | 0.04 | 0.058 | 0.498058581 | 1 |
| WBGene00004207 | ptb-1 | 1.475913327 | 1.319284865 | 0.225967105 | 0.295 | 0.489 | 0.065201068 | 1 |
| WBGene00012929 | rsks-1 | 0.39170376 | 0.234951216 | 0.226146119 | 0.04 | 0.053 | 0.638955928 | 1 |
| WBGene00007703 | gbf-1 | 0.677978649 | 0.521193472 | 0.226193196 | 0.098 | 0.126 | 0.539735124 | 1 |
| WBGene00008622 | F09C8.2 | 0.420002791 | 0.263031594 | 0.226461567 | 0.069 | 0.068 | 0.874920295 | 1 |
| WBGene00005325 | srh-106 | 0.373633016 | 0.216592983 | 0.226560877 | 0.064 | 0.058 | 0.728209264 | 1 |
| WBGene00006911 | vha-2 | 1.233709436 | 1.076662348 | 0.226571055 | 0.272 | 0.337 | 0.607401685 | 1 |
| WBGene00017979 | F32B5.6 | 0.519465667 | 0.361869901 | 0.22736263 | 0.069 | 0.095 | 0.452118378 | 1 |
| WBGene00015921 | C17H11.1 | 1.378827877 | 1.221052595 | 0.227621617 | 0.289 | 0.353 | 0.615673532 | 1 |
| WBGene00016115 | mdt-26 | 1.957471746 | 1.799596571 | 0.227765733 | 0.434 | 0.589 | 0.227826883 | 1 |
| WBGene00022522 | ZC132.3 | 0.836087629 | 0.678030881 | 0.228027686 | 0.156 | 0.195 | 0.64181855 | 1 |
| WBGene00003415 | mars-1 | 0.30190459 | 0.143753052 | 0.228164441 | 0.058 | 0.042 | 0.441658121 | 1 |
| WBGene00022077 | vamp-7 | 0.663268589 | 0.504641967 | 0.22884984 | 0.121 | 0.147 | 0.740392582 | 1 |
| WBGene00003609 | nhr-10 | 0.573134728 | 0.414049009 | 0.229512178 | 0.098 | 0.111 | 0.854194751 | 1 |
| WBGene00019163 | ubxn-6 | 0.304834795 | 0.144948781 | 0.23066676 | 0.052 | 0.037 | 0.434564313 | 1 |
| WBGene00011418 | igdb-1 | 0.433069433 | 0.272638203 | 0.23145334 | 0.069 | 0.084 | 0.747852308 | 1 |
| WBGene00001577 | gem-4 | 0.835327397 | 0.674188239 | 0.232474663 | 0.121 | 0.137 | 0.802753724 | 1 |
| WBGene00016603 | met-1 | 0.872395969 | 0.711245479 | 0.232491013 | 0.145 | 0.179 | 0.66302151 | 1 |
| WBGene00004056 | pmk-2 | 0.748122867 | 0.58687404 | 0.232632884 | 0.121 | 0.168 | 0.36935319 | 1 |
| WBGene00016215 | glb-10 | 0.985429036 | 0.824174487 | 0.232641139 | 0.156 | 0.221 | 0.292581007 | 1 |
| WBGene00017826 | tag-340 | 0.293070771 | 0.131674158 | 0.232846093 | 0.052 | 0.042 | 0.59277848 | 1 |

| | | | | | | | | |
|---|---|---|---|---|---|---|---|---|
| WBGene00019388 | K04F1.9 | 0.707326113 | 0.545872683 | 0.232928062 | 0.116 | 0.158 | 0.429617248 | 1 |
| WBGene00022452 | Y110A2AR.1 | 0.402687892 | 0.241079233 | 0.233152012 | 0.064 | 0.063 | 0.887775141 | 1 |
| WBGene00001648 | goa-1 | 1.017589176 | 0.855833235 | 0.233364494 | 0.214 | 0.263 | 0.654533819 | 1 |
| WBGene00008453 | E02A10.3 | 2.608374929 | 2.446373566 | 0.233718564 | 0.723 | 0.858 | 0.355518641 | 1 |
| WBGene00009674 | nucb-1 | 0.377516393 | 0.215489702 | 0.233755104 | 0.058 | 0.058 | 0.911312855 | 1 |
| WBGene00000086 | aex-3 | 0.753011218 | 0.590780808 | 0.234049007 | 0.121 | 0.153 | 0.610743548 | 1 |
| WBGene00185118 | K02E2.11 | 1.087047039 | 0.924567208 | 0.234408847 | 0.243 | 0.295 | 0.82707358 | 1 |
| WBGene00021563 | Y45G12B.2 | 0.419947604 | 0.257376184 | 0.234540981 | 0.064 | 0.068 | 0.961983642 | 1 |
| WBGene00005892 | srx-1 | 0.663045024 | 0.500211902 | 0.234918536 | 0.116 | 0.142 | 0.680327495 | 1 |
| WBGene00020251 | camt-1 | 0.751982432 | 0.588717731 | 0.235541175 | 0.11 | 0.121 | 0.956143304 | 1 |
| WBGene00004751 | sea-2 | 1.273525056 | 1.110011306 | 0.235900476 | 0.266 | 0.321 | 0.807535127 | 1 |
| WBGene00000006 | aat-5 | 0.906979614 | 0.743396518 | 0.236000521 | 0.162 | 0.226 | 0.364506894 | 1 |
| WBGene00018625 | prp-17 | 0.436610784 | 0.27279174 | 0.236340923 | 0.069 | 0.074 | 1 | 1 |
| WBGene00006964 | xrn-2 | 0.425641955 | 0.261769751 | 0.236417617 | 0.04 | 0.074 | 0.217250187 | 1 |
| WBGene00001684 | gpd-2 | 1.493810441 | 1.329784213 | 0.236639825 | 0.318 | 0.463 | 0.249423714 | 1 |
| WBGene00019995 | R10F2.4 | 1.453622786 | 1.2892691 | 0.237112248 | 0.306 | 0.421 | 0.42491005 | 1 |
| WBGene00011735 | hip-1 | 0.766395775 | 0.601568539 | 0.237795436 | 0.127 | 0.168 | 0.504543079 | 1 |
| WBGene00008939 | F18H3.1 | 0.343406001 | 0.17841805 | 0.238027298 | 0.058 | 0.053 | 0.750592032 | 1 |
| WBGene00015561 | C07A12.7 | 0.996931782 | 0.831931466 | 0.238045137 | 0.173 | 0.232 | 0.312030374 | 1 |
| WBGene00007886 | ethe-1 | 0.447745263 | 0.282641802 | 0.238193945 | 0.075 | 0.068 | 0.717509123 | 1 |
| WBGene00019012 | F57C9.6 | 0.794943447 | 0.628655095 | 0.23990338 | 0.098 | 0.121 | 0.577318426 | 1 |
| WBGene00023475 | srz-69 | 0.340754789 | 0.17428343 | 0.240167403 | 0.052 | 0.042 | 0.623395287 | 1 |
| WBGene00006528 | tba-1 | 0.946744177 | 0.78001414 | 0.240540598 | 0.173 | 0.258 | 0.235562387 | 1 |
| WBGene00022647 | slc-17.1 | 0.707531567 | 0.540740045 | 0.240629301 | 0.092 | 0.168 | 0.080025887 | 1 |
| WBGene00005341 | srh-123 | 1.164878552 | 0.998057816 | 0.240671448 | 0.202 | 0.289 | 0.224702558 | 1 |
| WBGene00019115 | nhr-195 | 0.367855727 | 0.199317932 | 0.243148641 | 0.058 | 0.053 | 0.746752574 | 1 |
| WBGene00008311 | rskn-2 | 0.336124376 | 0.167443046 | 0.243355718 | 0.052 | 0.053 | 0.941046578 | 1 |
| WBGene00016739 | pitr-1 | 0.606414869 | 0.437004677 | 0.244407244 | 0.127 | 0.121 | 0.657416239 | 1 |
| WBGene00009051 | nduf-6 | 0.353769409 | 0.184197529 | 0.244640511 | 0.064 | 0.053 | 0.571681391 | 1 |
| WBGene00004245 | puf-9 | 0.708389474 | 0.538570162 | 0.24499748 | 0.092 | 0.137 | 0.294412578 | 1 |
| WBGene00021970 | srz-64 | 0.595470887 | 0.425497042 | 0.245220423 | 0.064 | 0.1 | 0.270400828 | 1 |
| WBGene00017350 | F10E7.9 | 1.038986988 | 0.868921042 | 0.245353296 | 0.185 | 0.289 | 0.150225532 | 1 |
| WBGene00005301 | srh-80 | 0.361576261 | 0.191198033 | 0.245803825 | 0.035 | 0.053 | 0.452995517 | 1 |
| WBGene00022170 | cox-6B | 1.185060055 | 1.014575956 | 0.245956564 | 0.243 | 0.332 | 0.424101808 | 1 |
| WBGene00004704 | rsp-7 | 0.497201774 | 0.326126972 | 0.246808768 | 0.075 | 0.095 | 0.636846933 | 1 |
| WBGene00016384 | cdgs-1 | 0.878663879 | 0.707454128 | 0.247003458 | 0.156 | 0.189 | 0.667039594 | 1 |
| WBGene00007217 | tads-1 | 0.534045423 | 0.362450097 | 0.247559726 | 0.11 | 0.111 | 0.798597045 | 1 |
| WBGene00003966 | pdl-1 | 0.898687309 | 0.727000632 | 0.247691516 | 0.173 | 0.247 | 0.330181582 | 1 |
| WBGene00005393 | srh-178 | 0.685277888 | 0.513443947 | 0.247903974 | 0.116 | 0.132 | 0.862498464 | 1 |
| WBGene00017605 | F19F10.9 | 1.17921753 | 1.007178602 | 0.248199709 | 0.243 | 0.3 | 0.704555975 | 1 |
| WBGene00007772 | egrh-1 | 0.398314657 | 0.225746738 | 0.248962881 | 0.064 | 0.068 | 0.984232178 | 1 |
| WBGene00007030 | epc-1 | 1.288189648 | 1.115539719 | 0.249081197 | 0.26 | 0.342 | 0.37995403 | 1 |
| WBGene00019492 | K07E3.4 | 0.668059875 | 0.495389318 | 0.249110957 | 0.098 | 0.137 | 0.390691081 | 1 |
| WBGene00020107 | R151.2 | 0.464497537 | 0.291707044 | 0.249283987 | 0.087 | 0.074 | 0.538852053 | 1 |
| WBGene00044452 | Y102E9.5 | 0.732060503 | 0.558860145 | 0.249875297 | 0.081 | 0.132 | 0.18713395 | 1 |
| WBGene00014001 | pyk-2 | 1.553372236 | 1.379947523 | 0.250198973 | 0.387 | 0.453 | 0.959530003 | 1 |
| WBGene00004147 | larp-5 | 0.828291374 | 0.654796633 | 0.250300001 | 0.15 | 0.211 | 0.366445413 | 1 |
| WBGene00016415 | ampd-1 | 0.702704321 | 0.529173241 | 0.250352428 | 0.121 | 0.147 | 0.723898135 | 1 |
| WBGene00015901 | nhr-158 | 1.572161556 | 1.398283551 | 0.250852936 | 0.341 | 0.453 | 0.341600523 | 1 |
| WBGene00001423 | fib-1 | 0.451779824 | 0.277864973 | 0.250906092 | 0.058 | 0.068 | 0.756027795 | 1 |
| WBGene00021333 | Y34D9A.8 | 0.742559319 | 0.568141752 | 0.251631359 | 0.098 | 0.147 | 0.26026503 | 1 |
| WBGene00044132 | C23H4.8 | 1.170537442 | 0.995514942 | 0.252504092 | 0.243 | 0.311 | 0.676604236 | 1 |
| WBGene00017652 | F21C10.1 | 0.963208986 | 0.788067933 | 0.25267513 | 0.168 | 0.232 | 0.356547301 | 1 |

| | | | | | | | | |
|---|---|---|---|---|---|---|---|---|
| WBGene00019607 | K10B2.4 | 1.12012255 | 0.94493425 | 0.252743292 | 0.26 | 0.295 | 0.79934804 | 1 |
| WBGene00016761 | tkr-2 | 1.870012939 | 1.694434762 | 0.253305765 | 0.399 | 0.574 | 0.242742182 | 1 |
| WBGene00007363 | stdh-1 | 0.58971245 | 0.413485602 | 0.254241599 | 0.087 | 0.074 | 0.576048576 | 1 |
| WBGene00005481 | srh-276 | 1.189659886 | 1.013252965 | 0.25450139 | 0.179 | 0.279 | 0.125127565 | 1 |
| WBGene00004913 | snn-1 | 0.544945658 | 0.368210812 | 0.254974486 | 0.069 | 0.111 | 0.24167928 | 1 |
| WBGene00000793 | crh-1 | 1.669180381 | 1.49200166 | 0.255614862 | 0.37 | 0.537 | 0.326789598 | 1 |
| WBGene00019408 | K05F1.6 | 0.411132807 | 0.233898731 | 0.255694722 | 0.064 | 0.063 | 0.895274626 | 1 |
| WBGene00003149 | mbk-1 | 0.510010058 | 0.332482929 | 0.256117508 | 0.069 | 0.084 | 0.705610888 | 1 |
| WBGene00045207 | F13E9.14 | 0.376339049 | 0.197963925 | 0.257340906 | 0.058 | 0.058 | 0.911312855 | 1 |
| WBGene00016278 | erp-44.1 | 0.441317863 | 0.262581828 | 0.257861592 | 0.046 | 0.074 | 0.33574282 | 1 |
| WBGene00004739 | scd-1 | 1.180346737 | 1.00134707 | 0.258241932 | 0.266 | 0.295 | 0.792706421 | 1 |
| WBGene00018259 | F41B4.3 | 0.438607725 | 0.259453539 | 0.258464856 | 0.081 | 0.063 | 0.442925059 | 1 |
| WBGene00020726 | T23C6.4 | 0.914690205 | 0.735525033 | 0.258480705 | 0.133 | 0.189 | 0.275478529 | 1 |
| WBGene00019457 | vms-1 | 0.377952398 | 0.198554775 | 0.258816061 | 0.064 | 0.047 | 0.44660436 | 1 |
| WBGene00016016 | C23G10.10 | 1.330327637 | 1.150641956 | 0.259231642 | 0.312 | 0.316 | 0.486755427 | 1 |
| WBGene00004128 | pqn-41 | 0.833369771 | 0.653451962 | 0.25956653 | 0.162 | 0.205 | 0.614793553 | 1 |
| WBGene00003642 | nhr-52 | 0.468904248 | 0.288931738 | 0.259645448 | 0.087 | 0.084 | 0.787444429 | 1 |
| WBGene00012645 | mrpl-22 | 0.986992121 | 0.806902502 | 0.2598144 | 0.168 | 0.2 | 0.681266848 | 1 |
| WBGene00003928 | pas-7 | 0.480594772 | 0.300482033 | 0.259847756 | 0.075 | 0.079 | 0.987046125 | 1 |
| WBGene00021349 | arl-13 | 1.167800341 | 0.98764949 | 0.259902738 | 0.283 | 0.342 | 1 | 1 |
| WBGene00000547 | clp-7 | 0.493085393 | 0.312644866 | 0.260320654 | 0.058 | 0.084 | 0.412907711 | 1 |
| WBGene00011280 | pelo-1 | 0.482538203 | 0.301713914 | 0.260874306 | 0.069 | 0.079 | 0.818910239 | 1 |
| WBGene00003559 | ncl-1 | 0.485945876 | 0.305017665 | 0.261024232 | 0.081 | 0.079 | 0.827993376 | 1 |
| WBGene00044476 | F56D6.14 | 1.252364487 | 1.071252918 | 0.261288763 | 0.243 | 0.316 | 0.474929457 | 1 |
| WBGene00010591 | K05D4.9 | 0.439807541 | 0.258220031 | 0.2619754 | 0.064 | 0.063 | 0.889649097 | 1 |
| WBGene00006565 | tfg-1 | 0.778727496 | 0.597086752 | 0.2620522 | 0.139 | 0.205 | 0.290882309 | 1 |
| WBGene00005560 | sri-48 | 0.440179049 | 0.25843878 | 0.262195785 | 0.069 | 0.063 | 0.728139057 | 1 |
| WBGene00199110 | C32D5.17 | 0.395148734 | 0.213403214 | 0.26220336 | 0.035 | 0.068 | 0.17889803 | 1 |
| WBGene00004053 | tank-1 | 0.826904455 | 0.644535827 | 0.263102315 | 0.121 | 0.168 | 0.355602761 | 1 |
| WBGene00005438 | srh-230 | 3.108679027 | 2.925774656 | 0.263875228 | 0.884 | 0.921 | 0.000357177 | 1 |
| WBGene00022302 | Y76B12C.8 | 2.36946187 | 2.186491002 | 0.263971164 | 0.642 | 0.8 | 0.702783661 | 1 |
| WBGene00011329 | zipt-9 | 0.376525124 | 0.193381381 | 0.264220569 | 0.058 | 0.053 | 0.760217548 | 1 |
| WBGene00015176 | vps-51 | 0.617159388 | 0.433805992 | 0.264523035 | 0.11 | 0.105 | 0.718459683 | 1 |
| WBGene00019565 | cyp-35A3 | 0.712360441 | 0.528919036 | 0.264650005 | 0.11 | 0.142 | 0.536743584 | 1 |
| WBGene00007816 | C30F2.2 | 2.254207538 | 2.07073122 | 0.264700375 | 0.659 | 0.7 | 0.289620468 | 1 |
| WBGene00012897 | pisy-1 | 0.748030766 | 0.564354277 | 0.264989159 | 0.127 | 0.142 | 0.821242196 | 1 |
| WBGene00022829 | ZK867.2 | 0.579786503 | 0.396093957 | 0.265012325 | 0.092 | 0.105 | 0.844616867 | 1 |
| WBGene00005535 | sri-23 | 0.827103515 | 0.643333636 | 0.265123893 | 0.156 | 0.184 | 0.806332663 | 1 |
| WBGene00021637 | Y47G6A.7 | 0.403801146 | 0.219925211 | 0.265276899 | 0.052 | 0.068 | 0.606830247 | 1 |
| WBGene00271808 | F21C10.19 | 0.688335536 | 0.504411417 | 0.265346415 | 0.092 | 0.126 | 0.397244705 | 1 |
| WBGene00011510 | pdha-1 | 0.392134705 | 0.208078775 | 0.265536577 | 0.069 | 0.058 | 0.565990081 | 1 |
| WBGene00018037 | chtl-1 | 1.623072634 | 1.438626618 | 0.266099352 | 0.312 | 0.421 | 0.334192769 | 1 |
| WBGene00004132 | ifet-1 | 0.548249039 | 0.363033376 | 0.267209719 | 0.058 | 0.1 | 0.184479966 | 1 |
| WBGene00007809 | C29F4.3 | 2.155155033 | 1.96984044 | 0.267352444 | 0.59 | 0.726 | 0.944213051 | 1 |
| WBGene00001527 | gcs-1 | 0.46512416 | 0.279665818 | 0.267559831 | 0.064 | 0.079 | 0.670373702 | 1 |
| WBGene00016378 | imp-3 | 0.446488448 | 0.260057872 | 0.268962467 | 0.04 | 0.058 | 0.493006756 | 1 |
| WBGene00019871 | R04E5.8 | 0.768994814 | 0.582440914 | 0.269140386 | 0.127 | 0.168 | 0.4881564 | 1 |
| WBGene00005334 | srh-115 | 0.595952008 | 0.409205862 | 0.269417738 | 0.081 | 0.105 | 0.555570256 | 1 |
| WBGene00003639 | nhr-49 | 1.083788695 | 0.896822955 | 0.269734546 | 0.214 | 0.263 | 0.720459792 | 1 |
| WBGene00015798 | C15H9.2 | 0.453576869 | 0.266276816 | 0.270216858 | 0.075 | 0.084 | 0.900829072 | 1 |
| WBGene00008310 | C54E10.6 | 0.453389399 | 0.265949984 | 0.270417915 | 0.075 | 0.079 | 0.96115203 | 1 |
| WBGene00011499 | T05G5.4 | 0.393755665 | 0.205986132 | 0.270894174 | 0.035 | 0.063 | 0.248461696 | 1 |
| WBGene00044009 | ZK822.6 | 3.139678157 | 2.95158388 | 0.271362681 | 0.85 | 0.926 | 0.140169902 | 1 |

| | | | | | | | | |
|---|---|---|---|---|---|---|---|---|
| WBGene00000475 | cey-4 | 0.853641645 | 0.665345514 | 0.271653895 | 0.15 | 0.205 | 0.409888964 | 1 |
| WBGene00045386 | nlp-76 | 1.55047831 | 1.362004919 | 0.271909626 | 0.243 | 0.332 | 0.242165232 | 1 |
| WBGene00021965 | Y57G7A.1 | 0.378188165 | 0.189084547 | 0.272818851 | 0.035 | 0.053 | 0.451301926 | 1 |
| WBGene00019628 | pup-2 | 0.498619319 | 0.309056712 | 0.273481033 | 0.075 | 0.079 | 0.988773186 | 1 |
| WBGene00012607 | Y38F1A.4 | 2.001791453 | 1.812221265 | 0.27349197 | 0.474 | 0.642 | 0.357408126 | 1 |
| WBGene00020008 | R11F4.2 | 0.51659193 | 0.326409617 | 0.274375079 | 0.087 | 0.079 | 0.674111456 | 1 |
| WBGene00005378 | srh-162 | 0.748110243 | 0.557899866 | 0.274415568 | 0.11 | 0.158 | 0.335221745 | 1 |
| WBGene00016165 | wdr-60 | 1.25575681 | 1.065265749 | 0.274820509 | 0.243 | 0.311 | 0.569196225 | 1 |
| WBGene00005305 | srh-84 | 0.570154516 | 0.379516957 | 0.275031861 | 0.092 | 0.084 | 0.69654703 | 1 |
| WBGene00017198 | nhr-36 | 0.854366163 | 0.663665936 | 0.275122272 | 0.173 | 0.179 | 0.772776899 | 1 |
| WBGene00001426 | fkb-1 | 0.640409711 | 0.449604114 | 0.275274289 | 0.11 | 0.126 | 0.845074084 | 1 |
| WBGene00020301 | T07D1.2 | 0.546384083 | 0.355136531 | 0.275911895 | 0.087 | 0.095 | 0.943165151 | 1 |
| WBGene00005542 | sri-30 | 1.792348859 | 1.601080793 | 0.27594149 | 0.37 | 0.516 | 0.297686976 | 1 |
| WBGene00002129 | inx-7 | 0.467280502 | 0.275897207 | 0.276107731 | 0.069 | 0.074 | 0.979447075 | 1 |
| WBGene00000065 | act-3 | 0.611850174 | 0.420418002 | 0.276178246 | 0.075 | 0.105 | 0.399534059 | 1 |
| WBGene00004109 | stam-1 | 0.729168039 | 0.537578426 | 0.276405384 | 0.116 | 0.163 | 0.378773088 | 1 |
| WBGene00050939 | C05G5.7 | 2.353846623 | 2.161825543 | 0.27702786 | 0.405 | 0.474 | 0.926650481 | 1 |
| WBGene00005555 | sri-43 | 1.567916041 | 1.375714995 | 0.277287496 | 0.312 | 0.453 | 0.266715231 | 1 |
| WBGene00011279 | asd-1 | 1.024540798 | 0.83230175 | 0.277342321 | 0.214 | 0.279 | 0.600857685 | 1 |
| WBGene00011110 | prdx-3 | 0.324418044 | 0.132027168 | 0.277561362 | 0.058 | 0.037 | 0.314074776 | 1 |
| WBGene00008975 | F20D1.3 | 0.936198413 | 0.743307111 | 0.278283325 | 0.191 | 0.184 | 0.568576883 | 1 |
| WBGene00005300 | srh-79 | 0.647265237 | 0.45435086 | 0.278316615 | 0.104 | 0.116 | 0.94345359 | 1 |
| WBGene00018238 | F40F4.7 | 0.392136374 | 0.19898376 | 0.278660319 | 0.064 | 0.063 | 0.878414971 | 1 |
| WBGene00004336 | ret-1 | 0.976471043 | 0.783218642 | 0.278804281 | 0.197 | 0.221 | 0.983709333 | 1 |
| WBGene00015018 | srz-85 | 1.927069962 | 1.733582509 | 0.279143389 | 0.497 | 0.611 | 0.764330941 | 1 |
| WBGene00012967 | kvs-4 | 1.817124549 | 1.623520475 | 0.279311637 | 0.462 | 0.637 | 0.747774112 | 1 |
| WBGene00000379 | cct-4 | 0.743780119 | 0.550032006 | 0.279519442 | 0.127 | 0.174 | 0.422152974 | 1 |
| WBGene00005521 | sri-9 | 0.39811072 | 0.204220596 | 0.279724321 | 0.058 | 0.047 | 0.598357668 | 1 |
| WBGene00195084 | C43C3.4 | 0.959251266 | 0.765244611 | 0.279892439 | 0.197 | 0.211 | 0.847113231 | 1 |
| WBGene00010736 | istr-1 | 0.475387308 | 0.281276637 | 0.280042503 | 0.087 | 0.074 | 0.538852053 | 1 |
| WBGene00009042 | F22B5.4 | 1.047698042 | 0.853165839 | 0.280650645 | 0.208 | 0.221 | 0.841981882 | 1 |
| WBGene00001043 | dnj-25 | 0.485384383 | 0.290046723 | 0.281812673 | 0.075 | 0.089 | 0.787540937 | 1 |
| WBGene00018788 | shc-1 | 0.96542818 | 0.770077055 | 0.281832099 | 0.156 | 0.216 | 0.338138474 | 1 |
| WBGene00021953 | Y55H10A.2 | 2.574694547 | 2.378866917 | 0.28251955 | 0.624 | 0.763 | 0.673006228 | 1 |
| WBGene00010960 | atp-6 | 3.187421763 | 2.991305962 | 0.282935294 | 0.879 | 0.921 | 0.133630103 | 1 |
| WBGene00023068 | rpl-41.2 | 3.781203557 | 3.584672532 | 0.283534336 | 0.908 | 0.963 | 0.003700277 | 1 |
| WBGene00014079 | ZK792.7 | 0.868821398 | 0.672213632 | 0.28364505 | 0.15 | 0.205 | 0.411167028 | 1 |
| WBGene00004783 | seu-1 | 0.498860486 | 0.301965048 | 0.284060072 | 0.069 | 0.068 | 0.867734252 | 1 |
| WBGene00022675 | gbb-2 | 0.54456048 | 0.347656876 | 0.284071853 | 0.075 | 0.089 | 0.745911203 | 1 |
| WBGene00045208 | F13E9.15 | 4.391914561 | 4.194988702 | 0.28410396 | 0.977 | 0.995 | 0.004362274 | 1 |
| WBGene00006651 | tts-2 | 1.502784769 | 1.305761349 | 0.284244712 | 0.341 | 0.426 | 0.977666138 | 1 |
| WBGene00022381 | ubxn-2 | 1.220824426 | 1.023731689 | 0.284344714 | 0.214 | 0.332 | 0.135456841 | 1 |
| WBGene00003082 | lsm-8 | 0.443993392 | 0.246772845 | 0.284529106 | 0.069 | 0.063 | 0.715949304 | 1 |
| WBGene00003836 | nxt-1 | 0.435524334 | 0.238095289 | 0.284829904 | 0.046 | 0.074 | 0.333317008 | 1 |
| WBGene00077486 | sls-1.9 | 0.530219069 | 0.33236318 | 0.285445709 | 0.035 | 0.068 | 0.16584645 | 1 |
| WBGene00006747 | unc-7 | 0.675914412 | 0.47743738 | 0.28634183 | 0.11 | 0.142 | 0.566266609 | 1 |
| WBGene00012288 | sre-32 | 0.52041488 | 0.321869476 | 0.28644047 | 0.075 | 0.063 | 0.558439232 | 1 |
| WBGene00003391 | mog-3 | 0.567884591 | 0.368983425 | 0.286953725 | 0.075 | 0.116 | 0.279943767 | 1 |
| WBGene00010964 | ctc-1 | 2.469956722 | 2.270946255 | 0.287111414 | 0.624 | 0.774 | 0.802448384 | 1 |
| WBGene00007683 | C18E9.2 | 1.36859388 | 1.168996847 | 0.28795765 | 0.283 | 0.368 | 0.618857897 | 1 |
| WBGene00020092 | pcf-11 | 0.469709883 | 0.270041168 | 0.288061065 | 0.081 | 0.089 | 0.95386669 | 1 |
| WBGene00008040 | ttr-5 | 0.794466257 | 0.59457585 | 0.288380899 | 0.156 | 0.168 | 0.91866488 | 1 |
| WBGene00007173 | B0393.8 | 0.375660626 | 0.175735597 | 0.288430848 | 0.064 | 0.053 | 0.561626178 | 1 |

| | | | | | | | | |
|---|---|---|---|---|---|---|---|---|
| WBGene00016432 | C35B1.2 | 0.555900997 | 0.355975817 | 0.288431066 | 0.075 | 0.1 | 0.517902356 | 1 |
| WBGene00002189 | kin-1 | 1.220849408 | 1.020037618 | 0.289710174 | 0.283 | 0.342 | 0.88629815 | 1 |
| WBGene00016471 | srab-8 | 0.669728506 | 0.468873413 | 0.289772648 | 0.092 | 0.116 | 0.61943783 | 1 |
| WBGene00006229 | str-187 | 0.457246967 | 0.255949299 | 0.290411148 | 0.081 | 0.053 | 0.236161108 | 1 |
| WBGene00018572 | lin-42 | 2.009434941 | 1.807638799 | 0.291130293 | 0.439 | 0.511 | 0.777209315 | 1 |
| WBGene00002783 | let-607 | 0.706698842 | 0.504561769 | 0.291622153 | 0.145 | 0.168 | 0.922516209 | 1 |
| WBGene00014004 | fic-1 | 0.597879031 | 0.394691148 | 0.293138151 | 0.104 | 0.132 | 0.646501877 | 1 |
| WBGene00021024 | W04C9.2 | 0.297124716 | 0.093913324 | 0.293172067 | 0.058 | 0.021 | 0.064650119 | 1 |
| WBGene00003885 | osm-5 | 1.604943525 | 1.401703465 | 0.293213427 | 0.399 | 0.489 | 0.94217119 | 1 |
| WBGene00015012 | bath-20 | 1.293786461 | 1.090077668 | 0.293889665 | 0.249 | 0.321 | 0.661842297 | 1 |
| WBGene00194780 | C12D8.20 | 0.388666115 | 0.184899594 | 0.29397295 | 0.075 | 0.053 | 0.32091294 | 1 |
| WBGene00018637 | F49E8.6 | 2.093454953 | 1.889173035 | 0.294716511 | 0.538 | 0.616 | 0.455963881 | 1 |
| WBGene00018764 | F53F10.2 | 1.605192917 | 1.400502328 | 0.295306098 | 0.358 | 0.479 | 0.690526704 | 1 |
| WBGene00018714 | F52H2.1 | 0.396102571 | 0.19138194 | 0.295349439 | 0.052 | 0.047 | 0.786990731 | 1 |
| WBGene00014025 | asna-1 | 0.441070042 | 0.236320901 | 0.29539057 | 0.064 | 0.063 | 0.882157075 | 1 |
| WBGene00000908 | daf-12 | 0.410620801 | 0.205565031 | 0.295832942 | 0.058 | 0.047 | 0.601967007 | 1 |
| WBGene00016157 | ari-1.2 | 0.515074632 | 0.309704821 | 0.296286007 | 0.087 | 0.095 | 0.960758553 | 1 |
| WBGene00219721 | linc-104 | 1.442024015 | 1.236276763 | 0.296830541 | 0.324 | 0.389 | 0.913522325 | 1 |
| WBGene00007223 | C01F6.9 | 0.528521752 | 0.322708953 | 0.296925105 | 0.087 | 0.089 | 0.903555966 | 1 |
| WBGene00000224 | atgp-1 | 0.779831541 | 0.573174667 | 0.298142847 | 0.116 | 0.163 | 0.36195161 | 1 |
| WBGene00005422 | srh-212 | 1.035228188 | 0.828192209 | 0.29868978 | 0.225 | 0.247 | 0.813079437 | 1 |
| WBGene00013278 | Y57A10B.6 | 0.464829174 | 0.257782717 | 0.298704898 | 0.064 | 0.053 | 0.592045161 | 1 |
| WBGene00012472 | Y17G7B.21 | 0.607901866 | 0.400239104 | 0.299594037 | 0.098 | 0.111 | 0.907719297 | 1 |
| WBGene00008757 | F13E9.8 | 2.138588892 | 1.930673649 | 0.29995829 | 0.59 | 0.647 | 0.205851924 | 1 |
| WBGene00004184 | T19D2.2 | 1.472183935 | 1.264133618 | 0.300153161 | 0.387 | 0.447 | 0.520995369 | 1 |
| WBGene00012163 | VZK822L.2 | 0.464107541 | 0.255266618 | 0.301293764 | 0.075 | 0.074 | 0.828333764 | 1 |
| WBGene00004699 | rsp-2 | 0.937441521 | 0.728545902 | 0.301372674 | 0.185 | 0.226 | 0.778191218 | 1 |
| WBGene00009732 | cope-1 | 1.066879425 | 0.857774917 | 0.301674036 | 0.168 | 0.279 | 0.097238459 | 1 |
| WBGene00007408 | C07B5.3 | 2.669542312 | 2.460370165 | 0.301771618 | 0.775 | 0.858 | 0.257394325 | 1 |
| WBGene00005342 | srh-125 | 1.665473712 | 1.456208771 | 0.301905492 | 0.364 | 0.532 | 0.170421826 | 1 |
| WBGene00018252 | F40H3.6 | 2.053227571 | 1.84331651 | 0.302837646 | 0.312 | 0.411 | 0.339334962 | 1 |
| WBGene00000161 | apa-2 | 0.811128432 | 0.601186277 | 0.302882506 | 0.121 | 0.179 | 0.294863215 | 1 |
| WBGene00013111 | sta-1 | 1.624437913 | 1.414488433 | 0.302893073 | 0.387 | 0.458 | 0.991561145 | 1 |
| WBGene00021888 | manf-1 | 0.814544917 | 0.604191218 | 0.30347624 | 0.156 | 0.163 | 0.766055954 | 1 |
| WBGene00001538 | gcy-12 | 0.550080565 | 0.339617012 | 0.303634724 | 0.081 | 0.084 | 0.960690624 | 1 |
| WBGene00019900 | vdac-1 | 1.821558646 | 1.610728939 | 0.304162973 | 0.445 | 0.574 | 0.962620785 | 1 |
| WBGene00002002 | hsb-1 | 0.642943366 | 0.431950198 | 0.304398797 | 0.104 | 0.137 | 0.547471912 | 1 |
| WBGene00011681 | T10B10.4 | 0.750201284 | 0.539171584 | 0.304451502 | 0.121 | 0.158 | 0.527696167 | 1 |
| WBGene00006698 | uaf-2 | 0.763032073 | 0.551032387 | 0.305850896 | 0.139 | 0.153 | 1 | 1 |
| WBGene00004186 | prpf-4 | 0.446535957 | 0.234415548 | 0.306025062 | 0.058 | 0.058 | 0.919167812 | 1 |
| WBGene00012258 | W04G3.5 | 0.401033094 | 0.188758999 | 0.306246785 | 0.064 | 0.058 | 0.717323209 | 1 |
| WBGene00007006 | npr-3 | 0.937590166 | 0.725098358 | 0.306560877 | 0.162 | 0.189 | 0.770470009 | 1 |
| WBGene00012973 | spat-2 | 0.83699211 | 0.624447769 | 0.306636666 | 0.145 | 0.174 | 0.672187656 | 1 |
| WBGene00021395 | pnc-1 | 1.603512887 | 1.39095337 | 0.306658561 | 0.382 | 0.5 | 0.727950969 | 1 |
| WBGene00016905 | ztf-3 | 0.381238349 | 0.168485247 | 0.306937845 | 0.052 | 0.053 | 0.928665571 | 1 |
| WBGene00005552 | sri-40 | 1.518398853 | 1.305397979 | 0.307295304 | 0.376 | 0.437 | 0.59147302 | 1 |
| WBGene00018145 | F37C4.5 | 1.048392343 | 0.835364632 | 0.307334023 | 0.197 | 0.247 | 0.755095695 | 1 |
| WBGene00019564 | K09D9.1 | 0.641865408 | 0.428484809 | 0.307843131 | 0.075 | 0.074 | 0.878399754 | 1 |
| WBGene00044557 | ZC13.10 | 0.496669738 | 0.28229284 | 0.309280487 | 0.069 | 0.079 | 0.861934944 | 1 |
| WBGene00015152 | B0348.2 | 0.283502933 | 0.068988599 | 0.309478765 | 0.058 | 0.016 | 0.028974253 | 1 |
| WBGene00007345 | C05E7.3 | 3.580584723 | 3.366035886 | 0.309528543 | 0.931 | 0.984 | 0.022551974 | 1 |
| WBGene00015126 | B0303.4 | 0.868847441 | 0.654208123 | 0.30965908 | 0.127 | 0.158 | 0.6341384 | 1 |
| WBGene00022503 | ZC13.2 | 1.560372911 | 1.345596757 | 0.309856492 | 0.364 | 0.426 | 0.991407229 | 1 |

| | | | | | | | | |
|---|---|---|---|---|---|---|---|---|
| WBGene00003516 | nab-1 | 0.462922061 | 0.247960757 | 0.310123608 | 0.069 | 0.079 | 0.879267844 | 1 |
| WBGene00014745 | F26D2.8 | 0.836919622 | 0.621732789 | 0.310448977 | 0.15 | 0.163 | 0.941375146 | 1 |
| WBGene00003406 | mrg-1 | 0.523278047 | 0.30767312 | 0.311052159 | 0.075 | 0.1 | 0.541911088 | 1 |
| WBGene00009534 | F38B7.3 | 2.73161156 | 2.515491131 | 0.311795872 | 0.769 | 0.811 | 0.215899597 | 1 |
| WBGene00022435 | cfap-36 | 0.569031746 | 0.352889622 | 0.311827171 | 0.069 | 0.1 | 0.398083329 | 1 |
| WBGene00013557 | Y75B8A.24 | 0.410688078 | 0.194492376 | 0.311904467 | 0.069 | 0.042 | 0.231415921 | 1 |
| WBGene00011090 | R07B7.6 | 0.314724912 | 0.097642605 | 0.313183567 | 0.064 | 0.026 | 0.076296226 | 1 |
| WBGene00009326 | F32D8.1 | 0.753582732 | 0.535786622 | 0.314213367 | 0.127 | 0.158 | 0.660122346 | 1 |
| WBGene00012135 | T28F3.9 | 0.489423531 | 0.271270611 | 0.314728135 | 0.092 | 0.074 | 0.424853644 | 1 |
| WBGene00012035 | T26C5.5 | 2.271307899 | 2.052850845 | 0.315166908 | 0.555 | 0.595 | 0.310376483 | 1 |
| WBGene00044699 | nhr-286 | 0.836720377 | 0.618257783 | 0.315174902 | 0.197 | 0.168 | 0.303138222 | 1 |
| WBGene00013466 | srz-47 | 0.669602683 | 0.450441704 | 0.316182458 | 0.133 | 0.121 | 0.55169754 | 1 |
| WBGene00006801 | unc-68 | 1.139546036 | 0.920297753 | 0.316308411 | 0.191 | 0.263 | 0.369599039 | 1 |
| WBGene00006755 | unc-16 | 0.694242082 | 0.474942331 | 0.316382664 | 0.116 | 0.142 | 0.709497251 | 1 |
| WBGene00004887 | smn-1 | 0.397094076 | 0.177775734 | 0.316409484 | 0.046 | 0.058 | 0.70286966 | 1 |
| WBGene00019521 | dmd-7 | 0.27487823 | 0.055132229 | 0.317026466 | 0.052 | 0.016 | 0.049496627 | 1 |
| WBGene00001135 | eat-4 | 0.546632343 | 0.326800086 | 0.317150907 | 0.092 | 0.1 | 0.98985826 | 1 |
| WBGene00003915 | pan-1 | 1.556294574 | 1.336245018 | 0.317464404 | 0.318 | 0.411 | 0.596097062 | 1 |
| WBGene00044283 | spp-22 | 0.789025305 | 0.568961029 | 0.317485639 | 0.156 | 0.137 | 0.413481329 | 1 |
| WBGene00019494 | K07E8.1 | 0.426221377 | 0.205872131 | 0.317896764 | 0.064 | 0.047 | 0.45420412 | 1 |
| WBGene00016345 | C33E10.1 | 0.854186747 | 0.63315033 | 0.318888143 | 0.139 | 0.142 | 0.850123598 | 1 |
| WBGene00017837 | F26G1.1 | 0.456384384 | 0.234058993 | 0.320747739 | 0.046 | 0.074 | 0.338790848 | 1 |
| WBGene00017536 | F17A9.4 | 0.87700512 | 0.654648157 | 0.320793289 | 0.185 | 0.184 | 0.645287001 | 1 |
| WBGene00001080 | dpy-21 | 0.966715271 | 0.744156345 | 0.321084659 | 0.162 | 0.216 | 0.456095861 | 1 |
| WBGene00016650 | ubr-4 | 1.013371346 | 0.789678024 | 0.322721247 | 0.191 | 0.232 | 0.709381822 | 1 |
| WBGene00003080 | lsm-6 | 0.571404379 | 0.347079894 | 0.323631822 | 0.087 | 0.095 | 0.947960965 | 1 |
| WBGene00019953 | wapl-1 | 0.370929185 | 0.14616988 | 0.324259135 | 0.064 | 0.037 | 0.215239427 | 1 |
| WBGene00020142 | aak-2 | 0.931325232 | 0.706285041 | 0.324664367 | 0.156 | 0.216 | 0.372981082 | 1 |
| WBGene00009116 | snx-27 | 0.789591647 | 0.564407246 | 0.324872419 | 0.139 | 0.179 | 0.58965475 | 1 |
| WBGene00007010 | alx-1 | 0.797861312 | 0.572314715 | 0.325394957 | 0.15 | 0.158 | 0.806789532 | 1 |
| WBGene00009294 | F31D4.9 | 0.643019745 | 0.417439237 | 0.325443879 | 0.121 | 0.126 | 0.830308084 | 1 |
| WBGene00006372 | syx-2 | 0.657544731 | 0.431941562 | 0.325476573 | 0.075 | 0.132 | 0.13823323 | 1 |
| WBGene00015913 | C17F4.7 | 0.721963106 | 0.49596934 | 0.326040085 | 0.121 | 0.111 | 0.539249319 | 1 |
| WBGene00017367 | nhr-263 | 0.879803132 | 0.653336006 | 0.326723 | 0.15 | 0.174 | 0.868742621 | 1 |
| WBGene00044646 | B0205.14 | 1.362872457 | 1.13635202 | 0.32679991 | 0.283 | 0.311 | 0.880829185 | 1 |
| WBGene00020090 | R119.5 | 0.429263567 | 0.202688721 | 0.326878407 | 0.04 | 0.063 | 0.386820615 | 1 |
| WBGene00019890 | R05F9.6 | 0.347761151 | 0.121163394 | 0.326911461 | 0.052 | 0.037 | 0.429613825 | 1 |
| WBGene00009660 | F43G6.8 | 1.726993252 | 1.500389007 | 0.326920821 | 0.41 | 0.526 | 0.653698531 | 1 |
| WBGene00020287 | lsy-13 | 0.520211233 | 0.293598315 | 0.326933333 | 0.081 | 0.079 | 0.819703231 | 1 |
| WBGene00020694 | dhhc-11 | 2.223932564 | 1.997169831 | 0.32714947 | 0.509 | 0.611 | 0.992068467 | 1 |
| WBGene00005172 | srg-15 | 1.892905999 | 1.666055493 | 0.3272761 | 0.491 | 0.521 | 0.275976982 | 1 |
| WBGene00021732 | Y49G5B.1 | 0.827925503 | 0.600627897 | 0.327921128 | 0.104 | 0.163 | 0.192433541 | 1 |
| WBGene00044701 | AH9.6 | 0.75487577 | 0.527494615 | 0.328041665 | 0.133 | 0.132 | 0.797270325 | 1 |
| WBGene00005365 | srh-149 | 0.580656447 | 0.353051029 | 0.328365208 | 0.081 | 0.068 | 0.565583163 | 1 |
| WBGene00020682 | T22D1.3 | 0.473366097 | 0.245578662 | 0.328627803 | 0.046 | 0.063 | 0.544122287 | 1 |
| WBGene00008930 | F18A11.3 | 0.880459576 | 0.652637315 | 0.328678045 | 0.191 | 0.211 | 0.86651641 | 1 |
| WBGene00015091 | B0261.1 | 0.390453083 | 0.161844014 | 0.329813171 | 0.046 | 0.053 | 0.863660503 | 1 |
| WBGene00003884 | osm-3 | 0.768454974 | 0.53965345 | 0.330090824 | 0.139 | 0.158 | 0.919459214 | 1 |
| WBGene00013989 | ZK512.11 | 0.377338248 | 0.148299319 | 0.330433328 | 0.064 | 0.037 | 0.213274771 | 1 |
| WBGene00007730 | C25G4.2 | 0.420862076 | 0.191752798 | 0.330534819 | 0.064 | 0.047 | 0.436087272 | 1 |
| WBGene00269386 | F41F3.10 | 0.958717129 | 0.728391285 | 0.332289952 | 0.15 | 0.163 | 0.962157622 | 1 |
| WBGene00006005 | srx-114 | 0.78815455 | 0.556506337 | 0.334197727 | 0.139 | 0.137 | 0.698768692 | 1 |
| WBGene00020029 | R12C12.9 | 1.152014322 | 0.920052066 | 0.334650797 | 0.225 | 0.263 | 0.87875003 | 1 |

| | | | | | | | | |
|---|---|---|---|---|---|---|---|---|
| WBGene00011677 | cyp-13A1 | 0.464464042 | 0.232496367 | 0.334658613 | 0.069 | 0.053 | 0.44484664 | 1 |
| WBGene00012179 | W01D2.1 | 2.744361036 | 2.511942847 | 0.335308568 | 0.786 | 0.879 | 0.080808791 | 1 |
| WBGene00004321 | rcan-1 | 1.07389156 | 0.841455977 | 0.335333663 | 0.173 | 0.253 | 0.228119928 | 1 |
| WBGene00006311 | sun-1 | 0.447306368 | 0.214691965 | 0.335591646 | 0.064 | 0.058 | 0.742809371 | 1 |
| WBGene00017016 | snap-1 | 1.058753707 | 0.824452411 | 0.338025319 | 0.191 | 0.263 | 0.384303303 | 1 |
| WBGene00018512 | trpp-8 | 0.649916414 | 0.415143976 | 0.338705031 | 0.104 | 0.121 | 0.845401431 | 1 |
| WBGene00005360 | srh-144 | 0.360990621 | 0.126173963 | 0.338768828 | 0.058 | 0.032 | 0.203597855 | 1 |
| WBGene00018758 | F53E10.1 | 0.631618273 | 0.396358867 | 0.339407578 | 0.087 | 0.111 | 0.592070058 | 1 |
| WBGene00000829 | ctb-1 | 1.770653531 | 1.535371386 | 0.339440384 | 0.376 | 0.479 | 0.568065855 | 1 |
| WBGene00003014 | lin-28 | 1.305484002 | 1.069593048 | 0.34031871 | 0.277 | 0.321 | 0.937975212 | 1 |
| WBGene00000123 | ama-1 | 0.776145497 | 0.539888758 | 0.340846427 | 0.127 | 0.158 | 0.696220672 | 1 |
| WBGene00019301 | K02D10.1 | 0.713167792 | 0.476808602 | 0.340994231 | 0.139 | 0.142 | 0.729021042 | 1 |
| WBGene00008021 | C39B10.1 | 0.510356653 | 0.273283162 | 0.34202475 | 0.04 | 0.068 | 0.294966852 | 1 |
| WBGene00016626 | clhm-1 | 1.007517134 | 0.768255943 | 0.345180934 | 0.197 | 0.226 | 0.979879981 | 1 |
| WBGene00045161 | lfor-2 | 0.714872576 | 0.475088963 | 0.345934629 | 0.127 | 0.116 | 0.588172798 | 1 |
| WBGene00021355 | Y37E3.17 | 0.988125229 | 0.748200212 | 0.346138633 | 0.202 | 0.237 | 0.88201138 | 1 |
| WBGene00019402 | syf-2 | 0.593762973 | 0.35304635 | 0.347280679 | 0.092 | 0.084 | 0.683043061 | 1 |
| WBGene00000396 | cdh-4 | 0.842204846 | 0.601486662 | 0.34728293 | 0.156 | 0.153 | 0.653784605 | 1 |
| WBGene00007553 | C13G3.1 | 4.125997936 | 3.884603138 | 0.348259078 | 0.792 | 0.826 | 0.102466132 | 1 |
| WBGene00000896 | dad-1 | 1.249480163 | 1.007538522 | 0.349048005 | 0.283 | 0.374 | 0.703471578 | 1 |
| WBGene00010326 | F59C6.5 | 0.340122147 | 0.098168723 | 0.349065005 | 0.069 | 0.026 | 0.046723633 | 1 |
| WBGene00022800 | ZK688.5 | 0.512349974 | 0.270348827 | 0.349133854 | 0.064 | 0.068 | 0.954573182 | 1 |
| WBGene00003728 | nhr-138 | 1.441394234 | 1.199281718 | 0.349294526 | 0.353 | 0.395 | 0.661500076 | 1 |
| WBGene00012125 | T28D6.5 | 0.503929433 | 0.261696608 | 0.349468096 | 0.069 | 0.074 | 0.997318892 | 1 |
| WBGene00005096 | srd-18 | 0.721653493 | 0.478049137 | 0.351446796 | 0.081 | 0.121 | 0.296789634 | 1 |
| WBGene00006730 | uev-1 | 0.795944285 | 0.551836991 | 0.352172383 | 0.133 | 0.153 | 0.873307501 | 1 |
| WBGene00022280 | Y74C10AL.2 | 0.525174452 | 0.280539519 | 0.352933604 | 0.064 | 0.068 | 0.949017877 | 1 |
| WBGene00011582 | ntr-1 | 0.610258187 | 0.365554093 | 0.353033383 | 0.075 | 0.089 | 0.723004193 | 1 |
| WBGene00022865 | ufbp-1 | 0.455841826 | 0.210384037 | 0.354120736 | 0.064 | 0.047 | 0.455732773 | 1 |
| WBGene00011045 | gpx-2 | 0.936739071 | 0.691199661 | 0.354238489 | 0.191 | 0.211 | 0.902654575 | 1 |
| WBGene00011146 | pde-2 | 1.49007348 | 1.244455652 | 0.354351623 | 0.318 | 0.411 | 0.578924235 | 1 |
| WBGene00019673 | K12C11.1 | 0.633933132 | 0.388105641 | 0.354654102 | 0.087 | 0.111 | 0.610888661 | 1 |
| WBGene00018080 | F36A4.1 | 0.556843029 | 0.308916363 | 0.357682571 | 0.092 | 0.079 | 0.512490392 | 1 |
| WBGene00002997 | lin-8 | 0.536421478 | 0.287743174 | 0.358766956 | 0.046 | 0.089 | 0.139938249 | 1 |
| WBGene00003221 | mes-3 | 0.46283611 | 0.214093699 | 0.358859443 | 0.058 | 0.053 | 0.746752574 | 1 |
| WBGene00017504 | F16B4.2 | 0.498282998 | 0.249411491 | 0.359045689 | 0.081 | 0.058 | 0.316583103 | 1 |
| WBGene00010683 | K08F8.7 | 0.481562514 | 0.232435662 | 0.359414074 | 0.075 | 0.063 | 0.543204283 | 1 |
| WBGene00000552 | cmd-1 | 2.712787481 | 2.463098148 | 0.360225562 | 0.763 | 0.816 | 0.206893082 | 1 |
| WBGene00016367 | nhr-162 | 0.751685119 | 0.501981081 | 0.360246777 | 0.121 | 0.126 | 0.864561539 | 1 |
| WBGene00002696 | let-504 | 0.42386051 | 0.173635363 | 0.360998579 | 0.058 | 0.047 | 0.589377583 | 1 |
| WBGene00001065 | dpy-3 | 0.844457732 | 0.593914252 | 0.361457836 | 0.139 | 0.111 | 0.328105378 | 1 |
| WBGene00008671 | F11A5.3 | 1.178118633 | 0.927259855 | 0.361912714 | 0.231 | 0.247 | 0.814385193 | 1 |
| WBGene00007365 | C06B3.6 | 2.11951719 | 1.868568128 | 0.362042967 | 0.468 | 0.532 | 0.594874717 | 1 |
| WBGene00022667 | ZK154.5 | 0.815985977 | 0.564678818 | 0.362559592 | 0.121 | 0.158 | 0.563496219 | 1 |
| WBGene00005413 | srh-202 | 1.038763709 | 0.786637446 | 0.363741309 | 0.208 | 0.221 | 0.736358252 | 1 |
| WBGene00022552 | ZC196.9 | 0.740697527 | 0.488399202 | 0.363989541 | 0.104 | 0.153 | 0.326113583 | 1 |
| WBGene00009264 | sac-1 | 0.63277366 | 0.380401748 | 0.364095706 | 0.058 | 0.105 | 0.141218731 | 1 |
| WBGene00015866 | C16E9.2 | 0.407535132 | 0.153398844 | 0.366641163 | 0.052 | 0.047 | 0.760495736 | 1 |
| WBGene00003153 | mca-3 | 1.204259733 | 0.949724733 | 0.367216381 | 0.225 | 0.311 | 0.382562421 | 1 |
| WBGene00022817 | upp-1 | 1.330662648 | 1.075997265 | 0.367404486 | 0.324 | 0.368 | 0.621346021 | 1 |
| WBGene00004466 | rpn-10 | 0.822623693 | 0.567509617 | 0.368051812 | 0.145 | 0.132 | 0.537896555 | 1 |
| WBGene00001037 | dnj-19 | 1.0286374 | 0.773462234 | 0.368139945 | 0.208 | 0.263 | 0.807871622 | 1 |
| WBGene00014300 | D2023.1 | 0.630379333 | 0.374987683 | 0.368452267 | 0.069 | 0.084 | 0.705610888 | 1 |

| | | | | | | | | |
|---|---|---|---|---|---|---|---|---|
| WBGene00011026 | R05D7.1 | 0.496514518 | 0.240839714 | 0.368860773 | 0.081 | 0.058 | 0.329992866 | 1 |
| WBGene00006780 | unc-44 | 1.487655091 | 1.23133671 | 0.369789258 | 0.277 | 0.305 | 0.997512873 | 1 |
| WBGene00000502 | chp-1 | 0.764804662 | 0.507398371 | 0.37135878 | 0.139 | 0.147 | 0.827571505 | 1 |
| WBGene00011230 | nud-2 | 0.688508795 | 0.430887221 | 0.371669367 | 0.11 | 0.1 | 0.616781503 | 1 |
| WBGene00010037 | F54C8.4 | 1.432176792 | 1.173088622 | 0.373785217 | 0.329 | 0.411 | 0.906699783 | 1 |
| WBGene00006923 | vhp-1 | 1.656619518 | 1.396806721 | 0.374830633 | 0.364 | 0.468 | 0.744686231 | 1 |
| WBGene00007366 | C06B3.7 | 1.234437823 | 0.974213373 | 0.375424524 | 0.237 | 0.205 | 0.316506921 | 1 |
| WBGene00000289 | cam-1 | 0.772313203 | 0.511891743 | 0.37570875 | 0.098 | 0.126 | 0.537328233 | 1 |
| WBGene00010958 | ndfl-4 | 1.480178612 | 1.219512695 | 0.376061427 | 0.295 | 0.411 | 0.410281487 | 1 |
| WBGene00007216 | C01A2.4 | 0.483724164 | 0.222883839 | 0.376313044 | 0.046 | 0.058 | 0.693267969 | 1 |
| WBGene00012043 | irld-52 | 0.58424058 | 0.323394648 | 0.376321133 | 0.052 | 0.089 | 0.220853859 | 1 |
| WBGene00021611 | nhr-238 | 0.534043957 | 0.273151404 | 0.376388392 | 0.092 | 0.068 | 0.325050715 | 1 |
| WBGene00002250 | lap-2 | 1.225626544 | 0.964280082 | 0.377043244 | 0.133 | 0.168 | 0.439527074 | 1 |
| WBGene00013124 | Y52B11A.4 | 0.469151773 | 0.207550642 | 0.377410655 | 0.058 | 0.047 | 0.60016111 | 1 |
| WBGene00019480 | K07C11.8 | 1.014268279 | 0.752109087 | 0.378215766 | 0.168 | 0.205 | 0.63723505 | 1 |
| WBGene00010632 | K07C5.9 | 3.326117181 | 3.063441629 | 0.378960716 | 0.879 | 0.932 | 0.0198386 | 1 |
| WBGene00016961 | vps-32.1 | 1.472012184 | 1.209309183 | 0.379000317 | 0.272 | 0.374 | 0.263066053 | 1 |
| WBGene00010071 | sre-45 | 0.760137077 | 0.497419627 | 0.379021162 | 0.11 | 0.142 | 0.556339595 | 1 |
| WBGene00194982 | Y17D7B.10 | 2.224562002 | 1.961727869 | 0.3791895 | 0.416 | 0.5 | 0.678843284 | 1 |
| WBGene00009783 | rer-1 | 0.843796478 | 0.580029248 | 0.380535674 | 0.127 | 0.137 | 0.959974628 | 1 |
| WBGene00003618 | nhr-19 | 0.530202327 | 0.266294805 | 0.380738072 | 0.064 | 0.074 | 0.823979353 | 1 |
| WBGene00219617 | linc-25 | 0.656703913 | 0.392425072 | 0.381273773 | 0.104 | 0.116 | 0.940521427 | 1 |
| WBGene00019394 | K04F10.1 | 0.46526576 | 0.200958815 | 0.38131432 | 0.075 | 0.042 | 0.15719735 | 1 |
| WBGene00006180 | str-131 | 1.260356461 | 0.995828605 | 0.381633027 | 0.225 | 0.3 | 0.486227774 | 1 |
| WBGene00044927 | F32B5.9 | 0.641711879 | 0.376908438 | 0.382030611 | 0.058 | 0.068 | 0.732701517 | 1 |
| WBGene00022864 | ZK1236.5 | 0.449062159 | 0.183917091 | 0.382523474 | 0.069 | 0.042 | 0.222649123 | 1 |
| WBGene00014679 | srz-25 | 0.485326994 | 0.220108581 | 0.38262929 | 0.052 | 0.063 | 0.744813968 | 1 |
| WBGene00003564 | ncs-2 | 1.514257231 | 1.248143386 | 0.383921126 | 0.329 | 0.442 | 0.630759711 | 1 |
| WBGene00011121 | R07E5.17 | 1.062643783 | 0.795849142 | 0.384903307 | 0.15 | 0.253 | 0.089732787 | 1 |
| WBGene00003901 | paa-1 | 1.03954722 | 0.772666309 | 0.385027767 | 0.185 | 0.242 | 0.580899495 | 1 |
| WBGene00010187 | F57B1.1 | 1.742386144 | 1.475368305 | 0.385225312 | 0.393 | 0.537 | 0.500224546 | 1 |
| WBGene00015175 | srz-4 | 2.148463028 | 1.881068371 | 0.385768946 | 0.48 | 0.574 | 0.830306047 | 1 |
| WBGene00009589 | F40F11.4 | 0.428955983 | 0.161431191 | 0.385956691 | 0.052 | 0.032 | 0.305968884 | 1 |
| WBGene00010574 | K04H4.5 | 0.887655529 | 0.619704575 | 0.386571513 | 0.133 | 0.147 | 0.937244491 | 1 |
| WBGene00015364 | npr-19 | 0.802689316 | 0.534518516 | 0.386888684 | 0.15 | 0.153 | 0.742077397 | 1 |
| WBGene00044143 | K09E9.4 | 0.642859577 | 0.374374677 | 0.387341834 | 0.116 | 0.105 | 0.547492406 | 1 |
| WBGene00010579 | K05C4.2 | 0.559138314 | 0.289934069 | 0.388379629 | 0.116 | 0.079 | 0.163223098 | 1 |
| WBGene00009373 | F34D10.3 | 0.452749286 | 0.182768991 | 0.389499233 | 0.075 | 0.053 | 0.304991398 | 1 |
| WBGene00006261 | str-230 | 0.819023862 | 0.548820546 | 0.389820984 | 0.168 | 0.153 | 0.420094805 | 1 |
| WBGene00003903 | pab-2 | 1.377017226 | 1.106543666 | 0.390210865 | 0.324 | 0.363 | 0.598147628 | 1 |
| WBGene00006462 | svh-5 | 0.754946043 | 0.483788404 | 0.39119778 | 0.116 | 0.132 | 0.899763815 | 1 |
| WBGene00016395 | spcs-1 | 0.824929065 | 0.553458555 | 0.391649159 | 0.185 | 0.158 | 0.245379587 | 1 |
| WBGene00021956 | Y57E12AL.1 | 0.804217073 | 0.532374638 | 0.392185734 | 0.145 | 0.168 | 0.902448162 | 1 |
| WBGene00001209 | egl-45 | 0.561743897 | 0.28984094 | 0.392273048 | 0.087 | 0.074 | 0.531839401 | 1 |
| WBGene00003889 | osm-9 | 0.779903417 | 0.50541728 | 0.395999788 | 0.104 | 0.142 | 0.417945388 | 1 |
| WBGene00002263 | lea-1 | 0.83078732 | 0.556294021 | 0.396010121 | 0.139 | 0.137 | 0.72994976 | 1 |
| WBGene00268192 | ZK697.17 | 1.263365646 | 0.98802783 | 0.397228501 | 0.243 | 0.284 | 0.984553778 | 1 |
| WBGene00006584 | tni-1 | 1.021619021 | 0.745290377 | 0.398657966 | 0.162 | 0.237 | 0.253642164 | 1 |
| WBGene00010957 | nduo-6 | 3.969883023 | 3.693522879 | 0.398703409 | 0.96 | 0.958 | 0.005114663 | 1 |
| WBGene00000231 | atx-2 | 0.533017878 | 0.256436486 | 0.399022603 | 0.064 | 0.068 | 0.963836796 | 1 |
| WBGene00007696 | tram-1 | 1.727504109 | 1.449847297 | 0.400574105 | 0.399 | 0.474 | 0.817907844 | 1 |
| WBGene00021187 | Y9D1A.2 | 1.678120348 | 1.40009124 | 0.401111215 | 0.353 | 0.389 | 0.749517933 | 1 |
| WBGene00020812 | acdh-7 | 0.684710027 | 0.406666521 | 0.401131987 | 0.116 | 0.105 | 0.582362864 | 1 |

| | | | | | | | | |
|---|---|---|---|---|---|---|---|---|
| WBGene00017066 | maco-1 | 2.001346011 | 1.723138125 | 0.401369138 | 0.503 | 0.611 | 0.704980437 | 1 |
| WBGene00010280 | rde-12 | 0.576807457 | 0.298204252 | 0.401939462 | 0.075 | 0.079 | 0.987046125 | 1 |
| WBGene00018393 | msra-1 | 0.600292682 | 0.320777911 | 0.403254574 | 0.081 | 0.079 | 0.806481308 | 1 |
| WBGene00206423 | Y41C4A.29 | 0.566764439 | 0.287213007 | 0.403307465 | 0.052 | 0.079 | 0.372567396 | 1 |
| WBGene00015687 | chdp-1 | 0.692392251 | 0.412516321 | 0.403775616 | 0.11 | 0.116 | 0.937541574 | 1 |
| WBGene00020033 | R12E2.7 | 0.553779928 | 0.273610643 | 0.404198838 | 0.052 | 0.032 | 0.318483204 | 1 |
| WBGene00011733 | T12D8.5 | 0.771290729 | 0.490332091 | 0.405337634 | 0.052 | 0.079 | 0.362677326 | 1 |
| WBGene00015868 | C16H3.3 | 0.791922467 | 0.510517177 | 0.405982017 | 0.15 | 0.174 | 0.99004893 | 1 |
| WBGene00015547 | ain-1 | 0.854043204 | 0.571760595 | 0.40724772 | 0.116 | 0.126 | 0.985904706 | 1 |
| WBGene00008550 | F07B10.4 | 0.941354275 | 0.657735295 | 0.409175697 | 0.162 | 0.158 | 0.665328535 | 1 |
| WBGene00001187 | egl-19 | 0.879457313 | 0.595027013 | 0.410346184 | 0.121 | 0.132 | 0.965446353 | 1 |
| WBGene00012660 | Y39A1A.24 | 0.570127131 | 0.285666262 | 0.410390285 | 0.081 | 0.079 | 0.791672605 | 1 |
| WBGene00015167 | B0403.3 | 1.346745993 | 1.06107171 | 0.412140871 | 0.277 | 0.321 | 0.862712558 | 1 |
| WBGene00000066 | act-4 | 1.773849172 | 1.48811944 | 0.412220867 | 0.41 | 0.495 | 0.975995195 | 1 |
| WBGene00003989 | pfn-1 | 0.848237269 | 0.561613196 | 0.413511129 | 0.121 | 0.121 | 0.757865688 | 1 |
| WBGene00001179 | egl-10 | 0.794837262 | 0.507986473 | 0.413838211 | 0.121 | 0.111 | 0.571280213 | 1 |
| WBGene00005444 | srh-237 | 0.499609909 | 0.212570005 | 0.414111046 | 0.069 | 0.053 | 0.427722753 | 1 |
| WBGene00005370 | srh-154 | 1.922730596 | 1.635187265 | 0.414837337 | 0.509 | 0.584 | 0.29514267 | 1 |
| WBGene00002008 | hsp-4 | 1.270982001 | 0.982776942 | 0.415792009 | 0.197 | 0.211 | 0.873941942 | 1 |
| WBGene00044353 | ZK418.10 | 1.806567123 | 1.518119752 | 0.416141592 | 0.422 | 0.537 | 0.828265516 | 1 |
| WBGene00004338 | rfc-2 | 1.010372969 | 0.720822872 | 0.417732488 | 0.179 | 0.168 | 0.523399725 | 1 |
| WBGene00004134 | myrf-1 | 1.535608661 | 1.245950443 | 0.417888474 | 0.329 | 0.332 | 0.409192841 | 1 |
| WBGene00015955 | C18B2.4 | 0.81898529 | 0.529184518 | 0.418094137 | 0.162 | 0.147 | 0.4410892 | 1 |
| WBGene00014175 | ZK971.1 | 0.771604447 | 0.481206642 | 0.418955473 | 0.133 | 0.153 | 0.946917579 | 1 |
| WBGene00020374 | serp-1.2 | 0.788654455 | 0.497490183 | 0.42006125 | 0.15 | 0.147 | 0.604097296 | 1 |
| WBGene00008142 | C47E8.3 | 0.500338269 | 0.208952401 | 0.420380948 | 0.058 | 0.047 | 0.598357668 | 1 |
| WBGene00011529 | T06D8.9 | 0.535412661 | 0.244003902 | 0.42041397 | 0.046 | 0.053 | 0.842779777 | 1 |
| WBGene00012764 | Y41E3.6 | 0.736127935 | 0.444462528 | 0.420784236 | 0.081 | 0.089 | 0.886635064 | 1 |
| WBGene00015611 | C08F1.8 | 0.6432519 | 0.351529789 | 0.420866043 | 0.087 | 0.063 | 0.342158382 | 1 |
| WBGene00012602 | Y38E10A.24 | 0.563806892 | 0.272025895 | 0.420950996 | 0.127 | 0.079 | 0.080607899 | 1 |
| WBGene00014255 | ZK1320.5 | 1.017700599 | 0.725659898 | 0.421325671 | 0.179 | 0.211 | 0.831637712 | 1 |
| WBGene00219950 | F46A8.13 | 1.851812701 | 1.558097016 | 0.423742162 | 0.353 | 0.405 | 0.864257467 | 1 |
| WBGene00010284 | aman-2 | 0.44759824 | 0.152778643 | 0.42533477 | 0.064 | 0.047 | 0.424244408 | 1 |
| WBGene00017557 | nep-11 | 0.931241094 | 0.635022841 | 0.427352604 | 0.139 | 0.168 | 0.682775566 | 1 |
| WBGene00002019 | hsp-16.48 | 1.237499275 | 0.939596569 | 0.429779871 | 0.197 | 0.253 | 0.585251202 | 1 |
| WBGene00021469 | Y39G10AR.11 | 0.852526915 | 0.554492875 | 0.429972231 | 0.145 | 0.163 | 0.998093111 | 1 |
| WBGene00003048 | lit-1 | 1.944653929 | 1.645867389 | 0.431057859 | 0.428 | 0.453 | 0.360146434 | 1 |
| WBGene00001685 | gpd-3 | 1.382200227 | 1.082742109 | 0.432026742 | 0.289 | 0.332 | 0.817099675 | 1 |
| WBGene00013228 | Y56A3A.7 | 1.009951785 | 0.710062834 | 0.432648303 | 0.173 | 0.184 | 0.922414826 | 1 |
| WBGene00023401 | ZK380.t2 | 1.198170372 | 0.897569949 | 0.433674739 | 0.052 | 0.068 | 0.565193315 | 1 |
| WBGene00006957 | wsp-1 | 0.517178278 | 0.216322442 | 0.434043222 | 0.052 | 0.063 | 0.740136083 | 1 |
| WBGene00010527 | srz-74 | 0.800172008 | 0.497989612 | 0.435957044 | 0.15 | 0.126 | 0.288848351 | 1 |
| WBGene00020256 | T05C3.6 | 0.980401255 | 0.677641465 | 0.436790048 | 0.231 | 0.179 | 0.078265577 | 1 |
| WBGene00008783 | F14B6.2 | 0.50088297 | 0.198005542 | 0.436959764 | 0.064 | 0.058 | 0.713707117 | 1 |
| WBGene00009574 | F40E10.6 | 1.114053424 | 0.811020048 | 0.43718475 | 0.197 | 0.247 | 0.582558168 | 1 |
| WBGene00012148 | inos-1 | 1.074068548 | 0.770155083 | 0.438454449 | 0.185 | 0.221 | 0.853246979 | 1 |
| WBGene00012287 | sre-33 | 0.536312539 | 0.231467407 | 0.439798559 | 0.052 | 0.074 | 0.481913512 | 1 |
| WBGene00000227 | atm-1 | 0.89281498 | 0.587753436 | 0.440110777 | 0.127 | 0.168 | 0.496313669 | 1 |
| WBGene00018772 | F53G12.4 | 0.547064727 | 0.241981098 | 0.440142638 | 0.087 | 0.068 | 0.413807643 | 1 |
| WBGene00023506 | srz-11 | 1.408091171 | 1.100405552 | 0.443896517 | 0.277 | 0.316 | 0.871551125 | 1 |
| WBGene00017243 | F08C6.5 | 0.783416269 | 0.475663848 | 0.443992891 | 0.133 | 0.142 | 0.870530829 | 1 |
| WBGene00003561 | ncr-1 | 1.122760899 | 0.814334437 | 0.444965326 | 0.208 | 0.242 | 0.984177142 | 1 |
| WBGene00012760 | Y41C4A.17 | 4.626913474 | 4.317422189 | 0.446501542 | 0.832 | 0.863 | 0.565056618 | 1 |

| | | | | | | | | |
|---|---|---|---|---|---|---|---|---|
| WBGene00008914 | F17C11.2 | 1.425899346 | 1.113510598 | 0.450681698 | 0.277 | 0.216 | 0.095155527 | 1 |
| WBGene00012348 | pptr-1 | 1.683859045 | 1.3710155 | 0.451337831 | 0.382 | 0.479 | 0.829833705 | 1 |
| WBGene00022185 | Y71H2AM.20 | 0.463002482 | 0.150025341 | 0.451530569 | 0.064 | 0.037 | 0.210352109 | 1 |
| WBGene00077783 | D1054.18 | 0.872736111 | 0.558188577 | 0.453796167 | 0.15 | 0.121 | 0.290405665 | 1 |
| WBGene00219609 | linc-7 | 4.372932751 | 4.057330101 | 0.455318377 | 0.607 | 0.532 | 0.041596323 | 1 |
| WBGene00002097 | ins-14 | 2.722662669 | 2.406230071 | 0.45651574 | 0.659 | 0.726 | 0.085939208 | 1 |
| WBGene00011453 | ugt-56 | 0.423410552 | 0.106604776 | 0.457054121 | 0.052 | 0.026 | 0.18605588 | 1 |
| WBGene00008558 | F07H5.7 | 0.453974531 | 0.136794518 | 0.457594033 | 0.064 | 0.042 | 0.305983525 | 1 |
| WBGene00002094 | ins-11 | 0.609039028 | 0.29125867 | 0.458460146 | 0.075 | 0.042 | 0.176949852 | 1 |
| WBGene00001565 | gei-8 | 1.254415913 | 0.936389341 | 0.458815357 | 0.249 | 0.232 | 0.29643779 | 1 |
| WBGene00011308 | pnn-1 | 0.582093062 | 0.263734833 | 0.459293838 | 0.081 | 0.079 | 0.788391762 | 1 |
| WBGene00045388 | lfor-1 | 0.78058019 | 0.461347552 | 0.460555344 | 0.11 | 0.116 | 0.961483762 | 1 |
| WBGene00016783 | irg-2 | 0.751124866 | 0.430504107 | 0.46255798 | 0.121 | 0.074 | 0.107310161 | 1 |
| WBGene00009579 | arrd-6 | 0.967231034 | 0.646448871 | 0.462790835 | 0.173 | 0.2 | 0.965965236 | 1 |
| WBGene00008307 | ncs-5 | 1.298661017 | 0.977745189 | 0.462983674 | 0.266 | 0.316 | 0.944829746 | 1 |
| WBGene00008131 | C47B2.2 | 0.41278161 | 0.089536614 | 0.466343952 | 0.052 | 0.021 | 0.104070121 | 1 |
| WBGene00017640 | F20D6.11 | 0.683375393 | 0.359917306 | 0.466651379 | 0.121 | 0.105 | 0.423181763 | 1 |
| WBGene00013136 | Y53C10A.5 | 0.787948943 | 0.464423552 | 0.466748478 | 0.116 | 0.137 | 0.811678969 | 1 |
| WBGene00013639 | Y105C5A.15 | 0.961234409 | 0.637243568 | 0.46741998 | 0.156 | 0.132 | 0.344114612 | 1 |
| WBGene00012885 | iscu-1 | 0.757204535 | 0.432426008 | 0.468556371 | 0.098 | 0.137 | 0.440224413 | 1 |
| WBGene00017220 | F07F6.8 | 0.846755085 | 0.521494382 | 0.469252002 | 0.121 | 0.142 | 0.81637188 | 1 |
| WBGene00007390 | lmtr-3 | 1.051533228 | 0.725011292 | 0.471071577 | 0.139 | 0.221 | 0.138625053 | 1 |
| WBGene00011723 | T12A7.2 | 0.931124219 | 0.603532573 | 0.472614843 | 0.156 | 0.153 | 0.627565389 | 1 |
| WBGene00001534 | gcy-7 | 0.559211816 | 0.230751946 | 0.473867425 | 0.075 | 0.053 | 0.327899779 | 1 |
| WBGene00001562 | lin-66 | 0.713814404 | 0.384536428 | 0.475047704 | 0.081 | 0.1 | 0.668830526 | 1 |
| WBGene00020929 | W02C12.2 | 0.418550839 | 0.089032416 | 0.475394594 | 0.052 | 0.026 | 0.183076197 | 1 |
| WBGene00000223 | atf-7 | 1.348539694 | 1.01734995 | 0.477805801 | 0.272 | 0.332 | 0.919082374 | 1 |
| WBGene00010424 | lep-5 | 0.533628349 | 0.202154361 | 0.478215879 | 0.017 | 0.063 | 0.03291594 | 1 |
| WBGene00220257 | C02H6.3 | 2.460569767 | 2.128676922 | 0.478820162 | 0.578 | 0.679 | 0.311669186 | 1 |
| WBGene00045383 | sup-46 | 0.669303249 | 0.337396639 | 0.47884002 | 0.081 | 0.095 | 0.809304926 | 1 |
| WBGene00010639 | K07F5.15 | 0.698235732 | 0.364623613 | 0.481300549 | 0.075 | 0.084 | 0.862173807 | 1 |
| WBGene00005715 | srv-4 | 1.057993743 | 0.7243441 | 0.481354686 | 0.214 | 0.184 | 0.219062115 | 1 |
| WBGene00021562 | nuo-5 | 0.460200021 | 0.125895035 | 0.482300146 | 0.092 | 0.026 | 0.006006238 | 1 |
| WBGene00022606 | dmsr-9 | 0.931939231 | 0.595542395 | 0.485318047 | 0.162 | 0.147 | 0.488759018 | 1 |
| WBGene00018626 | F49D11.2 | 0.755649615 | 0.418940951 | 0.48576792 | 0.121 | 0.116 | 0.631832303 | 1 |
| WBGene00004453 | rpl-39 | 2.96734816 | 2.628858661 | 0.488337121 | 0.821 | 0.874 | 8.78E-05 | 0.54416786 |
| WBGene00010244 | F58D5.5 | 1.234438546 | 0.895857608 | 0.488469039 | 0.225 | 0.226 | 0.49017679 | 1 |
| WBGene00015902 | nhr-159 | 0.917932612 | 0.577104046 | 0.491711682 | 0.145 | 0.147 | 0.823857039 | 1 |
| WBGene00005231 | srh-5 | 0.945201501 | 0.603997862 | 0.492252797 | 0.191 | 0.179 | 0.395963512 | 1 |
| WBGene00003605 | nhr-6 | 1.813284738 | 1.47157609 | 0.492981371 | 0.277 | 0.326 | 0.824586198 | 1 |
| WBGene00010877 | lact-4 | 0.822243883 | 0.480340775 | 0.493261919 | 0.116 | 0.084 | 0.252587162 | 1 |
| WBGene00021254 | Y22D7AL.16 | 2.454419462 | 2.111906168 | 0.49414223 | 0.671 | 0.711 | 0.103105717 | 1 |
| WBGene00004208 | ptc-1 | 2.388235958 | 2.044642662 | 0.495700345 | 0.555 | 0.642 | 0.87295392 | 1 |
| WBGene00022500 | lfi-1 | 1.468189249 | 1.124305249 | 0.496119741 | 0.353 | 0.363 | 0.289815495 | 1 |
| WBGene00013845 | fbxa-37 | 0.518563316 | 0.173320501 | 0.498080097 | 0.075 | 0.037 | 0.096363782 | 1 |
| WBGene00009814 | F47B10.3 | 1.7709102 | 1.422451142 | 0.502720155 | 0.41 | 0.511 | 0.815647352 | 1 |
| WBGene00002179 | jph-1 | 0.692158019 | 0.342117488 | 0.505001738 | 0.104 | 0.1 | 0.681740617 | 1 |
| WBGene00010918 | M117.1 | 1.896327801 | 1.546041621 | 0.505356135 | 0.434 | 0.511 | 0.770699092 | 1 |
| WBGene00005410 | srh-199 | 1.479218285 | 1.128032548 | 0.506653922 | 0.306 | 0.353 | 0.757913928 | 1 |
| WBGene00015654 | srz-77 | 0.80737596 | 0.455116042 | 0.508203636 | 0.139 | 0.084 | 0.071142653 | 1 |
| WBGene00009622 | F41E7.6 | 0.412798333 | 0.060461174 | 0.508315072 | 0.069 | 0.016 | 0.009343447 | 1 |
| WBGene00013216 | Y54G11A.7 | 0.503127603 | 0.145815401 | 0.515492541 | 0.052 | 0.037 | 0.446242709 | 1 |
| WBGene00013490 | Y70C5A.2 | 0.880004864 | 0.522524447 | 0.515735224 | 0.127 | 0.126 | 0.772954891 | 1 |

| | | | | | | | | |
|---|---|---|---|---|---|---|---|---|
| WBGene00044771 | tmem-218 | 1.911925396 | 1.554154699 | 0.516154009 | 0.491 | 0.532 | 0.218818785 | 1 |
| WBGene00011631 | T08G11.4 | 0.740855234 | 0.382774218 | 0.516601706 | 0.11 | 0.111 | 0.785818659 | 1 |
| WBGene00021952 | vha-19 | 0.862902707 | 0.503731468 | 0.518174566 | 0.133 | 0.147 | 0.997354937 | 1 |
| WBGene00010965 | ctc-2 | 1.985983452 | 1.623282519 | 0.523266837 | 0.497 | 0.537 | 0.247990552 | 1 |
| WBGene00194851 | K07C10.3 | 0.818165064 | 0.455000817 | 0.523935258 | 0.173 | 0.1 | 0.017430082 | 1 |
| WBGene00020952 | kel-8 | 0.566934797 | 0.201806907 | 0.526768196 | 0.064 | 0.053 | 0.571681539 | 1 |
| WBGene00013389 | Y62H9A.1 | 1.581141188 | 1.215865559 | 0.526981339 | 0.312 | 0.384 | 0.897072106 | 1 |
| WBGene00011955 | cdka-1 | 0.660271404 | 0.294019501 | 0.528389803 | 0.069 | 0.084 | 0.738031456 | 1 |
| WBGene00019568 | K09D9.11 | 0.945806972 | 0.578076991 | 0.530522219 | 0.168 | 0.163 | 0.534198897 | 1 |
| WBGene00015799 | C15H9.3 | 1.040175547 | 0.669216411 | 0.535180906 | 0.168 | 0.189 | 0.959967377 | 1 |
| WBGene00009918 | gcsh-2 | 0.886152908 | 0.515111543 | 0.535299537 | 0.185 | 0.174 | 0.335705069 | 1 |
| WBGene00018736 | F53B1.3 | 1.347727843 | 0.976079898 | 0.536174647 | 0.225 | 0.274 | 0.800550128 | 1 |
| WBGene00020366 | acdh-10 | 0.649818502 | 0.277733456 | 0.53680525 | 0.075 | 0.068 | 0.712492735 | 1 |
| WBGene00020737 | T23F2.4 | 0.787492606 | 0.414962683 | 0.537447072 | 0.104 | 0.089 | 0.504871016 | 1 |
| WBGene00005363 | srh-147 | 1.595279098 | 1.219605177 | 0.541982903 | 0.353 | 0.426 | 0.722102099 | 1 |
| WBGene00194689 | Y38H6C.24 | 0.941316857 | 0.564984432 | 0.542932924 | 0.121 | 0.105 | 0.492645808 | 1 |
| WBGene00014473 | MTCE.36 | 0.721723817 | 0.344238381 | 0.544596365 | 0.121 | 0.089 | 0.225177702 | 1 |
| WBGene00004443 | rpl-29 | 2.244137723 | 1.864396004 | 0.547851495 | 0.636 | 0.653 | 0.000709926 | 1 |
| WBGene00019841 | R02F11.3 | 1.623285455 | 1.243214754 | 0.548326117 | 0.295 | 0.432 | 0.351994832 | 1 |
| WBGene00003025 | lin-40 | 1.204156036 | 0.823238003 | 0.549548558 | 0.254 | 0.242 | 0.244458316 | 1 |
| WBGene00001177 | egl-8 | 1.29305282 | 0.911753799 | 0.550098207 | 0.162 | 0.247 | 0.126514631 | 1 |
| WBGene00021362 | fmil-1 | 0.984419991 | 0.602864855 | 0.550467703 | 0.179 | 0.158 | 0.31569013 | 1 |
| WBGene00000097 | aip-1 | 1.665573614 | 1.28372063 | 0.550897407 | 0.37 | 0.405 | 0.472497703 | 1 |
| WBGene00005207 | srg-50 | 0.851566373 | 0.465231158 | 0.557363899 | 0.179 | 0.132 | 0.09108867 | 1 |
| WBGene00021239 | Y20F4.4 | 0.521065507 | 0.134247899 | 0.558059844 | 0.064 | 0.042 | 0.303537758 | 1 |
| WBGene00020524 | dmsr-13 | 1.758711073 | 1.371727985 | 0.558298581 | 0.37 | 0.426 | 0.732571778 | 1 |
| WBGene00009140 | F26A3.1 | 1.550494671 | 1.162380053 | 0.559931034 | 0.364 | 0.379 | 0.145211173 | 1 |
| WBGene00045195 | F53F4.17 | 2.269828158 | 1.880917164 | 0.561079961 | 0.578 | 0.626 | 0.03932239 | 1 |
| WBGene00018068 | F35H10.2 | 1.858421078 | 1.469373422 | 0.561277124 | 0.341 | 0.426 | 0.588081951 | 1 |
| WBGene00017637 | F20D6.8 | 0.78034737 | 0.391077766 | 0.561597326 | 0.15 | 0.1 | 0.077532608 | 1 |
| WBGene00003947 | pbs-1 | 0.982187203 | 0.592316379 | 0.562464705 | 0.15 | 0.153 | 0.804730978 | 1 |
| WBGene00005423 | srh-213 | 1.008665647 | 0.617541641 | 0.564272664 | 0.197 | 0.142 | 0.075218871 | 1 |
| WBGene00007520 | C11E4.6 | 0.734614274 | 0.340893664 | 0.568018772 | 0.121 | 0.095 | 0.264013917 | 1 |
| WBGene00201745 | C14F5.11 | 0.640800067 | 0.246315457 | 0.56912099 | 0.087 | 0.053 | 0.159593381 | 1 |
| WBGene00007506 | fbxc-58 | 0.458161513 | 0.060538959 | 0.573648088 | 0.058 | 0.005 | 0.003768022 | 1 |
| WBGene00014213 | ZK1073.1 | 0.932657063 | 0.532659178 | 0.577074964 | 0.173 | 0.153 | 0.347172717 | 1 |
| WBGene00001994 | hpk-1 | 1.875836919 | 1.474885235 | 0.578451007 | 0.509 | 0.468 | 0.019519203 | 1 |
| WBGene00010059 | F54E4.3 | 0.751427139 | 0.34934988 | 0.580074867 | 0.052 | 0.095 | 0.155313457 | 1 |
| WBGene00001336 | qars-1 | 0.970828572 | 0.56713262 | 0.582410149 | 0.191 | 0.184 | 0.398769999 | 1 |
| WBGene00043060 | R144.12 | 0.562337282 | 0.157148545 | 0.584563781 | 0.064 | 0.037 | 0.209384358 | 1 |
| WBGene00006594 | tom-1 | 0.626788381 | 0.21959144 | 0.587461008 | 0.075 | 0.068 | 0.674460189 | 1 |
| WBGene00019569 | K09D9.12 | 1.532843364 | 1.125584557 | 0.587550261 | 0.329 | 0.321 | 0.232091457 | 1 |
| WBGene00044644 | B0205.13 | 2.772697805 | 2.364667232 | 0.588663684 | 0.266 | 0.305 | 0.818838337 | 1 |
| WBGene00021050 | W05H9.3 | 2.008819066 | 1.600322469 | 0.589336015 | 0.474 | 0.453 | 0.068268523 | 1 |
| WBGene00195209 | C07B5.8 | 0.532710676 | 0.122637621 | 0.591610363 | 0.087 | 0.021 | 0.004658682 | 1 |
| WBGene00016788 | C49G7.10 | 2.1809228 | 1.76751296 | 0.596424327 | 0.202 | 0.242 | 0.563384658 | 1 |
| WBGene00002637 | let-418 | 0.8053588 | 0.390845117 | 0.598016835 | 0.133 | 0.111 | 0.316913266 | 1 |
| WBGene00000284 | cah-6 | 0.864335722 | 0.445342849 | 0.60447894 | 0.127 | 0.121 | 0.637926777 | 1 |
| WBGene00002295 | let-19 | 0.78022045 | 0.35849909 | 0.608415315 | 0.139 | 0.116 | 0.302125879 | 1 |
| WBGene00003678 | nhr-88 | 1.093884896 | 0.671341511 | 0.609601245 | 0.202 | 0.211 | 0.592129716 | 1 |
| WBGene00010828 | M02B1.3 | 1.000041016 | 0.575188871 | 0.612932083 | 0.185 | 0.142 | 0.135092443 | 1 |
| WBGene00000993 | dhs-30 | 0.653334046 | 0.226633242 | 0.615599133 | 0.098 | 0.058 | 0.11133771 | 1 |
| WBGene00018476 | F45E12.6 | 1.742502209 | 1.313510244 | 0.618904581 | 0.405 | 0.421 | 0.192668863 | 1 |

| | | | | | | | | |
|---|---|---|---|---|---|---|---|---|
| WBGene00019079 | rpa-4 | 1.040488462 | 0.611051641 | 0.619546372 | 0.173 | 0.168 | 0.547798901 | 1 |
| WBGene00012471 | Y17G7B.20 | 1.325662843 | 0.895522192 | 0.620561784 | 0.283 | 0.279 | 0.240496778 | 1 |
| WBGene00004140 | ebax-1 | 0.683793887 | 0.247843868 | 0.628942931 | 0.104 | 0.063 | 0.110710133 | 1 |
| WBGene00003636 | nhr-46 | 1.568068836 | 1.123621914 | 0.64120137 | 0.301 | 0.326 | 0.645078337 | 1 |
| WBGene00009917 | glb-18 | 1.375085466 | 0.925342254 | 0.648842301 | 0.237 | 0.284 | 0.937526794 | 1 |
| WBGene00002163 | ist-1 | 0.613472675 | 0.163098544 | 0.649752525 | 0.11 | 0.042 | 0.009784407 | 1 |
| WBGene00008741 | ctsa-1.2 | 0.766831627 | 0.315374497 | 0.651314963 | 0.116 | 0.084 | 0.221962627 | 1 |
| WBGene00016310 | C32D5.1 | 0.665656353 | 0.207611672 | 0.66081879 | 0.116 | 0.058 | 0.031488303 | 1 |
| WBGene00249817 | F42G8.19 | 2.067154922 | 1.600266102 | 0.673578185 | 0.549 | 0.611 | 0.021460024 | 1 |
| WBGene00022526 | ZC132.8 | 1.196970282 | 0.726549076 | 0.678674342 | 0.145 | 0.2 | 0.394168169 | 1 |
| WBGene00001439 | fkh-7 | 1.257934542 | 0.772420317 | 0.700448965 | 0.266 | 0.205 | 0.047002615 | 1 |
| WBGene00017510 | nhr-178 | 0.69848363 | 0.211623197 | 0.702391132 | 0.098 | 0.042 | 0.027944673 | 1 |
| WBGene00013273 | 5-Mar | 0.623056493 | 0.129032637 | 0.712725767 | 0.087 | 0.026 | 0.010037644 | 1 |
| WBGene00017060 | D2063.1 | 1.728667376 | 1.231256652 | 0.717611985 | 0.243 | 0.237 | 0.49086485 | 1 |
| WBGene00007645 | C17E4.6 | 1.168151804 | 0.666733936 | 0.723393071 | 0.185 | 0.195 | 0.662901284 | 1 |
| WBGene00012928 | aakb-2 | 0.930248562 | 0.424739816 | 0.729294962 | 0.179 | 0.121 | 0.048316665 | 1 |
| WBGene00017317 | attf-2 | 1.815492695 | 1.303113176 | 0.739207391 | 0.295 | 0.4 | 0.492957416 | 1 |
| WBGene00219650 | linc-56 | 2.167695019 | 1.627690657 | 0.779061615 | 0.468 | 0.511 | 0.289830427 | 1 |
| WBGene00022407 | srsx-4 | 0.959141712 | 0.409677614 | 0.792709129 | 0.185 | 0.116 | 0.024680342 | 1 |
| WBGene00018298 | nlp-50 | 1.122830555 | 0.572922885 | 0.793349069 | 0.092 | 0.132 | 0.352688251 | 1 |
| WBGene00020596 | oga-1 | 0.800182205 | 0.235222931 | 0.815063943 | 0.15 | 0.053 | 0.001075231 | 1 |
| WBGene00219602 | linc-151 | 1.492674379 | 0.92548702 | 0.81827839 | 0.197 | 0.205 | 0.776992021 | 1 |
| WBGene00022389 | pde-6 | 1.167150045 | 0.58744102 | 0.836343335 | 0.168 | 0.179 | 0.791159093 | 1 |
| WBGene00022404 | srsx-5 | 0.948318033 | 0.347227155 | 0.867190829 | 0.179 | 0.089 | 0.004509167 | 1 |
| WBGene00008592 | F08H9.4 | 0.973339749 | 0.355025877 | 0.892038358 | 0.15 | 0.068 | 0.006786427 | 1 |
| WBGene00009623 | F41E7.7 | 0.787905054 | 0.156247585 | 0.911289098 | 0.173 | 0.042 | 2.11E-05 | 0.130710633 |
| WBGene00304812 | ZK380.6 | 5.045097573 | 4.41299833 | 0.911926444 | 0.798 | 0.779 | 0.000184116 | 1 |
| WBGene00304815 | Y40A1A.6 | 2.308111388 | 1.672748789 | 0.916634471 | 0.156 | 0.205 | 0.357817387 | 1 |
| WBGene00017570 | F18E9.4 | 0.74429969 | 0.100945668 | 0.928163656 | 0.098 | 0.021 | 0.001438643 | 1 |
| WBGene00009888 | F49E2.5 | 1.874266849 | 1.228332273 | 0.931886609 | 0.387 | 0.363 | 0.061717139 | 1 |
| WBGene00001452 | flp-9 | 0.950721399 | 0.279335609 | 0.96860495 | 0.098 | 0.047 | 0.047041817 | 1 |
| WBGene00009988 | F53F4.4 | 0.949121393 | 0.233654761 | 1.032200162 | 0.069 | 0.058 | 0.569204443 | 1 |
| WBGene00005206 | srg-49 | 1.351696325 | 0.635825683 | 1.032783025 | 0.277 | 0.179 | 0.003057939 | 1 |
| WBGene00012382 | ttr-16 | 1.682550647 | 0.855401453 | 1.19332404 | 0.335 | 0.232 | 0.002307969 | 1 |
| WBGene00005205 | srg-48 | 1.940065866 | 1.059075836 | 1.270999947 | 0.486 | 0.374 | 5.00E-06 | 0.030992568 |
