## Supplemental Data S5 for "The forkhead transcription factor FKH-7/FOXP acts in chemosensory neurons to regulate developmental decision-making"

**Data S5. Single-cell RNA-seq ADF gene list table.**

This comparison is between all *fkh-7(R612C)* ADF cells and all *fkh-7(wild-type)* ADF cells.

| id | gene_name | fkh_7.avg | wt.avg | avg_log2FC | fkh7.pct | wt.pct | p_val | p_val_adj |
|---|---|---|---|---|---|---|---|---|
| WBGene00015514 | nlp-77 | 0.177027505 | 1.11163825 | -1.348358287 | 0.011 | 0.11 | 4.16E-12 | 2.65E-08 |
| WBGene00017068 | D2092.8 | 0.254692878 | 1.161271838 | -1.30791697 | 0.011 | 0.087 | 4.51E-09 | 2.88E-05 |
| WBGene00015354 | snet-1 | 0.49948596 | 1.360693213 | -1.242459433 | 0.044 | 0.054 | 0.497266889 | 1 |
| WBGene00004451 | rpl-37 | 1.869197391 | 2.567063251 | -1.006807616 | 0.345 | 0.644 | 2.34E-20 | 1.49E-16 |
| WBGene00008570 | kcnl-2 | 1.388635491 | 2.023899491 | -0.916492223 | 0.18 | 0.421 | 4.60E-15 | 2.93E-11 |
| WBGene00005834 | srw-87 | 0.251537613 | 0.851107636 | -0.8649967 | 0.021 | 0.138 | 1.19E-12 | 7.58E-09 |
| WBGene00017613 | F20A1.1 | 0.181764661 | 0.731208318 | -0.792679639 | 0.014 | 0.065 | 1.45E-05 | 0.092457965 |
| WBGene00044079 | tag-241 | 0.330488153 | 0.871197604 | -0.780078844 | 0.028 | 0.092 | 1.03E-05 | 0.065836897 |
| WBGene00008382 | D1081.4 | 0.699562895 | 1.238543766 | -0.77758503 | 0.062 | 0.171 | 7.77E-08 | 0.000495513 |
| WBGene00022405 | srsx-6 | 0.931868293 | 1.469531684 | -0.775684308 | 0.113 | 0.277 | 6.89E-10 | 4.40E-06 |
| WBGene00007588 | C14C10.5 | 0.158119841 | 0.6883091 | -0.764901416 | 0.014 | 0.052 | 0.000476987 | 1 |
| WBGene00017858 | F27C1.11 | 1.254613572 | 1.780114228 | -0.75813719 | 0.152 | 0.35 | 1.23E-11 | 7.84E-08 |
| WBGene00002017 | hsp-16.11 | 3.696248148 | 4.199664371 | -0.726276088 | 0.721 | 0.877 | 2.04E-20 | 1.30E-16 |
| WBGene00006352 | sur-6 | 0.628398558 | 1.127733092 | -0.720387456 | 0.067 | 0.177 | 1.62E-07 | 0.001030538 |
| WBGene00012967 | kvs-4 | 0.732567537 | 1.228265855 | -0.715141506 | 0.072 | 0.206 | 1.34E-09 | 8.58E-06 |
| WBGene00022887 | ZK1290.5 | 0.760852903 | 1.247963619 | -0.702752214 | 0.071 | 0.217 | 7.92E-11 | 5.05E-07 |
| WBGene00010647 | K08C7.4 | 0.596735519 | 1.082991039 | -0.701518429 | 0.064 | 0.152 | 8.24E-06 | 0.052594982 |
| WBGene00016532 | C39D10.5 | 0.652799867 | 1.134418126 | -0.694828273 | 0.053 | 0.167 | 5.41E-09 | 3.45E-05 |
| WBGene00003412 | mrp-6 | 0.404016427 | 0.885450328 | -0.694562302 | 0.039 | 0.094 | 0.000330627 | 1 |
| WBGene00013639 | Y105C5A.15 | 0.837602792 | 1.317942569 | -0.692983814 | 0.08 | 0.19 | 7.97E-07 | 0.005082728 |
| WBGene00000920 | daf-28 | 3.633477158 | 4.111381157 | -0.68946973 | 0.712 | 0.888 | 8.07E-16 | 5.15E-12 |
| WBGene00015478 | mapk-15 | 1.075880889 | 1.553732265 | -0.689393811 | 0.108 | 0.275 | 6.39E-11 | 4.08E-07 |
| WBGene00010516 | K02E11.7 | 0.250021084 | 0.702389506 | -0.652629679 | 0.011 | 0.073 | 2.29E-07 | 0.001459418 |
| WBGene00008516 | F02C12.2 | 1.073964168 | 1.520606491 | -0.644368665 | 0.134 | 0.319 | 1.18E-10 | 7.54E-07 |
| WBGene00009995 | F53F4.13 | 0.505561727 | 0.947142245 | -0.637066024 | 0.028 | 0.083 | 0.00011184 | 0.713649622 |
| WBGene00004435 | rpl-23 | 1.93195925 | 2.372173335 | -0.635094677 | 0.332 | 0.594 | 4.48E-13 | 2.86E-09 |
| WBGene00044353 | ZK418.10 | 0.71095604 | 1.146434678 | -0.628262871 | 0.071 | 0.215 | 9.45E-11 | 6.03E-07 |
| WBGene00012528 | pap-1 | 1.333545913 | 1.763090504 | -0.619701851 | 0.171 | 0.346 | 6.42E-09 | 4.10E-05 |
| WBGene00018467 | sdpn-1 | 1.412551025 | 1.839386513 | -0.615793441 | 0.18 | 0.396 | 2.13E-11 | 1.36E-07 |
| WBGene00020844 | T27A10.2 | 0.855843549 | 1.282558239 | -0.615619167 | 0.081 | 0.212 | 8.07E-09 | 5.15E-05 |
| WBGene00006801 | unc-68 | 0.679009452 | 1.104139125 | -0.61333247 | 0.065 | 0.185 | 1.68E-08 | 0.000107442 |
| WBGene00018009 | dhhc-3 | 0.080480946 | 0.505504518 | -0.6131794 | 0.007 | 0.05 | 1.79E-05 | 0.114135975 |
| WBGene00004286 | rab-39 | 0.396044223 | 0.814974303 | -0.604388349 | 0.035 | 0.092 | 0.000210859 | 1 |
| WBGene00014016 | ZK632.9 | 1.309413406 | 1.728299342 | -0.604324663 | 0.178 | 0.327 | 1.24E-06 | 0.007909118 |
| WBGene00001668 | gpa-6 | 0.569380351 | 0.987920346 | -0.603825575 | 0.057 | 0.14 | 1.04E-05 | 0.06638007 |
| WBGene00007330 | C05C10.3 | 0.470037113 | 0.885513114 | -0.599405166 | 0.039 | 0.144 | 4.19E-09 | 2.68E-05 |
| WBGene00010006 | F53H2.3 | 0.572259119 | 0.984268916 | -0.594404491 | 0.062 | 0.131 | 0.000231107 | 1 |
| WBGene00001480 | fmo-5 | 1.432907428 | 1.841773476 | -0.589869021 | 0.145 | 0.256 | 1.22E-05 | 0.077820433 |
| WBGene00022790 | ZK682.7 | 1.25525181 | 1.661821384 | -0.586555907 | 0.166 | 0.302 | 3.89E-06 | 0.02484719 |
| WBGene00004567 | rrn-2.1 | 1.940960014 | 2.34545312 | -0.583560197 | 0.284 | 0.49 | 1.43E-09 | 9.16E-06 |
| WBGene00021382 | Y37F4.1 | 2.830430724 | 3.233312672 | -0.58123579 | 0.481 | 0.623 | 4.35E-05 | 0.277673127 |
| WBGene00016855 | C50H11.8 | 0.33866204 | 0.740543439 | -0.579792301 | 0.021 | 0.052 | 0.006939965 | 1 |
| WBGene00015263 | B0563.6 | 0.333274304 | 0.734916228 | -0.579446812 | 0.021 | 0.098 | 1.09E-07 | 0.000695397 |
| WBGene00007696 | tram-1 | 0.381908524 | 0.782225716 | -0.577535627 | 0.027 | 0.092 | 6.14E-06 | 0.039165331 |
| WBGene00004363 | ric-3 | 0.382742609 | 0.777021562 | -0.568824289 | 0.041 | 0.088 | 0.001807346 | 1 |
| WBGene00018782 | cct-3 | 0.129799324 | 0.523175489 | -0.567521842 | 0.012 | 0.054 | 0.000127092 | 0.810973395 |
| WBGene00015735 | pdfr-1 | 1.175416942 | 1.568019201 | -0.566405333 | 0.118 | 0.271 | 1.92E-08 | 0.000122329 |
| WBGene00011300 | R107.5 | 1.615485183 | 2.00712452 | -0.56501613 | 0.219 | 0.394 | 2.04E-07 | 0.001299472 |
| WBGene00011955 | cdka-1 | 0.56942176 | 0.958834394 | -0.561803676 | 0.057 | 0.142 | 6.34E-06 | 0.04045303 |
| WBGene00009122 | tct-1 | 0.671170971 | 1.059899005 | -0.560816008 | 0.076 | 0.181 | 1.61E-06 | 0.010277904 |
| WBGene00018281 | tbc-18 | 0.447760809 | 0.833846894 | -0.557004479 | 0.044 | 0.096 | 0.001269126 | 1 |
| WBGene00001340 | etr-1 | 1.85568833 | 2.241563718 | -0.556700508 | 0.306 | 0.542 | 5.69E-11 | 3.63E-07 |
| WBGene00003072 | lrp-2 | 0.2514571 | 0.636864055 | -0.556024703 | 0.025 | 0.087 | 1.06E-05 | 0.067891714 |
| WBGene00008518 | irld-2 | 0.382529211 | 0.767766286 | -0.555779618 | 0.037 | 0.11 | 8.31E-06 | 0.053054916 |
| WBGene00007862 | kvs-2 | 1.295094343 | 1.67726255 | -0.551352178 | 0.154 | 0.327 | 2.60E-09 | 1.66E-05 |
| WBGene00000056 | acr-17 | 0.382950045 | 0.763028486 | -0.548337282 | 0.023 | 0.1 | 1.56E-07 | 0.00099371 |
| WBGene00004481 | rps-12 | 2.238397112 | 2.613207268 | -0.540736754 | 0.412 | 0.66 | 1.04E-09 | 6.62E-06 |
| WBGene00002344 | let-70 | 1.749001021 | 2.122459971 | -0.538787376 | 0.299 | 0.523 | 4.24E-09 | 2.71E-05 |
| WBGene00001708 | grk-1 | 0.231590911 | 0.603591247 | -0.53668304 | 0.021 | 0.073 | 6.41E-05 | 0.40905838 |
| WBGene00050944 | C14A11.9 | 0.399713213 | 0.771099966 | -0.535797827 | 0.039 | 0.129 | 2.00E-07 | 0.001274272 |

| Legend | |
|---|---|
| id | WormBase GeneID |
| gene_name | Gene symbol (WS273) |
| fkh_7.avg | Log of averaged normalized expression across all *fkh-7(R612C)* ADF neurons |
| wt.avg | Log of averaged normalized expression across all *fkh-7(wild-type)* ADF neurons |
| avg_log2FC | log fold-change (*fkh-7(R612C) - fkh-7(wild-type)* ) |
| fkh7.pct | Percent of *fkh-7(R612C)* ADF neurons where gene is detected ($\geq$ 1 UMI) |
| wt.pct | Percent of *fkh-7(wild-type)* ADF neurons where gene is detected ($\geq$ 1 UMI) |
| p_val | p-value |
| p_val_adj | Adjusted p-value, based on bonferroni correction |

| | | | | | | | | |
|---|---|---|---|---|---|---|---|---|
| WBGene00001426 | fkb-1 | 0.22637395 | 0.597400858 | -0.535278681 | 0.021 | 0.06 | 0.001484044 | 1 |
| WBGene00001677 | gpa-15 | 0.299609902 | 0.668799 | -0.53262728 | 0.027 | 0.063 | 0.00361982 | 1 |
| WBGene00004760 | sel-2 | 0.463941662 | 0.83282328 | -0.53218368 | 0.042 | 0.102 | 0.000238926 | 1 |
| WBGene00044995 | Y105C5A.28 | 0.421365801 | 0.789581642 | -0.531223167 | 0.035 | 0.125 | 9.93E-08 | 0.000633734 |
| WBGene00003808 | npr-2 | 0.550542996 | 0.91776933 | -0.529795611 | 0.035 | 0.121 | 2.39E-07 | 0.001528031 |
| WBGene00011372 | npr-26 | 1.342100922 | 1.70864945 | -0.528817744 | 0.159 | 0.34 | 1.07E-09 | 6.85E-06 |
| WBGene00006780 | unc-44 | 1.51281969 | 1.879088401 | -0.528414053 | 0.191 | 0.356 | 4.33E-07 | 0.002764044 |
| WBGene00009688 | F44E5.1 | 0.482312458 | 0.847801616 | -0.527289395 | 0.053 | 0.106 | 0.002527329 | 1 |
| WBGene00003476 | mtm-3 | 1.106387797 | 1.469481522 | -0.523833517 | 0.138 | 0.3 | 1.12E-08 | 7.18E-05 |
| WBGene00005040 | sra-14 | 0.817959023 | 1.180521229 | -0.523066696 | 0.083 | 0.198 | 4.11E-07 | 0.002621216 |
| WBGene00001303 | sec-61.G | 1.248185302 | 1.610429799 | -0.522608339 | 0.155 | 0.325 | 1.86E-08 | 0.000119001 |
| WBGene00019847 | R03G5.6 | 0.379386866 | 0.741156469 | -0.521923212 | 0.03 | 0.104 | 1.96E-06 | 0.012503013 |
| WBGene00000481 | cha-1 | 0.82174787 | 1.183002367 | -0.521180071 | 0.101 | 0.202 | 3.20E-05 | 0.204130675 |
| WBGene00017439 | F13H8.6 | 0.768137191 | 1.129315447 | -0.521070079 | 0.069 | 0.185 | 6.80E-08 | 0.000434106 |
| WBGene00011250 | R11H6.2 | 0.401144593 | 0.759190051 | -0.516550408 | 0.032 | 0.058 | 0.048028591 | 1 |
| WBGene00019691 | K12H6.12 | 0.748806673 | 1.103045419 | -0.511058481 | 0.076 | 0.165 | 3.62E-05 | 0.231059644 |
| WBGene00020333 | sre-12 | 0.448301582 | 0.800462163 | -0.508060323 | 0.042 | 0.087 | 0.004062258 | 1 |
| WBGene00009194 | F27E5.5 | 0.335056655 | 0.685391726 | -0.505426669 | 0.021 | 0.065 | 0.000359463 | 1 |
| WBGene00020181 | T02H6.11 | 0.583800505 | 0.933166805 | -0.50402903 | 0.06 | 0.129 | 0.000242808 | 1 |
| WBGene00020058 | R13D11.4 | 0.764917866 | 1.114159483 | -0.503849149 | 0.067 | 0.185 | 3.53E-08 | 0.000224939 |
| WBGene00021078 | unc-132 | 0.342357148 | 0.691240551 | -0.503332356 | 0.032 | 0.071 | 0.004563687 | 1 |
| WBGene00015377 | C03B1.6 | 0.130183711 | 0.478556817 | -0.502596152 | 0.011 | 0.065 | 2.03E-06 | 0.012925451 |
| WBGene00044732 | T03G11.10 | 0.215828036 | 0.562393638 | -0.499988476 | 0.021 | 0.056 | 0.003377644 | 1 |
| WBGene00000164 | apm-3 | 0.290217532 | 0.63643654 | -0.499488447 | 0.025 | 0.06 | 0.005444844 | 1 |
| WBGene00022732 | ZK418.2 | 0.90253913 | 1.247593785 | -0.497808639 | 0.087 | 0.206 | 2.47E-07 | 0.001576005 |
| WBGene00003831 | nuo-1 | 0.225076462 | 0.56838541 | -0.495290118 | 0.019 | 0.067 | 0.000116196 | 0.741444263 |
| WBGene00006479 | tmbi-4 | 0.315560625 | 0.657341879 | -0.49308612 | 0.025 | 0.102 | 2.53E-07 | 0.001615242 |
| WBGene00010146 | glb-19 | 1.691719375 | 2.033370331 | -0.492898141 | 0.228 | 0.446 | 7.80E-11 | 4.98E-07 |
| WBGene00015423 | C04E6.7 | 1.157118618 | 1.497449349 | -0.490993458 | 0.145 | 0.3 | 6.58E-08 | 0.000420152 |
| WBGene00012645 | mrpl-22 | 0.377879886 | 0.716688424 | -0.488797398 | 0.041 | 0.094 | 0.000590813 | 1 |
| WBGene00013490 | Y70C5A.2 | 0.663707174 | 1.001374792 | -0.487151397 | 0.072 | 0.187 | 1.72E-07 | 0.001098221 |
| WBGene00000488 | che-7 | 0.513692428 | 0.851316054 | -0.487087931 | 0.041 | 0.098 | 0.000275139 | 1 |
| WBGene00045419 | lsy-12 | 0.339249111 | 0.674585104 | -0.483787573 | 0.032 | 0.063 | 0.017119648 | 1 |
| WBGene00011046 | R05H10.7 | 0.096203163 | 0.430648572 | -0.482502733 | 0.005 | 0.056 | 9.90E-07 | 0.006319285 |
| WBGene00008977 | iglr-1 | 0.527797209 | 0.860733989 | -0.480326241 | 0.044 | 0.102 | 0.000373923 | 1 |
| WBGene00006529 | tba-2 | 0.943147135 | 1.274959887 | -0.478704612 | 0.111 | 0.202 | 0.00022937 | 1 |
| WBGene00004921 | snt-1 | 0.684517885 | 1.013272167 | -0.474292172 | 0.065 | 0.154 | 1.13E-05 | 0.071909889 |
| WBGene00018468 | cla-1 | 1.666484584 | 1.995096928 | -0.4740874 | 0.216 | 0.433 | 2.43E-10 | 1.55E-06 |
| WBGene00009784 | F46C5.9 | 0.525748519 | 0.852933775 | -0.472028547 | 0.055 | 0.137 | 1.29E-05 | 0.082325079 |
| WBGene00009100 | rgef-1 | 0.23779386 | 0.564924648 | -0.471949965 | 0.025 | 0.077 | 0.00011427 | 0.729156373 |
| WBGene00006756 | unc-17 | 1.424079004 | 1.751184285 | -0.471913166 | 0.196 | 0.367 | 3.48E-07 | 0.002220403 |
| WBGene00004433 | rpl-21 | 2.387877095 | 2.714939909 | -0.4718519 | 0.47 | 0.715 | 1.31E-10 | 8.38E-07 |
| WBGene00018849 | spt-16 | 0.47132371 | 0.795886534 | -0.468245177 | 0.048 | 0.112 | 0.000177086 | 1 |
| WBGene00044396 | ZK1290.14 | 0.381693387 | 0.705597953 | -0.467295512 | 0.021 | 0.058 | 0.00222434 | 1 |
| WBGene00006862 | pnk-4 | 0.347930741 | 0.670515778 | -0.465391833 | 0.037 | 0.075 | 0.008483979 | 1 |
| WBGene00011824 | T18D3.7 | 0.976636814 | 1.299009555 | -0.465085556 | 0.122 | 0.26 | 3.29E-07 | 0.002098379 |
| WBGene00004265 | qui-1 | 1.054318577 | 1.376364395 | -0.464613904 | 0.111 | 0.212 | 4.96E-05 | 0.31656279 |
| WBGene00001980 | hmr-1 | 0.698922534 | 1.018265321 | -0.460714255 | 0.074 | 0.158 | 6.44E-05 | 0.410984438 |
| WBGene00004482 | rps-13 | 2.292511371 | 2.610031746 | -0.45808507 | 0.424 | 0.69 | 1.58E-12 | 1.01E-08 |
| WBGene00009278 | npr-33 | 0.433784495 | 0.74963305 | -0.455673144 | 0.035 | 0.087 | 0.000555888 | 1 |
| WBGene00016734 | C46H11.10 | 0.71530786 | 1.029108994 | -0.452719339 | 0.083 | 0.188 | 3.05E-06 | 0.019474411 |
| WBGene00004469 | rps-0 | 2.237865393 | 2.551303985 | -0.452196303 | 0.451 | 0.665 | 3.20E-08 | 0.00020403 |
| WBGene00014938 | mdt-9 | 0.28177869 | 0.595212874 | -0.452189943 | 0.019 | 0.05 | 0.006374925 | 1 |
| WBGene00018704 | ztf-13 | 0.398872494 | 0.711546594 | -0.451093374 | 0.041 | 0.112 | 1.89E-05 | 0.120695275 |
| WBGene00014308 | F07H5.4 | 2.036819844 | 2.348870843 | -0.450194429 | 0.383 | 0.567 | 2.06E-05 | 0.131399252 |
| WBGene00006786 | unc-51 | 0.347376744 | 0.657379585 | -0.44723956 | 0.034 | 0.09 | 0.000159619 | 1 |
| WBGene00077491 | K08H10.10 | 2.02349324 | 2.332414732 | -0.445679505 | 0.35 | 0.56 | 2.70E-08 | 0.000172369 |
| WBGene00016405 | hecd-1 | 0.461848841 | 0.770186804 | -0.444837651 | 0.048 | 0.106 | 0.000442914 | 1 |
| WBGene00008890 | ser-5 | 0.579761178 | 0.887808829 | -0.444418819 | 0.044 | 0.138 | 1.72E-07 | 0.001096176 |
| WBGene00004484 | rps-15 | 2.277784833 | 2.585575824 | -0.444048536 | 0.422 | 0.677 | 6.65E-11 | 4.24E-07 |
| WBGene00001675 | gpa-13 | 0.28457012 | 0.592294814 | -0.44395289 | 0.027 | 0.071 | 0.000738516 | 1 |
| WBGene00020112 | pfd-5 | 0.269674812 | 0.577023032 | -0.443409753 | 0.019 | 0.062 | 0.000471387 | 1 |
| WBGene00015350 | fbxl-1 | 1.456809469 | 1.763971277 | -0.443140817 | 0.208 | 0.371 | 1.32E-06 | 0.008407637 |
| WBGene00006706 | ubc-9 | 0.417483986 | 0.72267242 | -0.44029384 | 0.039 | 0.092 | 0.000608506 | 1 |

| | | | | | | | | |
|---|---|---|---|---|---|---|---|---|
| WBGene00000263 | F23H11.5 | 1.070829683 | 1.375562255 | -0.439636171 | 0.133 | 0.262 | 3.09E-06 | 0.019697029 |
| WBGene00021607 | Y46H3C.7 | 0.513382267 | 0.817599116 | -0.43889214 | 0.039 | 0.11 | 1.40E-05 | 0.089523355 |
| WBGene00004622 | rrn-3.1 | 4.63423995 | 4.938014324 | -0.438253783 | 0.965 | 0.983 | 0.036122364 | 1 |
| WBGene00001226 | eif-3.C | 0.369703592 | 0.673260014 | -0.437939344 | 0.034 | 0.096 | 4.37E-05 | 0.27916423 |
| WBGene00015285 | gmeb-1 | 0.438913808 | 0.742301939 | -0.437696552 | 0.046 | 0.11 | 0.000179387 | 1 |
| WBGene00019603 | K09H11.6 | 0.961179962 | 1.263812593 | -0.436606597 | 0.083 | 0.196 | 4.27E-07 | 0.00272638 |
| WBGene00004454 | rpl-36.A | 2.464556193 | 2.766985661 | -0.436313494 | 0.512 | 0.712 | 4.05E-06 | 0.025858615 |
| WBGene00018188 | F38E9.6 | 1.863697145 | 2.16540217 | -0.435268343 | 0.3 | 0.487 | 4.01E-07 | 0.002559454 |
| WBGene00007041 | tag-180 | 0.137519788 | 0.439180786 | -0.435204826 | 0.009 | 0.052 | 3.15E-05 | 0.200985017 |
| WBGene00001747 | gsp-1 | 1.037426642 | 1.338514689 | -0.434378233 | 0.113 | 0.235 | 2.22E-06 | 0.014163833 |
| WBGene00010730 | ensa-1 | 0.82647488 | 1.126836853 | -0.433330729 | 0.087 | 0.198 | 1.75E-06 | 0.011146363 |
| WBGene00001031 | dnj-13 | 0.971407626 | 1.271325764 | -0.432690411 | 0.12 | 0.244 | 1.53E-06 | 0.009771455 |
| WBGene00017240 | calu-2 | 0.362705025 | 0.662438115 | -0.432423443 | 0.037 | 0.098 | 0.000107125 | 0.683563182 |
| WBGene00018879 | F55D10.4 | 0.351239949 | 0.65082641 | -0.432211902 | 0.028 | 0.079 | 0.000273486 | 1 |
| WBGene00021799 | Y53G8AL.1 | 0.720689815 | 1.017853301 | -0.428716287 | 0.06 | 0.098 | 0.02810254 | 1 |
| WBGene00015650 | pkg-2 | 1.248123588 | 1.543677801 | -0.426394596 | 0.148 | 0.302 | 2.16E-07 | 0.001377041 |
| WBGene00019554 | K09C4.10 | 0.444960333 | 0.740392102 | -0.426217948 | 0.041 | 0.1 | 0.000204773 | 1 |
| WBGene00020260 | T05E7.4 | 0.420236104 | 0.715243373 | -0.425605523 | 0.034 | 0.119 | 2.06E-07 | 0.001313194 |
| WBGene00006527 | tax-6 | 1.426520217 | 1.72149422 | -0.425557531 | 0.198 | 0.313 | 0.000242206 | 1 |
| WBGene00018214 | F39H12.3 | 1.422258892 | 1.715889882 | -0.423619973 | 0.191 | 0.373 | 4.06E-08 | 0.000259191 |
| WBGene00044588 | ZK1236.9 | 0.898450425 | 1.19140594 | -0.422645469 | 0.104 | 0.213 | 1.01E-05 | 0.064228947 |
| WBGene00013897 | ZC443.2 | 0.542713101 | 0.83451974 | -0.42098799 | 0.053 | 0.123 | 0.000103409 | 0.65985581 |
| WBGene00006604 | tra-1 | 0.479037262 | 0.770199742 | -0.420058667 | 0.048 | 0.112 | 0.000185796 | 1 |
| WBGene00017580 | lgc-4 | 0.313850129 | 0.604936189 | -0.419948416 | 0.023 | 0.069 | 0.000323276 | 1 |
| WBGene00012030 | T25G3.3 | 0.151287294 | 0.441461827 | -0.41863336 | 0.014 | 0.071 | 3.17E-06 | 0.020221289 |
| WBGene00012825 | Y43F8C.3 | 0.494402781 | 0.784572364 | -0.418626219 | 0.049 | 0.094 | 0.007392731 | 1 |
| WBGene00019871 | R04E5.8 | 0.705663252 | 0.995824618 | -0.418614363 | 0.071 | 0.165 | 7.56E-06 | 0.048265015 |
| WBGene00001475 | fmi-1 | 0.7180429 | 1.007697707 | -0.417883554 | 0.071 | 0.16 | 1.54E-05 | 0.098257602 |
| WBGene00016984 | npr-8 | 0.808042694 | 1.096880983 | -0.416705568 | 0.088 | 0.158 | 0.001698818 | 1 |
| WBGene00004492 | rps-23 | 2.742151112 | 3.03025864 | -0.415651302 | 0.606 | 0.785 | 5.19E-09 | 3.31E-05 |
| WBGene00003124 | mai-1 | 0.44868611 | 0.735822316 | -0.414249981 | 0.032 | 0.098 | 1.47E-05 | 0.093881101 |
| WBGene00008249 | srsx-28 | 1.837828705 | 2.124876441 | -0.414122344 | 0.295 | 0.5 | 6.68E-07 | 0.00426494 |
| WBGene00008956 | nekl-3 | 0.913651987 | 1.197815638 | -0.40996149 | 0.106 | 0.229 | 1.08E-06 | 0.006915792 |
| WBGene00004460 | rpn-3 | 0.664984538 | 0.948585593 | -0.409149836 | 0.081 | 0.152 | 0.000942776 | 1 |
| WBGene00000137 | amx-1 | 0.160477006 | 0.443441118 | -0.408230921 | 0.018 | 0.062 | 0.000211916 | 1 |
| WBGene00006595 | top-1 | 0.953965 | 1.236553742 | -0.407689376 | 0.11 | 0.213 | 5.82E-05 | 0.371505427 |
| WBGene00006924 | vig-1 | 0.321551603 | 0.603583995 | -0.406886733 | 0.027 | 0.067 | 0.001657028 | 1 |
| WBGene00019364 | mksr-1 | 0.576676557 | 0.858366364 | -0.406392487 | 0.055 | 0.096 | 0.017566013 | 1 |
| WBGene00019023 | F58A6.2 | 1.680024106 | 1.961166308 | -0.40560246 | 0.253 | 0.465 | 5.75E-09 | 3.67E-05 |
| WBGene00003407 | mrp-1 | 0.47923931 | 0.759947502 | -0.404976317 | 0.041 | 0.1 | 0.000201762 | 1 |
| WBGene00006483 | dgk-4 | 1.067390302 | 1.347645703 | -0.404323077 | 0.117 | 0.256 | 1.64E-07 | 0.00104596 |
| WBGene00004479 | rps-10 | 2.564084091 | 2.842902639 | -0.402250136 | 0.562 | 0.773 | 9.56E-10 | 6.10E-06 |
| WBGene00004416 | rpl-5 | 2.306276076 | 2.584364785 | -0.401197201 | 0.463 | 0.687 | 2.45E-06 | 0.01560577 |
| WBGene00009575 | F40F8.1 | 0.320320607 | 0.597783297 | -0.400294046 | 0.028 | 0.058 | 0.02075204 | 1 |
| WBGene00002018 | hsp-16.41 | 3.735677817 | 4.012415687 | -0.399248353 | 0.507 | 0.621 | 8.13E-05 | 0.51856563 |
| WBGene00004475 | rps-6 | 2.025759671 | 2.301900357 | -0.398386799 | 0.364 | 0.563 | 5.25E-06 | 0.033475147 |
| WBGene00013711 | Y106G6G.2 | 0.475280086 | 0.75127326 | -0.398173983 | 0.049 | 0.11 | 0.000484614 | 1 |
| WBGene00009691 | F44E5.4 | 4.221833618 | 4.497551443 | -0.397776738 | 0.867 | 0.91 | 5.74E-08 | 0.000366001 |
| WBGene00008366 | tpra-1 | 0.640410719 | 0.915556304 | -0.39695117 | 0.067 | 0.148 | 5.57E-05 | 0.355681086 |
| WBGene00000134 | amt-2 | 0.486848888 | 0.761550627 | -0.396310836 | 0.039 | 0.108 | 2.53E-05 | 0.16164838 |
| WBGene00044324 | ufm-1 | 0.139686641 | 0.413829319 | -0.395504282 | 0.014 | 0.062 | 4.11E-05 | 0.262557192 |
| WBGene00021697 | gcn-1 | 0.469738627 | 0.74341425 | -0.394830464 | 0.042 | 0.069 | 0.063855523 | 1 |
| WBGene00000491 | che-12 | 1.261225582 | 1.534075178 | -0.393638759 | 0.141 | 0.3 | 4.19E-08 | 0.000267239 |
| WBGene00012074 | T27A8.2 | 0.270018889 | 0.541795485 | -0.392090746 | 0.023 | 0.077 | 5.10E-05 | 0.325439231 |
| WBGene00009917 | glb-18 | 1.66012013 | 1.931325043 | -0.391265984 | 0.24 | 0.435 | 9.15E-08 | 0.000583883 |
| WBGene00219950 | F46A8.13 | 2.855391627 | 3.126145198 | -0.390614834 | 0.458 | 0.521 | 0.181838802 | 1 |
| WBGene00012340 | dct-15 | 0.598032928 | 0.868098607 | -0.389622416 | 0.065 | 0.152 | 1.91E-05 | 0.122002342 |
| WBGene00018193 | F39C12.1 | 0.420175146 | 0.688771718 | -0.387502942 | 0.028 | 0.065 | 0.00460553 | 1 |
| WBGene00005743 | srv-32 | 0.772121071 | 1.040472547 | -0.387149343 | 0.067 | 0.156 | 1.16E-05 | 0.074202758 |
| WBGene00004413 | rpl-2 | 2.491508109 | 2.758906491 | -0.385774319 | 0.53 | 0.738 | 4.53E-08 | 0.000289135 |
| WBGene00006603 | tps-2 | 0.765954208 | 1.032741545 | -0.384892767 | 0.087 | 0.162 | 0.000759921 | 1 |
| WBGene00006646 | tsp-20 | 0.341118288 | 0.607721578 | -0.384627244 | 0.028 | 0.096 | 5.72E-06 | 0.036483838 |
| WBGene00004344 | rgs-1 | 1.778266031 | 2.044593902 | -0.384229899 | 0.27 | 0.494 | 8.46E-09 | 5.40E-05 |
| WBGene00004758 | sek-1 | 0.873403531 | 1.139248871 | -0.383533754 | 0.101 | 0.223 | 1.05E-06 | 0.00668693 |

| | | | | | | | | |
|---|---|---|---|---|---|---|---|---|
| WBGene00015139 | B0310.3 | 0.985752876 | 1.251571072 | -0.383494594 | 0.113 | 0.235 | 3.12E-06 | 0.019901465 |
| WBGene00304791 | Y62H9A.16 | 0.157335434 | 0.422691958 | -0.382828541 | 0.012 | 0.05 | 0.000359805 | 1 |
| WBGene00022410 | Y97E10C.1 | 1.456673799 | 1.721376679 | -0.381885532 | 0.184 | 0.338 | 3.68E-06 | 0.023475098 |
| WBGene00007943 | C34F6.9 | 0.523735087 | 0.787249577 | -0.380171048 | 0.051 | 0.127 | 3.18E-05 | 0.202719986 |
| WBGene00009692 | F44E5.5 | 4.199271927 | 4.46199538 | -0.379029823 | 0.869 | 0.919 | 3.13E-11 | 2.00E-07 |
| WBGene00008439 | mfb-1 | 2.231252956 | 2.493702398 | -0.378634509 | 0.417 | 0.623 | 9.03E-07 | 0.005759839 |
| WBGene00004736 | sca-1 | 0.343239169 | 0.605435707 | -0.378269645 | 0.03 | 0.069 | 0.003626541 | 1 |
| WBGene00006805 | unc-73 | 0.289995141 | 0.551126864 | -0.376733443 | 0.028 | 0.058 | 0.019735644 | 1 |
| WBGene00022235 | sqd-1 | 0.972503711 | 1.233415055 | -0.376415502 | 0.104 | 0.194 | 0.00019494 | 1 |
| WBGene00009742 | F45H10.5 | 0.693538464 | 0.954299932 | -0.376199277 | 0.058 | 0.142 | 1.42E-05 | 0.090562455 |
| WBGene00021953 | Y55H10A.2 | 0.755931124 | 1.016523121 | -0.375954783 | 0.081 | 0.163 | 0.00012061 | 0.769613346 |
| WBGene00015424 | C04E6.8 | 4.569213669 | 4.829591113 | -0.375645247 | 0.892 | 0.933 | 1.67E-10 | 1.07E-06 |
| WBGene00001080 | dpy-21 | 0.884461374 | 1.14449538 | -0.375149771 | 0.11 | 0.15 | 0.099495838 | 1 |
| WBGene00004349 | rgs-6 | 0.626409982 | 0.886413486 | -0.375105766 | 0.06 | 0.127 | 0.000312148 | 1 |
| WBGene00019180 | H10D18.5 | 1.136956973 | 1.395263611 | -0.372657706 | 0.134 | 0.285 | 1.57E-07 | 0.001001606 |
| WBGene00017085 | E01A2.2 | 0.377051332 | 0.634915875 | -0.372019897 | 0.021 | 0.05 | 0.01101913 | 1 |
| WBGene00010642 | mks-6 | 0.874100795 | 1.131964252 | -0.37201833 | 0.097 | 0.206 | 7.04E-06 | 0.044935431 |
| WBGene00009676 | F44A6.4 | 0.238299287 | 0.496004494 | -0.371790023 | 0.019 | 0.056 | 0.001704763 | 1 |
| WBGene00004401 | rom-2 | 0.432049771 | 0.689165745 | -0.370939942 | 0.044 | 0.123 | 6.62E-06 | 0.042270234 |
| WBGene00004390 | rnp-7 | 0.986150773 | 1.243265821 | -0.370938605 | 0.133 | 0.219 | 0.001440586 | 1 |
| WBGene00006772 | unc-36 | 0.434969384 | 0.691729088 | -0.370425952 | 0.041 | 0.077 | 0.014391106 | 1 |
| WBGene00000241 | bbs-1 | 0.257453942 | 0.513527221 | -0.36943565 | 0.025 | 0.065 | 0.001505847 | 1 |
| WBGene00002016 | hsp-16.2 | 3.92337546 | 4.179027177 | -0.368827464 | 0.445 | 0.552 | 0.000808367 | 1 |
| WBGene00003905 | pad-1 | 0.218677691 | 0.473980744 | -0.368324448 | 0.019 | 0.065 | 0.000190716 | 1 |
| WBGene00017359 | F10E9.10 | 0.40843512 | 0.662352162 | -0.366324857 | 0.042 | 0.065 | 0.116194321 | 1 |
| WBGene00006752 | unc-13 | 1.107516774 | 1.361274433 | -0.366094916 | 0.138 | 0.275 | 1.93E-06 | 0.012336385 |
| WBGene00010677 | gtbp-1 | 0.704694531 | 0.958317355 | -0.36590039 | 0.071 | 0.16 | 2.11E-05 | 0.134567007 |
| WBGene00004431 | rpl-19 | 2.387641082 | 2.639773765 | -0.363750571 | 0.451 | 0.704 | 2.87E-10 | 1.83E-06 |
| WBGene00012281 | W05E10.2 | 3.360483082 | 3.611699131 | -0.362428148 | 0.707 | 0.863 | 2.79E-08 | 0.00017796 |
| WBGene00017724 | F22F7.7 | 0.531198313 | 0.782394343 | -0.362399265 | 0.051 | 0.096 | 0.007058233 | 1 |
| WBGene00001574 | gei-17 | 0.43050424 | 0.681500616 | -0.362111227 | 0.042 | 0.083 | 0.008654623 | 1 |
| WBGene00022134 | hrpu-2 | 0.224097265 | 0.475083528 | -0.362096638 | 0.019 | 0.06 | 0.000734099 | 1 |
| WBGene00016293 | C31H2.3 | 1.449969276 | 1.698590522 | -0.358684639 | 0.175 | 0.321 | 8.28E-07 | 0.005284257 |
| WBGene00009337 | uig-1 | 0.389482823 | 0.637295857 | -0.357518636 | 0.034 | 0.088 | 0.000214752 | 1 |
| WBGene00003724 | nhr-134 | 0.600384518 | 0.848121586 | -0.35740904 | 0.058 | 0.119 | 0.000848165 | 1 |
| WBGene00002179 | jph-1 | 0.32605168 | 0.573691599 | -0.357268884 | 0.03 | 0.075 | 0.001151554 | 1 |
| WBGene00019841 | R02F11.3 | 0.535846638 | 0.782669521 | -0.356090149 | 0.051 | 0.117 | 0.000211661 | 1 |
| WBGene00018846 | eef-1B.1 | 2.470846419 | 2.717111721 | -0.35528573 | 0.488 | 0.708 | 8.12E-08 | 0.000517941 |
| WBGene00017571 | jmjd-3.1 | 0.64761285 | 0.893866807 | -0.355269364 | 0.057 | 0.152 | 1.17E-06 | 0.00749524 |
| WBGene00016817 | C50E3.6 | 0.35820459 | 0.603921381 | -0.354494397 | 0.028 | 0.071 | 0.001383122 | 1 |
| WBGene00000295 | cat-1 | 1.598474544 | 1.844001326 | -0.35422027 | 0.231 | 0.413 | 2.61E-06 | 0.016658944 |
| WBGene00006741 | unc-1 | 0.316434322 | 0.561019075 | -0.35286121 | 0.028 | 0.067 | 0.003063056 | 1 |
| WBGene00199110 | C32D5.17 | 0.283227858 | 0.526940545 | -0.351603085 | 0.023 | 0.075 | 8.30E-05 | 0.529748299 |
| WBGene00000518 | ckk-1 | 0.715952748 | 0.959522795 | -0.351397299 | 0.076 | 0.173 | 8.56E-06 | 0.054595245 |
| WBGene00022139 | tub-2 | 0.359743193 | 0.602439317 | -0.350136495 | 0.034 | 0.058 | 0.067756289 | 1 |
| WBGene00014051 | spv-1 | 1.304525538 | 1.546535669 | -0.349146816 | 0.187 | 0.298 | 0.001212145 | 1 |
| WBGene00044029 | C05D12.7 | 1.867364772 | 2.10871588 | -0.348196047 | 0.29 | 0.487 | 2.17E-07 | 0.00138226 |
| WBGene00006702 | ubc-3 | 1.758353939 | 1.998649255 | -0.346672862 | 0.249 | 0.431 | 3.36E-07 | 0.002145482 |
| WBGene00001193 | egl-26 | 1.064178303 | 1.303554211 | -0.345346436 | 0.124 | 0.294 | 2.24E-09 | 1.43E-05 |
| WBGene00004418 | rpl-7 | 2.489164016 | 2.726734341 | -0.34274153 | 0.528 | 0.712 | 4.76E-07 | 0.003036439 |
| WBGene00009454 | F36A2.7 | 0.562847485 | 0.800054754 | -0.342217751 | 0.057 | 0.117 | 0.000768538 | 1 |
| WBGene00017557 | nep-11 | 0.542930586 | 0.779359481 | -0.341094795 | 0.044 | 0.121 | 1.04E-05 | 0.066365099 |
| WBGene00007405 | C07A9.9 | 0.505600578 | 0.741919518 | -0.340936163 | 0.041 | 0.104 | 0.000105043 | 0.670278819 |
| WBGene00009732 | cope-1 | 0.521785823 | 0.757367482 | -0.339872491 | 0.053 | 0.098 | 0.008942848 | 1 |
| WBGene00004504 | rpt-4 | 0.499575213 | 0.734637944 | -0.339123836 | 0.049 | 0.113 | 0.000251794 | 1 |
| WBGene00000833 | cts-1 | 0.71021384 | 0.944924316 | -0.33861564 | 0.076 | 0.152 | 0.000278926 | 1 |
| WBGene00004411 | rpc-1 | 0.53937044 | 0.773534359 | -0.337827124 | 0.057 | 0.106 | 0.004972915 | 1 |
| WBGene00009180 | nurf-1 | 1.746670057 | 1.980506241 | -0.337354304 | 0.261 | 0.379 | 0.002397363 | 1 |
| WBGene00017411 | F13A2.2 | 0.936813074 | 1.168738555 | -0.334597741 | 0.092 | 0.208 | 5.83E-07 | 0.003721644 |
| WBGene00020951 | sna-1 | 0.276062431 | 0.507088277 | -0.333299841 | 0.027 | 0.071 | 0.000868937 | 1 |
| WBGene00045336 | R11D1.12 | 2.399310229 | 2.630124589 | -0.332994731 | 0.392 | 0.563 | 7.32E-06 | 0.046718841 |
| WBGene00003559 | ncl-1 | 0.525646335 | 0.756249574 | -0.33269015 | 0.044 | 0.11 | 0.000116774 | 0.745133003 |
| WBGene00012768 | eef-1B.2 | 0.953803721 | 1.184255792 | -0.33247206 | 0.11 | 0.208 | 0.000120143 | 0.766630079 |
| WBGene00004486 | rps-17 | 2.66790293 | 2.898262801 | -0.332339043 | 0.576 | 0.765 | 1.30E-06 | 0.008269385 |

| | | | | | | | | |
|---|---|---|---|---|---|---|---|---|
| WBGene00005909 | srx-18 | 0.797523037 | 1.027697284 | -0.332071245 | 0.09 | 0.142 | 0.018668664 | 1 |
| WBGene00019177 | H10D18.1 | 1.09758747 | 1.327279708 | -0.331375853 | 0.133 | 0.267 | 1.68E-06 | 0.010688737 |
| WBGene00009503 | F37A8.5 | 1.611314124 | 1.840989998 | -0.331352244 | 0.24 | 0.363 | 0.001460606 | 1 |
| WBGene00003511 | mxl-3 | 2.087122894 | 2.316419409 | -0.330804945 | 0.341 | 0.554 | 2.41E-08 | 0.000153736 |
| WBGene00014215 | obr-3 | 0.777030178 | 1.006207161 | -0.330632497 | 0.064 | 0.121 | 0.002466608 | 1 |
| WBGene00004278 | rab-19 | 0.652990228 | 0.882137037 | -0.330588965 | 0.067 | 0.137 | 0.000431932 | 1 |
| WBGene00011108 | R07E3.7 | 0.248098424 | 0.477170001 | -0.330480429 | 0.019 | 0.054 | 0.002714806 | 1 |
| WBGene00000915 | hsp-90 | 2.537893395 | 2.765526394 | -0.328404999 | 0.528 | 0.717 | 2.11E-05 | 0.134459207 |
| WBGene00001854 | hil-3 | 0.479250504 | 0.70669604 | -0.328134547 | 0.039 | 0.094 | 0.000385507 | 1 |
| WBGene00017925 | F29B9.11 | 0.886352722 | 1.113610433 | -0.327863573 | 0.104 | 0.198 | 0.000174494 | 1 |
| WBGene00015919 | C17G10.7 | 2.426258712 | 2.653343501 | -0.3276141 | 0.438 | 0.646 | 1.03E-06 | 0.006556185 |
| WBGene00022412 | Y102A11A.2 | 0.52470656 | 0.751297373 | -0.326901443 | 0.049 | 0.087 | 0.022645978 | 1 |
| WBGene00002233 | kqt-1 | 0.849531681 | 1.076049687 | -0.326796404 | 0.09 | 0.169 | 0.000371429 | 1 |
| WBGene00004489 | rps-20 | 2.5888177 | 2.815233482 | -0.326648925 | 0.588 | 0.767 | 3.29E-06 | 0.020984351 |
| WBGene00003043 | lip-1 | 0.30681755 | 0.532523713 | -0.325625162 | 0.019 | 0.058 | 0.001179833 | 1 |
| WBGene00004445 | rpl-31 | 2.796745578 | 3.022295108 | -0.325399189 | 0.602 | 0.779 | 4.50E-06 | 0.028684492 |
| WBGene00008121 | C46F11.6 | 0.923578386 | 1.148724387 | -0.324817018 | 0.095 | 0.208 | 2.66E-06 | 0.016993147 |
| WBGene00013307 | Y57G11C.9 | 1.574526437 | 1.798642948 | -0.323331778 | 0.237 | 0.385 | 8.70E-05 | 0.554857386 |
| WBGene00007683 | C18E9.2 | 0.778029944 | 1.001059729 | -0.321763964 | 0.085 | 0.133 | 0.019884916 | 1 |
| WBGene00004472 | rps-3 | 2.31680791 | 2.538550892 | -0.319907499 | 0.454 | 0.667 | 6.50E-07 | 0.004149157 |
| WBGene00270303 | T28D9.16 | 0.528151907 | 0.748922241 | -0.318504265 | 0.051 | 0.113 | 0.00036819 | 1 |
| WBGene00015943 | tiar-1 | 0.352807119 | 0.573455408 | -0.318328192 | 0.03 | 0.062 | 0.015053014 | 1 |
| WBGene00018218 | F40A3.3 | 0.344520842 | 0.565096131 | -0.318222875 | 0.028 | 0.05 | 0.073452921 | 1 |
| WBGene00012621 | Y38H6C.8 | 1.324440537 | 1.544494922 | -0.317471369 | 0.127 | 0.208 | 0.001201959 | 1 |
| WBGene00002295 | let-19 | 0.413887396 | 0.633930111 | -0.317454533 | 0.037 | 0.081 | 0.002610348 | 1 |
| WBGene00010133 | F55H12.2 | 1.366718404 | 1.586327238 | -0.316828576 | 0.175 | 0.296 | 4.87E-05 | 0.310939841 |
| WBGene00004419 | rpl-7A | 2.142926467 | 2.362498865 | -0.31677601 | 0.398 | 0.606 | 2.05E-05 | 0.130852257 |
| WBGene00022199 | pfk-1.1 | 0.229497797 | 0.448872249 | -0.316490434 | 0.021 | 0.06 | 0.001430671 | 1 |
| WBGene00006739 | ulp-4 | 0.518024775 | 0.736583408 | -0.315313456 | 0.041 | 0.083 | 0.00584724 | 1 |
| WBGene00002151 | irk-3 | 1.302297587 | 1.52072901 | -0.315129931 | 0.161 | 0.323 | 1.27E-07 | 0.000807433 |
| WBGene00012031 | gpdh-3 | 0.227671603 | 0.445249678 | -0.313898809 | 0.019 | 0.063 | 0.000306406 | 1 |
| WBGene00017579 | F18G5.1 | 0.506703943 | 0.724248681 | -0.313850715 | 0.053 | 0.112 | 0.000874737 | 1 |
| WBGene00018985 | F56F10.2 | 0.353167407 | 0.569980766 | -0.312795557 | 0.034 | 0.069 | 0.009383095 | 1 |
| WBGene00002045 | icd-1 | 1.524072022 | 1.74078044 | -0.31264416 | 0.23 | 0.377 | 0.000177552 | 1 |
| WBGene00016794 | C49H3.9 | 0.110472711 | 0.326786607 | -0.312074986 | 0.011 | 0.052 | 8.78E-05 | 0.560526634 |
| WBGene00000906 | daf-10 | 0.83221393 | 1.048092295 | -0.311446646 | 0.083 | 0.142 | 0.005828322 | 1 |
| WBGene00015507 | C06A6.2 | 0.36935561 | 0.585084706 | -0.311231296 | 0.035 | 0.098 | 5.16E-05 | 0.32910092 |
| WBGene00001026 | dnj-8 | 0.230757239 | 0.446279311 | -0.310932625 | 0.021 | 0.052 | 0.007477873 | 1 |
| WBGene00000161 | apa-2 | 0.163603874 | 0.378874934 | -0.310570492 | 0.016 | 0.058 | 0.000269507 | 1 |
| WBGene00001209 | egl-45 | 0.269016516 | 0.484125267 | -0.310336329 | 0.025 | 0.056 | 0.010216158 | 1 |
| WBGene00004747 | sdc-3 | 0.281234803 | 0.496039627 | -0.309897855 | 0.028 | 0.067 | 0.003070948 | 1 |
| WBGene00016790 | C49H3.3 | 0.922079129 | 1.136799346 | -0.309775791 | 0.094 | 0.188 | 2.92E-05 | 0.186511413 |
| WBGene00005210 | srg-53 | 1.248983446 | 1.463234917 | -0.309099535 | 0.133 | 0.283 | 1.91E-07 | 0.001221486 |
| WBGene00016173 | fust-1 | 0.913722267 | 1.127699154 | -0.308703393 | 0.085 | 0.2 | 7.21E-07 | 0.004602886 |
| WBGene00021369 | siah-1 | 0.793425589 | 1.007392823 | -0.308689467 | 0.092 | 0.163 | 0.001812397 | 1 |
| WBGene00000912 | daf-16 | 1.194568978 | 1.407434865 | -0.307100559 | 0.141 | 0.26 | 5.36E-05 | 0.342236907 |
| WBGene00003902 | pab-1 | 1.365319458 | 1.57816409 | -0.307069896 | 0.186 | 0.342 | 3.15E-06 | 0.020081927 |
| WBGene00019285 | cbn-1 | 2.147842149 | 2.360683271 | -0.307064831 | 0.39 | 0.608 | 7.61E-06 | 0.048545003 |
| WBGene00012337 | scyl-1 | 0.580367905 | 0.793198529 | -0.307049687 | 0.049 | 0.112 | 0.000354046 | 1 |
| WBGene00004187 | prp-8 | 0.822661766 | 1.035455265 | -0.306996126 | 0.088 | 0.15 | 0.004786337 | 1 |
| WBGene00000230 | atp-3 | 0.737216442 | 0.949878095 | -0.306805912 | 0.078 | 0.163 | 8.45E-05 | 0.539371184 |
| WBGene00004274 | rab-11.1 | 1.079460961 | 1.291880973 | -0.306457299 | 0.127 | 0.246 | 2.05E-05 | 0.130938195 |
| WBGene00018765 | mpc-2 | 0.562324151 | 0.774679265 | -0.306363669 | 0.053 | 0.102 | 0.004120739 | 1 |
| WBGene00004822 | skr-16 | 0.335176325 | 0.547338528 | -0.306085359 | 0.032 | 0.077 | 0.00151149 | 1 |
| WBGene00020506 | dop-3 | 0.923041491 | 1.135113708 | -0.305955536 | 0.094 | 0.173 | 0.000711283 | 1 |
| WBGene00004494 | rps-25 | 3.174223925 | 3.386255957 | -0.305897561 | 0.724 | 0.863 | 2.12E-08 | 0.000135112 |
| WBGene00004476 | rps-7 | 2.565881555 | 2.777249891 | -0.30494005 | 0.539 | 0.708 | 0.000341212 | 1 |
| WBGene00022127 | yop-1 | 0.38962789 | 0.600568244 | -0.304322602 | 0.039 | 0.079 | 0.007233599 | 1 |
| WBGene00004414 | rpl-3 | 2.390391806 | 2.600834213 | -0.303604216 | 0.445 | 0.673 | 8.42E-09 | 5.37E-05 |
| WBGene00044134 | D1025.10 | 0.375031952 | 0.585429817 | -0.303539957 | 0.035 | 0.075 | 0.005525537 | 1 |
| WBGene00004345 | rgs-2 | 2.414527641 | 2.624912623 | -0.303521371 | 0.479 | 0.671 | 4.01E-06 | 0.025582736 |
| WBGene00003357 | mir-264 | 0.671009665 | 0.880896332 | -0.302802454 | 0.064 | 0.148 | 1.81E-05 | 0.115790104 |
| WBGene00016899 | C53C11.1 | 0.301116996 | 0.510730858 | -0.302408879 | 0.028 | 0.058 | 0.019072987 | 1 |
| WBGene00007772 | egrh-1 | 1.145715327 | 1.355024439 | -0.301969218 | 0.14 | 0.227 | 0.002151326 | 1 |

| | | | | | | | | |
|---|---|---|---|---|---|---|---|---|
| WBGene00004488 | rps-19 | 2.599304445 | 2.807409327 | -0.300231882 | 0.535 | 0.744 | 6.38E-07 | 0.004072061 |
| WBGene00011520 | nhr-213 | 0.541839794 | 0.748528712 | -0.298189077 | 0.055 | 0.112 | 0.001595652 | 1 |
| WBGene00045517 | R11D1.13 | 0.170077229 | 0.376664185 | -0.298041978 | 0.014 | 0.054 | 0.000317798 | 1 |
| WBGene00044484 | hot-9 | 2.459462254 | 2.665366346 | -0.297056811 | 0.463 | 0.663 | 4.12E-06 | 0.02631603 |
| WBGene00022592 | klu-2 | 0.810120923 | 1.015121116 | -0.295752762 | 0.085 | 0.146 | 0.004410105 | 1 |
| WBGene00001329 | epn-1 | 1.63889244 | 1.843190415 | -0.294739676 | 0.224 | 0.373 | 1.48E-05 | 0.094478822 |
| WBGene00000287 | cal-3 | 0.51996498 | 0.723631458 | -0.293828619 | 0.055 | 0.081 | 0.09185343 | 1 |
| WBGene00018710 | F52G3.1 | 0.473181387 | 0.676840004 | -0.293817276 | 0.037 | 0.087 | 0.001100604 | 1 |
| WBGene00022042 | icd-2 | 1.765798068 | 1.969315858 | -0.293614106 | 0.281 | 0.471 | 2.57E-06 | 0.016380931 |
| WBGene00016706 | C46C11.3 | 0.657343862 | 0.860688291 | -0.293364 | 0.062 | 0.102 | 0.031627542 | 1 |
| WBGene00004368 | ric-19 | 0.523746914 | 0.72703956 | -0.293289291 | 0.048 | 0.1 | 0.001745414 | 1 |
| WBGene00006789 | unc-54 | 5.137953262 | 5.340634866 | -0.292407745 | 0.984 | 0.983 | 4.03E-13 | 2.57E-09 |
| WBGene00010191 | dmsr-1 | 1.022700374 | 1.22523504 | -0.292195758 | 0.092 | 0.206 | 1.30E-06 | 0.008290683 |
| WBGene00016002 | gyf-1 | 1.32086288 | 1.523289249 | -0.292039519 | 0.189 | 0.319 | 0.000125537 | 0.801051679 |
| WBGene00006594 | tom-1 | 0.313021668 | 0.515315905 | -0.291848893 | 0.027 | 0.065 | 0.002612487 | 1 |
| WBGene00008167 | acox-1.5 | 0.30591295 | 0.507511961 | -0.290845893 | 0.027 | 0.073 | 0.000546728 | 1 |
| WBGene00003681 | nhr-91 | 0.312119483 | 0.513200445 | -0.290098507 | 0.018 | 0.056 | 0.000928538 | 1 |
| WBGene00015368 | srt-65 | 1.985323731 | 2.185297094 | -0.288500579 | 0.286 | 0.45 | 1.09E-05 | 0.069247111 |
| WBGene00011737 | sqst-1 | 1.210219038 | 1.410187645 | -0.288493718 | 0.161 | 0.283 | 0.000106924 | 0.682281745 |
| WBGene00010357 | H03A11.2 | 0.392935402 | 0.592746314 | -0.288266212 | 0.035 | 0.067 | 0.019569414 | 1 |
| WBGene00003587 | ned-8 | 0.291114181 | 0.490888211 | -0.288213003 | 0.03 | 0.052 | 0.074665027 | 1 |
| WBGene00016314 | C32D5.6 | 0.539730849 | 0.739060725 | -0.287572222 | 0.048 | 0.092 | 0.006173588 | 1 |
| WBGene00017015 | D1014.2 | 0.14410996 | 0.343349376 | -0.287441717 | 0.012 | 0.052 | 0.000220289 | 1 |
| WBGene00016509 | adss-1 | 0.269066522 | 0.468117761 | -0.287170235 | 0.028 | 0.065 | 0.004548216 | 1 |
| WBGene00002224 | klp-13 | 0.55981796 | 0.758685574 | -0.286905321 | 0.058 | 0.113 | 0.002284874 | 1 |
| WBGene00007223 | C01F6.9 | 0.277265718 | 0.475527148 | -0.286030781 | 0.027 | 0.063 | 0.0038133 | 1 |
| WBGene00004428 | rpl-16 | 2.399003054 | 2.597256475 | -0.286019227 | 0.5 | 0.658 | 0.000137571 | 0.87784064 |
| WBGene00010731 | K10C3.4 | 1.388132722 | 1.586085949 | -0.28558614 | 0.177 | 0.298 | 8.65E-05 | 0.55172783 |
| WBGene00013159 | Y53F4B.12 | 0.876726567 | 1.074361337 | -0.285126703 | 0.076 | 0.185 | 9.14E-07 | 0.005829588 |
| WBGene00000484 | che-2 | 0.249391342 | 0.446983326 | -0.285064976 | 0.019 | 0.054 | 0.002857272 | 1 |
| WBGene00006538 | tbb-4 | 1.327317924 | 1.524611866 | -0.284634991 | 0.166 | 0.237 | 0.0317797 | 1 |
| WBGene00004705 | rsp-8 | 0.451510798 | 0.648259021 | -0.283847686 | 0.042 | 0.106 | 0.000135008 | 0.861486809 |
| WBGene00044638 | F23A7.8 | 1.315098276 | 1.511772551 | -0.283741001 | 0.064 | 0.096 | 0.055992296 | 1 |
| WBGene00012850 | Y44A6C.1 | 0.334556978 | 0.530078723 | -0.282078251 | 0.034 | 0.083 | 0.000819624 | 1 |
| WBGene00004923 | snt-3 | 1.05517865 | 1.250281325 | -0.281473662 | 0.117 | 0.258 | 1.51E-07 | 0.000966059 |
| WBGene00004699 | rsp-2 | 0.944877748 | 1.138464331 | -0.279286402 | 0.102 | 0.177 | 0.001964477 | 1 |
| WBGene00004277 | rab-18 | 0.473208263 | 0.666670414 | -0.279106885 | 0.048 | 0.1 | 0.001933008 | 1 |
| WBGene00011299 | ikke-1 | 0.607165259 | 0.80056281 | -0.279013689 | 0.057 | 0.123 | 0.000326012 | 1 |
| WBGene00021547 | Y44E3A.1 | 0.354326712 | 0.547542683 | -0.278751723 | 0.032 | 0.079 | 0.00095742 | 1 |
| WBGene00003668 | nhr-78 | 0.166201032 | 0.359355658 | -0.278663221 | 0.016 | 0.054 | 0.000688755 | 1 |
| WBGene00000379 | cct-4 | 0.667285091 | 0.8599096 | -0.277898423 | 0.058 | 0.108 | 0.005670759 | 1 |
| WBGene00008168 | C48B4.3 | 1.270242222 | 1.462777945 | -0.277770332 | 0.173 | 0.296 | 0.000106469 | 0.679379952 |
| WBGene00021740 | Y50D4B.1 | 2.464164951 | 2.656377518 | -0.277304116 | 0.489 | 0.683 | 1.87E-06 | 0.011921513 |
| WBGene00044354 | ZK418.11 | 0.57555692 | 0.767581477 | -0.277032875 | 0.044 | 0.115 | 2.81E-05 | 0.179129649 |
| WBGene00016407 | unk-1 | 0.546126092 | 0.736437173 | -0.274560853 | 0.051 | 0.1 | 0.003801095 | 1 |
| WBGene00001168 | eef-1A.1 | 2.550588952 | 2.74066455 | -0.274221123 | 0.527 | 0.742 | 4.09E-06 | 0.026106776 |
| WBGene00019643 | K11C4.2 | 1.368050756 | 1.558009784 | -0.274052948 | 0.178 | 0.262 | 0.011574171 | 1 |
| WBGene00007647 | C17E4.10 | 1.589405641 | 1.778965698 | -0.273477354 | 0.226 | 0.354 | 0.000279655 | 1 |
| WBGene00006384 | taf-3 | 0.333526452 | 0.523016853 | -0.273376862 | 0.028 | 0.073 | 0.000973176 | 1 |
| WBGene00001999 | hrp-1 | 1.532450955 | 1.720552246 | -0.2713728 | 0.253 | 0.394 | 0.000363856 | 1 |
| WBGene00018344 | npr-27 | 0.670398188 | 0.857872633 | -0.270468452 | 0.064 | 0.14 | 0.000102463 | 0.653817395 |
| WBGene00004493 | rps-24 | 2.215329547 | 2.402644325 | -0.270238102 | 0.436 | 0.608 | 0.000147441 | 0.940823822 |
| WBGene00019888 | btbd-10 | 0.544905749 | 0.731952385 | -0.269851254 | 0.062 | 0.11 | 0.008354428 | 1 |
| WBGene00004436 | rpl-24.1 | 3.010138625 | 3.196802495 | -0.269299041 | 0.689 | 0.84 | 1.21E-05 | 0.077040282 |
| WBGene00012235 | W04A4.6 | 1.475248428 | 1.661896365 | -0.269276052 | 0.207 | 0.338 | 0.000252767 | 1 |
| WBGene00020827 | T26A8.4 | 0.415403786 | 0.601987525 | -0.269183434 | 0.037 | 0.075 | 0.009075792 | 1 |
| WBGene00001187 | egl-19 | 0.489963267 | 0.676433847 | -0.269020181 | 0.039 | 0.069 | 0.033448357 | 1 |
| WBGene00008144 | gasr-8 | 0.4295064 | 0.615878376 | -0.268877926 | 0.041 | 0.108 | 5.31E-05 | 0.338971119 |
| WBGene00004896 | smu-2 | 0.409858023 | 0.595910584 | -0.268417108 | 0.032 | 0.069 | 0.006264499 | 1 |
| WBGene00011729 | set-16 | 0.264194117 | 0.450135136 | -0.268256186 | 0.018 | 0.05 | 0.003525052 | 1 |
| WBGene00174531 | 21ur-10538 | 0.689751101 | 0.875494698 | -0.267971367 | 0.057 | 0.127 | 0.000127705 | 0.814886376 |
| WBGene00004321 | rcan-1 | 0.751148102 | 0.93679615 | -0.267833518 | 0.083 | 0.137 | 0.012632459 | 1 |
| WBGene00016294 | C31H2.4 | 0.343295456 | 0.528895177 | -0.267763797 | 0.023 | 0.085 | 9.40E-06 | 0.059980453 |
| WBGene00004425 | rpl-13 | 2.712490074 | 2.897850005 | -0.267417854 | 0.576 | 0.773 | 1.78E-07 | 0.001132832 |

| | | | | | | | | |
|---|---|---|---|---|---|---|---|---|
| WBGene00004719 | sad-1 | 0.349023248 | 0.534276335 | -0.26726371 | 0.023 | 0.065 | 0.000732724 | 1 |
| WBGene00016975 | nhr-173 | 0.871251496 | 1.056099766 | -0.266679683 | 0.064 | 0.133 | 0.000333156 | 1 |
| WBGene00016603 | met-1 | 0.957615578 | 1.142292271 | -0.266432149 | 0.102 | 0.179 | 0.00169287 | 1 |
| WBGene00001352 | evl-14 | 0.223337571 | 0.407729828 | -0.266021795 | 0.014 | 0.05 | 0.000822877 | 1 |
| WBGene00016803 | bath-42 | 0.270820628 | 0.454390177 | -0.264834878 | 0.027 | 0.063 | 0.004142485 | 1 |
| WBGene00004316 | rbf-1 | 0.393113775 | 0.576312229 | -0.264299501 | 0.039 | 0.069 | 0.034115244 | 1 |
| WBGene00002889 | let-765 | 0.640900996 | 0.824061236 | -0.26424437 | 0.062 | 0.1 | 0.034859948 | 1 |
| WBGene00015376 | C03B1.5 | 0.274600742 | 0.457730535 | -0.264200444 | 0.028 | 0.056 | 0.027563989 | 1 |
| WBGene00002583 | let-363 | 0.257332245 | 0.440408314 | -0.264122937 | 0.025 | 0.062 | 0.003339388 | 1 |
| WBGene00007016 | mdt-15 | 0.593109117 | 0.775821793 | -0.263598671 | 0.053 | 0.117 | 0.000261895 | 1 |
| WBGene00020135 | aakg-4 | 0.420374897 | 0.602795367 | -0.263177108 | 0.037 | 0.073 | 0.012521636 | 1 |
| WBGene00003690 | nhr-100 | 1.532988931 | 1.715232566 | -0.262921988 | 0.193 | 0.367 | 4.61E-07 | 0.002943548 |
| WBGene00017193 | F07C3.2 | 0.166780423 | 0.348956426 | -0.262824416 | 0.014 | 0.05 | 0.000815291 | 1 |
| WBGene00015303 | rga-6 | 1.337074448 | 1.519178269 | -0.26272028 | 0.163 | 0.292 | 3.37E-05 | 0.215278976 |
| WBGene00018855 | F55A4.3 | 1.064067131 | 1.245982273 | -0.262448074 | 0.136 | 0.233 | 0.001282461 | 1 |
| WBGene00009353 | sdhd-1 | 0.353748897 | 0.535276319 | -0.261888712 | 0.03 | 0.06 | 0.021921552 | 1 |
| WBGene00001207 | egl-43 | 0.743523822 | 0.924315691 | -0.260827533 | 0.076 | 0.14 | 0.00192818 | 1 |
| WBGene00015293 | C01C4.3 | 1.752154803 | 1.932658541 | -0.260411848 | 0.254 | 0.437 | 9.59E-07 | 0.006121263 |
| WBGene00006961 | xnp-1 | 0.420620674 | 0.600677538 | -0.259767144 | 0.039 | 0.083 | 0.003998665 | 1 |
| WBGene00006975 | zfp-1 | 0.718613645 | 0.898602201 | -0.259668598 | 0.081 | 0.158 | 0.000520276 | 1 |
| WBGene00020586 | frpr-18 | 1.548844554 | 1.728699843 | -0.259476334 | 0.21 | 0.348 | 0.00012846 | 0.819700916 |
| WBGene00003797 | npp-11 | 0.170093072 | 0.349930245 | -0.259450198 | 0.014 | 0.05 | 0.000865783 | 1 |
| WBGene00013924 | rad-23 | 0.372744106 | 0.552551675 | -0.259407487 | 0.032 | 0.054 | 0.082499999 | 1 |
| WBGene00003850 | odr-3 | 1.963563776 | 2.143315319 | -0.259326661 | 0.359 | 0.51 | 0.00473494 | 1 |
| WBGene00007971 | rpb-3 | 0.259498209 | 0.439091458 | -0.25909829 | 0.019 | 0.054 | 0.002857272 | 1 |
| WBGene00022248 | Y73B6BL.27 | 0.600910782 | 0.780363185 | -0.258895092 | 0.064 | 0.127 | 0.001017379 | 1 |
| WBGene00013354 | snx-6 | 0.437963143 | 0.61679424 | -0.257998738 | 0.035 | 0.052 | 0.199695066 | 1 |
| WBGene00011230 | nud-2 | 0.700992091 | 0.87967716 | -0.257788064 | 0.069 | 0.135 | 0.000953266 | 1 |
| WBGene00000554 | cnb-1 | 0.756352415 | 0.93408568 | -0.2564149 | 0.08 | 0.158 | 0.000285815 | 1 |
| WBGene00020868 | eif-1 | 0.948859795 | 1.126559479 | -0.256366454 | 0.101 | 0.165 | 0.007336 | 1 |
| WBGene00016212 | C29F5.3 | 0.672287967 | 0.849973041 | -0.256345376 | 0.069 | 0.121 | 0.0074067 | 1 |
| WBGene00001560 | gei-3 | 0.528010437 | 0.70508738 | -0.255468028 | 0.035 | 0.085 | 0.0008672 | 1 |
| WBGene00020051 | mfsd-6 | 0.413328239 | 0.590170835 | -0.255129936 | 0.046 | 0.077 | 0.043645878 | 1 |
| WBGene00011015 | R04F11.2 | 1.063198881 | 1.239938865 | -0.254981898 | 0.111 | 0.225 | 7.89E-06 | 0.050341658 |
| WBGene00021959 | Y57E12AL.6 | 0.521914686 | 0.698636131 | -0.254955152 | 0.053 | 0.108 | 0.001797863 | 1 |
| WBGene00044989 | Y37A1B.17 | 0.241828358 | 0.417946034 | -0.254084097 | 0.019 | 0.054 | 0.002983118 | 1 |
| WBGene00006715 | ubc-20 | 0.378726884 | 0.554837639 | -0.254074113 | 0.039 | 0.083 | 0.003723979 | 1 |
| WBGene00017338 | F10D7.1 | 0.222473091 | 0.398473415 | -0.253914795 | 0.018 | 0.052 | 0.002245407 | 1 |
| WBGene00000953 | del-2 | 0.431911767 | 0.607665309 | -0.253558763 | 0.037 | 0.09 | 0.000522781 | 1 |
| WBGene00013415 | ilcr-1 | 0.437382608 | 0.61285267 | -0.253149788 | 0.032 | 0.094 | 3.91E-05 | 0.249501735 |
| WBGene00004470 | rps-1 | 2.752945899 | 2.928377441 | -0.253094215 | 0.599 | 0.794 | 1.19E-05 | 0.0757445 |
| WBGene00022336 | Y82E9BR.3 | 2.129373437 | 2.304298449 | -0.252363446 | 0.382 | 0.579 | 3.91E-05 | 0.249542564 |
| WBGene00004471 | rps-2 | 2.1428995 | 2.317670321 | -0.252140996 | 0.424 | 0.583 | 0.003089859 | 1 |
| WBGene00019719 | M01H9.3 | 3.071563343 | 3.246190246 | -0.251933367 | 0.673 | 0.831 | 9.04E-05 | 0.576600997 |
| WBGene00016511 | C37H5.13 | 0.346664561 | 0.521001059 | -0.251514401 | 0.035 | 0.073 | 0.008410022 | 1 |
| WBGene00013899 | ZC443.4 | 1.918162791 | 2.092383914 | -0.25134795 | 0.311 | 0.479 | 4.03E-05 | 0.257036036 |
| WBGene00010936 | ztf-14 | 2.478317373 | 2.65231658 | -0.251027794 | 0.505 | 0.687 | 1.60E-05 | 0.102171224 |
| WBGene00008940 | F18H3.4 | 0.403373038 | 0.577035497 | -0.250541969 | 0.044 | 0.073 | 0.053239424 | 1 |
| WBGene00010967 | nduo-5 | 1.63538967 | 1.809007781 | -0.250477988 | 0.21 | 0.321 | 0.002793614 | 1 |
| WBGene00012885 | iscu-1 | 0.577046065 | 0.750589856 | -0.250370766 | 0.057 | 0.113 | 0.001644205 | 1 |
| WBGene00206362 | B0041.11 | 0.422761939 | 0.596138936 | -0.250130134 | 0.037 | 0.063 | 0.060413558 | 1 |
| WBGene00022400 | rpb-9 | 0.410645997 | 0.583961444 | -0.250041337 | 0.035 | 0.067 | 0.02208615 | 1 |
| WBGene00012548 | Y37D8A.6 | 2.381351811 | 2.554445274 | -0.24972108 | 0.481 | 0.675 | 1.00E-05 | 0.063841517 |
| WBGene00002245 | lag-1 | 1.725919684 | 1.898884914 | -0.24953608 | 0.269 | 0.454 | 2.65E-05 | 0.169406342 |
| WBGene00003564 | ncs-2 | 0.751080742 | 0.923781618 | -0.249154697 | 0.076 | 0.127 | 0.011832031 | 1 |
| WBGene00015803 | C15H9.9 | 0.202771535 | 0.374762763 | -0.248130891 | 0.019 | 0.058 | 0.001153389 | 1 |
| WBGene00016097 | C25E10.8 | 1.617642015 | 1.787677146 | -0.24530884 | 0.242 | 0.385 | 0.000181718 | 1 |
| WBGene00007997 | sre-13 | 2.153359632 | 2.322874875 | -0.2445588 | 0.366 | 0.544 | 5.04E-05 | 0.321800607 |
| WBGene00018213 | F39H12.2 | 1.548036511 | 1.717336383 | -0.244248086 | 0.224 | 0.408 | 2.65E-06 | 0.01692182 |
| WBGene00015745 | C13F10.6 | 0.388950163 | 0.557881593 | -0.243716537 | 0.03 | 0.063 | 0.012039142 | 1 |
| WBGene00015484 | atgl-1 | 0.301859998 | 0.470585609 | -0.243419602 | 0.023 | 0.058 | 0.003980563 | 1 |
| WBGene00000282 | cah-4 | 1.186702214 | 1.354573204 | -0.242186646 | 0.143 | 0.269 | 1.64E-05 | 0.104468216 |
| WBGene00001955 | hlh-11 | 0.464935653 | 0.632647687 | -0.24195732 | 0.042 | 0.075 | 0.030991895 | 1 |
| WBGene00004826 | skr-20 | 0.236765644 | 0.404198713 | -0.241554858 | 0.025 | 0.063 | 0.002342035 | 1 |

| | | | | | | | | |
|---|---|---|---|---|---|---|---|---|
| WBGene00016138 | flh-2 | 0.69265729 | 0.859950348 | -0.241352865 | 0.078 | 0.131 | 0.010642022 | 1 |
| WBGene00004430 | rpl-18 | 2.455852284 | 2.62282387 | -0.240889079 | 0.479 | 0.694 | 3.72E-06 | 0.023723488 |
| WBGene00018508 | F46F11.1 | 0.771354409 | 0.937973599 | -0.240380679 | 0.069 | 0.123 | 0.005257747 | 1 |
| WBGene00015750 | C14A11.5 | 1.758343587 | 1.924900115 | -0.240290276 | 0.272 | 0.421 | 0.000210975 | 1 |
| WBGene00004744 | scp-1 | 1.493130978 | 1.659603516 | -0.240169105 | 0.129 | 0.169 | 0.078101039 | 1 |
| WBGene00006703 | ubc-6 | 0.353357883 | 0.519770778 | -0.240083058 | 0.037 | 0.083 | 0.002240345 | 1 |
| WBGene00015799 | C15H9.3 | 2.11931857 | 2.285728676 | -0.240079035 | 0.36 | 0.569 | 4.82E-05 | 0.307725167 |
| WBGene00009625 | F41E7.9 | 0.696175683 | 0.862577922 | -0.240067685 | 0.072 | 0.106 | 0.111335537 | 1 |
| WBGene00002203 | kin-20 | 0.346932937 | 0.51295582 | -0.23952039 | 0.03 | 0.058 | 0.031831228 | 1 |
| WBGene00018803 | fbxa-24 | 2.798272616 | 2.964073382 | -0.239199943 | 0.505 | 0.625 | 0.000868455 | 1 |
| WBGene00011121 | R07E5.17 | 0.718544218 | 0.884049705 | -0.238773946 | 0.078 | 0.131 | 0.011292747 | 1 |
| WBGene00012826 | dyf-19 | 0.739574758 | 0.904604181 | -0.238087131 | 0.081 | 0.14 | 0.005789126 | 1 |
| WBGene00004238 | ptr-24 | 0.395883127 | 0.560275341 | -0.237167832 | 0.037 | 0.06 | 0.095719301 | 1 |
| WBGene00003839 | ocr-2 | 1.558716731 | 1.723038199 | -0.237065767 | 0.191 | 0.369 | 7.86E-08 | 0.000501396 |
| WBGene00000378 | cct-2 | 0.450309655 | 0.613471673 | -0.235393034 | 0.032 | 0.081 | 0.000605765 | 1 |
| WBGene00019162 | eif-1.A | 0.346113911 | 0.5091354 | -0.235190295 | 0.037 | 0.071 | 0.016404353 | 1 |
| WBGene00022250 | Y73B6BL.29 | 0.849442472 | 1.011992605 | -0.23451027 | 0.083 | 0.171 | 0.000106675 | 0.680691998 |
| WBGene00002244 | laf-1 | 0.348621568 | 0.511154432 | -0.234485357 | 0.025 | 0.058 | 0.007074116 | 1 |
| WBGene00004700 | rsp-3 | 1.114309648 | 1.276749852 | -0.234351677 | 0.141 | 0.25 | 0.000133644 | 0.852784807 |
| WBGene00010028 | F54B11.5 | 0.462754878 | 0.624803102 | -0.233786169 | 0.039 | 0.075 | 0.015059753 | 1 |
| WBGene00013548 | lury-1 | 1.213440203 | 1.375449614 | -0.233730174 | 0.134 | 0.237 | 0.00013711 | 0.874898038 |
| WBGene00012019 | dkf-2 | 0.636810976 | 0.798325451 | -0.233016132 | 0.065 | 0.123 | 0.002731181 | 1 |
| WBGene00013311 | sec-61.A | 0.688876757 | 0.850230243 | -0.232783874 | 0.06 | 0.131 | 0.000229735 | 1 |
| WBGene00018582 | F48A9.2 | 0.460331668 | 0.621381329 | -0.232345547 | 0.044 | 0.088 | 0.00496991 | 1 |
| WBGene00010626 | K07C5.3 | 0.231507716 | 0.391837969 | -0.231307661 | 0.019 | 0.052 | 0.004318839 | 1 |
| WBGene00011512 | T05H10.8 | 1.084308876 | 1.244461414 | -0.231051272 | 0.118 | 0.202 | 0.001009552 | 1 |
| WBGene00012366 | maea-1 | 0.347337386 | 0.506991294 | -0.230331902 | 0.027 | 0.058 | 0.012532685 | 1 |
| WBGene00008262 | ril-1 | 0.787925726 | 0.947157374 | -0.22972271 | 0.08 | 0.148 | 0.001077806 | 1 |
| WBGene00001853 | hil-2 | 1.877546912 | 2.036460166 | -0.229263364 | 0.258 | 0.425 | 4.31E-06 | 0.02748746 |
| WBGene00000914 | daf-19 | 0.636421287 | 0.795095014 | -0.228917799 | 0.058 | 0.079 | 0.235789142 | 1 |
| WBGene00012897 | pisy-1 | 0.431532136 | 0.590086957 | -0.228746254 | 0.048 | 0.063 | 0.293500373 | 1 |
| WBGene00009547 | F38H4.3 | 0.430137883 | 0.588662796 | -0.228703107 | 0.041 | 0.087 | 0.002803984 | 1 |
| WBGene00015776 | dct-1 | 1.726215876 | 1.884646615 | -0.228567241 | 0.276 | 0.437 | 8.24E-05 | 0.525489502 |
| WBGene00004053 | tank-1 | 0.464788097 | 0.623010105 | -0.228266107 | 0.041 | 0.063 | 0.102386175 | 1 |
| WBGene00004273 | rab-10 | 0.82243317 | 0.980466024 | -0.227993215 | 0.092 | 0.162 | 0.001839459 | 1 |
| WBGene00001971 | hmg-1.1 | 1.212186442 | 1.370116188 | -0.227844461 | 0.145 | 0.252 | 0.000311107 | 1 |
| WBGene00018285 | smk-1 | 0.875734662 | 1.033091098 | -0.22701735 | 0.101 | 0.146 | 0.060061694 | 1 |
| WBGene00000210 | asg-2 | 0.823986548 | 0.981200574 | -0.226811895 | 0.062 | 0.138 | 7.07E-05 | 0.450942094 |
| WBGene00022475 | Y119C1B.10 | 0.334381489 | 0.491523652 | -0.226708218 | 0.025 | 0.056 | 0.010743224 | 1 |
| WBGene00022817 | upp-1 | 1.268469067 | 1.424997809 | -0.22582324 | 0.177 | 0.281 | 0.00184782 | 1 |
| WBGene00004421 | rpl-10 | 2.860646294 | 3.01714948 | -0.22578637 | 0.647 | 0.802 | 0.000228139 | 1 |
| WBGene00007416 | ceh-57 | 1.67458257 | 1.830427388 | -0.224836546 | 0.249 | 0.421 | 1.44E-05 | 0.092062818 |
| WBGene00020334 | epg-7 | 0.521676538 | 0.67749838 | -0.224803399 | 0.032 | 0.069 | 0.006656591 | 1 |
| WBGene00004440 | rpl-26 | 2.229107725 | 2.384329525 | -0.223937721 | 0.424 | 0.619 | 0.000371715 | 1 |
| WBGene00016250 | hsp-110 | 2.034503146 | 2.189713741 | -0.223921557 | 0.35 | 0.519 | 0.000318565 | 1 |
| WBGene00019408 | K05F1.6 | 0.406232403 | 0.560818098 | -0.223020016 | 0.039 | 0.071 | 0.02556126 | 1 |
| WBGene00219592 | linc-84 | 0.26896599 | 0.423508013 | -0.222957009 | 0.025 | 0.056 | 0.010233068 | 1 |
| WBGene00010556 | rack-1 | 2.072525291 | 2.226904478 | -0.222722089 | 0.39 | 0.6 | 2.91E-05 | 0.185870306 |
| WBGene00022603 | gck-2 | 0.441648002 | 0.595596256 | -0.222100382 | 0.035 | 0.065 | 0.028440772 | 1 |
| WBGene00001840 | hel-1 | 1.175647005 | 1.329538728 | -0.222018826 | 0.125 | 0.212 | 0.001237985 | 1 |
| WBGene00020777 | T24E12.10 | 1.562935836 | 1.715941444 | -0.220740433 | 0.189 | 0.323 | 5.58E-05 | 0.356301217 |
| WBGene00007801 | trpa-1 | 0.329729635 | 0.482662479 | -0.220635456 | 0.03 | 0.067 | 0.005719505 | 1 |
| WBGene00011287 | glb-24 | 0.776544106 | 0.929311229 | -0.220396371 | 0.08 | 0.175 | 2.36E-05 | 0.150753527 |
| WBGene00022401 | Y97E10AR.6 | 0.603823882 | 0.756384866 | -0.220098975 | 0.067 | 0.119 | 0.006939218 | 1 |
| WBGene00017834 | F26F12.3 | 2.315887859 | 2.468390492 | -0.220014793 | 0.422 | 0.581 | 0.00077076 | 1 |
| WBGene00017311 | F09G2.2 | 0.360825796 | 0.513312913 | -0.219992407 | 0.034 | 0.079 | 0.001704069 | 1 |
| WBGene00012602 | Y38E10A.24 | 0.565997285 | 0.717856672 | -0.219086785 | 0.055 | 0.083 | 0.104881678 | 1 |
| WBGene00019748 | atg-2 | 1.087245426 | 1.238835228 | -0.218697855 | 0.136 | 0.194 | 0.035651494 | 1 |
| WBGene00002363 | let-92 | 0.856321142 | 1.007737557 | -0.218447711 | 0.095 | 0.15 | 0.017685859 | 1 |
| WBGene00021952 | vha-19 | 0.361245267 | 0.512634819 | -0.218408956 | 0.021 | 0.069 | 0.00016711 | 1 |
| WBGene00004148 | pqn-65 | 0.389386813 | 0.540766842 | -0.218395217 | 0.032 | 0.075 | 0.002159497 | 1 |
| WBGene00008405 | ttll-12 | 0.886908371 | 1.038046299 | -0.218045939 | 0.087 | 0.129 | 0.047951626 | 1 |
| WBGene00004704 | rsp-7 | 0.461019503 | 0.612035578 | -0.217870143 | 0.034 | 0.087 | 0.000317143 | 1 |
| WBGene00010868 | somi-1 | 0.366322913 | 0.516892055 | -0.217225354 | 0.032 | 0.056 | 0.060998756 | 1 |

| | | | | | | | | |
|---|---|---|---|---|---|---|---|---|
| WBGene00019733 | M02E1.1 | 0.965562269 | 1.115583536 | -0.216434939 | 0.104 | 0.156 | 0.03543734 | 1 |
| WBGene00004680 | rars-2 | 0.413531946 | 0.563453262 | -0.21629074 | 0.035 | 0.087 | 0.000632195 | 1 |
| WBGene00004423 | rpl-11.2 | 2.757088848 | 2.906867285 | -0.216084609 | 0.608 | 0.775 | 4.66E-06 | 0.029735701 |
| WBGene00014248 | ZK1307.7 | 1.529099248 | 1.677755401 | -0.214465494 | 0.2 | 0.34 | 2.98E-05 | 0.190029742 |
| WBGene00004046 | plp-1 | 0.951742108 | 1.100336879 | -0.21437694 | 0.095 | 0.167 | 0.002269245 | 1 |
| WBGene00015798 | C15H9.2 | 0.363167806 | 0.511302445 | -0.213713109 | 0.035 | 0.075 | 0.005729898 | 1 |
| WBGene00009579 | arrd-6 | 1.761028415 | 1.909108354 | -0.213634194 | 0.258 | 0.45 | 1.58E-06 | 0.010110261 |
| WBGene00006807 | unc-75 | 0.328053052 | 0.476014828 | -0.21346372 | 0.028 | 0.069 | 0.002344512 | 1 |
| WBGene00015552 | C06G3.6 | 0.21285244 | 0.360537059 | -0.213063868 | 0.021 | 0.05 | 0.011587991 | 1 |
| WBGene00006608 | tre-2 | 0.451038981 | 0.598605009 | -0.212892776 | 0.039 | 0.088 | 0.001322811 | 1 |
| WBGene00000475 | cey-4 | 0.548568143 | 0.696030698 | -0.212743496 | 0.049 | 0.098 | 0.004223437 | 1 |
| WBGene00000229 | atp-2 | 1.071494743 | 1.218921106 | -0.212691282 | 0.131 | 0.229 | 0.000478121 | 1 |
| WBGene00004464 | rpn-8 | 0.604565808 | 0.751123187 | -0.211437604 | 0.057 | 0.135 | 3.90E-05 | 0.248768624 |
| WBGene00011695 | T10G3.4 | 1.366625443 | 1.512998916 | -0.211172284 | 0.17 | 0.262 | 0.005757583 | 1 |
| WBGene00009050 | F22D6.2 | 2.216489383 | 2.362838188 | -0.211136696 | 0.431 | 0.619 | 0.000344887 | 1 |
| WBGene00004439 | rpl-25.2 | 1.294955727 | 1.440700966 | -0.210265934 | 0.18 | 0.288 | 0.002042749 | 1 |
| WBGene00009262 | F30A10.3 | 0.584767755 | 0.730144104 | -0.209733739 | 0.039 | 0.094 | 0.000410035 | 1 |
| WBGene00009380 | F34H10.3 | 0.555540392 | 0.700781683 | -0.20953889 | 0.051 | 0.112 | 0.000592173 | 1 |
| WBGene00007108 | gspd-1 | 0.32983125 | 0.474953112 | -0.20936659 | 0.032 | 0.071 | 0.00448061 | 1 |
| WBGene00022951 | C30E1.9 | 5.043635133 | 5.188648688 | -0.209210337 | 0.929 | 0.925 | 2.66E-05 | 0.170029665 |
| WBGene00206380 | C42D4.18 | 1.45432185 | 1.599255436 | -0.209094966 | 0.18 | 0.344 | 6.63E-07 | 0.004229027 |
| WBGene00021350 | Y37E3.8 | 3.13060127 | 3.275336375 | -0.208808618 | 0.728 | 0.842 | 0.000318464 | 1 |
| WBGene00005127 | srd-49 | 1.146189431 | 1.290906222 | -0.208782195 | 0.141 | 0.231 | 0.00220675 | 1 |
| WBGene00045192 | aho-3 | 0.971678031 | 1.116196051 | -0.20849543 | 0.12 | 0.202 | 0.002938691 | 1 |
| WBGene00003123 | mag-1 | 0.598755823 | 0.742836216 | -0.207864068 | 0.049 | 0.096 | 0.005573141 | 1 |
| WBGene00011240 | mask-1 | 0.566797291 | 0.710543688 | -0.207382214 | 0.053 | 0.087 | 0.042838369 | 1 |
| WBGene00006731 | uev-2 | 0.483799272 | 0.627540254 | -0.207374401 | 0.044 | 0.102 | 0.000563077 | 1 |
| WBGene00013136 | Y53C10A.5 | 0.886207704 | 1.029896572 | -0.207299217 | 0.102 | 0.16 | 0.015190937 | 1 |
| WBGene00014454 | MTCE.7 | 2.978155294 | 3.121765629 | -0.207185918 | 0.61 | 0.742 | 0.018532546 | 1 |
| WBGene00012893 | Y45F10D.15 | 0.364995995 | 0.508501941 | -0.207035316 | 0.032 | 0.071 | 0.004775615 | 1 |
| WBGene00002999 | lin-10 | 0.464267523 | 0.607524594 | -0.206676265 | 0.037 | 0.083 | 0.001972569 | 1 |
| WBGene00003920 | par-5 | 2.298284299 | 2.441289025 | -0.206312209 | 0.466 | 0.65 | 0.000963372 | 1 |
| WBGene00009716 | F44G4.7 | 1.140956634 | 1.283928289 | -0.206264498 | 0.131 | 0.244 | 5.89E-05 | 0.376076781 |
| WBGene00001994 | hpk-1 | 1.243770709 | 1.386697758 | -0.206200144 | 0.145 | 0.267 | 3.22E-05 | 0.205296613 |
| WBGene00002215 | klc-2 | 0.524998247 | 0.667655021 | -0.205810221 | 0.051 | 0.106 | 0.001550252 | 1 |
| WBGene00000280 | cah-2 | 0.502295416 | 0.644313435 | -0.204888691 | 0.035 | 0.094 | 0.000143379 | 0.914899594 |
| WBGene00004512 | rrn-1.1 | 1.241207443 | 1.382803146 | -0.204279418 | 0.11 | 0.219 | 1.93E-05 | 0.122850452 |
| WBGene00013766 | prmt-1 | 0.654301003 | 0.795707758 | -0.204006823 | 0.058 | 0.106 | 0.008683971 | 1 |
| WBGene00007554 | pptr-2 | 0.285808641 | 0.427072324 | -0.203800414 | 0.019 | 0.06 | 0.000759244 | 1 |
| WBGene00004480 | rps-11 | 2.97933117 | 3.120561951 | -0.203752948 | 0.677 | 0.812 | 0.000118313 | 0.754956273 |
| WBGene00020806 | T25F10.3 | 0.386914607 | 0.52799571 | -0.203537007 | 0.032 | 0.056 | 0.059721627 | 1 |
| WBGene00009584 | drap-1 | 1.781320027 | 1.922370741 | -0.203493165 | 0.269 | 0.437 | 5.83E-05 | 0.371959148 |
| WBGene00020917 | W01B11.6 | 1.805364742 | 1.946237693 | -0.203236707 | 0.286 | 0.433 | 0.000404896 | 1 |
| WBGene00011564 | ugt-50 | 0.262888355 | 0.403677654 | -0.203116024 | 0.021 | 0.05 | 0.011251439 | 1 |
| WBGene00003715 | nhr-125 | 0.504688543 | 0.644882385 | -0.202256961 | 0.048 | 0.075 | 0.081141972 | 1 |
| WBGene00006911 | vha-2 | 0.833445028 | 0.973012182 | -0.201352841 | 0.09 | 0.152 | 0.007267952 | 1 |
| WBGene00008342 | frpr-5 | 0.819906628 | 0.959413357 | -0.201265666 | 0.083 | 0.144 | 0.005086571 | 1 |
| WBGene00004267 | rab-3 | 1.201448279 | 1.340514513 | -0.200630166 | 0.141 | 0.269 | 1.04E-05 | 0.066340796 |
| WBGene00017150 | EGAP5.1 | 1.77340781 | 1.912156369 | -0.200171857 | 0.251 | 0.415 | 1.38E-05 | 0.08796776 |
| WBGene00000221 | atf-4 | 1.208563085 | 1.34718065 | -0.199982873 | 0.155 | 0.292 | 1.61E-05 | 0.102534075 |
| WBGene00001091 | drh-2 | 0.575510087 | 0.713930771 | -0.199698834 | 0.049 | 0.058 | 0.597241194 | 1 |
| WBGene00003754 | nlp-16 | 0.959109458 | 1.097339758 | -0.199424168 | 0.092 | 0.175 | 0.000316105 | 1 |
| WBGene00004364 | ric-4 | 1.383554919 | 1.521054403 | -0.198369823 | 0.186 | 0.298 | 0.001267337 | 1 |
| WBGene00003948 | pbs-2 | 0.57181984 | 0.709248912 | -0.19826824 | 0.046 | 0.096 | 0.002279658 | 1 |
| WBGene00000085 | aex-2 | 1.575638723 | 1.712429519 | -0.197347403 | 0.228 | 0.373 | 0.000115355 | 0.736080729 |
| WBGene00003015 | lin-29 | 0.694576887 | 0.831307243 | -0.197260207 | 0.064 | 0.125 | 0.001076326 | 1 |
| WBGene00006941 | wnk-1 | 0.689675643 | 0.825924383 | -0.196565382 | 0.055 | 0.104 | 0.004985473 | 1 |
| WBGene00011953 | ppm-2 | 0.724342221 | 0.860169484 | -0.195957318 | 0.074 | 0.121 | 0.021391209 | 1 |
| WBGene00004912 | sng-1 | 0.57124676 | 0.706695099 | -0.195410647 | 0.044 | 0.102 | 0.000489486 | 1 |
| WBGene00018373 | F43B10.1 | 0.309966542 | 0.445142361 | -0.195017483 | 0.025 | 0.054 | 0.015339509 | 1 |
| WBGene00006712 | ubc-17 | 0.504860591 | 0.639819916 | -0.194705149 | 0.046 | 0.083 | 0.020098156 | 1 |
| WBGene00000778 | cpn-2 | 1.499299684 | 1.634213324 | -0.19463924 | 0.219 | 0.358 | 9.81E-05 | 0.626111457 |
| WBGene00014309 | F07H5.5 | 2.299227108 | 2.433499773 | -0.193714507 | 0.442 | 0.619 | 0.000424804 | 1 |
| WBGene00000288 | cal-4 | 1.989949699 | 2.124030337 | -0.193437471 | 0.348 | 0.535 | 2.75E-05 | 0.175712062 |

| | | | | | | | | |
|---|---|---|---|---|---|---|---|---|
| WBGene00004410 | rla-2 | 2.491168116 | 2.625073091 | -0.193184043 | 0.507 | 0.698 | 0.000278837 | 1 |
| WBGene00009661 | patr-1 | 0.254327811 | 0.388029511 | -0.19289078 | 0.023 | 0.052 | 0.01352462 | 1 |
| WBGene00013042 | cccp-1 | 0.332970681 | 0.466518001 | -0.192668056 | 0.03 | 0.06 | 0.02337485 | 1 |
| WBGene00007861 | C32A9.1 | 0.404240196 | 0.537770549 | -0.192643577 | 0.025 | 0.079 | 7.67E-05 | 0.489630322 |
| WBGene00018788 | shc-1 | 0.419948444 | 0.553078909 | -0.192066662 | 0.041 | 0.062 | 0.141624539 | 1 |
| WBGene00003079 | lsm-5 | 0.366435737 | 0.499186617 | -0.191519036 | 0.028 | 0.069 | 0.002213416 | 1 |
| WBGene00011722 | rbm-22 | 0.203711155 | 0.335687062 | -0.190400988 | 0.018 | 0.05 | 0.003581937 | 1 |
| WBGene00000101 | aka-1 | 0.413087885 | 0.54467748 | -0.189843656 | 0.028 | 0.069 | 0.00219415 | 1 |
| WBGene00016703 | rdl-1 | 0.888440937 | 1.019981859 | -0.189773436 | 0.095 | 0.142 | 0.050662521 | 1 |
| WBGene00000567 | cnx-1 | 0.698609972 | 0.829887421 | -0.189393325 | 0.076 | 0.138 | 0.002962342 | 1 |
| WBGene00004185 | pro-1 | 0.619609367 | 0.750580058 | -0.188950766 | 0.049 | 0.102 | 0.002063978 | 1 |
| WBGene00020838 | trak-1 | 1.024449827 | 1.155188938 | -0.188616668 | 0.09 | 0.185 | 4.86E-05 | 0.309887593 |
| WBGene00004319 | rbr-2 | 0.960835053 | 1.09144086 | -0.188424349 | 0.102 | 0.162 | 0.016849324 | 1 |
| WBGene00001680 | gpb-2 | 0.323188458 | 0.453322067 | -0.187743112 | 0.027 | 0.056 | 0.018600434 | 1 |
| WBGene00002178 | jnk-1 | 0.725414274 | 0.855169433 | -0.187197126 | 0.064 | 0.14 | 8.88E-05 | 0.566411968 |
| WBGene00013343 | prp-6 | 0.34782319 | 0.477449333 | -0.187010993 | 0.027 | 0.058 | 0.012512879 | 1 |
| WBGene00020790 | T25B6.4 | 1.440001992 | 1.569256887 | -0.186475396 | 0.12 | 0.3 | 7.05E-11 | 4.50E-07 |
| WBGene00001784 | gst-36 | 1.157738538 | 1.286835352 | -0.186247333 | 0.124 | 0.194 | 0.004431335 | 1 |
| WBGene00015373 | C03B1.2 | 1.088842042 | 1.217907305 | -0.186201816 | 0.115 | 0.212 | 0.000266213 | 1 |
| WBGene00013962 | ZK287.1 | 0.605588026 | 0.734219615 | -0.185576156 | 0.051 | 0.115 | 0.000351791 | 1 |
| WBGene00006415 | tag-30 | 0.297042876 | 0.425407758 | -0.185191379 | 0.025 | 0.067 | 0.001068974 | 1 |
| WBGene00017663 | F21D12.3 | 1.308545756 | 1.436881821 | -0.185149804 | 0.171 | 0.3 | 8.25E-05 | 0.52634434 |
| WBGene00219763 | linc-94 | 0.370474285 | 0.498506251 | -0.184711083 | 0.034 | 0.063 | 0.027424983 | 1 |
| WBGene00000900 | daf-4 | 0.522635667 | 0.650657932 | -0.184697087 | 0.042 | 0.085 | 0.006224747 | 1 |
| WBGene00002210 | kin-29 | 0.469099126 | 0.596278103 | -0.18348048 | 0.042 | 0.079 | 0.016169262 | 1 |
| WBGene00004189 | pars-1 | 0.304040915 | 0.430914218 | -0.183039485 | 0.023 | 0.058 | 0.004381334 | 1 |
| WBGene00004740 | scd-2 | 2.223173453 | 2.349835191 | -0.182734261 | 0.392 | 0.594 | 0.000143492 | 0.915624081 |
| WBGene00010898 | nphp-1 | 0.622142023 | 0.748612633 | -0.182458521 | 0.069 | 0.11 | 0.032984867 | 1 |
| WBGene00012894 | tbc-17 | 0.865762427 | 0.992010989 | -0.182138174 | 0.099 | 0.163 | 0.006444898 | 1 |
| WBGene00001431 | fkb-6 | 1.078569859 | 1.204605929 | -0.181831613 | 0.138 | 0.21 | 0.011529983 | 1 |
| WBGene00016493 | C37A2.7 | 2.323044578 | 2.448731169 | -0.181327422 | 0.458 | 0.637 | 0.001631557 | 1 |
| WBGene00011282 | R74.8 | 0.492129213 | 0.617745696 | -0.181226277 | 0.037 | 0.06 | 0.103743194 | 1 |
| WBGene00003822 | nsy-1 | 0.850118318 | 0.975717649 | -0.181201532 | 0.087 | 0.154 | 0.002476456 | 1 |
| WBGene00004446 | rpl-32 | 2.832360043 | 2.957771024 | -0.1809298 | 0.617 | 0.779 | 0.00014983 | 0.95606476 |
| WBGene00044077 | pacs-1 | 0.405864472 | 0.530912573 | -0.180406276 | 0.041 | 0.081 | 0.008508689 | 1 |
| WBGene00015249 | otub-4 | 0.278059397 | 0.40305285 | -0.180327436 | 0.021 | 0.056 | 0.00336491 | 1 |
| WBGene00003858 | ogt-1 | 1.088061481 | 1.212844073 | -0.180023227 | 0.122 | 0.21 | 0.001287078 | 1 |
| WBGene00000140 | anc-1 | 0.75929664 | 0.883896288 | -0.179759295 | 0.074 | 0.115 | 0.037415958 | 1 |
| WBGene00004412 | rpl-1 | 2.494605787 | 2.618346948 | -0.178520759 | 0.518 | 0.704 | 0.000311647 | 1 |
| WBGene00006725 | ubl-1 | 2.524501622 | 2.648165296 | -0.17840897 | 0.518 | 0.698 | 0.000641055 | 1 |
| WBGene00017984 | F32D1.5 | 0.40006531 | 0.523630006 | -0.178266174 | 0.03 | 0.067 | 0.005765543 | 1 |
| WBGene00013898 | ZC443.3 | 1.496243249 | 1.619648964 | -0.178036813 | 0.191 | 0.354 | 1.45E-06 | 0.009243445 |
| WBGene00000135 | amt-3 | 0.567818193 | 0.691059123 | -0.177799079 | 0.055 | 0.102 | 0.007155787 | 1 |
| WBGene00007621 | C16C2.4 | 0.217869872 | 0.341051952 | -0.177714175 | 0.016 | 0.05 | 0.00176699 | 1 |
| WBGene00006791 | unc-57 | 0.67413309 | 0.797058213 | -0.177343465 | 0.062 | 0.123 | 0.001311749 | 1 |
| WBGene00044677 | ZK353.10 | 2.063503241 | 2.18639057 | -0.177288941 | 0.375 | 0.508 | 0.006375207 | 1 |
| WBGene00022522 | ZC132.3 | 0.602178961 | 0.72491026 | -0.177063836 | 0.06 | 0.113 | 0.0038942 | 1 |
| WBGene00004884 | smg-6 | 0.517677611 | 0.640408206 | -0.17706282 | 0.049 | 0.06 | 0.539798545 | 1 |
| WBGene00001664 | gpa-2 | 1.646667956 | 1.769335922 | -0.176972466 | 0.246 | 0.417 | 1.29E-05 | 0.082466699 |
| WBGene00022853 | ZK1127.6 | 0.321454021 | 0.444012677 | -0.176814765 | 0.028 | 0.05 | 0.070025272 | 1 |
| WBGene00017350 | F10E7.9 | 0.560180221 | 0.682389571 | -0.176310823 | 0.053 | 0.1 | 0.005967615 | 1 |
| WBGene00004487 | rps-18 | 2.600152588 | 2.722192609 | -0.176066534 | 0.549 | 0.717 | 0.000347406 | 1 |
| WBGene00019242 | srbc-40 | 1.228987257 | 1.351011596 | -0.176043908 | 0.147 | 0.235 | 0.002519853 | 1 |
| WBGene00017714 | F22F1.2 | 0.318158052 | 0.440094719 | -0.175917425 | 0.03 | 0.056 | 0.043530299 | 1 |
| WBGene00022794 | snu-23 | 0.247958283 | 0.369886894 | -0.175905802 | 0.019 | 0.05 | 0.006629474 | 1 |
| WBGene00020031 | suco-1 | 0.372727726 | 0.494481061 | -0.175652932 | 0.03 | 0.06 | 0.02311053 | 1 |
| WBGene00006960 | xbx-1 | 0.547704589 | 0.669295604 | -0.175418755 | 0.041 | 0.065 | 0.085388042 | 1 |
| WBGene00004491 | rps-22 | 2.525198364 | 2.646528745 | -0.175042739 | 0.534 | 0.715 | 0.000209278 | 1 |
| WBGene00010244 | F58D5.5 | 1.186119992 | 1.30730013 | -0.174825984 | 0.133 | 0.208 | 0.006680276 | 1 |
| WBGene00003630 | nhr-40 | 0.488786953 | 0.609912814 | -0.174747678 | 0.041 | 0.085 | 0.004013189 | 1 |
| WBGene00044387 | C27A2.8 | 1.87762366 | 1.998742229 | -0.174737158 | 0.295 | 0.442 | 0.000290721 | 1 |
| WBGene00011720 | T11G6.5 | 0.391344133 | 0.51220524 | -0.174365719 | 0.034 | 0.067 | 0.014101536 | 1 |
| WBGene00001788 | gst-40 | 2.042115542 | 2.162976445 | -0.174365424 | 0.318 | 0.5 | 7.28E-06 | 0.046423318 |
| WBGene00020854 | T27C4.1 | 1.702825494 | 1.823418206 | -0.173978508 | 0.281 | 0.402 | 0.013991173 | 1 |

| | | | | | | | | |
|---|---|---|---|---|---|---|---|---|
| WBGene00006831 | unc-104 | 0.750202587 | 0.869963237 | -0.172778095 | 0.067 | 0.121 | 0.004645855 | 1 |
| WBGene00001858 | hil-7 | 0.447663458 | 0.567067169 | -0.172263141 | 0.03 | 0.054 | 0.059449557 | 1 |
| WBGene00000377 | cct-1 | 0.43935541 | 0.558747295 | -0.172246081 | 0.037 | 0.079 | 0.00503639 | 1 |
| WBGene00003616 | nhr-17 | 0.823810416 | 0.943117002 | -0.17212302 | 0.081 | 0.14 | 0.005031167 | 1 |
| WBGene00004483 | rps-14 | 3.252875043 | 3.371942911 | -0.171778623 | 0.749 | 0.856 | 6.32E-05 | 0.403346578 |
| WBGene00021522 | nhr-274 | 0.809715339 | 0.928762678 | -0.171749006 | 0.083 | 0.156 | 0.00089521 | 1 |
| WBGene00003182 | mef-2 | 0.925958594 | 1.044843564 | -0.171514757 | 0.102 | 0.177 | 0.001914798 | 1 |
| WBGene00009367 | F33H2.3 | 0.551850514 | 0.670100725 | -0.170598993 | 0.046 | 0.1 | 0.001177119 | 1 |
| WBGene00195073 | F02E11.7 | 1.884649139 | 2.002695253 | -0.170304542 | 0.29 | 0.483 | 8.49E-06 | 0.054160503 |
| WBGene00003739 | nlp-1 | 0.733077696 | 0.850802375 | -0.169840811 | 0.076 | 0.117 | 0.044578453 | 1 |
| WBGene00006601 | tpi-1 | 0.453054114 | 0.570038674 | -0.168773045 | 0.037 | 0.085 | 0.001691131 | 1 |
| WBGene00013735 | Y111B2A.12 | 0.548713587 | 0.665621 | -0.168661744 | 0.046 | 0.077 | 0.045481857 | 1 |
| WBGene00018953 | F56C9.10 | 0.76092563 | 0.87775241 | -0.168545417 | 0.08 | 0.119 | 0.052950235 | 1 |
| WBGene00010971 | R01E6.7 | 0.554599641 | 0.671396125 | -0.168501708 | 0.042 | 0.094 | 0.001242915 | 1 |
| WBGene00006599 | tpa-1 | 0.534085578 | 0.64962042 | -0.166681544 | 0.051 | 0.079 | 0.079655831 | 1 |
| WBGene00004916 | snr-3 | 0.607982001 | 0.723242716 | -0.166286062 | 0.065 | 0.108 | 0.025513092 | 1 |
| WBGene00018764 | F53F10.2 | 1.447381248 | 1.562587718 | -0.166207803 | 0.226 | 0.3 | 0.094890243 | 1 |
| WBGene00004375 | rme-4 | 0.494874318 | 0.609435085 | -0.16527625 | 0.048 | 0.085 | 0.02220089 | 1 |
| WBGene00021539 | Y42H9AR.4 | 0.422903606 | 0.537390158 | -0.165169181 | 0.041 | 0.06 | 0.181606833 | 1 |
| WBGene00010631 | cash-1 | 0.346932553 | 0.460941439 | -0.164480054 | 0.025 | 0.054 | 0.016013439 | 1 |
| WBGene00219300 | W10G6.6 | 0.761535425 | 0.875165574 | -0.163933653 | 0.071 | 0.15 | 0.000114611 | 0.731334636 |
| WBGene00008920 | eef-1G | 2.21139557 | 2.324796836 | -0.163603445 | 0.415 | 0.594 | 0.000242003 | 1 |
| WBGene00000183 | arf-3 | 0.711954048 | 0.824651329 | -0.162587809 | 0.072 | 0.142 | 0.000740455 | 1 |
| WBGene00003825 | ntl-2 | 0.858274419 | 0.97093422 | -0.162533736 | 0.095 | 0.181 | 0.000355877 | 1 |
| WBGene00004474 | rps-5 | 2.680643132 | 2.79307666 | -0.162207294 | 0.572 | 0.75 | 0.000984166 | 1 |
| WBGene00002163 | ist-1 | 0.566569584 | 0.678915982 | -0.162081591 | 0.053 | 0.106 | 0.003013018 | 1 |
| WBGene00022271 | cpx-1 | 1.27624351 | 1.388325257 | -0.161699781 | 0.168 | 0.281 | 0.001131523 | 1 |
| WBGene00020397 | pcbd-1 | 0.650443338 | 0.76155813 | -0.160304759 | 0.06 | 0.102 | 0.020857719 | 1 |
| WBGene00007750 | syg-2 | 0.401069792 | 0.511968829 | -0.159993491 | 0.027 | 0.05 | 0.047564188 | 1 |
| WBGene00044294 | C01B10.11 | 0.411592621 | 0.522205528 | -0.159580693 | 0.034 | 0.058 | 0.067715788 | 1 |
| WBGene00003925 | pas-4 | 0.366984952 | 0.477419483 | -0.15932335 | 0.028 | 0.065 | 0.004785393 | 1 |
| WBGene00009920 | abts-1 | 1.344265603 | 1.454544344 | -0.159098592 | 0.166 | 0.258 | 0.003337229 | 1 |
| WBGene00004438 | rpl-25.1 | 1.338348793 | 1.448364718 | -0.158719429 | 0.177 | 0.313 | 3.62E-05 | 0.230814364 |
| WBGene00019829 | R02E12.5 | 0.998026913 | 1.107998257 | -0.158655113 | 0.101 | 0.169 | 0.005227917 | 1 |
| WBGene00004417 | rpl-6 | 2.337439734 | 2.447297158 | -0.158490761 | 0.428 | 0.656 | 1.13E-06 | 0.007222807 |
| WBGene00009374 | F34D10.4 | 1.56775177 | 1.677599908 | -0.158477363 | 0.184 | 0.275 | 0.002463717 | 1 |
| WBGene00006460 | ppm-1.A | 1.043308935 | 1.152907229 | -0.158116916 | 0.124 | 0.219 | 0.000531655 | 1 |
| WBGene00004831 | slo-2 | 0.571925956 | 0.681353552 | -0.15787065 | 0.049 | 0.113 | 0.000278956 | 1 |
| WBGene00014261 | shk-1 | 0.505718878 | 0.614982703 | -0.157634378 | 0.051 | 0.077 | 0.113970958 | 1 |
| WBGene00022748 | oaz-1 | 1.992883865 | 2.102119522 | -0.15759374 | 0.364 | 0.508 | 0.00621128 | 1 |
| WBGene00018466 | F45E1.5 | 1.157811855 | 1.266179564 | -0.156341557 | 0.14 | 0.235 | 0.001313466 | 1 |
| WBGene00021597 | spsb-1 | 0.345982446 | 0.454259538 | -0.156210823 | 0.027 | 0.05 | 0.052108245 | 1 |
| WBGene00001911 | his-37 | 1.884635425 | 1.992715404 | -0.155926451 | 0.297 | 0.452 | 0.000339408 | 1 |
| WBGene00020696 | pygl-1 | 0.473549406 | 0.581458765 | -0.155680296 | 0.046 | 0.062 | 0.28802687 | 1 |
| WBGene00005606 | srj-18 | 1.576494986 | 1.684334109 | -0.155578968 | 0.207 | 0.381 | 2.08E-06 | 0.013295747 |
| WBGene00020581 | T19D12.6 | 1.213879281 | 1.321466687 | -0.155215817 | 0.152 | 0.256 | 0.000733805 | 1 |
| WBGene00012149 | VF13D12L.3 | 0.333960214 | 0.441098429 | -0.154567771 | 0.03 | 0.063 | 0.012185436 | 1 |
| WBGene00001345 | fos-1 | 1.061897848 | 1.169002703 | -0.154519643 | 0.131 | 0.213 | 0.003044713 | 1 |
| WBGene00005111 | srd-33 | 1.333493184 | 1.440535801 | -0.154429853 | 0.166 | 0.26 | 0.003118607 | 1 |
| WBGene00006777 | unc-41 | 0.395216129 | 0.501862477 | -0.153858157 | 0.028 | 0.05 | 0.073452915 | 1 |
| WBGene00004495 | rps-26 | 3.149807027 | 3.256276897 | -0.153603553 | 0.747 | 0.842 | 0.000615795 | 1 |
| WBGene00012187 | rpb-11 | 0.363696036 | 0.469977439 | -0.153331654 | 0.032 | 0.067 | 0.009535334 | 1 |
| WBGene00010480 | tasp-1 | 1.484848002 | 1.590931673 | -0.153046386 | 0.207 | 0.335 | 0.000466336 | 1 |
| WBGene00017833 | F26F12.2 | 0.459142222 | 0.565188913 | -0.152993035 | 0.037 | 0.075 | 0.008794333 | 1 |
| WBGene00018700 | lmd-3 | 1.300549621 | 1.406509664 | -0.152868029 | 0.161 | 0.263 | 0.00100002 | 1 |
| WBGene00009141 | ncbp-2 | 0.660991255 | 0.766564345 | -0.152309774 | 0.071 | 0.14 | 0.000772062 | 1 |
| WBGene00016194 | C28H8.3 | 0.308359687 | 0.413788244 | -0.152101256 | 0.028 | 0.05 | 0.072895452 | 1 |
| WBGene00004427 | rpl-15 | 2.515150893 | 2.620516928 | -0.152011055 | 0.502 | 0.713 | 8.99E-06 | 0.057393157 |
| WBGene00019243 | fbxb-72 | 0.64877456 | 0.754126386 | -0.151990557 | 0.06 | 0.113 | 0.003879411 | 1 |
| WBGene00007235 | ahsa-1 | 0.651417546 | 0.756158456 | -0.151109192 | 0.067 | 0.11 | 0.02911202 | 1 |
| WBGene00194905 | Y53F4B.51 | 0.579615482 | 0.683809596 | -0.150320331 | 0.042 | 0.117 | 1.46E-05 | 0.09285851 |
| WBGene00001679 | gpb-1 | 1.685489599 | 1.789498775 | -0.150053523 | 0.279 | 0.387 | 0.022336209 | 1 |
| WBGene00006798 | unc-64 | 0.768436696 | 0.872405051 | -0.149994631 | 0.083 | 0.131 | 0.024821375 | 1 |
| WBGene00006412 | nlg-1 | 1.773030483 | 1.876895351 | -0.14984533 | 0.277 | 0.383 | 0.017276676 | 1 |

| | | | | | | | | |
|---|---|---|---|---|---|---|---|---|
| WBGene00004728 | sax-2 | 0.354605581 | 0.457919247 | -0.149050113 | 0.028 | 0.067 | 0.003405131 | 1 |
| WBGene00015248 | mai-2 | 1.083230547 | 1.186287292 | -0.148679455 | 0.104 | 0.223 | 3.22E-06 | 0.02053106 |
| WBGene00022836 | ZK973.11 | 0.308498211 | 0.411544224 | -0.148663972 | 0.034 | 0.073 | 0.005830725 | 1 |
| WBGene00017422 | F13C5.1 | 0.54731719 | 0.650337998 | -0.14862761 | 0.053 | 0.117 | 0.000415194 | 1 |
| WBGene00022027 | vps-20 | 0.42963898 | 0.532593408 | -0.148531843 | 0.035 | 0.067 | 0.021909023 | 1 |
| WBGene00008000 | slc-17.2 | 0.6784845 | 0.781374905 | -0.148439477 | 0.064 | 0.112 | 0.009574899 | 1 |
| WBGene00011059 | R06C1.4 | 2.372156975 | 2.474783952 | -0.14805943 | 0.424 | 0.627 | 0.000155108 | 0.98974514 |
| WBGene00001225 | eif-3.B | 0.265582011 | 0.368138002 | -0.147957019 | 0.016 | 0.05 | 0.001836703 | 1 |
| WBGene00019061 | F58F12.1 | 0.632524855 | 0.735064556 | -0.147933517 | 0.071 | 0.112 | 0.03293646 | 1 |
| WBGene00010367 | H05L14.2 | 0.581047358 | 0.683556538 | -0.147889486 | 0.055 | 0.102 | 0.007443212 | 1 |
| WBGene00007855 | C31H5.4 | 0.612147773 | 0.714638484 | -0.147862841 | 0.064 | 0.129 | 0.000807249 | 1 |
| WBGene00018880 | acc-3 | 0.610777606 | 0.713115999 | -0.147643092 | 0.062 | 0.087 | 0.154197426 | 1 |
| WBGene00003172 | mec-8 | 0.439810199 | 0.541727624 | -0.147035764 | 0.03 | 0.071 | 0.002512763 | 1 |
| WBGene00044386 | C27A2.7 | 0.375812445 | 0.477252602 | -0.146347211 | 0.037 | 0.065 | 0.043013845 | 1 |
| WBGene00004951 | spc-1 | 0.894570277 | 0.996008588 | -0.146344548 | 0.088 | 0.137 | 0.028794097 | 1 |
| WBGene00017098 | mek-5 | 0.761078768 | 0.862516497 | -0.146343709 | 0.065 | 0.104 | 0.039752331 | 1 |
| WBGene00010624 | mrps-15 | 0.437620385 | 0.538989341 | -0.146244491 | 0.039 | 0.081 | 0.00568786 | 1 |
| WBGene00008398 | D2005.3 | 0.411069794 | 0.511940378 | -0.145525491 | 0.041 | 0.079 | 0.011913101 | 1 |
| WBGene00007511 | C10C5.7 | 0.312596218 | 0.413361255 | -0.145373219 | 0.023 | 0.058 | 0.004075658 | 1 |
| WBGene00304822 | T14G8.6 | 0.413688865 | 0.514447387 | -0.14536382 | 0.035 | 0.081 | 0.002165965 | 1 |
| WBGene00006839 | unc-115 | 0.607947494 | 0.70817711 | -0.14460077 | 0.046 | 0.085 | 0.015233523 | 1 |
| WBGene00003000 | lin-11 | 1.142799385 | 1.242790743 | -0.144257037 | 0.145 | 0.233 | 0.003444182 | 1 |
| WBGene00302980 | F53E2.2 | 0.596764002 | 0.696506772 | -0.1438984 | 0.055 | 0.096 | 0.017841017 | 1 |
| WBGene00000396 | cdh-4 | 0.545632632 | 0.644993428 | -0.143347329 | 0.049 | 0.102 | 0.002243148 | 1 |
| WBGene00019466 | tos-1 | 3.626371784 | 3.725506812 | -0.143021614 | 0.806 | 0.846 | 0.02890641 | 1 |
| WBGene00010648 | K08C7.6 | 0.277401593 | 0.376516617 | -0.142992755 | 0.023 | 0.05 | 0.020308212 | 1 |
| WBGene00005742 | srv-31 | 0.354497278 | 0.453585318 | -0.142953823 | 0.032 | 0.063 | 0.019065978 | 1 |
| WBGene00008414 | D2030.4 | 0.34945094 | 0.448469271 | -0.142853256 | 0.025 | 0.058 | 0.007751206 | 1 |
| WBGene00019230 | ttll-11 | 1.036725584 | 1.135284762 | -0.142190838 | 0.102 | 0.2 | 9.04E-05 | 0.577098309 |
| WBGene00077437 | Y59H11AR.6 | 0.497232494 | 0.595138376 | -0.141248329 | 0.044 | 0.087 | 0.007688986 | 1 |
| WBGene00014096 | ZK829.7 | 0.309922892 | 0.407635537 | -0.140969549 | 0.03 | 0.062 | 0.016066268 | 1 |
| WBGene00004276 | rab-14 | 1.070933451 | 1.168134497 | -0.140231466 | 0.129 | 0.219 | 0.00169336 | 1 |
| WBGene00010239 | F58B4.6 | 2.429856458 | 2.526221359 | -0.139025165 | 0.47 | 0.652 | 0.001661263 | 1 |
| WBGene00001682 | gpc-2 | 2.093452765 | 2.189766602 | -0.138951494 | 0.376 | 0.563 | 0.000411985 | 1 |
| WBGene00017288 | F09E5.10 | 0.621574442 | 0.717836583 | -0.138876914 | 0.06 | 0.1 | 0.025638428 | 1 |
| WBGene00001665 | gpa-3 | 1.530487085 | 1.626607971 | -0.138673125 | 0.21 | 0.36 | 2.14E-05 | 0.136379503 |
| WBGene00015387 | kcnl-3 | 0.410623207 | 0.506550938 | -0.138394461 | 0.039 | 0.065 | 0.062841623 | 1 |
| WBGene00016998 | D1005.4 | 1.713935252 | 1.809571805 | -0.137974381 | 0.244 | 0.358 | 0.007540413 | 1 |
| WBGene00000479 | cgh-1 | 0.558855264 | 0.653920519 | -0.137150172 | 0.055 | 0.054 | 0.860083212 | 1 |
| WBGene00004432 | rpl-20 | 2.508294122 | 2.603229371 | -0.136962613 | 0.523 | 0.692 | 0.003433774 | 1 |
| WBGene00004434 | rpl-22 | 2.484819115 | 2.579545056 | -0.136660645 | 0.523 | 0.685 | 0.006954435 | 1 |
| WBGene00044509 | K07H8.12 | 0.862251422 | 0.956919914 | -0.136577764 | 0.088 | 0.179 | 9.35E-05 | 0.596472709 |
| WBGene00021004 | sftb-2 | 0.321017559 | 0.415497901 | -0.136306321 | 0.025 | 0.06 | 0.004923498 | 1 |
| WBGene00002189 | kin-1 | 1.123288509 | 1.217704682 | -0.136213745 | 0.145 | 0.192 | 0.181564738 | 1 |
| WBGene00000881 | cyn-5 | 0.471571399 | 0.565982984 | -0.136207126 | 0.041 | 0.075 | 0.019804951 | 1 |
| WBGene00013317 | Y57G11C.22 | 0.432477534 | 0.526466775 | -0.135597811 | 0.037 | 0.069 | 0.024402889 | 1 |
| WBGene00011060 | R06C1.6 | 2.426295806 | 2.52016466 | -0.135424129 | 0.493 | 0.663 | 0.001870325 | 1 |
| WBGene00020279 | glb-25 | 1.703836541 | 1.797564221 | -0.135220458 | 0.27 | 0.408 | 0.001104787 | 1 |
| WBGene00012289 | W05H12.1 | 0.813718549 | 0.907044991 | -0.134641594 | 0.081 | 0.138 | 0.007345756 | 1 |
| WBGene00001374 | exp-2 | 0.831665824 | 0.924888489 | -0.134491877 | 0.101 | 0.142 | 0.074237224 | 1 |
| WBGene00001087 | dpy-28 | 0.605281631 | 0.698450272 | -0.134413937 | 0.065 | 0.104 | 0.037981476 | 1 |
| WBGene00006965 | xtr-1 | 0.604222696 | 0.697284363 | -0.134259606 | 0.053 | 0.09 | 0.025440615 | 1 |
| WBGene00004285 | rab-37 | 1.577843334 | 1.670244743 | -0.133307056 | 0.228 | 0.365 | 0.000735181 | 1 |
| WBGene00045339 | T10H4.13 | 0.284771364 | 0.377003277 | -0.133062524 | 0.028 | 0.052 | 0.053841352 | 1 |
| WBGene00013638 | Y105C5A.14 | 0.288056149 | 0.38018352 | -0.132911701 | 0.025 | 0.05 | 0.033776177 | 1 |
| WBGene00011259 | miz-1 | 0.423613585 | 0.51568587 | -0.132832228 | 0.039 | 0.073 | 0.019326694 | 1 |
| WBGene00016494 | C37A2.8 | 0.590936281 | 0.682829647 | -0.132574103 | 0.057 | 0.092 | 0.036556786 | 1 |
| WBGene00018898 | F55F10.1 | 0.331248271 | 0.422464078 | -0.131596592 | 0.025 | 0.052 | 0.02256878 | 1 |
| WBGene00010639 | K07F5.15 | 0.34775468 | 0.438702933 | -0.131210593 | 0.03 | 0.067 | 0.00595305 | 1 |
| WBGene00006655 | tub-1 | 0.911374353 | 1.0022334 | -0.131081897 | 0.108 | 0.156 | 0.057556918 | 1 |
| WBGene00009783 | rer-1 | 0.381377338 | 0.472207383 | -0.131040055 | 0.032 | 0.056 | 0.065447009 | 1 |
| WBGene00000292 | cap-1 | 0.332980479 | 0.423651118 | -0.130810081 | 0.03 | 0.067 | 0.005547572 | 1 |
| WBGene00012484 | car-1 | 0.470279387 | 0.560837458 | -0.130647679 | 0.039 | 0.06 | 0.132011411 | 1 |
| WBGene00019396 | K04F10.3 | 0.358908524 | 0.449202933 | -0.130267297 | 0.035 | 0.065 | 0.030471941 | 1 |

| | | | | | | | | |
|---|---|---|---|---|---|---|---|---|
| WBGene00003884 | osm-3 | 0.734124659 | 0.824269574 | -0.130051621 | 0.076 | 0.127 | 0.011505548 | 1 |
| WBGene00019612 | gpcp-1 | 0.423335104 | 0.513466484 | -0.130032095 | 0.039 | 0.069 | 0.037168091 | 1 |
| WBGene00004477 | rps-8 | 2.748412503 | 2.838414239 | -0.129845058 | 0.594 | 0.746 | 0.003467705 | 1 |
| WBGene00006727 | ubq-1 | 1.787441088 | 1.876604464 | -0.128635561 | 0.283 | 0.444 | 0.00026565 | 1 |
| WBGene00009174 | F26H9.2 | 0.281610469 | 0.370711723 | -0.128545938 | 0.025 | 0.06 | 0.005398012 | 1 |
| WBGene00012094 | T27E9.2 | 0.76594516 | 0.854975469 | -0.128443586 | 0.069 | 0.117 | 0.011760968 | 1 |
| WBGene00010406 | math-33 | 0.367717414 | 0.456659633 | -0.128316498 | 0.037 | 0.056 | 0.167330035 | 1 |
| WBGene00003661 | nhr-71 | 0.246150686 | 0.334488673 | -0.127444776 | 0.021 | 0.05 | 0.01213044 | 1 |
| WBGene00003656 | nhr-66 | 0.764911869 | 0.852969787 | -0.127040722 | 0.072 | 0.119 | 0.018459203 | 1 |
| WBGene00018872 | F55C12.5 | 0.907063744 | 0.995058719 | -0.126949914 | 0.094 | 0.14 | 0.044816012 | 1 |
| WBGene00001976 | hmg-11 | 0.915199083 | 1.003153615 | -0.126891566 | 0.099 | 0.169 | 0.004307707 | 1 |
| WBGene00006067 | sto-5 | 0.531333744 | 0.618877168 | -0.126298465 | 0.035 | 0.094 | 0.000125906 | 0.803408469 |
| WBGene00010959 | nduo-1 | 1.685673593 | 1.773038995 | -0.126041632 | 0.237 | 0.308 | 0.129599624 | 1 |
| WBGene00002000 | hrp-2 | 0.94372676 | 1.031061721 | -0.125997715 | 0.094 | 0.14 | 0.035784726 | 1 |
| WBGene00011990 | T24D5.2 | 1.568855515 | 1.656162118 | -0.125956803 | 0.226 | 0.369 | 0.000307435 | 1 |
| WBGene00017814 | F26A10.2 | 0.52208874 | 0.609305785 | -0.125827598 | 0.037 | 0.071 | 0.016583395 | 1 |
| WBGene00000371 | cox-5B | 1.397061324 | 1.48413526 | -0.125621134 | 0.191 | 0.306 | 0.001283577 | 1 |
| WBGene00001565 | gei-8 | 0.457676888 | 0.544582585 | -0.125378418 | 0.044 | 0.067 | 0.123589739 | 1 |
| WBGene00004281 | rab-28 | 1.277484899 | 1.36423285 | -0.125150839 | 0.17 | 0.294 | 0.000205186 | 1 |
| WBGene00022738 | fubl-2 | 0.511759864 | 0.598155978 | -0.124643245 | 0.049 | 0.085 | 0.031280043 | 1 |
| WBGene00004128 | pqn-41 | 0.97270214 | 1.058969167 | -0.124457012 | 0.118 | 0.158 | 0.158809602 | 1 |
| WBGene00004893 | sms-2 | 0.387492712 | 0.473669267 | -0.124326487 | 0.028 | 0.06 | 0.015423063 | 1 |
| WBGene00006531 | tba-5 | 1.692462243 | 1.778322537 | -0.123870219 | 0.254 | 0.39 | 0.001785296 | 1 |
| WBGene00010697 | uda-1 | 0.849409658 | 0.935148041 | -0.123694341 | 0.083 | 0.131 | 0.029070929 | 1 |
| WBGene00004426 | rpl-14 | 2.474489236 | 2.560014456 | -0.123386811 | 0.479 | 0.662 | 0.000150099 | 0.957782283 |
| WBGene00004346 | rgs-3 | 2.130994583 | 2.216492727 | -0.123347749 | 0.371 | 0.544 | 0.001141063 | 1 |
| WBGene00004408 | rla-0 | 3.433591603 | 3.51905302 | -0.123294763 | 0.772 | 0.873 | 0.002008894 | 1 |
| WBGene00003003 | lin-14 | 0.594081938 | 0.679128476 | -0.122696219 | 0.055 | 0.1 | 0.009104509 | 1 |
| WBGene00001898 | his-24 | 1.217462774 | 1.302496452 | -0.122677666 | 0.131 | 0.238 | 0.000125477 | 0.800665855 |
| WBGene00018395 | mtch-1 | 1.019549576 | 1.104448503 | -0.122483261 | 0.122 | 0.171 | 0.103977582 | 1 |
| WBGene00016559 | C41A3.2 | 1.272850052 | 1.357080838 | -0.121519336 | 0.175 | 0.275 | 0.003429431 | 1 |
| WBGene00004478 | rps-9 | 2.839145526 | 2.923230651 | -0.121309193 | 0.62 | 0.777 | 0.016271562 | 1 |
| WBGene00043064 | nbet-1 | 0.565153669 | 0.649145123 | -0.121174053 | 0.051 | 0.092 | 0.014204094 | 1 |
| WBGene00019667 | spe-49 | 0.400977442 | 0.484962437 | -0.121164736 | 0.032 | 0.067 | 0.009621959 | 1 |
| WBGene00000403 | casy-1 | 2.434477535 | 2.518395214 | -0.12106762 | 0.484 | 0.654 | 0.002722569 | 1 |
| WBGene00008368 | D1053.3 | 1.37189594 | 1.455628844 | -0.120801044 | 0.155 | 0.298 | 3.99E-06 | 0.025468366 |
| WBGene00004415 | rpl-4 | 2.156861819 | 2.240513232 | -0.120683478 | 0.387 | 0.573 | 8.64E-05 | 0.551391417 |
| WBGene00009012 | F21D5.7 | 0.361641502 | 0.444621675 | -0.119715084 | 0.032 | 0.054 | 0.086907635 | 1 |
| WBGene00006697 | uaf-1 | 0.563032403 | 0.645959035 | -0.119637841 | 0.062 | 0.079 | 0.359378717 | 1 |
| WBGene00007042 | pbrm-1 | 0.588533244 | 0.671277858 | -0.119375246 | 0.06 | 0.108 | 0.009623258 | 1 |
| WBGene00004462 | rpn-6.1 | 0.353145583 | 0.435615919 | -0.118979545 | 0.034 | 0.06 | 0.052927232 | 1 |
| WBGene00004313 | rbc-1 | 0.452082468 | 0.534031507 | -0.118227472 | 0.035 | 0.069 | 0.015412108 | 1 |
| WBGene00007604 | xbx-9 | 1.260538931 | 1.342272998 | -0.117917334 | 0.177 | 0.283 | 0.002389485 | 1 |
| WBGene00001088 | dpy-30 | 0.662613743 | 0.744335567 | -0.11789967 | 0.06 | 0.137 | 8.76E-05 | 0.558724681 |
| WBGene00020114 | pde-4 | 1.419153865 | 1.50084572 | -0.117856435 | 0.178 | 0.292 | 0.000438329 | 1 |
| WBGene00011725 | lips-5 | 0.62238255 | 0.703857515 | -0.117543528 | 0.049 | 0.09 | 0.012089632 | 1 |
| WBGene00016200 | dpff-1 | 0.357829808 | 0.43908284 | -0.117223347 | 0.03 | 0.05 | 0.109678211 | 1 |
| WBGene00019322 | ahcy-1 | 0.687325531 | 0.768426866 | -0.117004493 | 0.076 | 0.112 | 0.07848838 | 1 |
| WBGene00004420 | rpl-9 | 2.39458155 | 2.475638072 | -0.116939843 | 0.475 | 0.673 | 0.000138981 | 0.886837633 |
| WBGene00005359 | srh-142 | 0.724577874 | 0.805158458 | -0.116253209 | 0.071 | 0.137 | 0.001065012 | 1 |
| WBGene00011605 | sftb-1 | 0.305532806 | 0.385323544 | -0.115113703 | 0.023 | 0.06 | 0.003069849 | 1 |
| WBGene00013577 | Y76A2B.5 | 0.907258745 | 0.986964644 | -0.114991305 | 0.09 | 0.169 | 0.000455108 | 1 |
| WBGene00018772 | F53G12.4 | 0.64353943 | 0.723122806 | -0.114814541 | 0.072 | 0.092 | 0.340200596 | 1 |
| WBGene00015568 | sre-3 | 0.972461179 | 1.051961982 | -0.114695415 | 0.11 | 0.188 | 0.0017018 | 1 |
| WBGene00000160 | apb-1 | 0.327029548 | 0.406290277 | -0.114349061 | 0.027 | 0.056 | 0.018123669 | 1 |
| WBGene00014004 | fic-1 | 0.530903068 | 0.610002814 | -0.114116811 | 0.055 | 0.094 | 0.022868202 | 1 |
| WBGene00022788 | slc-17.7 | 0.314535735 | 0.393559103 | -0.114006621 | 0.03 | 0.058 | 0.031962795 | 1 |
| WBGene00012896 | snrp-200 | 0.722418242 | 0.801440706 | -0.114005318 | 0.08 | 0.125 | 0.03188788 | 1 |
| WBGene00009574 | F40E10.6 | 0.402949784 | 0.481969011 | -0.114000647 | 0.041 | 0.058 | 0.218414093 | 1 |
| WBGene00004485 | rps-16 | 1.991431962 | 2.070408889 | -0.113939621 | 0.341 | 0.475 | 0.004630499 | 1 |
| WBGene00006572 | tin-9.1 | 0.610653487 | 0.68952733 | -0.113790902 | 0.064 | 0.112 | 0.011475641 | 1 |
| WBGene00022347 | Y82E9BR.14 | 0.620878515 | 0.699656176 | -0.113652141 | 0.071 | 0.119 | 0.016023228 | 1 |
| WBGene00012644 | rabx-5 | 0.333372506 | 0.41164231 | -0.112919457 | 0.028 | 0.063 | 0.007412108 | 1 |
| WBGene00008480 | eif-2A | 0.411060158 | 0.48919053 | -0.112718301 | 0.039 | 0.079 | 0.007550118 | 1 |

| | | | | | | | | |
|---|---|---|---|---|---|---|---|---|
| WBGene00015063 | B0228.6 | 0.946220234 | 1.023940848 | -0.112127145 | 0.092 | 0.138 | 0.043489067 | 1 |
| WBGene00016500 | memo-1 | 0.763299886 | 0.84082065 | -0.111838822 | 0.064 | 0.135 | 0.000341219 | 1 |
| WBGene00006707 | ubc-12 | 0.287717253 | 0.365109061 | -0.111652778 | 0.025 | 0.054 | 0.016713263 | 1 |
| WBGene00021527 | aakg-3 | 0.324939641 | 0.402267372 | -0.111560335 | 0.027 | 0.058 | 0.012914345 | 1 |
| WBGene00004424 | rpl-12 | 3.645503311 | 3.72261202 | -0.111244352 | 0.823 | 0.879 | 0.008265736 | 1 |
| WBGene00009780 | F46C5.4 | 0.348814201 | 0.425707993 | -0.110934292 | 0.032 | 0.052 | 0.117482112 | 1 |
| WBGene00012813 | Y43F8B.2 | 1.666357123 | 1.743176125 | -0.110826394 | 0.207 | 0.287 | 0.022826439 | 1 |
| WBGene00003830 | num-1 | 1.022958156 | 1.099599613 | -0.110570249 | 0.122 | 0.196 | 0.005858774 | 1 |
| WBGene00044728 | Y53F4B.45 | 1.831190398 | 1.907686752 | -0.110360911 | 0.23 | 0.342 | 0.001502444 | 1 |
| WBGene00001117 | dyf-1 | 0.990689923 | 1.067136129 | -0.110288563 | 0.117 | 0.2 | 0.001899669 | 1 |
| WBGene00004204 | swsn-4 | 0.752037289 | 0.82841105 | -0.110184047 | 0.072 | 0.113 | 0.03673355 | 1 |
| WBGene00007192 | B0491.5 | 0.538622916 | 0.614922853 | -0.11007754 | 0.044 | 0.067 | 0.123113427 | 1 |
| WBGene00002027 | hsr-9 | 0.505543992 | 0.581719654 | -0.109898249 | 0.037 | 0.092 | 0.000378117 | 1 |
| WBGene00006833 | unc-108 | 2.474155378 | 2.550218925 | -0.109736502 | 0.525 | 0.685 | 0.004286414 | 1 |
| WBGene00018187 | twf-2 | 0.874343658 | 0.950204685 | -0.109444327 | 0.088 | 0.154 | 0.003870387 | 1 |
| WBGene00001462 | flp-19 | 0.772357736 | 0.848145028 | -0.10933795 | 0.078 | 0.108 | 0.143412764 | 1 |
| WBGene00010484 | dph-3 | 0.377277564 | 0.452885231 | -0.109078807 | 0.035 | 0.065 | 0.030079435 | 1 |
| WBGene00015075 | B0238.11 | 0.274350584 | 0.349865844 | -0.10894549 | 0.021 | 0.05 | 0.01197468 | 1 |
| WBGene00017353 | F10E9.1 | 2.951699336 | 3.027039816 | -0.108693337 | 0.64 | 0.804 | 0.021813624 | 1 |
| WBGene00249815 | F20D12.12 | 0.492517531 | 0.56774225 | -0.108526329 | 0.049 | 0.065 | 0.329501985 | 1 |
| WBGene00006600 | tph-1 | 2.399118171 | 2.474306631 | -0.108474018 | 0.475 | 0.638 | 0.00588859 | 1 |
| WBGene00004409 | rla-1 | 2.543535377 | 2.618603097 | -0.108299828 | 0.495 | 0.687 | 0.000642514 | 1 |
| WBGene00012561 | Y37D8A.26 | 0.385671457 | 0.460549605 | -0.108026334 | 0.034 | 0.077 | 0.002581929 | 1 |
| WBGene00002083 | inf-1 | 0.505185795 | 0.580010681 | -0.107949493 | 0.049 | 0.081 | 0.051646996 | 1 |
| WBGene00018776 | F53H1.1 | 0.563259213 | 0.637749039 | -0.107466102 | 0.055 | 0.098 | 0.013325134 | 1 |
| WBGene00008530 | F02E9.5 | 0.654235656 | 0.728511373 | -0.107157208 | 0.067 | 0.123 | 0.004361458 | 1 |
| WBGene00021890 | Y54G2A.26 | 0.885774026 | 0.9598424 | -0.106858075 | 0.102 | 0.179 | 0.001769772 | 1 |
| WBGene00022435 | cfap-36 | 0.500130579 | 0.574168809 | -0.106814587 | 0.044 | 0.073 | 0.057591617 | 1 |
| WBGene00003640 | nhr-50 | 0.657901886 | 0.731803281 | -0.106617176 | 0.064 | 0.112 | 0.010923206 | 1 |
| WBGene00021643 | ccep-290 | 0.566109338 | 0.640007844 | -0.106613008 | 0.048 | 0.083 | 0.027777585 | 1 |
| WBGene00001167 | eef-2 | 2.302177604 | 2.375740655 | -0.106129049 | 0.435 | 0.621 | 0.001217955 | 1 |
| WBGene00006565 | tfg-1 | 0.356522477 | 0.429637073 | -0.105482064 | 0.03 | 0.056 | 0.045982637 | 1 |
| WBGene00003009 | lin-23 | 0.348271142 | 0.421260804 | -0.105301823 | 0.021 | 0.054 | 0.005420003 | 1 |
| WBGene00019776 | M6.4 | 0.565067055 | 0.637077905 | -0.103889697 | 0.051 | 0.102 | 0.003395637 | 1 |
| WBGene00017605 | F19F10.9 | 1.158885156 | 1.23079651 | -0.103746155 | 0.14 | 0.244 | 0.000468369 | 1 |
| WBGene00001440 | fkh-8 | 1.483747366 | 1.555354722 | -0.103307577 | 0.2 | 0.317 | 0.00117778 | 1 |
| WBGene00010507 | K02E2.6 | 0.905308982 | 0.976694449 | -0.102987459 | 0.097 | 0.167 | 0.003267339 | 1 |
| WBGene00012344 | ola-1 | 0.342652113 | 0.413811144 | -0.102660782 | 0.028 | 0.056 | 0.029895951 | 1 |
| WBGene00004207 | ptb-1 | 0.93941114 | 1.010096286 | -0.101977109 | 0.11 | 0.171 | 0.014927744 | 1 |
| WBGene00017711 | F22E5.13 | 0.661801421 | 0.732461872 | -0.101941482 | 0.051 | 0.106 | 0.001584002 | 1 |
| WBGene00004915 | snr-2 | 0.658974128 | 0.729634388 | -0.101941207 | 0.065 | 0.121 | 0.004319852 | 1 |
| WBGene00017039 | trk-1 | 1.455882278 | 1.526507445 | -0.101890579 | 0.214 | 0.29 | 0.066447552 | 1 |
| WBGene00006669 | twk-14 | 0.963660754 | 1.034102513 | -0.101625976 | 0.097 | 0.156 | 0.014660209 | 1 |
| WBGene00000380 | cct-5 | 1.611073873 | 1.6813597 | -0.101401014 | 0.239 | 0.36 | 0.002065211 | 1 |
| WBGene00001119 | dyf-3 | 0.982204571 | 1.052490166 | -0.101400679 | 0.104 | 0.19 | 0.000673816 | 1 |
| WBGene00016935 | ifta-1 | 1.32190668 | 1.391878644 | -0.100948205 | 0.164 | 0.283 | 0.000143259 | 0.914137527 |
| WBGene00008250 | srsx-29 | 1.856216975 | 1.926073364 | -0.100781467 | 0.297 | 0.454 | 0.000812509 | 1 |
| WBGene00019167 | H06I04.1 | 0.504731007 | 0.574257769 | -0.100305915 | 0.048 | 0.1 | 0.00210835 | 1 |
| WBGene00017637 | F20D6.8 | 1.825592293 | 1.755204591 | 0.101547989 | 0.283 | 0.404 | 0.022418475 | 1 |
| WBGene00012332 | rpi-1 | 0.954036116 | 0.883544176 | 0.101698371 | 0.106 | 0.171 | 0.013392917 | 1 |
| WBGene00006808 | unc-76 | 0.793845367 | 0.723343386 | 0.101712859 | 0.083 | 0.123 | 0.075322419 | 1 |
| WBGene00000910 | daf-14 | 1.214880824 | 1.144378472 | 0.101713393 | 0.154 | 0.198 | 0.204114357 | 1 |
| WBGene00022069 | eel-1 | 0.889640478 | 0.819102463 | 0.101764844 | 0.101 | 0.125 | 0.356897702 | 1 |
| WBGene00004184 | T19D2.2 | 1.79266207 | 1.721374679 | 0.102845966 | 0.297 | 0.398 | 0.164206367 | 1 |
| WBGene00003966 | pdl-1 | 1.730571685 | 1.659259601 | 0.10288159 | 0.263 | 0.373 | 0.039976547 | 1 |
| WBGene00009562 | flp-22 | 2.485806403 | 2.414388473 | 0.103034293 | 0.495 | 0.602 | 0.333005663 | 1 |
| WBGene00009116 | snx-27 | 0.653019171 | 0.581546448 | 0.103113343 | 0.06 | 0.071 | 0.56059707 | 1 |
| WBGene00004751 | sea-2 | 1.295752937 | 1.224273439 | 0.103123118 | 0.178 | 0.24 | 0.140992075 | 1 |
| WBGene00009660 | F43G6.8 | 1.879299958 | 1.80764495 | 0.103376324 | 0.313 | 0.413 | 0.079997762 | 1 |
| WBGene00018252 | F40H3.6 | 0.506228169 | 0.434559868 | 0.103395502 | 0.051 | 0.062 | 0.562727518 | 1 |
| WBGene00016100 | C25E10.11 | 1.630325503 | 1.557972032 | 0.104383992 | 0.244 | 0.288 | 0.657600541 | 1 |
| WBGene00020390 | T10B5.4 | 2.076288685 | 2.003853656 | 0.104501658 | 0.343 | 0.492 | 0.023683208 | 1 |
| WBGene00003568 | ncx-3 | 0.743288988 | 0.670770093 | 0.10462265 | 0.076 | 0.102 | 0.226318214 | 1 |
| WBGene00004463 | rpn-7 | 0.626272049 | 0.553723365 | 0.104665626 | 0.058 | 0.087 | 0.115376118 | 1 |

| | | | | | | | | |
|---|---|---|---|---|---|---|---|---|
| WBGene00003163 | mdl-1 | 1.428154548 | 1.35551326 | 0.104799226 | 0.193 | 0.244 | 0.200810342 | 1 |
| WBGene00003964 | pdi-3 | 1.039260138 | 0.966588249 | 0.104843374 | 0.115 | 0.146 | 0.269209615 | 1 |
| WBGene00304815 | Y40A1A.6 | 1.665144941 | 1.59214655 | 0.105314417 | 0.069 | 0.079 | 0.60515349 | 1 |
| WBGene00007129 | B0272.3 | 0.622531688 | 0.549044179 | 0.106020065 | 0.069 | 0.083 | 0.532611435 | 1 |
| WBGene00002002 | hsb-1 | 0.463890728 | 0.390152257 | 0.106382127 | 0.044 | 0.056 | 0.45624871 | 1 |
| WBGene00010963 | nduo-4 | 1.902139109 | 1.828291259 | 0.106539926 | 0.297 | 0.377 | 0.3202681 | 1 |
| WBGene00004919 | snr-6 | 0.839994364 | 0.765946378 | 0.106828663 | 0.09 | 0.125 | 0.139983635 | 1 |
| WBGene00004701 | rsp-4 | 0.46311435 | 0.389016299 | 0.106900891 | 0.032 | 0.06 | 0.037034996 | 1 |
| WBGene00000482 | chd-3 | 0.734540129 | 0.660272151 | 0.107146043 | 0.078 | 0.081 | 0.997359886 | 1 |
| WBGene00001496 | frm-10 | 1.621943467 | 1.547578196 | 0.107286407 | 0.269 | 0.294 | 0.821067578 | 1 |
| WBGene00018509 | xrep-4 | 0.739722599 | 0.665194907 | 0.107520731 | 0.078 | 0.117 | 0.069424002 | 1 |
| WBGene00008587 | F08G12.3 | 0.697971818 | 0.62290113 | 0.108304109 | 0.067 | 0.081 | 0.503206106 | 1 |
| WBGene00017735 | did-2 | 0.742885951 | 0.667707106 | 0.108460146 | 0.067 | 0.085 | 0.37633517 | 1 |
| WBGene00008999 | myrf-2 | 1.376702153 | 1.301131919 | 0.109024801 | 0.184 | 0.24 | 0.162953905 | 1 |
| WBGene00011156 | rbm-3.2 | 2.729596271 | 2.653980177 | 0.109090965 | 0.585 | 0.692 | 0.443418817 | 1 |
| WBGene00009804 | F47B8.3 | 1.183648666 | 1.107496054 | 0.109864996 | 0.148 | 0.221 | 0.02915985 | 1 |
| WBGene00000938 | dcp-66 | 0.856804518 | 0.780464816 | 0.110134909 | 0.081 | 0.117 | 0.089990624 | 1 |
| WBGene00012139 | T28F4.4 | 0.497695324 | 0.421281284 | 0.110242157 | 0.039 | 0.054 | 0.282870624 | 1 |
| WBGene00011270 | R31.2 | 0.810691262 | 0.734160655 | 0.110410327 | 0.085 | 0.106 | 0.396488764 | 1 |
| WBGene00003916 | par-1 | 0.638612516 | 0.561612141 | 0.11108806 | 0.053 | 0.085 | 0.060667869 | 1 |
| WBGene00011723 | T12A7.2 | 0.801440309 | 0.724379354 | 0.111175459 | 0.065 | 0.115 | 0.008721804 | 1 |
| WBGene00008746 | F13E6.2 | 0.440932415 | 0.363810679 | 0.111263147 | 0.039 | 0.05 | 0.437075301 | 1 |
| WBGene00004888 | smo-1 | 1.328458179 | 1.251210177 | 0.11144531 | 0.163 | 0.238 | 0.025445433 | 1 |
| WBGene00016397 | maph-9 | 0.838183304 | 0.760732238 | 0.111738269 | 0.088 | 0.117 | 0.236167622 | 1 |
| WBGene00000122 | aly-3 | 1.078514662 | 1.000989531 | 0.111845122 | 0.115 | 0.162 | 0.103287701 | 1 |
| WBGene00012471 | Y17G7B.20 | 1.667689006 | 1.589953329 | 0.112148876 | 0.237 | 0.321 | 0.119503649 | 1 |
| WBGene00002065 | iff-2 | 1.74262139 | 1.66448423 | 0.112728094 | 0.269 | 0.342 | 0.352981462 | 1 |
| WBGene00001082 | dpy-23 | 0.708719182 | 0.630307637 | 0.113123948 | 0.065 | 0.085 | 0.331702466 | 1 |
| WBGene00016463 | C35E7.11 | 1.266421847 | 1.187943549 | 0.113220252 | 0.171 | 0.215 | 0.349944251 | 1 |
| WBGene00012543 | nkcc-1 | 0.534115331 | 0.455630892 | 0.113229112 | 0.039 | 0.058 | 0.185109502 | 1 |
| WBGene00001946 | his-72 | 2.166927459 | 2.08767136 | 0.114342381 | 0.383 | 0.49 | 0.249977456 | 1 |
| WBGene00001748 | gsp-2 | 0.969191052 | 0.889934049 | 0.114343685 | 0.104 | 0.171 | 0.010205461 | 1 |
| WBGene00000065 | act-3 | 0.635556562 | 0.556077783 | 0.114663641 | 0.069 | 0.087 | 0.402793689 | 1 |
| WBGene00010425 | lpin-1 | 1.046603797 | 0.966936797 | 0.114935185 | 0.11 | 0.15 | 0.102399055 | 1 |
| WBGene00021956 | Y57E12AL.1 | 0.601819574 | 0.520809915 | 0.116872234 | 0.062 | 0.075 | 0.492358753 | 1 |
| WBGene00019185 | H10E21.5 | 1.230684633 | 1.148804034 | 0.118128733 | 0.145 | 0.227 | 0.009712775 | 1 |
| WBGene00001909 | his-35 | 1.924829663 | 1.842492428 | 0.118787521 | 0.329 | 0.417 | 0.20605467 | 1 |
| WBGene00008149 | pyp-1 | 0.909947964 | 0.827545494 | 0.118881635 | 0.09 | 0.119 | 0.224418133 | 1 |
| WBGene00015729 | C13A10.2 | 0.588686704 | 0.506169196 | 0.1190476 | 0.062 | 0.065 | 0.926275764 | 1 |
| WBGene00016016 | C23G10.10 | 0.884954389 | 0.802423001 | 0.119067624 | 0.083 | 0.115 | 0.151135834 | 1 |
| WBGene00001070 | dpy-8 | 0.567482976 | 0.484841461 | 0.119226504 | 0.048 | 0.052 | 0.820617592 | 1 |
| WBGene00020092 | pcf-11 | 0.630278495 | 0.547334671 | 0.119662644 | 0.064 | 0.088 | 0.186861074 | 1 |
| WBGene00011302 | R166.2 | 0.549421209 | 0.466467012 | 0.119677608 | 0.057 | 0.067 | 0.558896786 | 1 |
| WBGene00012016 | inpp-1 | 0.81942713 | 0.736260631 | 0.119983895 | 0.074 | 0.125 | 0.014082236 | 1 |
| WBGene00016235 | C29H12.2 | 1.236205235 | 1.15280387 | 0.120322736 | 0.155 | 0.227 | 0.033946567 | 1 |
| WBGene00020374 | serp-1.2 | 0.504740865 | 0.421234539 | 0.120474162 | 0.053 | 0.065 | 0.477043857 | 1 |
| WBGene00018757 | tag-304 | 0.874148557 | 0.790527922 | 0.120639076 | 0.095 | 0.115 | 0.494979493 | 1 |
| WBGene00007848 | cytb-5.1 | 1.088097313 | 1.004138271 | 0.121127293 | 0.118 | 0.156 | 0.184690743 | 1 |
| WBGene00014031 | ZK637.14 | 0.925650637 | 0.841414386 | 0.121527222 | 0.104 | 0.142 | 0.165545563 | 1 |
| WBGene00003060 | lpd-3 | 0.57659777 | 0.492167815 | 0.121806678 | 0.042 | 0.065 | 0.12268246 | 1 |
| WBGene00001149 | bcat-1 | 1.1337327 | 1.049240085 | 0.121897076 | 0.117 | 0.192 | 0.006157263 | 1 |
| WBGene00010505 | K02D3.2 | 0.671542107 | 0.586985677 | 0.121989142 | 0.057 | 0.085 | 0.109181773 | 1 |
| WBGene00219609 | linc-7 | 4.301924922 | 4.217023253 | 0.122487217 | 0.56 | 0.615 | 0.611116962 | 1 |
| WBGene00044026 | cisd-1 | 0.622226019 | 0.537200441 | 0.12266598 | 0.065 | 0.092 | 0.165302837 | 1 |
| WBGene00003034 | lin-49 | 0.804824732 | 0.719659871 | 0.122866923 | 0.078 | 0.117 | 0.062961274 | 1 |
| WBGene00012221 | W03C9.5 | 1.281098469 | 1.195669457 | 0.123248011 | 0.161 | 0.221 | 0.090606966 | 1 |
| WBGene00004783 | seu-1 | 0.627932252 | 0.542333993 | 0.123492184 | 0.058 | 0.083 | 0.174752954 | 1 |
| WBGene00010408 | mboa-2 | 0.798477078 | 0.712681767 | 0.12377647 | 0.08 | 0.117 | 0.083416103 | 1 |
| WBGene00044418 | C08G5.7 | 0.913100196 | 0.826949319 | 0.124289442 | 0.097 | 0.133 | 0.160953688 | 1 |
| WBGene00020682 | T22D1.3 | 0.511389229 | 0.425158015 | 0.124405345 | 0.042 | 0.065 | 0.127576352 | 1 |
| WBGene00002078 | xpo-1 | 0.932149819 | 0.845667392 | 0.124767768 | 0.085 | 0.135 | 0.024154581 | 1 |
| WBGene00016244 | C30B5.2 | 0.676229836 | 0.589734176 | 0.12478686 | 0.055 | 0.087 | 0.064826097 | 1 |
| WBGene00006632 | tsp-6 | 0.924892745 | 0.838350164 | 0.124854552 | 0.095 | 0.133 | 0.130545641 | 1 |
| WBGene00000262 | bra-1 | 0.745173675 | 0.658493088 | 0.125053653 | 0.062 | 0.092 | 0.0927905 | 1 |

| | | | | | | | | |
|---|---|---|---|---|---|---|---|---|
| WBGene00018160 | odr-8 | 0.87625019 | 0.789328201 | 0.125401923 | 0.095 | 0.135 | 0.105641731 | 1 |
| WBGene00004698 | rsp-1 | 1.257761937 | 1.170817262 | 0.125434651 | 0.133 | 0.215 | 0.004147747 | 1 |
| WBGene00010236 | F58B4.3 | 3.757413667 | 3.670362156 | 0.125588783 | 0.846 | 0.902 | 0.395741954 | 1 |
| WBGene00020100 | mks-1 | 1.072372481 | 0.985106679 | 0.125897939 | 0.102 | 0.183 | 0.001518427 | 1 |
| WBGene00006068 | sto-6 | 0.752596676 | 0.665016057 | 0.126352125 | 0.074 | 0.11 | 0.08966424 | 1 |
| WBGene00003649 | nhr-59 | 0.747210952 | 0.659420821 | 0.126654387 | 0.074 | 0.115 | 0.048354747 | 1 |
| WBGene00008613 | fbxb-69 | 1.099141283 | 1.010828897 | 0.127407841 | 0.134 | 0.171 | 0.300543158 | 1 |
| WBGene00012766 | Y41E3.8 | 0.64273893 | 0.553076542 | 0.129355482 | 0.067 | 0.085 | 0.409339275 | 1 |
| WBGene00003934 | pat-10 | 0.709786587 | 0.619928971 | 0.129637137 | 0.071 | 0.079 | 0.794208855 | 1 |
| WBGene00185118 | K02E2.11 | 1.087835829 | 0.997817588 | 0.12986887 | 0.122 | 0.16 | 0.263447132 | 1 |
| WBGene00004897 | snb-1 | 1.800973355 | 1.710591412 | 0.130393581 | 0.263 | 0.377 | 0.025907814 | 1 |
| WBGene00012779 | cdkl-1 | 1.138847158 | 1.048264297 | 0.130683445 | 0.118 | 0.179 | 0.023666763 | 1 |
| WBGene00000896 | dad-1 | 0.806306226 | 0.715674235 | 0.130754325 | 0.088 | 0.121 | 0.166837477 | 1 |
| WBGene00021469 | Y39G10AR.11 | 0.54947004 | 0.458442788 | 0.131324564 | 0.046 | 0.058 | 0.454024111 | 1 |
| WBGene00001081 | dpy-22 | 0.557448824 | 0.465455815 | 0.132717857 | 0.044 | 0.054 | 0.51267762 | 1 |
| WBGene00005719 | srv-8 | 1.023965604 | 0.931948663 | 0.132752384 | 0.108 | 0.183 | 0.003958018 | 1 |
| WBGene00003387 | mod-5 | 0.654729849 | 0.562467084 | 0.133107033 | 0.071 | 0.079 | 0.760800096 | 1 |
| WBGene00010882 | atg-7 | 0.548607237 | 0.456307291 | 0.133160675 | 0.055 | 0.071 | 0.350561474 | 1 |
| WBGene00015619 | C08G9.1 | 1.878030042 | 1.784959377 | 0.134272586 | 0.272 | 0.363 | 0.037703026 | 1 |
| WBGene00006803 | unc-70 | 0.808144185 | 0.715000952 | 0.134377281 | 0.081 | 0.11 | 0.213095654 | 1 |
| WBGene00010012 | saeg-1 | 1.128836026 | 1.035609908 | 0.134496858 | 0.097 | 0.162 | 0.007882581 | 1 |
| WBGene00021313 | paqr-2 | 0.52075313 | 0.427523294 | 0.134502222 | 0.046 | 0.05 | 0.837617562 | 1 |
| WBGene00017120 | tctn-1 | 0.809868607 | 0.716502651 | 0.134698602 | 0.072 | 0.108 | 0.084864633 | 1 |
| WBGene00017914 | F29A7.4 | 0.499100544 | 0.405462682 | 0.13509088 | 0.046 | 0.05 | 0.839663086 | 1 |
| WBGene00019492 | K07E3.4 | 0.735075244 | 0.641164892 | 0.135484 | 0.08 | 0.1 | 0.374597961 | 1 |
| WBGene00015901 | nhr-158 | 2.171176218 | 2.077248146 | 0.135509563 | 0.392 | 0.485 | 0.395658374 | 1 |
| WBGene00022503 | ZC13.2 | 0.589113906 | 0.494892133 | 0.135933285 | 0.06 | 0.058 | 0.785152591 | 1 |
| WBGene00007484 | C09G1.2 | 2.012519291 | 1.918161795 | 0.136129092 | 0.327 | 0.417 | 0.175001153 | 1 |
| WBGene00018476 | F45E12.6 | 4.091061981 | 3.996570744 | 0.136322039 | 0.887 | 0.892 | 0.315194939 | 1 |
| WBGene00006528 | tba-1 | 1.039098946 | 0.944287216 | 0.136784412 | 0.104 | 0.163 | 0.023540954 | 1 |
| WBGene00009126 | pyk-1 | 1.015614935 | 0.92013652 | 0.137746235 | 0.111 | 0.152 | 0.130338996 | 1 |
| WBGene00003953 | pbs-7 | 1.068831133 | 0.972456427 | 0.13903931 | 0.104 | 0.158 | 0.031313506 | 1 |
| WBGene00045388 | lfor-1 | 0.641511896 | 0.544556537 | 0.139877016 | 0.057 | 0.083 | 0.13740343 | 1 |
| WBGene00000490 | che-11 | 1.176776734 | 1.079701147 | 0.140050468 | 0.14 | 0.179 | 0.269075265 | 1 |
| WBGene00044143 | K09E9.4 | 0.707801636 | 0.610723383 | 0.140054314 | 0.065 | 0.094 | 0.135076008 | 1 |
| WBGene00012372 | W09H1.1 | 1.634453297 | 1.537360614 | 0.140075134 | 0.224 | 0.306 | 0.063944427 | 1 |
| WBGene00001030 | dnj-12 | 0.82379768 | 0.726691299 | 0.140094895 | 0.092 | 0.102 | 0.84532479 | 1 |
| WBGene00007534 | fubl-1 | 1.333202543 | 1.23562575 | 0.140773556 | 0.186 | 0.208 | 0.902273932 | 1 |
| WBGene00016289 | lntl-1 | 2.86953771 | 2.771316677 | 0.141702997 | 0.62 | 0.733 | 0.699167469 | 1 |
| WBGene00001684 | gpd-2 | 1.401081047 | 1.302349869 | 0.142438981 | 0.194 | 0.24 | 0.385062057 | 1 |
| WBGene00001464 | flp-21 | 2.186111154 | 2.087302175 | 0.142551224 | 0.311 | 0.41 | 0.063114515 | 1 |
| WBGene00009078 | rpb-12 | 0.483790421 | 0.384907752 | 0.142657536 | 0.034 | 0.052 | 0.163091225 | 1 |
| WBGene00010958 | ndfl-4 | 1.636212516 | 1.536381607 | 0.144025557 | 0.21 | 0.267 | 0.267298048 | 1 |
| WBGene00013094 | Y51H1A.3 | 0.782209649 | 0.680983963 | 0.146037796 | 0.078 | 0.087 | 0.751727596 | 1 |
| WBGene00016415 | ampd-1 | 0.604506871 | 0.503016329 | 0.146419901 | 0.057 | 0.073 | 0.355305947 | 1 |
| WBGene00249817 | F42G8.19 | 1.154535041 | 1.052910846 | 0.146612722 | 0.136 | 0.185 | 0.165645292 | 1 |
| WBGene00045483 | oxy-5 | 0.839109195 | 0.737253964 | 0.146946037 | 0.087 | 0.119 | 0.164302685 | 1 |
| WBGene00007653 | cysl-1 | 2.278313817 | 2.175609768 | 0.148170622 | 0.436 | 0.51 | 0.9262915 | 1 |
| WBGene00019767 | rpa-2 | 0.966686028 | 0.863695001 | 0.148584643 | 0.087 | 0.129 | 0.056264616 | 1 |
| WBGene00016111 | C25H3.1 | 1.396310388 | 1.293004299 | 0.149039183 | 0.189 | 0.252 | 0.177304515 | 1 |
| WBGene00003065 | lpd-9 | 0.539004272 | 0.435650438 | 0.149108065 | 0.041 | 0.054 | 0.354353685 | 1 |
| WBGene00000182 | arf-1.2 | 0.807271797 | 0.703656758 | 0.149484904 | 0.083 | 0.113 | 0.179800458 | 1 |
| WBGene00017616 | F20A1.4 | 2.126859507 | 2.023214312 | 0.14952841 | 0.341 | 0.488 | 0.017347926 | 1 |
| WBGene00018886 | npr-28 | 0.801036059 | 0.697299015 | 0.149660918 | 0.092 | 0.115 | 0.369439091 | 1 |
| WBGene00007877 | nfki-1 | 1.206464997 | 1.102289532 | 0.150293428 | 0.152 | 0.183 | 0.508858031 | 1 |
| WBGene00001336 | qars-1 | 1.365707536 | 1.261108542 | 0.150904451 | 0.171 | 0.26 | 0.017780972 | 1 |
| WBGene00006755 | unc-16 | 0.987470121 | 0.882740516 | 0.151092882 | 0.101 | 0.131 | 0.232462397 | 1 |
| WBGene00006773 | unc-37 | 0.445739414 | 0.340984918 | 0.151128793 | 0.035 | 0.056 | 0.13388889 | 1 |
| WBGene00001166 | eftu-2 | 0.570956617 | 0.466070763 | 0.151318301 | 0.049 | 0.065 | 0.324155286 | 1 |
| WBGene00010957 | nduo-6 | 3.923207759 | 3.818050652 | 0.151709637 | 0.848 | 0.844 | 0.033291651 | 1 |
| WBGene00000199 | arx-1 | 0.539764185 | 0.434179201 | 0.152326933 | 0.053 | 0.069 | 0.354531236 | 1 |
| WBGene00003961 | pct-1 | 1.001897873 | 0.896230489 | 0.152445812 | 0.12 | 0.152 | 0.313437422 | 1 |
| WBGene00008520 | F02D8.1 | 2.056834342 | 1.95032849 | 0.153655464 | 0.339 | 0.444 | 0.093064136 | 1 |
| WBGene00012929 | rsks-1 | 0.716130657 | 0.60920899 | 0.154255358 | 0.076 | 0.081 | 0.928594054 | 1 |

| | | | | | | | | |
|---|---|---|---|---|---|---|---|---|
| WBGene00017239 | F08B6.1 | 0.783207473 | 0.676187013 | 0.154397888 | 0.076 | 0.11 | 0.105032768 | 1 |
| WBGene00004429 | rpl-17 | 2.928010569 | 2.820574696 | 0.154997202 | 0.645 | 0.748 | 0.625615603 | 1 |
| WBGene00009483 | F36G3.2 | 0.574420735 | 0.466755959 | 0.15532744 | 0.053 | 0.071 | 0.29561235 | 1 |
| WBGene00021355 | Y37E3.17 | 0.709577375 | 0.60189451 | 0.155353534 | 0.072 | 0.096 | 0.25698263 | 1 |
| WBGene00000244 | bbs-8 | 1.820323812 | 1.71253221 | 0.155510409 | 0.274 | 0.385 | 0.042341793 | 1 |
| WBGene00016841 | magu-3 | 0.479526886 | 0.37094795 | 0.156646293 | 0.034 | 0.052 | 0.157473342 | 1 |
| WBGene00000793 | crh-1 | 1.845556241 | 1.736167408 | 0.157814727 | 0.311 | 0.387 | 0.555584197 | 1 |
| WBGene00017973 | ift-81 | 1.799677429 | 1.690212548 | 0.157924442 | 0.292 | 0.34 | 0.880663183 | 1 |
| WBGene00004378 | rme-8 | 0.448805598 | 0.339043762 | 0.158352856 | 0.035 | 0.052 | 0.219639458 | 1 |
| WBGene00003150 | mbk-2 | 1.799686029 | 1.689496542 | 0.158969827 | 0.288 | 0.379 | 0.116033657 | 1 |
| WBGene00006753 | unc-14 | 1.105529168 | 0.995133291 | 0.159267584 | 0.124 | 0.154 | 0.393616828 | 1 |
| WBGene00016158 | ari-1.1 | 0.583372033 | 0.472620249 | 0.15978105 | 0.06 | 0.075 | 0.438190272 | 1 |
| WBGene00011558 | ostf-4 | 0.602512754 | 0.491456915 | 0.160219709 | 0.053 | 0.069 | 0.340371757 | 1 |
| WBGene00021260 | Y22D7AR.6 | 0.810601872 | 0.699506964 | 0.160276072 | 0.078 | 0.11 | 0.129660981 | 1 |
| WBGene00008576 | F08G2.4 | 0.630003963 | 0.518885752 | 0.160309692 | 0.06 | 0.069 | 0.686011528 | 1 |
| WBGene00016964 | C56C10.7 | 0.501747928 | 0.3899187 | 0.161335473 | 0.041 | 0.067 | 0.077471765 | 1 |
| WBGene00014159 | arrd-4 | 1.806170361 | 1.693798167 | 0.162118807 | 0.277 | 0.385 | 0.071393775 | 1 |
| WBGene00003598 | nhl-2 | 0.690228409 | 0.577167232 | 0.163112799 | 0.065 | 0.096 | 0.117812051 | 1 |
| WBGene00022635 | ZC581.9 | 1.679061592 | 1.565967381 | 0.163160458 | 0.228 | 0.319 | 0.051782487 | 1 |
| WBGene00015913 | C17F4.7 | 1.175875152 | 1.062668129 | 0.16332321 | 0.099 | 0.085 | 0.315421719 | 1 |
| WBGene00004095 | pqe-1 | 0.503717951 | 0.390352704 | 0.163551479 | 0.051 | 0.054 | 0.948253067 | 1 |
| WBGene00013751 | Y113G7A.14 | 1.103676291 | 0.990116907 | 0.16383156 | 0.134 | 0.173 | 0.25552193 | 1 |
| WBGene00022678 | sar-1 | 0.651411164 | 0.537788486 | 0.163922874 | 0.048 | 0.09 | 0.009420953 | 1 |
| WBGene00017084 | E01A2.1 | 0.761810115 | 0.647757033 | 0.164543816 | 0.076 | 0.106 | 0.174835278 | 1 |
| WBGene00004917 | snr-4 | 0.998185997 | 0.884053308 | 0.164658665 | 0.129 | 0.16 | 0.404275007 | 1 |
| WBGene00015352 | C02F5.12 | 0.679096745 | 0.564902638 | 0.164747272 | 0.067 | 0.09 | 0.245545171 | 1 |
| WBGene00002191 | kin-3 | 0.742171719 | 0.627724794 | 0.165112012 | 0.076 | 0.085 | 0.762193626 | 1 |
| WBGene00022254 | nlp-71 | 2.743416465 | 2.628737917 | 0.165446174 | 0.498 | 0.544 | 0.958077034 | 1 |
| WBGene00018511 | F46F11.8 | 0.816460763 | 0.701213645 | 0.166266446 | 0.08 | 0.098 | 0.437487626 | 1 |
| WBGene00017380 | F11D5.1 | 0.734366493 | 0.618981788 | 0.166464942 | 0.069 | 0.09 | 0.290706683 | 1 |
| WBGene00022776 | perm-3 | 0.532792248 | 0.417352034 | 0.166545023 | 0.037 | 0.052 | 0.271490687 | 1 |
| WBGene00018204 | F39F10.4 | 0.937647937 | 0.821933996 | 0.166939929 | 0.087 | 0.123 | 0.091801498 | 1 |
| WBGene00022089 | Y69A2AR.18 | 1.038344732 | 0.922594383 | 0.166992454 | 0.106 | 0.142 | 0.155034522 | 1 |
| WBGene00269432 | ZC132.11 | 1.324452043 | 1.208682906 | 0.167019559 | 0.141 | 0.242 | 0.000799456 | 1 |
| WBGene00016378 | imp-3 | 0.536118577 | 0.420286482 | 0.167110389 | 0.053 | 0.071 | 0.307895082 | 1 |
| WBGene00009888 | F49E2.5 | 0.682576594 | 0.566194838 | 0.167903382 | 0.067 | 0.081 | 0.524572821 | 1 |
| WBGene00003653 | nhr-63 | 0.445414757 | 0.329025244 | 0.167914574 | 0.042 | 0.056 | 0.395044021 | 1 |
| WBGene00008555 | F07C6.4 | 0.917361421 | 0.800922578 | 0.167985742 | 0.092 | 0.123 | 0.220178541 | 1 |
| WBGene00016961 | vps-32.1 | 1.284722227 | 1.167937613 | 0.168484584 | 0.175 | 0.213 | 0.437328426 | 1 |
| WBGene00014021 | svop-1 | 0.545883008 | 0.428820989 | 0.169317603 | 0.053 | 0.065 | 0.491271254 | 1 |
| WBGene00045161 | lfor-2 | 0.919544567 | 0.801286665 | 0.170610087 | 0.097 | 0.102 | 0.976792214 | 1 |
| WBGene00014213 | ZK1073.1 | 1.30981056 | 1.191373358 | 0.170868764 | 0.163 | 0.225 | 0.090153654 | 1 |
| WBGene00007684 | C18E9.4 | 0.459189083 | 0.34074924 | 0.170872575 | 0.042 | 0.054 | 0.463957112 | 1 |
| WBGene00021921 | rbm-39 | 0.789519894 | 0.670831806 | 0.171230716 | 0.088 | 0.104 | 0.601028835 | 1 |
| WBGene00011511 | gpcp-2 | 0.664901377 | 0.545161926 | 0.172747513 | 0.065 | 0.085 | 0.321948664 | 1 |
| WBGene00003603 | nhr-4 | 0.909509866 | 0.78957977 | 0.173022555 | 0.097 | 0.123 | 0.311946018 | 1 |
| WBGene00014166 | ZK945.4 | 0.536907117 | 0.416210923 | 0.174127801 | 0.049 | 0.069 | 0.23634991 | 1 |
| WBGene00017699 | flcn-1 | 0.800982317 | 0.680236302 | 0.174199677 | 0.081 | 0.108 | 0.265617727 | 1 |
| WBGene00010560 | eif-2beta | 0.908085678 | 0.787326343 | 0.174218894 | 0.083 | 0.094 | 0.704120113 | 1 |
| WBGene00018968 | F56D2.8 | 0.596313058 | 0.475255974 | 0.174648454 | 0.044 | 0.071 | 0.079809627 | 1 |
| WBGene00010756 | K10G9.2 | 0.730826595 | 0.609669181 | 0.174793201 | 0.069 | 0.106 | 0.068637572 | 1 |
| WBGene00006513 | rimb-1 | 0.785257 | 0.663374019 | 0.175839971 | 0.078 | 0.092 | 0.572035099 | 1 |
| WBGene00008023 | C39B10.3 | 0.691759779 | 0.569740792 | 0.176036187 | 0.074 | 0.081 | 0.844699785 | 1 |
| WBGene00020196 | T03G11.3 | 1.643905451 | 1.521044748 | 0.177250527 | 0.251 | 0.34 | 0.115364496 | 1 |
| WBGene00021960 | tmem-258 | 0.929765052 | 0.806804369 | 0.177394767 | 0.095 | 0.125 | 0.251360885 | 1 |
| WBGene00010965 | ctc-2 | 1.981325763 | 1.857156856 | 0.179137867 | 0.323 | 0.398 | 0.475219712 | 1 |
| WBGene00004167 | pqn-87 | 0.951833836 | 0.827456237 | 0.179438945 | 0.092 | 0.125 | 0.183319282 | 1 |
| WBGene00012472 | Y17G7B.21 | 0.70594961 | 0.581487042 | 0.17956153 | 0.071 | 0.083 | 0.59222045 | 1 |
| WBGene00001127 | dyf-11 | 0.727009606 | 0.602485321 | 0.179650568 | 0.062 | 0.088 | 0.144269508 | 1 |
| WBGene00021936 | zipt-15 | 1.006187828 | 0.881070801 | 0.180505714 | 0.106 | 0.129 | 0.445662289 | 1 |
| WBGene00003225 | mev-1 | 0.688031306 | 0.562727963 | 0.180774511 | 0.06 | 0.062 | 0.98526022 | 1 |
| WBGene00012113 | T28B8.1 | 1.781248456 | 1.655529725 | 0.181373789 | 0.253 | 0.337 | 0.103933025 | 1 |
| WBGene00001122 | dyf-6 | 1.832341688 | 1.706425787 | 0.181658246 | 0.272 | 0.402 | 0.009921431 | 1 |
| WBGene00008745 | F13E6.1 | 0.820340602 | 0.693495937 | 0.18299817 | 0.09 | 0.108 | 0.532538413 | 1 |

| | | | | | | | | |
|---|---|---|---|---|---|---|---|---|
| WBGene00003636 | nhr-46 | 0.572123283 | 0.445190197 | 0.183125735 | 0.048 | 0.063 | 0.319996659 | 1 |
| WBGene00011517 | cgt-1 | 0.803542592 | 0.676305054 | 0.183564965 | 0.071 | 0.112 | 0.041943954 | 1 |
| WBGene00003148 | mbf-1 | 1.138536761 | 1.010801334 | 0.184283266 | 0.145 | 0.198 | 0.139178195 | 1 |
| WBGene00007883 | C33A12.4 | 5.051192191 | 4.923352967 | 0.184433015 | 0.981 | 0.985 | 0.004213755 | 1 |
| WBGene00006306 | sue-1 | 1.044816847 | 0.916521656 | 0.185090836 | 0.127 | 0.138 | 0.932045612 | 1 |
| WBGene00009727 | lips-1 | 0.847214673 | 0.718832551 | 0.185216251 | 0.055 | 0.067 | 0.432695791 | 1 |
| WBGene00003218 | mep-1 | 0.936219886 | 0.807785729 | 0.185291322 | 0.088 | 0.129 | 0.088859265 | 1 |
| WBGene00017121 | cyc-2.1 | 1.715958891 | 1.585764589 | 0.187830675 | 0.261 | 0.362 | 0.073769405 | 1 |
| WBGene00021474 | dot-1.1 | 0.438141731 | 0.306916245 | 0.189318359 | 0.037 | 0.052 | 0.29490226 | 1 |
| WBGene00077526 | C25A1.16 | 0.588160042 | 0.456812138 | 0.18949497 | 0.048 | 0.069 | 0.185521364 | 1 |
| WBGene00010961 | nduo-2 | 1.35492448 | 1.222859027 | 0.190530173 | 0.154 | 0.208 | 0.10670692 | 1 |
| WBGene00013272 | aqp-12 | 1.078228496 | 0.946128879 | 0.190579462 | 0.127 | 0.162 | 0.299700704 | 1 |
| WBGene00003637 | nhr-47 | 1.178564438 | 1.046328403 | 0.190776272 | 0.147 | 0.212 | 0.057704626 | 1 |
| WBGene00019128 | F59G1.4 | 0.855743261 | 0.723017199 | 0.191483232 | 0.076 | 0.094 | 0.401715144 | 1 |
| WBGene00008930 | F18A11.3 | 0.52595862 | 0.393022181 | 0.191786742 | 0.034 | 0.054 | 0.123546075 | 1 |
| WBGene00003885 | osm-5 | 1.29640232 | 1.162594744 | 0.193043526 | 0.161 | 0.221 | 0.101660822 | 1 |
| WBGene00003904 | pabp-2 | 0.581113262 | 0.447197363 | 0.193199803 | 0.044 | 0.069 | 0.105045816 | 1 |
| WBGene00018042 | mks-3 | 0.80515242 | 0.671168672 | 0.193297689 | 0.067 | 0.108 | 0.033819894 | 1 |
| WBGene00008254 | C51E3.9 | 0.454533374 | 0.320100372 | 0.193945826 | 0.039 | 0.052 | 0.368917394 | 1 |
| WBGene00022376 | Y94H6A.3 | 0.629697472 | 0.495130642 | 0.194138898 | 0.051 | 0.063 | 0.456778291 | 1 |
| WBGene00018491 | mdh-1 | 0.620441687 | 0.485652219 | 0.194460097 | 0.065 | 0.069 | 0.94912551 | 1 |
| WBGene00002264 | lec-1 | 0.982286903 | 0.847387625 | 0.194618519 | 0.118 | 0.125 | 0.892373358 | 1 |
| WBGene00020929 | W02C12.2 | 0.501493391 | 0.366498617 | 0.194756291 | 0.039 | 0.054 | 0.289618307 | 1 |
| WBGene00006699 | uba-1 | 0.783737379 | 0.648672372 | 0.194857616 | 0.074 | 0.098 | 0.263750645 | 1 |
| WBGene00021926 | Y55F3AM.9 | 0.512053391 | 0.376666322 | 0.195322253 | 0.053 | 0.063 | 0.59220109 | 1 |
| WBGene00020151 | T01G6.1 | 0.60063406 | 0.463536356 | 0.197790177 | 0.044 | 0.065 | 0.160764928 | 1 |
| WBGene00020481 | T13C2.6 | 0.565530657 | 0.428200044 | 0.198126194 | 0.055 | 0.056 | 0.943844082 | 1 |
| WBGene00045056 | F13H10.9 | 0.95848793 | 0.820453028 | 0.199142268 | 0.097 | 0.137 | 0.117024883 | 1 |
| WBGene00009580 | xbx-6 | 1.833236006 | 1.69478316 | 0.199745234 | 0.269 | 0.346 | 0.231892595 | 1 |
| WBGene00007495 | C09G9.5 | 1.003581706 | 0.864621297 | 0.200477494 | 0.099 | 0.137 | 0.136653057 | 1 |
| WBGene00004920 | snr-7 | 1.224358592 | 1.085324257 | 0.200584145 | 0.166 | 0.183 | 0.985499677 | 1 |
| WBGene00016316 | C32D5.8 | 1.708816532 | 1.569038396 | 0.201657223 | 0.269 | 0.331 | 0.555700717 | 1 |
| WBGene00009004 | pfd-6 | 0.681730557 | 0.541366927 | 0.202501912 | 0.067 | 0.088 | 0.309225454 | 1 |
| WBGene00011288 | R102.1 | 4.758081678 | 4.617188558 | 0.203265805 | 0.747 | 0.779 | 0.209652346 | 1 |
| WBGene00010500 | K02B12.7 | 0.582797306 | 0.441789174 | 0.203431733 | 0.053 | 0.071 | 0.297654326 | 1 |
| WBGene00022812 | uvs-1 | 0.46806962 | 0.326454905 | 0.204306847 | 0.035 | 0.05 | 0.280735047 | 1 |
| WBGene00003025 | lin-40 | 1.034474247 | 0.892685954 | 0.204557267 | 0.124 | 0.129 | 0.833916341 | 1 |
| WBGene00001237 | elc-2 | 3.594953944 | 3.45228345 | 0.205830014 | 0.814 | 0.831 | 0.059079712 | 1 |
| WBGene00020447 | mob-3 | 0.729593326 | 0.586362139 | 0.206638923 | 0.067 | 0.092 | 0.203203266 | 1 |
| WBGene00020052 | R13A5.10 | 1.021764926 | 0.878017237 | 0.207384079 | 0.102 | 0.163 | 0.016984777 | 1 |
| WBGene00012674 | bed-1 | 0.470647755 | 0.32676003 | 0.207586108 | 0.037 | 0.052 | 0.289916588 | 1 |
| WBGene00017198 | nhr-36 | 0.723802614 | 0.579421391 | 0.208298074 | 0.062 | 0.079 | 0.36352582 | 1 |
| WBGene00022527 | ZC132.9 | 1.008420575 | 0.863805736 | 0.208635111 | 0.111 | 0.148 | 0.177302121 | 1 |
| WBGene00011772 | T14G8.4 | 1.30428784 | 1.159581673 | 0.208766869 | 0.161 | 0.194 | 0.477733645 | 1 |
| WBGene00003129 | map-1 | 0.735690669 | 0.590671208 | 0.209218858 | 0.065 | 0.079 | 0.507150444 | 1 |
| WBGene00023491 | tsp-21 | 1.851013195 | 1.705413182 | 0.210056416 | 0.314 | 0.381 | 0.559475424 | 1 |
| WBGene00021187 | Y9D1A.2 | 2.142878573 | 1.996268743 | 0.211513275 | 0.336 | 0.402 | 0.890645211 | 1 |
| WBGene00017511 | srbc-41 | 2.667781991 | 2.521054611 | 0.211682863 | 0.493 | 0.635 | 0.182015653 | 1 |
| WBGene00000066 | act-4 | 2.247397655 | 2.100527716 | 0.211888534 | 0.428 | 0.5 | 0.581629438 | 1 |
| WBGene00003406 | mrg-1 | 0.619600199 | 0.472187025 | 0.212672256 | 0.057 | 0.065 | 0.661716437 | 1 |
| WBGene00006720 | ubc-25 | 0.671000454 | 0.52348714 | 0.212816726 | 0.046 | 0.065 | 0.20465691 | 1 |
| WBGene00020444 | T12B3.1 | 0.56267195 | 0.414940534 | 0.213131383 | 0.049 | 0.067 | 0.281144529 | 1 |
| WBGene00010908 | imph-1 | 0.61239836 | 0.464520131 | 0.213343187 | 0.039 | 0.056 | 0.230502878 | 1 |
| WBGene00007882 | C33A12.3 | 1.544977993 | 1.396870559 | 0.213673861 | 0.233 | 0.279 | 0.670972086 | 1 |
| WBGene00004732 | sax-7 | 0.605389808 | 0.457249041 | 0.21372195 | 0.053 | 0.065 | 0.480284245 | 1 |
| WBGene00009717 | dep-1 | 1.005834451 | 0.8576581 | 0.213773287 | 0.097 | 0.144 | 0.055472556 | 1 |
| WBGene00006066 | sto-4 | 0.694124141 | 0.545946087 | 0.213775743 | 0.055 | 0.062 | 0.746032445 | 1 |
| WBGene00013734 | Y111B2A.10 | 0.864196478 | 0.715600747 | 0.214378324 | 0.09 | 0.119 | 0.241890437 | 1 |
| WBGene00006523 | tam-1 | 0.864202949 | 0.715012789 | 0.215235905 | 0.102 | 0.117 | 0.680342989 | 1 |
| WBGene00007009 | wwp-1 | 0.694707332 | 0.545152245 | 0.215762382 | 0.069 | 0.077 | 0.763676082 | 1 |
| WBGene00015755 | C14B9.10 | 0.96162069 | 0.811723158 | 0.216256425 | 0.101 | 0.142 | 0.115628789 | 1 |
| WBGene00016305 | fbxc-57 | 1.135504842 | 0.985434323 | 0.216505993 | 0.148 | 0.194 | 0.249160169 | 1 |
| WBGene00012132 | ifta-2 | 1.652549868 | 1.502100089 | 0.217053151 | 0.221 | 0.287 | 0.137714267 | 1 |
| WBGene00008443 | pde-3 | 0.611167424 | 0.460580621 | 0.217250834 | 0.058 | 0.065 | 0.740717006 | 1 |

| | | | | | | | | |
|---|---|---|---|---|---|---|---|---|
| WBGene00007645 | C17E4.6 | 0.766655302 | 0.615814632 | 0.217617087 | 0.081 | 0.096 | 0.579436089 | 1 |
| WBGene00012468 | Y17G7B.17 | 1.171382656 | 1.020048235 | 0.218329419 | 0.127 | 0.217 | 0.002040444 | 1 |
| WBGene00019983 | sti-1 | 0.938703173 | 0.787129219 | 0.218674992 | 0.097 | 0.133 | 0.173510178 | 1 |
| WBGene00003014 | lin-28 | 0.527433031 | 0.375847346 | 0.218691916 | 0.051 | 0.054 | 0.9722146 | 1 |
| WBGene00018145 | F37C4.5 | 1.525676311 | 1.373989998 | 0.218837091 | 0.207 | 0.29 | 0.072540229 | 1 |
| WBGene00023417 | swm-1 | 1.558049281 | 1.406314456 | 0.218907079 | 0.159 | 0.223 | 0.054291439 | 1 |
| WBGene00001488 | frm-1 | 1.041399645 | 0.88931278 | 0.219414966 | 0.118 | 0.133 | 0.855935061 | 1 |
| WBGene00006750 | unc-10 | 0.529122217 | 0.377005362 | 0.219458233 | 0.046 | 0.05 | 0.851137659 | 1 |
| WBGene00007476 | C09E9.1 | 1.017544649 | 0.865346971 | 0.219574835 | 0.12 | 0.152 | 0.34823904 | 1 |
| WBGene00003827 | ntl-4 | 0.840715996 | 0.688147942 | 0.220109175 | 0.058 | 0.102 | 0.015633672 | 1 |
| WBGene00006139 | str-79 | 1.745263998 | 1.592545352 | 0.220326433 | 0.283 | 0.354 | 0.580327512 | 1 |
| WBGene00019301 | K02D10.1 | 0.725287737 | 0.572267139 | 0.220762058 | 0.071 | 0.075 | 0.93845861 | 1 |
| WBGene00004930 | sod-1 | 0.891872226 | 0.738791346 | 0.220849026 | 0.088 | 0.115 | 0.255348162 | 1 |
| WBGene00016140 | rpb-2 | 0.600570824 | 0.447383739 | 0.221002249 | 0.041 | 0.062 | 0.151983915 | 1 |
| WBGene00010724 | K09E9.1 | 0.698684263 | 0.545154467 | 0.221496676 | 0.065 | 0.079 | 0.511176051 | 1 |
| WBGene00016005 | ift-74 | 1.252019511 | 1.097604576 | 0.222773661 | 0.159 | 0.187 | 0.601571638 | 1 |
| WBGene00015517 | C06A8.8 | 0.775787863 | 0.621142102 | 0.223106672 | 0.08 | 0.083 | 0.970616992 | 1 |
| WBGene00022062 | Y67D8B.2 | 0.732160447 | 0.576863761 | 0.22404576 | 0.062 | 0.09 | 0.123359388 | 1 |
| WBGene00004918 | snr-5 | 0.724264652 | 0.568571119 | 0.224618288 | 0.051 | 0.085 | 0.045399592 | 1 |
| WBGene00019465 | acl-14 | 0.927310734 | 0.771267865 | 0.225122273 | 0.099 | 0.117 | 0.553262074 | 1 |
| WBGene00008206 | set-6 | 0.587428517 | 0.431125589 | 0.225497459 | 0.06 | 0.056 | 0.658350731 | 1 |
| WBGene00000510 | cka-2 | 0.929794097 | 0.773413768 | 0.225609126 | 0.101 | 0.125 | 0.371839976 | 1 |
| WBGene00000492 | che-13 | 0.7419451 | 0.585313403 | 0.225971773 | 0.064 | 0.09 | 0.156593745 | 1 |
| WBGene00001173 | egl-4 | 0.555440402 | 0.398697511 | 0.226132192 | 0.046 | 0.05 | 0.847445842 | 1 |
| WBGene00016458 | C35E7.6 | 0.710157247 | 0.55340376 | 0.226147479 | 0.076 | 0.102 | 0.262872637 | 1 |
| WBGene00017565 | ddo-2 | 0.840368222 | 0.683454737 | 0.226378307 | 0.076 | 0.096 | 0.346712212 | 1 |
| WBGene00009534 | F38B7.3 | 2.881238569 | 2.724280556 | 0.226442547 | 0.631 | 0.69 | 0.267073809 | 1 |
| WBGene00017982 | hpo-18 | 0.841671615 | 0.684615584 | 0.226583957 | 0.087 | 0.115 | 0.236746125 | 1 |
| WBGene00000293 | cap-2 | 0.696150594 | 0.538139166 | 0.227962304 | 0.076 | 0.083 | 0.858350972 | 1 |
| WBGene00003080 | lsm-6 | 0.538576902 | 0.380520377 | 0.228027365 | 0.055 | 0.058 | 0.970381156 | 1 |
| WBGene00018756 | plpp-1.1 | 0.603695145 | 0.445158892 | 0.228719465 | 0.06 | 0.056 | 0.662333391 | 1 |
| WBGene00003619 | nhr-20 | 1.580271228 | 1.421619385 | 0.228886227 | 0.246 | 0.304 | 0.507560027 | 1 |
| WBGene00018067 | npr-7 | 0.834010771 | 0.675217628 | 0.22909008 | 0.046 | 0.1 | 0.001285164 | 1 |
| WBGene00007104 | B0024.15 | 0.580597234 | 0.42179972 | 0.229096386 | 0.049 | 0.054 | 0.838809757 | 1 |
| WBGene00011279 | asd-1 | 1.22000979 | 1.060640718 | 0.22992097 | 0.154 | 0.187 | 0.460266542 | 1 |
| WBGene00007184 | ctr-9 | 0.738829531 | 0.578829418 | 0.230831369 | 0.06 | 0.073 | 0.507445742 | 1 |
| WBGene00015955 | C18B2.4 | 0.923342185 | 0.762500336 | 0.232045739 | 0.097 | 0.131 | 0.207693852 | 1 |
| WBGene00007574 | C14B1.3 | 1.464495977 | 1.302705658 | 0.233414091 | 0.2 | 0.271 | 0.145187276 | 1 |
| WBGene00021416 | mksr-2 | 1.367117625 | 1.204416288 | 0.234728413 | 0.196 | 0.227 | 0.86020361 | 1 |
| WBGene00021693 | Y48G8AR.2 | 0.652914492 | 0.489761468 | 0.235380058 | 0.053 | 0.071 | 0.281374125 | 1 |
| WBGene00007350 | sucl-1 | 0.737681112 | 0.574341006 | 0.235649961 | 0.072 | 0.073 | 0.869319766 | 1 |
| WBGene00019079 | rpa-4 | 0.745470925 | 0.581346203 | 0.236781923 | 0.071 | 0.09 | 0.359110527 | 1 |
| WBGene00000188 | arl-3 | 2.429018021 | 2.264688363 | 0.237077583 | 0.47 | 0.579 | 0.823660175 | 1 |
| WBGene00003045 | lir-2 | 0.548150438 | 0.383415448 | 0.237662353 | 0.044 | 0.06 | 0.316138478 | 1 |
| WBGene00021427 | sec-61.B | 0.703051959 | 0.537900097 | 0.238263772 | 0.057 | 0.083 | 0.144721381 | 1 |
| WBGene00003962 | pdi-1 | 0.535916622 | 0.370585492 | 0.238522402 | 0.049 | 0.058 | 0.667409462 | 1 |
| WBGene00044176 | C30G7.2 | 1.725250615 | 1.559574092 | 0.239020699 | 0.279 | 0.285 | 0.373076665 | 1 |
| WBGene00000058 | acr-19 | 0.668242489 | 0.502187487 | 0.239566729 | 0.055 | 0.087 | 0.073073171 | 1 |
| WBGene00004106 | pqn-15 | 1.143875267 | 0.977772307 | 0.239635917 | 0.115 | 0.167 | 0.060986443 | 1 |
| WBGene00017615 | srsx-32 | 1.844272404 | 1.677938871 | 0.239968563 | 0.293 | 0.338 | 0.979365841 | 1 |
| WBGene00014233 | ZK1128.7 | 0.479814942 | 0.313286698 | 0.240249473 | 0.046 | 0.052 | 0.76568245 | 1 |
| WBGene00016938 | C54H2.4 | 0.658203316 | 0.490932959 | 0.241320114 | 0.051 | 0.069 | 0.267164241 | 1 |
| WBGene00001172 | egl-3 | 1.852429293 | 1.685010609 | 0.241534106 | 0.267 | 0.387 | 0.034990514 | 1 |
| WBGene00013604 | Y87G2A.18 | 0.521468661 | 0.353493208 | 0.242337353 | 0.051 | 0.056 | 0.870099684 | 1 |
| WBGene00007246 | nspb-10 | 0.818535245 | 0.650170326 | 0.242899234 | 0.078 | 0.085 | 0.877538097 | 1 |
| WBGene00018349 | F42C5.9 | 0.748860045 | 0.577872728 | 0.246682555 | 0.06 | 0.075 | 0.414454884 | 1 |
| WBGene00015367 | srt-64 | 2.001413759 | 1.829828558 | 0.247545119 | 0.313 | 0.396 | 0.350115213 | 1 |
| WBGene00002101 | ins-18 | 2.67941008 | 2.506747948 | 0.249098802 | 0.465 | 0.525 | 0.98964647 | 1 |
| WBGene00000508 | cit-1.2 | 0.563339634 | 0.390631713 | 0.24916486 | 0.048 | 0.058 | 0.557127653 | 1 |
| WBGene00219313 | C17E4.20 | 0.590689252 | 0.416118971 | 0.251851679 | 0.051 | 0.067 | 0.356287167 | 1 |
| WBGene00016633 | C44B11.1 | 0.507641581 | 0.332566382 | 0.252580122 | 0.053 | 0.046 | 0.511272728 | 1 |
| WBGene00010864 | xbx-3 | 3.360879633 | 3.185735211 | 0.25267999 | 0.781 | 0.815 | 0.071779846 | 1 |
| WBGene00008586 | snrp-40.1 | 0.689890387 | 0.514715176 | 0.252724408 | 0.064 | 0.071 | 0.778121289 | 1 |
| WBGene00000192 | arl-8 | 0.694644192 | 0.519201439 | 0.253110389 | 0.071 | 0.067 | 0.692835156 | 1 |

| | | | | | | | | |
|---|---|---|---|---|---|---|---|---|
| WBGene00001085 | dpy-26 | 0.722311658 | 0.546382197 | 0.25381256 | 0.053 | 0.079 | 0.124283784 | 1 |
| WBGene00004357 | rho-1 | 1.238104953 | 1.061973263 | 0.254104316 | 0.152 | 0.181 | 0.513848629 | 1 |
| WBGene00019069 | lgc-30 | 0.640811606 | 0.464396947 | 0.254512553 | 0.057 | 0.075 | 0.315971646 | 1 |
| WBGene00016017 | C23G10.11 | 0.712481306 | 0.536055118 | 0.254529186 | 0.057 | 0.062 | 0.824071273 | 1 |
| WBGene00006331 | sup-26 | 0.905106423 | 0.728457896 | 0.254849954 | 0.071 | 0.075 | 0.906864934 | 1 |
| WBGene00001137 | eat-6 | 1.381483829 | 1.20474825 | 0.254975543 | 0.17 | 0.223 | 0.143220277 | 1 |
| WBGene00017156 | F01E11.3 | 0.632060442 | 0.455248742 | 0.255085362 | 0.053 | 0.062 | 0.649508767 | 1 |
| WBGene00003040 | lin-59 | 0.848100021 | 0.670867174 | 0.255692949 | 0.095 | 0.098 | 0.876037667 | 1 |
| WBGene00004490 | rps-21 | 2.712318405 | 2.534648445 | 0.25632357 | 0.602 | 0.658 | 0.112790788 | 1 |
| WBGene00018164 | sup-36 | 0.917477887 | 0.739597831 | 0.256626675 | 0.09 | 0.131 | 0.090424546 | 1 |
| WBGene00007933 | C34E7.3 | 0.961894986 | 0.783589624 | 0.257240261 | 0.106 | 0.125 | 0.575556949 | 1 |
| WBGene00007247 | C01G12.7 | 2.933850605 | 2.755363299 | 0.257502751 | 0.622 | 0.687 | 0.413849162 | 1 |
| WBGene00007703 | gbf-1 | 0.558561804 | 0.379992059 | 0.257621685 | 0.048 | 0.063 | 0.347507913 | 1 |
| WBGene00004367 | ric-8 | 0.713098253 | 0.533995857 | 0.258390138 | 0.057 | 0.069 | 0.498072278 | 1 |
| WBGene00014079 | ZK792.7 | 0.761675634 | 0.582067094 | 0.259120349 | 0.09 | 0.087 | 0.606777726 | 1 |
| WBGene00016119 | C25H3.10 | 2.929450328 | 2.749455973 | 0.259676964 | 0.595 | 0.679 | 0.95875113 | 1 |
| WBGene00010966 | nduo-3 | 0.820469729 | 0.639639498 | 0.260882877 | 0.08 | 0.096 | 0.518296019 | 1 |
| WBGene00015418 | pac-1 | 1.319023702 | 1.137854476 | 0.261371945 | 0.175 | 0.192 | 0.93741898 | 1 |
| WBGene00006434 | prdx-2 | 1.593630902 | 1.411902005 | 0.262179378 | 0.228 | 0.277 | 0.662220043 | 1 |
| WBGene00010964 | ctc-1 | 2.726007332 | 2.544191763 | 0.262304419 | 0.557 | 0.581 | 0.079714339 | 1 |
| WBGene00016126 | nhr-258 | 0.775951078 | 0.593899079 | 0.262645517 | 0.078 | 0.104 | 0.260351436 | 1 |
| WBGene00017811 | F26A1.13 | 0.767201277 | 0.584912075 | 0.262987727 | 0.06 | 0.094 | 0.060693065 | 1 |
| WBGene00017792 | F25E5.16 | 0.739155569 | 0.556748495 | 0.263157781 | 0.069 | 0.088 | 0.348707532 | 1 |
| WBGene00007752 | wdfy-3 | 0.651037609 | 0.468184534 | 0.263801224 | 0.046 | 0.077 | 0.050716949 | 1 |
| WBGene00001558 | gdi-1 | 1.326476409 | 1.143048313 | 0.264630805 | 0.136 | 0.198 | 0.040356236 | 1 |
| WBGene00001145 | eat-16 | 1.357345004 | 1.173480292 | 0.265260708 | 0.154 | 0.221 | 0.048700998 | 1 |
| WBGene00044650 | Y71G10AR.4 | 1.17308735 | 0.987828576 | 0.267271915 | 0.138 | 0.165 | 0.51724902 | 1 |
| WBGene00008957 | glb-13 | 1.529470019 | 1.344161287 | 0.267343988 | 0.205 | 0.265 | 0.268217393 | 1 |
| WBGene00014772 | F56D5.4 | 1.227549911 | 1.041943432 | 0.267773547 | 0.154 | 0.173 | 0.902568719 | 1 |
| WBGene00004836 | sls-2.4 | 1.243980739 | 1.057991505 | 0.268325746 | 0.127 | 0.158 | 0.330464836 | 1 |
| WBGene00004739 | scd-1 | 0.634607052 | 0.448563626 | 0.268403928 | 0.053 | 0.056 | 0.945555951 | 1 |
| WBGene00008265 | kcnl-4 | 0.565002759 | 0.378486006 | 0.269086794 | 0.067 | 0.05 | 0.186244813 | 1 |
| WBGene00000157 | aps-2 | 0.503124235 | 0.316277228 | 0.269563249 | 0.053 | 0.048 | 0.622086006 | 1 |
| WBGene00015861 | C16D9.6 | 1.522278164 | 1.334192766 | 0.271349871 | 0.219 | 0.229 | 0.586491395 | 1 |
| WBGene00019900 | vdac-1 | 1.754034633 | 1.565752259 | 0.271634048 | 0.277 | 0.34 | 0.639441561 | 1 |
| WBGene00016634 | C44B11.4 | 2.757846224 | 2.568490546 | 0.273182498 | 0.599 | 0.617 | 0.029171321 | 1 |
| WBGene00011500 | T05G5.5 | 1.927920251 | 1.737565732 | 0.274623521 | 0.332 | 0.388 | 0.919932293 | 1 |
| WBGene00003707 | nhr-117 | 0.675401862 | 0.484914159 | 0.274815665 | 0.062 | 0.077 | 0.450518109 | 1 |
| WBGene00006651 | tts-2 | 2.22764856 | 2.036618415 | 0.275598243 | 0.417 | 0.417 | 0.096427185 | 1 |
| WBGene00014569 | bkip-1 | 0.843011192 | 0.651395324 | 0.276443263 | 0.081 | 0.108 | 0.251191161 | 1 |
| WBGene00004108 | pqn-18 | 1.035868849 | 0.843114117 | 0.278086295 | 0.122 | 0.14 | 0.692316963 | 1 |
| WBGene00004405 | rop-1 | 0.765566828 | 0.572788661 | 0.278120106 | 0.071 | 0.071 | 0.844542496 | 1 |
| WBGene00021811 | ral-1 | 0.771750638 | 0.577360276 | 0.28044601 | 0.08 | 0.065 | 0.293353225 | 1 |
| WBGene00003609 | nhr-10 | 0.760059082 | 0.564466296 | 0.282180742 | 0.067 | 0.085 | 0.415123731 | 1 |
| WBGene00003892 | osm-12 | 1.208441956 | 1.012165115 | 0.283167624 | 0.133 | 0.198 | 0.040354243 | 1 |
| WBGene00020391 | cct-7 | 0.932717142 | 0.736133496 | 0.283610251 | 0.094 | 0.119 | 0.343165142 | 1 |
| WBGene00016653 | ssb-1 | 0.587882378 | 0.391293515 | 0.283617778 | 0.058 | 0.046 | 0.300246288 | 1 |
| WBGene00003175 | mec-12 | 0.607573466 | 0.408721832 | 0.286882266 | 0.051 | 0.05 | 0.849211692 | 1 |
| WBGene00194866 | Y41C4A.22 | 1.09958899 | 0.900234602 | 0.287607587 | 0.124 | 0.123 | 0.659544856 | 1 |
| WBGene00003081 | lsm-7 | 0.522819491 | 0.32312923 | 0.288092149 | 0.048 | 0.052 | 0.861827828 | 1 |
| WBGene00012907 | cpt-1 | 1.29501272 | 1.094993953 | 0.288566083 | 0.155 | 0.208 | 0.164519204 | 1 |
| WBGene00003618 | nhr-19 | 0.816278144 | 0.616244785 | 0.288587135 | 0.08 | 0.087 | 0.86761305 | 1 |
| WBGene00021345 | Y37B11A.3 | 0.514932412 | 0.314671527 | 0.288915386 | 0.044 | 0.052 | 0.652194954 | 1 |
| WBGene00016115 | mdt-26 | 2.495052272 | 2.294306102 | 0.289615504 | 0.466 | 0.527 | 0.568052378 | 1 |
| WBGene00000962 | dhc-1 | 0.7362444 | 0.534716577 | 0.290743192 | 0.071 | 0.071 | 0.875999785 | 1 |
| WBGene00043992 | bbs-4 | 1.140178797 | 0.938380772 | 0.291133009 | 0.136 | 0.152 | 0.867850579 | 1 |
| WBGene00004459 | rpn-2 | 0.540356691 | 0.338517271 | 0.291192729 | 0.053 | 0.05 | 0.705974822 | 1 |
| WBGene00001505 | fut-1 | 0.569084045 | 0.366157686 | 0.292760852 | 0.048 | 0.052 | 0.848513349 | 1 |
| WBGene00008690 | F11C1.1 | 0.558219013 | 0.354306096 | 0.294184154 | 0.048 | 0.054 | 0.758376022 | 1 |
| WBGene00000996 | dif-1 | 1.18429981 | 0.980038823 | 0.294686314 | 0.154 | 0.165 | 0.984597561 | 1 |
| WBGene00013020 | Y48G10A.3 | 0.691028315 | 0.485904548 | 0.295931042 | 0.067 | 0.065 | 0.789665737 | 1 |
| WBGene00004743 | scm-1 | 0.748756339 | 0.543533266 | 0.29607431 | 0.076 | 0.087 | 0.710633851 | 1 |
| WBGene00021349 | arl-13 | 1.034849003 | 0.829554909 | 0.296176772 | 0.115 | 0.152 | 0.232427711 | 1 |
| WBGene00010318 | F59B10.2 | 0.747346885 | 0.542036352 | 0.296200488 | 0.065 | 0.094 | 0.150221259 | 1 |

| | | | | | | | | |
|---|---|---|---|---|---|---|---|---|
| WBGene00018608 | F48E3.9 | 0.937848539 | 0.731851151 | 0.29719141 | 0.081 | 0.121 | 0.071378896 | 1 |
| WBGene00009881 | F49C12.12 | 0.695569018 | 0.488885944 | 0.298180646 | 0.049 | 0.077 | 0.096053048 | 1 |
| WBGene00000829 | ctb-1 | 1.977867112 | 1.771058773 | 0.298361366 | 0.29 | 0.365 | 0.46525331 | 1 |
| WBGene00017934 | F30B5.4 | 0.554382751 | 0.347565367 | 0.298374414 | 0.051 | 0.052 | 0.914677168 | 1 |
| WBGene00010579 | K05C4.2 | 0.549365917 | 0.342267858 | 0.298779344 | 0.039 | 0.05 | 0.437556693 | 1 |
| WBGene00004220 | ptr-5 | 0.967467723 | 0.759490268 | 0.300048044 | 0.104 | 0.123 | 0.620676629 | 1 |
| WBGene00023068 | rpl-41.2 | 3.860522861 | 3.652445305 | 0.300192459 | 0.853 | 0.877 | 5.22E-05 | 0.333307555 |
| WBGene00007191 | lgc-20 | 1.728355651 | 1.519943321 | 0.300675436 | 0.281 | 0.323 | 0.801128254 | 1 |
| WBGene00019995 | R10F2.4 | 2.346663712 | 2.138162 | 0.300804386 | 0.428 | 0.485 | 0.49031884 | 1 |
| WBGene00007180 | B0457.6 | 1.908923511 | 1.700307842 | 0.300968791 | 0.316 | 0.348 | 0.613726225 | 1 |
| WBGene00006476 | rhgf-2 | 0.894958544 | 0.685776074 | 0.301786511 | 0.072 | 0.106 | 0.107716464 | 1 |
| WBGene00015547 | ain-1 | 0.754160906 | 0.544774455 | 0.302080795 | 0.053 | 0.073 | 0.228588098 | 1 |
| WBGene00003404 | mpz-1 | 1.036636044 | 0.827093018 | 0.302306684 | 0.108 | 0.131 | 0.441687052 | 1 |
| WBGene00022791 | ZK686.1 | 0.647068797 | 0.437365936 | 0.302537278 | 0.053 | 0.067 | 0.421733132 | 1 |
| WBGene00009164 | hrdl-1 | 0.974024505 | 0.764047206 | 0.302933208 | 0.102 | 0.112 | 0.942691797 | 1 |
| WBGene00019032 | srsx-17 | 1.045842943 | 0.835442187 | 0.303544128 | 0.11 | 0.137 | 0.357652101 | 1 |
| WBGene00015390 | C03F11.4 | 0.86218369 | 0.649504795 | 0.306830787 | 0.08 | 0.094 | 0.530624191 | 1 |
| WBGene00018327 | F42A6.5 | 0.679514637 | 0.466785869 | 0.306902739 | 0.058 | 0.056 | 0.740902437 | 1 |
| WBGene00003153 | mca-3 | 1.288184345 | 1.074032781 | 0.3089554 | 0.127 | 0.175 | 0.102442171 | 1 |
| WBGene00008454 | E02A10.4 | 0.694640525 | 0.479923052 | 0.309771833 | 0.065 | 0.077 | 0.620520454 | 1 |
| WBGene00007346 | frpr-2 | 0.615845851 | 0.400256502 | 0.311029684 | 0.062 | 0.054 | 0.498917322 | 1 |
| WBGene00008446 | E01G4.3 | 1.754874082 | 1.539233171 | 0.311104073 | 0.228 | 0.269 | 0.712944375 | 1 |
| WBGene00004245 | puf-9 | 0.744298888 | 0.528292766 | 0.31163096 | 0.057 | 0.085 | 0.114511753 | 1 |
| WBGene00004980 | spk-1 | 0.622295466 | 0.40580077 | 0.312335825 | 0.048 | 0.067 | 0.234544744 | 1 |
| WBGene00020762 | zer-1 | 0.811971148 | 0.594146468 | 0.314254586 | 0.065 | 0.094 | 0.137305 | 1 |
| WBGene00000509 | cka-1 | 0.871188388 | 0.653252819 | 0.314414564 | 0.092 | 0.108 | 0.622899234 | 1 |
| WBGene00022500 | lfi-1 | 1.351180719 | 1.133131501 | 0.314578526 | 0.2 | 0.181 | 0.132837694 | 1 |
| WBGene00000170 | aqp-2 | 1.731912584 | 1.512961255 | 0.315879997 | 0.269 | 0.31 | 0.804962607 | 1 |
| WBGene00008208 | nhr-167 | 1.876193499 | 1.65614126 | 0.317468273 | 0.306 | 0.362 | 0.933060149 | 1 |
| WBGene00001604 | gln-3 | 2.144463404 | 1.924214589 | 0.317751874 | 0.376 | 0.452 | 0.849557012 | 1 |
| WBGene00003602 | nhr-3 | 0.821965542 | 0.600240058 | 0.319882256 | 0.095 | 0.081 | 0.281941962 | 1 |
| WBGene00044004 | C24A8.6 | 1.307280922 | 1.085120065 | 0.320510367 | 0.182 | 0.225 | 0.458401247 | 1 |
| WBGene00022599 | daf-41 | 2.072875059 | 1.850347871 | 0.321038871 | 0.35 | 0.452 | 0.339714016 | 1 |
| WBGene00010732 | K10C3.5 | 0.603798835 | 0.380394775 | 0.322303929 | 0.057 | 0.058 | 0.930719724 | 1 |
| WBGene00009216 | F28D1.6 | 2.856260758 | 2.632692549 | 0.322540747 | 0.574 | 0.65 | 0.762281696 | 1 |
| WBGene00000102 | akt-1 | 1.209859495 | 0.985712594 | 0.323375623 | 0.143 | 0.198 | 0.11950939 | 1 |
| WBGene00012193 | vps-25 | 2.029953836 | 1.805752894 | 0.323453586 | 0.371 | 0.44 | 0.722540928 | 1 |
| WBGene00006438 | nrfl-1 | 0.665724884 | 0.441014753 | 0.324188192 | 0.065 | 0.067 | 0.924689573 | 1 |
| WBGene00022479 | fbxa-36 | 1.243829718 | 1.015724334 | 0.329086507 | 0.124 | 0.165 | 0.184303105 | 1 |
| WBGene00010974 | bbs-5 | 1.661125366 | 1.43262595 | 0.329654975 | 0.24 | 0.304 | 0.468957693 | 1 |
| WBGene00021922 | atg-3 | 0.567068655 | 0.338529891 | 0.329711741 | 0.055 | 0.054 | 0.811193508 | 1 |
| WBGene00020736 | T23F2.3 | 2.813410535 | 2.58468989 | 0.32997414 | 0.608 | 0.644 | 0.029723984 | 1 |
| WBGene00022824 | ZK813.5 | 1.45750308 | 1.228086563 | 0.330978072 | 0.208 | 0.248 | 0.720238947 | 1 |
| WBGene00021461 | nekl-1 | 0.738052718 | 0.508414519 | 0.331297891 | 0.065 | 0.071 | 0.870042592 | 1 |
| WBGene00194909 | F41E6.17 | 0.869057747 | 0.639266072 | 0.33151931 | 0.076 | 0.096 | 0.365956955 | 1 |
| WBGene00004138 | pqn-53 | 0.933359481 | 0.702879773 | 0.332511931 | 0.085 | 0.11 | 0.312316408 | 1 |
| WBGene00011292 | allo-1 | 1.801886933 | 1.571048034 | 0.333030136 | 0.279 | 0.338 | 0.588023881 | 1 |
| WBGene00009848 | F48F5.6 | 1.181559266 | 0.950364656 | 0.333543317 | 0.155 | 0.177 | 0.86417295 | 1 |
| WBGene00195233 | F31E9.11 | 0.530431462 | 0.298204296 | 0.33503298 | 0.053 | 0.046 | 0.514584735 | 1 |
| WBGene00044771 | tmem-218 | 0.579686046 | 0.34696857 | 0.335740348 | 0.051 | 0.056 | 0.872448213 | 1 |
| WBGene00006918 | vha-9 | 0.981710737 | 0.748738737 | 0.33610755 | 0.081 | 0.14 | 0.00748838 | 1 |
| WBGene00004350 | rgs-7 | 1.343037432 | 1.10998574 | 0.336222519 | 0.168 | 0.179 | 0.807361883 | 1 |
| WBGene00011763 | T14B1.1 | 1.726322862 | 1.492499896 | 0.337335235 | 0.242 | 0.265 | 0.645857684 | 1 |
| WBGene00004503 | rpt-3 | 0.686888705 | 0.45290231 | 0.337571011 | 0.053 | 0.056 | 0.963033317 | 1 |
| WBGene00020694 | dhhc-11 | 1.869759634 | 1.635723827 | 0.337642298 | 0.272 | 0.356 | 0.274047597 | 1 |
| WBGene00019324 | K02F2.5 | 2.576700902 | 2.341198863 | 0.339757624 | 0.488 | 0.577 | 0.748407448 | 1 |
| WBGene00004302 | ran-1 | 1.302560607 | 1.066992597 | 0.339852799 | 0.152 | 0.183 | 0.528868382 | 1 |
| WBGene00010887 | M7.12 | 1.645111266 | 1.409417695 | 0.340033946 | 0.226 | 0.256 | 0.970313748 | 1 |
| WBGene00020037 | mrps-6 | 0.659022464 | 0.422879613 | 0.34068212 | 0.064 | 0.065 | 0.93400162 | 1 |
| WBGene00000207 | asb-2 | 0.826072134 | 0.589495317 | 0.341308202 | 0.072 | 0.094 | 0.31352955 | 1 |
| WBGene00016037 | dop-6 | 1.169355293 | 0.932037776 | 0.342376805 | 0.15 | 0.165 | 0.933411505 | 1 |
| WBGene00016099 | C25E10.10 | 2.884520772 | 2.646353964 | 0.343602072 | 0.601 | 0.66 | 0.116932462 | 1 |
| WBGene00011146 | pde-2 | 1.264120561 | 1.023652057 | 0.346922719 | 0.157 | 0.177 | 0.862720479 | 1 |
| WBGene00011631 | T08G11.4 | 0.857109614 | 0.615737766 | 0.348225969 | 0.09 | 0.092 | 0.864637761 | 1 |

| | | | | | | | | |
|---|---|---|---|---|---|---|---|---|
| WBGene00007443 | pfd-1 | 0.89156001 | 0.649917332 | 0.348616693 | 0.072 | 0.112 | 0.055868842 | 1 |
| WBGene00006713 | ubc-18 | 0.907721072 | 0.665744791 | 0.349097981 | 0.083 | 0.108 | 0.303714038 | 1 |
| WBGene00010333 | copz-1 | 0.785367618 | 0.542864706 | 0.349857749 | 0.042 | 0.092 | 0.001892314 | 1 |
| WBGene00006498 | ten-1 | 0.841073471 | 0.597906442 | 0.350815866 | 0.064 | 0.075 | 0.559938831 | 1 |
| WBGene00005346 | srh-129 | 1.676191663 | 1.432255338 | 0.351925727 | 0.237 | 0.263 | 0.867308486 | 1 |
| WBGene00010504 | K02D3.1 | 2.391481397 | 2.144286839 | 0.356626364 | 0.337 | 0.452 | 0.085174262 | 1 |
| WBGene00010828 | M02B1.3 | 0.563744754 | 0.315508057 | 0.358129852 | 0.051 | 0.035 | 0.158903863 | 1 |
| WBGene00001749 | gst-1 | 0.594557274 | 0.346319361 | 0.358131606 | 0.046 | 0.052 | 0.756986948 | 1 |
| WBGene00004131 | pqn-44 | 2.412681303 | 2.164119494 | 0.358598889 | 0.422 | 0.475 | 0.598425077 | 1 |
| WBGene00017220 | F07F6.8 | 1.503733719 | 1.254843699 | 0.359072396 | 0.2 | 0.244 | 0.4886854 | 1 |
| WBGene00004859 | sma-5 | 0.65776701 | 0.408318698 | 0.359877844 | 0.065 | 0.058 | 0.472611899 | 1 |
| WBGene00007030 | epc-1 | 1.482376424 | 1.23213616 | 0.361020388 | 0.205 | 0.237 | 0.931025503 | 1 |
| WBGene00000802 | crt-1 | 1.442574384 | 1.192048682 | 0.361432188 | 0.159 | 0.212 | 0.21078721 | 1 |
| WBGene00009165 | gpx-1 | 1.807387186 | 1.55632933 | 0.362199924 | 0.247 | 0.34 | 0.096258131 | 1 |
| WBGene00012179 | W01D2.1 | 2.888393524 | 2.636877717 | 0.362860607 | 0.647 | 0.662 | 0.001661183 | 1 |
| WBGene00006462 | svh-5 | 1.337277061 | 1.084999051 | 0.363960234 | 0.163 | 0.173 | 0.772868121 | 1 |
| WBGene00021254 | Y22D7AL.16 | 1.076509635 | 0.823815896 | 0.364560003 | 0.133 | 0.138 | 0.786656321 | 1 |
| WBGene00022019 | sut-2 | 0.583026297 | 0.330204428 | 0.364744856 | 0.055 | 0.048 | 0.51443826 | 1 |
| WBGene00016395 | spcs-1 | 0.714264965 | 0.460897222 | 0.365532388 | 0.071 | 0.075 | 0.993307881 | 1 |
| WBGene00019686 | K12H6.6 | 0.872745348 | 0.618847027 | 0.366297849 | 0.095 | 0.079 | 0.242627421 | 1 |
| WBGene00010700 | nipi-3 | 1.332171349 | 1.077105971 | 0.367981555 | 0.17 | 0.196 | 0.827843772 | 1 |
| WBGene00001196 | egl-30 | 1.895002851 | 1.639791964 | 0.368191481 | 0.318 | 0.348 | 0.412301711 | 1 |
| WBGene00012140 | T28F4.5 | 2.71192294 | 2.456651573 | 0.368278735 | 0.466 | 0.533 | 0.786927426 | 1 |
| WBGene00019749 | M03A8.3 | 1.01724531 | 0.76153176 | 0.368916671 | 0.111 | 0.108 | 0.587323152 | 1 |
| WBGene00016292 | tbc-7 | 0.687971928 | 0.431453974 | 0.37007718 | 0.049 | 0.056 | 0.731186464 | 1 |
| WBGene00017184 | ncam-1 | 0.673177487 | 0.415285997 | 0.372058774 | 0.051 | 0.062 | 0.560437011 | 1 |
| WBGene00016982 | C56G2.9 | 1.648707811 | 1.390793845 | 0.3720912 | 0.249 | 0.292 | 0.888931585 | 1 |
| WBGene00044544 | F26F12.8 | 1.118169702 | 0.86016332 | 0.372224528 | 0.115 | 0.123 | 0.9106676 | 1 |
| WBGene00009931 | F52D10.4 | 1.037058396 | 0.778140131 | 0.373540097 | 0.12 | 0.115 | 0.533003867 | 1 |
| WBGene00017014 | dylt-2 | 0.741729837 | 0.482007216 | 0.374700538 | 0.074 | 0.077 | 0.917837144 | 1 |
| WBGene00020997 | gpa-18 | 1.743187156 | 1.483022022 | 0.37533895 | 0.247 | 0.31 | 0.362779593 | 1 |
| WBGene00000390 | cdc-42 | 0.757267884 | 0.496862268 | 0.375685891 | 0.081 | 0.075 | 0.539496611 | 1 |
| WBGene00009726 | tbc-13 | 0.588490868 | 0.324994776 | 0.380144505 | 0.051 | 0.05 | 0.797061363 | 1 |
| WBGene00018018 | F33G12.6 | 0.684302391 | 0.419484495 | 0.382051466 | 0.046 | 0.071 | 0.112163692 | 1 |
| WBGene00013298 | Y57G11B.6 | 1.005404014 | 0.740407609 | 0.382308999 | 0.117 | 0.125 | 0.956586375 | 1 |
| WBGene00016321 | tppp-1 | 1.966122494 | 1.700992297 | 0.382502021 | 0.297 | 0.348 | 0.917398318 | 1 |
| WBGene00012444 | Y15E3A.5 | 2.019771971 | 1.754326803 | 0.382956428 | 0.265 | 0.267 | 0.439865008 | 1 |
| WBGene00018519 | F46H5.3 | 1.029868174 | 0.763131319 | 0.384819938 | 0.122 | 0.11 | 0.286951845 | 1 |
| WBGene00077771 | C41G7.9 | 0.644121294 | 0.377298757 | 0.384943551 | 0.067 | 0.06 | 0.479113838 | 1 |
| WBGene00012413 | Y7A5A.7 | 1.870184715 | 1.60326582 | 0.385082567 | 0.304 | 0.285 | 0.005967459 | 1 |
| WBGene00022252 | Y73B6BL.31 | 1.037345505 | 0.770284555 | 0.385287508 | 0.11 | 0.117 | 0.992698163 | 1 |
| WBGene00012359 | W09D10.1 | 0.667069761 | 0.398979123 | 0.386773034 | 0.055 | 0.062 | 0.785983889 | 1 |
| WBGene00019870 | R04E5.7 | 1.053675786 | 0.782880137 | 0.39067554 | 0.125 | 0.123 | 0.529111954 | 1 |
| WBGene00006392 | taf-10 | 0.793713951 | 0.520495878 | 0.394170359 | 0.081 | 0.077 | 0.618783212 | 1 |
| WBGene00003915 | pan-1 | 1.402112982 | 1.128541826 | 0.39467975 | 0.161 | 0.183 | 0.813624408 | 1 |
| WBGene00002086 | ins-3 | 4.126412709 | 3.850953437 | 0.397403726 | 0.747 | 0.771 | 0.07172525 | 1 |
| WBGene00007055 | tag-196 | 0.846777972 | 0.570117687 | 0.399136422 | 0.09 | 0.077 | 0.299912753 | 1 |
| WBGene00017659 | F21C10.10 | 0.720646743 | 0.443917015 | 0.399236606 | 0.08 | 0.069 | 0.358969705 | 1 |
| WBGene00016258 | vha-16 | 1.156707808 | 0.876033373 | 0.404927614 | 0.092 | 0.138 | 0.046239162 | 1 |
| WBGene00011382 | dop-5 | 0.886324398 | 0.605501593 | 0.405141669 | 0.094 | 0.096 | 0.832979477 | 1 |
| WBGene00003610 | nhr-11 | 0.911479319 | 0.629646226 | 0.406599206 | 0.072 | 0.079 | 0.867870279 | 1 |
| WBGene00004453 | rpl-39 | 3.123815925 | 2.841567942 | 0.407197765 | 0.71 | 0.765 | 0.000655513 | 1 |
| WBGene00010110 | F55D12.1 | 1.655837332 | 1.373535731 | 0.40727512 | 0.244 | 0.285 | 0.900038585 | 1 |
| WBGene00019482 | K07C11.10 | 1.67233835 | 1.389341237 | 0.408278531 | 0.274 | 0.292 | 0.391486129 | 1 |
| WBGene00015868 | C16H3.3 | 1.021055387 | 0.737256175 | 0.409435717 | 0.08 | 0.113 | 0.116217261 | 1 |
| WBGene00017004 | nrd-1 | 0.560877932 | 0.2767614 | 0.409893512 | 0.051 | 0.035 | 0.1557592 | 1 |
| WBGene00219912 | F22H10.10 | 0.597108535 | 0.312191271 | 0.411048724 | 0.06 | 0.037 | 0.058260814 | 1 |
| WBGene00007485 | C09G1.3 | 2.064665088 | 1.779351805 | 0.411620059 | 0.35 | 0.373 | 0.487685886 | 1 |
| WBGene00003019 | lin-33 | 1.057035918 | 0.769690989 | 0.414551103 | 0.101 | 0.119 | 0.597323177 | 1 |
| WBGene00021854 | Y54F10AR.1 | 0.69946781 | 0.411781248 | 0.415043976 | 0.067 | 0.054 | 0.292530814 | 1 |
| WBGene00015626 | C09B9.1 | 1.9440439 | 1.655787615 | 0.415865913 | 0.299 | 0.367 | 0.662029244 | 1 |
| WBGene00000004 | aat-3 | 0.712746666 | 0.424127029 | 0.41639012 | 0.08 | 0.069 | 0.359638293 | 1 |
| WBGene00021610 | nhr-237 | 1.125766361 | 0.837091798 | 0.416469361 | 0.14 | 0.144 | 0.693229983 | 1 |
| WBGene00003954 | pcm-1 | 0.998761828 | 0.708815429 | 0.418304232 | 0.115 | 0.112 | 0.543501248 | 1 |

| | | | | | | | | |
|---|---|---|---|---|---|---|---|---|
| WBGene00008075 | C43F9.7 | 1.966507766 | 1.675910362 | 0.419243433 | 0.279 | 0.323 | 0.832842577 | 1 |
| WBGene00002855 | let-721 | 0.569617734 | 0.278221332 | 0.420396144 | 0.051 | 0.042 | 0.401320126 | 1 |
| WBGene00006372 | syx-2 | 0.60133125 | 0.309566308 | 0.420927835 | 0.053 | 0.048 | 0.597276274 | 1 |
| WBGene00003903 | pab-2 | 1.693986269 | 1.402133356 | 0.42105475 | 0.279 | 0.256 | 0.046454937 | 1 |
| WBGene00012339 | W07G4.5 | 1.071690967 | 0.779811246 | 0.421093426 | 0.125 | 0.14 | 0.883223802 | 1 |
| WBGene00004347 | rgs-4 | 0.716609318 | 0.424556351 | 0.421343367 | 0.062 | 0.06 | 0.744781816 | 1 |
| WBGene00019536 | rpi-2 | 0.686572104 | 0.393350678 | 0.423029098 | 0.057 | 0.062 | 0.861935536 | 1 |
| WBGene00021333 | Y34D9A.8 | 0.940118444 | 0.645060263 | 0.425678973 | 0.115 | 0.098 | 0.201415065 | 1 |
| WBGene00008444 | tbc-14 | 0.69338487 | 0.396827776 | 0.427841449 | 0.057 | 0.06 | 0.962062569 | 1 |
| WBGene00000973 | dhs-9 | 0.539708423 | 0.24229795 | 0.429072615 | 0.053 | 0.037 | 0.153487851 | 1 |
| WBGene00045190 | R01H2.8 | 1.792549547 | 1.495019255 | 0.429245477 | 0.288 | 0.319 | 0.545313087 | 1 |
| WBGene00015070 | srt-70 | 0.867468623 | 0.569038201 | 0.430544089 | 0.048 | 0.071 | 0.139057546 | 1 |
| WBGene00006923 | vhp-1 | 1.801014002 | 1.502581145 | 0.430547603 | 0.277 | 0.304 | 0.57541634 | 1 |
| WBGene00009774 | F46B6.9 | 0.794053485 | 0.495359012 | 0.430925035 | 0.055 | 0.06 | 0.860068487 | 1 |
| WBGene00000908 | daf-12 | 0.802677317 | 0.503581023 | 0.431504739 | 0.051 | 0.06 | 0.633410419 | 1 |
| WBGene00304812 | ZK380.6 | 4.493570063 | 4.193276975 | 0.433231349 | 0.627 | 0.612 | 0.056455005 | 1 |
| WBGene00008021 | C39B10.1 | 0.528086113 | 0.226750345 | 0.434735617 | 0.051 | 0.027 | 0.033560995 | 1 |
| WBGene00012625 | Y38H6C.13 | 0.78933151 | 0.485974406 | 0.43765179 | 0.064 | 0.05 | 0.26339154 | 1 |
| WBGene00000442 | ceh-19 | 2.242086533 | 1.938704877 | 0.43768721 | 0.413 | 0.413 | 0.047712198 | 1 |
| WBGene00019872 | R04E5.9 | 0.953763314 | 0.650365183 | 0.437710978 | 0.09 | 0.094 | 0.90349975 | 1 |
| WBGene00013111 | sta-1 | 1.439262492 | 1.134283362 | 0.439991878 | 0.194 | 0.202 | 0.500997109 | 1 |
| WBGene00013295 | Y57G11B.2 | 0.593848325 | 0.288754438 | 0.440157437 | 0.051 | 0.042 | 0.410271626 | 1 |
| WBGene00000414 | cec-1 | 1.40306031 | 1.097538667 | 0.44077456 | 0.168 | 0.179 | 0.755497815 | 1 |
| WBGene00006578 | tli-1 | 0.856690349 | 0.550276811 | 0.442061291 | 0.064 | 0.077 | 0.50677067 | 1 |
| WBGene00001470 | baz-2 | 0.936763373 | 0.630107082 | 0.44241151 | 0.074 | 0.098 | 0.269852 | 1 |
| WBGene00044443 | ZC21.10 | 1.254709135 | 0.947073439 | 0.443824494 | 0.115 | 0.125 | 0.88310909 | 1 |
| WBGene00001527 | gcs-1 | 0.675213074 | 0.366103808 | 0.445950405 | 0.051 | 0.038 | 0.269033635 | 1 |
| WBGene00008643 | tmem-138 | 0.639875955 | 0.329149522 | 0.448283484 | 0.055 | 0.054 | 0.786819986 | 1 |
| WBGene00000901 | daf-5 | 0.951227912 | 0.639762533 | 0.449349557 | 0.095 | 0.092 | 0.629817787 | 1 |
| WBGene00008343 | C56A3.4 | 0.530145917 | 0.216276411 | 0.45281798 | 0.053 | 0.035 | 0.110635649 | 1 |
| WBGene00007553 | C13G3.1 | 1.271181341 | 0.957140515 | 0.453065142 | 0.099 | 0.052 | 0.002379966 | 1 |
| WBGene00003687 | nhr-97 | 2.114185452 | 1.799463523 | 0.454047766 | 0.39 | 0.377 | 0.005799689 | 1 |
| WBGene00012764 | Y41E3.6 | 2.868522498 | 2.551907047 | 0.456779541 | 0.364 | 0.327 | 0.055606426 | 1 |
| WBGene00000092 | ags-3 | 2.160133063 | 1.840607101 | 0.460978521 | 0.39 | 0.396 | 0.087171119 | 1 |
| WBGene00017082 | DC2.5 | 0.692226911 | 0.371579496 | 0.462596435 | 0.057 | 0.058 | 0.918859259 | 1 |
| WBGene00019150 | H04J21.1 | 0.79693632 | 0.475095191 | 0.464318601 | 0.087 | 0.067 | 0.145025411 | 1 |
| WBGene00021931 | Y55F3AM.14 | 0.660487482 | 0.33861643 | 0.464361771 | 0.067 | 0.044 | 0.074116757 | 1 |
| WBGene00000552 | cmd-1 | 2.576972751 | 2.255050882 | 0.464435084 | 0.539 | 0.571 | 0.012036218 | 1 |
| WBGene00020738 | T23F2.5 | 1.923462094 | 1.60038596 | 0.466100337 | 0.304 | 0.36 | 0.897042837 | 1 |
| WBGene00003196 | mel-11 | 0.742546082 | 0.418758706 | 0.467126441 | 0.058 | 0.062 | 0.95979965 | 1 |
| WBGene00271808 | F21C10.19 | 0.972527065 | 0.647853458 | 0.468405004 | 0.088 | 0.098 | 0.840371703 | 1 |
| WBGene00003401 | mpk-1 | 1.496252014 | 1.168568309 | 0.472747656 | 0.203 | 0.223 | 0.722712026 | 1 |
| WBGene00019339 | K02F6.5 | 2.001103086 | 1.673387696 | 0.472793369 | 0.346 | 0.387 | 0.41581794 | 1 |
| WBGene00003634 | nhr-44 | 2.051714651 | 1.722845001 | 0.474458613 | 0.348 | 0.392 | 0.377713417 | 1 |
| WBGene00004389 | rnp-6 | 0.756467091 | 0.42642483 | 0.476150333 | 0.051 | 0.069 | 0.29025473 | 1 |
| WBGene00016304 | fbxc-41 | 1.430024973 | 1.098381468 | 0.478460441 | 0.173 | 0.212 | 0.49233236 | 1 |
| WBGene00010879 | taap-1 | 0.810978447 | 0.476714437 | 0.482241029 | 0.087 | 0.081 | 0.497187343 | 1 |
| WBGene00011935 | scrm-1 | 1.349607921 | 1.012103174 | 0.486916425 | 0.17 | 0.206 | 0.585404805 | 1 |
| WBGene00003963 | pdi-2 | 0.748968882 | 0.409718709 | 0.489434542 | 0.071 | 0.042 | 0.034453706 | 1 |
| WBGene00010329 | pcdr-1 | 1.219763993 | 0.877098297 | 0.4943621 | 0.148 | 0.146 | 0.461413837 | 1 |
| WBGene00012648 | Y39A1A.9 | 1.028595993 | 0.684988796 | 0.495720398 | 0.104 | 0.102 | 0.601055495 | 1 |
| WBGene00018634 | pigx-1 | 0.606582712 | 0.261840262 | 0.497358223 | 0.053 | 0.037 | 0.155201721 | 1 |
| WBGene00010918 | M117.1 | 1.348301739 | 1.003501099 | 0.497442174 | 0.161 | 0.179 | 0.956747244 | 1 |
| WBGene00022243 | Y73B6BL.22 | 1.576320355 | 1.23097943 | 0.49822164 | 0.201 | 0.235 | 0.811903414 | 1 |
| WBGene00018946 | F56C3.9 | 0.685022101 | 0.337992007 | 0.500658595 | 0.053 | 0.046 | 0.535026811 | 1 |
| WBGene00021043 | pck-1 | 0.593850349 | 0.245072236 | 0.503180454 | 0.062 | 0.038 | 0.057103503 | 1 |
| WBGene00017467 | F14F9.4 | 0.739668177 | 0.384981282 | 0.511705024 | 0.06 | 0.044 | 0.196965302 | 1 |
| WBGene00013878 | atfs-1 | 0.559280538 | 0.203262684 | 0.513625193 | 0.053 | 0.031 | 0.054186413 | 1 |
| WBGene00012412 | Y7A5A.6 | 0.773759715 | 0.41741805 | 0.514092353 | 0.083 | 0.069 | 0.247870463 | 1 |
| WBGene00006840 | unc-116 | 0.929901112 | 0.573538076 | 0.514123185 | 0.092 | 0.088 | 0.594930826 | 1 |
| WBGene00022648 | ZK54.3 | 1.048204509 | 0.691299068 | 0.514905711 | 0.081 | 0.092 | 0.706714858 | 1 |
| WBGene00015561 | C07A12.7 | 0.916017131 | 0.558636233 | 0.515591649 | 0.097 | 0.073 | 0.088085117 | 1 |
| WBGene00003408 | mrp-2 | 1.631967009 | 1.273748695 | 0.516799785 | 0.214 | 0.227 | 0.641861476 | 1 |
| WBGene00013228 | Y56A3A.7 | 0.788236406 | 0.429319404 | 0.517807779 | 0.071 | 0.071 | 0.821446882 | 1 |

| | | | | | | | | |
|---|---|---|---|---|---|---|---|---|
| WBGene00018290 | vps-60 | 0.902811776 | 0.543474412 | 0.518414233 | 0.087 | 0.081 | 0.512505737 | 1 |
| WBGene00219650 | linc-56 | 1.670382054 | 1.308511874 | 0.522068314 | 0.2 | 0.2 | 0.420830624 | 1 |
| WBGene00012002 | T24H10.4 | 0.984924472 | 0.619559869 | 0.5271097 | 0.097 | 0.073 | 0.09629514 | 1 |
| WBGene00011681 | T10B10.4 | 0.952178796 | 0.586666599 | 0.527322635 | 0.095 | 0.098 | 0.838087072 | 1 |
| WBGene00016448 | C35D10.12 | 1.133604048 | 0.764014483 | 0.533205033 | 0.08 | 0.112 | 0.1419024 | 1 |
| WBGene00001262 | emb-8 | 1.616946322 | 1.246493397 | 0.534450597 | 0.171 | 0.219 | 0.287540977 | 1 |
| WBGene00013683 | zoo-1 | 0.510363815 | 0.139569222 | 0.53494352 | 0.051 | 0.019 | 0.003762987 | 1 |
| WBGene00004373 | rme-1 | 1.16017362 | 0.788824234 | 0.535743918 | 0.148 | 0.137 | 0.225812519 | 1 |
| WBGene00012760 | Y41C4A.17 | 4.73219317 | 4.357338964 | 0.540800304 | 0.829 | 0.779 | 1.49E-06 | 0.009489534 |
| WBGene00016748 | aexr-3 | 1.09916154 | 0.723981243 | 0.541270753 | 0.111 | 0.106 | 0.483934593 | 1 |
| WBGene00010736 | istr-1 | 0.714454834 | 0.338574152 | 0.542281196 | 0.053 | 0.05 | 0.700724356 | 1 |
| WBGene00000298 | cat-4 | 2.538874843 | 2.161178643 | 0.544900435 | 0.489 | 0.548 | 0.121331127 | 1 |
| WBGene00019304 | K02D10.4 | 1.040369997 | 0.65547799 | 0.555281789 | 0.127 | 0.1 | 0.072201341 | 1 |
| WBGene00004208 | ptc-1 | 2.027694217 | 1.639081072 | 0.560650256 | 0.341 | 0.331 | 0.018471273 | 1 |
| WBGene00020366 | acdh-10 | 0.638003722 | 0.24751549 | 0.563355437 | 0.067 | 0.035 | 0.010877217 | 1 |
| WBGene00016025 | xbx-4 | 0.911556062 | 0.520323859 | 0.564428759 | 0.067 | 0.075 | 0.789777738 | 1 |
| WBGene00019402 | syf-2 | 0.776015597 | 0.384371878 | 0.565022452 | 0.06 | 0.054 | 0.55210598 | 1 |
| WBGene00001439 | fkh-7 | 1.727321569 | 1.332229201 | 0.5699978 | 0.281 | 0.263 | 0.03556169 | 1 |
| WBGene00016739 | pitr-1 | 1.011731814 | 0.612370647 | 0.576156375 | 0.102 | 0.077 | 0.074027926 | 1 |
| WBGene00003632 | nhr-42 | 0.709188422 | 0.308843004 | 0.57757635 | 0.055 | 0.048 | 0.505836168 | 1 |
| WBGene00189933 | F38B7.11 | 1.096097495 | 0.695016169 | 0.57863804 | 0.129 | 0.113 | 0.223267402 | 1 |
| WBGene00018750 | F53C3.6 | 0.672863862 | 0.270192783 | 0.580931569 | 0.067 | 0.04 | 0.034982355 | 1 |
| WBGene00010742 | K10D6.2 | 1.143490653 | 0.739474461 | 0.582872157 | 0.124 | 0.119 | 0.479429765 | 1 |
| WBGene00044007 | ZK1320.13 | 0.928191584 | 0.518767198 | 0.590674531 | 0.097 | 0.067 | 0.045907449 | 1 |
| WBGene00011434 | T04D3.5 | 0.963653145 | 0.553339531 | 0.591957416 | 0.11 | 0.079 | 0.049924462 | 1 |
| WBGene00007217 | tads-1 | 0.815070172 | 0.398245429 | 0.601350988 | 0.067 | 0.058 | 0.403700144 | 1 |
| WBGene00007981 | glb-11 | 1.61579925 | 1.198783609 | 0.601626398 | 0.239 | 0.21 | 0.024568337 | 1 |
| WBGene00000097 | aip-1 | 1.518025615 | 1.098462426 | 0.605301731 | 0.194 | 0.227 | 0.978297517 | 1 |
| WBGene00021611 | nhr-238 | 1.048063886 | 0.627510171 | 0.606730759 | 0.118 | 0.085 | 0.02887272 | 1 |
| WBGene00008310 | C54E10.6 | 0.894320825 | 0.471698564 | 0.60971504 | 0.083 | 0.073 | 0.380820012 | 1 |
| WBGene00001745 | gsa-1 | 1.472495377 | 1.047292655 | 0.613437859 | 0.173 | 0.169 | 0.40437141 | 1 |
| WBGene00016507 | abhd-5.2 | 2.130163165 | 1.704048725 | 0.61475319 | 0.364 | 0.381 | 0.052513787 | 1 |
| WBGene00010837 | M03B6.5 | 0.682267884 | 0.254781947 | 0.616731842 | 0.058 | 0.033 | 0.034681991 | 1 |
| WBGene00017907 | F28F9.3 | 2.124625479 | 1.697136095 | 0.616736814 | 0.345 | 0.346 | 0.063165694 | 1 |
| WBGene00018572 | lin-42 | 2.580484213 | 2.151439415 | 0.618980803 | 0.44 | 0.417 | 0.00463131 | 1 |
| WBGene00077536 | F38B7.10 | 1.348028003 | 0.916765473 | 0.622180314 | 0.159 | 0.162 | 0.4813746 | 1 |
| WBGene00019431 | K06A5.3 | 0.75852845 | 0.323680606 | 0.627352827 | 0.072 | 0.05 | 0.086279271 | 1 |
| WBGene00044742 | Y55F3BR.10 | 0.869606643 | 0.433867387 | 0.628638863 | 0.08 | 0.058 | 0.11324217 | 1 |
| WBGene00013647 | Y105C5B.5 | 0.963479238 | 0.526877523 | 0.62988313 | 0.076 | 0.058 | 0.182023546 | 1 |
| WBGene00015162 | algn-11 | 0.89064974 | 0.452291653 | 0.632417038 | 0.088 | 0.071 | 0.182866403 | 1 |
| WBGene00002232 | kpc-1 | 1.089387352 | 0.650699989 | 0.632892083 | 0.12 | 0.088 | 0.046964861 | 1 |
| WBGene00018869 | rfip-1 | 2.628153922 | 2.186744915 | 0.636818585 | 0.442 | 0.49 | 0.102643652 | 1 |
| WBGene00017675 | F21F3.6 | 1.647251588 | 1.203488878 | 0.640214261 | 0.239 | 0.233 | 0.140914591 | 1 |
| WBGene00007786 | rmd-2 | 0.885886844 | 0.437663701 | 0.646649305 | 0.087 | 0.073 | 0.253665087 | 1 |
| WBGene00003048 | lit-1 | 2.18547507 | 1.735723954 | 0.648853705 | 0.378 | 0.362 | 0.011743736 | 1 |
| WBGene00017597 | F19C7.8 | 2.134199119 | 1.683041545 | 0.650882796 | 0.314 | 0.34 | 0.386209908 | 1 |
| WBGene00020751 | srbc-69 | 0.583369546 | 0.130083826 | 0.65395306 | 0.06 | 0.017 | 0.000243993 | 1 |
| WBGene00002008 | hsp-4 | 0.889643197 | 0.432723137 | 0.659196305 | 0.058 | 0.038 | 0.113186634 | 1 |
| WBGene00007010 | alx-1 | 0.733989749 | 0.270465903 | 0.668723554 | 0.062 | 0.04 | 0.078881603 | 1 |
| WBGene00010262 | droe-4 | 3.541586635 | 3.076755209 | 0.670609993 | 0.353 | 0.321 | 0.038114931 | 1 |
| WBGene00005655 | srr-4 | 1.973451508 | 1.49342232 | 0.692535729 | 0.304 | 0.312 | 0.090118423 | 1 |
| WBGene00021011 | W03G1.5 | 0.985470129 | 0.50243126 | 0.696877781 | 0.094 | 0.062 | 0.030637224 | 1 |
| WBGene00017080 | lec-12 | 0.816899285 | 0.333330647 | 0.697642075 | 0.08 | 0.044 | 0.009734666 | 1 |
| WBGene00020812 | acdh-7 | 0.725853099 | 0.239881022 | 0.701109506 | 0.055 | 0.029 | 0.02721637 | 1 |
| WBGene00019770 | M04F3.4 | 0.830911311 | 0.341015423 | 0.706770368 | 0.083 | 0.04 | 0.002515882 | 1 |
| WBGene00050939 | C05G5.7 | 2.389709484 | 1.898277953 | 0.708985833 | 0.33 | 0.29 | 0.009544539 | 1 |
| WBGene00000534 | cpi-2 | 1.486882987 | 0.989368923 | 0.717761074 | 0.216 | 0.183 | 0.015869969 | 1 |
| WBGene00016175 | C27H5.6 | 1.870542043 | 1.367926858 | 0.725120436 | 0.247 | 0.233 | 0.093468402 | 1 |
| WBGene00009803 | F47B8.2 | 2.420388505 | 1.914379873 | 0.730016143 | 0.366 | 0.367 | 0.038272778 | 1 |
| WBGene00016488 | C36C9.5 | 1.419146482 | 0.906736249 | 0.739251703 | 0.187 | 0.152 | 0.018263636 | 1 |
| WBGene00009069 | F23A7.4 | 1.845828783 | 1.331183843 | 0.742475703 | 0.085 | 0.098 | 0.576073104 | 1 |
| WBGene00003647 | nhr-57 | 0.680617616 | 0.16211789 | 0.748036983 | 0.06 | 0.015 | 0.000107981 | 0.689026408 |
| WBGene00006959 | xbp-1 | 3.704979864 | 3.16860258 | 0.773828848 | 0.714 | 0.71 | 3.74E-05 | 0.238372721 |
| WBGene00019342 | K02F6.8 | 1.94040746 | 1.403518724 | 0.774566716 | 0.329 | 0.306 | 0.004705753 | 1 |

| | | | | | | | | |
|---|---|---|---|---|---|---|---|---|
| WBGene00021039 | W05F2.7 | 1.657914577 | 1.116225689 | 0.781491873 | 0.235 | 0.198 | 0.010925561 | 1 |
| WBGene00015168 | pdi-6 | 1.06356656 | 0.519769471 | 0.784533364 | 0.099 | 0.083 | 0.195207377 | 1 |
| WBGene00043308 | tmem-107 | 0.885143984 | 0.340621998 | 0.785579168 | 0.078 | 0.042 | 0.009456335 | 1 |
| WBGene00008865 | F15G9.1 | 0.561473973 | 0.013518104 | 0.790533215 | 0.053 | 0.002 | 4.27E-07 | 0.002725031 |
| WBGene00004134 | myrf-1 | 1.410617281 | 0.857844838 | 0.797482062 | 0.161 | 0.133 | 0.075416586 | 1 |
| WBGene00194851 | K07C10.3 | 0.90191868 | 0.346159388 | 0.801791174 | 0.092 | 0.048 | 0.002541647 | 1 |
| WBGene00009901 | F49E12.8 | 1.094001998 | 0.534341958 | 0.807418764 | 0.125 | 0.073 | 0.001690629 | 1 |
| WBGene00013845 | fbxa-37 | 0.610706954 | 0.050543908 | 0.80814445 | 0.051 | 0.008 | 2.56E-05 | 0.163087397 |
| WBGene00013266 | Y57A10A.26 | 1.770992048 | 1.207218513 | 0.813353283 | 0.272 | 0.248 | 0.009710042 | 1 |
| WBGene00011480 | enpl-1 | 1.130019048 | 0.566078466 | 0.81359428 | 0.095 | 0.09 | 0.498345544 | 1 |
| WBGene00022667 | ZK154.5 | 1.066008 | 0.50106397 | 0.81504195 | 0.101 | 0.071 | 0.044344446 | 1 |
| WBGene00008681 | scrm-4 | 2.0818274 | 1.510313319 | 0.82452053 | 0.353 | 0.288 | 3.61E-05 | 0.230261566 |
| WBGene00002113 | ins-30 | 1.861509073 | 1.28371739 | 0.833577196 | 0.247 | 0.165 | 1.87E-05 | 0.119051639 |
| WBGene00194710 | mks-2 | 1.548002694 | 0.937380334 | 0.880941849 | 0.201 | 0.152 | 0.00372708 | 1 |
| WBGene00194982 | Y17D7B.10 | 2.154440951 | 1.541172959 | 0.884758691 | 0.254 | 0.246 | 0.132325661 | 1 |
| WBGene00016506 | abhd-5.1 | 0.97881445 | 0.363869753 | 0.887177665 | 0.11 | 0.06 | 0.001043659 | 1 |
| WBGene00019006 | spg-20 | 1.153793253 | 0.522918067 | 0.910160503 | 0.134 | 0.081 | 0.001336911 | 1 |
| WBGene00014473 | MTCE.36 | 0.98340355 | 0.348726441 | 0.915645518 | 0.085 | 0.038 | 0.001088624 | 1 |
| WBGene00007366 | C06B3.7 | 0.908926918 | 0.268577119 | 0.92382948 | 0.092 | 0.038 | 0.00019724 | 1 |
| WBGene00011423 | ipla-7 | 0.846546132 | 0.205453094 | 0.924901747 | 0.072 | 0.033 | 0.002140052 | 1 |
| WBGene00015164 | ykt-6 | 1.682768293 | 1.026235625 | 0.947176424 | 0.201 | 0.177 | 0.040927521 | 1 |
| WBGene00000993 | dhs-30 | 0.685778141 | 0.024388554 | 0.954183477 | 0.069 | 0.004 | 1.73E-08 | 0.000110242 |
| WBGene00020596 | oga-1 | 1.322313651 | 0.645049493 | 0.977085642 | 0.184 | 0.094 | 2.27E-06 | 0.014458969 |
| WBGene00002267 | lec-4 | 0.709208887 | 0.010733863 | 1.007686454 | 0.076 | 0.002 | 6.04E-10 | 3.85E-06 |
| WBGene00002107 | ins-24 | 2.872049936 | 2.162240994 | 1.024037841 | 0.539 | 0.512 | 5.10E-07 | 0.003257326 |
| WBGene00019967 | cyp-33C8 | 1.698664145 | 0.9794944 | 1.037542624 | 0.208 | 0.133 | 8.94E-05 | 0.570315382 |
| WBGene00019975 | flp-34 | 1.293959147 | 0.559547435 | 1.059532136 | 0.141 | 0.085 | 0.000733713 | 1 |
| WBGene00017618 | F20A1.6 | 0.826982755 | 0.087525701 | 1.066811025 | 0.064 | 0.012 | 7.10E-06 | 0.045335845 |
| WBGene00001787 | gst-39 | 1.584123988 | 0.843921925 | 1.067885846 | 0.18 | 0.146 | 0.018609812 | 1 |
| WBGene00004443 | rpl-29 | 2.439736959 | 1.693906703 | 1.076005612 | 0.514 | 0.396 | 1.19E-13 | 7.59E-10 |
| WBGene00011045 | gpx-2 | 1.192488459 | 0.445910127 | 1.077084857 | 0.143 | 0.067 | 8.90E-06 | 0.056822472 |
| WBGene00019327 | zip-2 | 2.31979188 | 1.572472919 | 1.078153358 | 0.36 | 0.327 | 0.000385231 | 1 |
| WBGene00011232 | pck-2 | 1.094614563 | 0.343546944 | 1.083561529 | 0.085 | 0.05 | 0.012561516 | 1 |
| WBGene00022668 | ZK154.6 | 1.775640677 | 1.012684567 | 1.100712997 | 0.21 | 0.146 | 0.00041607 | 1 |
| WBGene00010833 | M03B6.1 | 2.6591928 | 1.883432126 | 1.119186077 | 0.44 | 0.427 | 0.000199698 | 1 |
| WBGene00006470 | tag-120 | 1.169808053 | 0.378252024 | 1.141973958 | 0.118 | 0.054 | 5.51E-05 | 0.351483289 |
| WBGene00005097 | srd-19 | 1.11435278 | 0.31389803 | 1.154812098 | 0.111 | 0.048 | 4.34E-05 | 0.276650361 |
| WBGene00008922 | F17C11.11 | 2.59761895 | 1.767747614 | 1.197251261 | 0.304 | 0.306 | 0.054371317 | 1 |
| WBGene00001755 | gst-7 | 1.913215563 | 1.073501752 | 1.211450952 | 0.178 | 0.146 | 0.034394069 | 1 |
| WBGene00008777 | F13H10.6 | 3.03116704 | 2.179950166 | 1.228046363 | 0.61 | 0.463 | 4.16E-18 | 2.66E-14 |
| WBGene00010019 | F54B8.4 | 1.073669208 | 0.216149994 | 1.237138717 | 0.118 | 0.029 | 1.09E-08 | 6.96E-05 |
| WBGene00016997 | cebp-1 | 1.476253442 | 0.603333122 | 1.259357817 | 0.187 | 0.1 | 2.60E-06 | 0.016591776 |
| WBGene00002263 | lea-1 | 1.046182057 | 0.171862349 | 1.261376706 | 0.101 | 0.017 | 6.30E-09 | 4.02E-05 |
| WBGene00021487 | comt-3 | 1.306580829 | 0.426301546 | 1.269974556 | 0.118 | 0.04 | 9.32E-07 | 0.005945444 |
| WBGene00043054 | T21B4.15 | 1.501461341 | 0.576583797 | 1.334316247 | 0.186 | 0.079 | 1.95E-08 | 0.000124699 |
| WBGene00003991 | pfn-3 | 1.207764874 | 0.22723802 | 1.41460123 | 0.115 | 0.025 | 5.41E-09 | 3.45E-05 |
| WBGene00015678 | mcp-1 | 1.701163038 | 0.708175839 | 1.432577708 | 0.217 | 0.087 | 1.36E-10 | 8.67E-07 |
| WBGene00007365 | C06B3.6 | 2.626298085 | 1.527359541 | 1.585433187 | 0.454 | 0.194 | 7.74E-23 | 4.94E-19 |
| WBGene00010291 | F58H10.1 | 1.567419212 | 0.347468941 | 1.760016206 | 0.157 | 0.042 | 9.03E-11 | 5.76E-07 |
| WBGene00012128 | nra-1 | 1.711233606 | 0.335189981 | 1.985211315 | 0.136 | 0.04 | 1.27E-08 | 8.11E-05 |
| WBGene00018702 | F52E4.5 | 1.420777167 | 0.035911417 | 1.997938949 | 0.129 | 0.006 | 1.38E-15 | 8.79E-12 |
| WBGene00194689 | Y38H6C.24 | 1.982176516 | 0.544828481 | 2.073654882 | 0.166 | 0.05 | 1.62E-10 | 1.03E-06 |
| WBGene00044644 | B0205.13 | 2.307606742 | 0.642989713 | 2.401534733 | 0.115 | 0.023 | 3.16E-09 | 2.02E-05 |
| WBGene00001753 | gst-5 | 2.467075744 | 0.688446366 | 2.566019783 | 0.196 | 0.081 | 2.44E-09 | 1.56E-05 |

This comparison is between all ADF cells in Cluster 2 and all ADF cells in Cluster 1.

| id | gene_name | c1.avg | c2.avg | avg_log2FC | c1.pct | c2.pct | p_val | p_val_adj |
|---|---|---|---|---|---|---|---|---|
| WBGene00014248 | ZK1307.7 | 1.69988503 | 0.298689147 | -2.021498352 | 0.299 | 0.038 | 1.95E-10 | 1.24E-06 |
| WBGene00011059 | R06C1.4 | 2.517017283 | 1.186775554 | -1.919133147 | 0.567 | 0.195 | 7.77E-16 | 4.96E-12 |
| WBGene00012281 | W05E10.2 | 3.580977837 | 2.307581291 | -1.837122881 | 0.823 | 0.489 | 5.45E-28 | 3.48E-24 |
| WBGene00012216 | W02D9.10 | 3.108369494 | 1.98023724 | -1.627550809 | 0.698 | 0.368 | 1.81E-17 | 1.16E-13 |
| WBGene00011372 | npr-26 | 1.618922371 | 0.534142381 | -1.565006712 | 0.272 | 0.06 | 1.71E-07 | 0.001088981 |
| WBGene00005903 | srx-12 | 2.492856309 | 1.4705784 | -1.47483527 | 0.572 | 0.195 | 1.79E-14 | 1.14E-10 |
| WBGene00005739 | srv-28 | 2.197058623 | 1.182901875 | -1.463118911 | 0.473 | 0.143 | 2.31E-11 | 1.47E-07 |
| WBGene00009503 | F37A8.5 | 1.809064097 | 0.796452208 | -1.46089015 | 0.327 | 0.098 | 1.29E-07 | 0.000821928 |
| WBGene00010146 | glb-19 | 1.95055386 | 0.948389819 | -1.445817092 | 0.363 | 0.113 | 1.62E-08 | 0.000103489 |
| WBGene00002245 | lag-1 | 1.892116522 | 0.911092968 | -1.415317815 | 0.392 | 0.105 | 9.33E-10 | 5.95E-06 |
| WBGene00005040 | sra-14 | 1.087164842 | 0.114812746 | -1.402807548 | 0.156 | 0.008 | 3.91E-06 | 0.024934238 |
| WBGene00004265 | qui-1 | 1.299943448 | 0.342023617 | -1.38198619 | 0.176 | 0.038 | 4.16E-05 | 0.265487604 |
| WBGene00021740 | Y50D4B.1 | 2.639207893 | 1.682986917 | -1.379535261 | 0.619 | 0.316 | 2.62E-12 | 1.67E-08 |
| WBGene00044484 | hot-9 | 2.640995772 | 1.699327201 | -1.358540577 | 0.593 | 0.316 | 1.66E-10 | 1.06E-06 |
| WBGene00044029 | C05D12.7 | 2.066595508 | 1.148962374 | -1.323864772 | 0.414 | 0.165 | 4.64E-08 | 0.000295871 |
| WBGene00044803 | C02G6.3 | 1.522843218 | 0.605640013 | -1.323244516 | 0.236 | 0.068 | 1.10E-05 | 0.069975289 |
| WBGene00010426 | H37A05.2 | 1.539833931 | 0.623919321 | -1.321385466 | 0.243 | 0.068 | 6.88E-06 | 0.04389869 |
| WBGene00002017 | hsp-16.11 | 4.044457391 | 3.140796637 | -1.303706889 | 0.818 | 0.632 | 1.15E-15 | 7.31E-12 |
| WBGene00015919 | C17G10.7 | 2.616871375 | 1.717130907 | -1.298051112 | 0.575 | 0.271 | 9.31E-11 | 5.94E-07 |
| WBGene00017439 | F13H8.6 | 1.031591744 | 0.153877758 | -1.266273614 | 0.14 | 0.015 | 5.30E-05 | 0.337969613 |
| WBGene00002265 | lec-2 | 1.014897702 | 0.151063846 | -1.246248821 | 0.123 | 0.015 | 0.000206595 | 1 |
| WBGene00002018 | hsp-16.41 | 3.950911241 | 3.092716065 | -1.238113924 | 0.578 | 0.444 | 2.45E-05 | 0.15649843 |
| WBGene00003728 | nhr-138 | 2.302548247 | 1.4514484 | -1.227877529 | 0.537 | 0.233 | 3.39E-10 | 2.16E-06 |
| WBGene00002016 | hsp-16.2 | 4.12570445 | 3.292413313 | -1.202184991 | 0.51 | 0.398 | 0.000652296 | 1 |
| WBGene00008570 | kcnl-2 | 1.814400334 | 0.983496526 | -1.198740803 | 0.32 | 0.12 | 4.28E-06 | 0.027334069 |
| WBGene00005346 | srh-129 | 1.63810678 | 0.812121557 | -1.191644784 | 0.27 | 0.105 | 3.98E-05 | 0.254062214 |
| WBGene00022479 | fbxa-36 | 1.211996961 | 0.393014119 | -1.181542485 | 0.157 | 0.045 | 0.000555547 | 1 |
| WBGene00003639 | nhr-49 | 0.976983358 | 0.15901579 | -1.180077755 | 0.134 | 0.015 | 8.37E-05 | 0.533842461 |
| WBGene00003808 | npr-2 | 0.813830352 | 0 | -1.174109014 | 0.087 | 0 | 0.00040656 | 1 |
| WBGene00009619 | npr-6 | 1.287566499 | 0.492778216 | -1.146637115 | 0.182 | 0.053 | 0.000242171 | 1 |
| WBGene00005210 | srg-53 | 1.426803802 | 0.632264927 | -1.146277294 | 0.225 | 0.06 | 1.70E-05 | 0.10845703 |
| WBGene00018187 | twf-2 | 0.980524161 | 0.19204049 | -1.137541483 | 0.133 | 0.023 | 0.000228928 | 1 |
| WBGene00003953 | pbs-7 | 1.092825401 | 0.306925728 | -1.13381356 | 0.143 | 0.038 | 0.000706543 | 1 |
| WBGene00017411 | F13A2.2 | 1.122891091 | 0.34822132 | -1.117612238 | 0.163 | 0.038 | 0.000163352 | 1 |
| WBGene00015735 | pdfr-1 | 1.450502844 | 0.682841229 | -1.107501605 | 0.209 | 0.068 | 0.000185241 | 1 |
| WBGene00016984 | npr-8 | 1.024373477 | 0.261155652 | -1.101090571 | 0.135 | 0.023 | 0.000227135 | 1 |
| WBGene00000209 | asg-1 | 2.152368987 | 1.390885746 | -1.098588096 | 0.473 | 0.195 | 2.76E-08 | 0.000176017 |
| WBGene00004744 | scp-1 | 1.643249518 | 0.890894423 | -1.085418964 | 0.157 | 0.083 | 0.018108883 | 1 |
| WBGene00000883 | cyn-7 | 2.694710586 | 1.944211031 | -1.082741986 | 0.636 | 0.331 | 5.10E-10 | 3.26E-06 |
| WBGene00008368 | D1053.3 | 1.479669375 | 0.729599052 | -1.082122736 | 0.244 | 0.075 | 3.01E-05 | 0.192337449 |
| WBGene00015293 | C01C4.3 | 1.909029316 | 1.165881589 | -1.07213554 | 0.371 | 0.128 | 1.24E-07 | 0.000788968 |
| WBGene00014215 | obr-3 | 0.959655038 | 0.217239049 | -1.071079865 | 0.101 | 0.023 | 0.003582371 | 1 |
| WBGene00004481 | rps-12 | 2.501479726 | 1.762803533 | -1.065684479 | 0.558 | 0.331 | 2.79E-07 | 0.001782179 |
| WBGene00000232 | avr-14 | 0.738239235 | 0 | -1.065054084 | 0.055 | 0 | 0.005815119 | 1 |
| WBGene00007892 | C33B4.4 | 2.98083159 | 2.248447986 | -1.056606195 | 0.688 | 0.429 | 1.97E-10 | 1.26E-06 |
| WBGene00003163 | mdl-1 | 1.459441806 | 0.731659806 | -1.049967481 | 0.235 | 0.09 | 0.000168773 | 1 |
| WBGene00017715 | F22F1.3 | 0.958504921 | 0.230867853 | -1.04975839 | 0.12 | 0.015 | 0.00029945 | 1 |
| WBGene00009691 | F44E5.4 | 4.428424915 | 3.705896326 | -1.042388413 | 0.892 | 0.857 | 4.85E-14 | 3.09E-10 |
| WBGene00009692 | F44E5.5 | 4.3983593 | 3.682259753 | -1.033113265 | 0.897 | 0.865 | 2.45E-17 | 1.56E-13 |
| WBGene00008248 | srsx-27 | 2.850798069 | 2.13473209 | -1.033064837 | 0.685 | 0.391 | 2.50E-10 | 1.60E-06 |
| WBGene00020790 | T25B6.4 | 1.568452997 | 0.855419994 | -1.028689179 | 0.221 | 0.098 | 0.001407594 | 1 |
| WBGene00000518 | ckk-1 | 0.904407849 | 0.192235821 | -1.027447054 | 0.137 | 0.015 | 6.75E-05 | 0.430452008 |
| WBGene00003511 | mxl-3 | 2.267883357 | 1.555987536 | -1.027048571 | 0.47 | 0.248 | 1.57E-06 | 0.009989462 |
| WBGene00003690 | nhr-100 | 1.688770488 | 0.977356031 | -1.026354109 | 0.302 | 0.09 | 9.38E-07 | 0.00598446 |
| WBGene00195073 | F02E11.7 | 2.007246503 | 1.29640256 | -1.025531032 | 0.409 | 0.188 | 2.94E-06 | 0.018777719 |
| WBGene00022410 | Y97E10C.1 | 1.656418603 | 0.946886911 | -1.023637854 | 0.278 | 0.113 | 7.28E-05 | 0.464299276 |
| WBGene00000239 | bas-1 | 1.350109 | 0.642440822 | -1.020949371 | 0.219 | 0.068 | 6.63E-05 | 0.42324488 |
| WBGene00020058 | R13D11.4 | 1.011446674 | 0.30442965 | -1.020009955 | 0.136 | 0.03 | 0.000474965 | 1 |
| WBGene00006801 | unc-68 | 0.969047873 | 0.264016022 | -1.017145955 | 0.136 | 0.023 | 0.000212303 | 1 |
| WBGene00010669 | bath-37 | 0.921700154 | 0.218320884 | -1.014761785 | 0.126 | 0.023 | 0.00048959 | 1 |
| WBGene00009119 | ndk-1 | 2.674287517 | 1.973645375 | -1.010812944 | 0.602 | 0.376 | 3.83E-07 | 0.00244074 |

| Legend | |
|---|---|
| id | WormBase GeneID |
| gene_name | Gene symbol (WS273) |
| c1.avg | Log of averaged normalized expression across all Cluster 1 ADF neurons |
| c2.avg | Log of averaged normalized expression across all Cluster 2 ADF neurons |
| avg_log2FC | log fold-change (Cluster 2 - Cluster 1) |
| c1.pct | Percent of Cluster 1 ADF neurons where gene is detected ($\geq$ 1 UMI) |
| c2.pct | Percent of Cluster 2 ADF neurons where gene is detected ($\geq$ 1 UMI) |
| p_val | p-value |
| p_val_adj | Adjusted p-value, based on bonferroni correction |

| | | | | | | | | |
|---|---|---|---|---|---|---|---|---|
| WBGene00009742 | F45H10.5 | 0.890508792 | 0.190722535 | -1.009578163 | 0.11 | 0.015 | 0.000649724 | 1 |
| WBGene00013490 | Y70C5A.2 | 0.903190246 | 0.204132157 | -1.008527639 | 0.142 | 0.023 | 0.000130525 | 0.832878205 |
| WBGene00019023 | F58A6.2 | 1.888005333 | 1.190151547 | -1.006790196 | 0.384 | 0.143 | 2.73E-07 | 0.001743028 |
| WBGene00015650 | pkg-2 | 1.463937301 | 0.767691481 | -1.004470392 | 0.24 | 0.09 | 0.00017015 | 1 |
| WBGene00015368 | srt-65 | 2.149450775 | 1.453369381 | -1.004233175 | 0.389 | 0.188 | 1.89E-05 | 0.120558399 |
| WBGene00017068 | D2092.8 | 0.852047176 | 0.164626048 | -0.991739053 | 0.051 | 0.015 | 0.062235783 | 1 |
| WBGene00010191 | dmsr-1 | 1.187388781 | 0.503441279 | -0.986727669 | 0.161 | 0.045 | 0.000463965 | 1 |
| WBGene00007862 | kvs-2 | 1.558624585 | 0.878588221 | -0.981085091 | 0.254 | 0.113 | 0.000342339 | 1 |
| WBGene00017926 | cox-6C | 1.053891382 | 0.37472783 | -0.97982589 | 0.131 | 0.038 | 0.002142991 | 1 |
| WBGene00008439 | mfb-1 | 2.427632221 | 1.750881867 | -0.97634438 | 0.548 | 0.286 | 6.00E-08 | 0.00038257 |
| WBGene00004344 | rgs-1 | 1.976689365 | 1.300977886 | -0.9748456 | 0.404 | 0.188 | 8.21E-06 | 0.052368283 |
| WBGene00007997 | sre-13 | 2.299971093 | 1.627093524 | -0.970757131 | 0.476 | 0.271 | 1.28E-05 | 0.081765594 |
| WBGene00008925 | F17H10.1 | 1.229719102 | 0.560026548 | -0.966162127 | 0.191 | 0.075 | 0.001231131 | 1 |
| WBGene00008249 | srsx-28 | 2.046932166 | 1.381292581 | -0.960314929 | 0.419 | 0.211 | 1.13E-05 | 0.072144164 |
| WBGene00009579 | arrd-6 | 1.895938224 | 1.231861705 | -0.958059901 | 0.375 | 0.173 | 1.88E-05 | 0.119919084 |
| WBGene00004622 | rrn-3.1 | 4.852434247 | 4.189034879 | -0.957082979 | 0.973 | 0.977 | 0.003169318 | 1 |
| WBGene00015478 | mapk-15 | 1.39425599 | 0.731833116 | -0.955674196 | 0.201 | 0.09 | 0.003145186 | 1 |
| WBGene00015307 | C01G5.5 | 3.599771488 | 2.93989496 | -0.952000594 | 0.858 | 0.692 | 2.62E-13 | 1.67E-09 |
| WBGene00011977 | T24B8.3 | 0.962421232 | 0.304426319 | -0.949285998 | 0.135 | 0.038 | 0.001291525 | 1 |
| WBGene00044387 | C27A2.8 | 1.998275565 | 1.340764616 | -0.948587785 | 0.383 | 0.241 | 0.000504803 | 1 |
| WBGene00009920 | abts-1 | 1.459379957 | 0.802370347 | -0.947864507 | 0.224 | 0.113 | 0.003982268 | 1 |
| WBGene00021820 | nipa-1 | 1.757200219 | 1.101183551 | -0.946431993 | 0.314 | 0.158 | 0.000314953 | 1 |
| WBGene00015750 | C14A11.5 | 1.90219409 | 1.247514559 | -0.944502913 | 0.369 | 0.158 | 8.63E-06 | 0.055047138 |
| WBGene00009999 | srsx-35 | 0.860458793 | 0.211209951 | -0.936668085 | 0.102 | 0.015 | 0.00128664 | 1 |
| WBGene00174531 | 21ur-10538 | 0.843213923 | 0.194717382 | -0.935582744 | 0.1 | 0.023 | 0.003646742 | 1 |
| WBGene00022887 | ZK1290.5 | 1.083595603 | 0.438473627 | -0.930714276 | 0.154 | 0.045 | 0.000978417 | 1 |
| WBGene00015514 | nlp-77 | 0.79103505 | 0.146966057 | -0.929195143 | 0.064 | 0.015 | 0.023976864 | 1 |
| WBGene00011974 | xbx-5 | 1.42804257 | 0.786842643 | -0.925055955 | 0.212 | 0.098 | 0.002801924 | 1 |
| WBGene00000210 | asg-2 | 0.961925746 | 0.322810507 | -0.922048386 | 0.107 | 0.038 | 0.01234052 | 1 |
| WBGene00021799 | Y53G8AL.1 | 0.933294834 | 0.298275301 | -0.916139531 | 0.085 | 0.03 | 0.027753833 | 1 |
| WBGene00017858 | F27C1.11 | 1.599401012 | 0.97081777 | -0.906853926 | 0.263 | 0.128 | 0.000846429 | 1 |
| WBGene00018466 | F45E1.5 | 1.269993907 | 0.641722421 | -0.906404158 | 0.2 | 0.075 | 0.000639937 | 1 |
| WBGene00044353 | ZK418.10 | 1.001878724 | 0.37421353 | -0.905529463 | 0.153 | 0.045 | 0.000906239 | 1 |
| WBGene00008382 | D1081.4 | 1.052428822 | 0.425936058 | -0.903838003 | 0.124 | 0.045 | 0.008885532 | 1 |
| WBGene00015373 | C03B1.2 | 1.21141692 | 0.585042954 | -0.903666616 | 0.174 | 0.068 | 0.002145377 | 1 |
| WBGene00018018 | F33G12.6 | 0.624778447 | 0 | -0.901364767 | 0.066 | 0 | 0.002276844 | 1 |
| WBGene00013898 | ZC443.3 | 1.615818231 | 0.9910486 | -0.901352047 | 0.289 | 0.128 | 0.000156314 | 0.997440718 |
| WBGene00000472 | cey-1 | 2.090525096 | 1.470785391 | -0.894095399 | 0.443 | 0.226 | 9.94E-06 | 0.063395987 |
| WBGene00020114 | pde-4 | 1.517323267 | 0.897765228 | -0.89383331 | 0.247 | 0.135 | 0.004139364 | 1 |
| WBGene00012113 | T28B8.1 | 1.781168036 | 1.162627167 | -0.892365845 | 0.308 | 0.18 | 0.003144396 | 1 |
| WBGene00007405 | C07A9.9 | 0.683842742 | 0.065828387 | -0.891606244 | 0.08 | 0.008 | 0.002425604 | 1 |
| WBGene00001747 | gsp-1 | 1.250889338 | 0.634785385 | -0.888850118 | 0.186 | 0.068 | 0.001084518 | 1 |
| WBGene00006780 | unc-44 | 1.762797356 | 1.148265711 | -0.886581757 | 0.284 | 0.165 | 0.007364533 | 1 |
| WBGene00194905 | Y53F4B.51 | 0.688633178 | 0.075480753 | -0.884591963 | 0.088 | 0.008 | 0.001255191 | 1 |
| WBGene00015076 | B0238.12 | 0.838270758 | 0.225893926 | -0.883473018 | 0.095 | 0.023 | 0.005516798 | 1 |
| WBGene00017317 | attf-2 | 1.158913175 | 0.547971387 | -0.881402688 | 0.161 | 0.068 | 0.005386801 | 1 |
| WBGene00020586 | frpr-18 | 1.696301595 | 1.08987124 | -0.874894065 | 0.294 | 0.15 | 0.001070035 | 1 |
| WBGene00001980 | hmr-1 | 0.921820962 | 0.315887347 | -0.874177421 | 0.125 | 0.038 | 0.003159294 | 1 |
| WBGene00009216 | F28D1.6 | 2.8124861 | 2.209261211 | -0.870269555 | 0.632 | 0.459 | 1.42E-05 | 0.090428719 |
| WBGene00015263 | B0563.6 | 0.602732019 | 0 | -0.869558495 | 0.066 | 0 | 0.002276843 | 1 |
| WBGene00016099 | C25E10.10 | 2.834550988 | 2.231865947 | -0.86949072 | 0.651 | 0.474 | 1.03E-05 | 0.065785779 |
| WBGene00002065 | iff-2 | 1.762785796 | 1.163122397 | -0.865131412 | 0.325 | 0.15 | 0.000117492 | 0.749718757 |
| WBGene00022078 | epg-9 | 1.417550569 | 0.819282449 | -0.86311845 | 0.238 | 0.105 | 0.000859889 | 1 |
| WBGene00001588 | ggr-3 | 0.957695812 | 0.362598102 | -0.858544515 | 0.121 | 0.045 | 0.008771619 | 1 |
| WBGene00001303 | sec-61.G | 1.494244324 | 0.902649336 | -0.853491155 | 0.251 | 0.135 | 0.004686754 | 1 |
| WBGene00019537 | K08D12.3 | 1.309118725 | 0.719744298 | -0.850287563 | 0.215 | 0.105 | 0.002863549 | 1 |
| WBGene00004435 | rpl-23 | 2.222947658 | 1.633753382 | -0.85002766 | 0.484 | 0.271 | 2.10E-05 | 0.134201902 |
| WBGene00004677 | rrn-3.56 | 1.559572312 | 0.971603703 | -0.848259396 | 0.163 | 0.128 | 0.349158472 | 1 |
| WBGene00011073 | R07A4.2 | 1.172906524 | 0.585817933 | -0.846989799 | 0.161 | 0.075 | 0.01079784 | 1 |
| WBGene00017792 | F25E5.16 | 0.711809142 | 0.125532409 | -0.845818534 | 0.087 | 0.015 | 0.003645698 | 1 |
| WBGene00014308 | F07H5.4 | 2.254217793 | 1.668384222 | -0.845179188 | 0.493 | 0.316 | 0.000154204 | 0.983976037 |
| WBGene00002266 | lec-3 | 0.903825722 | 0.3229966 | -0.837959295 | 0.097 | 0.03 | 0.012663049 | 1 |
| WBGene00012235 | W04A4.6 | 1.624243045 | 1.046084852 | -0.834105957 | 0.285 | 0.158 | 0.002785009 | 1 |
| WBGene00001682 | gpc-2 | 2.195955707 | 1.618408186 | -0.833224944 | 0.492 | 0.278 | 1.98E-05 | 0.126264412 |

| | | | | | | | | |
|---|---|---|---|---|---|---|---|---|
| WBGene00013859 | ZC247.1 | 1.564509839 | 0.987268031 | -0.832783893 | 0.232 | 0.135 | 0.016315056 | 1 |
| WBGene00194866 | Y41C4A.22 | 1.064247098 | 0.48739854 | -0.832216554 | 0.133 | 0.053 | 0.008034005 | 1 |
| WBGene00001427 | fkb-2 | 1.58309646 | 1.007129201 | -0.830945108 | 0.286 | 0.135 | 0.000598707 | 1 |
| WBGene00009848 | F48F5.6 | 1.132597306 | 0.556717982 | -0.830818246 | 0.18 | 0.06 | 0.000608657 | 1 |
| WBGene00001091 | drh-2 | 0.699281361 | 0.123497786 | -0.830680108 | 0.06 | 0.008 | 0.01235635 | 1 |
| WBGene00010600 | nhr-196 | 0.700686592 | 0.127239966 | -0.827308603 | 0.093 | 0.015 | 0.00227012 | 1 |
| WBGene00006789 | unc-54 | 5.294894307 | 4.723744764 | -0.823994613 | 0.983 | 0.985 | 6.29E-27 | 4.02E-23 |
| WBGene00000282 | cah-4 | 1.325314169 | 0.7548463 | -0.823011165 | 0.216 | 0.113 | 0.005942999 | 1 |
| WBGene00015350 | fbxl-1 | 1.670345873 | 1.100256993 | -0.822464401 | 0.303 | 0.165 | 0.001663817 | 1 |
| WBGene00004431 | rpl-19 | 2.570969867 | 2.001163011 | -0.822057525 | 0.597 | 0.391 | 1.30E-05 | 0.083083187 |
| WBGene00009180 | nurf-1 | 1.920026702 | 1.351608803 | -0.820053684 | 0.332 | 0.218 | 0.010257455 | 1 |
| WBGene00015619 | C08G9.1 | 1.889065588 | 1.321200107 | -0.819256713 | 0.333 | 0.195 | 0.002712475 | 1 |
| WBGene00015938 | anat-1 | 0.909101679 | 0.341842242 | -0.818382376 | 0.129 | 0.038 | 0.002435018 | 1 |
| WBGene00000056 | acr-17 | 0.637338032 | 0.071325879 | -0.816582926 | 0.067 | 0.008 | 0.006789967 | 1 |
| WBGene00002000 | hrp-2 | 1.040865958 | 0.475072039 | -0.816268081 | 0.124 | 0.06 | 0.032093185 | 1 |
| WBGene00001005 | dlc-1 | 1.505762783 | 0.940375517 | -0.815681404 | 0.265 | 0.135 | 0.001900873 | 1 |
| WBGene00020390 | T10B5.4 | 2.096502465 | 1.532414973 | -0.813806228 | 0.442 | 0.218 | 1.60E-05 | 0.10190934 |
| WBGene00008920 | eef-1G | 2.321416816 | 1.759059539 | -0.811310054 | 0.525 | 0.331 | 3.90E-05 | 0.249066457 |
| WBGene00004451 | rpl-37 | 2.317479553 | 1.756986312 | -0.80862082 | 0.509 | 0.338 | 0.000131404 | 0.838485897 |
| WBGene00006702 | ubc-3 | 1.934549789 | 1.374776317 | -0.807582412 | 0.356 | 0.195 | 0.000527948 | 1 |
| WBGene00018572 | lin-42 | 2.451556066 | 1.89281994 | -0.806085839 | 0.446 | 0.308 | 0.00300236 | 1 |
| WBGene00022732 | ZK418.2 | 1.136440748 | 0.577997117 | -0.805663857 | 0.157 | 0.045 | 0.000645948 | 1 |
| WBGene00015074 | natc-2 | 1.858502619 | 1.300794506 | -0.804602729 | 0.376 | 0.203 | 0.000326879 | 1 |
| WBGene00004507 | rpy-1 | 0.689027456 | 0.134305423 | -0.800294726 | 0.077 | 0.015 | 0.008988132 | 1 |
| WBGene00018846 | eef-1B.1 | 2.649784168 | 2.096480106 | -0.798249027 | 0.622 | 0.383 | 1.21E-06 | 0.007735748 |
| WBGene00010677 | gtbp-1 | 0.887601278 | 0.334470847 | -0.79799853 | 0.123 | 0.045 | 0.007380814 | 1 |
| WBGene00017239 | F08B6.1 | 0.786811124 | 0.23384572 | -0.797760446 | 0.102 | 0.023 | 0.00348556 | 1 |
| WBGene00013639 | Y105C5A.15 | 1.149596617 | 0.598418583 | -0.795181816 | 0.142 | 0.068 | 0.023694406 | 1 |
| WBGene00006779 | unc-43 | 1.967295692 | 1.417952436 | -0.792534791 | 0.393 | 0.188 | 1.78E-05 | 0.1135923 |
| WBGene00001976 | hmg-11 | 1.011426586 | 0.462144948 | -0.792445896 | 0.146 | 0.038 | 0.000832151 | 1 |
| WBGene00001475 | fmi-1 | 0.920348701 | 0.371186157 | -0.792274078 | 0.124 | 0.038 | 0.003717231 | 1 |
| WBGene00006756 | unc-17 | 1.647013808 | 1.100277085 | -0.788774359 | 0.295 | 0.158 | 0.002011004 | 1 |
| WBGene00195086 | F21C10.13 | 0.927277674 | 0.383056755 | -0.785144821 | 0.126 | 0.038 | 0.00331569 | 1 |
| WBGene00013136 | Y53C10A.5 | 1.0101493 | 0.468694349 | -0.781154372 | 0.141 | 0.053 | 0.005136397 | 1 |
| WBGene00011273 | R53.4 | 1.405725199 | 0.864968113 | -0.780147566 | 0.226 | 0.12 | 0.00655152 | 1 |
| WBGene00010507 | K02E2.6 | 0.992533279 | 0.453127246 | -0.778198408 | 0.141 | 0.06 | 0.009865466 | 1 |
| WBGene00002045 | icd-1 | 1.685945768 | 1.14871947 | -0.775053715 | 0.318 | 0.173 | 0.00205476 | 1 |
| WBGene00004492 | rps-23 | 2.942617038 | 2.40661619 | -0.773285766 | 0.707 | 0.579 | 9.00E-07 | 0.005742036 |
| WBGene00010280 | rde-12 | 0.535774963 | 0 | -0.772959882 | 0.063 | 0 | 0.002941607 | 1 |
| WBGene00008250 | srsx-29 | 1.942401039 | 1.406723996 | -0.772818614 | 0.392 | 0.226 | 0.000566702 | 1 |
| WBGene00007696 | tram-1 | 0.645577823 | 0.111392355 | -0.770666725 | 0.064 | 0.015 | 0.022646056 | 1 |
| WBGene00004567 | rrn-2.1 | 2.20698655 | 1.673588578 | -0.769530608 | 0.398 | 0.278 | 0.006676771 | 1 |
| WBGene00003162 | mdh-2 | 0.751831099 | 0.219258341 | -0.768340076 | 0.077 | 0.023 | 0.022835401 | 1 |
| WBGene00004484 | rps-15 | 2.488707125 | 1.958393419 | -0.765080954 | 0.568 | 0.376 | 3.97E-05 | 0.253355601 |
| WBGene00003839 | ocr-2 | 1.692482138 | 1.162278444 | -0.764922239 | 0.292 | 0.165 | 0.004238912 | 1 |
| WBGene00015070 | srt-70 | 0.787269901 | 0.257604938 | -0.764145015 | 0.064 | 0.023 | 0.060599062 | 1 |
| WBGene00044699 | nhr-286 | 1.056357504 | 0.52827924 | -0.761855893 | 0.158 | 0.068 | 0.006267783 | 1 |
| WBGene00004467 | rpn-11 | 1.053580149 | 0.528237794 | -0.75790881 | 0.147 | 0.053 | 0.004013387 | 1 |
| WBGene00044764 | W01A8.7 | 1.38663721 | 0.861819448 | -0.757151982 | 0.216 | 0.113 | 0.008760726 | 1 |
| WBGene00013159 | Y53F4B.12 | 1.027505632 | 0.503131735 | -0.75651162 | 0.136 | 0.068 | 0.029019893 | 1 |
| WBGene00007861 | C32A9.1 | 0.521449211 | 0 | -0.75229219 | 0.058 | 0 | 0.004504593 | 1 |
| WBGene00017098 | mek-5 | 0.861893101 | 0.341565902 | -0.750673468 | 0.09 | 0.038 | 0.04184663 | 1 |
| WBGene00009367 | F33H2.3 | 0.661085608 | 0.141944343 | -0.748962529 | 0.08 | 0.015 | 0.007152118 | 1 |
| WBGene00022336 | Y82E9BR.3 | 2.26775037 | 1.749536874 | -0.74762404 | 0.496 | 0.331 | 0.000726756 | 1 |
| WBGene00015767 | hex-2 | 0.993247535 | 0.477276523 | -0.74438882 | 0.137 | 0.06 | 0.013598989 | 1 |
| WBGene00006309 | sul-2 | 0.675436405 | 0.160453535 | -0.742963232 | 0.073 | 0.015 | 0.011829165 | 1 |
| WBGene00009772 | ztf-7 | 1.329477255 | 0.815730472 | -0.741179936 | 0.192 | 0.113 | 0.030761974 | 1 |
| WBGene00020275 | atp-4 | 0.763960429 | 0.250640056 | -0.740564756 | 0.119 | 0.03 | 0.002232142 | 1 |
| WBGene00016448 | C35D10.12 | 1.023910757 | 0.51067685 | -0.740440013 | 0.102 | 0.045 | 0.043371503 | 1 |
| WBGene00006599 | tpa-1 | 0.641455839 | 0.128626012 | -0.739857047 | 0.071 | 0.015 | 0.012943117 | 1 |
| WBGene00022405 | srsx-6 | 1.275526893 | 0.763746881 | -0.738342486 | 0.204 | 0.105 | 0.011975898 | 1 |
| WBGene00004703 | rsp-6 | 0.96710865 | 0.457943765 | -0.734569654 | 0.137 | 0.06 | 0.011990379 | 1 |
| WBGene00020930 | hlh-30 | 1.548286505 | 1.043794759 | -0.72782774 | 0.242 | 0.135 | 0.011384965 | 1 |
| WBGene00010200 | F57C7.4 | 1.529241105 | 1.025019334 | -0.727438248 | 0.251 | 0.128 | 0.003689932 | 1 |

| | | | | | | | | |
|---|---|---|---|---|---|---|---|---|
| WBGene00000501 | cho-1 | 0.844152439 | 0.344767246 | -0.720460541 | 0.102 | 0.038 | 0.017991573 | 1 |
| WBGene00004680 | rars-2 | 0.537454225 | 0.038198386 | -0.720273924 | 0.067 | 0.008 | 0.006238608 | 1 |
| WBGene00019177 | H10D18.1 | 1.263464894 | 0.764502627 | -0.719850389 | 0.211 | 0.098 | 0.002736552 | 1 |
| WBGene00009995 | F53F4.13 | 0.790582876 | 0.29172502 | -0.719699755 | 0.058 | 0.03 | 0.189828816 | 1 |
| WBGene00021078 | unc-132 | 0.573855647 | 0.075480753 | -0.719002988 | 0.057 | 0.008 | 0.01597892 | 1 |
| WBGene00000122 | aly-3 | 1.091300071 | 0.594241158 | -0.717104429 | 0.148 | 0.06 | 0.007299879 | 1 |
| WBGene00017724 | F22F7.7 | 0.708399061 | 0.212808588 | -0.714985917 | 0.08 | 0.023 | 0.01823123 | 1 |
| WBGene00004704 | rsp-7 | 0.585139163 | 0.090585025 | -0.713490803 | 0.066 | 0.008 | 0.00747968 | 1 |
| WBGene00016500 | memo-1 | 0.850125586 | 0.355626515 | -0.713411357 | 0.106 | 0.038 | 0.015241861 | 1 |
| WBGene00000879 | cyn-3 | 1.486934354 | 0.993447115 | -0.711951592 | 0.271 | 0.12 | 0.000469959 | 1 |
| WBGene00022250 | Y73B6BL.29 | 0.979407118 | 0.48650429 | -0.711108466 | 0.134 | 0.06 | 0.018286352 | 1 |
| WBGene00004512 | rrn-1.1 | 1.360238253 | 0.868660585 | -0.709196664 | 0.173 | 0.083 | 0.011967571 | 1 |
| WBGene00000058 | acr-19 | 0.640832787 | 0.149951986 | -0.708191296 | 0.078 | 0.015 | 0.008550669 | 1 |
| WBGene00019603 | K09H11.6 | 1.166077331 | 0.676656766 | -0.706084623 | 0.145 | 0.083 | 0.056596537 | 1 |
| WBGene00016097 | C25E10.8 | 1.751054055 | 1.263834096 | -0.702909818 | 0.327 | 0.188 | 0.002094758 | 1 |
| WBGene00269383 | ZK384.7 | 0.911854426 | 0.425303714 | -0.7019443 | 0.109 | 0.038 | 0.011612146 | 1 |
| WBGene00021697 | gcn-1 | 0.658434437 | 0.172254378 | -0.701409559 | 0.061 | 0.015 | 0.031165543 | 1 |
| WBGene00004430 | rpl-18 | 2.587525645 | 2.102112705 | -0.700302841 | 0.608 | 0.398 | 1.08E-05 | 0.068789183 |
| WBGene00012528 | pap-1 | 1.610340245 | 1.126280651 | -0.698350375 | 0.268 | 0.165 | 0.01499259 | 1 |
| WBGene00004505 | rpt-5 | 0.776000832 | 0.292312446 | -0.697814835 | 0.073 | 0.03 | 0.067462384 | 1 |
| WBGene00016706 | C46C11.3 | 0.807927126 | 0.324896229 | -0.696866281 | 0.088 | 0.03 | 0.024656427 | 1 |
| WBGene00011156 | rbm-3.2 | 2.742121157 | 2.259510888 | -0.696259442 | 0.656 | 0.496 | 3.50E-05 | 0.223486289 |
| WBGene00044588 | ZK1236.9 | 1.097381812 | 0.615724607 | -0.694884461 | 0.17 | 0.06 | 0.001513024 | 1 |
| WBGene00010357 | H03A11.2 | 0.541390036 | 0.061618982 | -0.692163321 | 0.057 | 0.008 | 0.015442815 | 1 |
| WBGene00016748 | aexr-3 | 0.984635782 | 0.505432399 | -0.691344345 | 0.116 | 0.053 | 0.028355466 | 1 |
| WBGene00019719 | M01H9.3 | 3.206632288 | 2.728736244 | -0.689458253 | 0.766 | 0.624 | 5.72E-07 | 0.003649568 |
| WBGene00009161 | cox-7C | 1.012440102 | 0.537533488 | -0.685145418 | 0.145 | 0.06 | 0.009212439 | 1 |
| WBGene00001945 | his-71 | 2.636107997 | 2.161211694 | -0.685130541 | 0.605 | 0.459 | 0.000309649 | 1 |
| WBGene00006529 | tba-2 | 1.163074977 | 0.689373324 | -0.683407025 | 0.164 | 0.09 | 0.029689716 | 1 |
| WBGene00003407 | mrp-1 | 0.670740571 | 0.197740473 | -0.682394897 | 0.077 | 0.015 | 0.009531924 | 1 |
| WBGene00008438 | DH11.5 | 0.707085005 | 0.235234363 | -0.68073658 | 0.09 | 0.03 | 0.018297463 | 1 |
| WBGene00000287 | cal-3 | 0.669769035 | 0.19865405 | -0.679675252 | 0.073 | 0.023 | 0.026608695 | 1 |
| WBGene00007943 | C34F6.9 | 0.705535425 | 0.236292695 | -0.676974159 | 0.098 | 0.015 | 0.001859744 | 1 |
| WBGene00015248 | mai-2 | 1.180540811 | 0.715207422 | -0.671334173 | 0.17 | 0.098 | 0.039592157 | 1 |
| WBGene00004836 | sls-2.4 | 1.205731985 | 0.742009031 | -0.669010807 | 0.152 | 0.068 | 0.011939369 | 1 |
| WBGene00004478 | rps-9 | 2.926798358 | 2.463075472 | -0.669010708 | 0.718 | 0.534 | 1.92E-05 | 0.122281002 |
| WBGene00004285 | rab-37 | 1.669568921 | 1.206993234 | -0.66735565 | 0.312 | 0.165 | 0.001481933 | 1 |
| WBGene00010967 | nduo-5 | 1.768695856 | 1.306206719 | -0.667230785 | 0.271 | 0.211 | 0.192270372 | 1 |
| WBGene00017961 | nhr-180 | 0.577178882 | 0.114812746 | -0.667053332 | 0.051 | 0.008 | 0.024731134 | 1 |
| WBGene00004426 | rpl-14 | 2.562736947 | 2.100564327 | -0.666774147 | 0.589 | 0.406 | 0.000227645 | 1 |
| WBGene00303425 | R02E12.12 | 0.615313735 | 0.154169908 | -0.665289912 | 0.076 | 0.015 | 0.009902906 | 1 |
| WBGene00001171 | egl-2 | 0.733776875 | 0.273052449 | -0.664684845 | 0.077 | 0.023 | 0.024609167 | 1 |
| WBGene00000371 | cox-5B | 1.48595469 | 1.025380976 | -0.664467414 | 0.261 | 0.135 | 0.002393156 | 1 |
| WBGene00006717 | ubc-22 | 0.5570697 | 0.097018464 | -0.663713637 | 0.066 | 0.008 | 0.007593363 | 1 |
| WBGene00019492 | K07E3.4 | 0.737392497 | 0.277666499 | -0.663244417 | 0.099 | 0.023 | 0.004524975 | 1 |
| WBGene00004705 | rsp-8 | 0.59671596 | 0.137270916 | -0.662839087 | 0.082 | 0.008 | 0.002168421 | 1 |
| WBGene00018344 | npr-27 | 0.810667197 | 0.351846926 | -0.66193773 | 0.109 | 0.038 | 0.010823787 | 1 |
| WBGene00016033 | C24A3.2 | 0.655873833 | 0.197318518 | -0.661555479 | 0.08 | 0.015 | 0.007647243 | 1 |
| WBGene00011282 | R74.8 | 0.600247397 | 0.142812505 | -0.659939051 | 0.052 | 0.015 | 0.059436363 | 1 |
| WBGene00004417 | rpl-6 | 2.43746685 | 1.981037555 | -0.65848828 | 0.562 | 0.353 | 9.35E-05 | 0.59678587 |
| WBGene00004700 | rsp-3 | 1.241269981 | 0.78525247 | -0.657894202 | 0.206 | 0.105 | 0.006640006 | 1 |
| WBGene00002019 | hsp-16.48 | 1.466447272 | 1.011987253 | -0.655647215 | 0.194 | 0.135 | 0.103565813 | 1 |
| WBGene00012853 | Y44A6D.2 | 2.082690444 | 1.628422325 | -0.655370363 | 0.375 | 0.241 | 0.00432811 | 1 |
| WBGene00004440 | rpl-26 | 2.352089647 | 1.898967579 | -0.65371696 | 0.536 | 0.383 | 0.001058631 | 1 |
| WBGene00016258 | vha-16 | 1.07770149 | 0.624836362 | -0.653346273 | 0.122 | 0.06 | 0.04253383 | 1 |
| WBGene00008453 | E02A10.3 | 2.876150685 | 2.424579912 | -0.651478914 | 0.695 | 0.504 | 3.47E-05 | 0.221359349 |
| WBGene00019607 | K10B2.4 | 0.688004961 | 0.237139617 | -0.650461197 | 0.082 | 0.023 | 0.015914926 | 1 |
| WBGene00016165 | wdr-60 | 0.99456314 | 0.544018932 | -0.649997894 | 0.141 | 0.068 | 0.022287786 | 1 |
| WBGene00018879 | F55D10.4 | 0.551330405 | 0.100793223 | -0.649987758 | 0.059 | 0.008 | 0.013577673 | 1 |
| WBGene00018508 | F46F11.1 | 0.899977534 | 0.450217659 | -0.648866342 | 0.101 | 0.053 | 0.077620741 | 1 |
| WBGene00017571 | jmjd-3.1 | 0.818458799 | 0.368864994 | -0.648626752 | 0.111 | 0.038 | 0.010267253 | 1 |
| WBGene00001193 | egl-26 | 1.231180719 | 0.783216414 | -0.646275881 | 0.221 | 0.09 | 0.000955881 | 1 |
| WBGene00019944 | R07G3.8 | 0.761437661 | 0.315772338 | -0.642959151 | 0.102 | 0.038 | 0.01793497 | 1 |
| WBGene00008149 | pyp-1 | 0.916351206 | 0.470997377 | -0.64250976 | 0.111 | 0.053 | 0.039763039 | 1 |

| | | | | | | | | |
|---|---|---|---|---|---|---|---|---|
| WBGene00019983 | sti-1 | 0.913996655 | 0.468652399 | -0.64249595 | 0.123 | 0.053 | 0.019822447 | 1 |
| WBGene00004349 | rgs-6 | 0.804345611 | 0.359176772 | -0.642242877 | 0.099 | 0.045 | 0.046439294 | 1 |
| WBGene00002242 | kvs-1 | 1.121938579 | 0.67858312 | -0.639626721 | 0.153 | 0.083 | 0.03685877 | 1 |
| WBGene00004286 | rab-39 | 0.66330113 | 0.220592473 | -0.638693584 | 0.069 | 0.015 | 0.017474298 | 1 |
| WBGene00004267 | rab-3 | 1.314836352 | 0.877120118 | -0.631491041 | 0.213 | 0.128 | 0.029765343 | 1 |
| WBGene00000908 | daf-12 | 0.714702248 | 0.279645787 | -0.627653799 | 0.059 | 0.03 | 0.178721382 | 1 |
| WBGene00004415 | rpl-4 | 2.241952536 | 1.806914124 | -0.627627759 | 0.496 | 0.331 | 0.000682126 | 1 |
| WBGene00003619 | nhr-20 | 1.551584658 | 1.116712525 | -0.62738787 | 0.29 | 0.158 | 0.002907057 | 1 |
| WBGene00017984 | F32D1.5 | 0.505034743 | 0.073201617 | -0.623003509 | 0.054 | 0.008 | 0.020357381 | 1 |
| WBGene00004204 | swsn-4 | 0.833229901 | 0.401420761 | -0.622968905 | 0.099 | 0.045 | 0.045605381 | 1 |
| WBGene00016250 | hsp-110 | 2.155715124 | 1.724031601 | -0.622787678 | 0.451 | 0.286 | 0.00133209 | 1 |
| WBGene00017198 | nhr-36 | 0.701091388 | 0.270085267 | -0.621810393 | 0.076 | 0.03 | 0.054416455 | 1 |
| WBGene00005704 | sru-41 | 0.983276618 | 0.552501531 | -0.621477082 | 0.129 | 0.068 | 0.045147934 | 1 |
| WBGene00044544 | F26F12.8 | 1.04667993 | 0.616058257 | -0.621255752 | 0.127 | 0.06 | 0.028465022 | 1 |
| WBGene00001854 | hil-3 | 0.638351345 | 0.208483028 | -0.62016889 | 0.072 | 0.015 | 0.013229272 | 1 |
| WBGene00020196 | T03G11.3 | 1.630672136 | 1.201012855 | -0.619867314 | 0.312 | 0.165 | 0.001243094 | 1 |
| WBGene00006705 | ubc-8 | 1.157470594 | 0.72834454 | -0.619098029 | 0.176 | 0.083 | 0.010830842 | 1 |
| WBGene00015056 | B0222.5 | 0.500604531 | 0.071529532 | -0.619024373 | 0.051 | 0.008 | 0.023758108 | 1 |
| WBGene00044303 | hst-3.2 | 0.579094962 | 0.150646474 | -0.61812051 | 0.068 | 0.015 | 0.017612944 | 1 |
| WBGene00008262 | ril-1 | 0.910951199 | 0.482509978 | -0.618110025 | 0.121 | 0.053 | 0.021256193 | 1 |
| WBGene00004425 | rpl-13 | 2.849065221 | 2.421653544 | -0.616624706 | 0.687 | 0.549 | 6.33E-05 | 0.403764768 |
| WBGene00044557 | ZC13.10 | 2.21262761 | 1.7858684 | -0.615683395 | 0.468 | 0.278 | 0.000125427 | 0.800347913 |
| WBGene00012484 | car-1 | 0.558008105 | 0.132815464 | -0.613423315 | 0.054 | 0.015 | 0.053556328 | 1 |
| WBGene00013311 | sec-61.A | 0.812620133 | 0.388765199 | -0.61149341 | 0.102 | 0.038 | 0.01964102 | 1 |
| WBGene00018468 | cla-1 | 1.880530651 | 1.456771653 | -0.611355006 | 0.334 | 0.218 | 0.021757057 | 1 |
| WBGene00014307 | F07H5.3 | 3.356386027 | 2.932788256 | -0.611122403 | 0.796 | 0.684 | 1.03E-06 | 0.006567533 |
| WBGene00004419 | rpl-7A | 2.297138986 | 1.875398818 | -0.60844245 | 0.52 | 0.331 | 0.000416433 | 1 |
| WBGene00006839 | unc-115 | 0.700187878 | 0.279171702 | -0.607397949 | 0.07 | 0.023 | 0.038224634 | 1 |
| WBGene00015799 | C15H9.3 | 2.245352742 | 1.825522419 | -0.605687126 | 0.48 | 0.323 | 0.00306414 | 1 |
| WBGene00011531 | rsbp-1 | 1.087456136 | 0.667826355 | -0.605397805 | 0.162 | 0.06 | 0.00340162 | 1 |
| WBGene00018214 | F39H12.3 | 1.616449309 | 1.19738557 | -0.604581179 | 0.291 | 0.188 | 0.019046609 | 1 |
| WBGene00006527 | tax-6 | 1.621393148 | 1.203032059 | -0.603567469 | 0.263 | 0.18 | 0.036401859 | 1 |
| WBGene00012548 | Y37D8A.6 | 2.510649487 | 2.093435879 | -0.601912003 | 0.596 | 0.414 | 3.78E-05 | 0.241100801 |
| WBGene00006918 | vha-9 | 0.919534698 | 0.502712389 | -0.601347478 | 0.118 | 0.053 | 0.032703536 | 1 |
| WBGene00006862 | pnk-4 | 0.557996377 | 0.14144867 | -0.600951311 | 0.061 | 0.015 | 0.031311679 | 1 |
| WBGene00009380 | F34H10.3 | 0.670304962 | 0.254150123 | -0.600384523 | 0.088 | 0.023 | 0.009974184 | 1 |
| WBGene00022042 | icd-2 | 1.911002492 | 1.494883625 | -0.600332626 | 0.393 | 0.218 | 0.000376373 | 1 |
| WBGene00001329 | epn-1 | 1.784499611 | 1.36850597 | -0.600151964 | 0.313 | 0.173 | 0.001307678 | 1 |
| WBGene00004701 | rsp-4 | 0.470884822 | 0.054951936 | -0.600064312 | 0.05 | 0.008 | 0.025885428 | 1 |
| WBGene00015113 | B0281.6 | 0.834902256 | 0.419021659 | -0.599988875 | 0.09 | 0.045 | 0.088480764 | 1 |
| WBGene00006352 | sur-6 | 0.941039519 | 0.525412299 | -0.599623329 | 0.128 | 0.06 | 0.028375395 | 1 |
| WBGene00004485 | rps-16 | 2.072450011 | 1.658208615 | -0.597624009 | 0.419 | 0.308 | 0.015635378 | 1 |
| WBGene00004888 | smo-1 | 1.334565483 | 0.921171095 | -0.596402033 | 0.21 | 0.12 | 0.026172254 | 1 |
| WBGene00001648 | goa-1 | 0.961963207 | 0.550137675 | -0.594138653 | 0.141 | 0.06 | 0.012090931 | 1 |
| WBGene00000263 | F23H11.5 | 1.270498842 | 0.859315396 | -0.593212318 | 0.206 | 0.113 | 0.017493729 | 1 |
| WBGene00002178 | jnk-1 | 0.831742097 | 0.421379974 | -0.5920274 | 0.107 | 0.053 | 0.051809113 | 1 |
| WBGene00014472 | MTCE.33 | 1.895439914 | 1.485330411 | -0.591662946 | 0.28 | 0.226 | 0.20280107 | 1 |
| WBGene00015354 | snet-1 | 1.044782065 | 0.634919858 | -0.591306174 | 0.047 | 0.06 | 0.504906856 | 1 |
| WBGene00010731 | K10C3.4 | 1.529809321 | 1.120601347 | -0.590362314 | 0.247 | 0.15 | 0.023866055 | 1 |
| WBGene00016493 | C37A2.7 | 2.427156168 | 2.018424458 | -0.589675212 | 0.561 | 0.414 | 0.001558608 | 1 |
| WBGene00011720 | T11G6.5 | 0.492860772 | 0.085727584 | -0.587369031 | 0.056 | 0.008 | 0.017334397 | 1 |
| WBGene00011824 | T18D3.7 | 1.185725443 | 0.77931677 | -0.586323777 | 0.199 | 0.105 | 0.012589924 | 1 |
| WBGene00008890 | ser-5 | 0.780845434 | 0.374673999 | -0.585981515 | 0.098 | 0.03 | 0.0123709 | 1 |
| WBGene00018803 | fbxa-24 | 2.922825736 | 2.516764059 | -0.585823168 | 0.574 | 0.481 | 0.019314339 | 1 |
| WBGene00013766 | prmt-1 | 0.766225426 | 0.360305394 | -0.585618818 | 0.087 | 0.038 | 0.05597719 | 1 |
| WBGene00000134 | amt-2 | 0.669419152 | 0.264713626 | -0.583866655 | 0.079 | 0.023 | 0.020947889 | 1 |
| WBGene00018067 | npr-7 | 0.802674598 | 0.398592378 | -0.582967415 | 0.079 | 0.023 | 0.02174762 | 1 |
| WBGene00001470 | baz-2 | 0.843111923 | 0.439231507 | -0.582676273 | 0.092 | 0.038 | 0.038739758 | 1 |
| WBGene00012967 | kvs-4 | 1.042019771 | 0.638194121 | -0.582597263 | 0.146 | 0.068 | 0.01680458 | 1 |
| WBGene00044509 | K07H8.12 | 0.950017625 | 0.548793035 | -0.578844727 | 0.143 | 0.053 | 0.00484678 | 1 |
| WBGene00011300 | R107.5 | 1.863433612 | 1.462402087 | -0.578566192 | 0.316 | 0.211 | 0.021960187 | 1 |
| WBGene00004427 | rpl-15 | 2.608242276 | 2.207770853 | -0.577758136 | 0.621 | 0.474 | 0.000545133 | 1 |
| WBGene00015139 | B0310.3 | 1.163102144 | 0.762713549 | -0.577638641 | 0.182 | 0.098 | 0.021484006 | 1 |
| WBGene00006831 | unc-104 | 0.850576523 | 0.450277967 | -0.577508742 | 0.098 | 0.06 | 0.16092477 | 1 |

| | | | | | | | | |
|---|---|---|---|---|---|---|---|---|
| WBGene00012372 | W09H1.1 | 1.630340102 | 1.230459396 | -0.576905912 | 0.278 | 0.158 | 0.005430178 | 1 |
| WBGene00014309 | F07H5.5 | 2.406941073 | 2.007451857 | -0.576341111 | 0.541 | 0.421 | 0.008949896 | 1 |
| WBGene00000182 | arf-1.2 | 0.800141985 | 0.400920683 | -0.575954592 | 0.105 | 0.045 | 0.031915578 | 1 |
| WBGene00000900 | daf-4 | 0.627022709 | 0.229042732 | -0.574163739 | 0.068 | 0.023 | 0.043248773 | 1 |
| WBGene00011198 | R10E4.1 | 0.789590975 | 0.392789088 | -0.572464114 | 0.113 | 0.03 | 0.004106901 | 1 |
| WBGene00016407 | unk-1 | 0.682675141 | 0.286331744 | -0.571802653 | 0.081 | 0.03 | 0.040177398 | 1 |
| WBGene00020279 | glb-25 | 1.790705219 | 1.39444182 | -0.57168724 | 0.35 | 0.233 | 0.011948866 | 1 |
| WBGene00007330 | C05C10.3 | 0.731251335 | 0.336477198 | -0.56953869 | 0.099 | 0.023 | 0.004893192 | 1 |
| WBGene00016119 | C25H3.10 | 2.888049941 | 2.493355105 | -0.569424283 | 0.64 | 0.602 | 0.011859465 | 1 |
| WBGene00007184 | ctr-9 | 0.706096811 | 0.312132997 | -0.56836964 | 0.071 | 0.03 | 0.077423051 | 1 |
| WBGene00044079 | tag-241 | 0.666310808 | 0.273426322 | -0.566812499 | 0.064 | 0.023 | 0.060797767 | 1 |
| WBGene00007352 | cdc-48.1 | 0.645826937 | 0.253281992 | -0.566322646 | 0.077 | 0.03 | 0.051095434 | 1 |
| WBGene00003519 | nac-3 | 0.724712178 | 0.333412598 | -0.564525963 | 0.093 | 0.038 | 0.035955045 | 1 |
| WBGene00006706 | ubc-9 | 0.615693732 | 0.224547808 | -0.564304285 | 0.07 | 0.023 | 0.038031246 | 1 |
| WBGene00044728 | Y53F4B.45 | 1.908969703 | 1.518193685 | -0.563770623 | 0.3 | 0.165 | 0.002692784 | 1 |
| WBGene00004499 | rps-30 | 2.887794649 | 2.49704477 | -0.563732913 | 0.697 | 0.579 | 0.000475727 | 1 |
| WBGene00000491 | che-12 | 1.441543023 | 1.051293131 | -0.563011584 | 0.226 | 0.158 | 0.095208282 | 1 |
| WBGene00006538 | tbb-4 | 1.467012234 | 1.076899571 | -0.562813604 | 0.205 | 0.165 | 0.308432585 | 1 |
| WBGene00011015 | R04F11.2 | 1.192083776 | 0.80205774 | -0.562688628 | 0.176 | 0.09 | 0.018536403 | 1 |
| WBGene00020893 | T28C12.6 | 0.673718937 | 0.285277541 | -0.560402476 | 0.076 | 0.03 | 0.05736956 | 1 |
| WBGene00003616 | nhr-17 | 0.922888618 | 0.53496965 | -0.559648771 | 0.115 | 0.068 | 0.093884366 | 1 |
| WBGene00004923 | snt-3 | 1.193526214 | 0.80561388 | -0.5596392 | 0.193 | 0.12 | 0.047063089 | 1 |
| WBGene00004494 | rps-25 | 3.321358184 | 2.935527191 | -0.55663646 | 0.808 | 0.669 | 1.45E-07 | 0.000926628 |
| WBGene00001423 | fib-1 | 0.614399055 | 0.228915494 | -0.556135221 | 0.058 | 0.03 | 0.179144674 | 1 |
| WBGene00020444 | T12B3.1 | 0.534593418 | 0.149333496 | -0.555812579 | 0.064 | 0.015 | 0.024620776 | 1 |
| WBGene00003081 | lsm-7 | 0.472087551 | 0.0869748 | -0.555600256 | 0.055 | 0.015 | 0.046361332 | 1 |
| WBGene00001085 | dpy-26 | 0.681756207 | 0.297735296 | -0.554025064 | 0.07 | 0.03 | 0.07973032 | 1 |
| WBGene00005834 | srw-87 | 0.622974715 | 0.239580624 | -0.553120753 | 0.085 | 0.023 | 0.012915375 | 1 |
| WBGene00010936 | ztf-14 | 2.605062046 | 2.223204173 | -0.550904459 | 0.613 | 0.444 | 0.000915105 | 1 |
| WBGene00009100 | rgef-1 | 0.447246388 | 0.067543965 | -0.547794803 | 0.056 | 0.008 | 0.017371285 | 1 |
| WBGene00017653 | F21C10.3 | 2.225506267 | 1.846469996 | -0.546833749 | 0.46 | 0.323 | 0.010335194 | 1 |
| WBGene00012825 | Y43F8C.3 | 0.683173202 | 0.305092061 | -0.545455787 | 0.077 | 0.03 | 0.054336203 | 1 |
| WBGene00004401 | rom-2 | 0.602676039 | 0.225296306 | -0.544443869 | 0.089 | 0.03 | 0.020082857 | 1 |
| WBGene00004464 | rpn-8 | 0.716654403 | 0.339584634 | -0.543996686 | 0.104 | 0.023 | 0.003132071 | 1 |
| WBGene00000434 | ceh-9 | 0.585434845 | 0.208535312 | -0.543751086 | 0.064 | 0.015 | 0.024830903 | 1 |
| WBGene00021427 | sec-61.B | 0.666578127 | 0.289710809 | -0.54370461 | 0.075 | 0.03 | 0.06309837 | 1 |
| WBGene00004482 | rps-13 | 2.496263842 | 2.120304479 | -0.542394709 | 0.569 | 0.429 | 0.000596632 | 1 |
| WBGene00011259 | miz-1 | 0.507880801 | 0.132050048 | -0.542209164 | 0.061 | 0.015 | 0.030788336 | 1 |
| WBGene00010419 | atp-1 | 1.655507519 | 1.280007536 | -0.541731963 | 0.274 | 0.18 | 0.033558223 | 1 |
| WBGene00014886 | Y41E3.5 | 3.272377894 | 2.897744533 | -0.540481692 | 0.742 | 0.639 | 5.91E-05 | 0.377335577 |
| WBGene00194657 | T07D10.10 | 0.591413434 | 0.217648194 | -0.539229259 | 0.063 | 0.023 | 0.063748816 | 1 |
| WBGene00022062 | Y67D8B.2 | 0.699729415 | 0.326109814 | -0.539019146 | 0.081 | 0.038 | 0.078666809 | 1 |
| WBGene00013899 | ZC443.4 | 2.04427581 | 1.670979412 | -0.538552863 | 0.407 | 0.278 | 0.007074837 | 1 |
| WBGene00011860 | pigc-1 | 0.714531828 | 0.342433346 | -0.536824635 | 0.08 | 0.038 | 0.085044901 | 1 |
| WBGene00008516 | F02C12.2 | 1.351459778 | 0.979404746 | -0.536761949 | 0.237 | 0.12 | 0.004602254 | 1 |
| WBGene00022738 | fubl-2 | 0.592832085 | 0.221002508 | -0.536436686 | 0.071 | 0.03 | 0.072177013 | 1 |
| WBGene00009716 | F44G4.7 | 1.250671796 | 0.879615915 | -0.535320479 | 0.194 | 0.12 | 0.050562244 | 1 |
| WBGene00016734 | C46H11.10 | 0.916529791 | 0.545786122 | -0.534870052 | 0.144 | 0.06 | 0.011025717 | 1 |
| WBGene00010647 | K08C7.4 | 0.897656734 | 0.526917861 | -0.534863134 | 0.112 | 0.06 | 0.075271139 | 1 |
| WBGene00010500 | K02B12.7 | 0.556377331 | 0.18623208 | -0.534006718 | 0.067 | 0.023 | 0.046039408 | 1 |
| WBGene00000288 | cal-4 | 2.09490679 | 1.726047002 | -0.532152187 | 0.46 | 0.278 | 0.001117249 | 1 |
| WBGene00020894 | T28D9.1 | 2.157849622 | 1.78970406 | -0.531121776 | 0.443 | 0.308 | 0.007300944 | 1 |
| WBGene00010110 | F55D12.1 | 1.568960038 | 1.201097853 | -0.53071295 | 0.273 | 0.195 | 0.079210848 | 1 |
| WBGene00009357 | F33C8.2 | 0.53914864 | 0.172283995 | -0.529273805 | 0.057 | 0.015 | 0.044308263 | 1 |
| WBGene00020683 | ribo-1 | 0.560630245 | 0.194104025 | -0.52878556 | 0.052 | 0.023 | 0.13495601 | 1 |
| WBGene00006697 | uaf-1 | 0.641871053 | 0.275710796 | -0.528257588 | 0.076 | 0.03 | 0.056976978 | 1 |
| WBGene00016100 | C25E10.11 | 1.634576944 | 1.268749328 | -0.527777687 | 0.276 | 0.188 | 0.047868679 | 1 |
| WBGene00004470 | rps-1 | 2.879013199 | 2.513426029 | -0.527430797 | 0.71 | 0.564 | 0.000281525 | 1 |
| WBGene00001117 | dyf-1 | 1.06622897 | 0.700834831 | -0.527152312 | 0.166 | 0.09 | 0.031302583 | 1 |
| WBGene00018467 | sdpn-1 | 1.67773717 | 1.313414353 | -0.52560672 | 0.295 | 0.203 | 0.056873586 | 1 |
| WBGene00004491 | rps-22 | 2.623231301 | 2.259044552 | -0.525410416 | 0.648 | 0.421 | 0.000117133 | 0.747425856 |
| WBGene00003124 | mai-1 | 0.634558257 | 0.270452108 | -0.525294134 | 0.069 | 0.023 | 0.040694205 | 1 |
| WBGene00001971 | hmg-1.1 | 1.328969944 | 0.965359828 | -0.524578511 | 0.207 | 0.12 | 0.028157169 | 1 |
| WBGene00000379 | cct-4 | 0.802153193 | 0.439077768 | -0.523807116 | 0.087 | 0.045 | 0.106577389 | 1 |

| | | | | | | | | |
|---|---|---|---|---|---|---|---|---|
| WBGene00008932 | F18A11.5 | 2.679208356 | 2.316441331 | -0.523362187 | 0.581 | 0.459 | 0.001030758 | 1 |
| WBGene00005605 | srj-17 | 0.519581998 | 0.157378723 | -0.522548869 | 0.059 | 0.015 | 0.03749901 | 1 |
| WBGene00012074 | T27A8.2 | 0.447283358 | 0.085241337 | -0.522316228 | 0.055 | 0.008 | 0.018804991 | 1 |
| WBGene00022790 | ZK682.7 | 1.508408894 | 1.14668134 | -0.521862548 | 0.241 | 0.158 | 0.054699236 | 1 |
| WBGene00007016 | mdt-15 | 0.722571825 | 0.361694854 | -0.520635416 | 0.089 | 0.045 | 0.085492909 | 1 |
| WBGene00000912 | daf-16 | 1.339941115 | 0.979270126 | -0.520338247 | 0.207 | 0.135 | 0.073901505 | 1 |
| WBGene00012094 | T27E9.2 | 0.847300948 | 0.487205106 | -0.519508485 | 0.098 | 0.053 | 0.105525049 | 1 |
| WBGene00011471 | T05C12.8 | 1.486090958 | 1.127052107 | -0.517983569 | 0.253 | 0.158 | 0.027593146 | 1 |
| WBGene00003045 | lir-2 | 0.510278907 | 0.151350079 | -0.51782484 | 0.057 | 0.015 | 0.043182087 | 1 |
| WBGene00010642 | mks-6 | 1.043505437 | 0.684605422 | -0.517783272 | 0.158 | 0.083 | 0.030657013 | 1 |
| WBGene00015868 | C16H3.3 | 0.932707769 | 0.574538749 | -0.51672867 | 0.101 | 0.06 | 0.146723864 | 1 |
| WBGene00011695 | T10G3.4 | 1.476938651 | 1.118940207 | -0.516482579 | 0.225 | 0.135 | 0.043697257 | 1 |
| WBGene00012100 | sipa-1 | 0.996418598 | 0.638946331 | -0.515723467 | 0.118 | 0.075 | 0.176649618 | 1 |
| WBGene00001086 | dpy-27 | 1.216842595 | 0.85949499 | -0.515543617 | 0.168 | 0.105 | 0.0761348 | 1 |
| WBGene00004487 | rps-18 | 2.697911877 | 2.340885974 | -0.515079499 | 0.646 | 0.511 | 0.001278374 | 1 |
| WBGene00001207 | egl-43 | 0.871633564 | 0.514652629 | -0.515014625 | 0.114 | 0.053 | 0.0362604 | 1 |
| WBGene00012148 | inos-1 | 0.693121146 | 0.336557759 | -0.514412229 | 0.081 | 0.038 | 0.082520005 | 1 |
| WBGene00010006 | F53H2.3 | 0.828094942 | 0.471760126 | -0.514082472 | 0.102 | 0.045 | 0.043461894 | 1 |
| WBGene00002233 | kqt-1 | 1.001790559 | 0.645611447 | -0.513857839 | 0.136 | 0.068 | 0.034314447 | 1 |
| WBGene00001340 | etr-1 | 2.096400457 | 1.740296225 | -0.513749809 | 0.437 | 0.293 | 0.007749283 | 1 |
| WBGene00012019 | dkf-2 | 0.754737829 | 0.399195956 | -0.512938497 | 0.101 | 0.038 | 0.022799937 | 1 |
| WBGene00001037 | dnj-19 | 0.863725118 | 0.509189991 | -0.511486069 | 0.111 | 0.068 | 0.127355357 | 1 |
| WBGene00004488 | rps-19 | 2.741569431 | 2.387388547 | -0.510975005 | 0.652 | 0.519 | 0.000400857 | 1 |
| WBGene00219950 | F46A8.13 | 3.031381843 | 2.677385637 | -0.510708571 | 0.492 | 0.459 | 0.396108997 | 1 |
| WBGene00003150 | mbk-2 | 1.785546329 | 1.432491975 | -0.509349765 | 0.342 | 0.256 | 0.050075259 | 1 |
| WBGene00019667 | spe-49 | 0.479124735 | 0.126707535 | -0.508430547 | 0.054 | 0.015 | 0.054047549 | 1 |
| WBGene00021068 | W06H8.6 | 0.917123798 | 0.56503872 | -0.507951396 | 0.115 | 0.06 | 0.065893842 | 1 |
| WBGene00017900 | F28E10.1 | 0.748574312 | 0.396661318 | -0.50770313 | 0.084 | 0.038 | 0.06945096 | 1 |
| WBGene00009278 | npr-33 | 0.634473751 | 0.282614684 | -0.507625331 | 0.064 | 0.03 | 0.125200566 | 1 |
| WBGene00020190 | guk-1 | 0.678338591 | 0.326827261 | -0.507123654 | 0.063 | 0.03 | 0.135865438 | 1 |
| WBGene00003630 | nhr-40 | 0.58545926 | 0.235396698 | -0.505033522 | 0.067 | 0.023 | 0.048323971 | 1 |
| WBGene00011069 | ceh-62 | 0.639135894 | 0.289501428 | -0.50441591 | 0.07 | 0.03 | 0.082126731 | 1 |
| WBGene00008505 | F01G4.6 | 1.202928282 | 0.853608941 | -0.50396128 | 0.156 | 0.113 | 0.212082772 | 1 |
| WBGene00000280 | cah-2 | 0.609576624 | 0.260425102 | -0.503719169 | 0.068 | 0.03 | 0.096229946 | 1 |
| WBGene00003618 | nhr-19 | 0.762186933 | 0.413501252 | -0.503047103 | 0.089 | 0.038 | 0.049438085 | 1 |
| WBGene00006439 | ant-1.1 | 2.932851771 | 2.584525029 | -0.502529262 | 0.695 | 0.609 | 0.001361104 | 1 |
| WBGene00004409 | rla-1 | 2.616849103 | 2.268750652 | -0.502199909 | 0.603 | 0.466 | 0.006975615 | 1 |
| WBGene00010480 | tasp-1 | 1.573711937 | 1.225644914 | -0.502154568 | 0.278 | 0.195 | 0.059693153 | 1 |
| WBGene00019536 | rpi-2 | 0.593383495 | 0.246398623 | -0.500593354 | 0.063 | 0.03 | 0.130908801 | 1 |
| WBGene00001560 | gei-3 | 0.653269366 | 0.306423954 | -0.500392156 | 0.063 | 0.03 | 0.137773088 | 1 |
| WBGene00194851 | K07C10.3 | 0.710083949 | 0.363698109 | -0.499729134 | 0.076 | 0.038 | 0.11186671 | 1 |
| WBGene00009126 | pyk-1 | 1.007511112 | 0.661509026 | -0.499175494 | 0.137 | 0.083 | 0.087123916 | 1 |
| WBGene00013242 | Y56A3A.30 | 0.542820655 | 0.196893075 | -0.499068005 | 0.056 | 0.023 | 0.107749518 | 1 |
| WBGene00015901 | nhr-158 | 2.163750328 | 1.818058272 | -0.498728216 | 0.458 | 0.286 | 0.002281837 | 1 |
| WBGene00006650 | tts-1 | 1.00569369 | 0.660120367 | -0.498556919 | 0.059 | 0.053 | 0.757422434 | 1 |
| WBGene00002295 | let-19 | 0.561724644 | 0.216276164 | -0.498376809 | 0.063 | 0.023 | 0.061186673 | 1 |
| WBGene00021928 | Y55F3AM.11 | 1.088471712 | 0.743151635 | -0.498191564 | 0.149 | 0.105 | 0.180295566 | 1 |
| WBGene00009440 | idhg-1 | 0.590534585 | 0.246366341 | -0.496529819 | 0.08 | 0.03 | 0.039916847 | 1 |
| WBGene00012766 | Y41E3.8 | 0.637102076 | 0.293144138 | -0.496226412 | 0.082 | 0.03 | 0.037648091 | 1 |
| WBGene00009142 | F26A3.4 | 1.573981511 | 1.230457942 | -0.49559975 | 0.288 | 0.18 | 0.019953145 | 1 |
| WBGene00045336 | R11D1.12 | 2.552681161 | 2.209791191 | -0.49468566 | 0.49 | 0.361 | 0.008614636 | 1 |
| WBGene00000914 | daf-19 | 0.751720284 | 0.40905604 | -0.494360006 | 0.072 | 0.038 | 0.146711884 | 1 |
| WBGene00001510 | fzr-1 | 0.919585573 | 0.577635589 | -0.493329546 | 0.129 | 0.068 | 0.049498308 | 1 |
| WBGene00020181 | T02H6.11 | 0.802410224 | 0.460624419 | -0.493092685 | 0.099 | 0.053 | 0.098339491 | 1 |
| WBGene00007327 | C05C9.3 | 0.932800721 | 0.591244185 | -0.492761919 | 0.113 | 0.068 | 0.133522119 | 1 |
| WBGene00249815 | F20D12.12 | 0.565202758 | 0.224954225 | -0.490874871 | 0.062 | 0.023 | 0.070074071 | 1 |
| WBGene00004454 | rpl-36.A | 2.6566701 | 2.317227975 | -0.489711471 | 0.618 | 0.534 | 0.019854948 | 1 |
| WBGene00000906 | daf-10 | 0.977246899 | 0.638270597 | -0.48903943 | 0.119 | 0.06 | 0.05226983 | 1 |
| WBGene00000492 | che-13 | 0.705728811 | 0.368005387 | -0.487231908 | 0.082 | 0.038 | 0.077993545 | 1 |
| WBGene00001708 | grk-1 | 0.46263363 | 0.125664295 | -0.486143988 | 0.05 | 0.015 | 0.069863101 | 1 |
| WBGene00003830 | num-1 | 1.096016978 | 0.759433497 | -0.485587319 | 0.168 | 0.083 | 0.017687749 | 1 |
| WBGene00004493 | rps-24 | 2.34497538 | 2.009002663 | -0.484706172 | 0.53 | 0.436 | 0.01821122 | 1 |
| WBGene00004444 | rpl-30 | 2.020973835 | 1.685020191 | -0.484678657 | 0.417 | 0.278 | 0.005750893 | 1 |
| WBGene00019847 | R03G5.6 | 0.604483976 | 0.268765692 | -0.484339103 | 0.071 | 0.023 | 0.036950495 | 1 |

| | | | | | | | | |
|---|---|---|---|---|---|---|---|---|
| WBGene00022592 | klu-2 | 0.949067134 | 0.613472805 | -0.484160274 | 0.121 | 0.068 | 0.08419893 | 1 |
| WBGene00016315 | C32D5.7 | 1.946402008 | 1.611274619 | -0.483486623 | 0.397 | 0.286 | 0.021328124 | 1 |
| WBGene00018349 | F42C5.9 | 0.70610749 | 0.37109755 | -0.483317178 | 0.071 | 0.038 | 0.147744561 | 1 |
| WBGene00005359 | srh-142 | 0.799401179 | 0.465011239 | -0.482422708 | 0.109 | 0.053 | 0.052528576 | 1 |
| WBGene00012187 | rpb-11 | 0.451364131 | 0.117648911 | -0.481449293 | 0.054 | 0.015 | 0.053752368 | 1 |
| WBGene00004736 | sca-1 | 0.512717322 | 0.179219112 | -0.481136214 | 0.054 | 0.015 | 0.056355851 | 1 |
| WBGene00000793 | crh-1 | 1.830000977 | 1.496670483 | -0.48089425 | 0.362 | 0.241 | 0.020227304 | 1 |
| WBGene00009120 | F25H2.6 | 1.743784932 | 1.410706648 | -0.48053039 | 0.35 | 0.233 | 0.018277005 | 1 |
| WBGene00006752 | unc-13 | 1.272367238 | 0.93935007 | -0.480442216 | 0.213 | 0.135 | 0.065621094 | 1 |
| WBGene00006725 | ubl-1 | 2.620928268 | 2.287915331 | -0.480436114 | 0.621 | 0.481 | 0.003961005 | 1 |
| WBGene00012340 | dct-15 | 0.771703595 | 0.439281715 | -0.479583397 | 0.115 | 0.045 | 0.017719253 | 1 |
| WBGene00019554 | K09C4.10 | 0.632510382 | 0.300693703 | -0.478710277 | 0.075 | 0.03 | 0.062623558 | 1 |
| WBGene00020208 | T04C4.1 | 1.040284208 | 0.708575225 | -0.478554905 | 0.131 | 0.083 | 0.120875766 | 1 |
| WBGene00006068 | sto-6 | 0.746801698 | 0.415092845 | -0.478554716 | 0.098 | 0.045 | 0.052827198 | 1 |
| WBGene00004486 | rps-17 | 2.819999126 | 2.488489666 | -0.478267054 | 0.689 | 0.504 | 0.000171715 | 1 |
| WBGene00008977 | iglr-1 | 0.736209464 | 0.404743615 | -0.478204137 | 0.077 | 0.038 | 0.107819815 | 1 |
| WBGene00000833 | cts-1 | 0.864527489 | 0.534316206 | -0.476394179 | 0.12 | 0.06 | 0.050791634 | 1 |
| WBGene00017790 | F25E5.9 | 0.540437827 | 0.210362327 | -0.476198286 | 0.052 | 0.015 | 0.060685745 | 1 |
| WBGene00003889 | osm-9 | 0.674096282 | 0.344903816 | -0.474924338 | 0.073 | 0.045 | 0.22164371 | 1 |
| WBGene00007443 | pfd-1 | 0.818040741 | 0.48900895 | -0.474692534 | 0.098 | 0.045 | 0.053117246 | 1 |
| WBGene00015251 | B0546.4 | 0.958379794 | 0.629370721 | -0.474659759 | 0.122 | 0.068 | 0.078159159 | 1 |
| WBGene00004442 | rpl-28 | 2.107988448 | 1.779097638 | -0.47448914 | 0.437 | 0.338 | 0.024034396 | 1 |
| WBGene00020146 | got-1.2 | 0.749488171 | 0.421541846 | -0.473126536 | 0.098 | 0.045 | 0.055977531 | 1 |
| WBGene00011980 | T24B8.7 | 0.64698708 | 0.320125497 | -0.471561585 | 0.057 | 0.038 | 0.36012324 | 1 |
| WBGene00008518 | irld-2 | 0.620126516 | 0.294458403 | -0.469839772 | 0.078 | 0.03 | 0.050559508 | 1 |
| WBGene00044927 | F32B5.9 | 0.783275595 | 0.457863578 | -0.469470303 | 0.073 | 0.053 | 0.383823138 | 1 |
| WBGene00001226 | eif-3.C | 0.561086782 | 0.236536409 | -0.468227214 | 0.068 | 0.03 | 0.093469592 | 1 |
| WBGene00022603 | gck-2 | 0.552679358 | 0.230027439 | -0.465488323 | 0.052 | 0.03 | 0.262031405 | 1 |
| WBGene00011955 | cdka-1 | 0.809105395 | 0.486871828 | -0.464884769 | 0.103 | 0.06 | 0.136422469 | 1 |
| WBGene00001168 | eef-1A.1 | 2.680205226 | 2.360449784 | -0.46130959 | 0.643 | 0.534 | 0.01336746 | 1 |
| WBGene00006651 | tts-2 | 2.174729818 | 1.855782545 | -0.460143648 | 0.428 | 0.338 | 0.093493121 | 1 |
| WBGene00000230 | atp-3 | 0.878602037 | 0.560815329 | -0.458469308 | 0.128 | 0.053 | 0.016086037 | 1 |
| WBGene00003916 | par-1 | 0.63622189 | 0.320270602 | -0.455821356 | 0.072 | 0.038 | 0.138794256 | 1 |
| WBGene00021838 | Y54E10BL.3 | 0.518926377 | 0.203237353 | -0.455442989 | 0.058 | 0.023 | 0.092372055 | 1 |
| WBGene00016938 | C54H2.4 | 0.615195454 | 0.300695295 | -0.45372782 | 0.065 | 0.023 | 0.058174719 | 1 |
| WBGene00022951 | C30E1.9 | 5.149273983 | 4.835093754 | -0.453266258 | 0.918 | 0.992 | 1.04E-06 | 0.006646548 |
| WBGene00015285 | gmeb-1 | 0.629238684 | 0.315169073 | -0.45310667 | 0.083 | 0.03 | 0.036748863 | 1 |
| WBGene00006476 | rhgf-2 | 0.833785822 | 0.519842239 | -0.452924851 | 0.094 | 0.045 | 0.069399242 | 1 |
| WBGene00003002 | lin-13 | 0.647783298 | 0.333920678 | -0.452808045 | 0.067 | 0.045 | 0.325008702 | 1 |
| WBGene00004497 | rps-28 | 3.040046447 | 2.726391835 | -0.452507952 | 0.752 | 0.699 | 0.013765204 | 1 |
| WBGene00010960 | atp-6 | 3.163901138 | 2.851042655 | -0.451359382 | 0.651 | 0.594 | 0.03726551 | 1 |
| WBGene00008957 | glb-13 | 1.478458032 | 1.165638538 | -0.451303132 | 0.244 | 0.158 | 0.045046901 | 1 |
| WBGene00007042 | pbrm-1 | 0.662383681 | 0.350371939 | -0.450137793 | 0.089 | 0.038 | 0.047886519 | 1 |
| WBGene00000481 | cha-1 | 1.044319496 | 0.732941376 | -0.44922367 | 0.156 | 0.098 | 0.095712475 | 1 |
| WBGene00009337 | uig-1 | 0.549047478 | 0.23853311 | -0.447977538 | 0.064 | 0.03 | 0.121876612 | 1 |
| WBGene00001853 | hil-2 | 1.990019244 | 1.679613895 | -0.447820257 | 0.352 | 0.241 | 0.027409197 | 1 |
| WBGene00021959 | Y57E12AL.6 | 0.643636496 | 0.333491344 | -0.447444872 | 0.085 | 0.038 | 0.064469957 | 1 |
| WBGene00004489 | rps-20 | 2.736832947 | 2.426721459 | -0.447396306 | 0.686 | 0.586 | 0.002630468 | 1 |
| WBGene00012221 | W03C9.5 | 1.274300438 | 0.964202591 | -0.447376626 | 0.197 | 0.135 | 0.127343244 | 1 |
| WBGene00004445 | rpl-31 | 2.944251819 | 2.634628651 | -0.446691809 | 0.695 | 0.632 | 0.005787629 | 1 |
| WBGene00044068 | syd-9 | 1.403796872 | 1.094317455 | -0.44648442 | 0.22 | 0.128 | 0.029892757 | 1 |
| WBGene00016603 | met-1 | 1.083415717 | 0.774238898 | -0.446047864 | 0.147 | 0.083 | 0.056444038 | 1 |
| WBGene00009050 | F22D6.2 | 2.322334423 | 2.013349991 | -0.445770308 | 0.535 | 0.421 | 0.032662418 | 1 |
| WBGene00013415 | ilcr-1 | 0.558245596 | 0.250301738 | -0.444269076 | 0.067 | 0.023 | 0.048808352 | 1 |
| WBGene00006911 | vha-2 | 0.935696153 | 0.627835065 | -0.444149666 | 0.125 | 0.083 | 0.177842453 | 1 |
| WBGene00018188 | F38E9.6 | 2.052477565 | 1.744999477 | -0.443597112 | 0.404 | 0.286 | 0.014653165 | 1 |
| WBGene00020868 | eif-1 | 1.070785921 | 0.763946745 | -0.442675357 | 0.135 | 0.105 | 0.35125852 | 1 |
| WBGene00001167 | eef-2 | 2.37090659 | 2.064768394 | -0.441664056 | 0.541 | 0.398 | 0.016401834 | 1 |
| WBGene00010838 | M03C11.1 | 0.550663618 | 0.244717188 | -0.441387396 | 0.06 | 0.023 | 0.081900375 | 1 |
| WBGene00044638 | F23A7.8 | 1.446712761 | 1.142923022 | -0.438275951 | 0.083 | 0.053 | 0.232578307 | 1 |
| WBGene00004469 | rps-0 | 2.43278 | 2.129414724 | -0.437663579 | 0.566 | 0.466 | 0.030401625 | 1 |
| WBGene00017925 | F29B9.11 | 1.034131968 | 0.731393224 | -0.436759684 | 0.157 | 0.09 | 0.053041296 | 1 |
| WBGene00006384 | taf-3 | 0.461234381 | 0.158731921 | -0.436418798 | 0.055 | 0.015 | 0.050091602 | 1 |
| WBGene00001672 | gpa-10 | 0.892497768 | 0.590637311 | -0.435492584 | 0.123 | 0.06 | 0.043159794 | 1 |

| | | | | | | | | |
|---|---|---|---|---|---|---|---|---|
| WBGene00044144 | M153.4 | 1.405413733 | 1.103968738 | -0.434893199 | 0.247 | 0.128 | 0.007691526 | 1 |
| WBGene00000553 | cmk-1 | 0.748652011 | 0.447490346 | -0.43448444 | 0.095 | 0.053 | 0.11614241 | 1 |
| WBGene00004699 | rsp-2 | 1.074602325 | 0.773603624 | -0.434249334 | 0.145 | 0.09 | 0.11452179 | 1 |
| WBGene00022400 | rpb-9 | 0.529729444 | 0.228793274 | -0.434159119 | 0.055 | 0.023 | 0.119207746 | 1 |
| WBGene00001187 | egl-19 | 0.615925523 | 0.315110842 | -0.433983849 | 0.057 | 0.03 | 0.206310212 | 1 |
| WBGene00021047 | zfp-3 | 0.888380645 | 0.587691459 | -0.433802797 | 0.119 | 0.075 | 0.156694067 | 1 |
| WBGene00006519 | cox-6A | 0.847648274 | 0.54723681 | -0.433402129 | 0.111 | 0.06 | 0.083039112 | 1 |
| WBGene00022170 | cox-6B | 0.968237351 | 0.669811626 | -0.430537313 | 0.132 | 0.09 | 0.184646568 | 1 |
| WBGene00004439 | rpl-25.2 | 1.399498994 | 1.101123389 | -0.430465006 | 0.243 | 0.15 | 0.033298552 | 1 |
| WBGene00000898 | daf-2 | 0.592642154 | 0.294503175 | -0.430123626 | 0.066 | 0.03 | 0.109256124 | 1 |
| WBGene00006067 | sto-5 | 0.606188042 | 0.309136033 | -0.428555461 | 0.07 | 0.015 | 0.015457627 | 1 |
| WBGene00017894 | F28B12.1 | 2.006010341 | 1.708958427 | -0.428555323 | 0.431 | 0.301 | 0.009468974 | 1 |
| WBGene00015063 | B0228.6 | 1.016132485 | 0.719462311 | -0.428004589 | 0.119 | 0.083 | 0.263304209 | 1 |
| WBGene00044650 | Y71G10AR.4 | 1.120543396 | 0.824371872 | -0.427285189 | 0.158 | 0.098 | 0.084199901 | 1 |
| WBGene00016650 | ubr-4 | 1.103748805 | 0.807975183 | -0.426711138 | 0.132 | 0.105 | 0.397441244 | 1 |
| WBGene00012641 | epg-6 | 0.660444142 | 0.364950329 | -0.426307459 | 0.081 | 0.038 | 0.081726068 | 1 |
| WBGene00004916 | snr-3 | 0.696635502 | 0.401441615 | -0.425874757 | 0.091 | 0.045 | 0.084822971 | 1 |
| WBGene00018395 | mtch-1 | 1.092884943 | 0.797914181 | -0.425552855 | 0.152 | 0.098 | 0.12156681 | 1 |
| WBGene00119203 | T04A8.18 | 0.959112209 | 0.664817292 | -0.424577817 | 0.131 | 0.09 | 0.213009191 | 1 |
| WBGene00012615 | dct-16 | 0.805993486 | 0.511983166 | -0.424167231 | 0.056 | 0.053 | 0.889731098 | 1 |
| WBGene00020827 | T26A8.4 | 0.540786397 | 0.246776403 | -0.424166761 | 0.059 | 0.03 | 0.172424263 | 1 |
| WBGene00004436 | rpl-24.1 | 3.135482056 | 2.842334521 | -0.422922495 | 0.772 | 0.684 | 0.001295844 | 1 |
| WBGene00002981 | lgg-2 | 2.18553518 | 1.893695366 | -0.421035853 | 0.474 | 0.346 | 0.013062181 | 1 |
| WBGene00003827 | ntl-4 | 0.802043565 | 0.510241895 | -0.420980822 | 0.083 | 0.053 | 0.250298689 | 1 |
| WBGene00044134 | D1025.10 | 0.512782562 | 0.221056532 | -0.420871697 | 0.059 | 0.023 | 0.088655494 | 1 |
| WBGene00016998 | D1005.4 | 1.792326364 | 1.500833009 | -0.420536017 | 0.306 | 0.241 | 0.204482834 | 1 |
| WBGene00016532 | C39D10.5 | 0.943648794 | 0.65297012 | -0.419360682 | 0.113 | 0.068 | 0.128486382 | 1 |
| WBGene00003948 | pbs-2 | 0.671334924 | 0.380944099 | -0.418945404 | 0.075 | 0.038 | 0.124832615 | 1 |
| WBGene00008256 | glb-12 | 0.552154602 | 0.262123037 | -0.418427099 | 0.062 | 0.015 | 0.030173682 | 1 |
| WBGene00015776 | dct-1 | 1.836389278 | 1.54783826 | -0.416291123 | 0.369 | 0.233 | 0.007520343 | 1 |
| WBGene00003634 | nhr-44 | 1.938770363 | 1.650486602 | -0.415905553 | 0.383 | 0.271 | 0.042867827 | 1 |
| WBGene00003754 | nlp-16 | 1.058832155 | 0.770593563 | -0.415840387 | 0.14 | 0.075 | 0.048778242 | 1 |
| WBGene00022240 | shl-1 | 0.799932608 | 0.511772204 | -0.415727585 | 0.104 | 0.053 | 0.06925029 | 1 |
| WBGene00018606 | F48E3.6 | 1.452494408 | 1.164529463 | -0.415445598 | 0.264 | 0.188 | 0.091501164 | 1 |
| WBGene00003678 | nhr-88 | 1.015099907 | 0.727205241 | -0.415344207 | 0.144 | 0.09 | 0.116843966 | 1 |
| WBGene00006728 | ubq-2 | 2.944962575 | 2.657114725 | -0.415276666 | 0.719 | 0.647 | 0.010670027 | 1 |
| WBGene00019286 | K01A12.2 | 1.396274118 | 1.108837711 | -0.41468308 | 0.233 | 0.15 | 0.056566769 | 1 |
| WBGene00003947 | pbs-1 | 0.537098002 | 0.250325404 | -0.413725405 | 0.052 | 0.03 | 0.271820096 | 1 |
| WBGene00019243 | fbxb-72 | 0.731562587 | 0.445340414 | -0.41293131 | 0.091 | 0.045 | 0.076746726 | 1 |
| WBGene00008956 | nekl-3 | 1.090733657 | 0.804733294 | -0.412611307 | 0.175 | 0.09 | 0.020604434 | 1 |
| WBGene00044677 | ZK353.10 | 2.155154967 | 1.869277687 | -0.412433735 | 0.456 | 0.308 | 0.003565741 | 1 |
| WBGene00011146 | pde-2 | 1.187006352 | 0.901872995 | -0.41136048 | 0.175 | 0.105 | 0.062133593 | 1 |
| WBGene00006066 | sto-4 | 0.656715683 | 0.372032462 | -0.410711072 | 0.061 | 0.038 | 0.291678356 | 1 |
| WBGene00006603 | tps-2 | 0.933391176 | 0.648766115 | -0.410627163 | 0.129 | 0.075 | 0.091054529 | 1 |
| WBGene00019230 | ttll-11 | 1.115830752 | 0.832317639 | -0.409022963 | 0.154 | 0.113 | 0.240075219 | 1 |
| WBGene00020838 | trak-1 | 1.11982068 | 0.836987355 | -0.408042236 | 0.143 | 0.083 | 0.083739093 | 1 |
| WBGene00008520 | F02D8.1 | 2.037777234 | 1.756072161 | -0.406414512 | 0.4 | 0.316 | 0.142178318 | 1 |
| WBGene00021607 | Y46H3C.7 | 0.700955141 | 0.421065354 | -0.403795607 | 0.077 | 0.045 | 0.200981657 | 1 |
| WBGene00002151 | irk-3 | 1.443128043 | 1.16400867 | -0.402684135 | 0.253 | 0.135 | 0.005576754 | 1 |
| WBGene00001679 | gpb-1 | 1.766901963 | 1.488008658 | -0.402357988 | 0.336 | 0.293 | 0.338666556 | 1 |
| WBGene00004477 | rps-8 | 2.822683613 | 2.544781105 | -0.400928569 | 0.686 | 0.526 | 0.001443569 | 1 |
| WBGene00006513 | rimb-1 | 0.758885244 | 0.481249936 | -0.400543082 | 0.09 | 0.045 | 0.091962837 | 1 |
| WBGene00004476 | rps-7 | 2.702805056 | 2.425213054 | -0.400480604 | 0.626 | 0.571 | 0.063996629 | 1 |
| WBGene00012468 | Y17G7B.17 | 1.131892347 | 0.854425684 | -0.400299779 | 0.178 | 0.113 | 0.086957931 | 1 |
| WBGene00009098 | ehbp-1 | 0.415142443 | 0.137685701 | -0.400285465 | 0.054 | 0.015 | 0.055543822 | 1 |
| WBGene00002717 | let-526 | 0.905000726 | 0.627567365 | -0.400251733 | 0.131 | 0.075 | 0.079769635 | 1 |
| WBGene00018281 | tbc-18 | 0.681284383 | 0.404353108 | -0.399527376 | 0.073 | 0.038 | 0.139640885 | 1 |
| WBGene00000262 | bra-1 | 0.73458295 | 0.458634539 | -0.398109404 | 0.081 | 0.045 | 0.158703242 | 1 |
| WBGene00002190 | kin-2 | 1.371976706 | 1.096316677 | -0.397693357 | 0.209 | 0.158 | 0.218562878 | 1 |
| WBGene00006604 | tra-1 | 0.658953894 | 0.383631938 | -0.39720562 | 0.084 | 0.038 | 0.066864441 | 1 |
| WBGene00007053 | chd-7 | 1.044853246 | 0.769644608 | -0.397042137 | 0.135 | 0.098 | 0.255860241 | 1 |
| WBGene00020397 | pcbd-1 | 0.735042681 | 0.460395537 | -0.396232073 | 0.084 | 0.053 | 0.223645947 | 1 |
| WBGene00017834 | F26F12.3 | 2.421640116 | 2.147281394 | -0.395815967 | 0.508 | 0.429 | 0.13630046 | 1 |
| WBGene00004951 | spc-1 | 0.974233221 | 0.700067032 | -0.395538202 | 0.116 | 0.075 | 0.195874038 | 1 |

| | | | | | | | | |
|---|---|---|---|---|---|---|---|---|
| WBGene00001042 | dnj-24 | 0.627583366 | 0.354402124 | -0.394117223 | 0.072 | 0.038 | 0.147465634 | 1 |
| WBGene00004418 | rpl-7 | 2.639630781 | 2.366957327 | -0.393384639 | 0.625 | 0.549 | 0.027729872 | 1 |
| WBGene00006601 | tpi-1 | 0.540384805 | 0.268305767 | -0.392527079 | 0.064 | 0.03 | 0.129137167 | 1 |
| WBGene00004411 | rpc-1 | 0.687902727 | 0.416299494 | -0.391840636 | 0.085 | 0.045 | 0.12239951 | 1 |
| WBGene00021346 | Y37E3.1 | 0.532499259 | 0.261508838 | -0.390956536 | 0.059 | 0.023 | 0.088655489 | 1 |
| WBGene00004433 | rpl-21 | 2.587320551 | 2.316460837 | -0.390767966 | 0.599 | 0.504 | 0.031391574 | 1 |
| WBGene00016016 | C23G10.10 | 0.875726181 | 0.605369235 | -0.390042626 | 0.103 | 0.068 | 0.223268195 | 1 |
| WBGene00004375 | rme-4 | 0.580797244 | 0.310559378 | -0.389870829 | 0.071 | 0.023 | 0.037075377 | 1 |
| WBGene00004446 | rpl-32 | 2.923792027 | 2.653674424 | -0.389697327 | 0.71 | 0.579 | 0.005134204 | 1 |
| WBGene00000380 | cct-5 | 1.674742446 | 1.404838245 | -0.389389452 | 0.312 | 0.188 | 0.00910856 | 1 |
| WBGene00015716 | sox-4 | 0.562960368 | 0.293311722 | -0.389020763 | 0.058 | 0.03 | 0.193921881 | 1 |
| WBGene00016653 | ssb-1 | 0.527917344 | 0.258351739 | -0.388900962 | 0.057 | 0.023 | 0.104786643 | 1 |
| WBGene00021254 | Y22D7AL.16 | 0.992726837 | 0.723893577 | -0.387844411 | 0.141 | 0.098 | 0.184136868 | 1 |
| WBGene00003924 | pas-3 | 0.653110596 | 0.384352964 | -0.387735303 | 0.078 | 0.038 | 0.104898889 | 1 |
| WBGene00019947 | htz-1 | 0.742470995 | 0.473979342 | -0.387351578 | 0.09 | 0.06 | 0.259067885 | 1 |
| WBGene00004448 | rpl-34 | 2.355978268 | 2.088614916 | -0.385723781 | 0.523 | 0.451 | 0.096611518 | 1 |
| WBGene00003695 | nhr-105 | 2.42109753 | 2.154221324 | -0.385020978 | 0.542 | 0.451 | 0.021154449 | 1 |
| WBGene00018849 | spt-16 | 0.668971444 | 0.402218222 | -0.38484355 | 0.083 | 0.045 | 0.132983474 | 1 |
| WBGene00001169 | eef-1A.2 | 2.243398752 | 1.976876264 | -0.384510672 | 0.485 | 0.391 | 0.105945543 | 1 |
| WBGene00016981 | rpn-13 | 0.691901718 | 0.425463166 | -0.384389578 | 0.084 | 0.045 | 0.12508437 | 1 |
| WBGene00016212 | C29F5.3 | 0.79022979 | 0.52528953 | -0.382227998 | 0.1 | 0.053 | 0.096524041 | 1 |
| WBGene00019829 | R02E12.5 | 1.081093828 | 0.816345709 | -0.381950797 | 0.139 | 0.098 | 0.230265659 | 1 |
| WBGene00018985 | F56F10.2 | 0.491703048 | 0.22748067 | -0.381192315 | 0.055 | 0.023 | 0.120687637 | 1 |
| WBGene00011517 | cgt-1 | 0.773479841 | 0.50928429 | -0.381153611 | 0.097 | 0.045 | 0.063299194 | 1 |
| WBGene00077486 | sls-1.9 | 0.673162621 | 0.409341494 | -0.380613433 | 0.064 | 0.038 | 0.251186664 | 1 |
| WBGene00012203 | rga-1 | 0.503472448 | 0.239869007 | -0.380299377 | 0.056 | 0.03 | 0.215030499 | 1 |
| WBGene00020151 | T01G6.1 | 0.566109843 | 0.302575797 | -0.380199261 | 0.059 | 0.023 | 0.088514857 | 1 |
| WBGene00017615 | srsx-32 | 1.796830084 | 1.533508498 | -0.379892746 | 0.327 | 0.226 | 0.045076409 | 1 |
| WBGene00002202 | kin-19 | 1.074396774 | 0.811249183 | -0.379641724 | 0.133 | 0.105 | 0.397341937 | 1 |
| WBGene00006462 | svh-5 | 1.253105137 | 0.990090362 | -0.379450111 | 0.178 | 0.09 | 0.020036993 | 1 |
| WBGene00045056 | F13H10.9 | 0.923417497 | 0.661219807 | -0.378271307 | 0.121 | 0.083 | 0.21204921 | 1 |
| WBGene00004368 | ric-19 | 0.654861196 | 0.39309584 | -0.37764758 | 0.078 | 0.038 | 0.105461852 | 1 |
| WBGene00004421 | rpl-10 | 2.967204642 | 2.705947338 | -0.376914617 | 0.729 | 0.662 | 0.010654014 | 1 |
| WBGene00017711 | F22E5.13 | 0.72481969 | 0.463566777 | -0.376908282 | 0.081 | 0.053 | 0.275934338 | 1 |
| WBGene00013711 | Y106G6G.2 | 0.645417097 | 0.385058452 | -0.375618126 | 0.082 | 0.053 | 0.252977466 | 1 |
| WBGene00016405 | hecd-1 | 0.649736155 | 0.390336202 | -0.374235026 | 0.08 | 0.045 | 0.165716599 | 1 |
| WBGene00004416 | rpl-5 | 2.477467117 | 2.218077538 | -0.374220059 | 0.588 | 0.444 | 0.017099234 | 1 |
| WBGene00000284 | cah-6 | 0.570110837 | 0.310867338 | -0.37400931 | 0.052 | 0.038 | 0.469600208 | 1 |
| WBGene00009004 | pfd-6 | 0.645329125 | 0.38618498 | -0.373865973 | 0.084 | 0.03 | 0.034650814 | 1 |
| WBGene00022401 | Y97E10AR.6 | 0.708110263 | 0.449242468 | -0.373467283 | 0.098 | 0.053 | 0.099643018 | 1 |
| WBGene00003079 | lsm-5 | 0.460483928 | 0.20209433 | -0.372777391 | 0.052 | 0.015 | 0.061901097 | 1 |
| WBGene00015547 | ain-1 | 0.687618839 | 0.429338979 | -0.372619074 | 0.066 | 0.038 | 0.214152561 | 1 |
| WBGene00021369 | siah-1 | 0.929860597 | 0.67166373 | -0.37249934 | 0.132 | 0.083 | 0.130169666 | 1 |
| WBGene00018160 | odr-8 | 0.863832683 | 0.605664158 | -0.372458451 | 0.121 | 0.068 | 0.085990412 | 1 |
| WBGene00206380 | C42D4.18 | 1.554602819 | 1.296452009 | -0.372432894 | 0.271 | 0.173 | 0.038919683 | 1 |
| WBGene00000854 | cux-7 | 0.635322471 | 0.377349665 | -0.372176087 | 0.087 | 0.038 | 0.056540337 | 1 |
| WBGene00004438 | rpl-25.1 | 1.420664838 | 1.163883275 | -0.370457486 | 0.253 | 0.165 | 0.043683055 | 1 |
| WBGene00018078 | F35H12.6 | 0.740030055 | 0.48382265 | -0.369629153 | 0.101 | 0.045 | 0.049970206 | 1 |
| WBGene00013317 | Y57G11C.22 | 0.506618423 | 0.250788371 | -0.369084747 | 0.057 | 0.023 | 0.104541634 | 1 |
| WBGene00006791 | unc-57 | 0.762896948 | 0.507234523 | -0.368842912 | 0.097 | 0.053 | 0.120346619 | 1 |
| WBGene00003952 | pbs-6 | 0.65687643 | 0.401573029 | -0.368324951 | 0.08 | 0.03 | 0.047121194 | 1 |
| WBGene00015755 | C14B9.10 | 0.920577598 | 0.665747433 | -0.367642216 | 0.126 | 0.083 | 0.183044269 | 1 |
| WBGene00001574 | gei-17 | 0.586486491 | 0.33169881 | -0.367580924 | 0.066 | 0.03 | 0.115336637 | 1 |
| WBGene00004463 | rpn-7 | 0.620113537 | 0.365411695 | -0.367457085 | 0.076 | 0.045 | 0.202151582 | 1 |
| WBGene00017568 | F18E9.1 | 0.60622145 | 0.351642209 | -0.367280209 | 0.078 | 0.03 | 0.049023644 | 1 |
| WBGene00022579 | ZC262.2 | 0.561149839 | 0.306687033 | -0.367112229 | 0.07 | 0.03 | 0.08224806 | 1 |
| WBGene00008398 | D2005.3 | 0.488522822 | 0.234233137 | -0.366862468 | 0.064 | 0.023 | 0.061648253 | 1 |
| WBGene00017511 | srbc-41 | 2.628058901 | 2.374060156 | -0.366442729 | 0.572 | 0.481 | 0.061781514 | 1 |
| WBGene00006786 | unc-51 | 0.535586569 | 0.282372032 | -0.365311357 | 0.065 | 0.03 | 0.120506629 | 1 |
| WBGene00003172 | mec-8 | 0.517638889 | 0.264905741 | -0.36461686 | 0.052 | 0.03 | 0.273532783 | 1 |
| WBGene00003153 | mca-3 | 1.219061625 | 0.96640736 | -0.364503055 | 0.153 | 0.128 | 0.584753126 | 1 |
| WBGene00003149 | mbk-1 | 1.121930142 | 0.869446189 | -0.364257346 | 0.172 | 0.12 | 0.16227008 | 1 |
| WBGene00004495 | rps-26 | 3.229897145 | 2.977541321 | -0.364072496 | 0.794 | 0.782 | 0.01069875 | 1 |
| WBGene00009374 | F34D10.4 | 1.649510902 | 1.397562629 | -0.363484523 | 0.234 | 0.18 | 0.183356154 | 1 |

| | | | | | | | | |
|---|---|---|---|---|---|---|---|---|
| WBGene00016289 | lntl-1 | 2.851319222 | 2.599821127 | -0.362835054 | 0.694 | 0.534 | 0.00138938 | 1 |
| WBGene00003563 | ncs-1 | 1.06862133 | 0.817489772 | -0.362306254 | 0.162 | 0.09 | 0.050595213 | 1 |
| WBGene00019188 | H11E01.3 | 0.628428831 | 0.377700231 | -0.361724908 | 0.07 | 0.03 | 0.084333122 | 1 |
| WBGene00004185 | pro-1 | 0.711992778 | 0.461323116 | -0.361639879 | 0.08 | 0.038 | 0.092449916 | 1 |
| WBGene00001502 | ftt-2 | 2.13215823 | 1.881595517 | -0.361485583 | 0.431 | 0.353 | 0.13599978 | 1 |
| WBGene00010898 | nphp-1 | 0.712189039 | 0.461896066 | -0.361096432 | 0.092 | 0.06 | 0.222225866 | 1 |
| WBGene00004822 | skr-16 | 0.469874744 | 0.21962635 | -0.361032117 | 0.058 | 0.023 | 0.09792639 | 1 |
| WBGene00000244 | bbs-8 | 1.797623983 | 1.547595891 | -0.360714289 | 0.338 | 0.248 | 0.09032006 | 1 |
| WBGene00001999 | hrp-1 | 1.654295658 | 1.405373667 | -0.359118523 | 0.333 | 0.233 | 0.042333372 | 1 |
| WBGene00003926 | pas-5 | 0.603560921 | 0.355346345 | -0.358097939 | 0.07 | 0.038 | 0.161080219 | 1 |
| WBGene00009483 | F36G3.2 | 0.551516972 | 0.304150781 | -0.356873976 | 0.068 | 0.015 | 0.01917233 | 1 |
| WBGene00003003 | lin-14 | 0.662836055 | 0.416245496 | -0.355754976 | 0.081 | 0.045 | 0.162716747 | 1 |
| WBGene00017671 | pgal-1 | 2.456127207 | 2.209913563 | -0.355211203 | 0.538 | 0.444 | 0.084077402 | 1 |
| WBGene00007180 | B0457.6 | 1.841495252 | 1.595636065 | -0.35469983 | 0.34 | 0.271 | 0.172711795 | 1 |
| WBGene00006479 | tmbi-4 | 0.5207886 | 0.275499096 | -0.353877951 | 0.066 | 0.03 | 0.116073985 | 1 |
| WBGene00006331 | sup-26 | 0.85135116 | 0.606587232 | -0.353119705 | 0.076 | 0.053 | 0.354376295 | 1 |
| WBGene00013548 | lury-1 | 1.321233282 | 1.076554025 | -0.352997552 | 0.188 | 0.15 | 0.314981758 | 1 |
| WBGene00006975 | zfp-1 | 0.835761544 | 0.591306739 | -0.352673736 | 0.124 | 0.075 | 0.128909086 | 1 |
| WBGene00006578 | tli-1 | 0.748497332 | 0.504182022 | -0.352472486 | 0.072 | 0.053 | 0.42230181 | 1 |
| WBGene00000188 | arl-3 | 2.380591805 | 2.136320373 | -0.352409183 | 0.531 | 0.459 | 0.126534255 | 1 |
| WBGene00020777 | T24E12.10 | 1.666022689 | 1.421756111 | -0.35240218 | 0.259 | 0.211 | 0.318977917 | 1 |
| WBGene00003961 | pct-1 | 0.97952111 | 0.735978656 | -0.351357491 | 0.144 | 0.075 | 0.041689443 | 1 |
| WBGene00002005 | hsp-1 | 2.333397557 | 2.090205558 | -0.350851892 | 0.511 | 0.383 | 0.06583254 | 1 |
| WBGene00001664 | gpa-2 | 1.734073757 | 1.49092103 | -0.350795234 | 0.345 | 0.203 | 0.00456069 | 1 |
| WBGene00004428 | rpl-16 | 2.525561509 | 2.283114044 | -0.349777755 | 0.59 | 0.474 | 0.012431266 | 1 |
| WBGene00004268 | rab-5 | 0.948873745 | 0.706749766 | -0.349311065 | 0.13 | 0.083 | 0.145408164 | 1 |
| WBGene00012885 | iscu-1 | 0.690584855 | 0.44854913 | -0.349183741 | 0.089 | 0.045 | 0.098888428 | 1 |
| WBGene00004053 | tank-1 | 0.57033037 | 0.328548621 | -0.348817331 | 0.054 | 0.038 | 0.434408751 | 1 |
| WBGene00020917 | W01B11.6 | 1.901811768 | 1.661445046 | -0.346775878 | 0.367 | 0.278 | 0.076040942 | 1 |
| WBGene00003357 | mir-264 | 0.803482732 | 0.563550804 | -0.346148602 | 0.11 | 0.06 | 0.086483861 | 1 |
| WBGene00008446 | E01G4.3 | 1.683792751 | 1.444601375 | -0.345080212 | 0.255 | 0.195 | 0.219314849 | 1 |
| WBGene00005640 | srn-1 | 2.008977503 | 1.769828278 | -0.345019401 | 0.411 | 0.293 | 0.023983537 | 1 |
| WBGene00004420 | rpl-9 | 2.460594669 | 2.221581717 | -0.3448228 | 0.582 | 0.481 | 0.085855782 | 1 |
| WBGene00013577 | Y76A2B.5 | 0.972549836 | 0.73402357 | -0.344120661 | 0.132 | 0.098 | 0.274112083 | 1 |
| WBGene00002344 | let-70 | 1.971495438 | 1.733494288 | -0.343363079 | 0.418 | 0.323 | 0.085537097 | 1 |
| WBGene00004474 | rps-5 | 2.762283341 | 2.524838647 | -0.342560282 | 0.672 | 0.556 | 0.020570495 | 1 |
| WBGene00001488 | frm-1 | 0.997612594 | 0.760847588 | -0.341579699 | 0.13 | 0.09 | 0.242102777 | 1 |
| WBGene00003401 | mpk-1 | 1.378721378 | 1.142604931 | -0.340644027 | 0.219 | 0.165 | 0.195723524 | 1 |
| WBGene00007235 | ahsa-1 | 0.729024664 | 0.49300086 | -0.340510371 | 0.093 | 0.045 | 0.07968588 | 1 |
| WBGene00044386 | C27A2.7 | 0.451725396 | 0.215990334 | -0.340093805 | 0.055 | 0.023 | 0.118290049 | 1 |
| WBGene00018727 | jbts-14 | 0.688563066 | 0.452852205 | -0.340058892 | 0.092 | 0.053 | 0.14089371 | 1 |
| WBGene00004347 | rgs-4 | 0.613247063 | 0.378909479 | -0.33807767 | 0.064 | 0.038 | 0.24737977 | 1 |
| WBGene00004473 | rps-4 | 2.696715998 | 2.463088531 | -0.337053188 | 0.644 | 0.534 | 0.023126572 | 1 |
| WBGene00000396 | cdh-4 | 0.620281379 | 0.386723345 | -0.336953017 | 0.078 | 0.053 | 0.311833494 | 1 |
| WBGene00004831 | slo-2 | 0.651514905 | 0.419389122 | -0.334886716 | 0.086 | 0.038 | 0.063133542 | 1 |
| WBGene00003920 | par-5 | 2.394998164 | 2.162988221 | -0.334719594 | 0.567 | 0.466 | 0.034126881 | 1 |
| WBGene00006703 | ubc-6 | 0.461953136 | 0.232267604 | -0.331366178 | 0.063 | 0.03 | 0.131555311 | 1 |
| WBGene00004143 | pqn-59 | 0.853067245 | 0.623389941 | -0.331354308 | 0.123 | 0.06 | 0.04037869 | 1 |
| WBGene00010639 | K07F5.15 | 0.417766337 | 0.188311068 | -0.331033979 | 0.052 | 0.015 | 0.062012566 | 1 |
| WBGene00021890 | Y54G2A.26 | 0.9473444 | 0.717997955 | -0.330876978 | 0.144 | 0.105 | 0.242625764 | 1 |
| WBGene00019691 | K12H6.12 | 0.959438357 | 0.730684849 | -0.330021551 | 0.125 | 0.075 | 0.121454027 | 1 |
| WBGene00004302 | ran-1 | 1.221987074 | 0.993477547 | -0.329669562 | 0.17 | 0.143 | 0.468177042 | 1 |
| WBGene00020562 | T19C3.4 | 0.550880218 | 0.322678466 | -0.329225536 | 0.077 | 0.03 | 0.058361468 | 1 |
| WBGene00194909 | F41E6.17 | 0.790861374 | 0.562916 | -0.32885566 | 0.088 | 0.068 | 0.449361727 | 1 |
| WBGene00003060 | lpd-3 | 0.562337898 | 0.334401283 | -0.328843024 | 0.056 | 0.038 | 0.397523491 | 1 |
| WBGene00002002 | hsb-1 | 0.454496606 | 0.226990114 | -0.328222488 | 0.054 | 0.023 | 0.128702875 | 1 |
| WBGene00003000 | lin-11 | 1.217116096 | 0.99006845 | -0.327560512 | 0.195 | 0.128 | 0.093955346 | 1 |
| WBGene00007350 | sucl-1 | 0.687934284 | 0.461323268 | -0.326930589 | 0.078 | 0.038 | 0.106381773 | 1 |
| WBGene00012645 | mrpl-22 | 0.579560713 | 0.353123817 | -0.326679388 | 0.07 | 0.038 | 0.161694614 | 1 |
| WBGene00000488 | che-7 | 0.714646239 | 0.488807171 | -0.325816903 | 0.072 | 0.038 | 0.145680312 | 1 |
| WBGene00020322 | T07F12.2 | 0.517435567 | 0.291809399 | -0.325509753 | 0.063 | 0.03 | 0.134260033 | 1 |
| WBGene00020825 | tmem-231 | 1.18856657 | 0.963119851 | -0.325250864 | 0.163 | 0.12 | 0.25522616 | 1 |
| WBGene00010130 | vha-14 | 0.645481136 | 0.420224245 | -0.324977001 | 0.08 | 0.038 | 0.097318856 | 1 |
| WBGene00185118 | K02E2.11 | 1.070738092 | 0.845631936 | -0.324759536 | 0.145 | 0.105 | 0.277090192 | 1 |

| | | | | | | | | |
|---|---|---|---|---|---|---|---|---|
| WBGene00195233 | F31E9.11 | 0.450855956 | 0.226425697 | -0.323784422 | 0.054 | 0.023 | 0.130088682 | 1 |
| WBGene00007247 | C01G12.7 | 2.877198672 | 2.653524861 | -0.322693098 | 0.66 | 0.602 | 0.066876922 | 1 |
| WBGene00008765 | ttx-7 | 1.673878024 | 1.451046363 | -0.321478132 | 0.322 | 0.218 | 0.054797026 | 1 |
| WBGene00004915 | snr-2 | 0.718178945 | 0.49554126 | -0.321198285 | 0.097 | 0.06 | 0.188993459 | 1 |
| WBGene00004479 | rps-10 | 2.73202644 | 2.509552566 | -0.320961955 | 0.672 | 0.602 | 0.106802355 | 1 |
| WBGene00012097 | abcf-2 | 0.628005668 | 0.405536132 | -0.320955697 | 0.076 | 0.038 | 0.116270105 | 1 |
| WBGene00009784 | F46C5.9 | 0.720477939 | 0.498248625 | -0.320609129 | 0.099 | 0.06 | 0.165265802 | 1 |
| WBGene00005111 | srd-33 | 1.410884205 | 1.188665822 | -0.320593359 | 0.219 | 0.15 | 0.091645921 | 1 |
| WBGene00011344 | T01G9.2 | 0.44859501 | 0.226393992 | -0.320568307 | 0.052 | 0.023 | 0.141257129 | 1 |
| WBGene00020260 | T05E7.4 | 0.597017436 | 0.375247124 | -0.319946929 | 0.081 | 0.03 | 0.043164286 | 1 |
| WBGene00022678 | sar-1 | 0.623184729 | 0.402303121 | -0.318664801 | 0.072 | 0.038 | 0.150318546 | 1 |
| WBGene00000403 | casy-1 | 2.500058623 | 2.279684353 | -0.317932866 | 0.576 | 0.489 | 0.079092508 | 1 |
| WBGene00016559 | C41A3.2 | 1.338522759 | 1.118792701 | -0.317003465 | 0.232 | 0.158 | 0.08671603 | 1 |
| WBGene00009122 | tct-1 | 0.900566112 | 0.681145069 | -0.31655765 | 0.132 | 0.083 | 0.129974968 | 1 |
| WBGene00007416 | ceh-57 | 1.776655509 | 1.557304102 | -0.316457187 | 0.34 | 0.271 | 0.24183537 | 1 |
| WBGene00000377 | cct-1 | 0.52270644 | 0.303493 | -0.316258144 | 0.061 | 0.03 | 0.160842173 | 1 |
| WBGene00015064 | B0228.7 | 1.023996669 | 0.804819041 | -0.316206476 | 0.143 | 0.105 | 0.300559596 | 1 |
| WBGene00015390 | C03F11.4 | 0.790282054 | 0.571921633 | -0.315027497 | 0.09 | 0.06 | 0.274440607 | 1 |
| WBGene00004346 | rgs-3 | 2.197153571 | 1.97888597 | -0.314893586 | 0.466 | 0.368 | 0.110139251 | 1 |
| WBGene00021522 | nhr-274 | 0.89275783 | 0.674866148 | -0.314351249 | 0.124 | 0.075 | 0.125817991 | 1 |
| WBGene00017121 | cyc-2.1 | 1.679891573 | 1.463059288 | -0.312822862 | 0.321 | 0.226 | 0.079309631 | 1 |
| WBGene00004424 | rpl-12 | 3.70727793 | 3.490960649 | -0.312079869 | 0.848 | 0.865 | 0.001890881 | 1 |
| WBGene00010865 | M04D8.7 | 2.159543903 | 1.943229875 | -0.312075177 | 0.455 | 0.353 | 0.083467008 | 1 |
| WBGene00022271 | cpx-1 | 1.355527681 | 1.139832777 | -0.311181968 | 0.231 | 0.158 | 0.108024802 | 1 |
| WBGene00015387 | kcnl-3 | 0.481683935 | 0.266668674 | -0.31020145 | 0.055 | 0.03 | 0.231849557 | 1 |
| WBGene00004432 | rpl-20 | 2.578807608 | 2.364270181 | -0.309512082 | 0.615 | 0.526 | 0.110412021 | 1 |
| WBGene00000095 | aha-1 | 0.494074914 | 0.280620468 | -0.307949671 | 0.062 | 0.03 | 0.149511374 | 1 |
| WBGene00013962 | ZK287.1 | 0.693043813 | 0.479824135 | -0.307610972 | 0.088 | 0.038 | 0.056169204 | 1 |
| WBGene00019871 | R04E5.8 | 0.878884143 | 0.665896074 | -0.30727683 | 0.123 | 0.068 | 0.078709703 | 1 |
| WBGene00006910 | vha-1 | 1.245336418 | 1.033239406 | -0.305991308 | 0.173 | 0.143 | 0.523824234 | 1 |
| WBGene00010962 | ctc-3 | 2.656279853 | 2.444548428 | -0.305463878 | 0.528 | 0.489 | 0.540736708 | 1 |
| WBGene00010971 | R01E6.7 | 0.6358683 | 0.424199888 | -0.305372969 | 0.07 | 0.045 | 0.291008529 | 1 |
| WBGene00007385 | atp-5 | 0.982459864 | 0.771143622 | -0.304864895 | 0.142 | 0.098 | 0.184921882 | 1 |
| WBGene00000378 | cct-2 | 0.555334736 | 0.34431812 | -0.304432626 | 0.059 | 0.03 | 0.185311648 | 1 |
| WBGene00001196 | egl-30 | 1.80444386 | 1.593510792 | -0.304312091 | 0.34 | 0.278 | 0.252099792 | 1 |
| WBGene00001665 | gpa-3 | 1.601214112 | 1.390449077 | -0.304069671 | 0.293 | 0.203 | 0.076538551 | 1 |
| WBGene00003014 | lin-28 | 0.481254494 | 0.270515935 | -0.304031475 | 0.056 | 0.03 | 0.223191347 | 1 |
| WBGene00019242 | srbc-40 | 1.312785464 | 1.102394018 | -0.303530696 | 0.198 | 0.12 | 0.047172134 | 1 |
| WBGene00004472 | rps-3 | 2.452570407 | 2.242819004 | -0.302607309 | 0.571 | 0.451 | 0.035282836 | 1 |
| WBGene00018704 | ztf-13 | 0.58408005 | 0.375860004 | -0.300398028 | 0.08 | 0.038 | 0.092953043 | 1 |
| WBGene00021304 | nphp-2 | 0.820364067 | 0.61243831 | -0.299973458 | 0.107 | 0.053 | 0.060043038 | 1 |
| WBGene00022489 | Y119D3B.12 | 0.602542663 | 0.395524046 | -0.298664733 | 0.07 | 0.038 | 0.177415151 | 1 |
| WBGene00015568 | sre-3 | 1.034479293 | 0.82751069 | -0.298592577 | 0.153 | 0.105 | 0.166373712 | 1 |
| WBGene00003404 | mpz-1 | 0.964846465 | 0.758570784 | -0.297592902 | 0.123 | 0.09 | 0.310585182 | 1 |
| WBGene00009262 | F30A10.3 | 0.680057532 | 0.474283409 | -0.296869307 | 0.068 | 0.045 | 0.336975962 | 1 |
| WBGene00003015 | lin-29 | 0.785307224 | 0.580657948 | -0.295246495 | 0.1 | 0.045 | 0.049497815 | 1 |
| WBGene00003559 | ncl-1 | 0.665532615 | 0.461854609 | -0.293845249 | 0.079 | 0.053 | 0.308719983 | 1 |
| WBGene00006917 | vha-8 | 0.41445867 | 0.211666064 | -0.292567887 | 0.05 | 0.023 | 0.160770573 | 1 |
| WBGene00022435 | cfap-36 | 0.558932158 | 0.356821873 | -0.291583505 | 0.061 | 0.038 | 0.29284668 | 1 |
| WBGene00001562 | lin-66 | 0.503310531 | 0.301432294 | -0.291248731 | 0.058 | 0.03 | 0.190591377 | 1 |
| WBGene00004884 | smg-6 | 0.600931545 | 0.399403451 | -0.290743582 | 0.057 | 0.038 | 0.379365459 | 1 |
| WBGene00022089 | Y69A2AR.18 | 1.007147559 | 0.806087043 | -0.290069009 | 0.126 | 0.105 | 0.579194064 | 1 |
| WBGene00016173 | fust-1 | 1.044377054 | 0.844094685 | -0.28894638 | 0.147 | 0.09 | 0.108278659 | 1 |
| WBGene00003476 | mtm-3 | 1.319167294 | 1.118889316 | -0.288940045 | 0.221 | 0.173 | 0.284293838 | 1 |
| WBGene00004363 | ric-3 | 0.613321753 | 0.413707501 | -0.287982492 | 0.066 | 0.045 | 0.37459575 | 1 |
| WBGene00001840 | hel-1 | 1.274670821 | 1.075364173 | -0.287538713 | 0.171 | 0.135 | 0.426208178 | 1 |
| WBGene00001669 | gpa-7 | 0.646862461 | 0.447799965 | -0.287186475 | 0.088 | 0.053 | 0.177824966 | 1 |
| WBGene00000464 | ceh-44 | 0.701552291 | 0.502593902 | -0.28703628 | 0.078 | 0.06 | 0.494668324 | 1 |
| WBGene00010012 | saeg-1 | 1.107609133 | 0.908784788 | -0.286842896 | 0.13 | 0.113 | 0.676739845 | 1 |
| WBGene00001121 | dyf-5 | 1.011163949 | 0.812614561 | -0.286446217 | 0.105 | 0.09 | 0.649808007 | 1 |
| WBGene00008168 | C48B4.3 | 1.389288654 | 1.191475046 | -0.285384711 | 0.239 | 0.18 | 0.18948265 | 1 |
| WBGene00003934 | pat-10 | 0.689976015 | 0.492318498 | -0.285159519 | 0.079 | 0.045 | 0.189954013 | 1 |
| WBGene00003072 | lrp-2 | 0.476664036 | 0.279627687 | -0.284263364 | 0.059 | 0.023 | 0.092230075 | 1 |
| WBGene00023417 | swm-1 | 1.510282392 | 1.314435435 | -0.282547433 | 0.193 | 0.165 | 0.559553472 | 1 |

| | | | | | | | | |
|---|---|---|---|---|---|---|---|---|
| WBGene00006807 | unc-75 | 0.423588392 | 0.228420435 | -0.281567845 | 0.051 | 0.023 | 0.154249137 | 1 |
| WBGene00017066 | maco-1 | 0.767396317 | 0.572420858 | -0.281290128 | 0.113 | 0.06 | 0.080937654 | 1 |
| WBGene00012194 | toe-4 | 1.008548214 | 0.814020419 | -0.280644285 | 0.151 | 0.113 | 0.284242352 | 1 |
| WBGene00002264 | lec-1 | 0.941719829 | 0.748416724 | -0.27887743 | 0.127 | 0.083 | 0.181424332 | 1 |
| WBGene00012813 | Y43F8B.2 | 1.725613089 | 1.532547606 | -0.278534616 | 0.249 | 0.218 | 0.456876063 | 1 |
| WBGene00020333 | sre-12 | 0.654049197 | 0.461780827 | -0.277384624 | 0.065 | 0.053 | 0.592862066 | 1 |
| WBGene00017579 | F18G5.1 | 0.638422726 | 0.446271151 | -0.277216124 | 0.085 | 0.053 | 0.221564663 | 1 |
| WBGene00020682 | T22D1.3 | 0.492666077 | 0.300536852 | -0.277183879 | 0.057 | 0.03 | 0.215146037 | 1 |
| WBGene00009085 | F23D12.1 | 0.555348648 | 0.363268719 | -0.277112762 | 0.064 | 0.03 | 0.128324216 | 1 |
| WBGene00001745 | gsa-1 | 1.312728323 | 1.121274396 | -0.27620963 | 0.176 | 0.135 | 0.308383615 | 1 |
| WBGene00001173 | egl-4 | 0.505006744 | 0.313625407 | -0.276104906 | 0.05 | 0.03 | 0.316582629 | 1 |
| WBGene00004423 | rpl-11.2 | 2.853127416 | 2.662167514 | -0.275496904 | 0.687 | 0.692 | 0.168358258 | 1 |
| WBGene00012337 | scyl-1 | 0.709441531 | 0.51859269 | -0.275336677 | 0.084 | 0.045 | 0.139830514 | 1 |
| WBGene00010887 | M7.12 | 1.560655579 | 1.36981353 | -0.275326878 | 0.244 | 0.211 | 0.525926328 | 1 |
| WBGene00013735 | Y111B2A.12 | 0.627889743 | 0.437198605 | -0.27510916 | 0.064 | 0.038 | 0.253846042 | 1 |
| WBGene00001448 | flp-5 | 0.58844256 | 0.398305201 | -0.274310225 | 0.051 | 0.045 | 0.737800191 | 1 |
| WBGene00004414 | rpl-3 | 2.518112443 | 2.32806835 | -0.274175671 | 0.565 | 0.481 | 0.1074178 | 1 |
| WBGene00004930 | sod-1 | 0.842826028 | 0.653621428 | -0.272964537 | 0.105 | 0.075 | 0.305345227 | 1 |
| WBGene00006772 | unc-36 | 0.587463698 | 0.398435328 | -0.272710291 | 0.06 | 0.045 | 0.515359348 | 1 |
| WBGene00016140 | rpb-2 | 0.551454181 | 0.362477768 | -0.272635333 | 0.052 | 0.038 | 0.472467918 | 1 |
| WBGene00003620 | nhr-21 | 1.806204238 | 1.617832417 | -0.271763092 | 0.344 | 0.218 | 0.014012147 | 1 |
| WBGene00018872 | F55C12.5 | 0.971331856 | 0.783769608 | -0.270595125 | 0.12 | 0.09 | 0.374183789 | 1 |
| WBGene00003600 | nhr-1 | 0.515992239 | 0.328787229 | -0.27007974 | 0.064 | 0.03 | 0.135221503 | 1 |
| WBGene00022235 | sqd-1 | 1.127047893 | 0.94000857 | -0.269840703 | 0.149 | 0.135 | 0.738274315 | 1 |
| WBGene00012354 | cox-4 | 0.85251201 | 0.665776131 | -0.269402926 | 0.119 | 0.075 | 0.164530022 | 1 |
| WBGene00304822 | T14G8.6 | 0.484268002 | 0.297683362 | -0.269184735 | 0.061 | 0.03 | 0.16459955 | 1 |
| WBGene00020762 | zer-1 | 0.734626226 | 0.548062504 | -0.269154556 | 0.082 | 0.06 | 0.426375722 | 1 |
| WBGene00004389 | rnp-6 | 0.63294077 | 0.446633589 | -0.268784447 | 0.062 | 0.045 | 0.468000631 | 1 |
| WBGene00002083 | inf-1 | 0.562725132 | 0.376654515 | -0.268443156 | 0.067 | 0.045 | 0.335338481 | 1 |
| WBGene00002363 | let-92 | 0.952568424 | 0.767755477 | -0.266628722 | 0.124 | 0.105 | 0.59385849 | 1 |
| WBGene00006536 | tbb-1 | 0.779465914 | 0.595082912 | -0.266008442 | 0.093 | 0.075 | 0.501181054 | 1 |
| WBGene00006392 | taf-10 | 0.692866482 | 0.509558137 | -0.264458039 | 0.082 | 0.06 | 0.402765751 | 1 |
| WBGene00006608 | tre-2 | 0.54509845 | 0.36221107 | -0.263850716 | 0.066 | 0.038 | 0.212214638 | 1 |
| WBGene00004471 | rps-2 | 2.251044873 | 2.068501856 | -0.263353906 | 0.511 | 0.421 | 0.080732607 | 1 |
| WBGene00020297 | nog-1 | 0.741645391 | 0.559217631 | -0.263187625 | 0.098 | 0.068 | 0.296974871 | 1 |
| WBGene00018757 | tag-304 | 0.855603911 | 0.673272618 | -0.263048452 | 0.11 | 0.068 | 0.161373237 | 1 |
| WBGene00003196 | mel-11 | 0.621091142 | 0.438834431 | -0.262940852 | 0.063 | 0.038 | 0.265858106 | 1 |
| WBGene00010958 | ndfl-4 | 1.610243646 | 1.428224009 | -0.262598828 | 0.239 | 0.226 | 0.830108234 | 1 |
| WBGene00017565 | ddo-2 | 0.788864372 | 0.607071517 | -0.26227165 | 0.09 | 0.053 | 0.158663237 | 1 |
| WBGene00000381 | cct-6 | 0.608628092 | 0.427483619 | -0.261336232 | 0.075 | 0.053 | 0.373425217 | 1 |
| WBGene00012765 | Y41E3.7 | 1.625607966 | 1.444891215 | -0.26071916 | 0.308 | 0.233 | 0.192665893 | 1 |
| WBGene00004447 | rpl-33 | 2.49044489 | 2.310320611 | -0.259864404 | 0.586 | 0.489 | 0.144293538 | 1 |
| WBGene00010408 | mboa-2 | 0.778703517 | 0.598611185 | -0.259818315 | 0.101 | 0.075 | 0.391390927 | 1 |
| WBGene00001177 | egl-8 | 0.742333567 | 0.562264823 | -0.259784284 | 0.092 | 0.06 | 0.251296051 | 1 |
| WBGene00017045 | atg-13 | 0.932206259 | 0.752477028 | -0.25929447 | 0.141 | 0.09 | 0.15448622 | 1 |
| WBGene00018213 | F39H12.2 | 1.653009266 | 1.473513642 | -0.258957447 | 0.323 | 0.233 | 0.100078306 | 1 |
| WBGene00002637 | let-418 | 0.756833341 | 0.577778738 | -0.258321189 | 0.097 | 0.075 | 0.457001868 | 1 |
| WBGene00004747 | sdc-3 | 0.410128334 | 0.231120901 | -0.258253136 | 0.05 | 0.023 | 0.163551455 | 1 |
| WBGene00011232 | pck-2 | 0.8233291 | 0.644589558 | -0.257866651 | 0.067 | 0.075 | 0.696158211 | 1 |
| WBGene00006803 | unc-70 | 0.784859927 | 0.606260915 | -0.25766391 | 0.099 | 0.068 | 0.295603558 | 1 |
| WBGene00003367 | mix-1 | 0.860284629 | 0.682194228 | -0.256930138 | 0.11 | 0.075 | 0.250049182 | 1 |
| WBGene00000064 | act-2 | 1.321263347 | 1.143183636 | -0.256914716 | 0.209 | 0.15 | 0.157346289 | 1 |
| WBGene00003609 | nhr-10 | 0.691306933 | 0.513520427 | -0.256491711 | 0.079 | 0.053 | 0.315815567 | 1 |
| WBGene00012896 | snrp-200 | 0.781179488 | 0.603443472 | -0.256418869 | 0.106 | 0.068 | 0.189062879 | 1 |
| WBGene00013307 | Y57G11C.9 | 1.708233612 | 1.530711287 | -0.256110578 | 0.314 | 0.263 | 0.453910558 | 1 |
| WBGene00011520 | nhr-213 | 0.666235685 | 0.488991684 | -0.25570904 | 0.086 | 0.053 | 0.212365062 | 1 |
| WBGene00001081 | dpy-22 | 0.534407877 | 0.358493274 | -0.253791125 | 0.05 | 0.038 | 0.535679928 | 1 |
| WBGene00000915 | hsp-90 | 2.673306591 | 2.497465759 | -0.253684698 | 0.63 | 0.541 | 0.132463972 | 1 |
| WBGene00019893 | sgt-1 | 0.903910311 | 0.728116029 | -0.253617538 | 0.124 | 0.09 | 0.29924517 | 1 |
| WBGene00016975 | nhr-173 | 0.983942866 | 0.808405298 | -0.253247179 | 0.097 | 0.098 | 0.920683283 | 1 |
| WBGene00000101 | aka-1 | 0.498158411 | 0.322754335 | -0.25305459 | 0.05 | 0.03 | 0.314659197 | 1 |
| WBGene00001994 | hpk-1 | 1.334642436 | 1.159393602 | -0.252830625 | 0.214 | 0.128 | 0.040061719 | 1 |
| WBGene00004450 | rpl-36 | 2.882931645 | 2.707817752 | -0.252635945 | 0.707 | 0.647 | 0.091598266 | 1 |
| WBGene00001030 | dnj-12 | 0.798330904 | 0.623348864 | -0.252445722 | 0.101 | 0.068 | 0.23860805 | 1 |

| | | | | | | | | |
|---|---|---|---|---|---|---|---|---|
| WBGene00015913 | C17F4.7 | 1.143102849 | 0.968286513 | -0.25220666 | 0.091 | 0.098 | 0.791185223 | 1 |
| WBGene00013900 | ugt-18 | 0.49034181 | 0.316276747 | -0.251122804 | 0.059 | 0.03 | 0.190434175 | 1 |
| WBGene00010239 | F58B4.6 | 2.496828474 | 2.323606192 | -0.249906928 | 0.569 | 0.474 | 0.113424605 | 1 |
| WBGene00022389 | pde-6 | 1.179674176 | 1.006501805 | -0.249834921 | 0.188 | 0.143 | 0.273816782 | 1 |
| WBGene00010131 | F55H2.5 | 0.418660028 | 0.245662125 | -0.249583216 | 0.057 | 0.023 | 0.106101145 | 1 |
| WBGene00004760 | sel-2 | 0.677081649 | 0.505097079 | -0.248121286 | 0.073 | 0.053 | 0.421272802 | 1 |
| WBGene00002182 | kap-1 | 0.609405695 | 0.437561384 | -0.247918935 | 0.067 | 0.038 | 0.210090008 | 1 |
| WBGene00001851 | hif-1 | 1.013195739 | 0.841551894 | -0.247629724 | 0.141 | 0.12 | 0.591911401 | 1 |
| WBGene00016138 | flh-2 | 0.795702152 | 0.624624965 | -0.246812209 | 0.107 | 0.075 | 0.292012893 | 1 |
| WBGene00019396 | K04F10.3 | 0.422537246 | 0.252120499 | -0.245859396 | 0.054 | 0.023 | 0.13028759 | 1 |
| WBGene00003923 | pas-2 | 0.495322058 | 0.324937826 | -0.245812486 | 0.059 | 0.038 | 0.32709303 | 1 |
| WBGene00000414 | cec-1 | 1.28766267 | 1.117581725 | -0.245374936 | 0.178 | 0.135 | 0.282049648 | 1 |
| WBGene00002335 | let-60 | 0.484526175 | 0.314539007 | -0.245239644 | 0.057 | 0.03 | 0.214585925 | 1 |
| WBGene00007104 | B0024.15 | 0.526927585 | 0.357893508 | -0.243864625 | 0.056 | 0.023 | 0.113690171 | 1 |
| WBGene00004408 | rla-0 | 3.494468478 | 3.327196331 | -0.241322697 | 0.824 | 0.797 | 0.214477971 | 1 |
| WBGene00011312 | trcs-2 | 0.508183187 | 0.341066142 | -0.241098932 | 0.056 | 0.038 | 0.408170951 | 1 |
| WBGene00013897 | ZC443.2 | 0.712056513 | 0.545302604 | -0.240575037 | 0.091 | 0.053 | 0.160697964 | 1 |
| WBGene00044354 | ZK418.11 | 0.691087076 | 0.524436081 | -0.240426564 | 0.083 | 0.045 | 0.139820563 | 1 |
| WBGene00007604 | xbx-9 | 1.319480043 | 1.152900667 | -0.240323239 | 0.236 | 0.165 | 0.126990949 | 1 |
| WBGene00003724 | nhr-134 | 0.74555869 | 0.579770246 | -0.239182166 | 0.09 | 0.068 | 0.409505326 | 1 |
| WBGene00004918 | snr-5 | 0.671550414 | 0.506395529 | -0.238268134 | 0.069 | 0.053 | 0.514374844 | 1 |
| WBGene00001701 | grd-12 | 1.176614939 | 1.012145124 | -0.237279787 | 0.17 | 0.135 | 0.322813367 | 1 |
| WBGene00017557 | nep-11 | 0.681834531 | 0.51767454 | -0.236832805 | 0.086 | 0.045 | 0.123451692 | 1 |
| WBGene00001955 | hlh-11 | 0.567406903 | 0.403828101 | -0.235994326 | 0.061 | 0.038 | 0.300942565 | 1 |
| WBGene00003851 | odr-4 | 1.517719237 | 1.354511152 | -0.235459496 | 0.271 | 0.211 | 0.288059426 | 1 |
| WBGene00023068 | rpl-41.2 | 3.78488126 | 3.621678172 | -0.235452285 | 0.861 | 0.887 | 0.052285113 | 1 |
| WBGene00013238 | trap-4 | 0.395953496 | 0.232758214 | -0.235441025 | 0.05 | 0.023 | 0.16550726 | 1 |
| WBGene00004483 | rps-14 | 3.330216527 | 3.167533919 | -0.234701391 | 0.801 | 0.797 | 0.025414115 | 1 |
| WBGene00003053 | lmp-1 | 0.829254988 | 0.668042759 | -0.232580083 | 0.124 | 0.068 | 0.07786167 | 1 |
| WBGene00015798 | C15H9.2 | 0.4551447 | 0.294843546 | -0.231265679 | 0.058 | 0.03 | 0.200446395 | 1 |
| WBGene00219650 | linc-56 | 1.531576566 | 1.371359708 | -0.231144066 | 0.205 | 0.165 | 0.43953792 | 1 |
| WBGene00004480 | rps-11 | 3.067715462 | 2.907765188 | -0.230759467 | 0.742 | 0.737 | 0.019197977 | 1 |
| WBGene00006600 | tph-1 | 2.454058028 | 2.294448722 | -0.230267554 | 0.565 | 0.474 | 0.076478285 | 1 |
| WBGene00004434 | rpl-22 | 2.549518564 | 2.390004123 | -0.230130692 | 0.607 | 0.556 | 0.341272239 | 1 |
| WBGene00006715 | ubc-20 | 0.485136825 | 0.325736645 | -0.22996585 | 0.064 | 0.03 | 0.133540596 | 1 |
| WBGene00000149 | apl-1 | 2.426099439 | 2.266786821 | -0.229839523 | 0.536 | 0.451 | 0.128591204 | 1 |
| WBGene00004274 | rab-11.1 | 1.204902266 | 1.046594372 | -0.228390015 | 0.189 | 0.15 | 0.379384595 | 1 |
| WBGene00004504 | rpt-4 | 0.636944573 | 0.48031451 | -0.225969416 | 0.083 | 0.06 | 0.399097492 | 1 |
| WBGene00006812 | unc-80 | 0.468633949 | 0.312682136 | -0.224990906 | 0.054 | 0.03 | 0.258420233 | 1 |
| WBGene00003412 | mrp-6 | 0.681065209 | 0.526551161 | -0.222916651 | 0.065 | 0.068 | 0.900240359 | 1 |
| WBGene00001480 | fmo-5 | 1.6671081 | 1.513498489 | -0.221611824 | 0.2 | 0.18 | 0.651917228 | 1 |
| WBGene00006433 | sdhb-1 | 0.625359069 | 0.472687404 | -0.220258654 | 0.07 | 0.053 | 0.468899498 | 1 |
| WBGene00001898 | his-24 | 1.27639868 | 1.125363383 | -0.217897874 | 0.186 | 0.158 | 0.562409723 | 1 |
| WBGene00013994 | ZK524.4 | 0.529267709 | 0.380366827 | -0.214818564 | 0.059 | 0.03 | 0.184200968 | 1 |
| WBGene00009625 | F41E7.9 | 0.796322612 | 0.648140636 | -0.213781401 | 0.09 | 0.075 | 0.632648112 | 1 |
| WBGene00009881 | F49C12.12 | 0.618915964 | 0.470779442 | -0.213715825 | 0.063 | 0.06 | 0.915761789 | 1 |
| WBGene00021260 | Y22D7AR.6 | 0.775939887 | 0.627897577 | -0.213579906 | 0.097 | 0.068 | 0.321936668 | 1 |
| WBGene00010849 | M04B2.7 | 0.57678611 | 0.429212186 | -0.212904167 | 0.076 | 0.045 | 0.219704013 | 1 |
| WBGene00002189 | kin-1 | 1.186532781 | 1.039173708 | -0.212594205 | 0.171 | 0.143 | 0.539052256 | 1 |
| WBGene00004449 | rpl-35 | 2.893586015 | 2.746289074 | -0.212504566 | 0.707 | 0.662 | 0.139498475 | 1 |
| WBGene00000140 | anc-1 | 0.837754812 | 0.690995899 | -0.211728356 | 0.095 | 0.083 | 0.662489274 | 1 |
| WBGene00000229 | atp-2 | 1.161626085 | 1.015170009 | -0.211291455 | 0.186 | 0.12 | 0.100670996 | 1 |
| WBGene00003649 | nhr-59 | 0.722757932 | 0.578184543 | -0.208575312 | 0.099 | 0.06 | 0.187735016 | 1 |
| WBGene00001972 | hmg-1.2 | 1.164679447 | 1.020310358 | -0.208280568 | 0.167 | 0.158 | 0.883874413 | 1 |
| WBGene00003687 | nhr-97 | 1.99232643 | 1.848126838 | -0.208036036 | 0.395 | 0.308 | 0.162570614 | 1 |
| WBGene00012621 | Y38H6C.8 | 1.452421065 | 1.308381597 | -0.207805027 | 0.165 | 0.173 | 0.888553254 | 1 |
| WBGene00004413 | rpl-2 | 2.64502634 | 2.501095579 | -0.207648195 | 0.634 | 0.602 | 0.497257937 | 1 |
| WBGene00000901 | daf-5 | 0.830635247 | 0.686827186 | -0.207471176 | 0.094 | 0.09 | 0.901842839 | 1 |
| WBGene00007801 | trpa-1 | 0.422384922 | 0.278893136 | -0.207014888 | 0.051 | 0.023 | 0.153558247 | 1 |
| WBGene00012348 | pptr-1 | 1.077632546 | 0.934281559 | -0.206811758 | 0.171 | 0.12 | 0.186810063 | 1 |
| WBGene00006843 | unc-119 | 1.435969406 | 1.292753022 | -0.206617567 | 0.233 | 0.188 | 0.413521093 | 1 |
| WBGene00015507 | C06A6.2 | 0.49493429 | 0.351779946 | -0.206528062 | 0.068 | 0.045 | 0.329406208 | 1 |
| WBGene00008366 | tpra-1 | 0.798069994 | 0.654999027 | -0.206407775 | 0.111 | 0.068 | 0.155959688 | 1 |
| WBGene00004456 | rpl-43 | 2.891807128 | 2.749010417 | -0.206012108 | 0.72 | 0.684 | 0.174005484 | 1 |

| | | | | | | | | |
|---|---|---|---|---|---|---|---|---|
| WBGene00004387 | rnp-4 | 0.80111945 | 0.658899516 | -0.205179993 | 0.102 | 0.075 | 0.373764968 | 1 |
| WBGene00006412 | nlg-1 | 1.840443118 | 1.698558906 | -0.204695649 | 0.329 | 0.316 | 0.947408339 | 1 |
| WBGene00003858 | ogt-1 | 1.166011002 | 1.024835136 | -0.203673721 | 0.169 | 0.128 | 0.333916603 | 1 |
| WBGene00000881 | cyn-5 | 0.534131841 | 0.393095178 | -0.203472894 | 0.06 | 0.038 | 0.319253151 | 1 |
| WBGene00020100 | mks-1 | 1.04777034 | 0.906798805 | -0.203378934 | 0.145 | 0.113 | 0.421937783 | 1 |
| WBGene00020391 | cct-7 | 0.859596959 | 0.718870754 | -0.203024998 | 0.108 | 0.09 | 0.590991169 | 1 |
| WBGene00019025 | srb-16 | 0.85395557 | 0.713962745 | -0.201966955 | 0.106 | 0.083 | 0.459109988 | 1 |
| WBGene00016121 | ift-43 | 1.205547667 | 1.065716959 | -0.201733069 | 0.185 | 0.165 | 0.707823736 | 1 |
| WBGene00003084 | lst-2 | 0.561647021 | 0.422958562 | -0.200085152 | 0.067 | 0.038 | 0.2054302 | 1 |
| WBGene00003403 | mps-1 | 1.086802765 | 0.948296071 | -0.19982292 | 0.171 | 0.12 | 0.195179176 | 1 |
| WBGene00016744 | bbs-9 | 0.973089078 | 0.835151162 | -0.199002347 | 0.156 | 0.113 | 0.262333128 | 1 |
| WBGene00013308 | nuo-3 | 1.273131521 | 1.13520798 | -0.198981608 | 0.215 | 0.165 | 0.294032353 | 1 |
| WBGene00020951 | sna-1 | 0.409242231 | 0.271418181 | -0.198838074 | 0.05 | 0.03 | 0.321230267 | 1 |
| WBGene00003883 | osm-1 | 1.329041935 | 1.191464406 | -0.198482419 | 0.197 | 0.188 | 0.911017325 | 1 |
| WBGene00000085 | aex-2 | 1.659235898 | 1.522626533 | -0.197085653 | 0.305 | 0.241 | 0.287543914 | 1 |
| WBGene00002152 | iars-1 | 0.503803206 | 0.368211577 | -0.195617371 | 0.051 | 0.038 | 0.501420454 | 1 |
| WBGene00044698 | K09F5.6 | 0.731064189 | 0.596025508 | -0.194819635 | 0.086 | 0.053 | 0.213048265 | 1 |
| WBGene00012850 | Y44A6C.1 | 0.448541217 | 0.313522107 | -0.1947914 | 0.061 | 0.03 | 0.166502838 | 1 |
| WBGene00022233 | ipla-6 | 0.492097347 | 0.35717848 | -0.19464678 | 0.061 | 0.03 | 0.165941334 | 1 |
| WBGene00008342 | frpr-5 | 0.904706223 | 0.769862615 | -0.194538205 | 0.116 | 0.083 | 0.317619668 | 1 |
| WBGene00010367 | H05L14.2 | 0.646908737 | 0.513023943 | -0.193154929 | 0.08 | 0.06 | 0.458315665 | 1 |
| WBGene00007055 | tag-196 | 0.739240897 | 0.605492251 | -0.192958509 | 0.086 | 0.068 | 0.504193236 | 1 |
| WBGene00001681 | gpc-1 | 2.082592901 | 1.948881801 | -0.192904341 | 0.446 | 0.398 | 0.515778912 | 1 |
| WBGene00219300 | W10G6.6 | 0.832911809 | 0.700040139 | -0.1916933 | 0.112 | 0.083 | 0.348098685 | 1 |
| WBGene00012894 | tbc-17 | 0.943447988 | 0.811595037 | -0.190223599 | 0.133 | 0.105 | 0.430391975 | 1 |
| WBGene00000292 | cap-1 | 0.392630251 | 0.261108428 | -0.189745881 | 0.05 | 0.03 | 0.3096943 | 1 |
| WBGene00022376 | Y94H6A.3 | 0.582711818 | 0.451315082 | -0.18956542 | 0.059 | 0.045 | 0.538511979 | 1 |
| WBGene00010484 | dph-3 | 0.429348571 | 0.29831648 | -0.189039348 | 0.054 | 0.023 | 0.134519522 | 1 |
| WBGene00012779 | cdkl-1 | 1.111565031 | 0.981319736 | -0.187904241 | 0.149 | 0.135 | 0.76771585 | 1 |
| WBGene00012117 | T28B11.1 | 2.236615331 | 2.106642963 | -0.187510491 | 0.48 | 0.353 | 0.045328102 | 1 |
| WBGene00016002 | gyf-1 | 1.437921971 | 1.308175535 | -0.187184539 | 0.257 | 0.211 | 0.404500911 | 1 |
| WBGene00000475 | cey-4 | 0.636888208 | 0.507314735 | -0.186935006 | 0.077 | 0.045 | 0.203999476 | 1 |
| WBGene00003850 | odr-3 | 2.068657503 | 1.939147378 | -0.186843615 | 0.437 | 0.391 | 0.634240552 | 1 |
| WBGene00006823 | unc-94 | 1.108387573 | 0.979760945 | -0.185568997 | 0.172 | 0.128 | 0.279692135 | 1 |
| WBGene00018967 | ddx-15 | 0.398633783 | 0.270198966 | -0.185292273 | 0.052 | 0.023 | 0.143407016 | 1 |
| WBGene00015790 | C15C7.4 | 2.239980122 | 2.114297465 | -0.181321745 | 0.366 | 0.286 | 0.138094237 | 1 |
| WBGene00016463 | C35E7.11 | 1.244152476 | 1.118738376 | -0.180934301 | 0.196 | 0.165 | 0.464299081 | 1 |
| WBGene00017014 | dylt-2 | 0.640149832 | 0.515851095 | -0.179325171 | 0.078 | 0.06 | 0.488379488 | 1 |
| WBGene00004313 | rbc-1 | 0.506528799 | 0.382679014 | -0.178677471 | 0.054 | 0.038 | 0.45878234 | 1 |
| WBGene00012179 | W01D2.1 | 2.790162926 | 2.666431824 | -0.178506246 | 0.66 | 0.609 | 0.220775085 | 1 |
| WBGene00022144 | pghm-1 | 1.502275832 | 1.379013888 | -0.177829395 | 0.242 | 0.203 | 0.468473483 | 1 |
| WBGene00006055 | ssr-2 | 1.366177518 | 1.242921684 | -0.177820581 | 0.237 | 0.158 | 0.106501645 | 1 |
| WBGene00019285 | cbn-1 | 2.269582005 | 2.147557812 | -0.176043699 | 0.503 | 0.436 | 0.365898823 | 1 |
| WBGene00016817 | C50E3.6 | 0.497528268 | 0.37586482 | -0.175523254 | 0.051 | 0.03 | 0.300408091 | 1 |
| WBGene00003025 | lin-40 | 0.983203793 | 0.861622583 | -0.175404609 | 0.129 | 0.105 | 0.536365878 | 1 |
| WBGene00016790 | C49H3.3 | 1.044741073 | 0.923425941 | -0.17502074 | 0.143 | 0.113 | 0.398591927 | 1 |
| WBGene00009547 | F38H4.3 | 0.523220341 | 0.402379478 | -0.174336514 | 0.065 | 0.045 | 0.391398239 | 1 |
| WBGene00001558 | gdi-1 | 1.256860816 | 1.136072496 | -0.17426071 | 0.165 | 0.173 | 0.657297658 | 1 |
| WBGene00016868 | C52B9.8 | 1.139092227 | 1.01858431 | -0.173856175 | 0.182 | 0.128 | 0.1930754 | 1 |
| WBGene00009974 | F53C11.4 | 0.863149417 | 0.743482871 | -0.172642332 | 0.133 | 0.09 | 0.218353131 | 1 |
| WBGene00021936 | zipt-15 | 0.962115083 | 0.842641452 | -0.172364016 | 0.122 | 0.083 | 0.250574326 | 1 |
| WBGene00005742 | srv-31 | 0.416844085 | 0.299296086 | -0.169585915 | 0.051 | 0.015 | 0.067733981 | 1 |
| WBGene00015423 | C04E6.7 | 1.348092525 | 1.231546881 | -0.168139822 | 0.225 | 0.18 | 0.326184232 | 1 |
| WBGene00012826 | dyf-19 | 0.835500772 | 0.719500084 | -0.167353617 | 0.111 | 0.098 | 0.692339496 | 1 |
| WBGene00019324 | K02F2.5 | 2.484286822 | 2.368614084 | -0.166880486 | 0.537 | 0.481 | 0.698884514 | 1 |
| WBGene00009688 | F44E5.1 | 0.687398673 | 0.572149984 | -0.166268713 | 0.08 | 0.068 | 0.710861688 | 1 |
| WBGene00019150 | H04J21.1 | 0.669056259 | 0.553878572 | -0.166166279 | 0.08 | 0.06 | 0.45928157 | 1 |
| WBGene00004184 | T19D2.2 | 1.772549762 | 1.657596289 | -0.165842805 | 0.352 | 0.301 | 0.491181295 | 1 |
| WBGene00001149 | bcat-1 | 1.107486623 | 0.993138091 | -0.164970061 | 0.157 | 0.12 | 0.368229501 | 1 |
| WBGene00195084 | C43C3.4 | 0.869806603 | 0.755539287 | -0.164852891 | 0.102 | 0.09 | 0.750851893 | 1 |
| WBGene00004441 | rpl-27 | 2.095858715 | 1.98227247 | -0.163870312 | 0.44 | 0.368 | 0.355526883 | 1 |
| WBGene00004475 | rps-6 | 2.180723634 | 2.06728664 | -0.163654989 | 0.466 | 0.414 | 0.43744171 | 1 |
| WBGene00022599 | daf-41 | 1.985667113 | 1.872449457 | -0.163338551 | 0.406 | 0.346 | 0.494255406 | 1 |
| WBGene00001194 | egl-27 | 1.061531654 | 0.948891629 | -0.162505206 | 0.159 | 0.128 | 0.447130749 | 1 |

| | | | | | | | | |
|---|---|---|---|---|---|---|---|---|
| WBGene00007855 | C31H5.4 | 0.675638969 | 0.563248095 | -0.162145756 | 0.1 | 0.06 | 0.181390404 | 1 |
| WBGene00019749 | M03A8.3 | 0.915945063 | 0.804197054 | -0.1612183 | 0.111 | 0.098 | 0.677143413 | 1 |
| WBGene00019364 | mksr-1 | 0.734492289 | 0.622780566 | -0.161165949 | 0.073 | 0.083 | 0.653665855 | 1 |
| WBGene00000485 | che-3 | 1.746597022 | 1.636442812 | -0.158918932 | 0.301 | 0.256 | 0.451844925 | 1 |
| WBGene00004410 | rla-2 | 2.570366105 | 2.460343267 | -0.158729403 | 0.603 | 0.564 | 0.860112431 | 1 |
| WBGene00004921 | snt-1 | 0.868259898 | 0.758252953 | -0.158706474 | 0.112 | 0.075 | 0.249264512 | 1 |
| WBGene00017321 | F09G8.7 | 0.47048687 | 0.36052558 | -0.158640607 | 0.055 | 0.038 | 0.440965773 | 1 |
| WBGene00004087 | ppk-1 | 0.581704838 | 0.471870441 | -0.15845754 | 0.072 | 0.053 | 0.436719975 | 1 |
| WBGene00001668 | gpa-6 | 0.804428462 | 0.694649552 | -0.15837749 | 0.101 | 0.068 | 0.278584527 | 1 |
| WBGene00044742 | Y55F3BR.10 | 0.697068904 | 0.587962633 | -0.157407076 | 0.072 | 0.045 | 0.277913296 | 1 |
| WBGene00020581 | T19D12.6 | 1.279473437 | 1.171323883 | -0.156026825 | 0.209 | 0.15 | 0.189328526 | 1 |
| WBGene00001088 | dpy-30 | 0.71515665 | 0.607496716 | -0.155320453 | 0.102 | 0.06 | 0.167610829 | 1 |
| WBGene00021044 | W05H7.1 | 0.621092689 | 0.514480076 | -0.153809487 | 0.076 | 0.06 | 0.559968423 | 1 |
| WBGene00002113 | ins-30 | 1.638066654 | 1.531527607 | -0.153703355 | 0.206 | 0.226 | 0.537463318 | 1 |
| WBGene00000778 | cpn-2 | 1.578607833 | 1.472238755 | -0.153458141 | 0.293 | 0.233 | 0.242755041 | 1 |
| WBGene00011558 | ostf-4 | 0.563305252 | 0.456955786 | -0.153429847 | 0.063 | 0.045 | 0.443363724 | 1 |
| WBGene00000763 | coq-3 | 0.520382867 | 0.414392144 | -0.15291229 | 0.061 | 0.045 | 0.496464653 | 1 |
| WBGene00015287 | osta-1 | 0.638190058 | 0.532259072 | -0.15282611 | 0.078 | 0.045 | 0.197760384 | 1 |
| WBGene00004896 | smu-2 | 0.515652872 | 0.409734672 | -0.152807663 | 0.054 | 0.023 | 0.134111958 | 1 |
| WBGene00011303 | R166.3 | 0.590772807 | 0.484948419 | -0.15267232 | 0.07 | 0.045 | 0.304573377 | 1 |
| WBGene00016454 | C35E7.2 | 1.106674471 | 1.000877827 | -0.152632294 | 0.158 | 0.143 | 0.67994787 | 1 |
| WBGene00017814 | F26A10.2 | 0.577155096 | 0.471502661 | -0.152424243 | 0.055 | 0.045 | 0.654137679 | 1 |
| WBGene00005719 | srv-8 | 0.99321588 | 0.888457263 | -0.151134737 | 0.146 | 0.128 | 0.690462243 | 1 |
| WBGene00008915 | F17C11.4 | 0.870299518 | 0.766293593 | -0.150048833 | 0.106 | 0.098 | 0.847134526 | 1 |
| WBGene00003825 | ntl-2 | 0.925969796 | 0.822009064 | -0.149983632 | 0.14 | 0.113 | 0.46196103 | 1 |
| WBGene00022527 | ZC132.9 | 0.953919581 | 0.850176562 | -0.14966954 | 0.133 | 0.098 | 0.327206054 | 1 |
| WBGene00077491 | K08H10.10 | 2.195462401 | 2.091747304 | -0.149629256 | 0.455 | 0.414 | 0.52097189 | 1 |
| WBGene00021534 | mvk-1 | 0.562835455 | 0.459261555 | -0.149425552 | 0.065 | 0.053 | 0.60937904 | 1 |
| WBGene00004496 | rps-27 | 3.091787404 | 2.988880108 | -0.148463847 | 0.762 | 0.759 | 0.742987122 | 1 |
| WBGene00020246 | clic-1 | 0.776772401 | 0.674468583 | -0.147593211 | 0.103 | 0.083 | 0.510203515 | 1 |
| WBGene00001565 | gei-8 | 0.512187713 | 0.410112267 | -0.147263739 | 0.058 | 0.038 | 0.365115058 | 1 |
| WBGene00018582 | F48A9.2 | 0.552582398 | 0.451031715 | -0.146506666 | 0.067 | 0.053 | 0.53941239 | 1 |
| WBGene00012893 | Y45F10D.15 | 0.448152376 | 0.346838353 | -0.146165239 | 0.054 | 0.03 | 0.264190954 | 1 |
| WBGene00014233 | ZK1128.7 | 0.415378519 | 0.314219261 | -0.145941959 | 0.051 | 0.03 | 0.300777209 | 1 |
| WBGene00004453 | rpl-39 | 3.010385316 | 2.909271834 | -0.145875919 | 0.742 | 0.699 | 0.474975085 | 1 |
| WBGene00001447 | flp-4 | 1.24096461 | 1.140023115 | -0.145627794 | 0.134 | 0.098 | 0.273633615 | 1 |
| WBGene00004032 | pkc-1 | 0.693531825 | 0.592879404 | -0.145210749 | 0.093 | 0.068 | 0.379503251 | 1 |
| WBGene00011545 | T06G6.3 | 0.93656414 | 0.836648523 | -0.144147765 | 0.128 | 0.113 | 0.666798766 | 1 |
| WBGene00012016 | inpp-1 | 0.792157608 | 0.692444689 | -0.143855333 | 0.103 | 0.068 | 0.249451807 | 1 |
| WBGene00020142 | aak-2 | 0.933553472 | 0.833843941 | -0.143850446 | 0.129 | 0.083 | 0.169158848 | 1 |
| WBGene00003950 | pbs-4 | 0.662599762 | 0.562993513 | -0.143701442 | 0.092 | 0.06 | 0.240563425 | 1 |
| WBGene00012274 | W05B5.1 | 0.986051616 | 0.886715061 | -0.143312356 | 0.127 | 0.113 | 0.712861243 | 1 |
| WBGene00010957 | nduo-6 | 3.885805814 | 3.787146103 | -0.142335876 | 0.837 | 0.91 | 0.559167033 | 1 |
| WBGene00019643 | K11C4.2 | 1.475072626 | 1.376544553 | -0.142145963 | 0.224 | 0.18 | 0.395739276 | 1 |
| WBGene00018008 | vpr-1 | 0.63465238 | 0.537672929 | -0.139911772 | 0.083 | 0.053 | 0.262444771 | 1 |
| WBGene00017350 | F10E7.9 | 0.631875115 | 0.535525512 | -0.139003096 | 0.079 | 0.053 | 0.314573919 | 1 |
| WBGene00000510 | cka-2 | 0.869261715 | 0.772988077 | -0.138893499 | 0.114 | 0.098 | 0.627476979 | 1 |
| WBGene00003822 | nsy-1 | 0.923497691 | 0.827536322 | -0.138442991 | 0.124 | 0.083 | 0.199979594 | 1 |
| WBGene00018608 | F48E3.9 | 0.855725615 | 0.75999858 | -0.13810492 | 0.102 | 0.09 | 0.735444839 | 1 |
| WBGene00001464 | flp-21 | 2.151142931 | 2.056438952 | -0.136628961 | 0.361 | 0.338 | 0.865972366 | 1 |
| WBGene00020052 | R13A5.10 | 0.966624605 | 0.872003403 | -0.13650954 | 0.136 | 0.098 | 0.299095973 | 1 |
| WBGene00011953 | ppm-2 | 0.80278785 | 0.708293468 | -0.136326577 | 0.099 | 0.083 | 0.603648282 | 1 |
| WBGene00015125 | B0303.3 | 0.608626257 | 0.514192965 | -0.136238441 | 0.076 | 0.053 | 0.358992963 | 1 |
| WBGene00001166 | eftu-2 | 0.533150782 | 0.43917325 | -0.135580921 | 0.06 | 0.038 | 0.322114463 | 1 |
| WBGene00004429 | rpl-17 | 2.889021401 | 2.795307069 | -0.135201203 | 0.695 | 0.692 | 0.802335962 | 1 |
| WBGene00010796 | clip-1 | 0.615543988 | 0.521988045 | -0.134972694 | 0.067 | 0.053 | 0.582090189 | 1 |
| WBGene00012973 | spat-2 | 0.608316131 | 0.51504318 | -0.134564424 | 0.061 | 0.045 | 0.504280164 | 1 |
| WBGene00019079 | rpa-4 | 0.681138381 | 0.58841482 | -0.133771822 | 0.082 | 0.068 | 0.598886419 | 1 |
| WBGene00019180 | H10D18.5 | 1.279852575 | 1.187341861 | -0.133464748 | 0.212 | 0.165 | 0.339680687 | 1 |
| WBGene00008555 | F07C6.4 | 0.874093845 | 0.782313395 | -0.1324112 | 0.109 | 0.09 | 0.569788985 | 1 |
| WBGene00020163 | T02G5.3 | 1.601157959 | 1.509650417 | -0.132017477 | 0.293 | 0.248 | 0.449333598 | 1 |
| WBGene00011398 | qdpr-1 | 0.581131496 | 0.489855526 | -0.131683389 | 0.062 | 0.045 | 0.47996785 | 1 |
| WBGene00004238 | ptr-24 | 0.488697752 | 0.397570557 | -0.131468751 | 0.05 | 0.03 | 0.317740375 | 1 |
| WBGene00014454 | MTCE.7 | 3.059971359 | 2.971037076 | -0.128305048 | 0.672 | 0.684 | 0.563262874 | 1 |

| | | | | | | | | |
|---|---|---|---|---|---|---|---|---|
| WBGene00015611 | C08F1.8 | 0.759411644 | 0.670650862 | -0.128054739 | 0.092 | 0.083 | 0.754977734 | 1 |
| WBGene00005743 | srv-32 | 0.920024659 | 0.831596779 | -0.127574464 | 0.112 | 0.09 | 0.500090761 | 1 |
| WBGene00002179 | jph-1 | 0.462654254 | 0.374672198 | -0.126931276 | 0.054 | 0.038 | 0.462996071 | 1 |
| WBGene00004740 | scd-2 | 2.296169176 | 2.208405927 | -0.126615604 | 0.499 | 0.414 | 0.238624168 | 1 |
| WBGene00009454 | F36A2.7 | 0.693686062 | 0.606915152 | -0.125183962 | 0.09 | 0.053 | 0.167091802 | 1 |
| WBGene00017240 | calu-2 | 0.527605165 | 0.441403935 | -0.124362088 | 0.069 | 0.045 | 0.333756694 | 1 |
| WBGene00014145 | ZK899.7 | 1.031825089 | 0.946278878 | -0.123417094 | 0.153 | 0.105 | 0.21635889 | 1 |
| WBGene00000223 | atf-7 | 1.241222354 | 1.155729979 | -0.123339426 | 0.187 | 0.143 | 0.371639248 | 1 |
| WBGene00011884 | enol-1 | 1.301152157 | 1.215888548 | -0.123009386 | 0.205 | 0.18 | 0.590172098 | 1 |
| WBGene00012768 | eef-1B.2 | 1.080840565 | 0.995592375 | -0.122987141 | 0.162 | 0.12 | 0.335328995 | 1 |
| WBGene00018117 | nlp-47 | 2.260624647 | 2.175485352 | -0.122830039 | 0.477 | 0.429 | 0.618907416 | 1 |
| WBGene00019875 | mca-2 | 0.614740225 | 0.529827751 | -0.122502805 | 0.079 | 0.045 | 0.180076713 | 1 |
| WBGene00004277 | rab-18 | 0.580458425 | 0.496240904 | -0.121500199 | 0.076 | 0.053 | 0.38335421 | 1 |
| WBGene00020628 | T20F5.6 | 0.552325698 | 0.469266081 | -0.119829698 | 0.057 | 0.045 | 0.603641228 | 1 |
| WBGene00007647 | C17E4.10 | 1.69444061 | 1.611630399 | -0.119469881 | 0.294 | 0.241 | 0.304076576 | 1 |
| WBGene00017220 | F07F6.8 | 1.402014883 | 1.319277639 | -0.119364612 | 0.229 | 0.165 | 0.15838279 | 1 |
| WBGene00007192 | B0491.5 | 0.585643075 | 0.503022511 | -0.119196278 | 0.057 | 0.045 | 0.613029237 | 1 |
| WBGene00003966 | pdl-1 | 1.706791209 | 1.624413006 | -0.118846625 | 0.321 | 0.278 | 0.604737328 | 1 |
| WBGene00016982 | C56G2.9 | 1.543047652 | 1.461934725 | -0.117021217 | 0.272 | 0.256 | 0.919530792 | 1 |
| WBGene00017699 | flcn-1 | 0.754546225 | 0.673631817 | -0.116734815 | 0.097 | 0.075 | 0.481709196 | 1 |
| WBGene00006919 | vha-10 | 0.65924312 | 0.578603931 | -0.116337758 | 0.071 | 0.068 | 0.867778595 | 1 |
| WBGene00003715 | nhr-125 | 0.583652613 | 0.504347641 | -0.11441289 | 0.063 | 0.045 | 0.448040668 | 1 |
| WBGene00004152 | pqn-70 | 0.632794309 | 0.553863104 | -0.113873657 | 0.084 | 0.053 | 0.252035558 | 1 |
| WBGene00001746 | gsk-3 | 0.82762962 | 0.74891761 | -0.113557426 | 0.082 | 0.098 | 0.510289513 | 1 |
| WBGene00001130 | dyn-1 | 0.481818689 | 0.4031571 | -0.113484685 | 0.056 | 0.038 | 0.402187597 | 1 |
| WBGene00000192 | arl-8 | 0.623770813 | 0.545121686 | -0.113466705 | 0.07 | 0.06 | 0.68755 | 1 |
| WBGene00002196 | kin-10 | 0.708012239 | 0.630723221 | -0.111504482 | 0.078 | 0.06 | 0.5351995 | 1 |
| WBGene00004461 | rpn-5 | 0.495394295 | 0.418192715 | -0.111378337 | 0.054 | 0.045 | 0.704276501 | 1 |
| WBGene00000962 | dhc-1 | 0.653898858 | 0.577003421 | -0.110936665 | 0.07 | 0.075 | 0.867256693 | 1 |
| WBGene00043992 | bbs-4 | 1.057711077 | 0.980870635 | -0.110857324 | 0.148 | 0.113 | 0.373420154 | 1 |
| WBGene00004452 | rpl-38 | 3.398495899 | 3.321750399 | -0.110720352 | 0.803 | 0.857 | 0.130760792 | 1 |
| WBGene00017288 | F09E5.10 | 0.677895005 | 0.601303021 | -0.110498876 | 0.082 | 0.06 | 0.437267529 | 1 |
| WBGene00006965 | xtr-1 | 0.658863636 | 0.582895806 | -0.109598411 | 0.073 | 0.053 | 0.418406123 | 1 |
| WBGene00004357 | rho-1 | 1.166607042 | 1.090795913 | -0.10937234 | 0.166 | 0.165 | 0.931180502 | 1 |
| WBGene00018700 | lmd-3 | 1.361604938 | 1.286355948 | -0.108561344 | 0.214 | 0.18 | 0.538990733 | 1 |
| WBGene00021350 | Y37E3.8 | 3.211415235 | 3.136360037 | -0.108281761 | 0.778 | 0.82 | 0.309900277 | 1 |
| WBGene00019395 | K04F10.2 | 0.850458097 | 0.775862446 | -0.107618776 | 0.112 | 0.09 | 0.495604214 | 1 |
| WBGene00017616 | F20A1.4 | 2.087381508 | 2.013051021 | -0.107236226 | 0.417 | 0.376 | 0.809350173 | 1 |
| WBGene00008224 | slrp-1 | 0.653589561 | 0.580168186 | -0.105924653 | 0.092 | 0.068 | 0.400931316 | 1 |
| WBGene00017778 | nono-1 | 0.690027785 | 0.617752182 | -0.104271653 | 0.08 | 0.053 | 0.294585392 | 1 |
| WBGene00001439 | fkh-7 | 1.565933758 | 1.493923391 | -0.103889 | 0.279 | 0.226 | 0.340112586 | 1 |
| WBGene00003656 | nhr-66 | 0.816544501 | 0.744926648 | -0.103322722 | 0.098 | 0.075 | 0.475160021 | 1 |
| WBGene00004919 | snr-6 | 0.813674096 | 0.742473566 | -0.102720653 | 0.111 | 0.075 | 0.252673102 | 1 |
| WBGene00015861 | C16D9.6 | 1.445016067 | 1.374258301 | -0.102081877 | 0.22 | 0.248 | 0.456933028 | 1 |
| WBGene00004412 | rpl-1 | 2.564062056 | 2.494238862 | -0.100733575 | 0.61 | 0.586 | 0.866387146 | 1 |
| WBGene00004466 | rpn-10 | 0.326121293 | 0.396849037 | 0.102038566 | 0.038 | 0.053 | 0.398764065 | 1 |
| WBGene00012760 | Y41C4A.17 | 4.561001424 | 4.632560348 | 0.103237705 | 0.793 | 0.887 | 0.07422669 | 1 |
| WBGene00000123 | ama-1 | 0.614435076 | 0.686676764 | 0.104222725 | 0.076 | 0.083 | 0.722166462 | 1 |
| WBGene00022127 | yop-1 | 0.487040932 | 0.559398421 | 0.104389791 | 0.058 | 0.06 | 0.853206274 | 1 |
| WBGene00002889 | let-765 | 0.723614247 | 0.796207352 | 0.104729712 | 0.078 | 0.098 | 0.365637976 | 1 |
| WBGene00009917 | glb-18 | 1.789877051 | 1.863294835 | 0.105919473 | 0.333 | 0.338 | 0.572308036 | 1 |
| WBGene00011304 | mnk-1 | 0.655740875 | 0.729505362 | 0.106419659 | 0.079 | 0.083 | 0.807539208 | 1 |
| WBGene00003725 | nhr-135 | 1.069813598 | 1.144051402 | 0.107102511 | 0.155 | 0.158 | 0.72323427 | 1 |
| WBGene00001189 | egl-21 | 1.723890941 | 1.799351021 | 0.108865883 | 0.344 | 0.301 | 0.762748492 | 1 |
| WBGene00010425 | lpin-1 | 0.999675367 | 1.075286291 | 0.109083505 | 0.127 | 0.143 | 0.528512539 | 1 |
| WBGene00018710 | F52G3.1 | 0.566196789 | 0.64264014 | 0.110284444 | 0.064 | 0.038 | 0.260127266 | 1 |
| WBGene00016792 | C49H3.6 | 0.548718886 | 0.628974839 | 0.115784866 | 0.073 | 0.075 | 0.861619772 | 1 |
| WBGene00001989 | hot-4 | 0.939153563 | 1.021229691 | 0.118410822 | 0.125 | 0.128 | 0.742538081 | 1 |
| WBGene00003646 | nhr-56 | 0.612247017 | 0.69484081 | 0.119157656 | 0.082 | 0.09 | 0.680541708 | 1 |
| WBGene00019408 | K05F1.6 | 0.472717138 | 0.555539891 | 0.119487974 | 0.054 | 0.06 | 0.74152267 | 1 |
| WBGene00001345 | fos-1 | 1.104018872 | 1.187422067 | 0.120325377 | 0.169 | 0.18 | 0.567435365 | 1 |
| WBGene00004336 | ret-1 | 0.720977822 | 0.804414197 | 0.120373244 | 0.083 | 0.068 | 0.619147545 | 1 |
| WBGene00019069 | lgc-30 | 0.549484007 | 0.633867888 | 0.121740208 | 0.065 | 0.068 | 0.852468096 | 1 |
| WBGene00021547 | Y44E3A.1 | 0.440699411 | 0.525703234 | 0.122634593 | 0.056 | 0.045 | 0.660819737 | 1 |

| | | | | | | | | |
|---|---|---|---|---|---|---|---|---|
| WBGene00007176 | kcnl-1 | 1.398435472 | 1.483916706 | 0.123323353 | 0.227 | 0.211 | 0.96472459 | 1 |
| WBGene00018476 | F45E12.6 | 4.035881313 | 4.12271487 | 0.125274342 | 0.88 | 0.955 | 0.346151958 | 1 |
| WBGene00004460 | rpn-3 | 0.79968617 | 0.887173845 | 0.126218035 | 0.113 | 0.128 | 0.549257829 | 1 |
| WBGene00012543 | nkcc-1 | 0.48602957 | 0.574563961 | 0.127728128 | 0.047 | 0.053 | 0.740611903 | 1 |
| WBGene00012513 | Y26E6A.2 | 0.86311445 | 0.953116636 | 0.129845708 | 0.119 | 0.12 | 0.837896507 | 1 |
| WBGene00235291 | C01G5.25 | 1.062196539 | 1.152460265 | 0.13022303 | 0.151 | 0.158 | 0.713672612 | 1 |
| WBGene00000408 | cdk-7 | 0.647542556 | 0.738097107 | 0.130642601 | 0.076 | 0.09 | 0.495206566 | 1 |
| WBGene00002845 | let-711 | 0.399218571 | 0.489941396 | 0.13088537 | 0.037 | 0.053 | 0.354517471 | 1 |
| WBGene00007752 | wdfy-3 | 0.556046675 | 0.646990379 | 0.131204031 | 0.061 | 0.06 | 0.975835448 | 1 |
| WBGene00015367 | srt-64 | 1.911153825 | 2.003372487 | 0.133043406 | 0.35 | 0.368 | 0.229815787 | 1 |
| WBGene00003568 | ncx-3 | 0.697355783 | 0.790353815 | 0.134167799 | 0.089 | 0.083 | 0.909331669 | 1 |
| WBGene00006646 | tsp-20 | 0.465740111 | 0.559072461 | 0.134650118 | 0.062 | 0.053 | 0.750582729 | 1 |
| WBGene00019128 | F59G1.4 | 0.782292816 | 0.876989231 | 0.136618049 | 0.083 | 0.098 | 0.513578592 | 1 |
| WBGene00014159 | arrd-4 | 1.741811244 | 1.836766735 | 0.136991816 | 0.333 | 0.301 | 0.989709291 | 1 |
| WBGene00010864 | xbx-3 | 3.268675419 | 3.363851885 | 0.137310616 | 0.787 | 0.872 | 0.201600137 | 1 |
| WBGene00002163 | ist-1 | 0.609748128 | 0.705204631 | 0.137714624 | 0.08 | 0.068 | 0.694877955 | 1 |
| WBGene00010959 | nduo-1 | 1.716228454 | 1.811977501 | 0.138136675 | 0.267 | 0.301 | 0.207918009 | 1 |
| WBGene00010756 | K10G9.2 | 0.66236797 | 0.758435548 | 0.138596219 | 0.087 | 0.083 | 0.959941414 | 1 |
| WBGene00015887 | C17C3.1 | 0.507128833 | 0.603567243 | 0.139131216 | 0.064 | 0.083 | 0.396903587 | 1 |
| WBGene00018968 | F56D2.8 | 0.527734943 | 0.625037173 | 0.140377444 | 0.054 | 0.083 | 0.157904819 | 1 |
| WBGene00007433 | swsn-7 | 0.761056204 | 0.858804149 | 0.141020476 | 0.11 | 0.098 | 0.810886299 | 1 |
| WBGene00019870 | R04E5.7 | 0.920548907 | 1.018704762 | 0.141608964 | 0.124 | 0.128 | 0.800003987 | 1 |
| WBGene00002027 | hsr-9 | 0.530187422 | 0.628374555 | 0.14165409 | 0.065 | 0.053 | 0.643010463 | 1 |
| WBGene00001911 | his-37 | 1.925254157 | 2.023707941 | 0.142038786 | 0.371 | 0.368 | 0.491469454 | 1 |
| WBGene00004698 | rsp-1 | 1.204432843 | 1.303250553 | 0.142563821 | 0.176 | 0.143 | 0.443247892 | 1 |
| WBGene00000102 | akt-1 | 1.096065721 | 1.19536097 | 0.143252763 | 0.169 | 0.173 | 0.705426208 | 1 |
| WBGene00004106 | pqn-15 | 1.055037433 | 1.154565791 | 0.143589068 | 0.142 | 0.128 | 0.813192129 | 1 |
| WBGene00022248 | Y73B6BL.27 | 0.678077269 | 0.777959937 | 0.14410023 | 0.093 | 0.098 | 0.76749658 | 1 |
| WBGene00021050 | W05H9.3 | 2.454200222 | 2.554280879 | 0.144385868 | 0.477 | 0.451 | 0.714006091 | 1 |
| WBGene00011028 | R05D7.3 | 0.555141555 | 0.655885319 | 0.145342529 | 0.063 | 0.075 | 0.558185889 | 1 |
| WBGene00004924 | snt-4 | 2.270691873 | 2.372278575 | 0.146558632 | 0.525 | 0.541 | 0.31396231 | 1 |
| WBGene00008940 | F18H3.4 | 0.477282868 | 0.5788775 | 0.146570071 | 0.055 | 0.083 | 0.189288703 | 1 |
| WBGene00004364 | ric-4 | 1.438712759 | 1.540535755 | 0.146899531 | 0.242 | 0.218 | 0.897388962 | 1 |
| WBGene00004273 | rab-10 | 0.888119024 | 0.990404925 | 0.147567362 | 0.123 | 0.143 | 0.453475257 | 1 |
| WBGene00008549 | din-1 | 0.796767316 | 0.899939054 | 0.148845355 | 0.087 | 0.12 | 0.192674303 | 1 |
| WBGene00019900 | vdac-1 | 1.654991937 | 1.758700781 | 0.149620234 | 0.307 | 0.308 | 0.580993529 | 1 |
| WBGene00004920 | snr-7 | 1.146895414 | 1.250616884 | 0.149638451 | 0.172 | 0.188 | 0.533783451 | 1 |
| WBGene00009975 | F53C11.5 | 0.359040354 | 0.464424846 | 0.152037684 | 0.039 | 0.06 | 0.24734166 | 1 |
| WBGene00003964 | pdi-3 | 0.991505 | 1.097615019 | 0.153084399 | 0.128 | 0.143 | 0.502517788 | 1 |
| WBGene00003741 | nlp-3 | 4.437721878 | 4.544384704 | 0.153881931 | 0.934 | 1 | 0.030460029 | 1 |
| WBGene00001685 | gpd-3 | 1.128477185 | 1.235193051 | 0.15395845 | 0.161 | 0.188 | 0.337133045 | 1 |
| WBGene00020135 | aakg-4 | 0.498180057 | 0.604903023 | 0.153968693 | 0.056 | 0.045 | 0.669553953 | 1 |
| WBGene00012620 | Y38H6C.7 | 0.492205115 | 0.599751192 | 0.155156192 | 0.06 | 0.075 | 0.475616385 | 1 |
| WBGene00016115 | mdt-26 | 2.390116304 | 2.497742064 | 0.15527115 | 0.486 | 0.564 | 0.065739692 | 1 |
| WBGene00007534 | fubl-1 | 1.273836457 | 1.381536292 | 0.155378018 | 0.195 | 0.203 | 0.618556604 | 1 |
| WBGene00016458 | C35E7.6 | 0.624282457 | 0.732307735 | 0.155847533 | 0.088 | 0.09 | 0.827294216 | 1 |
| WBGene00003818 | nsf-1 | 1.175239615 | 1.283808667 | 0.156632033 | 0.187 | 0.188 | 0.761794329 | 1 |
| WBGene00013298 | Y57G11B.6 | 0.873177579 | 0.982471224 | 0.1576774 | 0.12 | 0.128 | 0.681750794 | 1 |
| WBGene00011867 | chc-1 | 0.782715229 | 0.892693033 | 0.158664433 | 0.113 | 0.098 | 0.732766358 | 1 |
| WBGene00007684 | C18E9.4 | 0.390073847 | 0.500149875 | 0.15880614 | 0.046 | 0.06 | 0.443091565 | 1 |
| WBGene00004933 | sod-4 | 1.421241289 | 1.531644423 | 0.159278053 | 0.231 | 0.211 | 0.967500748 | 1 |
| WBGene00004743 | scm-1 | 0.641504283 | 0.752081316 | 0.159528937 | 0.081 | 0.083 | 0.881263952 | 1 |
| WBGene00006708 | ubc-13 | 0.342985623 | 0.45395881 | 0.160100467 | 0.041 | 0.053 | 0.509727938 | 1 |
| WBGene00021370 | nape-2 | 0.650011907 | 0.762750489 | 0.162647394 | 0.075 | 0.098 | 0.309976205 | 1 |
| WBGene00019428 | dgk-5 | 0.802633537 | 0.915407624 | 0.162698616 | 0.1 | 0.083 | 0.624528408 | 1 |
| WBGene00018294 | atg-18 | 0.684924971 | 0.798131453 | 0.163322431 | 0.103 | 0.113 | 0.605854535 | 1 |
| WBGene00022473 | bet-1 | 0.725276954 | 0.83857718 | 0.163457675 | 0.084 | 0.105 | 0.380564668 | 1 |
| WBGene00019185 | H10E21.5 | 1.177369143 | 1.293324507 | 0.167288229 | 0.186 | 0.173 | 0.963264037 | 1 |
| WBGene00021960 | tmem-258 | 0.857825077 | 0.973789189 | 0.167300849 | 0.111 | 0.098 | 0.743383004 | 1 |
| WBGene00001122 | dyf-6 | 1.758971994 | 1.875719019 | 0.168430354 | 0.336 | 0.323 | 0.665103767 | 1 |
| WBGene00015508 | mvb-12 | 0.284193844 | 0.40150561 | 0.169245104 | 0.03 | 0.053 | 0.183887852 | 1 |
| WBGene00007490 | mks-5 | 0.442246449 | 0.560919358 | 0.171208818 | 0.058 | 0.053 | 0.880978147 | 1 |
| WBGene00018042 | mks-3 | 0.72790079 | 0.846707008 | 0.171401141 | 0.085 | 0.098 | 0.541670006 | 1 |
| WBGene00003954 | pcm-1 | 0.854990738 | 0.973929512 | 0.171592379 | 0.113 | 0.113 | 0.900544444 | 1 |

| | | | | | | | | |
|---|---|---|---|---|---|---|---|---|
| WBGene00044988 | W01A8.8 | 0.740632627 | 0.860222027 | 0.172531034 | 0.101 | 0.113 | 0.612394345 | 1 |
| WBGene00006594 | tom-1 | 0.399549025 | 0.519210044 | 0.172634359 | 0.043 | 0.06 | 0.34365343 | 1 |
| WBGene00022027 | vps-20 | 0.464763002 | 0.584785542 | 0.173155923 | 0.049 | 0.06 | 0.546941718 | 1 |
| WBGene00018161 | F38A5.2 | 0.350105822 | 0.470486675 | 0.17367286 | 0.034 | 0.053 | 0.26012499 | 1 |
| WBGene00001837 | hda-4 | 0.913215708 | 1.034084363 | 0.174376609 | 0.131 | 0.098 | 0.371142993 | 1 |
| WBGene00019032 | srsx-17 | 0.93492399 | 1.056233643 | 0.175012834 | 0.115 | 0.173 | 0.050191126 | 1 |
| WBGene00010966 | nduo-3 | 0.72218039 | 0.844198116 | 0.17603437 | 0.087 | 0.09 | 0.788239054 | 1 |
| WBGene00000953 | del-2 | 0.504055211 | 0.626819989 | 0.177112137 | 0.06 | 0.083 | 0.287618279 | 1 |
| WBGene00020778 | T24E12.11 | 0.715751508 | 0.838806938 | 0.177531459 | 0.078 | 0.09 | 0.535753941 | 1 |
| WBGene00019431 | K06A5.3 | 0.557654808 | 0.680849715 | 0.177732682 | 0.06 | 0.075 | 0.457853013 | 1 |
| WBGene00009888 | F49E2.5 | 0.6126021 | 0.735842258 | 0.177797963 | 0.071 | 0.09 | 0.397287716 | 1 |
| WBGene00015729 | C13A10.2 | 0.534077909 | 0.657400819 | 0.177917351 | 0.063 | 0.068 | 0.791621628 | 1 |
| WBGene00011289 | R102.2 | 7.109328331 | 7.233267939 | 0.178807058 | 0.999 | 1 | 4.24E-05 | 0.270263019 |
| WBGene00010560 | eif-2beta | 0.835984896 | 0.96039261 | 0.179482393 | 0.082 | 0.135 | 0.032983606 | 1 |
| WBGene00006528 | tba-1 | 0.978615886 | 1.103835786 | 0.180654129 | 0.13 | 0.15 | 0.358572402 | 1 |
| WBGene00010840 | M03C11.3 | 0.515888123 | 0.641540603 | 0.181278209 | 0.035 | 0.068 | 0.060956438 | 1 |
| WBGene00008610 | F09C3.2 | 0.432309425 | 0.558137376 | 0.181531361 | 0.05 | 0.06 | 0.586910754 | 1 |
| WBGene00012764 | Y41E3.6 | 2.7129522 | 2.839374894 | 0.182389393 | 0.34 | 0.391 | 0.201788619 | 1 |
| WBGene00001080 | dpy-21 | 1.001036211 | 1.127532193 | 0.182495126 | 0.127 | 0.143 | 0.492213197 | 1 |
| WBGene00044007 | ZK1320.13 | 0.736279239 | 0.863837023 | 0.184026983 | 0.08 | 0.105 | 0.320312148 | 1 |
| WBGene00019748 | atg-2 | 1.146157723 | 1.273849202 | 0.184219864 | 0.159 | 0.195 | 0.209591554 | 1 |
| WBGene00018855 | F55A4.3 | 1.138676097 | 1.267004625 | 0.185138931 | 0.176 | 0.226 | 0.09490465 | 1 |
| WBGene00007645 | C17E4.6 | 0.68059014 | 0.809208558 | 0.185557153 | 0.085 | 0.113 | 0.239005724 | 1 |
| WBGene00003052 | lmn-1 | 0.933433225 | 1.062431765 | 0.186105554 | 0.126 | 0.128 | 0.81472768 | 1 |
| WBGene00017811 | F26A1.13 | 0.667312253 | 0.796406657 | 0.186243856 | 0.076 | 0.083 | 0.696891731 | 1 |
| WBGene00006537 | tbb-2 | 1.134276655 | 1.264954761 | 0.188528656 | 0.141 | 0.165 | 0.31431225 | 1 |
| WBGene00003123 | mag-1 | 0.653355451 | 0.784254399 | 0.188847262 | 0.073 | 0.06 | 0.655339752 | 1 |
| WBGene00007108 | gspd-1 | 0.384960653 | 0.515907213 | 0.188915952 | 0.049 | 0.06 | 0.575732261 | 1 |
| WBGene00010993 | vha-20 | 0.335352617 | 0.467264106 | 0.19030805 | 0.038 | 0.053 | 0.392689693 | 1 |
| WBGene00269432 | ZC132.11 | 1.253337796 | 1.386913138 | 0.192708483 | 0.193 | 0.165 | 0.649928797 | 1 |
| WBGene00007646 | nkb-1 | 0.661523503 | 0.795733647 | 0.193624309 | 0.073 | 0.083 | 0.629315542 | 1 |
| WBGene00001137 | eat-6 | 1.283135956 | 1.418568186 | 0.195387406 | 0.192 | 0.218 | 0.317725011 | 1 |
| WBGene00044771 | tmem-218 | 0.457292176 | 0.593292595 | 0.19620713 | 0.052 | 0.06 | 0.660710581 | 1 |
| WBGene00006713 | ubc-18 | 0.781361126 | 0.917911908 | 0.197001136 | 0.092 | 0.113 | 0.372830283 | 1 |
| WBGene00002191 | kin-3 | 0.671105437 | 0.808652481 | 0.198438438 | 0.081 | 0.075 | 0.915122358 | 1 |
| WBGene00002129 | inx-7 | 0.671867715 | 0.809939076 | 0.199194868 | 0.094 | 0.09 | 0.992090421 | 1 |
| WBGene00011481 | imp-2 | 0.404058181 | 0.54225488 | 0.199375693 | 0.045 | 0.053 | 0.640852078 | 1 |
| WBGene00000159 | aps-1 | 0.456090324 | 0.594534874 | 0.199733265 | 0.059 | 0.053 | 0.829280555 | 1 |
| WBGene00011763 | T14B1.1 | 1.602955643 | 1.742633712 | 0.201512858 | 0.25 | 0.278 | 0.218658862 | 1 |
| WBGene00010037 | F54C8.4 | 1.513312404 | 1.653147033 | 0.201738727 | 0.27 | 0.293 | 0.266870149 | 1 |
| WBGene00003628 | nhr-35 | 0.395923549 | 0.536881461 | 0.20335928 | 0.045 | 0.06 | 0.41520885 | 1 |
| WBGene00009069 | F23A7.4 | 1.613482878 | 1.754523896 | 0.203479177 | 0.099 | 0.038 | 0.02597777 | 1 |
| WBGene00012413 | Y7A5A.7 | 1.732774414 | 1.874203327 | 0.204038793 | 0.289 | 0.338 | 0.079321695 | 1 |
| WBGene00022019 | sut-2 | 0.451446208 | 0.592987762 | 0.204201299 | 0.051 | 0.053 | 0.910476753 | 1 |
| WBGene00008587 | F08G12.3 | 0.644108524 | 0.786879274 | 0.205974652 | 0.073 | 0.075 | 0.865205817 | 1 |
| WBGene00000910 | daf-14 | 1.162933409 | 1.307064451 | 0.207937141 | 0.171 | 0.203 | 0.257381061 | 1 |
| WBGene00011629 | T08G11.1 | 0.376450265 | 0.520721215 | 0.208138984 | 0.046 | 0.053 | 0.687871683 | 1 |
| WBGene00004220 | ptr-5 | 0.854330993 | 0.99929401 | 0.209137426 | 0.11 | 0.135 | 0.322239581 | 1 |
| WBGene00016977 | akap-1 | 0.920554931 | 1.066768204 | 0.210941164 | 0.12 | 0.12 | 0.800190337 | 1 |
| WBGene00017769 | hmgs-1 | 0.303367445 | 0.450577131 | 0.212378685 | 0.033 | 0.053 | 0.217406065 | 1 |
| WBGene00000829 | ctb-1 | 1.864907333 | 2.012191818 | 0.212486595 | 0.321 | 0.361 | 0.11979392 | 1 |
| WBGene00003082 | lsm-8 | 0.423889842 | 0.57117699 | 0.212490439 | 0.044 | 0.06 | 0.378977599 | 1 |
| WBGene00014213 | ZK1073.1 | 1.235508973 | 1.383443168 | 0.21342393 | 0.192 | 0.195 | 0.676880664 | 1 |
| WBGene00000249 | bir-1 | 0.408651492 | 0.556666983 | 0.213541214 | 0.043 | 0.053 | 0.580432384 | 1 |
| WBGene00020092 | pcf-11 | 0.572063857 | 0.720133534 | 0.213619389 | 0.075 | 0.083 | 0.652473057 | 1 |
| WBGene00009574 | F40E10.6 | 0.422176147 | 0.570482511 | 0.213960855 | 0.046 | 0.068 | 0.276364589 | 1 |
| WBGene00004751 | sea-2 | 1.242748353 | 1.391240686 | 0.214229152 | 0.208 | 0.211 | 0.612850047 | 1 |
| WBGene00006434 | prdx-2 | 1.491204978 | 1.640410686 | 0.215258335 | 0.247 | 0.286 | 0.138530821 | 1 |
| WBGene00004136 | pqn-51 | 0.610071181 | 0.759628814 | 0.215766056 | 0.077 | 0.09 | 0.530023228 | 1 |
| WBGene00006698 | uaf-2 | 0.631754146 | 0.781322654 | 0.215781745 | 0.08 | 0.083 | 0.793999611 | 1 |
| WBGene00013228 | Y56A3A.7 | 0.612704563 | 0.762398914 | 0.215963298 | 0.07 | 0.075 | 0.761297252 | 1 |
| WBGene00016200 | dpff-1 | 0.377845539 | 0.528481978 | 0.217322444 | 0.037 | 0.06 | 0.183390723 | 1 |
| WBGene00005726 | srv-15 | 0.580744289 | 0.73180103 | 0.217928811 | 0.069 | 0.075 | 0.730849517 | 1 |
| WBGene00006655 | tub-1 | 0.93606789 | 1.087623035 | 0.218647856 | 0.125 | 0.173 | 0.094876722 | 1 |

| | | | | | | | | |
|---|---|---|---|---|---|---|---|---|
| WBGene00019698 | riok-1 | 0.378848855 | 0.531620579 | 0.220403009 | 0.043 | 0.06 | 0.349962389 | 1 |
| WBGene00003661 | nhr-71 | 0.26934492 | 0.422586119 | 0.221080318 | 0.033 | 0.053 | 0.2268841 | 1 |
| WBGene00016415 | ampd-1 | 0.537083125 | 0.690554946 | 0.221413035 | 0.062 | 0.083 | 0.343576216 | 1 |
| WBGene00016494 | C37A2.8 | 0.61586533 | 0.769446911 | 0.221571385 | 0.07 | 0.098 | 0.222146576 | 1 |
| WBGene00012037 | T26E3.4 | 0.389526079 | 0.543234883 | 0.221754929 | 0.042 | 0.053 | 0.542256338 | 1 |
| WBGene00017982 | hpo-18 | 0.749344591 | 0.903404623 | 0.222261644 | 0.1 | 0.105 | 0.703062769 | 1 |
| WBGene00006773 | unc-37 | 0.376616478 | 0.531659899 | 0.223680375 | 0.043 | 0.06 | 0.34996238 | 1 |
| WBGene00016378 | imp-3 | 0.461995001 | 0.617040766 | 0.223683757 | 0.062 | 0.06 | 0.968950983 | 1 |
| WBGene00000840 | cul-5 | 0.452700363 | 0.608173878 | 0.224300869 | 0.049 | 0.068 | 0.345873122 | 1 |
| WBGene00021539 | Y42H9AR.4 | 0.458943201 | 0.614595861 | 0.22455932 | 0.049 | 0.053 | 0.814774455 | 1 |
| WBGene00011270 | R31.2 | 0.754316088 | 0.910284676 | 0.225015108 | 0.094 | 0.098 | 0.810727476 | 1 |
| WBGene00008034 | prcc-1 | 0.665761033 | 0.822121079 | 0.225579863 | 0.079 | 0.09 | 0.582653578 | 1 |
| WBGene00004095 | pqe-1 | 0.430502058 | 0.586996989 | 0.225774461 | 0.05 | 0.068 | 0.368318772 | 1 |
| WBGene00001336 | qars-1 | 1.29644201 | 1.453004459 | 0.225871868 | 0.215 | 0.203 | 0.893870153 | 1 |
| WBGene00000508 | cit-1.2 | 0.463796357 | 0.620439926 | 0.225988899 | 0.052 | 0.053 | 0.936695556 | 1 |
| WBGene00021355 | Y37E3.17 | 0.638862592 | 0.795810251 | 0.226427609 | 0.083 | 0.09 | 0.684317584 | 1 |
| WBGene00045483 | oxy-5 | 0.770933928 | 0.928586642 | 0.227444788 | 0.101 | 0.113 | 0.575868465 | 1 |
| WBGene00015112 | B0281.5 | 0.369080529 | 0.526973118 | 0.227790855 | 0.036 | 0.06 | 0.170749554 | 1 |
| WBGene00016703 | rdl-1 | 0.932852334 | 1.090774055 | 0.227832884 | 0.114 | 0.143 | 0.259483114 | 1 |
| WBGene00001008 | dlk-1 | 0.714299761 | 0.87252045 | 0.228264203 | 0.069 | 0.12 | 0.027822207 | 1 |
| WBGene00304812 | ZK380.6 | 4.340153795 | 4.498485131 | 0.228423833 | 0.609 | 0.699 | 0.066206411 | 1 |
| WBGene00013020 | Y48G10A.3 | 0.577179527 | 0.735971184 | 0.229087936 | 0.064 | 0.083 | 0.385000462 | 1 |
| WBGene00014021 | svop-1 | 0.470496939 | 0.629633794 | 0.229585952 | 0.057 | 0.075 | 0.359777534 | 1 |
| WBGene00195043 | Y57G11C.1135 | 0.434867483 | 0.594278582 | 0.229981602 | 0.06 | 0.068 | 0.650342083 | 1 |
| WBGene00017311 | F09G2.2 | 0.415712386 | 0.575750741 | 0.23088654 | 0.055 | 0.06 | 0.753379573 | 1 |
| WBGene00023491 | tsp-21 | 1.762782483 | 1.923669445 | 0.232110823 | 0.341 | 0.383 | 0.157317899 | 1 |
| WBGene00011723 | T12A7.2 | 0.744015975 | 0.905713162 | 0.23327973 | 0.086 | 0.113 | 0.260540015 | 1 |
| WBGene00017973 | ift-81 | 1.727453904 | 1.889403685 | 0.233644146 | 0.312 | 0.338 | 0.185953837 | 1 |
| WBGene00018164 | sup-36 | 0.814848668 | 0.977422429 | 0.234544358 | 0.106 | 0.135 | 0.242149883 | 1 |
| WBGene00017914 | F29A7.4 | 0.43390814 | 0.596902471 | 0.235151114 | 0.044 | 0.075 | 0.108765995 | 1 |
| WBGene00021382 | Y37F4.1 | 3.022098955 | 3.185255376 | 0.235384959 | 0.539 | 0.617 | 0.036818745 | 1 |
| WBGene00013295 | Y57G11B.2 | 0.437634107 | 0.602046986 | 0.237197645 | 0.044 | 0.068 | 0.21562553 | 1 |
| WBGene00006750 | unc-10 | 0.43744282 | 0.602364158 | 0.237931196 | 0.045 | 0.068 | 0.231174391 | 1 |
| WBGene00004014 | phb-1 | 0.38950591 | 0.554534573 | 0.238086033 | 0.033 | 0.053 | 0.225284292 | 1 |
| WBGene00014079 | ZK792.7 | 0.657831719 | 0.823688375 | 0.239280575 | 0.084 | 0.12 | 0.145921918 | 1 |
| WBGene00020629 | T20F5.7 | 0.346066551 | 0.512864702 | 0.240638866 | 0.041 | 0.053 | 0.487986916 | 1 |
| WBGene00006889 | pfd-3 | 0.388468872 | 0.556816213 | 0.242873874 | 0.038 | 0.06 | 0.214263027 | 1 |
| WBGene00004468 | rpn-12 | 0.333280907 | 0.501778159 | 0.24309015 | 0.03 | 0.053 | 0.167358905 | 1 |
| WBGene00020231 | T05A8.5 | 0.319479632 | 0.488606345 | 0.24399827 | 0.038 | 0.06 | 0.211407759 | 1 |
| WBGene00009141 | ncbp-2 | 0.69049704 | 0.860474551 | 0.245225712 | 0.105 | 0.098 | 0.957804591 | 1 |
| WBGene00022696 | ift-139 | 0.739985689 | 0.910521701 | 0.246031459 | 0.1 | 0.098 | 0.956489527 | 1 |
| WBGene00003901 | paa-1 | 0.697612856 | 0.8684448 | 0.246458397 | 0.086 | 0.075 | 0.732184497 | 1 |
| WBGene00000876 | cyl-1 | 0.503813454 | 0.675885372 | 0.248247303 | 0.061 | 0.068 | 0.699979912 | 1 |
| WBGene00015811 | C16A3.6 | 0.382963157 | 0.555179892 | 0.248456229 | 0.038 | 0.053 | 0.387390926 | 1 |
| WBGene00003674 | nhr-84 | 1.307496988 | 1.479769202 | 0.248536268 | 0.219 | 0.226 | 0.630394105 | 1 |
| WBGene00021926 | Y55F3AM.9 | 0.426735313 | 0.599076599 | 0.248635918 | 0.057 | 0.068 | 0.539635738 | 1 |
| WBGene00077761 | zip-9 | 0.368393994 | 0.540893721 | 0.248864501 | 0.034 | 0.068 | 0.055169649 | 1 |
| WBGene00002263 | lea-1 | 0.696020796 | 0.869465563 | 0.250227905 | 0.055 | 0.105 | 0.023186054 | 1 |
| WBGene00004804 | skn-1 | 0.824955614 | 0.999363668 | 0.251617634 | 0.105 | 0.113 | 0.615060621 | 1 |
| WBGene00012005 | 1-Jun | 1.271290306 | 1.445811046 | 0.251780206 | 0.206 | 0.211 | 0.60184674 | 1 |
| WBGene00003603 | nhr-4 | 0.830766218 | 1.005469491 | 0.252043546 | 0.108 | 0.12 | 0.584760642 | 1 |
| WBGene00000552 | cmd-1 | 2.412489124 | 2.587536634 | 0.252540175 | 0.55 | 0.586 | 0.049385923 | 1 |
| WBGene00005015 | spt-5 | 0.391502075 | 0.566733736 | 0.252805849 | 0.037 | 0.06 | 0.193365563 | 1 |
| WBGene00022712 | ZK355.2 | 0.355954983 | 0.531262467 | 0.252915238 | 0.047 | 0.068 | 0.286967445 | 1 |
| WBGene00019840 | R02F11.2 | 3.010476668 | 3.186046202 | 0.253293296 | 0.759 | 0.782 | 0.027124342 | 1 |
| WBGene00003406 | mrg-1 | 0.528450018 | 0.704274472 | 0.253661068 | 0.059 | 0.075 | 0.415486072 | 1 |
| WBGene00002215 | klc-2 | 0.572566414 | 0.748476555 | 0.253784688 | 0.076 | 0.09 | 0.494803788 | 1 |
| WBGene00009580 | xbx-6 | 1.746032624 | 1.922035791 | 0.253918897 | 0.304 | 0.316 | 0.297902185 | 1 |
| WBGene00021097 | cdc-37 | 0.612217222 | 0.788520865 | 0.254352391 | 0.073 | 0.09 | 0.439274137 | 1 |
| WBGene00016037 | dop-6 | 1.039333241 | 1.215914032 | 0.254752231 | 0.155 | 0.173 | 0.430889169 | 1 |
| WBGene00021461 | nekl-1 | 0.611122733 | 0.788695967 | 0.256184025 | 0.066 | 0.083 | 0.413252825 | 1 |
| WBGene00000509 | cka-1 | 0.749161172 | 0.927104793 | 0.256718379 | 0.095 | 0.128 | 0.197904839 | 1 |
| WBGene00010260 | ddx-17 | 0.958764889 | 1.137117207 | 0.257308005 | 0.14 | 0.135 | 0.942113223 | 1 |
| WBGene00010305 | F59A2.5 | 0.371430388 | 0.550697859 | 0.258628292 | 0.028 | 0.06 | 0.044554442 | 1 |

| | | | | | | | | |
|---|---|---|---|---|---|---|---|---|
| WBGene00011737 | sqst-1 | 1.287195912 | 1.466510536 | 0.258696319 | 0.218 | 0.226 | 0.561911432 | 1 |
| WBGene00010603 | nhr-198 | 0.718260926 | 0.899092364 | 0.260884618 | 0.088 | 0.105 | 0.449177045 | 1 |
| WBGene00002267 | lec-4 | 0.409870957 | 0.590824107 | 0.261060212 | 0.037 | 0.068 | 0.089571576 | 1 |
| WBGene00011512 | T05H10.8 | 1.140135281 | 1.321507175 | 0.261664332 | 0.154 | 0.188 | 0.234314258 | 1 |
| WBGene00001915 | his-41 | 3.035215106 | 3.21690582 | 0.262124292 | 0.696 | 0.774 | 0.019280733 | 1 |
| WBGene00044418 | C08G5.7 | 0.848608261 | 1.030702044 | 0.262705798 | 0.113 | 0.12 | 0.696226154 | 1 |
| WBGene00000170 | aqp-2 | 1.608779572 | 1.791432217 | 0.263512065 | 0.28 | 0.346 | 0.041229735 | 1 |
| WBGene00011051 | pfs-2 | 0.375965737 | 0.558618824 | 0.263512702 | 0.037 | 0.068 | 0.085566911 | 1 |
| WBGene00010262 | droe-4 | 3.32120291 | 3.504659951 | 0.264672564 | 0.331 | 0.391 | 0.120497675 | 1 |
| WBGene00013094 | Y51H1A.3 | 0.710642348 | 0.894190513 | 0.264804027 | 0.08 | 0.098 | 0.428725899 | 1 |
| WBGene00010406 | math-33 | 0.386897597 | 0.570590519 | 0.265012868 | 0.043 | 0.068 | 0.191394258 | 1 |
| WBGene00019552 | K09C4.7 | 0.462970706 | 0.647292112 | 0.265919577 | 0.056 | 0.075 | 0.315290358 | 1 |
| WBGene00002210 | kin-29 | 0.507491499 | 0.692081309 | 0.266306804 | 0.06 | 0.06 | 0.931639837 | 1 |
| WBGene00020710 | T23B3.2 | 1.683546065 | 1.869418498 | 0.268157237 | 0.315 | 0.323 | 0.390391102 | 1 |
| WBGene00007761 | rad-26 | 0.559818957 | 0.746122716 | 0.268779509 | 0.064 | 0.075 | 0.557774277 | 1 |
| WBGene00022069 | eel-1 | 0.831677751 | 1.018239217 | 0.269151302 | 0.114 | 0.098 | 0.730054635 | 1 |
| WBGene00015742 | kxd-1 | 0.237364339 | 0.424150527 | 0.269475507 | 0.024 | 0.053 | 0.060472643 | 1 |
| WBGene00011288 | R102.1 | 4.668217966 | 4.855219182 | 0.269785728 | 0.746 | 0.88 | 0.007019103 | 1 |
| WBGene00022201 | Y71H10B.1 | 0.320184486 | 0.507986021 | 0.270940343 | 0.037 | 0.053 | 0.354976848 | 1 |
| WBGene00008989 | F20G2.6 | 2.888029366 | 3.076368557 | 0.271716017 | 0.698 | 0.722 | 0.028698421 | 1 |
| WBGene00001070 | dpy-8 | 0.50366394 | 0.692250061 | 0.272072261 | 0.045 | 0.083 | 0.06090707 | 1 |
| WBGene00006003 | srx-112 | 0.21895002 | 0.407639933 | 0.272222003 | 0.023 | 0.053 | 0.04695124 | 1 |
| WBGene00022500 | lfi-1 | 1.227500247 | 1.416620764 | 0.272843232 | 0.18 | 0.263 | 0.01648582 | 1 |
| WBGene00000415 | ced-1 | 0.41833476 | 0.608865604 | 0.274877904 | 0.056 | 0.045 | 0.675859464 | 1 |
| WBGene00001604 | gln-3 | 2.019589072 | 2.210509558 | 0.275440038 | 0.409 | 0.436 | 0.242639343 | 1 |
| WBGene00020031 | suco-1 | 0.407410316 | 0.598543654 | 0.275747118 | 0.042 | 0.06 | 0.310614505 | 1 |
| WBGene00008348 | C56G7.3 | 0.301119827 | 0.492672022 | 0.276351402 | 0.03 | 0.053 | 0.177228438 | 1 |
| WBGene00019323 | teg-4 | 0.387333126 | 0.578966018 | 0.276467823 | 0.037 | 0.053 | 0.346765741 | 1 |
| WBGene00022243 | Y73B6BL.22 | 1.400078121 | 1.59228017 | 0.277288944 | 0.213 | 0.248 | 0.199705104 | 1 |
| WBGene00012359 | W09D10.1 | 0.521975056 | 0.714267671 | 0.277419602 | 0.056 | 0.075 | 0.328455264 | 1 |
| WBGene00010074 | irld-34 | 1.782151401 | 1.974983756 | 0.278198283 | 0.319 | 0.376 | 0.061494868 | 1 |
| WBGene00020846 | T27A10.6 | 0.373016859 | 0.566708608 | 0.279438125 | 0.048 | 0.068 | 0.304684098 | 1 |
| WBGene00003085 | ccar-1 | 0.708992005 | 0.903480393 | 0.280587433 | 0.092 | 0.105 | 0.541114314 | 1 |
| WBGene00044143 | K09E9.4 | 0.63635242 | 0.832130843 | 0.28244856 | 0.073 | 0.12 | 0.055359966 | 1 |
| WBGene00001145 | eat-16 | 1.247275081 | 1.44369452 | 0.28337335 | 0.182 | 0.218 | 0.147321033 | 1 |
| WBGene00007886 | ethe-1 | 0.485862161 | 0.68293455 | 0.284315359 | 0.051 | 0.075 | 0.230767211 | 1 |
| WBGene00002085 | ins-2 | 0.303931744 | 0.501166813 | 0.284550057 | 0.029 | 0.053 | 0.146299038 | 1 |
| WBGene00001431 | fkb-6 | 1.114389529 | 1.312738838 | 0.286157563 | 0.167 | 0.211 | 0.144204181 | 1 |
| WBGene00014051 | spv-1 | 1.40120005 | 1.599596515 | 0.286225596 | 0.235 | 0.278 | 0.100425711 | 1 |
| WBGene00006438 | nrfl-1 | 0.537864279 | 0.736342061 | 0.286342912 | 0.063 | 0.09 | 0.210471419 | 1 |
| WBGene00009012 | F21D5.7 | 0.375697805 | 0.574204716 | 0.286384935 | 0.04 | 0.06 | 0.259401721 | 1 |
| WBGene00011500 | T05G5.5 | 1.814721669 | 2.013357928 | 0.286571546 | 0.352 | 0.414 | 0.056183845 | 1 |
| WBGene00007772 | egrh-1 | 1.224707662 | 1.424353147 | 0.28802755 | 0.177 | 0.211 | 0.196200185 | 1 |
| WBGene00007227 | C01G6.5 | 0.693897154 | 0.895066399 | 0.290225872 | 0.073 | 0.105 | 0.162535015 | 1 |
| WBGene00304815 | Y40A1A.6 | 1.603917453 | 1.805185405 | 0.290368276 | 0.077 | 0.053 | 0.358878864 | 1 |
| WBGene00010726 | K09E9.3 | 0.295697757 | 0.497530016 | 0.291182399 | 0.026 | 0.053 | 0.087713381 | 1 |
| WBGene00018564 | F47D12.9 | 0.737431173 | 0.939499176 | 0.291522505 | 0.1 | 0.135 | 0.18750851 | 1 |
| WBGene00009294 | F31D4.9 | 0.315528757 | 0.517708634 | 0.291683906 | 0.038 | 0.053 | 0.378578269 | 1 |
| WBGene00003660 | nhr-70 | 0.298081045 | 0.500621966 | 0.292204781 | 0.031 | 0.053 | 0.195977582 | 1 |
| WBGene00010974 | bbs-5 | 1.530991717 | 1.734073935 | 0.292985709 | 0.267 | 0.301 | 0.133817666 | 1 |
| WBGene00013219 | Y54G11A.11 | 0.770458239 | 0.973856911 | 0.293442255 | 0.103 | 0.128 | 0.305592464 | 1 |
| WBGene00001127 | dyf-11 | 0.641985894 | 0.845930139 | 0.294229351 | 0.071 | 0.098 | 0.234730673 | 1 |
| WBGene00010269 | F58G1.7 | 0.463098056 | 0.667984554 | 0.295588735 | 0.052 | 0.06 | 0.670067077 | 1 |
| WBGene00010427 | hpo-11 | 0.426047236 | 0.631536674 | 0.296458593 | 0.04 | 0.068 | 0.121673569 | 1 |
| WBGene00004276 | rab-14 | 1.090953562 | 1.29744677 | 0.297906728 | 0.171 | 0.18 | 0.515209422 | 1 |
| WBGene00011434 | T04D3.5 | 0.760225054 | 0.966728871 | 0.297922033 | 0.088 | 0.143 | 0.042654067 | 1 |
| WBGene00010550 | K04B12.2 | 0.224921036 | 0.432458495 | 0.299413263 | 0.023 | 0.053 | 0.046717393 | 1 |
| WBGene00021469 | Y39G10AR.11 | 0.478870853 | 0.687708901 | 0.301289616 | 0.049 | 0.068 | 0.332640856 | 1 |
| WBGene00016304 | fbxc-41 | 1.256774048 | 1.465649055 | 0.301342936 | 0.188 | 0.218 | 0.278896951 | 1 |
| WBGene00012130 | slc-17.4 | 0.408427007 | 0.618628955 | 0.303257308 | 0.055 | 0.068 | 0.487371379 | 1 |
| WBGene00020812 | acdh-7 | 0.493804537 | 0.704656437 | 0.30419499 | 0.037 | 0.083 | 0.012748064 | 1 |
| WBGene00044442 | T08A9.13 | 1.85482334 | 2.066331782 | 0.305142181 | 0.38 | 0.361 | 0.634432564 | 1 |
| WBGene00004465 | rpn-9 | 0.230036267 | 0.441624953 | 0.305257948 | 0.03 | 0.053 | 0.168532738 | 1 |
| WBGene00021043 | pck-1 | 0.41309646 | 0.626930164 | 0.308496824 | 0.046 | 0.083 | 0.070573419 | 1 |

| | | | | | | | | |
|---|---|---|---|---|---|---|---|---|
| WBGene00006753 | unc-14 | 1.02527668 | 1.239991629 | 0.309768192 | 0.135 | 0.158 | 0.34682578 | 1 |
| WBGene00012602 | Y38E10A.24 | 0.612652246 | 0.827566079 | 0.310055121 | 0.066 | 0.083 | 0.393859137 | 1 |
| WBGene00015501 | C06A5.3 | 0.531885073 | 0.747251113 | 0.310707518 | 0.057 | 0.083 | 0.205930996 | 1 |
| WBGene00004459 | rpn-2 | 0.419595152 | 0.636116908 | 0.312374865 | 0.049 | 0.068 | 0.339215599 | 1 |
| WBGene00000920 | daf-28 | 3.861433673 | 4.078750229 | 0.313521518 | 0.796 | 0.797 | 0.171709164 | 1 |
| WBGene00007400 | bud-31 | 0.303680558 | 0.521786487 | 0.314660343 | 0.037 | 0.053 | 0.352226282 | 1 |
| WBGene00004345 | rgs-2 | 2.491220074 | 2.710424197 | 0.316244702 | 0.569 | 0.586 | 0.133153542 | 1 |
| WBGene00018586 | ubxn-3 | 0.338317118 | 0.557526008 | 0.316251579 | 0.041 | 0.06 | 0.278348438 | 1 |
| WBGene00006059 | stc-1 | 0.306084923 | 0.526579117 | 0.318105881 | 0.018 | 0.053 | 0.009785368 | 1 |
| WBGene00009137 | bath-39 | 0.559287643 | 0.780058146 | 0.318504511 | 0.075 | 0.06 | 0.627194311 | 1 |
| WBGene00004783 | seu-1 | 0.557904541 | 0.779678592 | 0.319952323 | 0.067 | 0.09 | 0.281019853 | 1 |
| WBGene00004437 | rpl-24.2 | 0.391885028 | 0.614527617 | 0.321205359 | 0.038 | 0.053 | 0.393415641 | 1 |
| WBGene00005182 | srg-25 | 0.365915209 | 0.588927552 | 0.321738802 | 0.035 | 0.06 | 0.136726767 | 1 |
| WBGene00003826 | ntl-3 | 0.314624505 | 0.537829798 | 0.322017169 | 0.041 | 0.053 | 0.477962487 | 1 |
| WBGene00004758 | sek-1 | 0.979318239 | 1.202769091 | 0.322371436 | 0.157 | 0.173 | 0.408134915 | 1 |
| WBGene00003885 | osm-5 | 1.204220086 | 1.428594237 | 0.323703475 | 0.185 | 0.226 | 0.138410628 | 1 |
| WBGene00020852 | nhr-228 | 0.34683541 | 0.57161606 | 0.324289929 | 0.043 | 0.06 | 0.332276969 | 1 |
| WBGene00003904 | pabp-2 | 0.488804629 | 0.713704143 | 0.324461414 | 0.054 | 0.075 | 0.25624775 | 1 |
| WBGene00003048 | lit-1 | 1.964557299 | 2.189604473 | 0.324674442 | 0.36 | 0.444 | 0.016347695 | 1 |
| WBGene00008208 | nhr-167 | 1.74637097 | 1.97160282 | 0.324940873 | 0.331 | 0.346 | 0.31986587 | 1 |
| WBGene00012193 | vps-25 | 1.898350966 | 2.123729409 | 0.325152362 | 0.405 | 0.398 | 0.444927858 | 1 |
| WBGene00001452 | flp-9 | 0.357293007 | 0.58309279 | 0.325760227 | 0.015 | 0.06 | 0.000563654 | 1 |
| WBGene00011302 | R166.2 | 0.479695999 | 0.707574631 | 0.328759373 | 0.059 | 0.083 | 0.263489997 | 1 |
| WBGene00022347 | Y82E9BR.14 | 0.628259135 | 0.857829396 | 0.331199877 | 0.09 | 0.12 | 0.203850306 | 1 |
| WBGene00004947 | sos-1 | 0.259461072 | 0.489168766 | 0.331398151 | 0.03 | 0.053 | 0.167066398 | 1 |
| WBGene00000424 | ced-10 | 0.33014875 | 0.56021091 | 0.331909537 | 0.023 | 0.053 | 0.047896499 | 1 |
| WBGene00001910 | C50F4.6 | 1.409713162 | 1.640136971 | 0.332431286 | 0.2 | 0.278 | 0.018503559 | 1 |
| WBGene00001561 | gei-4 | 0.850476123 | 1.081267288 | 0.332961268 | 0.112 | 0.135 | 0.310578304 | 1 |
| WBGene00022750 | tres-1 | 0.445764513 | 0.677064167 | 0.333694863 | 0.042 | 0.068 | 0.168523038 | 1 |
| WBGene00013272 | aqp-12 | 0.985620789 | 1.217949883 | 0.335180032 | 0.145 | 0.135 | 0.950421292 | 1 |
| WBGene00011299 | ikke-1 | 0.672907666 | 0.90527386 | 0.335233556 | 0.087 | 0.098 | 0.577854536 | 1 |
| WBGene00022434 | Y108G3AL.2 | 0.303746231 | 0.536967499 | 0.336467167 | 0.019 | 0.06 | 0.003216504 | 1 |
| WBGene00006731 | uev-2 | 0.52353257 | 0.756841102 | 0.336593062 | 0.069 | 0.09 | 0.315789289 | 1 |
| WBGene00017536 | F17A9.4 | 0.773031696 | 1.006367103 | 0.336631834 | 0.105 | 0.135 | 0.199012676 | 1 |
| WBGene00003218 | mep-1 | 0.845027172 | 1.078834572 | 0.337312778 | 0.103 | 0.143 | 0.117760717 | 1 |
| WBGene00000242 | bbs-2 | 0.829988093 | 1.0644439 | 0.338248231 | 0.104 | 0.12 | 0.443079044 | 1 |
| WBGene00013703 | rbm-12 | 0.257864438 | 0.492953858 | 0.33916234 | 0.029 | 0.053 | 0.141798191 | 1 |
| WBGene00019322 | ahcy-1 | 0.694994683 | 0.930379087 | 0.339587913 | 0.089 | 0.12 | 0.203500698 | 1 |
| WBGene00007786 | rmd-2 | 0.66391559 | 0.899789935 | 0.340294747 | 0.076 | 0.113 | 0.106175615 | 1 |
| WBGene00010236 | F58B4.3 | 3.684361438 | 3.921934971 | 0.342746158 | 0.861 | 0.955 | 1.90E-06 | 0.01215204 |
| WBGene00009804 | F47B8.3 | 1.115362736 | 1.354470723 | 0.344959907 | 0.183 | 0.188 | 0.545473695 | 1 |
| WBGene00006890 | vem-1 | 2.332780836 | 2.572422429 | 0.345729738 | 0.504 | 0.556 | 0.022715904 | 1 |
| WBGene00219609 | linc-7 | 4.229414772 | 4.469919367 | 0.346974787 | 0.58 | 0.632 | 0.113795149 | 1 |
| WBGene00007883 | C33A12.4 | 4.959142424 | 5.200418016 | 0.3480871 | 0.981 | 0.992 | 1.08E-05 | 0.068931924 |
| WBGene00010579 | K05C4.2 | 0.422569673 | 0.664428703 | 0.348928822 | 0.04 | 0.075 | 0.053751303 | 1 |
| WBGene00018266 | nhr-183 | 0.386750028 | 0.630573912 | 0.351763508 | 0.038 | 0.068 | 0.09289286 | 1 |
| WBGene00007683 | C18E9.2 | 0.857442627 | 1.103483768 | 0.354962335 | 0.103 | 0.143 | 0.119248475 | 1 |
| WBGene00012351 | W09C5.1 | 0.350607151 | 0.598783322 | 0.35804253 | 0.04 | 0.06 | 0.250053807 | 1 |
| WBGene00001089 | dre-1 | 0.151911047 | 0.400400801 | 0.358494937 | 0.013 | 0.053 | 0.001000286 | 1 |
| WBGene00007068 | cTel55X.1 | 0.36796482 | 0.617666216 | 0.360242967 | 0.034 | 0.075 | 0.019360837 | 1 |
| WBGene00235102 | Y54G2A.75 | 0.757700308 | 1.008146541 | 0.361317538 | 0.101 | 0.113 | 0.550853425 | 1 |
| WBGene00011983 | fbxa-81 | 0.250578745 | 0.501516265 | 0.362026315 | 0.025 | 0.053 | 0.071443611 | 1 |
| WBGene00011287 | glb-24 | 0.818238429 | 1.069400046 | 0.362349619 | 0.124 | 0.135 | 0.505265916 | 1 |
| WBGene00044077 | pacs-1 | 0.433292001 | 0.684665197 | 0.362654863 | 0.058 | 0.075 | 0.376879405 | 1 |
| WBGene00002855 | let-721 | 0.406169686 | 0.657768935 | 0.362980988 | 0.044 | 0.068 | 0.202495586 | 1 |
| WBGene00011681 | T10B10.4 | 0.759041677 | 1.011821577 | 0.364684309 | 0.091 | 0.135 | 0.08427319 | 1 |
| WBGene00003884 | osm-3 | 0.743522962 | 0.997380848 | 0.366239513 | 0.095 | 0.135 | 0.112445077 | 1 |
| WBGene00009563 | F39H2.3 | 0.217442665 | 0.471622787 | 0.366704402 | 0.024 | 0.053 | 0.057680056 | 1 |
| WBGene00007495 | C09G9.5 | 0.904593333 | 1.158936738 | 0.36693997 | 0.114 | 0.135 | 0.350318037 | 1 |
| WBGene00044313 | M02D8.7 | 0.227761913 | 0.48214078 | 0.366991131 | 0.022 | 0.053 | 0.034290354 | 1 |
| WBGene00004862 | sma-9 | 0.941271064 | 1.196135337 | 0.367691423 | 0.137 | 0.165 | 0.269025008 | 1 |
| WBGene00015205 | moa-2 | 0.43662386 | 0.691957919 | 0.368369181 | 0.047 | 0.083 | 0.07480343 | 1 |
| WBGene00001684 | gpd-2 | 1.319886529 | 1.576064877 | 0.369587232 | 0.213 | 0.241 | 0.253751002 | 1 |
| WBGene00007979 | imp-1 | 0.358843363 | 0.615735224 | 0.370616613 | 0.023 | 0.06 | 0.013497533 | 1 |

| | | | | | | | | |
|---|---|---|---|---|---|---|---|---|
| WBGene00012434 | Y11D7A.10 | 0.372841142 | 0.630088865 | 0.371130013 | 0.038 | 0.075 | 0.039254097 | 1 |
| WBGene00004378 | rme-8 | 0.362421167 | 0.619870511 | 0.371420891 | 0.041 | 0.06 | 0.270628703 | 1 |
| WBGene00016246 | daf-37 | 0.320667519 | 0.579097906 | 0.372836239 | 0.026 | 0.068 | 0.009316892 | 1 |
| WBGene00020447 | mob-3 | 0.627976977 | 0.887114507 | 0.37385643 | 0.075 | 0.113 | 0.103524049 | 1 |
| WBGene00000490 | che-11 | 1.095874512 | 1.35506088 | 0.373926888 | 0.152 | 0.203 | 0.079187869 | 1 |
| WBGene00007980 | C36E8.1 | 0.493399877 | 0.753244642 | 0.374876755 | 0.051 | 0.083 | 0.1162852 | 1 |
| WBGene00004367 | ric-8 | 0.595573133 | 0.855804064 | 0.375433874 | 0.058 | 0.098 | 0.063383876 | 1 |
| WBGene00019872 | R04E5.9 | 0.784101719 | 1.044510481 | 0.37569043 | 0.091 | 0.098 | 0.700820311 | 1 |
| WBGene00016397 | maph-9 | 0.765913683 | 1.027303279 | 0.377105475 | 0.098 | 0.135 | 0.13757328 | 1 |
| WBGene00020997 | gpa-18 | 1.590968385 | 1.853044859 | 0.378096429 | 0.271 | 0.323 | 0.078951198 | 1 |
| WBGene00006483 | dgk-4 | 1.175263028 | 1.437920727 | 0.37893496 | 0.18 | 0.203 | 0.314455349 | 1 |
| WBGene00001129 | dyf-13 | 1.078745057 | 1.341790036 | 0.379493686 | 0.17 | 0.18 | 0.527129526 | 1 |
| WBGene00015424 | C04E6.8 | 4.66607045 | 4.92975317 | 0.380413753 | 0.902 | 0.977 | 3.40E-05 | 0.217008528 |
| WBGene00017150 | EGAP5.1 | 1.805953762 | 2.069705465 | 0.380513273 | 0.322 | 0.383 | 0.044308926 | 1 |
| WBGene00020932 | dhod-1 | 0.205088143 | 0.4722883 | 0.385488341 | 0.027 | 0.068 | 0.012107485 | 1 |
| WBGene00009164 | hrdl-1 | 0.841882299 | 1.110703333 | 0.387826773 | 0.101 | 0.15 | 0.057620017 | 1 |
| WBGene00006720 | ubc-25 | 0.565955767 | 0.835058274 | 0.388232852 | 0.049 | 0.098 | 0.017327167 | 1 |
| WBGene00001946 | his-72 | 2.092385159 | 2.363145831 | 0.390625079 | 0.437 | 0.421 | 0.377526559 | 1 |
| WBGene00016496 | C37C3.2 | 0.741984897 | 1.013406245 | 0.391578233 | 0.09 | 0.128 | 0.126493581 | 1 |
| WBGene00019808 | R01B10.6 | 0.496998361 | 0.770303853 | 0.394296478 | 0.065 | 0.09 | 0.232676391 | 1 |
| WBGene00003129 | map-1 | 0.630944179 | 0.905249598 | 0.395739068 | 0.068 | 0.098 | 0.205554094 | 1 |
| WBGene00000567 | cnx-1 | 0.725456697 | 1.001308119 | 0.397969479 | 0.105 | 0.113 | 0.603645986 | 1 |
| WBGene00016558 | pks-1 | 0.371054256 | 0.648019883 | 0.399576936 | 0.04 | 0.068 | 0.119273036 | 1 |
| WBGene00077536 | F38B7.10 | 1.126052086 | 1.403222286 | 0.399872073 | 0.151 | 0.226 | 0.014837761 | 1 |
| WBGene00002980 | lgg-1 | 3.251190771 | 3.528413277 | 0.399947534 | 0.782 | 0.902 | 5.76E-06 | 0.036734455 |
| WBGene00009584 | drap-1 | 1.81281255 | 2.091020451 | 0.401369159 | 0.343 | 0.391 | 0.046659831 | 1 |
| WBGene00007009 | wwp-1 | 0.587265555 | 0.866006984 | 0.402138877 | 0.068 | 0.105 | 0.10116103 | 1 |
| WBGene00004350 | rgs-7 | 1.199481376 | 1.478810535 | 0.402986793 | 0.173 | 0.173 | 0.739765778 | 1 |
| WBGene00016931 | C54G6.2 | 0.229851298 | 0.513259588 | 0.408871734 | 0.028 | 0.06 | 0.047877868 | 1 |
| WBGene00008454 | E02A10.4 | 0.558146156 | 0.841972 | 0.409474137 | 0.067 | 0.098 | 0.156751595 | 1 |
| WBGene00008265 | kcnl-4 | 0.440185781 | 0.726723441 | 0.413386461 | 0.055 | 0.09 | 0.085852791 | 1 |
| WBGene00021474 | dot-1.1 | 0.337541773 | 0.624574709 | 0.414100994 | 0.041 | 0.068 | 0.13973701 | 1 |
| WBGene00010737 | K10C8.4 | 0.313386386 | 0.600571863 | 0.414321064 | 0.025 | 0.06 | 0.023811477 | 1 |
| WBGene00021922 | atg-3 | 0.424163532 | 0.711505071 | 0.414546214 | 0.052 | 0.068 | 0.426662288 | 1 |
| WBGene00017676 | F21F3.7 | 0.599726633 | 0.887843445 | 0.415664696 | 0.083 | 0.105 | 0.304953343 | 1 |
| WBGene00018756 | plpp-1.1 | 0.490721902 | 0.779532686 | 0.416665886 | 0.054 | 0.09 | 0.0835484 | 1 |
| WBGene00045190 | R01H2.8 | 1.620768224 | 1.910141463 | 0.417477338 | 0.295 | 0.361 | 0.029029189 | 1 |
| WBGene00018511 | F46F11.8 | 0.722540406 | 1.012641361 | 0.41852721 | 0.085 | 0.113 | 0.23097369 | 1 |
| WBGene00012897 | pisy-1 | 0.469854599 | 0.762156365 | 0.421702309 | 0.048 | 0.105 | 0.006354387 | 1 |
| WBGene00003393 | mog-5 | 0.274660544 | 0.567655987 | 0.422703073 | 0.027 | 0.053 | 0.102747385 | 1 |
| WBGene00018327 | F42A6.5 | 0.542396158 | 0.835592646 | 0.422993118 | 0.051 | 0.098 | 0.026727087 | 1 |
| WBGene00021333 | Y34D9A.8 | 0.768705549 | 1.062113241 | 0.423297822 | 0.104 | 0.128 | 0.304076327 | 1 |
| WBGene00003148 | mbf-1 | 1.038469493 | 1.332028568 | 0.423516221 | 0.166 | 0.203 | 0.185083662 | 1 |
| WBGene00011279 | asd-1 | 1.105616654 | 1.401120252 | 0.426321576 | 0.162 | 0.226 | 0.034880528 | 1 |
| WBGene00016622 | mafr-1 | 0.236862833 | 0.532600614 | 0.42665943 | 0.026 | 0.068 | 0.010016575 | 1 |
| WBGene00003653 | nhr-63 | 0.349914367 | 0.646776498 | 0.428281524 | 0.047 | 0.06 | 0.459354244 | 1 |
| WBGene00009850 | F48F7.5 | 0.400003201 | 0.697680313 | 0.429457294 | 0.057 | 0.068 | 0.538673039 | 1 |
| WBGene00019743 | nhr-203 | 0.306058872 | 0.60513189 | 0.43147116 | 0.037 | 0.068 | 0.083080756 | 1 |
| WBGene00011735 | hip-1 | 0.434989636 | 0.735143983 | 0.433031187 | 0.058 | 0.075 | 0.363842695 | 1 |
| WBGene00012332 | rpi-1 | 0.87845594 | 1.18155861 | 0.437284718 | 0.133 | 0.165 | 0.188502041 | 1 |
| WBGene00023414 | R01H2.7 | 0.231819253 | 0.535565895 | 0.438213774 | 0.029 | 0.06 | 0.055364738 | 1 |
| WBGene00010471 | cdr-6 | 0.239455646 | 0.543409189 | 0.438512269 | 0.025 | 0.068 | 0.006983546 | 1 |
| WBGene00021920 | mrps-25 | 1.113255729 | 1.417356222 | 0.438724273 | 0.166 | 0.195 | 0.2286696 | 1 |
| WBGene00014058 | ZK673.2 | 0.295996957 | 0.600860514 | 0.439825142 | 0.035 | 0.06 | 0.126435535 | 1 |
| WBGene00016294 | C31H2.4 | 0.393339531 | 0.700724944 | 0.443463411 | 0.054 | 0.045 | 0.75071622 | 1 |
| WBGene00235287 | Y55F3C.17 | 0.346032549 | 0.654954825 | 0.445680635 | 0.043 | 0.075 | 0.091200204 | 1 |
| WBGene00010732 | K10C3.5 | 0.459564247 | 0.769021243 | 0.446452074 | 0.052 | 0.09 | 0.065706116 | 1 |
| WBGene00000369 | ccf-1 | 0.232218713 | 0.545217447 | 0.451561722 | 0.022 | 0.053 | 0.0339268 | 1 |
| WBGene00013340 | tcc-1 | 0.260587174 | 0.5736369 | 0.451635287 | 0.022 | 0.068 | 0.002369795 | 1 |
| WBGene00009534 | F38B7.3 | 2.765017288 | 3.078668766 | 0.452503432 | 0.651 | 0.722 | 0.000124343 | 0.79343207 |
| WBGene00013920 | pssy-1 | 0.211186792 | 0.524966488 | 0.452688411 | 0.022 | 0.053 | 0.035167149 | 1 |
| WBGene00011511 | gpcp-2 | 0.565115753 | 0.879429696 | 0.453459167 | 0.07 | 0.105 | 0.115991943 | 1 |
| WBGene00011849 | T19H5.4 | 0.275514166 | 0.590006811 | 0.453716981 | 0.031 | 0.075 | 0.010453941 | 1 |
| WBGene00014036 | ZK643.5 | 0.610548763 | 0.925237337 | 0.453999646 | 0.079 | 0.12 | 0.07039885 | 1 |

| | | | | | | | | |
|---|---|---|---|---|---|---|---|---|
| WBGene00009558 | F39B2.8 | 0.268364344 | 0.584419035 | 0.455970535 | 0.027 | 0.053 | 0.104856305 | 1 |
| WBGene00021883 | Y54G2A.18 | 0.288456193 | 0.605099109 | 0.456819165 | 0.024 | 0.068 | 0.005212671 | 1 |
| WBGene00022419 | Y102A11A.9 | 1.050378198 | 1.367511113 | 0.457526084 | 0.149 | 0.165 | 0.374389003 | 1 |
| WBGene00022465 | ift-20 | 1.783887288 | 2.101082778 | 0.457616361 | 0.369 | 0.444 | 0.008468975 | 1 |
| WBGene00008023 | C39B10.3 | 0.590456231 | 0.907851051 | 0.457903933 | 0.07 | 0.128 | 0.015124281 | 1 |
| WBGene00002999 | lin-10 | 0.490679785 | 0.808163807 | 0.458032623 | 0.058 | 0.068 | 0.615792705 | 1 |
| WBGene00008683 | repo-1 | 0.183160747 | 0.500913113 | 0.458419763 | 0.021 | 0.053 | 0.025748722 | 1 |
| WBGene00007476 | C09E9.1 | 0.902593247 | 1.221389782 | 0.45992618 | 0.128 | 0.188 | 0.028239921 | 1 |
| WBGene00022817 | upp-1 | 1.301476555 | 1.620554732 | 0.460332504 | 0.217 | 0.293 | 0.01766568 | 1 |
| WBGene00016126 | nhr-258 | 0.64770965 | 0.967951834 | 0.46201181 | 0.087 | 0.113 | 0.237818025 | 1 |
| WBGene00022251 | blos-2 | 0.177480209 | 0.497864262 | 0.462216484 | 0.022 | 0.053 | 0.033120835 | 1 |
| WBGene00006590 | toc-1 | 0.230000955 | 0.551695609 | 0.464107281 | 0.028 | 0.053 | 0.116847787 | 1 |
| WBGene00009973 | F53C11.3 | 0.79502434 | 1.117575617 | 0.465343129 | 0.093 | 0.15 | 0.024444875 | 1 |
| WBGene00006802 | unc-69 | 0.94033297 | 1.263227324 | 0.465838083 | 0.128 | 0.18 | 0.047606527 | 1 |
| WBGene00000175 | aqp-7 | 0.333220303 | 0.656241262 | 0.466020736 | 0.027 | 0.068 | 0.012240382 | 1 |
| WBGene00022748 | oaz-1 | 2.001016706 | 2.324181037 | 0.466227578 | 0.419 | 0.534 | 0.000863392 | 1 |
| WBGene00017290 | F09E5.12 | 0.534980828 | 0.858410766 | 0.466610768 | 0.062 | 0.12 | 0.010282631 | 1 |
| WBGene00006864 | npr-16 | 0.574356768 | 0.900984767 | 0.471224594 | 0.059 | 0.105 | 0.03525893 | 1 |
| WBGene00017298 | toca-1 | 0.249961318 | 0.577733285 | 0.472874991 | 0.026 | 0.06 | 0.03049043 | 1 |
| WBGene00010096 | rskd-1 | 0.207774509 | 0.535663155 | 0.473043324 | 0.017 | 0.06 | 0.001296764 | 1 |
| WBGene00007630 | har-1 | 0.82702476 | 1.155198117 | 0.473454075 | 0.11 | 0.135 | 0.266531714 | 1 |
| WBGene00019466 | tos-1 | 3.628564691 | 3.956918167 | 0.473713931 | 0.811 | 0.925 | 1.17E-05 | 0.074957364 |
| WBGene00019759 | M03F4.6 | 0.503327511 | 0.832558777 | 0.474980315 | 0.059 | 0.105 | 0.030414194 | 1 |
| WBGene00015527 | C06E2.1 | 0.588592612 | 0.918645674 | 0.476165915 | 0.068 | 0.12 | 0.023448104 | 1 |
| WBGene00077771 | C41G7.9 | 0.478275897 | 0.809130881 | 0.477322845 | 0.061 | 0.083 | 0.280672547 | 1 |
| WBGene00021906 | Y55B1AR.2 | 0.214054513 | 0.545617621 | 0.478344451 | 0.021 | 0.053 | 0.025173211 | 1 |
| WBGene00021349 | arl-13 | 0.894624713 | 1.22701818 | 0.479542406 | 0.128 | 0.165 | 0.132145334 | 1 |
| WBGene00013538 | ari-1.4 | 0.304830945 | 0.637465048 | 0.47988957 | 0.029 | 0.06 | 0.057831734 | 1 |
| WBGene00018152 | acs-4 | 0.341786646 | 0.674974798 | 0.480688895 | 0.031 | 0.068 | 0.029403507 | 1 |
| WBGene00012648 | Y39A1A.9 | 0.831246082 | 1.165194989 | 0.481786432 | 0.099 | 0.135 | 0.138185141 | 1 |
| WBGene00021312 | szy-2 | 0.276861814 | 0.610934251 | 0.481964649 | 0.033 | 0.068 | 0.037342694 | 1 |
| WBGene00007848 | cytb-5.1 | 1.001233332 | 1.335965638 | 0.482916638 | 0.13 | 0.18 | 0.072592888 | 1 |
| WBGene00004109 | stam-1 | 0.393313169 | 0.733117786 | 0.490234435 | 0.044 | 0.068 | 0.20536178 | 1 |
| WBGene00009403 | F35C11.4 | 0.53257059 | 0.872590176 | 0.490544571 | 0.057 | 0.105 | 0.021251837 | 1 |
| WBGene00004443 | rpl-29 | 2.101292911 | 2.441866979 | 0.491344519 | 0.446 | 0.541 | 0.002096304 | 1 |
| WBGene00002107 | ins-24 | 2.544441582 | 2.886449183 | 0.493412671 | 0.513 | 0.617 | 0.000408911 | 1 |
| WBGene00018696 | pho-6 | 1.104056593 | 1.447398338 | 0.495337433 | 0.108 | 0.143 | 0.177772899 | 1 |
| WBGene00045171 | K09E9.5 | 0.16194138 | 0.506122677 | 0.496548652 | 0.019 | 0.06 | 0.003228114 | 1 |
| WBGene00015791 | C15C7.5 | 0.261551617 | 0.607507876 | 0.499109379 | 0.025 | 0.053 | 0.070484289 | 1 |
| WBGene00007364 | stdh-3 | 0.362089187 | 0.709417309 | 0.501088558 | 0.029 | 0.083 | 0.001796633 | 1 |
| WBGene00001094 | dars-1 | 0.233631777 | 0.581983752 | 0.502565666 | 0.024 | 0.075 | 0.001365621 | 1 |
| WBGene00012132 | ifta-2 | 1.533541055 | 1.881950548 | 0.502648649 | 0.237 | 0.361 | 0.00061748 | 1 |
| WBGene00010375 | H09F14.1 | 0.165283686 | 0.514226047 | 0.503417413 | 0.013 | 0.06 | 0.000137597 | 0.878003414 |
| WBGene00271642 | R106.8 | 1.427309626 | 1.776371266 | 0.503589497 | 0.256 | 0.323 | 0.024950862 | 1 |
| WBGene00000834 | cua-1 | 0.313289317 | 0.662560441 | 0.503891718 | 0.021 | 0.068 | 0.001664059 | 1 |
| WBGene00011624 | T08G5.3 | 0.325134518 | 0.674986352 | 0.504729505 | 0.03 | 0.09 | 0.000703527 | 1 |
| WBGene00011382 | dop-5 | 0.711454694 | 1.062218862 | 0.506045725 | 0.09 | 0.128 | 0.117120548 | 1 |
| WBGene00012290 | W05H12.2 | 0.412753959 | 0.763597923 | 0.506160847 | 0.052 | 0.083 | 0.122782108 | 1 |
| WBGene00014202 | mmcm-1 | 0.154984154 | 0.507387157 | 0.508410065 | 0.01 | 0.053 | 0.000243561 | 1 |
| WBGene00000390 | cdc-42 | 0.590159631 | 0.944859307 | 0.511723464 | 0.073 | 0.113 | 0.089940699 | 1 |
| WBGene00008576 | F08G2.4 | 0.527297073 | 0.883270307 | 0.513560818 | 0.061 | 0.09 | 0.163320626 | 1 |
| WBGene00044026 | cisd-1 | 0.531373581 | 0.887361968 | 0.51358268 | 0.075 | 0.105 | 0.164636325 | 1 |
| WBGene00021956 | Y57E12AL.1 | 0.512789745 | 0.868882226 | 0.513732857 | 0.064 | 0.098 | 0.11303901 | 1 |
| WBGene00016964 | C56C10.7 | 0.398482623 | 0.755885661 | 0.515623591 | 0.052 | 0.06 | 0.629730663 | 1 |
| WBGene00007129 | B0272.3 | 0.536510527 | 0.895298756 | 0.517621999 | 0.07 | 0.113 | 0.058986977 | 1 |
| WBGene00010632 | K07C5.9 | 2.456123457 | 2.815951069 | 0.519121512 | 0.532 | 0.729 | 3.08E-06 | 0.019668034 |
| WBGene00006373 | syx-5 | 0.282367214 | 0.64222427 | 0.51916399 | 0.03 | 0.06 | 0.069346185 | 1 |
| WBGene00001494 | frm-8 | 0.746065745 | 1.106069904 | 0.519376215 | 0.118 | 0.135 | 0.395983255 | 1 |
| WBGene00004902 | snf-3 | 0.136483927 | 0.496640961 | 0.519596767 | 0.014 | 0.06 | 0.000249974 | 1 |
| WBGene00006798 | unc-64 | 0.767831432 | 1.128005448 | 0.519621267 | 0.099 | 0.158 | 0.022705377 | 1 |
| WBGene00003903 | pab-2 | 1.513000553 | 1.873474895 | 0.520054545 | 0.258 | 0.338 | 0.017494213 | 1 |
| WBGene00014166 | ZK945.4 | 0.429095266 | 0.789843569 | 0.520449788 | 0.058 | 0.068 | 0.578174259 | 1 |
| WBGene00018886 | npr-28 | 0.700818098 | 1.06188542 | 0.520910034 | 0.099 | 0.135 | 0.137739743 | 1 |
| WBGene00003635 | nhr-45 | 0.482104222 | 0.84330629 | 0.521104433 | 0.051 | 0.098 | 0.024567124 | 1 |

| | | | | | | | | |
|---|---|---|---|---|---|---|---|---|
| WBGene00045495 | F11C1.7 | 0.425542706 | 0.786821123 | 0.521214581 | 0.051 | 0.09 | 0.055416635 | 1 |
| WBGene00001909 | his-35 | 1.834085711 | 2.196813179 | 0.523305118 | 0.362 | 0.436 | 0.016905597 | 1 |
| WBGene00020844 | T27A10.2 | 1.030450516 | 1.394579017 | 0.525326383 | 0.145 | 0.135 | 0.962564457 | 1 |
| WBGene00003391 | mog-3 | 0.504556848 | 0.869427234 | 0.526396697 | 0.061 | 0.098 | 0.085329033 | 1 |
| WBGene00000004 | aat-3 | 0.532335442 | 0.897254314 | 0.526466648 | 0.067 | 0.128 | 0.00961548 | 1 |
| WBGene00016303 | fbxc-32 | 0.073196359 | 0.43880848 | 0.527466794 | 0.008 | 0.053 | 3.96E-05 | 0.252937461 |
| WBGene00019166 | tat-2 | 0.956841929 | 1.322493239 | 0.527523331 | 0.125 | 0.135 | 0.527133439 | 1 |
| WBGene00004776 | ser-1 | 0.193467249 | 0.559520867 | 0.52810374 | 0.017 | 0.053 | 0.006707493 | 1 |
| WBGene00000970 | dhs-6 | 0.199730483 | 0.566444617 | 0.529056662 | 0.022 | 0.06 | 0.009505166 | 1 |
| WBGene00018252 | F40H3.6 | 0.419709328 | 0.786493042 | 0.529157046 | 0.051 | 0.09 | 0.053834152 | 1 |
| WBGene00219313 | C17E4.20 | 0.457848349 | 0.825816589 | 0.530865955 | 0.054 | 0.098 | 0.032975442 | 1 |
| WBGene00009562 | flp-22 | 2.398826373 | 2.769044858 | 0.534112373 | 0.536 | 0.617 | 0.003313513 | 1 |
| WBGene00000545 | clp-4 | 0.205517558 | 0.575991072 | 0.534480301 | 0.019 | 0.053 | 0.013669105 | 1 |
| WBGene00008000 | slc-17.2 | 0.675422896 | 1.047013174 | 0.536091451 | 0.079 | 0.143 | 0.008856136 | 1 |
| WBGene00020366 | acdh-10 | 0.416144542 | 0.787964486 | 0.53642279 | 0.045 | 0.098 | 0.009472114 | 1 |
| WBGene00010153 | srxa-2 | 1.256209534 | 1.628218675 | 0.536695743 | 0.179 | 0.271 | 0.005404945 | 1 |
| WBGene00017472 | cst-1 | 0.331477701 | 0.704511127 | 0.538173474 | 0.036 | 0.083 | 0.009392709 | 1 |
| WBGene00014066 | rev-1 | 0.282598917 | 0.657686263 | 0.541136654 | 0.02 | 0.068 | 0.001038911 | 1 |
| WBGene00016406 | C34D10.1 | 0.263213603 | 0.639525655 | 0.542903531 | 0.028 | 0.053 | 0.121718752 | 1 |
| WBGene00015002 | B0034.1 | 0.229035839 | 0.607853508 | 0.546518372 | 0.022 | 0.053 | 0.033165171 | 1 |
| WBGene00018145 | F37C4.5 | 1.400971387 | 1.78018035 | 0.54708289 | 0.234 | 0.338 | 0.001425973 | 1 |
| WBGene00010505 | K02D3.2 | 0.576979133 | 0.956365201 | 0.547338398 | 0.065 | 0.105 | 0.072486358 | 1 |
| WBGene00019741 | nhr-201 | 0.20125567 | 0.584012525 | 0.552201417 | 0.021 | 0.053 | 0.02567617 | 1 |
| WBGene00016005 | ift-74 | 1.124889274 | 1.511294189 | 0.557464454 | 0.165 | 0.226 | 0.028555316 | 1 |
| WBGene00016162 | crh-2 | 0.917960964 | 1.305776288 | 0.559499245 | 0.144 | 0.203 | 0.033270993 | 1 |
| WBGene00011725 | lips-5 | 0.605581783 | 0.994817551 | 0.561548512 | 0.065 | 0.098 | 0.127141387 | 1 |
| WBGene00019339 | K02F6.5 | 1.800760249 | 2.190370651 | 0.562088995 | 0.354 | 0.451 | 0.00067248 | 1 |
| WBGene00022280 | Y74C10AL.2 | 0.157762814 | 0.547406952 | 0.562137666 | 0.021 | 0.053 | 0.025316036 | 1 |
| WBGene00009925 | F52B11.2 | 0.287533048 | 0.677416309 | 0.562482648 | 0.029 | 0.06 | 0.055364738 | 1 |
| WBGene00001496 | frm-10 | 1.530266505 | 1.9201854 | 0.562534055 | 0.271 | 0.353 | 0.005853337 | 1 |
| WBGene00019365 | rei-2 | 0.869509215 | 1.261546573 | 0.565590352 | 0.11 | 0.195 | 0.001694015 | 1 |
| WBGene00007703 | gbf-1 | 0.419862446 | 0.812128042 | 0.565919631 | 0.051 | 0.083 | 0.104805215 | 1 |
| WBGene00012928 | aakb-2 | 0.260060517 | 0.652730072 | 0.566502421 | 0.03 | 0.06 | 0.066445158 | 1 |
| WBGene00007191 | lgc-20 | 1.576545814 | 1.9702813 | 0.568040233 | 0.289 | 0.391 | 0.003049839 | 1 |
| WBGene00013734 | Y111B2A.10 | 0.737931818 | 1.134283756 | 0.571814975 | 0.098 | 0.15 | 0.040005133 | 1 |
| WBGene00017016 | snap-1 | 1.162248868 | 1.55905588 | 0.572471507 | 0.179 | 0.248 | 0.020225693 | 1 |
| WBGene00017563 | F18C5.10 | 0.154556435 | 0.552598367 | 0.574253122 | 0.015 | 0.068 | 6.62E-05 | 0.422280057 |
| WBGene00020596 | oga-1 | 0.995323687 | 1.39454934 | 0.575960871 | 0.132 | 0.203 | 0.020974515 | 1 |
| WBGene00004134 | myrf-1 | 1.124935257 | 1.524230008 | 0.576060557 | 0.135 | 0.233 | 0.001743531 | 1 |
| WBGene00000514 | ckb-4 | 0.099535976 | 0.499544101 | 0.577089738 | 0.009 | 0.053 | 9.89E-05 | 0.631174704 |
| WBGene00002001 | hars-1 | 0.220763567 | 0.621605341 | 0.578292441 | 0.019 | 0.06 | 0.003310456 | 1 |
| WBGene00003907 | paf-2 | 0.323269105 | 0.72415682 | 0.578358719 | 0.037 | 0.053 | 0.33114897 | 1 |
| WBGene00021416 | mksr-2 | 1.233760388 | 1.635244009 | 0.579218428 | 0.197 | 0.308 | 0.0006399 | 1 |
| WBGene00021011 | W03G1.5 | 0.723777382 | 1.12700674 | 0.581736995 | 0.069 | 0.143 | 0.002508566 | 1 |
| WBGene00003915 | pan-1 | 1.221303314 | 1.624799297 | 0.582121654 | 0.161 | 0.248 | 0.004348204 | 1 |
| WBGene00019301 | K02D10.1 | 0.595753588 | 0.999743961 | 0.582834907 | 0.064 | 0.135 | 0.002397478 | 1 |
| WBGene00004321 | rcan-1 | 0.785055485 | 1.18970068 | 0.583779617 | 0.099 | 0.18 | 0.002478133 | 1 |
| WBGene00015295 | acl-12 | 0.171384527 | 0.576409156 | 0.584327024 | 0.02 | 0.06 | 0.004676548 | 1 |
| WBGene00044004 | C24A8.6 | 1.147191318 | 1.555014293 | 0.588364183 | 0.194 | 0.263 | 0.015150115 | 1 |
| WBGene00021328 | npr-23 | 0.256050964 | 0.66398744 | 0.588527931 | 0.022 | 0.075 | 0.000479896 | 1 |
| WBGene00001119 | dyf-3 | 0.956611526 | 1.364569127 | 0.588558408 | 0.143 | 0.165 | 0.261117192 | 1 |
| WBGene00011280 | pelo-1 | 0.247742144 | 0.65632946 | 0.589466894 | 0.024 | 0.075 | 0.001168404 | 1 |
| WBGene00008975 | F20D1.3 | 0.22448431 | 0.63369122 | 0.590360779 | 0.026 | 0.083 | 0.000583859 | 1 |
| WBGene00022568 | ZC239.5 | 0.773970033 | 1.183543867 | 0.59089014 | 0.1 | 0.143 | 0.090965773 | 1 |
| WBGene00021876 | Y54G2A.11 | 0.294039237 | 0.705298866 | 0.593322227 | 0.037 | 0.09 | 0.003755267 | 1 |
| WBGene00008310 | C54E10.6 | 0.65342215 | 1.065215374 | 0.594092043 | 0.072 | 0.12 | 0.04229485 | 1 |
| WBGene00003372 | mlc-4 | 0.230726014 | 0.642847293 | 0.594565326 | 0.027 | 0.075 | 0.003115784 | 1 |
| WBGene00007200 | vamp-8 | 0.80242271 | 1.216269857 | 0.597055227 | 0.103 | 0.165 | 0.017150075 | 1 |
| WBGene00016633 | C44B11.1 | 0.366557723 | 0.781414872 | 0.598512353 | 0.042 | 0.105 | 0.001369262 | 1 |
| WBGene00015161 | pho-5 | 0.320414065 | 0.737009874 | 0.601020708 | 0.023 | 0.068 | 0.003467792 | 1 |
| WBGene00003640 | nhr-50 | 0.632250816 | 1.050852602 | 0.603914722 | 0.084 | 0.105 | 0.316577242 | 1 |
| WBGene00017084 | E01A2.1 | 0.646901767 | 1.066636011 | 0.605548512 | 0.083 | 0.143 | 0.014631443 | 1 |
| WBGene00020347 | ech-1.2 | 0.218176 | 0.639762199 | 0.608220318 | 0.025 | 0.06 | 0.021882858 | 1 |
| WBGene00008973 | F20D1.1 | 0.279866902 | 0.701959095 | 0.608950313 | 0.035 | 0.075 | 0.020007236 | 1 |

| | | | | | | | | |
|---|---|---|---|---|---|---|---|---|
| WBGene00012929 | rsks-1 | 0.603867339 | 1.026872415 | 0.610267325 | 0.071 | 0.128 | 0.016125672 | 1 |
| WBGene00018847 | 6-Mar | 0.735391647 | 1.1585958 | 0.610554533 | 0.085 | 0.165 | 0.001866275 | 1 |
| WBGene00016293 | C31H2.3 | 1.513777186 | 1.93943617 | 0.614096105 | 0.235 | 0.316 | 0.011502173 | 1 |
| WBGene00021804 | Y53G8AM.4 | 0.582388568 | 1.008129598 | 0.614214472 | 0.071 | 0.12 | 0.032011079 | 1 |
| WBGene00016961 | vps-32.1 | 1.167514294 | 1.593342494 | 0.614340232 | 0.182 | 0.278 | 0.002894238 | 1 |
| WBGene00008214 | gem-1 | 0.310501237 | 0.738602027 | 0.617618886 | 0.031 | 0.068 | 0.031800755 | 1 |
| WBGene00012471 | Y17G7B.20 | 1.567443351 | 1.997751905 | 0.620804017 | 0.263 | 0.376 | 0.000367177 | 1 |
| WBGene00010882 | atg-7 | 0.441452745 | 0.873138164 | 0.622790414 | 0.059 | 0.09 | 0.128374576 | 1 |
| WBGene00009672 | F43G9.12 | 0.276345008 | 0.708764424 | 0.623849348 | 0.025 | 0.075 | 0.001608376 | 1 |
| WBGene00015208 | B0495.9 | 0.53725001 | 0.969752526 | 0.623969235 | 0.072 | 0.098 | 0.236872945 | 1 |
| WBGene00077526 | C25A1.16 | 0.462700607 | 0.898302254 | 0.628440336 | 0.054 | 0.09 | 0.065434618 | 1 |
| WBGene00022733 | tmem-17 | 0.459195895 | 0.894797783 | 0.628440683 | 0.058 | 0.09 | 0.116485225 | 1 |
| WBGene00004108 | pqn-18 | 0.883141844 | 1.32062305 | 0.631151967 | 0.119 | 0.218 | 0.000808843 | 1 |
| WBGene00020854 | T27C4.1 | 1.697300086 | 2.134984583 | 0.631445254 | 0.327 | 0.421 | 0.002038709 | 1 |
| WBGene00018509 | xrep-4 | 0.639523936 | 1.077889174 | 0.632427354 | 0.089 | 0.15 | 0.013144517 | 1 |
| WBGene00001178 | egl-9 | 0.814898376 | 1.253520329 | 0.632797715 | 0.103 | 0.211 | 0.000126449 | 0.806868069 |
| WBGene00019342 | K02F6.8 | 1.652798192 | 2.094588156 | 0.637368191 | 0.301 | 0.436 | 0.000117086 | 0.747125549 |
| WBGene00009116 | snx-27 | 0.553555369 | 0.995695096 | 0.637872791 | 0.06 | 0.105 | 0.035359359 | 1 |
| WBGene00022503 | ZC13.2 | 0.479176403 | 0.921597448 | 0.638278648 | 0.051 | 0.113 | 0.004040377 | 1 |
| WBGene00020294 | T06F4.1 | 0.181414102 | 0.625490831 | 0.640667294 | 0.02 | 0.075 | 0.00018407 | 1 |
| WBGene00004990 | spp-5 | 0.311323663 | 0.75636928 | 0.642065104 | 0.037 | 0.083 | 0.011308504 | 1 |
| WBGene00017735 | did-2 | 0.641170806 | 1.086358339 | 0.642269847 | 0.069 | 0.12 | 0.026610867 | 1 |
| WBGene00001812 | haf-2 | 0.195272982 | 0.641141848 | 0.643252801 | 0.008 | 0.068 | 2.27E-07 | 0.001446199 |
| WBGene00020482 | T13C2.7 | 0.87743505 | 1.323778316 | 0.643937216 | 0.122 | 0.195 | 0.007279493 | 1 |
| WBGene00012339 | W07G4.5 | 0.875527625 | 1.323687247 | 0.646557664 | 0.126 | 0.18 | 0.039710361 | 1 |
| WBGene00022254 | nlp-71 | 2.623047672 | 3.071994565 | 0.647693457 | 0.509 | 0.602 | 0.00457503 | 1 |
| WBGene00007107 | pfd-4 | 0.476854237 | 0.928646863 | 0.651798981 | 0.057 | 0.09 | 0.093051935 | 1 |
| WBGene00016634 | C44B11.4 | 2.603889774 | 3.056446444 | 0.652901264 | 0.586 | 0.767 | 3.62E-07 | 0.002312182 |
| WBGene00021039 | W05F2.7 | 1.366633294 | 1.819570767 | 0.653450647 | 0.205 | 0.308 | 0.001596833 | 1 |
| WBGene00000370 | ccg-1 | 0.060718062 | 0.515491592 | 0.656099518 | 0.005 | 0.053 | 9.93E-07 | 0.006334665 |
| WBGene00021352 | pcyt-2.1 | 0.34659552 | 0.801418389 | 0.656170699 | 0.04 | 0.083 | 0.021650376 | 1 |
| WBGene00012136 | T28F4.1 | 0.314895212 | 0.770271453 | 0.656969045 | 0.037 | 0.083 | 0.010696195 | 1 |
| WBGene00003962 | pdi-1 | 0.391572099 | 0.848760539 | 0.659583494 | 0.05 | 0.075 | 0.192519066 | 1 |
| WBGene00007260 | olrn-1 | 0.285041904 | 0.744332715 | 0.662616575 | 0.035 | 0.083 | 0.006671669 | 1 |
| WBGene00009112 | tag-353 | 0.19462028 | 0.654859713 | 0.663985148 | 0.021 | 0.068 | 0.001578982 | 1 |
| WBGene00020896 | T28D9.4 | 0.228679108 | 0.690965043 | 0.666937625 | 0.024 | 0.06 | 0.016573854 | 1 |
| WBGene00007653 | cysl-1 | 2.160908861 | 2.623413775 | 0.667253546 | 0.456 | 0.579 | 1.27E-05 | 0.080750319 |
| WBGene00007363 | stdh-1 | 0.129580307 | 0.59257732 | 0.667963494 | 0.013 | 0.068 | 1.43E-05 | 0.091304952 |
| WBGene00006363 | syd-1 | 0.203430157 | 0.666570507 | 0.668170285 | 0.022 | 0.06 | 0.009417895 | 1 |
| WBGene00016626 | clhm-1 | 0.448038604 | 0.916770261 | 0.676236837 | 0.055 | 0.105 | 0.015519431 | 1 |
| WBGene00003602 | nhr-3 | 0.651217373 | 1.120002149 | 0.676313472 | 0.08 | 0.15 | 0.004965262 | 1 |
| WBGene00011990 | T24D5.2 | 1.540341248 | 2.012338344 | 0.68094787 | 0.289 | 0.338 | 0.03803978 | 1 |
| WBGene00007485 | C09G1.3 | 1.866794356 | 2.33916698 | 0.681489642 | 0.339 | 0.519 | 1.81E-06 | 0.011548944 |
| WBGene00000293 | cap-2 | 0.552094611 | 1.025356884 | 0.682773134 | 0.07 | 0.143 | 0.002167296 | 1 |
| WBGene00019781 | M60.6 | 0.180654605 | 0.656658986 | 0.68672916 | 0.017 | 0.068 | 0.000211616 | 1 |
| WBGene00016305 | fbxc-57 | 0.99440717 | 1.470617987 | 0.687026985 | 0.163 | 0.226 | 0.024798838 | 1 |
| WBGene00020865 | T27E4.7 | 0.319668838 | 0.796067415 | 0.687297865 | 0.029 | 0.083 | 0.001879345 | 1 |
| WBGene00000221 | atf-4 | 1.205224603 | 1.681814826 | 0.687574351 | 0.215 | 0.263 | 0.045978102 | 1 |
| WBGene00020008 | R11F4.2 | 0.703506594 | 1.18191196 | 0.690193049 | 0.086 | 0.158 | 0.003673557 | 1 |
| WBGene00011309 | R186.8 | 0.112714694 | 0.592182915 | 0.691726425 | 0.01 | 0.068 | 2.23E-06 | 0.014211263 |
| WBGene00007030 | epc-1 | 1.297665969 | 1.777217227 | 0.691846222 | 0.21 | 0.293 | 0.006671699 | 1 |
| WBGene00016150 | hint-3 | 0.189448848 | 0.669374779 | 0.692386761 | 0.018 | 0.06 | 0.002159345 | 1 |
| WBGene00015517 | C06A8.8 | 0.631799388 | 1.112760197 | 0.693879774 | 0.07 | 0.158 | 0.000341375 | 1 |
| WBGene00003610 | nhr-11 | 0.713449426 | 1.194443622 | 0.693927941 | 0.062 | 0.173 | 3.37E-06 | 0.02149522 |
| WBGene00012472 | Y17G7B.21 | 0.575353776 | 1.05638723 | 0.693984578 | 0.07 | 0.12 | 0.026777044 | 1 |
| WBGene00011263 | R13H4.5 | 0.071575029 | 0.554366344 | 0.696520637 | 0.007 | 0.06 | 9.54E-07 | 0.006089557 |
| WBGene00017504 | F16B4.2 | 0.236164706 | 0.719719081 | 0.697621499 | 0.025 | 0.083 | 0.000354245 | 1 |
| WBGene00006704 | ubc-7 | 0.998546332 | 1.484980986 | 0.701776863 | 0.164 | 0.248 | 0.004154389 | 1 |
| WBGene00000896 | dad-1 | 0.689997901 | 1.176482688 | 0.701849189 | 0.095 | 0.165 | 0.005801847 | 1 |
| WBGene00015935 | C17H12.10 | 0.137443934 | 0.624854074 | 0.703184192 | 0.009 | 0.053 | 0.000103633 | 0.661280225 |
| WBGene00015206 | B0495.7 | 0.47192578 | 0.959618018 | 0.703591172 | 0.05 | 0.09 | 0.045995252 | 1 |
| WBGene00000899 | daf-3 | 0.544088306 | 1.031997483 | 0.703904149 | 0.075 | 0.105 | 0.155312949 | 1 |
| WBGene00016192 | bcl-7 | 0.087255278 | 0.576469474 | 0.705786894 | 0.01 | 0.06 | 2.45E-05 | 0.156486361 |
| WBGene00016475 | srab-9 | 0.009043983 | 0.498602955 | 0.706284301 | 0.001 | 0.06 | 1.86E-12 | 1.19E-08 |

| | | | | | | | | |
|---|---|---|---|---|---|---|---|---|
| WBGene00017004 | nrd-1 | 0.360185753 | 0.85052717 | 0.707413131 | 0.035 | 0.105 | 0.000136846 | 0.873212477 |
| WBGene00009512 | bigr-1 | 0.21759161 | 0.710029314 | 0.710437433 | 0.02 | 0.083 | 3.17E-05 | 0.202552345 |
| WBGene00007484 | C09G1.2 | 1.893208659 | 2.386722824 | 0.711990438 | 0.355 | 0.481 | 0.000155832 | 0.994362505 |
| WBGene00005087 | srd-9 | 0.275333115 | 0.769699018 | 0.713219236 | 0.029 | 0.06 | 0.055729099 | 1 |
| WBGene00010318 | F59B10.2 | 0.578863691 | 1.073302424 | 0.713324309 | 0.069 | 0.15 | 0.000632319 | 1 |
| WBGene00017180 | F02E11.2 | 2.612843025 | 3.107765619 | 0.714022372 | 0.57 | 0.759 | 1.55E-08 | 9.91E-05 |
| WBGene00010018 | fbxa-69 | 0.221315004 | 0.716789222 | 0.714818197 | 0.021 | 0.06 | 0.006826458 | 1 |
| WBGene00010470 | cdr-4 | 0.07953161 | 0.578151293 | 0.719356144 | 0.01 | 0.068 | 2.02E-06 | 0.012907421 |
| WBGene00006523 | tam-1 | 0.719203837 | 1.218579334 | 0.720446553 | 0.102 | 0.165 | 0.012540681 | 1 |
| WBGene00017385 | F11G11.5 | 0.116796154 | 0.618490355 | 0.723791736 | 0.014 | 0.053 | 0.001568356 | 1 |
| WBGene00007929 | dmd-10 | 0.36513058 | 0.868125736 | 0.725668617 | 0.037 | 0.105 | 0.000241263 | 1 |
| WBGene00020807 | T25F10.4 | 0.111804433 | 0.61626071 | 0.727776569 | 0.015 | 0.06 | 0.000410589 | 1 |
| WBGene00016321 | tppp-1 | 1.770177904 | 2.276886242 | 0.731025607 | 0.305 | 0.436 | 7.10E-05 | 0.453131999 |
| WBGene00013647 | Y105C5B.5 | 0.700004342 | 1.207608621 | 0.732318176 | 0.062 | 0.105 | 0.05357146 | 1 |
| WBGene00007969 | acs-19 | 0.355056652 | 0.865879594 | 0.736961724 | 0.033 | 0.098 | 0.000278371 | 1 |
| WBGene00016316 | C32D5.8 | 1.564719795 | 2.081444543 | 0.745476232 | 0.277 | 0.451 | 9.27E-07 | 0.005912021 |
| WBGene00014473 | MTCE.36 | 0.64884261 | 1.165968618 | 0.746055127 | 0.052 | 0.135 | 0.000174949 | 1 |
| WBGene00007048 | nfx-1 | 0.129102509 | 0.648467472 | 0.749285257 | 0.015 | 0.068 | 5.99E-05 | 0.382079747 |
| WBGene00004208 | ptc-1 | 1.779558697 | 2.301993533 | 0.753714146 | 0.319 | 0.459 | 5.68E-05 | 0.362261462 |
| WBGene00011298 | R107.2 | 0.104521599 | 0.627078214 | 0.753889837 | 0.016 | 0.083 | 1.67E-06 | 0.010629777 |
| WBGene00014255 | ZK1320.5 | 0.330653072 | 0.854252966 | 0.75539497 | 0.021 | 0.06 | 0.006738601 | 1 |
| WBGene00016506 | abhd-5.1 | 0.64951086 | 1.173200747 | 0.755524804 | 0.077 | 0.15 | 0.00234275 | 1 |
| WBGene00008530 | F02E9.5 | 0.608850748 | 1.136285579 | 0.760927615 | 0.086 | 0.15 | 0.009159456 | 1 |
| WBGene00020751 | srbc-69 | 0.309853773 | 0.837787744 | 0.761647722 | 0.028 | 0.12 | 3.53E-07 | 0.002252686 |
| WBGene00020857 | srt-62 | 0.144537944 | 0.675404956 | 0.765879206 | 0.017 | 0.068 | 0.000201422 | 1 |
| WBGene00021345 | Y37B11A.3 | 0.341568231 | 0.873272634 | 0.767087305 | 0.04 | 0.105 | 0.000661742 | 1 |
| WBGene00002232 | kpc-1 | 0.820201329 | 1.353749388 | 0.769747138 | 0.093 | 0.188 | 0.000534127 | 1 |
| WBGene00000684 | col-110 | 0.021239983 | 0.556917144 | 0.772818783 | 0.003 | 0.06 | 7.28E-10 | 4.65E-06 |
| WBGene00004859 | sma-5 | 0.462786334 | 0.998651129 | 0.773089483 | 0.054 | 0.12 | 0.001798072 | 1 |
| WBGene00001757 | gst-9 | 0.031325549 | 0.571402079 | 0.779165732 | 0.003 | 0.053 | 2.15E-08 | 0.000137149 |
| WBGene00015623 | C09B8.4 | 0.229683072 | 0.769805728 | 0.779232279 | 0.026 | 0.06 | 0.029960829 | 1 |
| WBGene00019767 | rpa-2 | 0.8342889 | 1.376097545 | 0.781664646 | 0.097 | 0.18 | 0.001212389 | 1 |
| WBGene00018413 | cyp-33C4 | 0.264631363 | 0.809758596 | 0.786452356 | 0.031 | 0.09 | 0.000702767 | 1 |
| WBGene00000066 | act-4 | 2.094679321 | 2.639964166 | 0.786679742 | 0.442 | 0.609 | 3.57E-07 | 0.002280482 |
| WBGene00016395 | spcs-1 | 0.515698795 | 1.061659549 | 0.787654872 | 0.065 | 0.128 | 0.005402385 | 1 |
| WBGene00001034 | dnj-16 | 0.273473952 | 0.824920784 | 0.795569611 | 0.031 | 0.105 | 3.54E-05 | 0.226203559 |
| WBGene00021610 | nhr-237 | 0.911360249 | 1.463969162 | 0.797246139 | 0.128 | 0.241 | 0.000160612 | 1 |
| WBGene00044437 | Y47G6A.32 | 0.33686806 | 0.892136243 | 0.801082653 | 0.037 | 0.105 | 0.000253151 | 1 |
| WBGene00011292 | allo-1 | 1.609993633 | 2.170124507 | 0.808098035 | 0.289 | 0.444 | 4.86E-06 | 0.031011475 |
| WBGene00015626 | C09B9.1 | 1.727978549 | 2.289926912 | 0.810720116 | 0.32 | 0.414 | 0.000848078 | 1 |
| WBGene00000092 | ags-3 | 1.930581791 | 2.496853202 | 0.816956956 | 0.374 | 0.534 | 6.24E-06 | 0.039805413 |
| WBGene00011935 | scrm-1 | 1.111527668 | 1.68472791 | 0.826953148 | 0.178 | 0.248 | 0.020720462 | 1 |
| WBGene00005647 | srp-6 | 0.296578526 | 0.872442278 | 0.830795779 | 0.03 | 0.09 | 0.000523103 | 1 |
| WBGene00008419 | wdr-23 | 0.780087809 | 1.356246659 | 0.831221515 | 0.11 | 0.18 | 0.006637762 | 1 |
| WBGene00020736 | T23F2.3 | 2.618853131 | 3.196972419 | 0.834049829 | 0.601 | 0.797 | 1.29E-09 | 8.25E-06 |
| WBGene00000279 | cah-1 | 0.313007411 | 0.892947459 | 0.836676631 | 0.031 | 0.105 | 3.93E-05 | 0.250793147 |
| WBGene00006139 | str-79 | 1.583035211 | 2.16351893 | 0.837460984 | 0.295 | 0.474 | 2.86E-07 | 0.001823011 |
| WBGene00006498 | ten-1 | 0.639101052 | 1.223929682 | 0.843729365 | 0.058 | 0.15 | 3.89E-05 | 0.24797996 |
| WBGene00008227 | C50B8.5 | 0.190914491 | 0.779785805 | 0.849561724 | 0.024 | 0.068 | 0.004275896 | 1 |
| WBGene00022824 | ZK813.5 | 1.260267977 | 1.850296306 | 0.851230944 | 0.209 | 0.361 | 4.52E-06 | 0.028821241 |
| WBGene00021237 | crtc-1 | 0.340839064 | 0.931408897 | 0.85201217 | 0.043 | 0.098 | 0.0045079 | 1 |
| WBGene00018795 | F54C4.4 | 0.189710432 | 0.781008013 | 0.853062088 | 0.021 | 0.083 | 5.03E-05 | 0.320698962 |
| WBGene00194713 | F19B10.13 | 0.209567259 | 0.801260504 | 0.853632911 | 0.023 | 0.075 | 0.000665078 | 1 |
| WBGene00000973 | dhs-9 | 0.313386183 | 0.908124194 | 0.858025579 | 0.036 | 0.113 | 4.15E-05 | 0.264896226 |
| WBGene00007882 | C33A12.3 | 1.381833736 | 1.97701616 | 0.858666731 | 0.243 | 0.338 | 0.001138409 | 1 |
| WBGene00000843 | cup-2 | 0.343573217 | 0.939055643 | 0.859099544 | 0.044 | 0.113 | 0.000555123 | 1 |
| WBGene00020775 | T24E12.6 | 0.062088301 | 0.658167019 | 0.859959811 | 0.007 | 0.06 | 8.60E-07 | 0.005486713 |
| WBGene00008206 | set-6 | 0.420422025 | 1.016830575 | 0.860435658 | 0.049 | 0.12 | 0.000684484 | 1 |
| WBGene00000534 | cpi-2 | 1.183815276 | 1.780246408 | 0.860468236 | 0.185 | 0.308 | 0.000127265 | 0.812075063 |
| WBGene00019664 | K11H12.8 | 0.213896212 | 0.816226575 | 0.868979027 | 0.022 | 0.09 | 1.46E-05 | 0.093319466 |
| WBGene00043308 | tmem-107 | 0.563960374 | 1.167891664 | 0.871288678 | 0.047 | 0.158 | 3.08E-07 | 0.001962877 |
| WBGene00249817 | F42G8.19 | 1.010230054 | 1.615154924 | 0.87272211 | 0.148 | 0.241 | 0.001188833 | 1 |
| WBGene00004057 | pmk-3 | 0.280821085 | 0.88582175 | 0.872831459 | 0.035 | 0.105 | 0.000120664 | 0.769956218 |
| WBGene00008681 | scrm-4 | 1.750896262 | 2.357016273 | 0.874446334 | 0.299 | 0.489 | 1.30E-07 | 0.000829138 |

| | | | | | | | | |
|---|---|---|---|---|---|---|---|---|
| WBGene00021854 | Y54F10AR.1 | 0.473891483 | 1.084427074 | 0.880816669 | 0.044 | 0.18 | 6.63E-10 | 4.23E-06 |
| WBGene00021915 | Y55D5A.1 | 0.213894144 | 0.831252361 | 0.890659639 | 0.027 | 0.083 | 0.000698835 | 1 |
| WBGene00003019 | lin-33 | 0.830064155 | 1.448084609 | 0.891615043 | 0.095 | 0.211 | 2.35E-05 | 0.150175164 |
| WBGene00008021 | C39B10.1 | 0.295291574 | 0.913981365 | 0.892580693 | 0.028 | 0.12 | 2.75E-07 | 0.001754465 |
| WBGene00000224 | atgp-1 | 0.380218326 | 1.005875414 | 0.902632378 | 0.054 | 0.135 | 0.000130101 | 0.830175584 |
| WBGene00021556 | Y45G5AM.3 | 0.027061107 | 0.653085469 | 0.903162243 | 0.003 | 0.068 | 2.45E-11 | 1.56E-07 |
| WBGene00018758 | F53E10.1 | 0.071367828 | 0.698378399 | 0.904585041 | 0.008 | 0.09 | 4.28E-11 | 2.73E-07 |
| WBGene00013692 | Y105E8B.7 | 0.270312503 | 0.897815858 | 0.905295979 | 0.034 | 0.105 | 7.18E-05 | 0.457909892 |
| WBGene00002219 | klp-7 | 0.130427535 | 0.758053027 | 0.905472185 | 0.013 | 0.083 | 1.16E-07 | 0.000743039 |
| WBGene00010329 | pcdr-1 | 0.968139649 | 1.59800838 | 0.908708495 | 0.13 | 0.271 | 3.31E-06 | 0.021112452 |
| WBGene00003040 | lin-59 | 0.664652345 | 1.296592053 | 0.911696283 | 0.082 | 0.203 | 3.84E-06 | 0.024477611 |
| WBGene00017907 | F28F9.3 | 1.838003017 | 2.479231394 | 0.925096999 | 0.328 | 0.466 | 1.57E-05 | 0.100169762 |
| WBGene00019994 | cdh-1 | 0.194016587 | 0.84251237 | 0.93558165 | 0.02 | 0.098 | 4.65E-07 | 0.002968293 |
| WBGene00021059 | atat-2 | 0.424199472 | 1.07377128 | 0.937134026 | 0.055 | 0.12 | 0.002088984 | 1 |
| WBGene00007553 | C13G3.1 | 1.026218678 | 1.678695029 | 0.941324396 | 0.06 | 0.195 | 2.69E-08 | 0.000171881 |
| WBGene00011474 | aldo-1 | 1.591549531 | 2.246843948 | 0.945390006 | 0.311 | 0.511 | 5.30E-09 | 3.38E-05 |
| WBGene00194982 | Y17D7B.10 | 1.799120669 | 2.454914759 | 0.946110881 | 0.238 | 0.338 | 0.002852411 | 1 |
| WBGene00020481 | T13C2.6 | 0.394613283 | 1.050456724 | 0.946182081 | 0.044 | 0.135 | 9.98E-06 | 0.063679685 |
| WBGene00007024 | plin-1 | 0.2133014 | 0.870224056 | 0.947739057 | 0.021 | 0.075 | 0.000276174 | 1 |
| WBGene00019776 | M6.4 | 0.492287419 | 1.15001765 | 0.948904142 | 0.068 | 0.128 | 0.008714517 | 1 |
| WBGene00009726 | tbc-13 | 0.361159212 | 1.027637271 | 0.96152459 | 0.042 | 0.113 | 0.000301399 | 1 |
| WBGene00017934 | F30B5.4 | 0.350360034 | 1.019342874 | 0.965138225 | 0.042 | 0.12 | 7.53E-05 | 0.480681444 |
| WBGene00003707 | nhr-117 | 0.478287906 | 1.147769154 | 0.965857277 | 0.056 | 0.165 | 1.23E-06 | 0.007837341 |
| WBGene00019305 | snap-29 | 0.369685258 | 1.039996818 | 0.967055163 | 0.04 | 0.075 | 0.053545806 | 1 |
| WBGene00000446 | ceh-23 | 0.993950711 | 1.665932891 | 0.969465358 | 0.14 | 0.286 | 2.68E-06 | 0.017103576 |
| WBGene00007981 | glb-11 | 1.326526053 | 1.998818177 | 0.969912513 | 0.206 | 0.361 | 2.40E-06 | 0.015288564 |
| WBGene00002783 | let-607 | 0.303958884 | 0.97719733 | 0.971277767 | 0.029 | 0.12 | 4.58E-07 | 0.002919627 |
| WBGene00015947 | C18A11.1 | 0.0492314 | 0.723220721 | 0.972361051 | 0.002 | 0.068 | 1.64E-12 | 1.05E-08 |
| WBGene00008579 | F08G2.8 | 0.242481782 | 0.918607005 | 0.975442506 | 0.027 | 0.09 | 0.000162108 | 1 |
| WBGene00003387 | mod-5 | 0.499172729 | 1.178082589 | 0.979459888 | 0.062 | 0.165 | 8.79E-06 | 0.056093266 |
| WBGene00010918 | M117.1 | 1.085091319 | 1.765681161 | 0.981883589 | 0.151 | 0.301 | 3.81E-06 | 0.024318593 |
| WBGene00004131 | pqn-44 | 2.186749688 | 2.875525527 | 0.993693488 | 0.42 | 0.647 | 1.15E-09 | 7.37E-06 |
| WBGene00007217 | tads-1 | 0.522017717 | 1.212124747 | 0.995613989 | 0.049 | 0.158 | 5.28E-07 | 0.003366518 |
| WBGene00021611 | nhr-238 | 0.753114359 | 1.44548064 | 0.9988734 | 0.085 | 0.226 | 1.77E-07 | 0.001127167 |
| WBGene00020694 | dhhc-11 | 1.648305154 | 2.344472897 | 1.00435775 | 0.293 | 0.451 | 1.60E-06 | 0.010219199 |
| WBGene00012009 | T25B9.3 | 0.680830184 | 1.378459558 | 1.006466439 | 0.092 | 0.241 | 9.61E-08 | 0.000613461 |
| WBGene00020737 | T23F2.4 | 0.222130249 | 0.923871874 | 1.012399163 | 0.021 | 0.098 | 9.49E-07 | 0.006053976 |
| WBGene00018634 | pigx-1 | 0.338313511 | 1.04220765 | 1.015504584 | 0.033 | 0.135 | 5.46E-08 | 0.000348388 |
| WBGene00012140 | T28F4.5 | 2.478305393 | 3.189375005 | 1.025856604 | 0.472 | 0.684 | 5.74E-12 | 3.66E-08 |
| WBGene00007010 | alx-1 | 0.418337549 | 1.132298584 | 1.030028044 | 0.04 | 0.135 | 1.67E-06 | 0.010655685 |
| WBGene00010827 | M02B1.2 | 0.270262585 | 0.985494782 | 1.031861943 | 0.034 | 0.135 | 9.55E-08 | 0.000609083 |
| WBGene00011425 | T04B2.7 | 0.005601615 | 0.72216771 | 1.033786351 | 0.001 | 0.083 | 3.09E-17 | 1.97E-13 |
| WBGene00005096 | srd-18 | 0.228941373 | 0.947296489 | 1.036367363 | 0.024 | 0.113 | 1.39E-07 | 0.000886623 |
| WBGene00000298 | cat-4 | 2.253228976 | 2.977475413 | 1.044866744 | 0.499 | 0.647 | 3.11E-07 | 0.001981451 |
| WBGene00021979 | Y58A7A.5 | 0 | 0.729940574 | 1.053081647 | 0 | 0.06 | 3.09E-14 | 1.97E-10 |
| WBGene00004793 | shw-3 | 0.307676607 | 1.04847305 | 1.068743354 | 0.027 | 0.128 | 2.10E-08 | 0.000133889 |
| WBGene00004405 | rop-1 | 0.550853571 | 1.295625785 | 1.07447918 | 0.058 | 0.165 | 2.56E-06 | 0.016338851 |
| WBGene00000996 | dif-1 | 0.964652746 | 1.709425957 | 1.074480617 | 0.144 | 0.271 | 2.04E-05 | 0.129870634 |
| WBGene00010828 | M02B1.3 | 0.325446122 | 1.070463857 | 1.07483339 | 0.028 | 0.15 | 7.04E-11 | 4.50E-07 |
| WBGene00019304 | K02D10.4 | 0.74700746 | 1.493173671 | 1.076490293 | 0.099 | 0.226 | 4.28E-06 | 0.027334391 |
| WBGene00012002 | T24H10.4 | 0.699053151 | 1.445510114 | 1.07690976 | 0.07 | 0.195 | 4.53E-07 | 0.002887467 |
| WBGene00009165 | gpx-1 | 1.567507019 | 2.314304596 | 1.077401162 | 0.278 | 0.391 | 0.000198887 | 1 |
| WBGene00010879 | taap-1 | 0.536862763 | 1.285375737 | 1.079875956 | 0.07 | 0.18 | 7.37E-06 | 0.047007838 |
| WBGene00001425 | fis-2 | 0.233137905 | 0.982244679 | 1.080732629 | 0.024 | 0.135 | 1.88E-10 | 1.20E-06 |
| WBGene00007343 | C05E7.1 | 0.159245695 | 0.909590203 | 1.082518301 | 0.008 | 0.12 | 2.45E-16 | 1.56E-12 |
| WBGene00012444 | Y15E3A.5 | 1.771610479 | 2.528519941 | 1.091989526 | 0.239 | 0.459 | 3.10E-09 | 1.98E-05 |
| WBGene00021811 | ral-1 | 0.552986757 | 1.312449928 | 1.095673751 | 0.058 | 0.18 | 1.43E-07 | 0.000913227 |
| WBGene00018748 | F53C3.4 | 0.107452492 | 0.870464721 | 1.100793959 | 0.012 | 0.113 | 7.33E-13 | 4.68E-09 |
| WBGene00007913 | cyp-36A1 | 3.793678419 | 4.561172988 | 1.107260609 | 0.754 | 0.977 | 2.88E-25 | 1.84E-21 |
| WBGene00019686 | K12H6.6 | 0.626584203 | 1.39559884 | 1.109453602 | 0.069 | 0.218 | 5.38E-09 | 3.43E-05 |
| WBGene00006840 | unc-116 | 0.641001318 | 1.415910313 | 1.117957364 | 0.076 | 0.195 | 1.86E-06 | 0.011881472 |
| WBGene00012907 | cpt-1 | 1.069302431 | 1.848320243 | 1.123885134 | 0.167 | 0.278 | 0.000276428 | 1 |
| WBGene00016017 | C23G10.11 | 0.496599833 | 1.277191621 | 1.126155901 | 0.05 | 0.12 | 0.000902341 | 1 |
| WBGene00017675 | F21F3.6 | 1.32287029 | 2.107083647 | 1.131380722 | 0.212 | 0.406 | 8.86E-09 | 5.65E-05 |

| | | | | | | | | |
|---|---|---|---|---|---|---|---|---|
| WBGene00019465 | acl-14 | 0.718502311 | 1.506977195 | 1.137528806 | 0.093 | 0.211 | 1.10E-05 | 0.070430442 |
| WBGene00015955 | C18B2.4 | 0.712222984 | 1.50155598 | 1.138766799 | 0.097 | 0.233 | 3.75E-07 | 0.002392277 |
| WBGene00194710 | mks-2 | 1.163704905 | 1.953175452 | 1.138965242 | 0.156 | 0.331 | 2.81E-08 | 0.000179554 |
| WBGene00002007 | hsp-3 | 0.936072277 | 1.726100453 | 1.139769732 | 0.112 | 0.256 | 6.54E-07 | 0.004170311 |
| WBGene00011631 | T08G11.4 | 0.610435582 | 1.404039227 | 1.144928043 | 0.073 | 0.218 | 2.10E-08 | 0.000134197 |
| WBGene00003408 | mrp-2 | 1.336858854 | 2.135139874 | 1.151676068 | 0.196 | 0.391 | 6.82E-09 | 4.35E-05 |
| WBGene00011480 | enpl-1 | 0.757338404 | 1.564400161 | 1.164343994 | 0.077 | 0.211 | 1.56E-07 | 0.000997201 |
| WBGene00018259 | F41B4.3 | 0.066105632 | 0.873512699 | 1.164842172 | 0.008 | 0.098 | 1.70E-12 | 1.08E-08 |
| WBGene00007366 | C06B3.7 | 0.508689424 | 1.324033905 | 1.176293439 | 0.049 | 0.188 | 8.28E-10 | 5.28E-06 |
| WBGene00006470 | tag-120 | 0.7219222 | 1.540524416 | 1.180993357 | 0.067 | 0.233 | 7.28E-11 | 4.65E-07 |
| WBGene00013878 | atfs-1 | 0.26018268 | 1.078800416 | 1.181015748 | 0.029 | 0.135 | 6.32E-09 | 4.03E-05 |
| WBGene00000802 | crt-1 | 1.184883552 | 2.008625447 | 1.188408347 | 0.169 | 0.293 | 2.85E-05 | 0.181832895 |
| WBGene00000442 | ceh-19 | 1.961900367 | 2.788934344 | 1.193157817 | 0.378 | 0.669 | 7.49E-15 | 4.78E-11 |
| WBGene00008343 | C56A3.4 | 0.245613221 | 1.073147125 | 1.19387906 | 0.03 | 0.143 | 1.73E-09 | 1.10E-05 |
| WBGene00001749 | gst-1 | 0.336686254 | 1.165122188 | 1.195180414 | 0.043 | 0.09 | 0.012282659 | 1 |
| WBGene00003647 | nhr-57 | 0.318094466 | 1.148762359 | 1.198400451 | 0.026 | 0.128 | 8.45E-09 | 5.39E-05 |
| WBGene00016783 | irg-2 | 0.11203163 | 0.954599423 | 1.215568375 | 0.002 | 0.06 | 5.31E-11 | 3.39E-07 |
| WBGene00003930 | pat-3 | 0.103105378 | 0.950869901 | 1.223065673 | 0.009 | 0.113 | 2.32E-14 | 1.48E-10 |
| WBGene00016175 | C27H5.6 | 1.506528047 | 2.366292935 | 1.24037854 | 0.211 | 0.451 | 1.07E-11 | 6.81E-08 |
| WBGene00000993 | dhs-30 | 0.266509929 | 1.131838942 | 1.248405876 | 0.023 | 0.143 | 1.12E-11 | 7.14E-08 |
| WBGene00003989 | pfn-1 | 0.272383379 | 1.152280425 | 1.269423105 | 0.024 | 0.135 | 1.68E-10 | 1.07E-06 |
| WBGene00008476 | E03H4.8 | 0.060380018 | 0.953353192 | 1.288287969 | 0.005 | 0.083 | 3.46E-12 | 2.21E-08 |
| WBGene00001779 | gst-31 | 0.207504317 | 1.100693938 | 1.288600236 | 0.019 | 0.128 | 1.72E-11 | 1.10E-07 |
| WBGene00189933 | F38B7.11 | 0.760501456 | 1.656266198 | 1.292315352 | 0.103 | 0.256 | 7.77E-08 | 0.000496025 |
| WBGene00011423 | ipla-7 | 0.424490691 | 1.32689376 | 1.301892431 | 0.037 | 0.173 | 3.73E-11 | 2.38E-07 |
| WBGene00006923 | vhp-1 | 1.501290944 | 2.414508875 | 1.31749498 | 0.254 | 0.549 | 3.63E-16 | 2.31E-12 |
| WBGene00008745 | F13E6.1 | 0.591883325 | 1.512333341 | 1.327928673 | 0.081 | 0.226 | 1.78E-08 | 0.000113667 |
| WBGene00018750 | F53C3.6 | 0.329630268 | 1.252721766 | 1.331739527 | 0.037 | 0.18 | 2.58E-12 | 1.65E-08 |
| WBGene00010837 | M03B6.5 | 0.326947973 | 1.260449936 | 1.346758653 | 0.029 | 0.165 | 1.27E-12 | 8.10E-09 |
| WBGene00019006 | spg-20 | 0.726223971 | 1.662789718 | 1.351178759 | 0.088 | 0.256 | 6.38E-10 | 4.07E-06 |
| WBGene00016488 | C36C9.5 | 1.031402292 | 1.970407904 | 1.35469874 | 0.148 | 0.331 | 6.02E-09 | 3.84E-05 |
| WBGene00010724 | K09E9.1 | 0.452427173 | 1.395624167 | 1.360745627 | 0.055 | 0.195 | 1.23E-09 | 7.82E-06 |
| WBGene00015561 | C07A12.7 | 0.583869824 | 1.531318273 | 1.366879178 | 0.066 | 0.226 | 1.59E-10 | 1.02E-06 |
| WBGene00011772 | T14G8.4 | 1.058341352 | 2.015068201 | 1.380265081 | 0.148 | 0.383 | 4.80E-13 | 3.07E-09 |
| WBGene00018290 | vps-60 | 0.567042476 | 1.525242856 | 1.382390936 | 0.067 | 0.203 | 2.48E-08 | 0.000158136 |
| WBGene00007574 | C14B1.3 | 1.207884144 | 2.176325127 | 1.397165004 | 0.206 | 0.436 | 2.68E-12 | 1.71E-08 |
| WBGene00001527 | gcs-1 | 0.356298028 | 1.325974687 | 1.398947707 | 0.033 | 0.135 | 6.07E-08 | 0.000387225 |
| WBGene00020738 | T23F2.5 | 1.59635044 | 2.575637129 | 1.412812051 | 0.297 | 0.571 | 1.31E-15 | 8.38E-12 |
| WBGene00017082 | DC2.5 | 0.365582613 | 1.346726382 | 1.41549125 | 0.045 | 0.143 | 2.11E-06 | 0.01349248 |
| WBGene00017467 | F14F9.4 | 0.399053137 | 1.382013893 | 1.418112608 | 0.031 | 0.203 | 4.74E-17 | 3.03E-13 |
| WBGene00007216 | C01A2.4 | 0.279260199 | 1.274370543 | 1.435640759 | 0.021 | 0.15 | 6.79E-14 | 4.33E-10 |
| WBGene00000097 | aip-1 | 1.137545488 | 2.174440686 | 1.49592356 | 0.191 | 0.346 | 3.22E-07 | 0.002051568 |
| WBGene00007365 | C06B3.6 | 2.040174535 | 3.07804915 | 1.497336561 | 0.283 | 0.662 | 2.81E-21 | 1.79E-17 |
| WBGene00006372 | syx-2 | 0.267171804 | 1.317226991 | 1.51490941 | 0.031 | 0.188 | 3.28E-15 | 2.09E-11 |
| WBGene00022252 | Y73B6BL.31 | 0.710766738 | 1.769725669 | 1.527754797 | 0.087 | 0.301 | 1.15E-14 | 7.35E-11 |
| WBGene00011045 | gpx-2 | 0.694357533 | 1.754753432 | 1.529827905 | 0.082 | 0.286 | 1.04E-13 | 6.63E-10 |
| WBGene00010019 | F54B8.4 | 0.539669001 | 1.612181834 | 1.547308945 | 0.058 | 0.203 | 7.99E-10 | 5.10E-06 |
| WBGene00021977 | Y58A7A.3 | 0.200992271 | 1.28130519 | 1.558562091 | 0.008 | 0.165 | 3.64E-25 | 2.32E-21 |
| WBGene00008726 | F13A7.11 | 0.033486483 | 1.122312277 | 1.570843574 | 0.002 | 0.135 | 1.09E-26 | 6.97E-23 |
| WBGene00007346 | frpr-2 | 0.299812273 | 1.396701326 | 1.582476397 | 0.034 | 0.233 | 6.97E-21 | 4.45E-17 |
| WBGene00003963 | pdi-2 | 0.381772189 | 1.480036263 | 1.584460133 | 0.035 | 0.218 | 6.02E-18 | 3.84E-14 |
| WBGene00010700 | nipi-3 | 0.998736293 | 2.098322134 | 1.586367039 | 0.154 | 0.383 | 3.25E-13 | 2.07E-09 |
| WBGene00002086 | ins-3 | 3.784000671 | 4.885436265 | 1.58903567 | 0.729 | 0.97 | 7.37E-35 | 4.70E-31 |
| WBGene00008075 | C43F9.7 | 1.613368726 | 2.730056298 | 1.611039622 | 0.262 | 0.571 | 5.42E-19 | 3.46E-15 |
| WBGene00015167 | B0403.3 | 0.006610956 | 1.125009562 | 1.613508122 | 0.001 | 0.135 | 1.76E-28 | 1.13E-24 |
| WBGene00008777 | F13H10.6 | 2.4828366 | 3.609863038 | 1.625955453 | 0.493 | 0.872 | 1.82E-34 | 1.16E-30 |
| WBGene00010504 | K02D3.1 | 2.049606111 | 3.188337665 | 1.642842366 | 0.362 | 0.609 | 1.78E-14 | 1.14E-10 |
| WBGene00019770 | M04F3.4 | 0.390343967 | 1.543346104 | 1.663430465 | 0.036 | 0.256 | 2.39E-23 | 1.52E-19 |
| WBGene00016507 | abhd-5.2 | 1.705774745 | 2.882280779 | 1.697339421 | 0.323 | 0.722 | 1.62E-28 | 1.04E-24 |
| WBGene00019975 | flp-34 | 0.76343943 | 1.944247822 | 1.703546411 | 0.087 | 0.308 | 2.14E-15 | 1.36E-11 |
| WBGene00022648 | ZK54.3 | 0.648967776 | 1.829849009 | 1.703651498 | 0.066 | 0.233 | 2.06E-11 | 1.31E-07 |
| WBGene00016739 | pitr-1 | 0.589320376 | 1.791717617 | 1.734692537 | 0.06 | 0.308 | 3.72E-22 | 2.37E-18 |
| WBGene00020363 | fbxa-3 | 0.057533459 | 1.2740521 | 1.75506541 | 0.006 | 0.188 | 3.25E-32 | 2.07E-28 |
| WBGene00008865 | F15G9.1 | 0.079995415 | 1.298020824 | 1.757239216 | 0.007 | 0.18 | 2.17E-29 | 1.38E-25 |

| | | | | | | | | |
|---|---|---|---|---|---|---|---|---|
| WBGene00013266 | Y57A10A.26 | 1.278924661 | 2.512867756 | 1.780203585 | 0.225 | 0.519 | 6.43E-19 | 4.10E-15 |
| WBGene00004373 | rme-1 | 0.737428222 | 1.974435293 | 1.784623967 | 0.104 | 0.421 | 1.59E-25 | 1.01E-21 |
| WBGene00044443 | ZC21.10 | 0.855014588 | 2.09862335 | 1.794148193 | 0.092 | 0.316 | 1.23E-14 | 7.85E-11 |
| WBGene00015168 | pdi-6 | 0.569141685 | 1.831393469 | 1.821044389 | 0.062 | 0.301 | 1.38E-20 | 8.78E-17 |
| WBGene00017080 | lec-12 | 0.33095532 | 1.631716105 | 1.876601134 | 0.038 | 0.241 | 1.36E-20 | 8.67E-17 |
| WBGene00010742 | K10D6.2 | 0.677873072 | 2.005479672 | 1.915331458 | 0.092 | 0.331 | 2.97E-17 | 1.89E-13 |
| WBGene00009803 | F47B8.2 | 1.916800827 | 3.245595126 | 1.917044945 | 0.319 | 0.707 | 2.75E-28 | 1.76E-24 |
| WBGene00022667 | ZK154.5 | 0.537764676 | 1.8782307 | 1.933883685 | 0.059 | 0.286 | 2.14E-19 | 1.36E-15 |
| WBGene00010736 | istr-1 | 0.248152992 | 1.608674775 | 1.962818029 | 0.033 | 0.188 | 5.04E-15 | 3.22E-11 |
| WBGene00013845 | fbxa-37 | 0.075102301 | 1.441242668 | 1.970923932 | 0.008 | 0.188 | 6.56E-30 | 4.19E-26 |
| WBGene00012128 | nra-1 | 0.96221829 | 2.329181675 | 1.972111297 | 0.055 | 0.346 | 2.17E-29 | 1.39E-25 |
| WBGene00015162 | algn-11 | 0.395834247 | 1.769936359 | 1.982410303 | 0.052 | 0.278 | 1.55E-20 | 9.90E-17 |
| WBGene00005655 | srr-4 | 1.442379547 | 2.874341305 | 2.065884127 | 0.251 | 0.714 | 1.04E-37 | 6.67E-34 |
| WBGene00018869 | rfip-1 | 2.109674299 | 3.545908984 | 2.072048658 | 0.411 | 0.85 | 5.20E-42 | 3.32E-38 |
| WBGene00010833 | M03B6.1 | 2.024057547 | 3.474037101 | 2.091878312 | 0.386 | 0.774 | 2.02E-31 | 1.29E-27 |
| WBGene00009901 | F49E12.8 | 0.527268711 | 1.979509479 | 2.095140554 | 0.064 | 0.361 | 1.44E-28 | 9.20E-25 |
| WBGene00006959 | xbp-1 | 3.144286491 | 4.601969577 | 2.102992159 | 0.674 | 0.985 | 7.69E-56 | 4.90E-52 |
| WBGene00001787 | gst-39 | 0.942461636 | 2.437976315 | 2.157571612 | 0.131 | 0.398 | 3.60E-18 | 2.30E-14 |
| WBGene00017597 | F19C7.8 | 1.587081422 | 3.089603002 | 2.167680432 | 0.28 | 0.662 | 2.49E-28 | 1.59E-24 |
| WBGene00002008 | hsp-4 | 0.339104821 | 1.844989408 | 2.172532226 | 0.025 | 0.218 | 1.50E-22 | 9.57E-19 |
| WBGene00235164 | K09F6.13 | 0.028276943 | 1.635845212 | 2.319230771 | 0.003 | 0.113 | 1.43E-20 | 9.16E-17 |
| WBGene00019967 | cyp-33C8 | 1.007964736 | 2.64211242 | 2.35757676 | 0.118 | 0.564 | 7.48E-42 | 4.77E-38 |
| WBGene00022668 | ZK154.6 | 1.049423842 | 2.735966154 | 2.43316623 | 0.128 | 0.549 | 3.32E-38 | 2.12E-34 |
| WBGene00021487 | comt-3 | 0.537773648 | 2.244732465 | 2.462621021 | 0.048 | 0.316 | 1.39E-27 | 8.87E-24 |
| WBGene00019327 | zip-2 | 1.576851026 | 3.312585976 | 2.504136204 | 0.28 | 0.805 | 9.86E-50 | 6.29E-46 |
| WBGene00016997 | cebp-1 | 0.689417577 | 2.44193491 | 2.528348066 | 0.084 | 0.586 | 3.52E-58 | 2.25E-54 |
| WBGene00015164 | ykt-6 | 0.952224821 | 2.723832335 | 2.555889375 | 0.131 | 0.609 | 3.09E-47 | 1.97E-43 |
| WBGene00005097 | srd-19 | 0.33912142 | 2.111761027 | 2.55737837 | 0.042 | 0.361 | 1.41E-38 | 9.01E-35 |
| WBGene00001755 | gst-7 | 1.103671819 | 2.92662077 | 2.629959411 | 0.13 | 0.398 | 3.99E-18 | 2.55E-14 |
| WBGene00010291 | F58H10.1 | 0.592564761 | 2.563203795 | 2.843031161 | 0.06 | 0.406 | 1.11E-37 | 7.10E-34 |
| WBGene00008922 | F17C11.11 | 1.701487946 | 3.708219074 | 2.895101047 | 0.241 | 0.759 | 5.22E-49 | 3.33E-45 |
| WBGene00003991 | pfn-3 | 0.247770407 | 2.299234011 | 2.959636368 | 0.026 | 0.398 | 1.55E-56 | 9.87E-53 |
| WBGene00018702 | F52E4.5 | 0.308095504 | 2.478351437 | 3.131017472 | 0.015 | 0.466 | 8.14E-82 | 5.20E-78 |
| WBGene00194689 | Y38H6C.24 | 0.730455011 | 3.124818063 | 3.454335702 | 0.06 | 0.474 | 3.59E-50 | 2.29E-46 |
| WBGene00016788 | C49G7.10 | 0.172514796 | 2.58383967 | 3.478806438 | 0.005 | 0.165 | 7.91E-29 | 5.05E-25 |
| WBGene00044644 | B0205.13 | 0.988060163 | 3.432735091 | 3.526920396 | 0.029 | 0.368 | 4.75E-47 | 3.03E-43 |
| WBGene00001753 | gst-5 | 1.022286667 | 3.638413597 | 3.774273348 | 0.086 | 0.534 | 1.12E-48 | 7.15E-45 |

This comparison is between *fkh-7(R612C)* ADF cells in Cluster 2 and *fkh-7(wild-type)* ADF cells in Cluster 1.

| id | gene_name | fkh-7_2 | wt_1 | avg_log2FC | fkh7.pc | wt.pct | p_val | p_val_adj |
|---|---|---|---|---|---|---|---|---|
| WBGene00014248 | ZK1307.7 | 0.304630157 | 1.682463107 | -1.987792763 | 0.038 | 0.342 | 1.15E-11 | 7.34E-08 |
| WBGene00012281 | W05E10.2 | 2.286041456 | 3.614468773 | -1.916515502 | 0.477 | 0.863 | 1.18E-29 | 7.51E-26 |
| WBGene00011059 | R06C1.4 | 1.175045185 | 2.478171608 | -1.880014029 | 0.185 | 0.627 | 5.50E-17 | 3.51E-13 |
| WBGene00011372 | npr-26 | 0.52024548 | 1.711582785 | -1.718736422 | 0.054 | 0.34 | 4.04E-10 | 2.58E-06 |
| WBGene00010146 | glb-19 | 0.920627981 | 2.034884702 | -1.607532645 | 0.1 | 0.445 | 5.37E-12 | 3.43E-08 |
| WBGene00002017 | hsp-16.11 | 3.129280613 | 4.202451919 | -1.548258921 | 0.623 | 0.876 | 6.24E-22 | 3.98E-18 |
| WBGene00009503 | F37A8.5 | 0.772922423 | 1.842692074 | -1.543351371 | 0.092 | 0.364 | 1.33E-08 | 8.51E-05 |
| WBGene00005040 | sra-14 | 0.117312701 | 1.184533756 | -1.539674523 | 0.008 | 0.199 | 1.51E-07 | 0.000961232 |
| WBGene00004265 | qui-1 | 0.320174641 | 1.378171136 | -1.526366297 | 0.031 | 0.211 | 2.02E-06 | 0.012869642 |
| WBGene00008570 | kcnl-2 | 0.983043891 | 2.027604954 | -1.506983066 | 0.115 | 0.422 | 1.03E-09 | 6.57E-06 |
| WBGene00012216 | W02D9.10 | 1.986840538 | 3.023880515 | -1.496132432 | 0.362 | 0.723 | 3.31E-16 | 2.12E-12 |
| WBGene00017068 | D2092.8 | 0.133871544 | 1.162121305 | -1.483450831 | 0.008 | 0.085 | 0.002050462 | 1 |
| WBGene00002245 | lag-1 | 0.879133966 | 1.899581145 | -1.472194085 | 0.1 | 0.455 | 8.78E-12 | 5.60E-08 |
| WBGene00044484 | hot-9 | 1.679854531 | 2.6671136 | -1.424313763 | 0.308 | 0.663 | 1.12E-12 | 7.16E-09 |
| WBGene00005739 | srv-28 | 1.14653211 | 2.133364924 | -1.423698806 | 0.138 | 0.526 | 1.08E-12 | 6.89E-09 |
| WBGene00021740 | Y50D4B.1 | 1.680860959 | 2.65978042 | -1.412282251 | 0.315 | 0.685 | 3.45E-14 | 2.20E-10 |
| WBGene00005903 | srx-12 | 1.44121145 | 2.417320098 | -1.408227107 | 0.177 | 0.615 | 1.85E-15 | 1.18E-11 |
| WBGene00017439 | F13H8.6 | 0.157163658 | 1.133234713 | -1.408172871 | 0.015 | 0.186 | 1.78E-06 | 0.011349107 |
| WBGene00015514 | nlp-77 | 0.150115171 | 1.115524193 | -1.392790809 | 0.015 | 0.11 | 0.000831634 | 1 |
| WBGene00044029 | C05D12.7 | 1.146886447 | 2.112090814 | -1.392495553 | 0.162 | 0.487 | 3.27E-10 | 2.09E-06 |
| WBGene00015919 | C17G10.7 | 1.717779009 | 2.656928397 | -1.354906164 | 0.269 | 0.648 | 8.45E-13 | 5.39E-09 |
| WBGene00003808 | npr-2 | 0 | 0.921248326 | -1.329080391 | 0 | 0.122 | 2.94E-05 | 0.18744647 |
| WBGene00002018 | hsp-16.41 | 3.10740658 | 4.017376303 | -1.312808806 | 0.446 | 0.623 | 1.81E-06 | 0.011538607 |
| WBGene00015735 | pdfr-1 | 0.694195457 | 1.572601774 | -1.267272438 | 0.069 | 0.273 | 3.55E-06 | 0.02262486 |
| WBGene00004481 | rps-12 | 1.738948348 | 2.613998055 | -1.262429872 | 0.315 | 0.658 | 9.54E-11 | 6.08E-07 |
| WBGene00002016 | hsp-16.2 | 3.314389352 | 4.184724763 | -1.255628582 | 0.408 | 0.555 | 0.00011966 | 0.763553439 |
| WBGene00010426 | H37A05.2 | 0.607212986 | 1.473742295 | -1.250137537 | 0.062 | 0.29 | 2.01E-07 | 0.00128279 |
| WBGene00005210 | srg-53 | 0.615893567 | 1.464729997 | -1.224612108 | 0.054 | 0.282 | 1.64E-07 | 0.001045009 |
| WBGene00004451 | rpl-37 | 1.723811997 | 2.566634969 | -1.215936523 | 0.323 | 0.642 | 6.03E-10 | 3.85E-06 |
| WBGene00020058 | R13D11.4 | 0.272432417 | 1.113856564 | -1.213918445 | 0.023 | 0.184 | 5.93E-06 | 0.037847689 |
| WBGene00044803 | C02G6.3 | 0.616068712 | 1.456050533 | -1.211837606 | 0.069 | 0.259 | 6.07E-06 | 0.038717904 |
| WBGene00006801 | unc-68 | 0.269356477 | 1.108010753 | -1.209922365 | 0.023 | 0.186 | 6.37E-06 | 0.040643204 |
| WBGene00013490 | Y70C5A.2 | 0.161621089 | 0.999846411 | -1.209303515 | 0.015 | 0.186 | 1.87E-06 | 0.011957722 |
| WBGene00016984 | npr-8 | 0.266445612 | 1.100738653 | -1.203630433 | 0.023 | 0.159 | 5.86E-05 | 0.37391743 |
| WBGene00017411 | F13A2.2 | 0.354984338 | 1.172730044 | -1.179757674 | 0.038 | 0.209 | 7.30E-06 | 0.046577243 |
| WBGene00015478 | mapk-15 | 0.743738281 | 1.558297512 | -1.175160563 | 0.092 | 0.277 | 3.30E-05 | 0.210744426 |
| WBGene00017858 | F27C1.11 | 0.97006473 | 1.783244571 | -1.173170524 | 0.123 | 0.35 | 2.33E-06 | 0.014870687 |
| WBGene00007862 | kvs-2 | 0.868657145 | 1.679334346 | -1.169559977 | 0.108 | 0.327 | 1.83E-06 | 0.011674877 |
| WBGene00008382 | D1081.4 | 0.433908239 | 1.242656347 | -1.166776884 | 0.046 | 0.172 | 0.000443846 | 1 |
| WBGene00022887 | ZK1290.5 | 0.446632097 | 1.252091901 | -1.162032865 | 0.046 | 0.219 | 1.27E-05 | 0.0811345 |
| WBGene00044353 | ZK418.10 | 0.34220542 | 1.145897075 | -1.159481966 | 0.038 | 0.215 | 5.19E-06 | 0.033115211 |
| WBGene00003511 | mxl-3 | 1.511171081 | 2.314355813 | -1.15875063 | 0.231 | 0.551 | 1.18E-09 | 7.51E-06 |
| WBGene00009691 | F44E5.4 | 3.698475331 | 4.499879914 | -1.156182418 | 0.854 | 0.909 | 1.17E-15 | 7.47E-12 |
| WBGene00002265 | lec-2 | 0.154294174 | 0.947543023 | -1.144416181 | 0.015 | 0.149 | 3.96E-05 | 0.252696333 |
| WBGene00009692 | F44E5.5 | 3.676132461 | 4.464430124 | -1.137273129 | 0.862 | 0.919 | 1.09E-19 | 6.94E-16 |
| WBGene00014215 | obr-3 | 0.221734984 | 1.009881623 | -1.137055248 | 0.023 | 0.122 | 0.00106985 | 1 |
| WBGene00018187 | twf-2 | 0.162773005 | 0.949890559 | -1.135570592 | 0.015 | 0.153 | 2.96E-05 | 0.188760656 |
| WBGene00003728 | nhr-138 | 1.459754616 | 2.240300221 | -1.126089274 | 0.231 | 0.598 | 9.14E-11 | 5.83E-07 |
| WBGene00022410 | Y97E10C.1 | 0.945556126 | 1.724346139 | -1.12355649 | 0.108 | 0.338 | 1.57E-06 | 0.010014204 |
| WBGene00003639 | nhr-49 | 0.16240275 | 0.938877293 | -1.120215973 | 0.015 | 0.166 | 9.99E-06 | 0.063746559 |
| WBGene00004744 | scp-1 | 0.888155193 | 1.662393239 | -1.11698939 | 0.077 | 0.168 | 0.008168191 | 1 |
| WBGene00015650 | pkg-2 | 0.779982942 | 1.548230704 | -1.108347237 | 0.092 | 0.304 | 4.53E-06 | 0.0289334 |
| WBGene00000518 | ckk-1 | 0.196263569 | 0.963096234 | -1.106305683 | 0.015 | 0.174 | 5.25E-06 | 0.033476102 |
| WBGene00008249 | srsx-28 | 1.361437216 | 2.125570129 | -1.102410764 | 0.192 | 0.497 | 4.73E-08 | 0.000301667 |
| WBGene00009742 | F45H10.5 | 0.194721563 | 0.957861773 | -1.100978597 | 0.015 | 0.143 | 6.78E-05 | 0.43272847 |
| WBGene00008439 | mfb-1 | 1.734524165 | 2.494802748 | -1.096850141 | 0.269 | 0.621 | 2.88E-10 | 1.84E-06 |
| WBGene00004435 | rpl-23 | 1.612419008 | 2.372666598 | -1.096805427 | 0.254 | 0.592 | 2.47E-09 | 1.58E-05 |
| WBGene00019023 | F58A6.2 | 1.206081485 | 1.966140218 | -1.096532965 | 0.146 | 0.468 | 1.14E-09 | 7.28E-06 |
| WBGene00015368 | srt-65 | 1.425736881 | 2.185012564 | -1.095403262 | 0.177 | 0.449 | 2.53E-07 | 0.001615997 |
| WBGene00015293 | C01C4.3 | 1.181641793 | 1.937608959 | -1.090630081 | 0.131 | 0.439 | 2.54E-09 | 1.62E-05 |
| WBGene00009619 | npr-6 | 0.501716761 | 1.252844579 | -1.083648378 | 0.054 | 0.215 | 4.82E-05 | 0.307669846 |

| Legend | |
|---|---|
| id | WormBase GeneID |
| gene_name | Gene symbol (WS273) |
| fkh-7_2 | Log of averaged normalized expression across all *fkh-7(R612C)* ADF neurons in cluster 2 |
| wt_1 | Log of averaged normalized expression across all *fkh-7(wild-type)* ADF neurons in cluster 1 |
| avg_log2FC | log fold-change (*fkh-7(R612C)* - *fkh-7(wild-type)*) |
| fkh7.pct | Percent of Cluster 2 *fkh-7(R612C)* ADF neurons where gene is detected ($\geq$ 1 UMI) |
| wt.pct | Percent of Cluster 1 *fkh-7(wild-type)* ADF neurons where gene is detected ($\geq$ 1 UMI) |
| p_val | p-value |
| p_val_adj | Adjusted p-value, based on bonferroni correction |

| | | | | | | | | |
|---|---|---|---|---|---|---|---|---|
| WBGene00004622 | rrn-3.1 | 4.194495482 | 4.941717395 | -1.078013348 | 0.977 | 0.983 | 0.001893562 | 1 |
| WBGene00010191 | dmsr-1 | 0.478241448 | 1.22517801 | -1.077601674 | 0.038 | 0.205 | 1.06E-05 | 0.067899061 |
| WBGene00004344 | rgs-1 | 1.302449907 | 2.047808968 | -1.075325822 | 0.185 | 0.495 | 2.64E-08 | 0.000168188 |
| WBGene00015354 | snet-1 | 0.618654453 | 1.361723848 | -1.072022531 | 0.054 | 0.052 | 0.907598112 | 1 |
| WBGene00006780 | unc-44 | 1.138706611 | 1.88094344 | -1.070821393 | 0.162 | 0.356 | 0.000162283 | 1 |
| WBGene00015263 | B0563.6 | 0 | 0.737931721 | -1.064610435 | 0 | 0.099 | 0.000196478 | 1 |
| WBGene00001980 | hmr-1 | 0.280488871 | 1.016859887 | -1.062358813 | 0.031 | 0.157 | 0.000194894 | 1 |
| WBGene00003690 | nhr-100 | 0.991646403 | 1.719979953 | -1.050763201 | 0.092 | 0.369 | 1.66E-08 | 0.000105958 |
| WBGene00001303 | sec-61.G | 0.89350933 | 1.612244375 | -1.036915484 | 0.131 | 0.325 | 5.77E-05 | 0.368389293 |
| WBGene00008368 | D1053.3 | 0.741479683 | 1.460068181 | -1.036704062 | 0.077 | 0.3 | 1.23E-06 | 0.007859143 |
| WBGene00009579 | arrd-6 | 1.187946751 | 1.90647141 | -1.036611962 | 0.154 | 0.447 | 6.96E-08 | 0.000443905 |
| WBGene00021799 | Y53G8AL.1 | 0.304209268 | 1.021552235 | -1.034907142 | 0.031 | 0.099 | 0.014333656 | 1 |
| WBGene00013639 | Y105C5A.15 | 0.608756952 | 1.322182973 | -1.029256182 | 0.069 | 0.191 | 0.001709182 | 1 |
| WBGene00000883 | cyn-7 | 1.938817123 | 2.645345157 | -1.019304491 | 0.323 | 0.694 | 9.37E-11 | 5.98E-07 |
| WBGene00020790 | T25B6.4 | 0.868599577 | 1.573840949 | -1.017448231 | 0.1 | 0.302 | 1.14E-05 | 0.072929766 |
| WBGene00195073 | F02E11.7 | 1.302254795 | 2.006356082 | -1.015803435 | 0.185 | 0.484 | 6.09E-08 | 0.000388798 |
| WBGene00007696 | tram-1 | 0.077624492 | 0.78079178 | -1.014455959 | 0.008 | 0.091 | 0.001275938 | 1 |
| WBGene00022405 | srsx-6 | 0.775996529 | 1.473989625 | -1.006991177 | 0.108 | 0.279 | 0.000220167 | 1 |
| WBGene00015350 | fbxl-1 | 1.064306239 | 1.762212168 | -1.006865423 | 0.154 | 0.369 | 2.12E-05 | 0.135042052 |
| WBGene00014308 | F07H5.4 | 1.651741017 | 2.349633344 | -1.006845799 | 0.3 | 0.565 | 2.57E-06 | 0.016427112 |
| WBGene00001747 | gsp-1 | 0.645572017 | 1.342786587 | -1.005868003 | 0.069 | 0.236 | 6.07E-05 | 0.387403382 |
| WBGene00000056 | acr-17 | 0.072913271 | 0.766120882 | -1.000087183 | 0.008 | 0.101 | 0.000597699 | 1 |
| WBGene00000239 | bas-1 | 0.626473987 | 1.31905131 | -0.999177869 | 0.062 | 0.263 | 2.79E-06 | 0.017797548 |
| WBGene00008248 | srsx-27 | 2.110784265 | 2.802389527 | -0.997775481 | 0.377 | 0.733 | 6.51E-11 | 4.15E-07 |
| WBGene00015750 | C14A11.5 | 1.234637041 | 1.926120425 | -0.99759965 | 0.154 | 0.422 | 4.13E-07 | 0.002634563 |
| WBGene00007588 | C14C10.5 | 0 | 0.691192223 | -0.997179593 | 0 | 0.052 | 0.007857927 | 1 |
| WBGene00017715 | F22F1.3 | 0.23561401 | 0.92645994 | -0.996679997 | 0.015 | 0.141 | 7.96E-05 | 0.507720792 |
| WBGene00007892 | C33B4.4 | 2.235667534 | 2.925684855 | -0.995484568 | 0.415 | 0.747 | 4.63E-11 | 2.95E-07 |
| WBGene00003953 | pbs-7 | 0.283443821 | 0.972276941 | -0.993776126 | 0.031 | 0.157 | 0.000198377 | 1 |
| WBGene00022732 | ZK418.2 | 0.565703386 | 1.248852429 | -0.985575736 | 0.038 | 0.205 | 1.26E-05 | 0.080244718 |
| WBGene00007997 | sre-13 | 1.645466022 | 2.328095316 | -0.984825897 | 0.277 | 0.547 | 7.67E-07 | 0.0048943 |
| WBGene00004567 | rrn-2.1 | 1.665735847 | 2.347285808 | -0.983268748 | 0.269 | 0.489 | 3.11E-05 | 0.19827513 |
| WBGene00174531 | 21ur-10538 | 0.198792133 | 0.878873957 | -0.981150675 | 0.023 | 0.128 | 0.000616441 | 1 |
| WBGene00007405 | C07A9.9 | 0.067297503 | 0.744954373 | -0.977652205 | 0.008 | 0.104 | 0.000439727 | 1 |
| WBGene00012528 | pap-1 | 1.091883491 | 1.761324722 | -0.965799544 | 0.154 | 0.344 | 0.000108916 | 0.694991429 |
| WBGene00044387 | C27A2.8 | 1.332624471 | 2.000482459 | -0.963515408 | 0.231 | 0.441 | 2.78E-05 | 0.177672863 |
| WBGene00004484 | rps-15 | 1.916694054 | 2.582790457 | -0.960973978 | 0.362 | 0.675 | 4.77E-09 | 3.05E-05 |
| WBGene00000209 | asg-1 | 1.408071345 | 2.074102439 | -0.960879757 | 0.2 | 0.513 | 4.11E-08 | 0.00026204 |
| WBGene00009999 | srsx-35 | 0.149218772 | 0.814274288 | -0.959472296 | 0.008 | 0.124 | 9.73E-05 | 0.621045737 |
| WBGene00000232 | avr-14 | 0 | 0.659605825 | -0.951610052 | 0 | 0.062 | 0.003669113 | 1 |
| WBGene00009180 | nurf-1 | 1.322806274 | 1.979408037 | -0.947276107 | 0.208 | 0.377 | 0.001012163 | 1 |
| WBGene00000210 | asg-2 | 0.329156993 | 0.984821511 | -0.945923948 | 0.038 | 0.139 | 0.00185476 | 1 |
| WBGene00009995 | F53F4.13 | 0.297547103 | 0.950688092 | -0.942283266 | 0.031 | 0.083 | 0.042441241 | 1 |
| WBGene00010677 | gtbp-1 | 0.304139633 | 0.956983938 | -0.941855242 | 0.038 | 0.159 | 0.000372652 | 1 |
| WBGene00006756 | unc-17 | 1.10241567 | 1.754236573 | -0.940378783 | 0.154 | 0.368 | 4.07E-05 | 0.259682517 |
| WBGene00017613 | F20A1.1 | 0.083346841 | 0.734213506 | -0.93900211 | 0.008 | 0.066 | 0.009310691 | 1 |
| WBGene00006702 | ubc-3 | 1.351449856 | 1.998235693 | -0.933114719 | 0.192 | 0.431 | 5.77E-06 | 0.036843041 |
| WBGene00020586 | frpr-18 | 1.086259982 | 1.730958183 | -0.930102897 | 0.146 | 0.348 | 6.75E-05 | 0.430474139 |
| WBGene00009920 | abts-1 | 0.815022315 | 1.458982218 | -0.929037759 | 0.115 | 0.259 | 0.001163022 | 1 |
| WBGene00018466 | F45E1.5 | 0.622854668 | 1.266357599 | -0.928378488 | 0.069 | 0.234 | 6.94E-05 | 0.443038999 |
| WBGene00004431 | rpl-19 | 2.000521981 | 2.642423897 | -0.92606871 | 0.385 | 0.704 | 7.74E-09 | 4.94E-05 |
| WBGene00003163 | mdl-1 | 0.716446309 | 1.356171775 | -0.922928757 | 0.085 | 0.244 | 0.000140066 | 0.893758271 |
| WBGene00006352 | sur-6 | 0.487004522 | 1.125167611 | -0.920674725 | 0.054 | 0.176 | 0.000846323 | 1 |
| WBGene00006789 | unc-54 | 4.70677009 | 5.341022423 | -0.915032696 | 0.985 | 0.983 | 6.74E-30 | 4.30E-26 |
| WBGene00018846 | eef-1B.1 | 2.084045715 | 2.718123866 | -0.914781404 | 0.377 | 0.708 | 2.43E-09 | 1.55E-05 |
| WBGene00001475 | fmi-1 | 0.378316472 | 1.011375317 | -0.913310856 | 0.038 | 0.161 | 0.000370501 | 1 |
| WBGene00013898 | ZC443.3 | 0.990773056 | 1.62231636 | -0.911124393 | 0.123 | 0.354 | 2.95E-06 | 0.018802705 |
| WBGene00005346 | srh-129 | 0.802307118 | 1.43361511 | -0.910784908 | 0.1 | 0.263 | 0.000183032 | 1 |
| WBGene00010669 | bath-37 | 0.222836807 | 0.852764621 | -0.908793733 | 0.023 | 0.147 | 0.000159292 | 1 |
| WBGene00011977 | T24B8.3 | 0.268336668 | 0.89759408 | -0.907826549 | 0.031 | 0.151 | 0.000271222 | 1 |
| WBGene00017926 | cox-6C | 0.342748694 | 0.971015286 | -0.906397096 | 0.031 | 0.153 | 0.000286987 | 1 |
| WBGene00004492 | rps-23 | 2.404703504 | 3.032690668 | -0.905993966 | 0.569 | 0.783 | 8.14E-10 | 5.19E-06 |
| WBGene00015373 | C03B1.2 | 0.595212257 | 1.221984965 | -0.904241878 | 0.069 | 0.213 | 0.00035408 | 1 |
| WBGene00019603 | K09H11.6 | 0.63540908 | 1.260736383 | -0.902156599 | 0.077 | 0.195 | 0.002045632 | 1 |

| | | | | | | | | |
|---|---|---|---|---|---|---|---|---|
| WBGene00009119 | ndk-1 | 1.979351986 | 2.599420214 | -0.894569358 | 0.377 | 0.631 | 1.17E-06 | 0.007492459 |
| WBGene00022479 | fbxa-36 | 0.400485693 | 1.019418816 | -0.892931746 | 0.046 | 0.166 | 0.000660334 | 1 |
| WBGene00021078 | unc-132 | 0.077157095 | 0.694132183 | -0.8901069 | 0.008 | 0.072 | 0.006014972 | 1 |
| WBGene00003839 | ocr-2 | 1.10020805 | 1.716872282 | -0.889658429 | 0.146 | 0.366 | 1.72E-05 | 0.109589568 |
| WBGene00015307 | C01G5.5 | 2.93603585 | 3.546596202 | -0.880852392 | 0.685 | 0.87 | 1.36E-11 | 8.67E-08 |
| WBGene00005834 | srw-87 | 0.244484936 | 0.854427409 | -0.879960981 | 0.023 | 0.139 | 0.000280506 | 1 |
| WBGene00194905 | Y53F4B.51 | 0.077157095 | 0.686679609 | -0.879355109 | 0.008 | 0.118 | 0.000154352 | 0.984918413 |
| WBGene00012235 | W04A4.6 | 1.060943669 | 1.666586825 | -0.873758377 | 0.162 | 0.34 | 0.000378283 | 1 |
| WBGene00008925 | F17H10.1 | 0.547084678 | 1.150569913 | -0.870645156 | 0.069 | 0.217 | 0.000285002 | 1 |
| WBGene00012967 | kvs-4 | 0.627985711 | 1.22943594 | -0.867709263 | 0.062 | 0.205 | 0.000246041 | 1 |
| WBGene00044079 | tag-241 | 0.278931869 | 0.874566487 | -0.859319111 | 0.023 | 0.093 | 0.009243544 | 1 |
| WBGene00006529 | tba-2 | 0.680838594 | 1.276341015 | -0.85912839 | 0.085 | 0.201 | 0.003134071 | 1 |
| WBGene00020114 | pde-4 | 0.911346144 | 1.505344628 | -0.856958668 | 0.138 | 0.294 | 0.000697232 | 1 |
| WBGene00004286 | rab-39 | 0.225150298 | 0.818203219 | -0.855594509 | 0.015 | 0.093 | 0.003698661 | 1 |
| WBGene00002045 | icd-1 | 1.151845472 | 1.743804099 | -0.854015776 | 0.169 | 0.377 | 0.000168902 | 1 |
| WBGene00000282 | cah-4 | 0.767000989 | 1.358869239 | -0.853885389 | 0.115 | 0.271 | 0.000522036 | 1 |
| WBGene00001091 | drh-2 | 0.126175193 | 0.716887428 | -0.852217612 | 0.008 | 0.058 | 0.01691714 | 1 |
| WBGene00021820 | nipa-1 | 1.099660765 | 1.68773393 | -0.848410238 | 0.154 | 0.35 | 7.40E-05 | 0.47235154 |
| WBGene00013136 | Y53C10A.5 | 0.441753907 | 1.028574595 | -0.846603296 | 0.046 | 0.159 | 0.001070514 | 1 |
| WBGene00001588 | ggr-3 | 0.341678855 | 0.925648866 | -0.84249064 | 0.038 | 0.132 | 0.003139297 | 1 |
| WBGene00010600 | nhr-196 | 0.084260344 | 0.667162037 | -0.840949382 | 0.008 | 0.11 | 0.000284919 | 1 |
| WBGene00000287 | cal-3 | 0.13554397 | 0.71688118 | -0.83869231 | 0.015 | 0.079 | 0.008843146 | 1 |
| WBGene00001682 | gpc-2 | 1.613209369 | 2.191554225 | -0.834375256 | 0.277 | 0.565 | 1.31E-06 | 0.008329477 |
| WBGene00011974 | xbx-5 | 0.799334752 | 1.375872235 | -0.831767769 | 0.1 | 0.24 | 0.001322841 | 1 |
| WBGene00010647 | K08C7.4 | 0.506096775 | 1.082494746 | -0.831566494 | 0.054 | 0.151 | 0.005003804 | 1 |
| WBGene00006527 | tax-6 | 1.139246699 | 1.714565935 | -0.830010209 | 0.162 | 0.309 | 0.001570458 | 1 |
| WBGene00022078 | epg-9 | 0.80731543 | 1.382617422 | -0.829985331 | 0.1 | 0.273 | 0.000130693 | 0.833954677 |
| WBGene00019537 | K08D12.3 | 0.684167775 | 1.259430565 | -0.829928773 | 0.092 | 0.25 | 0.000216258 | 1 |
| WBGene00015076 | B0238.12 | 0.230549165 | 0.803321593 | -0.826335942 | 0.023 | 0.112 | 0.002270588 | 1 |
| WBGene00021697 | gcn-1 | 0.175899304 | 0.746453219 | -0.823135304 | 0.015 | 0.07 | 0.019929429 | 1 |
| WBGene00004677 | rrn-3.56 | 0.964624907 | 1.534308755 | -0.821880063 | 0.123 | 0.188 | 0.149330376 | 1 |
| WBGene00044588 | ZK1236.9 | 0.626278352 | 1.195437682 | -0.821123344 | 0.062 | 0.215 | 0.00011893 | 0.758892028 |
| WBGene00017724 | F22F7.7 | 0.217222429 | 0.785538468 | -0.81990673 | 0.023 | 0.097 | 0.007054746 | 1 |
| WBGene00022336 | Y82E9BR.3 | 1.737975219 | 2.30510914 | -0.818201296 | 0.323 | 0.578 | 2.14E-05 | 0.136358807 |
| WBGene00013159 | Y53F4B.12 | 0.512214192 | 1.078175047 | -0.816508918 | 0.069 | 0.186 | 0.002115108 | 1 |
| WBGene00008920 | eef-1G | 1.762529602 | 2.327795863 | -0.815506832 | 0.323 | 0.594 | 2.51E-06 | 0.015992394 |
| WBGene00006599 | tpa-1 | 0.080806338 | 0.644992518 | -0.813948604 | 0.008 | 0.077 | 0.003701511 | 1 |
| WBGene00003831 | nuo-1 | 0 | 0.563638983 | -0.813159165 | 0 | 0.066 | 0.002704595 | 1 |
| WBGene00003407 | mrp-1 | 0.201872322 | 0.763031575 | -0.809581671 | 0.015 | 0.101 | 0.001958458 | 1 |
| WBGene00004680 | rars-2 | 0 | 0.56024576 | -0.80826378 | 0 | 0.085 | 0.000583123 | 1 |
| WBGene00018468 | cla-1 | 1.434902972 | 1.994337163 | -0.807092933 | 0.215 | 0.433 | 0.000138068 | 0.881009947 |
| WBGene00022790 | ZK682.7 | 1.097983063 | 1.65700565 | -0.806499114 | 0.154 | 0.302 | 0.002272069 | 1 |
| WBGene00016097 | C25E10.8 | 1.228489751 | 1.784867793 | -0.802683843 | 0.177 | 0.383 | 7.76E-05 | 0.494907728 |
| WBGene00019177 | H10D18.1 | 0.776760299 | 1.331534487 | -0.800369969 | 0.1 | 0.269 | 0.000127017 | 0.810493476 |
| WBGene00002000 | hrp-2 | 0.483760944 | 1.034788069 | -0.7949641 | 0.062 | 0.141 | 0.016980881 | 1 |
| WBGene00018879 | F55D10.4 | 0.103003395 | 0.653598495 | -0.794340821 | 0.008 | 0.079 | 0.003287557 | 1 |
| WBGene00001193 | egl-26 | 0.751331026 | 1.30121567 | -0.793315848 | 0.077 | 0.292 | 3.44E-06 | 0.021954088 |
| WBGene00010507 | K02E2.6 | 0.425386135 | 0.974982616 | -0.792900118 | 0.054 | 0.166 | 0.001581273 | 1 |
| WBGene00007943 | C34F6.9 | 0.241137489 | 0.790406515 | -0.7924277 | 0.015 | 0.128 | 0.000243519 | 1 |
| WBGene00018214 | F39H12.3 | 1.165242452 | 1.713498911 | -0.790966874 | 0.177 | 0.371 | 0.000194187 | 1 |
| WBGene00018467 | sdpn-1 | 1.291966893 | 1.838606418 | -0.788634132 | 0.192 | 0.395 | 0.000298234 | 1 |
| WBGene00017317 | attf-2 | 0.524911467 | 1.071309143 | -0.788285217 | 0.062 | 0.195 | 0.000474618 | 1 |
| WBGene00007330 | C05C10.3 | 0.343049045 | 0.888916393 | -0.787520117 | 0.023 | 0.145 | 0.000200939 | 1 |
| WBGene00008516 | F02C12.2 | 0.978855587 | 1.522948646 | -0.784960359 | 0.115 | 0.319 | 2.70E-05 | 0.17199701 |
| WBGene00001340 | etr-1 | 1.692863442 | 2.236783698 | -0.784711056 | 0.285 | 0.542 | 1.06E-05 | 0.067735805 |
| WBGene00016532 | C39D10.5 | 0.582088162 | 1.12596824 | -0.784653092 | 0.062 | 0.166 | 0.003575687 | 1 |
| WBGene00000472 | cey-1 | 1.465152628 | 2.005584373 | -0.779678198 | 0.223 | 0.491 | 2.50E-06 | 0.015960229 |
| WBGene00007861 | C32A9.1 | 0 | 0.540181283 | -0.779316858 | 0 | 0.079 | 0.000925897 | 1 |
| WBGene00000164 | apm-3 | 0.0991499 | 0.639164876 | -0.779076928 | 0.008 | 0.06 | 0.014913822 | 1 |
| WBGene00015938 | anat-1 | 0.319984981 | 0.859943544 | -0.77899554 | 0.031 | 0.145 | 0.000566729 | 1 |
| WBGene00011300 | R107.5 | 1.470867497 | 2.010794752 | -0.778950373 | 0.208 | 0.395 | 0.000641487 | 1 |
| WBGene00001708 | grk-1 | 0.055737149 | 0.595230712 | -0.778324688 | 0.008 | 0.072 | 0.005805711 | 1 |
| WBGene00011273 | R53.4 | 0.832682725 | 1.371201886 | -0.776918924 | 0.108 | 0.263 | 0.000594399 | 1 |
| WBGene00004719 | sad-1 | 0 | 0.536675235 | -0.7742587 | 0 | 0.066 | 0.002704595 | 1 |

| | | | | | | | | |
|---|---|---|---|---|---|---|---|---|
| WBGene00001976 | hmg-11 | 0.470648741 | 1.006821612 | -0.773533942 | 0.038 | 0.17 | 0.00024462 | 1 |
| WBGene00011824 | T18D3.7 | 0.76590553 | 1.299369201 | -0.769625392 | 0.1 | 0.259 | 0.000362609 | 1 |
| WBGene00016706 | C46C11.3 | 0.331277311 | 0.864031608 | -0.768601982 | 0.031 | 0.103 | 0.012644688 | 1 |
| WBGene00010357 | H03A11.2 | 0.062997084 | 0.595337883 | -0.76800543 | 0.008 | 0.068 | 0.007770661 | 1 |
| WBGene00008250 | srsx-29 | 1.397855966 | 1.927102765 | -0.763541731 | 0.215 | 0.453 | 4.25E-05 | 0.27102469 |
| WBGene00006862 | pnk-4 | 0.144487899 | 0.673346711 | -0.762981985 | 0.015 | 0.075 | 0.01302785 | 1 |
| WBGene00009367 | F33H2.3 | 0.144993469 | 0.672930429 | -0.761652034 | 0.015 | 0.101 | 0.00194405 | 1 |
| WBGene00000263 | F23H11.5 | 0.848736339 | 1.376322802 | -0.761146375 | 0.108 | 0.261 | 0.000713233 | 1 |
| WBGene00018344 | npr-27 | 0.330444345 | 0.856966476 | -0.759610867 | 0.031 | 0.139 | 0.000842639 | 1 |
| WBGene00004430 | rpl-18 | 2.098876198 | 2.624694742 | -0.758595806 | 0.392 | 0.694 | 1.02E-07 | 0.000648232 |
| WBGene00016855 | C50H11.8 | 0.218684218 | 0.7435745 | -0.757256607 | 0.015 | 0.052 | 0.071373382 | 1 |
| WBGene00020112 | pfd-5 | 0.055818059 | 0.579563896 | -0.755605521 | 0.008 | 0.062 | 0.012119687 | 1 |
| WBGene00004349 | rgs-6 | 0.366116079 | 0.889818912 | -0.75554348 | 0.046 | 0.128 | 0.009474212 | 1 |
| WBGene00004507 | rpy-1 | 0.137201465 | 0.659314782 | -0.753250293 | 0.015 | 0.099 | 0.002179931 | 1 |
| WBGene00004704 | rsp-7 | 0.092581569 | 0.61468827 | -0.753240748 | 0.008 | 0.087 | 0.001769851 | 1 |
| WBGene00017571 | jmjd-3.1 | 0.375958573 | 0.897289931 | -0.752122165 | 0.038 | 0.153 | 0.000794861 | 1 |
| WBGene00022250 | Y73B6BL.29 | 0.495354892 | 1.015679259 | -0.750669385 | 0.062 | 0.172 | 0.002900943 | 1 |
| WBGene00004512 | rrn-1.1 | 0.865339257 | 1.384635476 | -0.74918608 | 0.077 | 0.219 | 0.000681069 | 1 |
| WBGene00009575 | F40F8.1 | 0.081491558 | 0.600390941 | -0.748613567 | 0.008 | 0.058 | 0.017114036 | 1 |
| WBGene00017098 | mek-5 | 0.348220864 | 0.865864281 | -0.746801591 | 0.038 | 0.104 | 0.02340569 | 1 |
| WBGene00016250 | hsp-110 | 1.666971731 | 2.183098167 | -0.74461305 | 0.277 | 0.518 | 5.46E-05 | 0.348219594 |
| WBGene00003681 | nhr-91 | 0 | 0.515527078 | -0.743748358 | 0 | 0.056 | 0.005794642 | 1 |
| WBGene00010967 | nduo-5 | 1.29392903 | 1.809444869 | -0.743732146 | 0.2 | 0.319 | 0.033349164 | 1 |
| WBGene00004267 | rab-3 | 0.821062792 | 1.334021226 | -0.740042588 | 0.115 | 0.267 | 0.000865497 | 1 |
| WBGene00015139 | B0310.3 | 0.737554736 | 1.249981936 | -0.739276181 | 0.092 | 0.234 | 0.00092233 | 1 |
| WBGene00019719 | M01H9.3 | 2.737691346 | 3.249875692 | -0.738925817 | 0.623 | 0.832 | 2.89E-08 | 0.000184216 |
| WBGene00004705 | rsp-8 | 0.140226528 | 0.651023347 | -0.736924038 | 0.008 | 0.106 | 0.00039595 | 1 |
| WBGene00008890 | ser-5 | 0.381859352 | 0.891217578 | -0.734848587 | 0.031 | 0.139 | 0.000844065 | 1 |
| WBGene00195086 | F21C10.13 | 0.390373593 | 0.899505428 | -0.734521973 | 0.038 | 0.139 | 0.002113396 | 1 |
| WBGene00012825 | Y43F8C.3 | 0.273132136 | 0.781882295 | -0.733971331 | 0.023 | 0.093 | 0.009996464 | 1 |
| WBGene00001427 | fkb-2 | 1.021670666 | 1.529298179 | -0.732351696 | 0.138 | 0.327 | 0.000242246 | 1 |
| WBGene00010006 | F53H2.3 | 0.480401837 | 0.987896493 | -0.732160024 | 0.046 | 0.132 | 0.008592653 | 1 |
| WBGene00021952 | vha-19 | 0 | 0.506468874 | -0.730680132 | 0 | 0.068 | 0.002321677 | 1 |
| WBGene00004482 | rps-13 | 2.103866829 | 2.609766427 | -0.729858841 | 0.415 | 0.689 | 9.32E-08 | 0.000594825 |
| WBGene00015074 | natc-2 | 1.306721707 | 1.81068562 | -0.727066238 | 0.2 | 0.429 | 5.39E-05 | 0.343937544 |
| WBGene00015248 | mai-2 | 0.68207235 | 1.183346329 | -0.723185483 | 0.085 | 0.221 | 0.001220536 | 1 |
| WBGene00002065 | iff-2 | 1.16653269 | 1.667288563 | -0.722438014 | 0.146 | 0.342 | 0.000153826 | 0.981565816 |
| WBGene00004419 | rpl-7A | 1.861988558 | 2.362678384 | -0.722342729 | 0.323 | 0.605 | 7.09E-06 | 0.045235258 |
| WBGene00016407 | unk-1 | 0.230725254 | 0.729847398 | -0.720081042 | 0.023 | 0.099 | 0.006231863 | 1 |
| WBGene00009100 | rgef-1 | 0.069050062 | 0.567425949 | -0.719004421 | 0.008 | 0.077 | 0.003805652 | 1 |
| WBGene00013859 | ZC247.1 | 1.001642835 | 1.49966357 | -0.718492044 | 0.138 | 0.251 | 0.016132097 | 1 |
| WBGene00016165 | wdr-60 | 0.454984106 | 0.952584159 | -0.717885129 | 0.054 | 0.162 | 0.002290221 | 1 |
| WBGene00001005 | dlc-1 | 0.938885066 | 1.43605034 | -0.717257875 | 0.131 | 0.3 | 0.000605312 | 1 |
| WBGene00010731 | K10C3.4 | 1.085751496 | 1.582833324 | -0.717137489 | 0.146 | 0.298 | 0.001614354 | 1 |
| WBGene00001854 | hil-3 | 0.21281636 | 0.709632124 | -0.716753639 | 0.015 | 0.095 | 0.003069257 | 1 |
| WBGene00012113 | T28B8.1 | 1.160893232 | 1.657519763 | -0.716480634 | 0.177 | 0.337 | 0.002068656 | 1 |
| WBGene00006779 | unc-43 | 1.425760721 | 1.922366796 | -0.716451121 | 0.185 | 0.449 | 2.57E-06 | 0.016397914 |
| WBGene00006706 | ubc-9 | 0.229178361 | 0.725653736 | -0.716262561 | 0.023 | 0.093 | 0.00964485 | 1 |
| WBGene00020275 | atp-4 | 0.25574311 | 0.75122423 | -0.714828154 | 0.031 | 0.139 | 0.000795503 | 1 |
| WBGene00001329 | epn-1 | 1.34669559 | 1.842007964 | -0.714584706 | 0.162 | 0.371 | 2.91E-05 | 0.18541602 |
| WBGene00000134 | amt-2 | 0.270066374 | 0.764639033 | -0.713517523 | 0.023 | 0.108 | 0.003161796 | 1 |
| WBGene00011073 | R07A4.2 | 0.595997094 | 1.090263855 | -0.713076205 | 0.077 | 0.182 | 0.005966075 | 1 |
| WBGene00002266 | lec-3 | 0.329346176 | 0.823407548 | -0.712779891 | 0.031 | 0.114 | 0.005313734 | 1 |
| WBGene00004703 | rsp-6 | 0.44108175 | 0.934573667 | -0.711958342 | 0.054 | 0.159 | 0.0025971 | 1 |
| WBGene00006309 | sul-2 | 0.16386868 | 0.657189428 | -0.711711397 | 0.015 | 0.087 | 0.00550758 | 1 |
| WBGene00000501 | cho-1 | 0.314995006 | 0.804755837 | -0.706575522 | 0.031 | 0.122 | 0.002905378 | 1 |
| WBGene00004401 | rom-2 | 0.197775988 | 0.687024875 | -0.705836944 | 0.023 | 0.122 | 0.001112199 | 1 |
| WBGene00004440 | rpl-26 | 1.897379317 | 2.386313239 | -0.705382545 | 0.377 | 0.619 | 6.72E-05 | 0.428531486 |
| WBGene00019554 | K09C4.10 | 0.246249784 | 0.733851477 | -0.703460544 | 0.023 | 0.099 | 0.006211176 | 1 |
| WBGene00000491 | che-12 | 1.048509271 | 1.535864294 | -0.703104676 | 0.154 | 0.3 | 0.002971145 | 1 |
| WBGene00012340 | dct-15 | 0.372789342 | 0.859550735 | -0.702248248 | 0.038 | 0.151 | 0.000929543 | 1 |
| WBGene00004700 | rsp-3 | 0.797728066 | 1.280925209 | -0.697106122 | 0.108 | 0.251 | 0.000820176 | 1 |
| WBGene00018508 | F46F11.1 | 0.45854849 | 0.941498789 | -0.696750001 | 0.054 | 0.124 | 0.026833855 | 1 |
| WBGene00000879 | cyn-3 | 0.993227528 | 1.475132368 | -0.695241722 | 0.115 | 0.309 | 8.19E-05 | 0.522546292 |

| | | | | | | | | |
|---|---|---|---|---|---|---|---|---|
| WBGene00003162 | mdh-2 | 0.223791569 | 0.705451053 | -0.694887749 | 0.023 | 0.093 | 0.00955088 | 1 |
| WBGene00016500 | memo-1 | 0.362508909 | 0.844114891 | -0.694810562 | 0.038 | 0.135 | 0.003063627 | 1 |
| WBGene00015377 | C03B1.6 | 0 | 0.480761288 | -0.693591926 | 0 | 0.066 | 0.002704595 | 1 |
| WBGene00013311 | sec-61.A | 0.368999897 | 0.849272573 | -0.692887008 | 0.031 | 0.13 | 0.001806101 | 1 |
| WBGene00008262 | ril-1 | 0.466629667 | 0.946824574 | -0.692774811 | 0.046 | 0.147 | 0.002829897 | 1 |
| WBGene00011108 | R07E3.7 | 0 | 0.479369493 | -0.691583991 | 0 | 0.054 | 0.006747802 | 1 |
| WBGene00009772 | ztf-7 | 0.82851799 | 1.307093998 | -0.690439233 | 0.115 | 0.219 | 0.016188043 | 1 |
| WBGene00020390 | T10B5.4 | 1.527341774 | 2.005239783 | -0.689461088 | 0.208 | 0.491 | 3.24E-06 | 0.020699987 |
| WBGene00016734 | C46H11.10 | 0.555445791 | 1.032831312 | -0.688721723 | 0.062 | 0.19 | 0.000877717 | 1 |
| WBGene00019944 | R07G3.8 | 0.22258854 | 0.699716045 | -0.688349486 | 0.023 | 0.101 | 0.005503485 | 1 |
| WBGene00015619 | C08G9.1 | 1.309453777 | 1.785265919 | -0.686451817 | 0.185 | 0.362 | 0.000760894 | 1 |
| WBGene00004425 | rpl-13 | 2.425105144 | 2.900584693 | -0.685971987 | 0.538 | 0.772 | 2.86E-07 | 0.001821873 |
| WBGene00011282 | R74.8 | 0.145878954 | 0.620416255 | -0.684612611 | 0.015 | 0.06 | 0.041505724 | 1 |
| WBGene00010280 | rde-12 | 0 | 0.474273175 | -0.684231558 | 0 | 0.072 | 0.001710099 | 1 |
| WBGene00015767 | hex-2 | 0.485996754 | 0.959899571 | -0.683697244 | 0.062 | 0.166 | 0.003700223 | 1 |
| WBGene00001226 | eif-3.C | 0.19617547 | 0.668876645 | -0.68196364 | 0.023 | 0.095 | 0.008010546 | 1 |
| WBGene00004426 | rpl-14 | 2.08964715 | 2.56042743 | -0.679192375 | 0.392 | 0.66 | 7.01E-06 | 0.044741488 |
| WBGene00008518 | irld-2 | 0.300327265 | 0.770871431 | -0.678851734 | 0.031 | 0.11 | 0.006938209 | 1 |
| WBGene00044764 | W01A8.7 | 0.875060789 | 1.345490249 | -0.67868625 | 0.115 | 0.255 | 0.002410046 | 1 |
| WBGene00019847 | R03G5.6 | 0.274189671 | 0.744189221 | -0.67806602 | 0.023 | 0.104 | 0.004332156 | 1 |
| WBGene00004417 | rpl-6 | 1.979913933 | 2.449280454 | -0.677152752 | 0.346 | 0.656 | 7.85E-07 | 0.005006721 |
| WBGene00012548 | Y37D8A.6 | 2.086362146 | 2.555470227 | -0.676779903 | 0.408 | 0.675 | 4.94E-07 | 0.003149887 |
| WBGene00022042 | icd-2 | 1.503810832 | 1.972904476 | -0.676759074 | 0.215 | 0.472 | 8.38E-06 | 0.053489521 |
| WBGene00020181 | T02H6.11 | 0.469106279 | 0.93668109 | -0.674567861 | 0.054 | 0.13 | 0.020803117 | 1 |
| WBGene00009161 | cox-7C | 0.513980893 | 0.980788519 | -0.673461047 | 0.054 | 0.168 | 0.001702473 | 1 |
| WBGene00011955 | cdka-1 | 0.495727597 | 0.962406308 | -0.673275063 | 0.062 | 0.143 | 0.018013047 | 1 |
| WBGene00004478 | rps-9 | 2.458539921 | 2.924651546 | -0.67245693 | 0.531 | 0.778 | 5.32E-06 | 0.033968589 |
| WBGene00004285 | rab-37 | 1.207940954 | 1.672543565 | -0.670279883 | 0.162 | 0.366 | 0.000155596 | 0.992855231 |
| WBGene00009278 | npr-33 | 0.28827991 | 0.752689072 | -0.670000796 | 0.031 | 0.087 | 0.033165295 | 1 |
| WBGene00003124 | mai-1 | 0.275905647 | 0.738840323 | -0.667873561 | 0.023 | 0.099 | 0.006169983 | 1 |
| WBGene00004505 | rpt-5 | 0.298144593 | 0.760209948 | -0.666619397 | 0.031 | 0.087 | 0.035700238 | 1 |
| WBGene00001945 | his-71 | 2.146230666 | 2.607463044 | -0.665417665 | 0.446 | 0.683 | 1.24E-05 | 0.079024613 |
| WBGene00017792 | F25E5.16 | 0.092581569 | 0.553480217 | -0.664936194 | 0.008 | 0.087 | 0.001805637 | 1 |
| WBGene00000371 | cox-5B | 1.025893476 | 1.486349269 | -0.664297289 | 0.131 | 0.306 | 0.000309043 | 1 |
| WBGene00000912 | daf-16 | 0.944021623 | 1.403773624 | -0.663281931 | 0.123 | 0.257 | 0.004171306 | 1 |
| WBGene00010200 | F57C7.4 | 1.039708089 | 1.499325811 | -0.663088209 | 0.131 | 0.296 | 0.000786679 | 1 |
| WBGene00000481 | cha-1 | 0.725763402 | 1.183958205 | -0.661035369 | 0.092 | 0.201 | 0.008668952 | 1 |
| WBGene00012074 | T27A8.2 | 0.087125162 | 0.5442198 | -0.659448167 | 0.008 | 0.077 | 0.0037607 | 1 |
| WBGene00018803 | fbxa-24 | 2.511475212 | 2.965317365 | -0.654755824 | 0.477 | 0.625 | 0.001859605 | 1 |
| WBGene00015799 | C15H9.3 | 1.83576107 | 2.289495144 | -0.654599898 | 0.323 | 0.571 | 0.000213179 | 1 |
| WBGene00004454 | rpl-36.A | 2.316906262 | 2.769033818 | -0.652282183 | 0.531 | 0.712 | 0.000466005 | 1 |
| WBGene00008977 | iglr-1 | 0.412395468 | 0.864077417 | -0.651639307 | 0.038 | 0.103 | 0.025583302 | 1 |
| WBGene00017984 | F32D1.5 | 0.074829213 | 0.525992592 | -0.650891169 | 0.008 | 0.068 | 0.008081113 | 1 |
| WBGene00004923 | snt-3 | 0.800567802 | 1.251552147 | -0.650632878 | 0.115 | 0.257 | 0.001668439 | 1 |
| WBGene00004467 | rpn-11 | 0.537663011 | 0.987769912 | -0.649366994 | 0.054 | 0.155 | 0.004438158 | 1 |
| WBGene00004494 | rps-25 | 2.940509711 | 3.389157445 | -0.647261861 | 0.662 | 0.863 | 5.44E-10 | 3.47E-06 |
| WBGene00011015 | R04F11.2 | 0.791907771 | 1.240357527 | -0.64697624 | 0.085 | 0.224 | 0.000943481 | 1 |
| WBGene00006538 | tbb-4 | 1.078543893 | 1.526967811 | -0.646938963 | 0.162 | 0.236 | 0.134570445 | 1 |
| WBGene00269383 | ZK384.7 | 0.380226159 | 0.828291774 | -0.646422042 | 0.031 | 0.116 | 0.004646945 | 1 |
| WBGene00003616 | nhr-17 | 0.490404336 | 0.937760218 | -0.645398113 | 0.054 | 0.137 | 0.011688673 | 1 |
| WBGene00012031 | gpdh-3 | 0 | 0.447332627 | -0.645364563 | 0 | 0.064 | 0.003150306 | 1 |
| WBGene00018188 | F38E9.6 | 1.716150342 | 2.162693779 | -0.644226002 | 0.277 | 0.485 | 0.000148734 | 0.949069199 |
| WBGene00017580 | lgc-4 | 0.161038728 | 0.607566522 | -0.644203434 | 0.015 | 0.07 | 0.02010121 | 1 |
| WBGene00006831 | unc-104 | 0.422388883 | 0.867403357 | -0.642020176 | 0.054 | 0.12 | 0.034195402 | 1 |
| WBGene00009380 | F34H10.3 | 0.259315788 | 0.703700838 | -0.641112108 | 0.023 | 0.112 | 0.002310062 | 1 |
| WBGene00018849 | spt-16 | 0.343913405 | 0.78813787 | -0.640880434 | 0.031 | 0.108 | 0.007946991 | 1 |
| WBGene00013899 | ZC443.4 | 1.64679478 | 2.090435234 | -0.640037883 | 0.262 | 0.476 | 0.000179024 | 1 |
| WBGene00001971 | hmg-1.1 | 0.922763001 | 1.365039139 | -0.638069591 | 0.1 | 0.248 | 0.001121686 | 1 |
| WBGene00219950 | F46A8.13 | 2.688705607 | 3.130063189 | -0.636744395 | 0.446 | 0.518 | 0.243223487 | 1 |
| WBGene00044699 | nhr-286 | 0.537705015 | 0.979062351 | -0.63674404 | 0.069 | 0.174 | 0.004489258 | 1 |
| WBGene00017239 | F08B6.1 | 0.238646089 | 0.679034681 | -0.635346438 | 0.023 | 0.11 | 0.002721926 | 1 |
| WBGene00007016 | mdt-15 | 0.332828503 | 0.773056909 | -0.635115338 | 0.038 | 0.116 | 0.009652261 | 1 |
| WBGene00009194 | F27E5.5 | 0.248107018 | 0.688266355 | -0.635015693 | 0.023 | 0.066 | 0.063058538 | 1 |
| WBGene00010642 | mks-6 | 0.695979967 | 1.13588846 | -0.634653802 | 0.085 | 0.207 | 0.002943884 | 1 |

| | | | | | | | | |
|---|---|---|---|---|---|---|---|---|
| WBGene00017961 | nhr-180 | 0.117312701 | 0.556636991 | -0.633810975 | 0.008 | 0.064 | 0.011022903 | 1 |
| WBGene00016493 | C37A2.7 | 2.011400947 | 2.449615463 | -0.63220991 | 0.4 | 0.634 | 0.000135241 | 0.862972984 |
| WBGene00007750 | syg-2 | 0.079142692 | 0.514291191 | -0.627786582 | 0.008 | 0.05 | 0.030856924 | 1 |
| WBGene00010936 | ztf-14 | 2.219990004 | 2.653814732 | -0.625876785 | 0.438 | 0.687 | 2.25E-05 | 0.143708393 |
| WBGene00007041 | tag-180 | 0 | 0.43297296 | -0.624647942 | 0 | 0.05 | 0.009151173 | 1 |
| WBGene00011046 | R05H10.7 | 0 | 0.432676955 | -0.624220897 | 0 | 0.056 | 0.005794642 | 1 |
| WBGene00009716 | F44G4.7 | 0.848153884 | 1.280654727 | -0.623966821 | 0.108 | 0.242 | 0.002522673 | 1 |
| WBGene00001207 | egl-43 | 0.490001008 | 0.922199069 | -0.62353 | 0.046 | 0.139 | 0.005276488 | 1 |
| WBGene00013766 | prmt-1 | 0.367262751 | 0.798886867 | -0.622701971 | 0.038 | 0.106 | 0.021796068 | 1 |
| WBGene00014309 | F07H5.5 | 2.003809723 | 2.434914799 | -0.621953155 | 0.415 | 0.619 | 0.000725712 | 1 |
| WBGene00009337 | uig-1 | 0.202690697 | 0.633254235 | -0.621171882 | 0.023 | 0.087 | 0.014026645 | 1 |
| WBGene00012853 | Y44A6D.2 | 1.639096139 | 2.068778568 | -0.61990071 | 0.238 | 0.435 | 0.000501444 | 1 |
| WBGene00044509 | K07H8.12 | 0.525771669 | 0.955057766 | -0.619328923 | 0.046 | 0.178 | 0.000315448 | 1 |
| WBGene00002178 | jnk-1 | 0.429283869 | 0.858499215 | -0.619226851 | 0.054 | 0.141 | 0.008665719 | 1 |
| WBGene00008762 | ztf-2 | 0.084818126 | 0.513965968 | -0.619129463 | 0.008 | 0.054 | 0.022976989 | 1 |
| WBGene00002019 | hsp-16.48 | 1.008163396 | 1.437026097 | -0.618718092 | 0.131 | 0.203 | 0.087362811 | 1 |
| WBGene00012484 | car-1 | 0.135681507 | 0.563325284 | -0.616959555 | 0.015 | 0.06 | 0.040417434 | 1 |
| WBGene00004415 | rpl-4 | 1.816915047 | 2.244186195 | -0.616421967 | 0.331 | 0.574 | 4.51E-05 | 0.287940037 |
| WBGene00020930 | hlh-30 | 1.058635262 | 1.485789391 | -0.616253144 | 0.138 | 0.279 | 0.004621042 | 1 |
| WBGene00011720 | T11G6.5 | 0.087621691 | 0.514528422 | -0.615896223 | 0.008 | 0.068 | 0.008049571 | 1 |
| WBGene00006839 | unc-115 | 0.284777269 | 0.711117418 | -0.61507882 | 0.023 | 0.085 | 0.017108209 | 1 |
| WBGene00018281 | tbc-18 | 0.411998994 | 0.837123675 | -0.613325269 | 0.038 | 0.097 | 0.040366968 | 1 |
| WBGene00004736 | sca-1 | 0.182998361 | 0.608067618 | -0.61324531 | 0.015 | 0.07 | 0.020518746 | 1 |
| WBGene00304791 | Y62H9A.16 | 0 | 0.424690266 | -0.612698541 | 0 | 0.05 | 0.009151173 | 1 |
| WBGene00012366 | maea-1 | 0.084818126 | 0.509296343 | -0.612392618 | 0.008 | 0.058 | 0.016982541 | 1 |
| WBGene00015285 | gmeb-1 | 0.321388182 | 0.745337847 | -0.611630079 | 0.031 | 0.11 | 0.007359507 | 1 |
| WBGene00008438 | DH11.5 | 0.199192318 | 0.623102524 | -0.611573151 | 0.023 | 0.104 | 0.00399533 | 1 |
| WBGene00019607 | K10B2.4 | 0.241999761 | 0.665678938 | -0.611239848 | 0.023 | 0.091 | 0.011302602 | 1 |
| WBGene00002233 | kqt-1 | 0.656528367 | 1.079866727 | -0.610748153 | 0.069 | 0.17 | 0.00656487 | 1 |
| WBGene00004469 | rps-0 | 2.130311719 | 2.553446925 | -0.610455062 | 0.462 | 0.665 | 0.000322971 | 1 |
| WBGene00021607 | Y46H3C.7 | 0.391633287 | 0.814588592 | -0.610195521 | 0.038 | 0.108 | 0.017687657 | 1 |
| WBGene00004204 | swsn-4 | 0.409021761 | 0.831674138 | -0.609758489 | 0.046 | 0.114 | 0.02404004 | 1 |
| WBGene00006717 | ubc-22 | 0.0991499 | 0.521295769 | -0.609027752 | 0.008 | 0.079 | 0.003326888 | 1 |
| WBGene00018018 | F33G12.6 | 0 | 0.421470613 | -0.608053563 | 0 | 0.072 | 0.0017101 | 1 |
| WBGene00000900 | daf-4 | 0.233755581 | 0.653429509 | -0.605461495 | 0.023 | 0.085 | 0.01610454 | 1 |
| WBGene00008453 | E02A10.3 | 2.41503624 | 2.833730428 | -0.60404803 | 0.492 | 0.75 | 1.11E-05 | 0.071019152 |
| WBGene00018218 | F40A3.3 | 0.149346469 | 0.567597995 | -0.603409403 | 0.015 | 0.05 | 0.082686826 | 1 |
| WBGene00045336 | R11D1.12 | 2.215989031 | 2.633138184 | -0.601819014 | 0.354 | 0.563 | 0.0002789 | 1 |
| WBGene00009216 | F28D1.6 | 2.21978702 | 2.636430886 | -0.601090039 | 0.454 | 0.65 | 0.00027322 | 1 |
| WBGene00004427 | rpl-15 | 2.205958393 | 2.622176498 | -0.600475795 | 0.462 | 0.712 | 7.45E-06 | 0.047527229 |
| WBGene00000379 | cct-4 | 0.447245155 | 0.863251011 | -0.600169585 | 0.046 | 0.108 | 0.038399347 | 1 |
| WBGene00016099 | C25E10.10 | 2.232411483 | 2.648411483 | -0.600161138 | 0.462 | 0.658 | 0.000685274 | 1 |
| WBGene00002295 | let-19 | 0.220754288 | 0.636650762 | -0.60001178 | 0.023 | 0.081 | 0.019902173 | 1 |
| WBGene00004488 | rps-19 | 2.394811532 | 2.810625212 | -0.599892335 | 0.508 | 0.743 | 3.63E-06 | 0.023132767 |
| WBGene00044324 | ufm-1 | 0 | 0.415793846 | -0.599863719 | 0 | 0.062 | 0.003669115 | 1 |
| WBGene00044396 | ZK1290.14 | 0.28762031 | 0.701539777 | -0.597159562 | 0.031 | 0.056 | 0.251697297 | 1 |
| WBGene00022738 | fubl-2 | 0.179620113 | 0.592975076 | -0.596345156 | 0.023 | 0.083 | 0.018599669 | 1 |
| WBGene00004485 | rps-16 | 1.659603853 | 2.072599173 | -0.595826301 | 0.3 | 0.474 | 0.003241632 | 1 |
| WBGene00000914 | daf-19 | 0.378975751 | 0.791883386 | -0.595699797 | 0.031 | 0.077 | 0.067787146 | 1 |
| WBGene00005704 | sru-41 | 0.539284653 | 0.951971993 | -0.595381978 | 0.062 | 0.147 | 0.014592049 | 1 |
| WBGene00004486 | rps-17 | 2.48751905 | 2.900050138 | -0.595156555 | 0.492 | 0.764 | 2.20E-06 | 0.014059679 |
| WBGene00006752 | unc-13 | 0.953308767 | 1.365580427 | -0.594782279 | 0.138 | 0.277 | 0.005442387 | 1 |
| WBGene00006604 | tra-1 | 0.355915795 | 0.767386442 | -0.593626662 | 0.031 | 0.11 | 0.007003255 | 1 |
| WBGene00015943 | tiar-1 | 0.150165337 | 0.561487855 | -0.593412956 | 0.015 | 0.06 | 0.040687213 | 1 |
| WBGene00000288 | cal-4 | 1.71365374 | 2.123855292 | -0.591795745 | 0.269 | 0.534 | 2.57E-05 | 0.164003234 |
| WBGene00022199 | pfk-1.1 | 0.041197077 | 0.450968613 | -0.591175363 | 0.008 | 0.06 | 0.013481501 | 1 |
| WBGene00011156 | rbm-3.2 | 2.243441627 | 2.653169544 | -0.591112433 | 0.485 | 0.691 | 6.83E-05 | 0.435983707 |
| WBGene00001352 | evl-14 | 0 | 0.40967093 | -0.590030219 | 0 | 0.05 | 0.009151167 | 1 |
| WBGene00004464 | rpn-8 | 0.346207285 | 0.75418328 | -0.588584944 | 0.023 | 0.135 | 0.000417087 | 1 |
| WBGene00016033 | C24A3.2 | 0.201442408 | 0.609378663 | -0.588527612 | 0.015 | 0.103 | 0.001746236 | 1 |
| WBGene00194866 | Y41C4A.22 | 0.496261711 | 0.903672741 | -0.587769873 | 0.054 | 0.124 | 0.026734187 | 1 |
| WBGene00004470 | rps-1 | 2.524889037 | 2.932255069 | -0.587704955 | 0.569 | 0.797 | 7.63E-06 | 0.048674641 |
| WBGene00009848 | F48F5.6 | 0.543655461 | 0.950051509 | -0.586305563 | 0.054 | 0.176 | 0.001007684 | 1 |
| WBGene00002244 | laf-1 | 0.101479759 | 0.507461771 | -0.585708236 | 0.015 | 0.056 | 0.051232889 | 1 |

| | | | | | | | | |
|---|---|---|---|---|---|---|---|---|
| WBGene00014472 | MTCE.33 | 1.48514219 | 1.890610168 | -0.584966641 | 0.223 | 0.319 | 0.087756138 | 1 |
| WBGene00000833 | cts-1 | 0.543823105 | 0.948465183 | -0.583775119 | 0.062 | 0.153 | 0.010185819 | 1 |
| WBGene00006786 | unc-51 | 0.249117433 | 0.653534312 | -0.583450225 | 0.023 | 0.089 | 0.013193253 | 1 |
| WBGene00001677 | gpa-15 | 0.267657012 | 0.671624847 | -0.582802393 | 0.023 | 0.064 | 0.075422406 | 1 |
| WBGene00008956 | nekl-3 | 0.794672869 | 1.198002736 | -0.581882 | 0.085 | 0.228 | 0.000765027 | 1 |
| WBGene00000906 | daf-10 | 0.649099328 | 1.051853457 | -0.581051384 | 0.062 | 0.143 | 0.017771499 | 1 |
| WBGene00020279 | glb-25 | 1.397836845 | 1.800057697 | -0.580282029 | 0.231 | 0.408 | 0.001760781 | 1 |
| WBGene00004433 | rpl-21 | 2.315591959 | 2.716698867 | -0.578674947 | 0.5 | 0.716 | 9.90E-05 | 0.63160627 |
| WBGene00000122 | aly-3 | 0.60452697 | 1.004652935 | -0.577259744 | 0.062 | 0.162 | 0.005774798 | 1 |
| WBGene00006705 | ubc-8 | 0.740211371 | 1.139341797 | -0.575823486 | 0.085 | 0.205 | 0.004115397 | 1 |
| WBGene00001171 | egl-2 | 0.278551466 | 0.675872466 | -0.573213036 | 0.023 | 0.091 | 0.011375918 | 1 |
| WBGene00004489 | rps-20 | 2.420317279 | 2.816012849 | -0.570868037 | 0.577 | 0.766 | 3.34E-05 | 0.212948282 |
| WBGene00001560 | gei-3 | 0.31249607 | 0.70801887 | -0.570618782 | 0.031 | 0.085 | 0.039255911 | 1 |
| WBGene00045419 | lsy-12 | 0.275105128 | 0.67021527 | -0.570023442 | 0.023 | 0.062 | 0.086532282 | 1 |
| WBGene00022134 | hrpu-2 | 0.082452944 | 0.477275516 | -0.569608567 | 0.008 | 0.06 | 0.014741981 | 1 |
| WBGene00022592 | klu-2 | 0.623998739 | 1.018814335 | -0.569598502 | 0.069 | 0.147 | 0.027958991 | 1 |
| WBGene00012019 | dkf-2 | 0.406762811 | 0.801511384 | -0.569501808 | 0.038 | 0.124 | 0.006847961 | 1 |
| WBGene00006603 | tps-2 | 0.638910722 | 1.032912041 | -0.568423749 | 0.069 | 0.161 | 0.013385759 | 1 |
| WBGene00013711 | Y106G6G.2 | 0.353658003 | 0.747656675 | -0.56841993 | 0.046 | 0.108 | 0.039469318 | 1 |
| WBGene00011531 | rsbp-1 | 0.679006194 | 1.072415401 | -0.567569512 | 0.062 | 0.197 | 0.000561583 | 1 |
| WBGene00004445 | rpl-31 | 2.630112555 | 3.023390366 | -0.567379948 | 0.623 | 0.778 | 7.52E-05 | 0.479785557 |
| WBGene00044303 | hst-3.2 | 0.153868545 | 0.54711451 | -0.567334003 | 0.015 | 0.075 | 0.013164875 | 1 |
| WBGene00002242 | kvs-1 | 0.689888162 | 1.082827533 | -0.566891682 | 0.085 | 0.18 | 0.015017865 | 1 |
| WBGene00004493 | rps-24 | 2.012948015 | 2.405199845 | -0.565899771 | 0.438 | 0.609 | 0.001353675 | 1 |
| WBGene00001168 | eef-1A.1 | 2.348725554 | 2.740495982 | -0.565205254 | 0.523 | 0.741 | 0.000205704 | 1 |
| WBGene00020868 | eif-1 | 0.730966839 | 1.122640181 | -0.565065188 | 0.092 | 0.162 | 0.068947823 | 1 |
| WBGene00004491 | rps-22 | 2.256590622 | 2.647973437 | -0.564646047 | 0.415 | 0.716 | 6.32E-06 | 0.040301925 |
| WBGene00019230 | ttll-11 | 0.727089394 | 1.118104084 | -0.564114955 | 0.092 | 0.195 | 0.012054888 | 1 |
| WBGene00303425 | R02E12.12 | 0.157461568 | 0.546848202 | -0.561766166 | 0.015 | 0.087 | 0.005567149 | 1 |
| WBGene00006924 | vig-1 | 0.217961679 | 0.606210051 | -0.560124002 | 0.023 | 0.068 | 0.055844432 | 1 |
| WBGene00009122 | tct-1 | 0.672347187 | 1.060220362 | -0.559582707 | 0.077 | 0.18 | 0.008151469 | 1 |
| WBGene00044728 | Y53F4B.45 | 1.523868665 | 1.910524059 | -0.557825818 | 0.162 | 0.342 | 0.000368045 | 1 |
| WBGene00011695 | T10G3.4 | 1.134360112 | 1.517513381 | -0.55277332 | 0.138 | 0.263 | 0.013650569 | 1 |
| WBGene00011259 | miz-1 | 0.134900662 | 0.518021105 | -0.552725962 | 0.015 | 0.074 | 0.014781951 | 1 |
| WBGene00001026 | dnj-8 | 0.065575767 | 0.448366078 | -0.552249684 | 0.008 | 0.052 | 0.026171211 | 1 |
| WBGene00000230 | atp-3 | 0.570672505 | 0.953430069 | -0.552202438 | 0.054 | 0.164 | 0.002424792 | 1 |
| WBGene00018193 | F39C12.1 | 0.308926001 | 0.691656185 | -0.552162939 | 0.031 | 0.066 | 0.142498388 | 1 |
| WBGene00002344 | let-70 | 1.743450732 | 2.126028753 | -0.551943413 | 0.323 | 0.524 | 0.001691996 | 1 |
| WBGene00000280 | cah-2 | 0.265702142 | 0.647065797 | -0.550191454 | 0.031 | 0.095 | 0.02078869 | 1 |
| WBGene00004487 | rps-18 | 2.343754737 | 2.724522901 | -0.549332343 | 0.5 | 0.716 | 9.75E-05 | 0.62193517 |
| WBGene00001117 | dyf-1 | 0.687108452 | 1.066539117 | -0.547402738 | 0.085 | 0.199 | 0.005104698 | 1 |
| WBGene00006479 | tmbi-4 | 0.281040795 | 0.660133567 | -0.546915263 | 0.031 | 0.103 | 0.012362284 | 1 |
| WBGene00016603 | met-1 | 0.763143071 | 1.142158763 | -0.546804059 | 0.077 | 0.178 | 0.009148913 | 1 |
| WBGene00020827 | T26A8.4 | 0.220352533 | 0.599121461 | -0.546448054 | 0.023 | 0.074 | 0.037902665 | 1 |
| WBGene00004699 | rsp-2 | 0.759981 | 1.137875965 | -0.545187191 | 0.085 | 0.176 | 0.020246164 | 1 |
| WBGene00015056 | B0222.5 | 0.073121292 | 0.45067389 | -0.544693261 | 0.008 | 0.056 | 0.019414252 | 1 |
| WBGene00001037 | dnj-19 | 0.462802766 | 0.839920001 | -0.544065164 | 0.054 | 0.13 | 0.020387567 | 1 |
| WBGene00016405 | hecd-1 | 0.397766315 | 0.773298438 | -0.541778332 | 0.046 | 0.106 | 0.040104304 | 1 |
| WBGene00009784 | F46C5.9 | 0.476129491 | 0.850805998 | -0.540543939 | 0.054 | 0.135 | 0.014035799 | 1 |
| WBGene00017925 | F29B9.11 | 0.743293563 | 1.117500123 | -0.539865949 | 0.092 | 0.199 | 0.008846647 | 1 |
| WBGene00021369 | siah-1 | 0.627224839 | 1.001216001 | -0.539555195 | 0.069 | 0.161 | 0.013879295 | 1 |
| WBGene00001853 | hil-2 | 1.662403319 | 2.035197985 | -0.537829015 | 0.231 | 0.424 | 0.000732272 | 1 |
| WBGene00003412 | mrp-6 | 0.512369275 | 0.884726796 | -0.537198349 | 0.062 | 0.093 | 0.286859161 | 1 |
| WBGene00003630 | nhr-40 | 0.240225238 | 0.612558836 | -0.537163836 | 0.023 | 0.085 | 0.016946174 | 1 |
| WBGene00003519 | nac-3 | 0.339934182 | 0.711743185 | -0.536407005 | 0.038 | 0.114 | 0.012991372 | 1 |
| WBGene00011860 | pigc-1 | 0.308603895 | 0.679383601 | -0.534922044 | 0.031 | 0.093 | 0.02288872 | 1 |
| WBGene00004830 | slo-1 | 0.071158826 | 0.441801776 | -0.534724747 | 0.008 | 0.056 | 0.019791546 | 1 |
| WBGene00010868 | somi-1 | 0.148962948 | 0.519231457 | -0.534184541 | 0.015 | 0.056 | 0.054355386 | 1 |
| WBGene00006697 | uaf-1 | 0.281256182 | 0.648716393 | -0.530133024 | 0.031 | 0.079 | 0.05833929 | 1 |
| WBGene00001086 | dpy-27 | 0.834061936 | 1.201351861 | -0.529887353 | 0.092 | 0.197 | 0.01016059 | 1 |
| WBGene00004476 | rps-7 | 2.409031863 | 2.776132841 | -0.529614761 | 0.562 | 0.706 | 0.003466181 | 1 |
| WBGene00010480 | tasp-1 | 1.227001076 | 1.592942113 | -0.527941318 | 0.192 | 0.335 | 0.008796272 | 1 |
| WBGene00004416 | rpl-5 | 2.221298622 | 2.586704158 | -0.527168754 | 0.446 | 0.689 | 0.000512104 | 1 |
| WBGene00019691 | K12H6.12 | 0.742577412 | 1.10691495 | -0.52562796 | 0.077 | 0.166 | 0.019284557 | 1 |

| | | | | | | | | |
|---|---|---|---|---|---|---|---|---|
| WBGene00017653 | F21C10.3 | 1.823792686 | 2.187792467 | -0.525140678 | 0.315 | 0.526 | 0.000938693 | 1 |
| WBGene00022603 | gck-2 | 0.234758267 | 0.59819693 | -0.524331158 | 0.031 | 0.066 | 0.134272079 | 1 |
| WBGene00006384 | taf-3 | 0.162113312 | 0.525377336 | -0.524079206 | 0.015 | 0.074 | 0.014375629 | 1 |
| WBGene00044134 | D1025.10 | 0.225622905 | 0.587997891 | -0.522796596 | 0.023 | 0.075 | 0.033622692 | 1 |
| WBGene00012094 | T27E9.2 | 0.496065559 | 0.858304774 | -0.52260072 | 0.054 | 0.118 | 0.042407161 | 1 |
| WBGene00021959 | Y57E12AL.6 | 0.340014221 | 0.70154912 | -0.521584605 | 0.038 | 0.108 | 0.018860398 | 1 |
| WBGene00194657 | T07D10.10 | 0.222151691 | 0.58348097 | -0.521287959 | 0.023 | 0.068 | 0.057347294 | 1 |
| WBGene00019492 | K07E3.4 | 0.283245918 | 0.643907673 | -0.520324925 | 0.023 | 0.101 | 0.005708936 | 1 |
| WBGene00006911 | vha-2 | 0.611286261 | 0.971782984 | -0.520086833 | 0.077 | 0.151 | 0.045012204 | 1 |
| WBGene00009440 | idhg-1 | 0.219921771 | 0.580154103 | -0.519705399 | 0.023 | 0.097 | 0.007303925 | 1 |
| WBGene00013415 | ilcr-1 | 0.255398746 | 0.615507924 | -0.519527726 | 0.023 | 0.095 | 0.008639071 | 1 |
| WBGene00012645 | mrpl-22 | 0.359965966 | 0.719652883 | -0.518918531 | 0.038 | 0.095 | 0.041091816 | 1 |
| WBGene00005738 | srv-27 | 0.073121292 | 0.432748425 | -0.518832281 | 0.008 | 0.052 | 0.026372374 | 1 |
| WBGene00006725 | ubl-1 | 2.290808597 | 2.650426223 | -0.518818567 | 0.477 | 0.698 | 0.000378793 | 1 |
| WBGene00020196 | T03G11.3 | 1.154824159 | 1.514370177 | -0.518715257 | 0.162 | 0.34 | 0.000608396 | 1 |
| WBGene00009357 | F33C8.2 | 0.175929493 | 0.535091748 | -0.518161604 | 0.015 | 0.074 | 0.015463832 | 1 |
| WBGene00004436 | rpl-24.1 | 2.838997728 | 3.197946035 | -0.517852943 | 0.677 | 0.839 | 2.10E-05 | 0.134290424 |
| WBGene00004363 | ric-3 | 0.421495687 | 0.780151436 | -0.517430871 | 0.046 | 0.089 | 0.124818618 | 1 |
| WBGene00001423 | fib-1 | 0.188055542 | 0.546186963 | -0.516674425 | 0.023 | 0.064 | 0.070357875 | 1 |
| WBGene00001187 | egl-19 | 0.321328977 | 0.679282241 | -0.5164174 | 0.031 | 0.07 | 0.107895885 | 1 |
| WBGene00044638 | F23A7.8 | 1.158518799 | 1.516285454 | -0.51614818 | 0.054 | 0.097 | 0.132754681 | 1 |
| WBGene00019667 | spe-49 | 0.129450126 | 0.487189817 | -0.516109278 | 0.015 | 0.068 | 0.022873782 | 1 |
| WBGene00000488 | che-7 | 0.497690117 | 0.854636341 | -0.514964547 | 0.038 | 0.099 | 0.033792423 | 1 |
| WBGene00004499 | rps-30 | 2.485462232 | 2.84061487 | -0.512376949 | 0.569 | 0.752 | 0.000197526 | 1 |
| WBGene00001648 | goa-1 | 0.559854839 | 0.91436234 | -0.511446213 | 0.062 | 0.149 | 0.011848265 | 1 |
| WBGene00003072 | lrp-2 | 0.285241167 | 0.6395937 | -0.511222642 | 0.023 | 0.087 | 0.015194912 | 1 |
| WBGene00005605 | srj-17 | 0.160733541 | 0.514529668 | -0.510419918 | 0.015 | 0.072 | 0.018030824 | 1 |
| WBGene00003476 | mtm-3 | 1.117798797 | 1.471003379 | -0.509566499 | 0.169 | 0.3 | 0.011459299 | 1 |
| WBGene00022400 | rpb-9 | 0.233501566 | 0.586524782 | -0.509304843 | 0.023 | 0.068 | 0.058367771 | 1 |
| WBGene00004411 | rpc-1 | 0.424126871 | 0.776654943 | -0.5085905 | 0.046 | 0.106 | 0.043538695 | 1 |
| WBGene00012187 | rpb-11 | 0.120206988 | 0.472150996 | -0.507747875 | 0.015 | 0.068 | 0.022795744 | 1 |
| WBGene00004418 | rpl-7 | 2.379285404 | 2.730656633 | -0.506921529 | 0.546 | 0.712 | 0.00055829 | 1 |
| WBGene00000058 | acr-19 | 0.153160312 | 0.504475745 | -0.506841033 | 0.015 | 0.087 | 0.005667712 | 1 |
| WBGene00008149 | pyp-1 | 0.479628196 | 0.830806395 | -0.506643045 | 0.054 | 0.12 | 0.03593389 | 1 |
| WBGene00015113 | B0281.6 | 0.426890084 | 0.777660536 | -0.506054791 | 0.046 | 0.106 | 0.046758804 | 1 |
| WBGene00044557 | ZC13.10 | 1.804894299 | 2.154911587 | -0.504968205 | 0.285 | 0.52 | 6.19E-05 | 0.395020548 |
| WBGene00022951 | C30E1.9 | 4.841646658 | 5.191533459 | -0.504779951 | 0.992 | 0.925 | 1.88E-08 | 0.000119939 |
| WBGene00014307 | F07H5.3 | 2.938676716 | 3.2868352 | -0.502286519 | 0.677 | 0.822 | 5.62E-06 | 0.035857231 |
| WBGene00004409 | rla-1 | 2.274769509 | 2.621382119 | -0.500056293 | 0.462 | 0.687 | 0.000722718 | 1 |
| WBGene00003948 | pbs-2 | 0.36083719 | 0.707266263 | -0.499791506 | 0.031 | 0.095 | 0.02140504 | 1 |
| WBGene00001574 | gei-17 | 0.338192217 | 0.684363879 | -0.499420139 | 0.031 | 0.083 | 0.046006002 | 1 |
| WBGene00012030 | T25G3.3 | 0.097514596 | 0.443530697 | -0.499195713 | 0.008 | 0.072 | 0.006086221 | 1 |
| WBGene00006439 | ant-1.1 | 2.575991429 | 2.921969825 | -0.499141316 | 0.6 | 0.764 | 0.000191686 | 1 |
| WBGene00002151 | irk-3 | 1.179755602 | 1.525253272 | -0.498447774 | 0.138 | 0.325 | 0.000170459 | 1 |
| WBGene00000241 | bbs-1 | 0.170660344 | 0.515854986 | -0.498010599 | 0.023 | 0.066 | 0.062689498 | 1 |
| WBGene00009050 | F22D6.2 | 2.021196261 | 2.365941382 | -0.497362075 | 0.415 | 0.619 | 0.00391542 | 1 |
| WBGene00019871 | R04E5.8 | 0.650845525 | 0.994754183 | -0.496155316 | 0.062 | 0.164 | 0.00527401 | 1 |
| WBGene00001480 | fmo-5 | 1.497751046 | 1.840567245 | -0.49457923 | 0.177 | 0.255 | 0.085392909 | 1 |
| WBGene00015484 | atgl-1 | 0.130203422 | 0.472761366 | -0.494206647 | 0.015 | 0.058 | 0.044606505 | 1 |
| WBGene00004479 | rps-10 | 2.502245302 | 2.843235793 | -0.49194529 | 0.592 | 0.772 | 0.000644706 | 1 |
| WBGene00249815 | F20D12.12 | 0.229592235 | 0.570252807 | -0.491469318 | 0.023 | 0.066 | 0.067237739 | 1 |
| WBGene00018985 | F56F10.2 | 0.232164962 | 0.572498659 | -0.490997736 | 0.023 | 0.07 | 0.050630817 | 1 |
| WBGene00007352 | cdc-48.1 | 0.258432193 | 0.598408387 | -0.490481969 | 0.031 | 0.083 | 0.044131876 | 1 |
| WBGene00000434 | ceh-9 | 0.21286962 | 0.551592679 | -0.488674078 | 0.015 | 0.077 | 0.011380317 | 1 |
| WBGene00044294 | C01B10.11 | 0.186404125 | 0.524563226 | -0.487860458 | 0.023 | 0.058 | 0.10900842 | 1 |
| WBGene00020893 | T28C12.6 | 0.290988765 | 0.62839996 | -0.486781459 | 0.031 | 0.083 | 0.045185784 | 1 |
| WBGene00020260 | T05E7.4 | 0.382441502 | 0.718203746 | -0.484402524 | 0.031 | 0.12 | 0.003735046 | 1 |
| WBGene00018898 | F55F10.1 | 0.079717115 | 0.415162929 | -0.483946012 | 0.008 | 0.05 | 0.030856924 | 1 |
| WBGene00005359 | srh-142 | 0.473556287 | 0.808362119 | -0.483022715 | 0.054 | 0.137 | 0.013202997 | 1 |
| WBGene00017834 | F26F12.3 | 2.132585755 | 2.467322957 | -0.482923701 | 0.415 | 0.578 | 0.014375437 | 1 |
| WBGene00004701 | rsp-4 | 0.056185084 | 0.390884631 | -0.482869376 | 0.008 | 0.06 | 0.014571879 | 1 |
| WBGene00011198 | R10E4.1 | 0.400257183 | 0.734674906 | -0.482462791 | 0.031 | 0.126 | 0.002502309 | 1 |
| WBGene00019983 | sti-1 | 0.452221403 | 0.785730727 | -0.481152247 | 0.046 | 0.132 | 0.009924303 | 1 |
| WBGene00018395 | mtch-1 | 0.766848777 | 1.100311142 | -0.4810845 | 0.085 | 0.168 | 0.036921289 | 1 |

| | | | | | | | | |
|---|---|---|---|---|---|---|---|---|
| WBGene00020333 | sre-12 | 0.470279371 | 0.803653652 | -0.480957423 | 0.054 | 0.087 | 0.231984192 | 1 |
| WBGene00011564 | ugt-50 | 0.072432462 | 0.405603114 | -0.480663648 | 0.008 | 0.05 | 0.030739557 | 1 |
| WBGene00015776 | dct-1 | 1.555151093 | 1.887619314 | -0.479650253 | 0.231 | 0.437 | 0.000553925 | 1 |
| WBGene00020894 | T28D9.1 | 1.774866418 | 2.107296546 | -0.479595296 | 0.292 | 0.499 | 0.001078054 | 1 |
| WBGene00013924 | rad-23 | 0.222819676 | 0.555012014 | -0.479252239 | 0.015 | 0.054 | 0.063822734 | 1 |
| WBGene00003830 | num-1 | 0.771637233 | 1.103462523 | -0.4787227 | 0.085 | 0.197 | 0.005807563 | 1 |
| WBGene00016841 | magu-3 | 0.041197077 | 0.37274471 | -0.478322127 | 0.008 | 0.052 | 0.024704319 | 1 |
| WBGene00018704 | ztf-13 | 0.383064028 | 0.714496488 | -0.478155966 | 0.038 | 0.112 | 0.014240747 | 1 |
| WBGene00009676 | F44A6.4 | 0.167225717 | 0.49827102 | -0.477597418 | 0.023 | 0.056 | 0.11912028 | 1 |
| WBGene00010516 | K02E11.7 | 0.3743854 | 0.705313273 | -0.477428002 | 0.023 | 0.074 | 0.038714911 | 1 |
| WBGene00014886 | Y41E3.5 | 2.904913306 | 3.235720281 | -0.477253583 | 0.638 | 0.778 | 7.57E-05 | 0.483237933 |
| WBGene00001999 | hrp-1 | 1.389318672 | 1.719197901 | -0.475915129 | 0.223 | 0.393 | 0.003440507 | 1 |
| WBGene00004368 | ric-19 | 0.400568678 | 0.730033114 | -0.475316709 | 0.038 | 0.101 | 0.031696247 | 1 |
| WBGene00004439 | rpl-25.2 | 1.116409861 | 1.445120036 | -0.47422854 | 0.154 | 0.29 | 0.008011755 | 1 |
| WBGene00022401 | Y97E10AR.6 | 0.424812311 | 0.753451421 | -0.474126014 | 0.046 | 0.118 | 0.021109476 | 1 |
| WBGene00011471 | T05C12.8 | 1.124417727 | 1.45303716 | -0.474097627 | 0.154 | 0.304 | 0.003165833 | 1 |
| WBGene00021068 | W06H8.6 | 0.574950859 | 0.903045771 | -0.473340903 | 0.062 | 0.13 | 0.044841819 | 1 |
| WBGene00004444 | rpl-30 | 1.696365914 | 2.024403149 | -0.473257692 | 0.277 | 0.491 | 0.000612405 | 1 |
| WBGene00020683 | ribo-1 | 0.198167169 | 0.525781594 | -0.472647705 | 0.023 | 0.06 | 0.097547094 | 1 |
| WBGene00016899 | C53C11.1 | 0.185455666 | 0.513048922 | -0.472617166 | 0.015 | 0.058 | 0.048447897 | 1 |
| WBGene00004822 | skr-16 | 0.224166366 | 0.549781456 | -0.469763276 | 0.023 | 0.077 | 0.030372185 | 1 |
| WBGene00004472 | rps-3 | 2.208142043 | 2.533648426 | -0.469606444 | 0.438 | 0.665 | 0.000334209 | 1 |
| WBGene00003619 | nhr-20 | 1.095163199 | 1.419189623 | -0.467471314 | 0.154 | 0.304 | 0.003363063 | 1 |
| WBGene00004760 | sel-2 | 0.514206684 | 0.836097488 | -0.464390268 | 0.054 | 0.103 | 0.110481922 | 1 |
| WBGene00006650 | tts-1 | 0.671209673 | 0.993085594 | -0.464368794 | 0.054 | 0.066 | 0.612909761 | 1 |
| WBGene00020917 | W01B11.6 | 1.61785064 | 1.939585668 | -0.464165529 | 0.269 | 0.431 | 0.006006416 | 1 |
| WBGene00004888 | smo-1 | 0.934966592 | 1.255343836 | -0.462206662 | 0.123 | 0.24 | 0.012484039 | 1 |
| WBGene00016212 | C29F5.3 | 0.534674947 | 0.853290011 | -0.459664374 | 0.054 | 0.122 | 0.036126518 | 1 |
| WBGene00003889 | osm-9 | 0.323186199 | 0.641643635 | -0.459436963 | 0.038 | 0.091 | 0.053867824 | 1 |
| WBGene00003754 | nlp-16 | 0.78291568 | 1.101198313 | -0.459184776 | 0.077 | 0.176 | 0.009139405 | 1 |
| WBGene00003559 | ncl-1 | 0.434563989 | 0.752679882 | -0.458944221 | 0.046 | 0.108 | 0.040665345 | 1 |
| WBGene00017790 | F25E5.9 | 0.214730681 | 0.532214683 | -0.458032595 | 0.015 | 0.062 | 0.036140328 | 1 |
| WBGene00012148 | inos-1 | 0.343130925 | 0.660573807 | -0.457973271 | 0.038 | 0.093 | 0.053824302 | 1 |
| WBGene00007327 | C05C9.3 | 0.601492156 | 0.91877174 | -0.457737682 | 0.069 | 0.143 | 0.04064383 | 1 |
| WBGene00004916 | snr-3 | 0.409042935 | 0.726225633 | -0.457597907 | 0.046 | 0.108 | 0.041172674 | 1 |
| WBGene00007042 | pbrm-1 | 0.357169719 | 0.674111046 | -0.45724968 | 0.038 | 0.108 | 0.018860408 | 1 |
| WBGene00004442 | rpl-28 | 1.773448576 | 2.089595497 | -0.456103595 | 0.323 | 0.503 | 0.002403562 | 1 |
| WBGene00011069 | ceh-62 | 0.29528536 | 0.61059999 | -0.454902852 | 0.031 | 0.079 | 0.05833929 | 1 |
| WBGene00206380 | C42D4.18 | 1.28382416 | 1.598440122 | -0.453894888 | 0.162 | 0.342 | 0.000917596 | 1 |
| WBGene00003587 | ned-8 | 0.178782761 | 0.493136654 | -0.453516802 | 0.023 | 0.052 | 0.15863089 | 1 |
| WBGene00044677 | ZK353.10 | 1.874784156 | 2.18897524 | -0.453281918 | 0.3 | 0.507 | 0.000714197 | 1 |
| WBGene00004428 | rpl-16 | 2.285329684 | 2.599226239 | -0.452857004 | 0.469 | 0.658 | 0.000791194 | 1 |
| WBGene00001167 | eef-2 | 2.06330007 | 2.377020366 | -0.452602715 | 0.392 | 0.621 | 0.001783838 | 1 |
| WBGene00008930 | F18A11.3 | 0.081491558 | 0.394906205 | -0.452161757 | 0.008 | 0.054 | 0.022976989 | 1 |
| WBGene00013548 | lury-1 | 1.060719534 | 1.374018156 | -0.451994367 | 0.138 | 0.234 | 0.032707578 | 1 |
| WBGene00008505 | F01G4.6 | 0.842835534 | 1.156110788 | -0.451960655 | 0.108 | 0.168 | 0.144051386 | 1 |
| WBGene00016998 | D1005.4 | 1.497332354 | 1.810467467 | -0.451758475 | 0.231 | 0.356 | 0.047987078 | 1 |
| WBGene00004421 | rpl-10 | 2.70783737 | 3.019048001 | -0.448982033 | 0.654 | 0.801 | 0.000736118 | 1 |
| WBGene00006703 | ubc-6 | 0.205105852 | 0.516159821 | -0.448756018 | 0.023 | 0.081 | 0.022432943 | 1 |
| WBGene00003357 | mir-264 | 0.573443607 | 0.884288572 | -0.448454489 | 0.062 | 0.149 | 0.011878359 | 1 |
| WBGene00011250 | R11H6.2 | 0.44522189 | 0.755647957 | -0.447850148 | 0.038 | 0.056 | 0.457332275 | 1 |
| WBGene00008932 | F18A11.5 | 2.304935899 | 2.615219309 | -0.447644337 | 0.454 | 0.617 | 0.001047012 | 1 |
| WBGene00015364 | npr-19 | 0.185682562 | 0.495766012 | -0.447355855 | 0.008 | 0.052 | 0.027087004 | 1 |
| WBGene00001209 | egl-45 | 0.176391556 | 0.486349662 | -0.447175022 | 0.023 | 0.056 | 0.123483732 | 1 |
| WBGene00020146 | got-1.2 | 0.429448173 | 0.739264465 | -0.446970428 | 0.046 | 0.114 | 0.027406847 | 1 |
| WBGene00003172 | mec-8 | 0.226367218 | 0.53590638 | -0.446570614 | 0.023 | 0.07 | 0.051387088 | 1 |
| WBGene00045517 | R11D1.13 | 0.069050062 | 0.37848373 | -0.446418418 | 0.008 | 0.054 | 0.023065192 | 1 |
| WBGene00020838 | trak-1 | 0.849986808 | 1.159155926 | -0.446036753 | 0.085 | 0.186 | 0.01274945 | 1 |
| WBGene00015070 | srt-70 | 0.26283201 | 0.571553008 | -0.445390252 | 0.023 | 0.072 | 0.045087443 | 1 |
| WBGene00010626 | K07C5.3 | 0.079142692 | 0.386937679 | -0.444054302 | 0.008 | 0.05 | 0.030856924 | 1 |
| WBGene00004836 | sls-2.4 | 0.754025246 | 1.061772631 | -0.443985627 | 0.069 | 0.159 | 0.014435637 | 1 |
| WBGene00004446 | rpl-32 | 2.651131759 | 2.958820893 | -0.443901588 | 0.577 | 0.779 | 0.000278261 | 1 |
| WBGene00119203 | T04A8.18 | 0.646920798 | 0.954569798 | -0.443843686 | 0.085 | 0.168 | 0.032587705 | 1 |
| WBGene00010419 | atp-1 | 1.296530995 | 1.60359184 | -0.442995158 | 0.185 | 0.304 | 0.022418156 | 1 |

| | | | | | | | | |
|---|---|---|---|---|---|---|---|---|
| WBGene00017198 | nhr-36 | 0.27553238 | 0.581970041 | -0.442096094 | 0.031 | 0.079 | 0.057240244 | 1 |
| WBGene00006067 | sto-5 | 0.315253875 | 0.621551256 | -0.441893713 | 0.015 | 0.095 | 0.00305808 | 1 |
| WBGene00001679 | gpb-1 | 1.484163785 | 1.790295034 | -0.441654035 | 0.285 | 0.385 | 0.127543092 | 1 |
| WBGene00004477 | rps-8 | 2.5313388 | 2.837387057 | -0.441534303 | 0.515 | 0.745 | 0.00013049 | 0.832657228 |
| WBGene00016803 | bath-42 | 0.151067807 | 0.456506882 | -0.440655439 | 0.015 | 0.064 | 0.03188707 | 1 |
| WBGene00003002 | lin-13 | 0.299592874 | 0.603897454 | -0.439018708 | 0.038 | 0.074 | 0.159548593 | 1 |
| WBGene00044144 | M153.4 | 1.085466492 | 1.389640117 | -0.438829781 | 0.123 | 0.284 | 0.00115242 | 1 |
| WBGene00019243 | fbxb-72 | 0.453599914 | 0.757194664 | -0.437994641 | 0.046 | 0.114 | 0.024741181 | 1 |
| WBGene00021838 | Y54E10BL.3 | 0.207472576 | 0.510733679 | -0.43751329 | 0.023 | 0.068 | 0.05701045 | 1 |
| WBGene00010960 | atp-6 | 2.85774515 | 3.160999163 | -0.437503061 | 0.592 | 0.694 | 0.023513357 | 1 |
| WBGene00000137 | amx-1 | 0.144049022 | 0.445517352 | -0.434926864 | 0.015 | 0.062 | 0.035660705 | 1 |
| WBGene00009174 | F26H9.2 | 0.071158826 | 0.37250754 | -0.434754296 | 0.008 | 0.06 | 0.014741981 | 1 |
| WBGene00022235 | sqd-1 | 0.932058556 | 1.233405764 | -0.434752123 | 0.131 | 0.193 | 0.158093186 | 1 |
| WBGene00019829 | R02E12.5 | 0.806671069 | 1.107657592 | -0.434231765 | 0.092 | 0.168 | 0.056654194 | 1 |
| WBGene00017894 | F28B12.1 | 1.6825213 | 1.983002377 | -0.433502559 | 0.292 | 0.493 | 0.001117995 | 1 |
| WBGene00006975 | zfp-1 | 0.6015555 | 0.9020365 | -0.433502449 | 0.077 | 0.159 | 0.030207589 | 1 |
| WBGene00044068 | syd-9 | 1.10955232 | 1.40999565 | -0.433448103 | 0.131 | 0.273 | 0.003981382 | 1 |
| WBGene00011575 | madf-4 | 0.070506324 | 0.370775554 | -0.433196929 | 0.008 | 0.052 | 0.026574873 | 1 |
| WBGene00006601 | tpi-1 | 0.273721676 | 0.572556757 | -0.431127889 | 0.031 | 0.085 | 0.040709515 | 1 |
| WBGene00015803 | C15H9.9 | 0.077939251 | 0.376574743 | -0.430839944 | 0.008 | 0.058 | 0.017114027 | 1 |
| WBGene00000182 | arf-1.2 | 0.408514015 | 0.706584155 | -0.430024312 | 0.046 | 0.114 | 0.027442132 | 1 |
| WBGene00004497 | rps-28 | 2.726455933 | 3.024173416 | -0.429515536 | 0.692 | 0.803 | 0.006433598 | 1 |
| WBGene00004438 | rpl-25.1 | 1.149641455 | 1.447125325 | -0.429178504 | 0.154 | 0.311 | 0.002199718 | 1 |
| WBGene00012885 | iscu-1 | 0.456855598 | 0.753648492 | -0.428181637 | 0.046 | 0.114 | 0.028811796 | 1 |
| WBGene00015063 | B0228.6 | 0.731230926 | 1.027652462 | -0.427645879 | 0.085 | 0.139 | 0.14013685 | 1 |
| WBGene00000553 | cmk-1 | 0.455781334 | 0.75201764 | -0.42737865 | 0.054 | 0.11 | 0.070266629 | 1 |
| WBGene00013242 | Y56A3A.30 | 0.201008937 | 0.497096035 | -0.427163389 | 0.023 | 0.07 | 0.050256162 | 1 |
| WBGene00010013 | F54B3.1 | 0.089249349 | 0.385273088 | -0.427071981 | 0.008 | 0.052 | 0.026881125 | 1 |
| WBGene00020397 | pcbd-1 | 0.468874093 | 0.764646557 | -0.426709468 | 0.054 | 0.103 | 0.107145618 | 1 |
| WBGene00003905 | pad-1 | 0.180545676 | 0.476168759 | -0.426493956 | 0.015 | 0.066 | 0.027476149 | 1 |
| WBGene00004375 | rme-4 | 0.316701168 | 0.612079605 | -0.426141006 | 0.023 | 0.085 | 0.017602505 | 1 |
| WBGene00012372 | W09H1.1 | 1.246662226 | 1.541905698 | -0.425946294 | 0.162 | 0.308 | 0.003811877 | 1 |
| WBGene00003079 | lsm-5 | 0.206308105 | 0.501464345 | -0.425820444 | 0.015 | 0.07 | 0.020730335 | 1 |
| WBGene00017900 | F28E10.1 | 0.404189179 | 0.698943661 | -0.425240829 | 0.038 | 0.103 | 0.028902737 | 1 |
| WBGene00012615 | dct-16 | 0.497248048 | 0.791447167 | -0.42443961 | 0.046 | 0.06 | 0.574424853 | 1 |
| WBGene00012100 | sipa-1 | 0.649783208 | 0.94349243 | -0.423732838 | 0.077 | 0.133 | 0.115861881 | 1 |
| WBGene00009142 | F26A3.4 | 1.246660762 | 1.540109735 | -0.423357377 | 0.185 | 0.325 | 0.010757391 | 1 |
| WBGene00016173 | fust-1 | 0.835323389 | 1.127478367 | -0.421490538 | 0.085 | 0.199 | 0.005492457 | 1 |
| WBGene00007416 | ceh-57 | 1.535921325 | 1.827762455 | -0.421037752 | 0.254 | 0.418 | 0.013858355 | 1 |
| WBGene00000915 | hsp-90 | 2.469872658 | 2.761649034 | -0.420944331 | 0.531 | 0.716 | 0.004346485 | 1 |
| WBGene00000898 | daf-2 | 0.300372802 | 0.591652804 | -0.420228215 | 0.031 | 0.075 | 0.073234298 | 1 |
| WBGene00077491 | K08H10.10 | 2.031288801 | 2.322005411 | -0.419415412 | 0.4 | 0.557 | 0.010904137 | 1 |
| WBGene00004053 | tank-1 | 0.334990108 | 0.625697048 | -0.419401459 | 0.038 | 0.064 | 0.276667325 | 1 |
| WBGene00010838 | M03C11.1 | 0.249714085 | 0.539158134 | -0.417579495 | 0.023 | 0.072 | 0.044818637 | 1 |
| WBGene00017826 | tag-340 | 0.084260344 | 0.373571632 | -0.417387961 | 0.008 | 0.05 | 0.030974676 | 1 |
| WBGene00003149 | mbk-1 | 0.835090027 | 1.124191446 | -0.417085182 | 0.115 | 0.203 | 0.043938541 | 1 |
| WBGene00004414 | rpl-3 | 2.310181544 | 2.598823978 | -0.416423008 | 0.469 | 0.671 | 0.000865062 | 1 |
| WBGene00006772 | unc-36 | 0.405990492 | 0.694622136 | -0.416407441 | 0.046 | 0.077 | 0.240363606 | 1 |
| WBGene00021928 | Y55F3AM.11 | 0.728380919 | 1.01700464 | -0.416396011 | 0.1 | 0.174 | 0.057511391 | 1 |
| WBGene00013307 | Y57G11C.9 | 1.506922324 | 1.795480016 | -0.416300751 | 0.254 | 0.383 | 0.052775576 | 1 |
| WBGene00004951 | spc-1 | 0.711618091 | 0.999661382 | -0.415558627 | 0.077 | 0.137 | 0.094122053 | 1 |
| WBGene00006728 | ubq-2 | 2.65359212 | 2.941602588 | -0.415511274 | 0.638 | 0.772 | 0.00274782 | 1 |
| WBGene00003668 | nhr-78 | 0.073121292 | 0.361105812 | -0.415473839 | 0.008 | 0.054 | 0.023242517 | 1 |
| WBGene00022170 | cox-6B | 0.681014676 | 0.968527788 | -0.414793742 | 0.092 | 0.17 | 0.042888664 | 1 |
| WBGene00020190 | guk-1 | 0.333240307 | 0.620674522 | -0.414679917 | 0.031 | 0.068 | 0.125650368 | 1 |
| WBGene00016975 | nhr-173 | 0.761955096 | 1.04843476 | -0.41330279 | 0.085 | 0.13 | 0.209009271 | 1 |
| WBGene00000380 | cct-5 | 1.394580717 | 1.680815227 | -0.412949107 | 0.185 | 0.36 | 0.001296079 | 1 |
| WBGene00000101 | aka-1 | 0.244649635 | 0.530601618 | -0.412541507 | 0.023 | 0.068 | 0.057178666 | 1 |
| WBGene00001510 | fzr-1 | 0.587710274 | 0.873630578 | -0.412495805 | 0.069 | 0.143 | 0.033600037 | 1 |
| WBGene00001668 | gpa-6 | 0.706139077 | 0.991555798 | -0.411769287 | 0.069 | 0.141 | 0.043831209 | 1 |
| WBGene00018782 | cct-3 | 0.233962126 | 0.517939272 | -0.409692421 | 0.023 | 0.052 | 0.168204356 | 1 |
| WBGene00006791 | unc-57 | 0.516373592 | 0.800240845 | -0.409533878 | 0.054 | 0.124 | 0.033191325 | 1 |
| WBGene00004185 | pro-1 | 0.469815058 | 0.753638668 | -0.409470913 | 0.038 | 0.103 | 0.028439745 | 1 |
| WBGene00017714 | F22F1.2 | 0.159115421 | 0.442158495 | -0.408344839 | 0.008 | 0.056 | 0.020097975 | 1 |

| | | | | | | | | |
|---|---|---|---|---|---|---|---|---|
| WBGene00013897 | ZC443.2 | 0.554955869 | 0.837798211 | -0.408055245 | 0.054 | 0.124 | 0.031664088 | 1 |
| WBGene00003947 | pbs-1 | 0.255422835 | 0.538164848 | -0.407910501 | 0.031 | 0.062 | 0.183100815 | 1 |
| WBGene00000284 | cah-6 | 0.287571818 | 0.570089764 | -0.407587239 | 0.031 | 0.068 | 0.128184198 | 1 |
| WBGene00000396 | cdh-4 | 0.359167167 | 0.641027533 | -0.406638552 | 0.046 | 0.101 | 0.064762605 | 1 |
| WBGene00020777 | T24E12.10 | 1.439113294 | 1.720689568 | -0.406228694 | 0.215 | 0.325 | 0.061218988 | 1 |
| WBGene00010898 | nphp-1 | 0.470396271 | 0.751665864 | -0.405786247 | 0.062 | 0.11 | 0.113349571 | 1 |
| WBGene00019947 | htz-1 | 0.450730542 | 0.731589665 | -0.405194064 | 0.054 | 0.108 | 0.07462948 | 1 |
| WBGene00015901 | nhr-158 | 1.794096504 | 2.073973441 | -0.403777068 | 0.277 | 0.484 | 0.00191226 | 1 |
| WBGene00011520 | nhr-213 | 0.466445048 | 0.745526193 | -0.402628984 | 0.046 | 0.11 | 0.037977723 | 1 |
| WBGene00019242 | srbc-40 | 1.066572465 | 1.345372719 | -0.402223745 | 0.108 | 0.232 | 0.005088471 | 1 |
| WBGene00009098 | ehbp-1 | 0.140649631 | 0.418718309 | -0.401168302 | 0.015 | 0.064 | 0.031251321 | 1 |
| WBGene00009374 | F34D10.4 | 1.402338969 | 1.679924327 | -0.40047102 | 0.177 | 0.275 | 0.039274575 | 1 |
| WBGene00021047 | zfp-3 | 0.597894421 | 0.875255689 | -0.400147726 | 0.077 | 0.141 | 0.075936752 | 1 |
| WBGene00016100 | C25E10.11 | 1.285201332 | 1.562542447 | -0.400118651 | 0.192 | 0.29 | 0.084507441 | 1 |
| WBGene00003920 | par-5 | 2.166501502 | 2.443366506 | -0.399431769 | 0.454 | 0.648 | 0.004066985 | 1 |
| WBGene00001085 | dpy-26 | 0.265359553 | 0.541400189 | -0.398242457 | 0.023 | 0.077 | 0.029363225 | 1 |
| WBGene00007027 | ssl-1 | 0.254087587 | 0.530100713 | -0.398202768 | 0.023 | 0.052 | 0.165551402 | 1 |
| WBGene00004474 | rps-5 | 2.517000825 | 2.792940628 | -0.398096986 | 0.546 | 0.749 | 0.001660382 | 1 |
| WBGene00016315 | C32D5.7 | 1.621737148 | 1.897498509 | -0.397839548 | 0.285 | 0.443 | 0.011877109 | 1 |
| WBGene00015251 | B0546.4 | 0.640091652 | 0.915379421 | -0.397156298 | 0.069 | 0.126 | 0.092429967 | 1 |
| WBGene00009120 | F25H2.6 | 1.414417987 | 1.689644121 | -0.39706738 | 0.231 | 0.408 | 0.003963501 | 1 |
| WBGene00008398 | D2005.3 | 0.239040547 | 0.514262641 | -0.397061551 | 0.023 | 0.079 | 0.026437272 | 1 |
| WBGene00002202 | kin-19 | 0.798996651 | 1.073797567 | -0.396453917 | 0.1 | 0.153 | 0.175459373 | 1 |
| WBGene00001675 | gpa-13 | 0.320118671 | 0.594884939 | -0.396403931 | 0.038 | 0.072 | 0.182934871 | 1 |
| WBGene00003081 | lsm-7 | 0.041197077 | 0.315435536 | -0.395642466 | 0.008 | 0.05 | 0.02902452 | 1 |
| WBGene00004495 | rps-26 | 2.986396877 | 3.259376199 | -0.393825914 | 0.785 | 0.843 | 0.000950168 | 1 |
| WBGene00013317 | Y57G11C.22 | 0.255894076 | 0.528839075 | -0.393776397 | 0.023 | 0.07 | 0.051539471 | 1 |
| WBGene00017579 | F18G5.1 | 0.45454429 | 0.727234421 | -0.393408698 | 0.054 | 0.112 | 0.060706438 | 1 |
| WBGene00008168 | C48B4.3 | 1.192076958 | 1.464271059 | -0.392693081 | 0.177 | 0.296 | 0.023832954 | 1 |
| WBGene00018067 | npr-7 | 0.406149956 | 0.678062443 | -0.392286796 | 0.023 | 0.101 | 0.005872403 | 1 |
| WBGene00004413 | rpl-2 | 2.487370388 | 2.757575648 | -0.389823789 | 0.592 | 0.737 | 0.014994802 | 1 |
| WBGene00020208 | T04C4.1 | 0.700645981 | 0.970791315 | -0.389737333 | 0.077 | 0.143 | 0.063786626 | 1 |
| WBGene00003695 | nhr-105 | 2.133145973 | 2.402921047 | -0.389203163 | 0.438 | 0.594 | 0.009188194 | 1 |
| WBGene00021522 | nhr-274 | 0.65738502 | 0.926681259 | -0.38851235 | 0.069 | 0.155 | 0.019796469 | 1 |
| WBGene00010971 | R01E6.7 | 0.398542154 | 0.667685545 | -0.388291836 | 0.038 | 0.093 | 0.053006206 | 1 |
| WBGene00009688 | F44E5.1 | 0.58215414 | 0.851113212 | -0.38802592 | 0.069 | 0.106 | 0.272262744 | 1 |
| WBGene00018572 | lin-42 | 1.881734662 | 2.150621106 | -0.387921139 | 0.292 | 0.414 | 0.058681438 | 1 |
| WBGene00015387 | kcnl-3 | 0.232500883 | 0.501128912 | -0.387548326 | 0.023 | 0.064 | 0.07585753 | 1 |
| WBGene00007554 | pptr-2 | 0.154033613 | 0.422543916 | -0.387378482 | 0.015 | 0.058 | 0.047972718 | 1 |
| WBGene00012337 | scyl-1 | 0.527887258 | 0.796371081 | -0.387340279 | 0.046 | 0.112 | 0.033645199 | 1 |
| WBGene00021843 | Y54E10BR.4 | 0.101506946 | 0.36894378 | -0.385829793 | 0.008 | 0.052 | 0.026983902 | 1 |
| WBGene00015423 | C04E6.7 | 1.231466709 | 1.498787853 | -0.385662889 | 0.177 | 0.3 | 0.017252387 | 1 |
| WBGene00044732 | T03G11.10 | 0.298589 | 0.564886601 | -0.384186228 | 0.031 | 0.056 | 0.258596671 | 1 |
| WBGene00001664 | gpa-2 | 1.508643852 | 1.774138032 | -0.383027137 | 0.208 | 0.42 | 0.000291585 | 1 |
| WBGene00004893 | sms-2 | 0.210482232 | 0.475856159 | -0.382853649 | 0.023 | 0.06 | 0.102589376 | 1 |
| WBGene00000378 | cct-2 | 0.351017855 | 0.616128868 | -0.382474344 | 0.031 | 0.081 | 0.051607902 | 1 |
| WBGene00002981 | lgg-2 | 1.89796474 | 2.162775893 | -0.382041737 | 0.346 | 0.534 | 0.003925365 | 1 |
| WBGene00020444 | T12B3.1 | 0.152529572 | 0.416909313 | -0.381419341 | 0.015 | 0.068 | 0.023747632 | 1 |
| WBGene00013577 | Y76A2B.5 | 0.721511364 | 0.985832982 | -0.381335487 | 0.092 | 0.168 | 0.044800139 | 1 |
| WBGene00001426 | fkb-1 | 0.336561585 | 0.600007285 | -0.380071805 | 0.038 | 0.06 | 0.366447217 | 1 |
| WBGene00004448 | rpl-34 | 2.089959242 | 2.35337782 | -0.380032676 | 0.446 | 0.6 | 0.012892735 | 1 |
| WBGene00017711 | F22E5.13 | 0.47209105 | 0.735470548 | -0.379976296 | 0.054 | 0.106 | 0.086168549 | 1 |
| WBGene00007184 | ctr-9 | 0.318301223 | 0.581376148 | -0.37953689 | 0.031 | 0.074 | 0.086974668 | 1 |
| WBGene00004747 | sdc-3 | 0.235871672 | 0.498306278 | -0.378613105 | 0.023 | 0.068 | 0.059403287 | 1 |
| WBGene00001665 | gpa-3 | 1.359450737 | 1.621756199 | -0.37842679 | 0.192 | 0.358 | 0.003297539 | 1 |
| WBGene00003150 | mbk-2 | 1.428389784 | 1.690015415 | -0.377446001 | 0.246 | 0.377 | 0.025663591 | 1 |
| WBGene00003724 | nhr-134 | 0.58987227 | 0.851433975 | -0.377353774 | 0.069 | 0.12 | 0.116614171 | 1 |
| WBGene00006910 | vha-1 | 0.978218023 | 1.239254829 | -0.376596507 | 0.131 | 0.211 | 0.10199383 | 1 |
| WBGene00004420 | rpl-9 | 2.214567302 | 2.475576633 | -0.376556868 | 0.469 | 0.671 | 0.00594224 | 1 |
| WBGene00002190 | kin-2 | 1.111566738 | 1.372504217 | -0.376453208 | 0.162 | 0.251 | 0.073540336 | 1 |
| WBGene00006608 | tre-2 | 0.333372091 | 0.594173711 | -0.376257204 | 0.031 | 0.087 | 0.035599758 | 1 |
| WBGene00001672 | gpa-10 | 0.600877604 | 0.861550132 | -0.376070962 | 0.062 | 0.151 | 0.011413518 | 1 |
| WBGene00007621 | C16C2.4 | 0.082452944 | 0.342727402 | -0.37549667 | 0.008 | 0.05 | 0.031211322 | 1 |
| WBGene00014938 | mdt-9 | 0.338174862 | 0.597812325 | -0.37457768 | 0.031 | 0.05 | 0.3583968 | 1 |

| | | | | | | | | |
|---|---|---|---|---|---|---|---|---|
| WBGene00044386 | C27A2.7 | 0.220463168 | 0.479452391 | -0.373642468 | 0.023 | 0.066 | 0.066268414 | 1 |
| WBGene00015507 | C06A6.2 | 0.322384955 | 0.580513607 | -0.372400927 | 0.038 | 0.097 | 0.039840377 | 1 |
| WBGene00003003 | lin-14 | 0.424072057 | 0.681984787 | -0.372089416 | 0.046 | 0.101 | 0.064150755 | 1 |
| WBGene00007235 | ahsa-1 | 0.501942516 | 0.759232258 | -0.371190634 | 0.046 | 0.11 | 0.039321293 | 1 |
| WBGene00022579 | ZC262.2 | 0.31276359 | 0.569894573 | -0.370961595 | 0.031 | 0.091 | 0.025491182 | 1 |
| WBGene00004831 | slo-2 | 0.42726308 | 0.684216385 | -0.370705259 | 0.038 | 0.114 | 0.013518102 | 1 |
| WBGene00012766 | Y41E3.8 | 0.298990524 | 0.55553863 | -0.37012068 | 0.031 | 0.085 | 0.041974889 | 1 |
| WBGene00016790 | C49H3.3 | 0.87107522 | 1.127510573 | -0.369958011 | 0.1 | 0.186 | 0.030946431 | 1 |
| WBGene00011980 | T24B8.7 | 0.326427339 | 0.582814541 | -0.369888544 | 0.038 | 0.066 | 0.249471711 | 1 |
| WBGene00004915 | snr-2 | 0.469951244 | 0.725811134 | -0.369127795 | 0.054 | 0.12 | 0.040684898 | 1 |
| WBGene00017557 | nep-11 | 0.526956605 | 0.782495566 | -0.368664792 | 0.046 | 0.122 | 0.018046601 | 1 |
| WBGene00016794 | C49H3.9 | 0.073121292 | 0.328402877 | -0.368293477 | 0.008 | 0.052 | 0.026881125 | 1 |
| WBGene00044927 | F32B5.9 | 0.466305527 | 0.721250891 | -0.367808413 | 0.054 | 0.087 | 0.230682121 | 1 |
| WBGene00050944 | C14A11.9 | 0.513138485 | 0.76766882 | -0.367209651 | 0.054 | 0.128 | 0.025470444 | 1 |
| WBGene00015745 | C13F10.6 | 0.306034987 | 0.56035964 | -0.366912916 | 0.031 | 0.064 | 0.16457608 | 1 |
| WBGene00001840 | hel-1 | 1.076975617 | 1.331154311 | -0.366702341 | 0.131 | 0.211 | 0.095785625 | 1 |
| WBGene00005111 | srd-33 | 1.187570454 | 1.441613229 | -0.366506252 | 0.146 | 0.259 | 0.017934398 | 1 |
| WBGene00016119 | C25H3.10 | 2.497717495 | 2.751615626 | -0.366297575 | 0.592 | 0.677 | 0.028342047 | 1 |
| WBGene00010500 | K02B12.7 | 0.190145622 | 0.443859263 | -0.366031413 | 0.023 | 0.072 | 0.044684735 | 1 |
| WBGene00008256 | glb-12 | 0.267430091 | 0.520649813 | -0.365318836 | 0.015 | 0.075 | 0.013536507 | 1 |
| WBGene00008366 | tpra-1 | 0.666027773 | 0.919030182 | -0.365005321 | 0.069 | 0.149 | 0.025602925 | 1 |
| WBGene00003678 | nhr-88 | 0.739059523 | 0.991709163 | -0.364496382 | 0.092 | 0.159 | 0.08942819 | 1 |
| WBGene00019364 | mksr-1 | 0.603098569 | 0.855708723 | -0.364439417 | 0.077 | 0.095 | 0.632464434 | 1 |
| WBGene00016559 | C41A3.2 | 1.102808158 | 1.355167373 | -0.364077387 | 0.146 | 0.273 | 0.012861786 | 1 |
| WBGene00000377 | cct-1 | 0.309515557 | 0.561228209 | -0.363144594 | 0.031 | 0.079 | 0.060917297 | 1 |
| WBGene00009126 | pyk-1 | 0.672614699 | 0.923620976 | -0.362125512 | 0.085 | 0.153 | 0.065055372 | 1 |
| WBGene00018213 | F39H12.2 | 1.466723183 | 1.71725845 | -0.361445987 | 0.223 | 0.406 | 0.004094138 | 1 |
| WBGene00009262 | F30A10.3 | 0.48296109 | 0.733146327 | -0.360941 | 0.046 | 0.095 | 0.091192698 | 1 |
| WBGene00012641 | epg-6 | 0.344136107 | 0.594221634 | -0.36079715 | 0.031 | 0.095 | 0.021845904 | 1 |
| WBGene00021890 | Y54G2A.26 | 0.710361478 | 0.95958696 | -0.359556367 | 0.1 | 0.178 | 0.049861329 | 1 |
| WBGene00013962 | ZK287.1 | 0.488580481 | 0.737233176 | -0.35873001 | 0.038 | 0.116 | 0.012261542 | 1 |
| WBGene00004504 | rpt-4 | 0.489077796 | 0.737652665 | -0.358617731 | 0.062 | 0.114 | 0.102892212 | 1 |
| WBGene00010639 | K07F5.15 | 0.192264237 | 0.440761515 | -0.35850579 | 0.015 | 0.068 | 0.023990905 | 1 |
| WBGene00003924 | pas-3 | 0.391690035 | 0.640101267 | -0.358381653 | 0.038 | 0.093 | 0.05355044 | 1 |
| WBGene00015716 | sox-4 | 0.299160975 | 0.547458174 | -0.358217137 | 0.031 | 0.066 | 0.140753445 | 1 |
| WBGene00006920 | vha-11 | 0.165956736 | 0.413698002 | -0.357415096 | 0.015 | 0.058 | 0.048289073 | 1 |
| WBGene00020322 | T07F12.2 | 0.267605502 | 0.51518043 | -0.357175121 | 0.023 | 0.074 | 0.039601514 | 1 |
| WBGene00022271 | cpx-1 | 1.142781274 | 1.390179403 | -0.356920053 | 0.154 | 0.28 | 0.020807198 | 1 |
| WBGene00016448 | C35D10.12 | 0.519863235 | 0.767109537 | -0.356701014 | 0.046 | 0.112 | 0.033645199 | 1 |
| WBGene00009547 | F38H4.3 | 0.327361485 | 0.574464301 | -0.356494007 | 0.038 | 0.085 | 0.081179655 | 1 |
| WBGene00015798 | C15H9.2 | 0.258169507 | 0.505120469 | -0.356274928 | 0.023 | 0.074 | 0.039661242 | 1 |
| WBGene00022062 | Y67D8B.2 | 0.332510992 | 0.579404107 | -0.356191473 | 0.038 | 0.091 | 0.057060652 | 1 |
| WBGene00012194 | toe-4 | 0.760317166 | 1.006930239 | -0.355787458 | 0.092 | 0.188 | 0.02145575 | 1 |
| WBGene00016509 | adss-1 | 0.22373629 | 0.470284581 | -0.355693997 | 0.023 | 0.066 | 0.067432994 | 1 |
| WBGene00004921 | snt-1 | 0.770444088 | 1.016961509 | -0.355649461 | 0.077 | 0.155 | 0.039029833 | 1 |
| WBGene00004423 | rpl-11.2 | 2.661565372 | 2.908007664 | -0.355541073 | 0.692 | 0.776 | 0.004870418 | 1 |
| WBGene00018606 | F48E3.6 | 1.148502926 | 1.394129013 | -0.354363537 | 0.185 | 0.306 | 0.030566659 | 1 |
| WBGene00006807 | unc-75 | 0.233121913 | 0.478210167 | -0.353587609 | 0.023 | 0.07 | 0.052616777 | 1 |
| WBGene00006068 | sto-6 | 0.42290199 | 0.667830669 | -0.35335739 | 0.046 | 0.11 | 0.034006444 | 1 |
| WBGene00021427 | sec-61.B | 0.295498337 | 0.54031127 | -0.353190404 | 0.031 | 0.083 | 0.048078507 | 1 |
| WBGene00004143 | pqn-59 | 0.612680667 | 0.856943009 | -0.35239607 | 0.054 | 0.139 | 0.012862015 | 1 |
| WBGene00006741 | unc-1 | 0.319324228 | 0.563507501 | -0.352281997 | 0.031 | 0.068 | 0.124086692 | 1 |
| WBGene00016258 | vha-16 | 0.635501996 | 0.879413931 | -0.351890538 | 0.062 | 0.139 | 0.024363414 | 1 |
| WBGene00003015 | lin-29 | 0.590771325 | 0.834577636 | -0.351738155 | 0.046 | 0.126 | 0.012752519 | 1 |
| WBGene00004471 | rps-2 | 2.077157024 | 2.320649388 | -0.351285225 | 0.415 | 0.582 | 0.01842109 | 1 |
| WBGene00011729 | set-16 | 0.209150319 | 0.452236165 | -0.350698744 | 0.015 | 0.05 | 0.085627663 | 1 |
| WBGene00017671 | pgal-1 | 2.201180342 | 2.44399777 | -0.350311499 | 0.431 | 0.603 | 0.021614529 | 1 |
| WBGene00003000 | lin-11 | 1.004466956 | 1.246910421 | -0.349771984 | 0.131 | 0.234 | 0.027686757 | 1 |
| WBGene00016002 | gyf-1 | 1.275425135 | 1.517864046 | -0.349765415 | 0.2 | 0.317 | 0.047048964 | 1 |
| WBGene00004432 | rpl-20 | 2.361779561 | 2.60396994 | -0.349406858 | 0.515 | 0.691 | 0.018513645 | 1 |
| WBGene00018373 | F43B10.1 | 0.206185461 | 0.447224912 | -0.347746421 | 0.015 | 0.054 | 0.065070689 | 1 |
| WBGene00003009 | lin-23 | 0.182430665 | 0.423253678 | -0.347434167 | 0.015 | 0.054 | 0.064444196 | 1 |
| WBGene00001448 | flp-5 | 0.343480055 | 0.584180504 | -0.347257344 | 0.031 | 0.056 | 0.247169657 | 1 |
| WBGene00006918 | vha-9 | 0.511789047 | 0.751792313 | -0.346251522 | 0.054 | 0.141 | 0.011333259 | 1 |

| | | | | | | | | |
|---|---|---|---|---|---|---|---|---|
| WBGene00017085 | E01A2.2 | 0.397869755 | 0.63763955 | -0.345914695 | 0.023 | 0.05 | 0.189562706 | 1 |
| WBGene00022240 | shl-1 | 0.520973611 | 0.759942981 | -0.344759925 | 0.054 | 0.112 | 0.061540755 | 1 |
| WBGene00004475 | rps-6 | 2.063795465 | 2.302420639 | -0.344263355 | 0.4 | 0.561 | 0.033396872 | 1 |
| WBGene00016981 | rpn-13 | 0.399868133 | 0.638477149 | -0.344240044 | 0.038 | 0.097 | 0.042201734 | 1 |
| WBGene00003916 | par-1 | 0.32657486 | 0.564102525 | -0.342679985 | 0.038 | 0.085 | 0.079595775 | 1 |
| WBGene00004346 | rgs-3 | 1.980722199 | 2.218083068 | -0.34243935 | 0.362 | 0.544 | 0.017649564 | 1 |
| WBGene00002717 | let-526 | 0.617000202 | 0.854173345 | -0.342168518 | 0.069 | 0.145 | 0.035498849 | 1 |
| WBGene00044544 | F26F12.8 | 0.626616117 | 0.863505353 | -0.341758926 | 0.062 | 0.124 | 0.056269457 | 1 |
| WBGene00044354 | ZK418.11 | 0.533809953 | 0.770686125 | -0.341740078 | 0.046 | 0.116 | 0.02326217 | 1 |
| WBGene00001502 | ftt-2 | 1.888991586 | 2.125476089 | -0.34117502 | 0.354 | 0.485 | 0.059553756 | 1 |
| WBGene00006065 | sto-3 | 0.135286335 | 0.37162671 | -0.340967087 | 0.008 | 0.052 | 0.027190446 | 1 |
| WBGene00004884 | smg-6 | 0.406973494 | 0.64314868 | -0.34072877 | 0.038 | 0.06 | 0.369963646 | 1 |
| WBGene00001994 | hpk-1 | 1.150198551 | 1.386043801 | -0.340252772 | 0.123 | 0.267 | 0.002560967 | 1 |
| WBGene00003925 | pas-4 | 0.243828665 | 0.479619871 | -0.340174804 | 0.023 | 0.066 | 0.064741507 | 1 |
| WBGene00023067 | rpl-41.1 | 0.293720326 | 0.528905837 | -0.339300972 | 0.031 | 0.05 | 0.345724094 | 1 |
| WBGene00004463 | rpn-7 | 0.307883137 | 0.543017749 | -0.339227539 | 0.031 | 0.083 | 0.047422559 | 1 |
| WBGene00004274 | rab-11.1 | 1.061457257 | 1.296080537 | -0.338489842 | 0.154 | 0.248 | 0.066330239 | 1 |
| WBGene00021346 | Y37E3.1 | 0.26680504 | 0.500844676 | -0.337647822 | 0.023 | 0.066 | 0.066268432 | 1 |
| WBGene00006805 | unc-73 | 0.313830058 | 0.54780731 | -0.337557821 | 0.031 | 0.056 | 0.252267257 | 1 |
| WBGene00021304 | nphp-2 | 0.595266373 | 0.829240753 | -0.337553678 | 0.046 | 0.126 | 0.013944816 | 1 |
| WBGene00015568 | sre-3 | 0.818202631 | 1.051267058 | -0.336240892 | 0.1 | 0.188 | 0.030424242 | 1 |
| WBGene00020951 | sna-1 | 0.27688863 | 0.509393664 | -0.33543386 | 0.031 | 0.072 | 0.100960641 | 1 |
| WBGene00006519 | cox-6A | 0.556915674 | 0.789210409 | -0.335130462 | 0.062 | 0.132 | 0.038314359 | 1 |
| WBGene00019188 | H11E01.3 | 0.38493318 | 0.616382845 | -0.333911284 | 0.031 | 0.075 | 0.076005664 | 1 |
| WBGene00002363 | let-92 | 0.780047615 | 1.011415251 | -0.333792941 | 0.108 | 0.151 | 0.290268706 | 1 |
| WBGene00003926 | pas-5 | 0.362224238 | 0.593158778 | -0.333168115 | 0.038 | 0.081 | 0.103673374 | 1 |
| WBGene00004268 | rab-5 | 0.690558138 | 0.921463406 | -0.333125885 | 0.077 | 0.151 | 0.046059215 | 1 |
| WBGene00004424 | rpl-12 | 3.49352317 | 3.724322193 | -0.332972605 | 0.862 | 0.878 | 0.000343248 | 1 |
| WBGene00003045 | lir-2 | 0.154586067 | 0.385261733 | -0.332794639 | 0.015 | 0.06 | 0.042338135 | 1 |
| WBGene00016650 | ubr-4 | 0.820684239 | 1.050868183 | -0.332085234 | 0.108 | 0.145 | 0.340399421 | 1 |
| WBGene00001669 | gpa-7 | 0.423299968 | 0.653259724 | -0.331761799 | 0.046 | 0.104 | 0.052156792 | 1 |
| WBGene00002083 | inf-1 | 0.344783785 | 0.574627841 | -0.331594879 | 0.038 | 0.079 | 0.115061421 | 1 |
| WBGene00012768 | eef-1B.2 | 0.945961161 | 1.175130134 | -0.33062094 | 0.108 | 0.205 | 0.030544916 | 1 |
| WBGene00005640 | srn-1 | 1.772596869 | 2.001041361 | -0.329575736 | 0.285 | 0.462 | 0.007765151 | 1 |
| WBGene00016138 | flh-2 | 0.635288015 | 0.863291859 | -0.328940015 | 0.077 | 0.132 | 0.123709135 | 1 |
| WBGene00001680 | gpb-2 | 0.227708164 | 0.455434843 | -0.328540151 | 0.023 | 0.056 | 0.13222473 | 1 |
| WBGene00010730 | ensa-1 | 0.885214026 | 1.112624119 | -0.328083414 | 0.115 | 0.195 | 0.070223698 | 1 |
| WBGene00019286 | K01A12.2 | 1.105729428 | 1.332608133 | -0.327316782 | 0.146 | 0.255 | 0.031117283 | 1 |
| WBGene00000403 | casy-1 | 2.296187333 | 2.52291283 | -0.32709575 | 0.492 | 0.656 | 0.019711139 | 1 |
| WBGene00000793 | crh-1 | 1.514422571 | 1.740936322 | -0.326790265 | 0.246 | 0.389 | 0.036372019 | 1 |
| WBGene00012203 | rga-1 | 0.24477853 | 0.47037244 | -0.325463216 | 0.031 | 0.068 | 0.120852329 | 1 |
| WBGene00006715 | ubc-20 | 0.332131646 | 0.557305584 | -0.324857324 | 0.031 | 0.083 | 0.0467742 | 1 |
| WBGene00017568 | F18E9.1 | 0.330230371 | 0.554344894 | -0.323328911 | 0.023 | 0.091 | 0.011901129 | 1 |
| WBGene00012621 | Y38H6C.8 | 1.325081593 | 1.549048833 | -0.323116426 | 0.177 | 0.209 | 0.433225357 | 1 |
| WBGene00001955 | hlh-11 | 0.411465961 | 0.635364398 | -0.323017165 | 0.038 | 0.075 | 0.154943408 | 1 |
| WBGene00016817 | C50E3.6 | 0.38306892 | 0.606548505 | -0.32241289 | 0.031 | 0.072 | 0.098656598 | 1 |
| WBGene00000854 | cux-7 | 0.384577109 | 0.608032091 | -0.322377395 | 0.038 | 0.097 | 0.042422156 | 1 |
| WBGene00013735 | Y111B2A.12 | 0.445338238 | 0.668437411 | -0.321864071 | 0.038 | 0.077 | 0.139958195 | 1 |
| WBGene00219592 | linc-84 | 0.203857472 | 0.425509417 | -0.319776161 | 0.023 | 0.056 | 0.121454586 | 1 |
| WBGene00000095 | aha-1 | 0.286251164 | 0.507596771 | -0.319334209 | 0.031 | 0.074 | 0.087542836 | 1 |
| WBGene00012221 | W03C9.5 | 0.97837961 | 1.199708669 | -0.319310336 | 0.138 | 0.222 | 0.086012736 | 1 |
| WBGene00000908 | daf-12 | 0.285259581 | 0.505874161 | -0.318279561 | 0.031 | 0.06 | 0.203101613 | 1 |
| WBGene00018078 | F35H12.6 | 0.492635491 | 0.712938717 | -0.317830373 | 0.046 | 0.128 | 0.013066107 | 1 |
| WBGene00009353 | sdhd-1 | 0.318136708 | 0.53767861 | -0.316732014 | 0.031 | 0.06 | 0.203577323 | 1 |
| WBGene00003850 | odr-3 | 1.922202307 | 2.140939401 | -0.31557092 | 0.377 | 0.507 | 0.157035307 | 1 |
| WBGene00005743 | srv-32 | 0.822421826 | 1.039702486 | -0.31346973 | 0.085 | 0.155 | 0.061898162 | 1 |
| WBGene00012158 | ucr-2.1 | 0.153034606 | 0.370150996 | -0.313232739 | 0.015 | 0.05 | 0.085627663 | 1 |
| WBGene00007385 | atp-5 | 0.765106202 | 0.98174648 | -0.312545855 | 0.092 | 0.17 | 0.045797418 | 1 |
| WBGene00017240 | calu-2 | 0.449605604 | 0.665245046 | -0.311101953 | 0.046 | 0.099 | 0.076952021 | 1 |
| WBGene00004473 | rps-4 | 2.469209819 | 2.683795472 | -0.309581658 | 0.531 | 0.708 | 0.013025556 | 1 |
| WBGene00000262 | bra-1 | 0.441800472 | 0.656104303 | -0.309175074 | 0.038 | 0.091 | 0.06071479 | 1 |
| WBGene00017066 | maco-1 | 0.550494113 | 0.76439066 | -0.308587487 | 0.054 | 0.137 | 0.015352826 | 1 |
| WBGene00019285 | cbn-1 | 2.148853147 | 2.36205949 | -0.307591735 | 0.431 | 0.607 | 0.031509105 | 1 |
| WBGene00304822 | T14G8.6 | 0.303607248 | 0.516778339 | -0.307540875 | 0.031 | 0.081 | 0.054280269 | 1 |

| | | | | | | | | |
|---|---|---|---|---|---|---|---|---|
| WBGene00022404 | srsx-5 | 0.139920918 | 0.353077425 | -0.307519836 | 0.015 | 0.054 | 0.06423646 | 1 |
| WBGene00022794 | snu-23 | 0.15864321 | 0.371679411 | -0.30734627 | 0.015 | 0.05 | 0.085627663 | 1 |
| WBGene00022435 | cfap-36 | 0.363723452 | 0.576700383 | -0.307260761 | 0.038 | 0.074 | 0.169891933 | 1 |
| WBGene00000492 | che-13 | 0.375085339 | 0.587881102 | -0.306999392 | 0.038 | 0.091 | 0.059802427 | 1 |
| WBGene00012850 | Y44A6C.1 | 0.319713634 | 0.532463352 | -0.306932963 | 0.031 | 0.083 | 0.047815216 | 1 |
| WBGene00016748 | aexr-3 | 0.51454663 | 0.726966233 | -0.306456708 | 0.054 | 0.106 | 0.084488759 | 1 |
| WBGene00020562 | T19C3.4 | 0.32902276 | 0.541178412 | -0.306075908 | 0.031 | 0.095 | 0.022620056 | 1 |
| WBGene00077486 | sls-1.9 | 0.417063432 | 0.628425696 | -0.30493129 | 0.038 | 0.089 | 0.067415815 | 1 |
| WBGene00006415 | tag-30 | 0.216074442 | 0.427416359 | -0.304901936 | 0.023 | 0.068 | 0.059056383 | 1 |
| WBGene00000484 | che-2 | 0.238023042 | 0.449072702 | -0.304480298 | 0.023 | 0.054 | 0.148772231 | 1 |
| WBGene00012644 | rabx-5 | 0.203285094 | 0.413598454 | -0.303418041 | 0.015 | 0.064 | 0.032317027 | 1 |
| WBGene00007053 | chd-7 | 0.78195671 | 0.992228263 | -0.303357726 | 0.1 | 0.161 | 0.122450823 | 1 |
| WBGene00004873 | smc-3 | 0.173080046 | 0.382724408 | -0.302452881 | 0.015 | 0.054 | 0.06486131 | 1 |
| WBGene00014095 | gdh-1 | 0.203946072 | 0.413117068 | -0.301769959 | 0.023 | 0.05 | 0.188576412 | 1 |
| WBGene00004480 | rps-11 | 2.914403974 | 3.123025017 | -0.300976544 | 0.731 | 0.81 | 0.000859967 | 1 |
| WBGene00001042 | dnj-24 | 0.361264842 | 0.569456056 | -0.300356432 | 0.038 | 0.087 | 0.076687721 | 1 |
| WBGene00022776 | perm-3 | 0.211392354 | 0.419330025 | -0.299990646 | 0.023 | 0.052 | 0.165551402 | 1 |
| WBGene00009625 | F41E7.9 | 0.659087789 | 0.865925856 | -0.298404254 | 0.077 | 0.106 | 0.427533956 | 1 |
| WBGene00004483 | rps-14 | 3.166328178 | 3.372768338 | -0.297830195 | 0.792 | 0.855 | 0.000815837 | 1 |
| WBGene00012813 | Y43F8B.2 | 1.539619694 | 1.745720612 | -0.297340773 | 0.215 | 0.286 | 0.196244444 | 1 |
| WBGene00004447 | rpl-33 | 2.285403914 | 2.490678588 | -0.296148754 | 0.477 | 0.65 | 0.035044062 | 1 |
| WBGene00002005 | hsp-1 | 2.076201187 | 2.281409824 | -0.296053483 | 0.377 | 0.565 | 0.022245169 | 1 |
| WBGene00001031 | dnj-13 | 1.069197577 | 1.27269082 | -0.293578692 | 0.154 | 0.244 | 0.057224452 | 1 |
| WBGene00018872 | F55C12.5 | 0.796229781 | 0.998709484 | -0.292116463 | 0.092 | 0.141 | 0.206264338 | 1 |
| WBGene00018349 | F42C5.9 | 0.378226465 | 0.580416353 | -0.291698349 | 0.038 | 0.075 | 0.147072626 | 1 |
| WBGene00001169 | eef-1A.2 | 1.988890876 | 2.190894048 | -0.291428974 | 0.392 | 0.532 | 0.106824081 | 1 |
| WBGene00010239 | F58B4.6 | 2.326123746 | 2.527801957 | -0.290960156 | 0.462 | 0.65 | 0.017382473 | 1 |
| WBGene00000475 | cey-4 | 0.492400553 | 0.693944273 | -0.290766126 | 0.038 | 0.097 | 0.042754606 | 1 |
| WBGene00000140 | anc-1 | 0.682512798 | 0.883162599 | -0.289476473 | 0.077 | 0.114 | 0.270785769 | 1 |
| WBGene00007801 | trpa-1 | 0.284493867 | 0.484881651 | -0.289098463 | 0.023 | 0.068 | 0.059751828 | 1 |
| WBGene00018727 | jbts-14 | 0.461221421 | 0.661575921 | -0.289050444 | 0.054 | 0.103 | 0.107600998 | 1 |
| WBGene00012765 | Y41E3.7 | 1.422476251 | 1.622295382 | -0.28827807 | 0.231 | 0.366 | 0.046839899 | 1 |
| WBGene00003618 | nhr-19 | 0.421286315 | 0.618910658 | -0.285111659 | 0.038 | 0.087 | 0.077445861 | 1 |
| WBGene00010865 | M04D8.7 | 1.94117231 | 2.13806646 | -0.284058213 | 0.346 | 0.505 | 0.038369036 | 1 |
| WBGene00019180 | H10D18.5 | 1.203252361 | 1.399619096 | -0.283297314 | 0.169 | 0.286 | 0.030150041 | 1 |
| WBGene00007223 | C01F6.9 | 0.281512178 | 0.477720733 | -0.283069108 | 0.031 | 0.064 | 0.157177511 | 1 |
| WBGene00002179 | jph-1 | 0.381857522 | 0.576221617 | -0.280408115 | 0.038 | 0.075 | 0.151826257 | 1 |
| WBGene00019396 | K04F10.3 | 0.257249988 | 0.45130052 | -0.279955739 | 0.023 | 0.066 | 0.067237721 | 1 |
| WBGene00018160 | odr-8 | 0.594344999 | 0.787945546 | -0.279306549 | 0.062 | 0.133 | 0.036369574 | 1 |
| WBGene00012354 | cox-4 | 0.626859789 | 0.820197757 | -0.278927728 | 0.062 | 0.141 | 0.024729563 | 1 |
| WBGene00000229 | atp-2 | 1.029778003 | 1.223000498 | -0.278761135 | 0.123 | 0.23 | 0.021787514 | 1 |
| WBGene00003600 | nhr-1 | 0.335232655 | 0.527662256 | -0.277617231 | 0.031 | 0.083 | 0.047946704 | 1 |
| WBGene00001784 | gst-36 | 1.068088123 | 1.260169724 | -0.277115174 | 0.131 | 0.191 | 0.15448066 | 1 |
| WBGene00012894 | tbc-17 | 0.799354846 | 0.990413542 | -0.275639433 | 0.1 | 0.162 | 0.121120545 | 1 |
| WBGene00003952 | pbs-6 | 0.409176363 | 0.600049366 | -0.275371534 | 0.031 | 0.093 | 0.025040877 | 1 |
| WBGene00012896 | snrp-200 | 0.613844766 | 0.804634737 | -0.275251745 | 0.069 | 0.126 | 0.093168219 | 1 |
| WBGene00008957 | glb-13 | 1.151462922 | 1.342147313 | -0.275099425 | 0.146 | 0.263 | 0.024071366 | 1 |
| WBGene00005065 | sra-39 | 0.148236269 | 0.3388667 | -0.275021578 | 0.015 | 0.052 | 0.074652022 | 1 |
| WBGene00001851 | hif-1 | 0.830364057 | 1.020878733 | -0.274854578 | 0.115 | 0.174 | 0.168279613 | 1 |
| WBGene00004434 | rpl-22 | 2.390833107 | 2.580718798 | -0.273947145 | 0.554 | 0.685 | 0.101166249 | 1 |
| WBGene00016016 | C23G10.10 | 0.61579456 | 0.80561958 | -0.273859615 | 0.069 | 0.116 | 0.157708733 | 1 |
| WBGene00003563 | ncs-1 | 0.830295004 | 1.019990023 | -0.273672064 | 0.092 | 0.195 | 0.014917635 | 1 |
| WBGene00219300 | W10G6.6 | 0.683580689 | 0.87264918 | -0.272768175 | 0.077 | 0.149 | 0.049127663 | 1 |
| WBGene00010962 | ctc-3 | 2.447485016 | 2.635733842 | -0.271585648 | 0.485 | 0.574 | 0.42979559 | 1 |
| WBGene00004189 | pars-1 | 0.244967512 | 0.4329436 | -0.27119217 | 0.015 | 0.058 | 0.04844788 | 1 |
| WBGene00004462 | rpn-6.1 | 0.24445716 | 0.431175708 | -0.269377923 | 0.023 | 0.058 | 0.117145672 | 1 |
| WBGene00016938 | C54H2.4 | 0.306670409 | 0.49318156 | -0.269078713 | 0.023 | 0.07 | 0.052461718 | 1 |
| WBGene00003620 | nhr-21 | 1.61613784 | 1.802478726 | -0.268833072 | 0.215 | 0.412 | 0.000967972 | 1 |
| WBGene00009454 | F36A2.7 | 0.617359734 | 0.803245185 | -0.268176019 | 0.054 | 0.118 | 0.043471885 | 1 |
| WBGene00001121 | dyf-5 | 0.825370629 | 1.011245424 | -0.268160645 | 0.092 | 0.116 | 0.53897652 | 1 |
| WBGene00012097 | abcf-2 | 0.413200089 | 0.598960984 | -0.267996321 | 0.038 | 0.079 | 0.12003237 | 1 |
| WBGene00001470 | baz-2 | 0.447401162 | 0.632815973 | -0.267497028 | 0.038 | 0.099 | 0.037326301 | 1 |
| WBGene00003065 | lpd-9 | 0.252519384 | 0.437697603 | -0.267155698 | 0.031 | 0.054 | 0.284221025 | 1 |
| WBGene00000778 | cpn-2 | 1.444226689 | 1.629161062 | -0.266803902 | 0.215 | 0.354 | 0.016647918 | 1 |

| | | | | | | | | |
|---|---|---|---|---|---|---|---|---|
| WBGene00008414 | D2030.4 | 0.266392991 | 0.450564145 | -0.26570281 | 0.015 | 0.058 | 0.049410086 | 1 |
| WBGene00010110 | F55D12.1 | 1.188234174 | 1.371748399 | -0.264755063 | 0.185 | 0.282 | 0.091598305 | 1 |
| WBGene00011344 | T01G9.2 | 0.231058392 | 0.413981181 | -0.263901801 | 0.023 | 0.06 | 0.102019058 | 1 |
| WBGene00001562 | lin-66 | 0.27768933 | 0.460352096 | -0.263526667 | 0.023 | 0.066 | 0.068614344 | 1 |
| WBGene00007443 | pfd-1 | 0.463093849 | 0.645292538 | -0.262857145 | 0.038 | 0.11 | 0.016464873 | 1 |
| WBGene00002189 | kin-1 | 1.036073641 | 1.218002339 | -0.26246763 | 0.138 | 0.191 | 0.323564186 | 1 |
| WBGene00044650 | Y71G10AR.4 | 0.802147291 | 0.984002968 | -0.262362283 | 0.092 | 0.164 | 0.070956011 | 1 |
| WBGene00004896 | smu-2 | 0.417462588 | 0.598512261 | -0.261199466 | 0.023 | 0.07 | 0.052461733 | 1 |
| WBGene00008342 | frpr-5 | 0.782177019 | 0.962986554 | -0.260853019 | 0.085 | 0.145 | 0.114706949 | 1 |
| WBGene00003851 | odr-4 | 1.342516553 | 1.523240644 | -0.260729751 | 0.208 | 0.309 | 0.117784381 | 1 |
| WBGene00008765 | ttx-7 | 1.468561497 | 1.649124777 | -0.260497748 | 0.223 | 0.369 | 0.032905955 | 1 |
| WBGene00004450 | rpl-36 | 2.699914663 | 2.880320901 | -0.260271186 | 0.638 | 0.768 | 0.025787814 | 1 |
| WBGene00008520 | F02D8.1 | 1.774983246 | 1.955293548 | -0.260132779 | 0.323 | 0.447 | 0.10670913 | 1 |
| WBGene00010130 | vha-14 | 0.428110768 | 0.608334646 | -0.260008095 | 0.038 | 0.087 | 0.07936795 | 1 |
| WBGene00004408 | rla-0 | 3.343590408 | 3.523486879 | -0.259535746 | 0.808 | 0.876 | 0.058017895 | 1 |
| WBGene00015755 | C14B9.10 | 0.623773783 | 0.80354591 | -0.259356357 | 0.077 | 0.141 | 0.083541513 | 1 |
| WBGene00000160 | apb-1 | 0.228649679 | 0.408225829 | -0.259073621 | 0.023 | 0.056 | 0.131151174 | 1 |
| WBGene00014016 | ZK632.9 | 1.551385636 | 1.730795166 | -0.258833238 | 0.223 | 0.327 | 0.103275379 | 1 |
| WBGene00011312 | trcs-2 | 0.34771296 | 0.526819302 | -0.258395831 | 0.038 | 0.072 | 0.197483986 | 1 |
| WBGene00000244 | bbs-8 | 1.530847742 | 1.709869731 | -0.258274136 | 0.238 | 0.383 | 0.027450825 | 1 |
| WBGene00009085 | F23D12.1 | 0.37027337 | 0.549209973 | -0.258150949 | 0.031 | 0.079 | 0.057709117 | 1 |
| WBGene00020297 | nog-1 | 0.539813152 | 0.718628696 | -0.257976299 | 0.062 | 0.114 | 0.111434232 | 1 |
| WBGene00001898 | his-24 | 1.124428975 | 1.303241413 | -0.257971818 | 0.154 | 0.238 | 0.105948305 | 1 |
| WBGene00007647 | C17E4.10 | 1.598467257 | 1.777076486 | -0.257678649 | 0.231 | 0.352 | 0.037701223 | 1 |
| WBGene00007604 | xbx-9 | 1.16856837 | 1.34655058 | -0.256774051 | 0.169 | 0.284 | 0.037096412 | 1 |
| WBGene00003858 | ogt-1 | 1.039522388 | 1.216913054 | -0.255920635 | 0.131 | 0.211 | 0.101121687 | 1 |
| WBGene00012826 | dyf-19 | 0.731269118 | 0.908052569 | -0.255044608 | 0.1 | 0.141 | 0.298949075 | 1 |
| WBGene00019643 | K11C4.2 | 1.383713421 | 1.560478579 | -0.255018216 | 0.177 | 0.261 | 0.136466018 | 1 |
| WBGene00000085 | aex-2 | 1.540508656 | 1.71717399 | -0.254874201 | 0.246 | 0.375 | 0.058859782 | 1 |
| WBGene00008746 | F13E6.2 | 0.189214023 | 0.365578809 | -0.254440601 | 0.015 | 0.05 | 0.086171336 | 1 |
| WBGene00002203 | kin-20 | 0.339010754 | 0.515281605 | -0.254305082 | 0.038 | 0.058 | 0.401893874 | 1 |
| WBGene00006513 | rimb-1 | 0.490026453 | 0.666183741 | -0.254141245 | 0.046 | 0.093 | 0.108524005 | 1 |
| WBGene00002335 | let-60 | 0.32074757 | 0.496485451 | -0.253536169 | 0.031 | 0.064 | 0.16103944 | 1 |
| WBGene00013900 | ugt-18 | 0.322514382 | 0.497210097 | -0.252032642 | 0.031 | 0.066 | 0.146395757 | 1 |
| WBGene00020581 | T19D12.6 | 1.141231135 | 1.315844993 | -0.251914547 | 0.131 | 0.251 | 0.015768723 | 1 |
| WBGene00015900 | nhr-157 | 0.220753656 | 0.395054891 | -0.251463528 | 0.023 | 0.052 | 0.164674117 | 1 |
| WBGene00001701 | grd-12 | 0.972633296 | 1.146217206 | -0.250428646 | 0.115 | 0.203 | 0.038485069 | 1 |
| WBGene00004410 | rla-2 | 2.450399619 | 2.62384355 | -0.250226699 | 0.562 | 0.698 | 0.192873408 | 1 |
| WBGene00020825 | tmem-231 | 0.977287472 | 1.149987321 | -0.249153216 | 0.123 | 0.174 | 0.24796611 | 1 |
| WBGene00016289 | lntl-1 | 2.599739392 | 2.772213446 | -0.248827461 | 0.523 | 0.731 | 0.002281242 | 1 |
| WBGene00003822 | nsy-1 | 0.798085389 | 0.970209261 | -0.248322256 | 0.069 | 0.151 | 0.025225761 | 1 |
| WBGene00006412 | nlg-1 | 1.708055147 | 1.879846127 | -0.247841995 | 0.315 | 0.383 | 0.470139468 | 1 |
| WBGene00003827 | ntl-4 | 0.51942231 | 0.691030597 | -0.247578423 | 0.054 | 0.103 | 0.111768682 | 1 |
| WBGene00015064 | B0228.7 | 0.792336256 | 0.963605335 | -0.247089051 | 0.1 | 0.168 | 0.110607606 | 1 |
| WBGene00012468 | Y17G7B.17 | 0.850724639 | 1.020146794 | -0.244424502 | 0.108 | 0.217 | 0.015534445 | 1 |
| WBGene00016744 | bbs-9 | 0.801149849 | 0.970470788 | -0.244278479 | 0.1 | 0.191 | 0.037294179 | 1 |
| WBGene00018700 | lmd-3 | 1.224489217 | 1.393707899 | -0.244130953 | 0.162 | 0.259 | 0.062620829 | 1 |
| WBGene00006066 | sto-4 | 0.37917615 | 0.54838435 | -0.244115831 | 0.038 | 0.062 | 0.332133274 | 1 |
| WBGene00006600 | tph-1 | 2.309413452 | 2.478420464 | -0.243825578 | 0.477 | 0.64 | 0.022245204 | 1 |
| WBGene00019536 | rpi-2 | 0.219955683 | 0.388466628 | -0.243109905 | 0.023 | 0.06 | 0.101734859 | 1 |
| WBGene00022489 | Y119D3B.12 | 0.403034378 | 0.571521113 | -0.243074977 | 0.038 | 0.075 | 0.163856984 | 1 |
| WBGene00000881 | cyn-5 | 0.400568006 | 0.568487765 | -0.242257004 | 0.038 | 0.075 | 0.15079792 | 1 |
| WBGene00045339 | T10H4.13 | 0.21119128 | 0.378824169 | -0.241843138 | 0.023 | 0.052 | 0.160340101 | 1 |
| WBGene00045056 | F13H10.9 | 0.651775347 | 0.819290823 | -0.241673746 | 0.077 | 0.135 | 0.099116165 | 1 |
| WBGene00000064 | act-2 | 1.128151915 | 1.295337908 | -0.241198403 | 0.138 | 0.25 | 0.024399913 | 1 |
| WBGene00003961 | pct-1 | 0.728893013 | 0.895576978 | -0.240474129 | 0.069 | 0.151 | 0.026766161 | 1 |
| WBGene00006651 | tts-2 | 1.875064738 | 2.041651362 | -0.240333696 | 0.346 | 0.42 | 0.423888117 | 1 |
| WBGene00006917 | vha-8 | 0.216058674 | 0.381548988 | -0.238752055 | 0.023 | 0.054 | 0.149171713 | 1 |
| WBGene00010244 | F58D5.5 | 1.128063672 | 1.293303857 | -0.238391196 | 0.162 | 0.203 | 0.504229602 | 1 |
| WBGene00018582 | F48A9.2 | 0.459374418 | 0.624063212 | -0.237595706 | 0.054 | 0.089 | 0.215975141 | 1 |
| WBGene00004277 | rab-18 | 0.505227749 | 0.669489945 | -0.236980256 | 0.054 | 0.101 | 0.12826612 | 1 |
| WBGene00010367 | H05L14.2 | 0.522242495 | 0.686425812 | -0.236866458 | 0.062 | 0.103 | 0.190973248 | 1 |
| WBGene00004502 | rpt-2 | 0.248825316 | 0.41275177 | -0.236495883 | 0.023 | 0.05 | 0.194551239 | 1 |
| WBGene00002224 | klp-13 | 0.592145108 | 0.755772255 | -0.236064073 | 0.062 | 0.112 | 0.11307474 | 1 |

| | | | | | | | | |
|---|---|---|---|---|---|---|---|---|
| WBGene00010133 | F55H12.2 | 1.422428312 | 1.585701949 | -0.235554066 | 0.177 | 0.296 | 0.020499537 | 1 |
| WBGene00044995 | Y105C5A.28 | 0.631989914 | 0.792744711 | -0.231920149 | 0.046 | 0.126 | 0.014793363 | 1 |
| WBGene00011517 | cgt-1 | 0.518451554 | 0.679153069 | -0.23184328 | 0.046 | 0.112 | 0.033985358 | 1 |
| WBGene00002002 | hsb-1 | 0.231665429 | 0.392025045 | -0.231350022 | 0.023 | 0.056 | 0.131151144 | 1 |
| WBGene00008121 | C46F11.6 | 0.983994078 | 1.144132451 | -0.231030837 | 0.123 | 0.205 | 0.075162688 | 1 |
| WBGene00019888 | btbd-10 | 0.565347952 | 0.725306635 | -0.230771599 | 0.062 | 0.108 | 0.143288697 | 1 |
| WBGene00007192 | B0491.5 | 0.442282193 | 0.602206859 | -0.230722522 | 0.038 | 0.066 | 0.267154088 | 1 |
| WBGene00006476 | rhgf-2 | 0.529153805 | 0.688651824 | -0.230107002 | 0.046 | 0.106 | 0.0481487 | 1 |
| WBGene00000292 | cap-1 | 0.266397553 | 0.425653065 | -0.229757137 | 0.031 | 0.068 | 0.121310341 | 1 |
| WBGene00003825 | ntl-2 | 0.810138414 | 0.969178909 | -0.229446934 | 0.108 | 0.18 | 0.08790193 | 1 |
| WBGene00000464 | ceh-44 | 0.511668921 | 0.670578906 | -0.229258648 | 0.062 | 0.085 | 0.436072926 | 1 |
| WBGene00270303 | T28D9.16 | 0.593185797 | 0.751976319 | -0.229086299 | 0.062 | 0.114 | 0.104897521 | 1 |
| WBGene00009483 | F36G3.2 | 0.310184473 | 0.468917838 | -0.229003838 | 0.015 | 0.072 | 0.01840592 | 1 |
| WBGene00022788 | slc-17.7 | 0.230218138 | 0.388677288 | -0.228608231 | 0.015 | 0.056 | 0.056707961 | 1 |
| WBGene00006536 | tbb-1 | 0.574180937 | 0.732319983 | -0.228146417 | 0.069 | 0.106 | 0.240045002 | 1 |
| WBGene00004238 | ptr-24 | 0.405112418 | 0.562761309 | -0.227439273 | 0.031 | 0.06 | 0.202626712 | 1 |
| WBGene00012893 | Y45F10D.15 | 0.353578978 | 0.510812254 | -0.226839668 | 0.031 | 0.072 | 0.101089891 | 1 |
| WBGene00001177 | egl-8 | 0.57214089 | 0.729307885 | -0.226744044 | 0.062 | 0.108 | 0.144399979 | 1 |
| WBGene00020151 | T01G6.1 | 0.308582815 | 0.465686527 | -0.226652746 | 0.023 | 0.066 | 0.066461349 | 1 |
| WBGene00017193 | F07C3.2 | 0.193653123 | 0.350664306 | -0.226519255 | 0.008 | 0.05 | 0.031689318 | 1 |
| WBGene00199110 | C32D5.17 | 0.372694744 | 0.529314465 | -0.225954495 | 0.023 | 0.075 | 0.031915822 | 1 |
| WBGene00015868 | C16H3.3 | 0.584573665 | 0.740278152 | -0.224634092 | 0.062 | 0.114 | 0.101745309 | 1 |
| WBGene00013994 | ZK524.4 | 0.348704352 | 0.503800099 | -0.223755865 | 0.023 | 0.07 | 0.053084208 | 1 |
| WBGene00015328 | C02B10.3 | 0.235782423 | 0.390384836 | -0.223044134 | 0.023 | 0.058 | 0.117145672 | 1 |
| WBGene00003060 | lpd-3 | 0.340939099 | 0.494420789 | -0.221427273 | 0.038 | 0.066 | 0.267703045 | 1 |
| WBGene00010131 | F55H2.5 | 0.250676001 | 0.404049362 | -0.221270987 | 0.023 | 0.07 | 0.052772204 | 1 |
| WBGene00004187 | prp-8 | 0.886215927 | 1.039190651 | -0.220695875 | 0.1 | 0.151 | 0.209861714 | 1 |
| WBGene00021254 | Y22D7AL.16 | 0.656937985 | 0.809529917 | -0.220143624 | 0.085 | 0.135 | 0.162432099 | 1 |
| WBGene00014454 | MTCE.7 | 2.970034381 | 3.122360618 | -0.219760307 | 0.677 | 0.741 | 0.78294885 | 1 |
| WBGene00022075 | Y69A2AR.3 | 0.225385638 | 0.377326217 | -0.219203921 | 0.023 | 0.058 | 0.117467889 | 1 |
| WBGene00021350 | Y37E3.8 | 3.121015052 | 3.272753723 | -0.218912628 | 0.815 | 0.841 | 0.021168016 | 1 |
| WBGene00011605 | sftb-1 | 0.235481575 | 0.387177354 | -0.218850748 | 0.023 | 0.06 | 0.102019083 | 1 |
| WBGene00017814 | F26A10.2 | 0.455185581 | 0.606503079 | -0.218305004 | 0.038 | 0.07 | 0.206162106 | 1 |
| WBGene00004390 | rnp-7 | 1.092830806 | 1.243702358 | -0.217661641 | 0.146 | 0.219 | 0.163254218 | 1 |
| WBGene00010484 | dph-3 | 0.304251147 | 0.454996399 | -0.217479427 | 0.023 | 0.066 | 0.06941132 | 1 |
| WBGene00005742 | srv-31 | 0.305247421 | 0.455699062 | -0.217055836 | 0.015 | 0.064 | 0.032209075 | 1 |
| WBGene00009004 | pfd-6 | 0.393550599 | 0.543789799 | -0.216749348 | 0.031 | 0.089 | 0.033297411 | 1 |
| WBGene00012344 | ola-1 | 0.265887686 | 0.415775602 | -0.216242553 | 0.023 | 0.056 | 0.130084389 | 1 |
| WBGene00022475 | Y119C1B.10 | 0.344051971 | 0.493774345 | -0.216003727 | 0.031 | 0.056 | 0.257437727 | 1 |
| WBGene00021205 | Y17G9B.8 | 0.306064193 | 0.455425323 | -0.215482561 | 0.031 | 0.064 | 0.164972594 | 1 |
| WBGene00017852 | F27C1.2 | 0.184171867 | 0.333531062 | -0.215479769 | 0.015 | 0.052 | 0.074890843 | 1 |
| WBGene00004456 | rpl-43 | 2.741997511 | 2.891196034 | -0.215247969 | 0.677 | 0.766 | 0.097989172 | 1 |
| WBGene00000149 | apl-1 | 2.259264141 | 2.407818626 | -0.214318818 | 0.446 | 0.602 | 0.037458144 | 1 |
| WBGene00013238 | trap-4 | 0.237538804 | 0.385784371 | -0.213873144 | 0.023 | 0.062 | 0.090031045 | 1 |
| WBGene00021953 | Y55H10A.2 | 0.872210064 | 1.020219275 | -0.213532155 | 0.092 | 0.164 | 0.065460194 | 1 |
| WBGene00019893 | sgt-1 | 0.739980344 | 0.887936062 | -0.213454981 | 0.092 | 0.143 | 0.184969432 | 1 |
| WBGene00004449 | rpl-35 | 2.76056154 | 2.908209841 | -0.213011472 | 0.669 | 0.756 | 0.060505985 | 1 |
| WBGene00003367 | mix-1 | 0.693540996 | 0.840513225 | -0.212036106 | 0.077 | 0.124 | 0.179998744 | 1 |
| WBGene00001681 | gpc-1 | 1.94691659 | 2.093675369 | -0.211728163 | 0.4 | 0.503 | 0.240177999 | 1 |
| WBGene00004313 | rbc-1 | 0.389989952 | 0.536429576 | -0.211267719 | 0.038 | 0.07 | 0.213846726 | 1 |
| WBGene00003053 | lmp-1 | 0.679225131 | 0.825204189 | -0.210603263 | 0.069 | 0.141 | 0.041691202 | 1 |
| WBGene00009732 | cope-1 | 0.605564594 | 0.751035602 | -0.209870301 | 0.062 | 0.097 | 0.271294941 | 1 |
| WBGene00006843 | unc-119 | 1.270360333 | 1.415021291 | -0.208701647 | 0.177 | 0.265 | 0.142007951 | 1 |
| WBGene00007855 | C31H5.4 | 0.57313696 | 0.717597145 | -0.208411992 | 0.062 | 0.13 | 0.049026175 | 1 |
| WBGene00016393 | C34B2.8 | 0.193882799 | 0.338242079 | -0.208266416 | 0.015 | 0.052 | 0.075852375 | 1 |
| WBGene00011953 | ppm-2 | 0.719937324 | 0.863511531 | -0.207133797 | 0.085 | 0.122 | 0.304280272 | 1 |
| WBGene00044698 | K09F5.6 | 0.525936698 | 0.668806968 | -0.20611823 | 0.038 | 0.097 | 0.042977478 | 1 |
| WBGene00185118 | K02E2.11 | 0.858716284 | 1.001474241 | -0.205956197 | 0.108 | 0.161 | 0.24125218 | 1 |
| WBGene00004826 | skr-20 | 0.263463283 | 0.406126188 | -0.205819066 | 0.031 | 0.064 | 0.160261374 | 1 |
| WBGene00006460 | ppm-1.A | 0.997253286 | 1.139774553 | -0.205614726 | 0.131 | 0.217 | 0.080264327 | 1 |
| WBGene00012117 | T28B11.1 | 2.108332777 | 2.250556455 | -0.205185395 | 0.354 | 0.547 | 0.005825017 | 1 |
| WBGene00017511 | srbc-41 | 2.382297971 | 2.52364013 | -0.203913631 | 0.477 | 0.634 | 0.042310687 | 1 |
| WBGene00002637 | let-418 | 0.587855258 | 0.728362195 | -0.202708661 | 0.077 | 0.122 | 0.189668461 | 1 |
| WBGene00016318 | C32D5.11 | 0.382730894 | 0.522946196 | -0.202287921 | 0.038 | 0.06 | 0.367148897 | 1 |

| | | | | | | | | |
|---|---|---|---|---|---|---|---|---|
| WBGene00017350 | F10E7.9 | 0.545048601 | 0.685255435 | -0.202275703 | 0.054 | 0.101 | 0.121296336 | 1 |
| WBGene00010631 | cash-1 | 0.323104912 | 0.463082146 | -0.201944461 | 0.031 | 0.054 | 0.29182805 | 1 |
| WBGene00019025 | srb-16 | 0.725670104 | 0.865500875 | -0.20173316 | 0.085 | 0.116 | 0.387642613 | 1 |
| WBGene00004740 | scd-2 | 2.211572119 | 2.351291649 | -0.201572673 | 0.408 | 0.594 | 0.029123132 | 1 |
| WBGene00019875 | mca-2 | 0.48490305 | 0.623671326 | -0.200200305 | 0.031 | 0.099 | 0.017140584 | 1 |
| WBGene00003902 | pab-1 | 1.44122692 | 1.578635289 | -0.198238374 | 0.208 | 0.342 | 0.033686276 | 1 |
| WBGene00015007 | ain-2 | 0.460074998 | 0.596219216 | -0.196414588 | 0.046 | 0.054 | 0.749197323 | 1 |
| WBGene00006739 | ulp-4 | 0.603493388 | 0.739603523 | -0.196365417 | 0.046 | 0.083 | 0.17822352 | 1 |
| WBGene00016314 | C32D5.6 | 0.606434505 | 0.742087692 | -0.19570618 | 0.046 | 0.093 | 0.104256119 | 1 |
| WBGene00017045 | atg-13 | 0.764606295 | 0.900028354 | -0.195372733 | 0.092 | 0.168 | 0.063737007 | 1 |
| WBGene00004728 | sax-2 | 0.318540884 | 0.453705719 | -0.195001638 | 0.031 | 0.066 | 0.144614244 | 1 |
| WBGene00008144 | gasr-8 | 0.483379558 | 0.618543104 | -0.194999777 | 0.046 | 0.108 | 0.041685346 | 1 |
| WBGene00020506 | dop-3 | 1.004581662 | 1.139043777 | -0.193987827 | 0.115 | 0.174 | 0.187619138 | 1 |
| WBGene00021004 | sftb-2 | 0.283139499 | 0.417468811 | -0.193796232 | 0.031 | 0.06 | 0.196990864 | 1 |
| WBGene00003715 | nhr-125 | 0.5134469 | 0.647636475 | -0.193594635 | 0.046 | 0.075 | 0.272710149 | 1 |
| WBGene00017121 | cyc-2.1 | 1.449223501 | 1.583375724 | -0.193540746 | 0.215 | 0.36 | 0.025833459 | 1 |
| WBGene00004412 | rpl-1 | 2.482662778 | 2.616466195 | -0.193037526 | 0.577 | 0.702 | 0.178242925 | 1 |
| WBGene00010012 | saeg-1 | 0.902021667 | 1.034807784 | -0.191569873 | 0.108 | 0.161 | 0.225185378 | 1 |
| WBGene00006433 | sdhb-1 | 0.481342344 | 0.613765748 | -0.191046588 | 0.054 | 0.079 | 0.366628732 | 1 |
| WBGene00003014 | lin-28 | 0.236573277 | 0.368850311 | -0.19083542 | 0.023 | 0.052 | 0.172243412 | 1 |
| WBGene00003923 | pas-2 | 0.331319598 | 0.463372109 | -0.190511503 | 0.038 | 0.064 | 0.288202285 | 1 |
| WBGene00013343 | prp-6 | 0.333142921 | 0.464577902 | -0.189620595 | 0.031 | 0.054 | 0.291188957 | 1 |
| WBGene00017615 | srsx-32 | 1.551444135 | 1.682646774 | -0.189285397 | 0.231 | 0.34 | 0.095416046 | 1 |
| WBGene00004387 | rnp-4 | 0.669974414 | 0.800690474 | -0.188583411 | 0.077 | 0.112 | 0.302761443 | 1 |
| WBGene00013354 | snx-6 | 0.483348778 | 0.614055884 | -0.188570493 | 0.046 | 0.05 | 0.875106987 | 1 |
| WBGene00016868 | C52B9.8 | 1.03322039 | 1.163877408 | -0.188498232 | 0.131 | 0.221 | 0.06197353 | 1 |
| WBGene00022678 | sar-1 | 0.409917641 | 0.540199281 | -0.187956676 | 0.038 | 0.091 | 0.061483865 | 1 |
| WBGene00003403 | mps-1 | 0.940603879 | 1.070665255 | -0.187638901 | 0.115 | 0.209 | 0.039249677 | 1 |
| WBGene00206362 | B0041.11 | 0.468795128 | 0.598741341 | -0.187472758 | 0.046 | 0.064 | 0.502868331 | 1 |
| WBGene00006941 | wnk-1 | 0.68511865 | 0.814748318 | -0.187016079 | 0.062 | 0.101 | 0.217376772 | 1 |
| WBGene00011146 | pde-2 | 0.892707771 | 1.022285395 | -0.186940997 | 0.1 | 0.176 | 0.070342688 | 1 |
| WBGene00001088 | dpy-30 | 0.617948534 | 0.74737707 | -0.186725907 | 0.062 | 0.137 | 0.032127938 | 1 |
| WBGene00001565 | gei-8 | 0.417845922 | 0.547016272 | -0.186353423 | 0.038 | 0.068 | 0.242462908 | 1 |
| WBGene00010408 | mboa-2 | 0.580871065 | 0.709357029 | -0.185366062 | 0.069 | 0.116 | 0.171697149 | 1 |
| WBGene00006055 | ssr-2 | 1.173813691 | 1.30223602 | -0.185274258 | 0.146 | 0.267 | 0.022245924 | 1 |
| WBGene00004087 | ppk-1 | 0.455568035 | 0.583648482 | -0.184781026 | 0.046 | 0.095 | 0.10026702 | 1 |
| WBGene00022139 | tub-2 | 0.477132603 | 0.605061748 | -0.184562744 | 0.054 | 0.058 | 0.909130357 | 1 |
| WBGene00015303 | rga-6 | 1.391751535 | 1.519325632 | -0.184050517 | 0.223 | 0.292 | 0.323734017 | 1 |
| WBGene00002152 | iars-1 | 0.375294799 | 0.502102624 | -0.182945021 | 0.038 | 0.056 | 0.448343466 | 1 |
| WBGene00003883 | osm-1 | 1.151615509 | 1.277356433 | -0.181405808 | 0.177 | 0.236 | 0.296791376 | 1 |
| WBGene00003368 | mkk-4 | 0.326597428 | 0.452072041 | -0.181021602 | 0.031 | 0.064 | 0.156795155 | 1 |
| WBGene00000295 | cat-1 | 1.718706289 | 1.844132685 | -0.180952039 | 0.277 | 0.414 | 0.117649579 | 1 |
| WBGene00009974 | F53C11.4 | 0.755515076 | 0.880881912 | -0.180866113 | 0.092 | 0.164 | 0.07135317 | 1 |
| WBGene00022389 | pde-6 | 1.021038056 | 1.145798991 | -0.179991983 | 0.146 | 0.211 | 0.200328012 | 1 |
| WBGene00013308 | nuo-3 | 1.099433985 | 1.223354957 | -0.178780173 | 0.154 | 0.253 | 0.06421034 | 1 |
| WBGene00015790 | C15C7.4 | 2.116191919 | 2.239583523 | -0.178016455 | 0.277 | 0.422 | 0.02343451 | 1 |
| WBGene00017321 | F09G8.7 | 0.367486457 | 0.490847945 | -0.177973007 | 0.038 | 0.07 | 0.22220635 | 1 |
| WBGene00000188 | arl-3 | 2.143985966 | 2.267086364 | -0.177596334 | 0.454 | 0.578 | 0.19609602 | 1 |
| WBGene00012274 | W05B5.1 | 0.843347668 | 0.965784945 | -0.176639654 | 0.1 | 0.143 | 0.289302999 | 1 |
| WBGene00006595 | top-1 | 1.114213745 | 1.236562664 | -0.17651218 | 0.138 | 0.213 | 0.156753623 | 1 |
| WBGene00011240 | mask-1 | 0.591202447 | 0.713490733 | -0.176424703 | 0.069 | 0.087 | 0.56820097 | 1 |
| WBGene00013322 | Y57G11C.32 | 0.248809094 | 0.370192048 | -0.175118586 | 0.023 | 0.052 | 0.166652913 | 1 |
| WBGene00003934 | pat-10 | 0.501250619 | 0.622606336 | -0.175079291 | 0.046 | 0.079 | 0.236635618 | 1 |
| WBGene00009661 | patr-1 | 0.268760077 | 0.389893967 | -0.174759263 | 0.031 | 0.052 | 0.321823097 | 1 |
| WBGene00020682 | T22D1.3 | 0.306509275 | 0.42716567 | -0.174070383 | 0.031 | 0.066 | 0.148555679 | 1 |
| WBGene00010849 | M04B2.7 | 0.437233275 | 0.557137184 | -0.172984774 | 0.046 | 0.083 | 0.185715887 | 1 |
| WBGene00001488 | frm-1 | 0.773066398 | 0.892725106 | -0.172631024 | 0.092 | 0.133 | 0.311929077 | 1 |
| WBGene00018967 | ddx-15 | 0.275648071 | 0.394716493 | -0.171779422 | 0.023 | 0.064 | 0.079639561 | 1 |
| WBGene00016511 | C37H5.13 | 0.398699093 | 0.51740171 | -0.171251677 | 0.038 | 0.072 | 0.199199206 | 1 |
| WBGene00000381 | cct-6 | 0.435478923 | 0.554125593 | -0.171170962 | 0.054 | 0.085 | 0.284602844 | 1 |
| WBGene00002182 | kap-1 | 0.445706379 | 0.563866001 | -0.1704683 | 0.038 | 0.07 | 0.218462511 | 1 |
| WBGene00004046 | plp-1 | 0.977603786 | 1.095227624 | -0.169695327 | 0.115 | 0.164 | 0.261188419 | 1 |
| WBGene00021534 | mvk-1 | 0.431837518 | 0.549038751 | -0.169085638 | 0.046 | 0.074 | 0.306894369 | 1 |
| WBGene00001746 | gsk-3 | 0.723075894 | 0.840011011 | -0.168701713 | 0.092 | 0.093 | 0.929760002 | 1 |

| | | | | | | | | |
|---|---|---|---|---|---|---|---|---|
| WBGene00014145 | ZK899.7 | 0.938524135 | 1.055385671 | -0.168595558 | 0.1 | 0.182 | 0.05763775 | 1 |
| WBGene00016121 | ift-43 | 1.047573597 | 1.164156053 | -0.168192931 | 0.154 | 0.219 | 0.235896621 | 1 |
| WBGene00195084 | C43C3.4 | 0.691505144 | 0.808077383 | -0.168178192 | 0.077 | 0.118 | 0.251435834 | 1 |
| WBGene00015249 | otub-4 | 0.281893279 | 0.398272368 | -0.167899535 | 0.031 | 0.054 | 0.286741595 | 1 |
| WBGene00004441 | rpl-27 | 1.976413338 | 2.092697358 | -0.16776238 | 0.354 | 0.509 | 0.079397055 | 1 |
| WBGene00001447 | flp-4 | 1.119518448 | 1.235625419 | -0.167506951 | 0.092 | 0.153 | 0.10683399 | 1 |
| WBGene00004879 | smg-1 | 0.282609754 | 0.397992983 | -0.166462812 | 0.031 | 0.056 | 0.265035275 | 1 |
| WBGene00022233 | ipla-6 | 0.364085779 | 0.478877996 | -0.165610163 | 0.031 | 0.07 | 0.115540566 | 1 |
| WBGene00019162 | eif-1.A | 0.396699338 | 0.511447918 | -0.165547207 | 0.046 | 0.072 | 0.331065254 | 1 |
| WBGene00006331 | sup-26 | 0.617027732 | 0.731455414 | -0.165084249 | 0.054 | 0.075 | 0.425419811 | 1 |
| WBGene00006727 | ubq-1 | 1.759966325 | 1.874115483 | -0.164682424 | 0.338 | 0.443 | 0.268000089 | 1 |
| WBGene00006812 | unc-80 | 0.318859577 | 0.432271312 | -0.163618548 | 0.031 | 0.062 | 0.183535823 | 1 |
| WBGene00000554 | cnb-1 | 0.824650916 | 0.937602054 | -0.162954046 | 0.085 | 0.159 | 0.058779475 | 1 |
| WBGene00001030 | dnj-12 | 0.612638189 | 0.724843492 | -0.161878034 | 0.062 | 0.101 | 0.218350635 | 1 |
| WBGene00001462 | flp-19 | 0.73432476 | 0.845400269 | -0.160248086 | 0.085 | 0.106 | 0.553091278 | 1 |
| WBGene00020051 | mfsd-6 | 0.482445127 | 0.592754153 | -0.159142285 | 0.062 | 0.077 | 0.589948116 | 1 |
| WBGene00005909 | srx-18 | 0.921130687 | 1.031416684 | -0.159109061 | 0.108 | 0.143 | 0.414148099 | 1 |
| WBGene00015547 | ain-1 | 0.437361957 | 0.547208786 | -0.158475476 | 0.038 | 0.074 | 0.170270423 | 1 |
| WBGene00015477 | attf-4 | 0.309410173 | 0.418999837 | -0.158104466 | 0.031 | 0.05 | 0.365222623 | 1 |
| WBGene00018757 | tag-304 | 0.684516019 | 0.793693475 | -0.157509775 | 0.069 | 0.116 | 0.171083577 | 1 |
| WBGene00017288 | F09E5.10 | 0.61167755 | 0.720804285 | -0.157436599 | 0.062 | 0.101 | 0.216212237 | 1 |
| WBGene00001972 | hmg-1.2 | 1.016710239 | 1.12578283 | -0.157358486 | 0.154 | 0.19 | 0.559831074 | 1 |
| WBGene00021702 | Y48G9A.9 | 0.277790355 | 0.386565612 | -0.156929523 | 0.023 | 0.056 | 0.134756131 | 1 |
| WBGene00015552 | C06G3.6 | 0.253909347 | 0.362291988 | -0.156363098 | 0.023 | 0.05 | 0.194048094 | 1 |
| WBGene00022836 | ZK973.11 | 0.305930731 | 0.413499992 | -0.155189639 | 0.031 | 0.074 | 0.093031416 | 1 |
| WBGene00022089 | Y69A2AR.18 | 0.818776904 | 0.926084496 | -0.15481213 | 0.108 | 0.143 | 0.413644697 | 1 |
| WBGene00006965 | xtr-1 | 0.593037756 | 0.70019346 | -0.154593004 | 0.054 | 0.091 | 0.20860628 | 1 |
| WBGene00007350 | sucl-1 | 0.469815213 | 0.576873141 | -0.154451941 | 0.038 | 0.074 | 0.173513107 | 1 |
| WBGene00007180 | B0457.6 | 1.593384437 | 1.700375179 | -0.154355013 | 0.269 | 0.348 | 0.297588987 | 1 |
| WBGene00006468 | rhgf-1 | 0.313472919 | 0.420091379 | -0.153817924 | 0.023 | 0.052 | 0.170439427 | 1 |
| WBGene00019733 | M02E1.1 | 1.006443563 | 1.112915538 | -0.15360659 | 0.1 | 0.155 | 0.188042703 | 1 |
| WBGene00012149 | VF13D12L.3 | 0.336920078 | 0.443165946 | -0.153280386 | 0.038 | 0.064 | 0.308546465 | 1 |
| WBGene00000485 | che-3 | 1.631751629 | 1.73692845 | -0.151738077 | 0.254 | 0.348 | 0.184790371 | 1 |
| WBGene00010958 | ndfl-4 | 1.433549828 | 1.53817995 | -0.150949358 | 0.223 | 0.267 | 0.61356999 | 1 |
| WBGene00006823 | unc-94 | 0.994071881 | 1.098173079 | -0.150186283 | 0.131 | 0.209 | 0.106244756 | 1 |
| WBGene00006572 | tin-9.1 | 0.583817647 | 0.687389337 | -0.149422365 | 0.069 | 0.11 | 0.221803384 | 1 |
| WBGene00012348 | pptr-1 | 0.948195053 | 1.051500483 | -0.149038231 | 0.123 | 0.19 | 0.143407655 | 1 |
| WBGene00001081 | dpy-22 | 0.365421641 | 0.467612971 | -0.147430926 | 0.038 | 0.054 | 0.492482901 | 1 |
| WBGene00022144 | pghm-1 | 1.351642341 | 1.453582499 | -0.14706856 | 0.192 | 0.271 | 0.186359902 | 1 |
| WBGene00016454 | C35E7.2 | 0.960637038 | 1.062304718 | -0.146675457 | 0.123 | 0.18 | 0.211457034 | 1 |
| WBGene00016140 | rpb-2 | 0.341553141 | 0.443063768 | -0.146448877 | 0.031 | 0.06 | 0.210322553 | 1 |
| WBGene00006803 | unc-70 | 0.616697351 | 0.717960639 | -0.146092045 | 0.069 | 0.11 | 0.227526101 | 1 |
| WBGene00006919 | vha-10 | 0.556959195 | 0.658148378 | -0.145985134 | 0.062 | 0.085 | 0.398993307 | 1 |
| WBGene00007511 | C10C5.7 | 0.308058011 | 0.408689705 | -0.145180847 | 0.031 | 0.056 | 0.259177479 | 1 |
| WBGene00023417 | swm-1 | 1.303907266 | 1.404419796 | -0.145008929 | 0.162 | 0.222 | 0.26405992 | 1 |
| WBGene00003084 | lst-2 | 0.430886153 | 0.530767729 | -0.144098653 | 0.038 | 0.077 | 0.139002041 | 1 |
| WBGene00008224 | slrp-1 | 0.561657361 | 0.661259775 | -0.143695909 | 0.062 | 0.112 | 0.128369651 | 1 |
| WBGene00004207 | ptb-1 | 0.910695024 | 1.010137798 | -0.143465597 | 0.085 | 0.17 | 0.030620496 | 1 |
| WBGene00003656 | nhr-66 | 0.756974493 | 0.856294154 | -0.143287983 | 0.077 | 0.12 | 0.225183728 | 1 |
| WBGene00018776 | F53H1.1 | 0.533743462 | 0.632996115 | -0.143191311 | 0.054 | 0.097 | 0.157154187 | 1 |
| WBGene00008915 | F17C11.4 | 0.778570255 | 0.877434883 | -0.142631509 | 0.1 | 0.135 | 0.359158772 | 1 |
| WBGene00002583 | let-363 | 0.343686934 | 0.442473259 | -0.14251854 | 0.031 | 0.062 | 0.181800312 | 1 |
| WBGene00007247 | C01G12.7 | 2.658627105 | 2.757085362 | -0.142045239 | 0.6 | 0.687 | 0.203186772 | 1 |
| WBGene00003950 | pbs-4 | 0.550159485 | 0.648179317 | -0.141412726 | 0.054 | 0.104 | 0.101963437 | 1 |
| WBGene00010553 | K04C1.3 | 0.379865925 | 0.477748896 | -0.141215277 | 0.046 | 0.085 | 0.172091561 | 1 |
| WBGene00003153 | mca-3 | 0.980603484 | 1.077845842 | -0.140291067 | 0.131 | 0.176 | 0.386944483 | 1 |
| WBGene00017663 | F21D12.3 | 1.342183958 | 1.438922669 | -0.139564459 | 0.215 | 0.3 | 0.225857551 | 1 |
| WBGene00017422 | F13C5.1 | 0.556771829 | 0.653108609 | -0.138984595 | 0.062 | 0.118 | 0.091762296 | 1 |
| WBGene00020334 | epg-7 | 0.58449102 | 0.680349905 | -0.138295139 | 0.054 | 0.07 | 0.587672169 | 1 |
| WBGene00006462 | svh-5 | 0.989792424 | 1.085456223 | -0.138013689 | 0.085 | 0.172 | 0.031492434 | 1 |
| WBGene00219763 | linc-94 | 0.405462123 | 0.500781587 | -0.137516918 | 0.038 | 0.064 | 0.300658366 | 1 |
| WBGene00015287 | osta-1 | 0.541738383 | 0.636834683 | -0.137194961 | 0.046 | 0.091 | 0.114863624 | 1 |
| WBGene00000763 | coq-3 | 0.422190692 | 0.516568867 | -0.136158925 | 0.046 | 0.07 | 0.360787631 | 1 |
| WBGene00017359 | F10E9.10 | 0.571006989 | 0.665158836 | -0.135832402 | 0.054 | 0.066 | 0.679830057 | 1 |

| | | | | | | | | |
|---|---|---|---|---|---|---|---|---|
| WBGene00019395 | K04F10.2 | 0.764822571 | 0.858745591 | -0.135502274 | 0.085 | 0.13 | 0.21927849 | 1 |
| WBGene00012139 | T28F4.4 | 0.320591148 | 0.413964544 | -0.134709335 | 0.031 | 0.052 | 0.326696333 | 1 |
| WBGene00017060 | D2063.1 | 3.14274759 | 3.234882378 | -0.132922401 | 0.608 | 0.625 | 0.632674018 | 1 |
| WBGene00004281 | rab-28 | 1.273877222 | 1.365990926 | -0.132891984 | 0.177 | 0.294 | 0.050428451 | 1 |
| WBGene00003564 | ncs-2 | 0.830546841 | 0.92219711 | -0.13222339 | 0.092 | 0.126 | 0.365234296 | 1 |
| WBGene00008446 | E01G4.3 | 1.45021374 | 1.541042891 | -0.131038766 | 0.192 | 0.269 | 0.23348721 | 1 |
| WBGene00044989 | Y37A1B.17 | 0.329188559 | 0.41992629 | -0.130906874 | 0.031 | 0.054 | 0.297622283 | 1 |
| WBGene00004452 | rpl-38 | 3.305897256 | 3.396612895 | -0.130875003 | 0.854 | 0.851 | 0.030721882 | 1 |
| WBGene00002264 | lec-1 | 0.760502283 | 0.850698195 | -0.130125196 | 0.085 | 0.126 | 0.281355141 | 1 |
| WBGene00020246 | clic-1 | 0.685725895 | 0.775521134 | -0.129547147 | 0.085 | 0.122 | 0.294767519 | 1 |
| WBGene00011352 | rskn-1 | 0.908916736 | 0.998039392 | -0.128576813 | 0.108 | 0.153 | 0.295256354 | 1 |
| WBGene00012779 | cdkl-1 | 0.953171696 | 1.042093804 | -0.128287484 | 0.131 | 0.178 | 0.321823212 | 1 |
| WBGene00010806 | M01E5.3 | 0.252472698 | 0.340859199 | -0.127514767 | 0.023 | 0.05 | 0.19708133 | 1 |
| WBGene00019841 | R02F11.3 | 0.697785405 | 0.785814374 | -0.126998957 | 0.077 | 0.118 | 0.246471858 | 1 |
| WBGene00008405 | ttll-12 | 0.949956138 | 1.037260259 | -0.125953223 | 0.108 | 0.128 | 0.663806195 | 1 |
| WBGene00001194 | egl-27 | 0.947609788 | 1.034864611 | -0.1258821 | 0.123 | 0.201 | 0.102559615 | 1 |
| WBGene00001788 | gst-40 | 2.074140816 | 2.1612362 | -0.125652079 | 0.292 | 0.497 | 0.001701703 | 1 |
| WBGene00004278 | rab-19 | 0.799330888 | 0.885532248 | -0.124362275 | 0.1 | 0.137 | 0.363780401 | 1 |
| WBGene00012289 | W05H12.1 | 0.817098274 | 0.902406475 | -0.123073718 | 0.077 | 0.137 | 0.100005612 | 1 |
| WBGene00010648 | K08C7.6 | 0.293931621 | 0.378335575 | -0.121769166 | 0.023 | 0.05 | 0.195055341 | 1 |
| WBGene00015232 | B0511.6 | 0.295368313 | 0.379638027 | -0.121575498 | 0.031 | 0.05 | 0.368282267 | 1 |
| WBGene00015913 | C17F4.7 | 0.982498888 | 1.066458619 | -0.121128288 | 0.1 | 0.085 | 0.524547252 | 1 |
| WBGene00015611 | C08F1.8 | 0.652996026 | 0.736526603 | -0.120509149 | 0.077 | 0.106 | 0.383534632 | 1 |
| WBGene00006712 | ubc-17 | 0.559695565 | 0.642558595 | -0.119546083 | 0.054 | 0.083 | 0.313804062 | 1 |
| WBGene00011545 | T06G6.3 | 0.827637359 | 0.909495468 | -0.118096288 | 0.108 | 0.143 | 0.386760633 | 1 |
| WBGene00001173 | egl-4 | 0.319818665 | 0.40060366 | -0.116548112 | 0.031 | 0.05 | 0.372129134 | 1 |
| WBGene00020100 | mks-1 | 0.904467892 | 0.985002402 | -0.116186739 | 0.108 | 0.182 | 0.100789457 | 1 |
| WBGene00016463 | C35E7.11 | 1.104530295 | 1.185017091 | -0.116117901 | 0.154 | 0.213 | 0.283636428 | 1 |
| WBGene00004460 | rpn-3 | 0.863089303 | 0.943289916 | -0.115705027 | 0.115 | 0.149 | 0.455446143 | 1 |
| WBGene00018764 | F53F10.2 | 1.484926149 | 1.564207271 | -0.114378481 | 0.262 | 0.3 | 0.826214899 | 1 |
| WBGene00004302 | ran-1 | 0.987152858 | 1.065930352 | -0.113651901 | 0.138 | 0.182 | 0.376153893 | 1 |
| WBGene00012717 | Y39E4B.6 | 1.097341306 | 1.17594505 | -0.113401231 | 0.131 | 0.215 | 0.087055738 | 1 |
| WBGene00019240 | H24K24.3 | 0.426217636 | 0.50346489 | -0.11144423 | 0.054 | 0.044 | 0.611569353 | 1 |
| WBGene00010796 | clip-1 | 0.531328751 | 0.608557801 | -0.111417967 | 0.054 | 0.085 | 0.287902423 | 1 |
| WBGene00004930 | sod-1 | 0.664633831 | 0.74181757 | -0.111352597 | 0.077 | 0.116 | 0.249055302 | 1 |
| WBGene00015125 | B0303.3 | 0.523427511 | 0.600213291 | -0.110778465 | 0.054 | 0.093 | 0.177361316 | 1 |
| WBGene00001087 | dpy-28 | 0.625268217 | 0.701362726 | -0.109781172 | 0.069 | 0.104 | 0.284872932 | 1 |
| WBGene00021948 | kvs-5 | 0.594638553 | 0.670267717 | -0.109109821 | 0.062 | 0.104 | 0.175682845 | 1 |
| WBGene00003649 | nhr-59 | 0.588266264 | 0.662218736 | -0.106690865 | 0.062 | 0.116 | 0.104019283 | 1 |
| WBGene00008167 | acox-1.5 | 0.436360505 | 0.509818825 | -0.105977954 | 0.046 | 0.074 | 0.300673212 | 1 |
| WBGene00003634 | nhr-44 | 1.650685315 | 1.723296426 | -0.104755689 | 0.262 | 0.391 | 0.10340676 | 1 |
| WBGene00000479 | cgh-1 | 0.57787259 | 0.650041712 | -0.104118034 | 0.062 | 0.052 | 0.633098397 | 1 |
| WBGene00022412 | Y102A11A.2 | 0.677785573 | 0.749635803 | -0.103657971 | 0.077 | 0.085 | 0.857995761 | 1 |
| WBGene00019114 | nhr-194 | 0.644592825 | 0.716343436 | -0.103514252 | 0.062 | 0.11 | 0.144414244 | 1 |
| WBGene00020142 | aak-2 | 0.84681234 | 0.918356674 | -0.103216656 | 0.085 | 0.147 | 0.105609053 | 1 |
| WBGene00021044 | W05H7.1 | 0.499840376 | 0.571381464 | -0.103211973 | 0.054 | 0.091 | 0.216840617 | 1 |
| WBGene00021382 | Y37F4.1 | 3.15602483 | 3.226659076 | -0.101903676 | 0.608 | 0.621 | 0.811301194 | 1 |
| WBGene00011398 | qdpr-1 | 0.498753173 | 0.569257437 | -0.101716152 | 0.046 | 0.07 | 0.374512046 | 1 |
| WBGene00001440 | fkh-8 | 1.489448741 | 1.559921948 | -0.101671346 | 0.238 | 0.319 | 0.360542693 | 1 |
| WBGene00015390 | C03F11.4 | 0.581922844 | 0.652272889 | -0.101493661 | 0.062 | 0.095 | 0.28062567 | 1 |
| WBGene00001464 | flp-21 | 2.005147806 | 2.07521585 | -0.101086819 | 0.323 | 0.406 | 0.316403952 | 1 |
| WBGene00011962 | T23G5.2 | 0.243486525 | 0.313426246 | -0.100901688 | 0.023 | 0.05 | 0.196573394 | 1 |
| WBGene00014004 | fic-1 | 0.542832827 | 0.61264912 | -0.100723619 | 0.054 | 0.095 | 0.180757242 | 1 |
| WBGene00009917 | glb-18 | 1.86169683 | 1.931331582 | -0.100461711 | 0.331 | 0.433 | 0.255599911 | 1 |
| WBGene00010139 | F56A8.3 | 0.422474384 | 0.352859017 | 0.100433746 | 0.046 | 0.054 | 0.771944526 | 1 |
| WBGene00017311 | F09G2.2 | 0.585801236 | 0.515639936 | 0.10122136 | 0.062 | 0.079 | 0.553740318 | 1 |
| WBGene00016792 | C49H3.6 | 0.639690953 | 0.568191204 | 0.103152334 | 0.077 | 0.093 | 0.706484329 | 1 |
| WBGene00011028 | R05D7.3 | 0.640446427 | 0.568286664 | 0.104104533 | 0.069 | 0.074 | 0.957205992 | 1 |
| WBGene00016200 | dpff-1 | 0.504497764 | 0.431998779 | 0.104593927 | 0.054 | 0.048 | 0.740029892 | 1 |
| WBGene00002215 | klc-2 | 0.736480182 | 0.663908677 | 0.10469855 | 0.085 | 0.104 | 0.617713128 | 1 |
| WBGene00020374 | serp-1.2 | 0.495853993 | 0.423227314 | 0.10477815 | 0.054 | 0.066 | 0.686217395 | 1 |
| WBGene00016703 | rdl-1 | 1.092713143 | 1.020080039 | 0.104787419 | 0.138 | 0.141 | 0.786774499 | 1 |
| WBGene00001118 | dyf-2 | 0.785578582 | 0.71282455 | 0.104961882 | 0.092 | 0.097 | 0.966795915 | 1 |
| WBGene00016111 | C25H3.1 | 1.370604605 | 1.297205645 | 0.105892316 | 0.192 | 0.253 | 0.430637085 | 1 |

| | | | | | | | | |
|---|---|---|---|---|---|---|---|---|
| WBGene00004924 | snt-4 | 2.368839303 | 2.294214991 | 0.107660125 | 0.538 | 0.598 | 0.77305809 | 1 |
| WBGene00007971 | rpb-3 | 0.516370278 | 0.441151491 | 0.108517771 | 0.038 | 0.054 | 0.508680849 | 1 |
| WBGene00018316 | F41H10.3 | 0.620045918 | 0.544157557 | 0.109483761 | 0.077 | 0.07 | 0.6835248 | 1 |
| WBGene00009992 | F53F4.10 | 0.53042359 | 0.454414864 | 0.109657413 | 0.046 | 0.056 | 0.701247459 | 1 |
| WBGene00021248 | Y22D7AL.10 | 0.758150429 | 0.681358266 | 0.110787673 | 0.085 | 0.101 | 0.712586757 | 1 |
| WBGene00006655 | tub-1 | 1.076004715 | 0.998545905 | 0.111749441 | 0.169 | 0.155 | 0.467719401 | 1 |
| WBGene00008443 | pde-3 | 0.540544535 | 0.46272001 | 0.112277056 | 0.054 | 0.066 | 0.688046516 | 1 |
| WBGene00003175 | mec-12 | 0.488880178 | 0.410666754 | 0.112838119 | 0.054 | 0.05 | 0.827784429 | 1 |
| WBGene00016028 | flp-24 | 1.437713227 | 1.359474818 | 0.112874165 | 0.246 | 0.284 | 0.925469134 | 1 |
| WBGene00015978 | C18F10.7 | 0.491449886 | 0.412207989 | 0.114321891 | 0.062 | 0.041 | 0.287464046 | 1 |
| WBGene00008404 | D2013.6 | 0.513865755 | 0.434257016 | 0.114851133 | 0.046 | 0.062 | 0.553054226 | 1 |
| WBGene00009574 | F40E10.6 | 0.55791922 | 0.477994263 | 0.115307339 | 0.062 | 0.056 | 0.778634093 | 1 |
| WBGene00017120 | tctn-1 | 0.794066338 | 0.713212816 | 0.116646976 | 0.077 | 0.106 | 0.414754197 | 1 |
| WBGene00044988 | W01A8.8 | 0.834824897 | 0.753604897 | 0.117175692 | 0.1 | 0.122 | 0.620775989 | 1 |
| WBGene00010199 | bet-2 | 0.850019146 | 0.768426491 | 0.117713319 | 0.108 | 0.108 | 0.895824067 | 1 |
| WBGene00002078 | xpo-1 | 0.931014391 | 0.848973695 | 0.118359705 | 0.1 | 0.135 | 0.419288829 | 1 |
| WBGene00011512 | T05H10.8 | 1.327783442 | 1.245705822 | 0.118412976 | 0.185 | 0.201 | 0.975919154 | 1 |
| WBGene00003052 | lmn-1 | 1.044150819 | 0.961891259 | 0.118675458 | 0.115 | 0.155 | 0.422841103 | 1 |
| WBGene00003628 | nhr-35 | 0.516502465 | 0.433179214 | 0.120210041 | 0.054 | 0.064 | 0.742278035 | 1 |
| WBGene00021370 | nape-2 | 0.748722838 | 0.665277861 | 0.120385655 | 0.092 | 0.091 | 0.838006994 | 1 |
| WBGene00014017 | ZK632.10 | 0.970343994 | 0.886370481 | 0.121148172 | 0.131 | 0.157 | 0.721542432 | 1 |
| WBGene00012620 | Y38H6C.7 | 0.610106281 | 0.526070831 | 0.121237526 | 0.077 | 0.074 | 0.857437498 | 1 |
| WBGene00003818 | nsf-1 | 1.275323366 | 1.19017865 | 0.12283786 | 0.177 | 0.224 | 0.55194973 | 1 |
| WBGene00021539 | Y42H9AR.4 | 0.625135673 | 0.539799607 | 0.12311392 | 0.054 | 0.06 | 0.861438487 | 1 |
| WBGene00001745 | gsa-1 | 1.136711606 | 1.051052195 | 0.123580407 | 0.138 | 0.17 | 0.571916331 | 1 |
| WBGene00019150 | H04J21.1 | 0.563644962 | 0.477287221 | 0.124587885 | 0.062 | 0.068 | 0.888331323 | 1 |
| WBGene00004245 | puf-9 | 0.602028649 | 0.515497371 | 0.124838245 | 0.069 | 0.083 | 0.686903791 | 1 |
| WBGene00004490 | rps-21 | 2.622839972 | 2.5359372 | 0.125374199 | 0.638 | 0.656 | 0.374326577 | 1 |
| WBGene00015424 | C04E6.8 | 4.910712696 | 4.823720535 | 0.12550316 | 0.977 | 0.932 | 0.165534891 | 1 |
| WBGene00000123 | ama-1 | 0.698075116 | 0.610912836 | 0.125748589 | 0.085 | 0.083 | 0.864875441 | 1 |
| WBGene00020844 | T27A10.2 | 1.36088426 | 1.272501011 | 0.127510075 | 0.131 | 0.211 | 0.101042696 | 1 |
| WBGene00009141 | ncbp-2 | 0.85222447 | 0.763719591 | 0.12768555 | 0.092 | 0.139 | 0.264701856 | 1 |
| WBGene00020679 | ogdh-1 | 0.467715108 | 0.378349868 | 0.128926788 | 0.054 | 0.052 | 0.890407065 | 1 |
| WBGene00008601 | mob-2 | 0.955090902 | 0.86507324 | 0.129868034 | 0.123 | 0.132 | 0.996773071 | 1 |
| WBGene00004336 | ret-1 | 0.81708702 | 0.726007128 | 0.131400508 | 0.069 | 0.099 | 0.382880539 | 1 |
| WBGene00011304 | mnk-1 | 0.741384963 | 0.649880159 | 0.132013528 | 0.085 | 0.093 | 0.900457334 | 1 |
| WBGene00002262 | ldh-1 | 0.764496205 | 0.671985192 | 0.133465179 | 0.085 | 0.104 | 0.632676516 | 1 |
| WBGene00003927 | pas-6 | 0.644273887 | 0.551748268 | 0.133486252 | 0.062 | 0.077 | 0.601960783 | 1 |
| WBGene00007877 | nfki-1 | 1.191430763 | 1.09858022 | 0.133955018 | 0.177 | 0.18 | 0.702846351 | 1 |
| WBGene00022791 | ZK686.1 | 0.532597343 | 0.439419521 | 0.134427181 | 0.062 | 0.068 | 0.894097157 | 1 |
| WBGene00000408 | cdk-7 | 0.725731069 | 0.632493813 | 0.134512926 | 0.085 | 0.089 | 0.955723712 | 1 |
| WBGene00010965 | ctc-2 | 1.950522279 | 1.856927649 | 0.135028507 | 0.323 | 0.397 | 0.685152553 | 1 |
| WBGene00006483 | dgk-4 | 1.442149732 | 1.348263614 | 0.135449036 | 0.2 | 0.255 | 0.500729832 | 1 |
| WBGene00016235 | C29H12.2 | 1.246538713 | 1.152306542 | 0.135948286 | 0.2 | 0.226 | 0.919678333 | 1 |
| WBGene00004167 | pqn-87 | 0.92560081 | 0.830716912 | 0.13688853 | 0.115 | 0.126 | 0.974734934 | 1 |
| WBGene00015112 | B0281.5 | 0.536381277 | 0.441453354 | 0.136952043 | 0.062 | 0.046 | 0.469677017 | 1 |
| WBGene00001837 | hda-4 | 1.048846743 | 0.953031847 | 0.138231674 | 0.1 | 0.174 | 0.083625322 | 1 |
| WBGene00006699 | uba-1 | 0.747338944 | 0.651437949 | 0.13835589 | 0.092 | 0.099 | 0.984124046 | 1 |
| WBGene00016494 | C37A2.8 | 0.781756926 | 0.685696797 | 0.138585471 | 0.1 | 0.093 | 0.686737222 | 1 |
| WBGene00001989 | hot-4 | 1.035887467 | 0.93972368 | 0.138735018 | 0.131 | 0.161 | 0.651975032 | 1 |
| WBGene00020778 | T24E12.11 | 0.851824323 | 0.755339687 | 0.139197906 | 0.092 | 0.114 | 0.622085201 | 1 |
| WBGene00010097 | srpa-68 | 0.480703059 | 0.383882123 | 0.139683085 | 0.046 | 0.054 | 0.772983313 | 1 |
| WBGene00004345 | rgs-2 | 2.72581201 | 2.628634952 | 0.14019686 | 0.585 | 0.671 | 0.973695733 | 1 |
| WBGene00004138 | pqn-53 | 0.803636452 | 0.705804946 | 0.141141028 | 0.077 | 0.11 | 0.37670103 | 1 |
| WBGene00019482 | K07C11.10 | 1.486303541 | 1.388009702 | 0.141808034 | 0.254 | 0.29 | 0.907687195 | 1 |
| WBGene00009882 | vha-17 | 0.946739839 | 0.84834014 | 0.141960758 | 0.108 | 0.139 | 0.543128138 | 1 |
| WBGene00003637 | nhr-47 | 1.144200483 | 1.045596746 | 0.142255122 | 0.185 | 0.211 | 0.942812536 | 1 |
| WBGene00021187 | Y9D1A.2 | 2.100578808 | 2.001270673 | 0.143271354 | 0.362 | 0.404 | 0.625844379 | 1 |
| WBGene00000065 | act-3 | 0.658021104 | 0.558549848 | 0.143506688 | 0.062 | 0.087 | 0.425185122 | 1 |
| WBGene00001431 | fkb-6 | 1.300693658 | 1.200580796 | 0.14443233 | 0.2 | 0.207 | 0.783279812 | 1 |
| WBGene00007933 | C34E7.3 | 0.880762368 | 0.780273402 | 0.144974932 | 0.1 | 0.124 | 0.602789992 | 1 |
| WBGene00007683 | C18E9.2 | 1.100234494 | 0.999529035 | 0.145287267 | 0.138 | 0.132 | 0.661913524 | 1 |
| WBGene00003741 | nlp-3 | 4.546689259 | 4.445646719 | 0.145773571 | 1 | 0.938 | 0.058086327 | 1 |
| WBGene00017184 | ncam-1 | 0.520578642 | 0.417256097 | 0.149062923 | 0.046 | 0.062 | 0.559104147 | 1 |

| | | | | | | | | |
|---|---|---|---|---|---|---|---|---|
| WBGene00017659 | F21C10.10 | 0.549655237 | 0.445995017 | 0.149550085 | 0.062 | 0.07 | 0.848807349 | 1 |
| WBGene00002210 | kin-29 | 0.703541545 | 0.598880952 | 0.150993319 | 0.062 | 0.079 | 0.563744192 | 1 |
| WBGene00004933 | sod-4 | 1.549570935 | 1.444803477 | 0.151147492 | 0.215 | 0.284 | 0.44682837 | 1 |
| WBGene00010908 | imph-1 | 0.562559613 | 0.457761355 | 0.151191926 | 0.054 | 0.054 | 0.95088442 | 1 |
| WBGene00006306 | sue-1 | 1.015134883 | 0.910320359 | 0.151215394 | 0.131 | 0.135 | 0.929458917 | 1 |
| WBGene00010406 | math-33 | 0.558031118 | 0.452433438 | 0.152345248 | 0.062 | 0.054 | 0.686451844 | 1 |
| WBGene00013604 | Y87G2A.18 | 0.461134043 | 0.355219585 | 0.152802263 | 0.046 | 0.056 | 0.723205449 | 1 |
| WBGene00013638 | Y105C5A.14 | 0.488121392 | 0.382017026 | 0.153076243 | 0.046 | 0.05 | 0.913776711 | 1 |
| WBGene00021921 | rbm-39 | 0.779843824 | 0.673663674 | 0.153185577 | 0.1 | 0.104 | 0.953295595 | 1 |
| WBGene00010425 | lpin-1 | 1.073116956 | 0.965664698 | 0.15502084 | 0.138 | 0.149 | 0.957839452 | 1 |
| WBGene00008610 | F09C3.2 | 0.56795958 | 0.460366887 | 0.155223444 | 0.062 | 0.064 | 0.983636178 | 1 |
| WBGene00008613 | fbxb-69 | 1.122203404 | 1.014513104 | 0.155364261 | 0.154 | 0.172 | 0.942234149 | 1 |
| WBGene00007433 | swsn-7 | 0.872016441 | 0.76423139 | 0.155500958 | 0.1 | 0.132 | 0.512697182 | 1 |
| WBGene00018491 | mdh-1 | 0.596070667 | 0.487882057 | 0.156083171 | 0.062 | 0.07 | 0.835635581 | 1 |
| WBGene00012513 | Y26E6A.2 | 0.9455734 | 0.835940746 | 0.158166486 | 0.115 | 0.141 | 0.624280221 | 1 |
| WBGene00022473 | bet-1 | 0.829628322 | 0.719991483 | 0.158172524 | 0.1 | 0.104 | 0.984951891 | 1 |
| WBGene00004276 | rab-14 | 1.269162519 | 1.159314181 | 0.158477652 | 0.169 | 0.217 | 0.562420038 | 1 |
| WBGene00009717 | dep-1 | 0.971124425 | 0.860993994 | 0.158884627 | 0.092 | 0.145 | 0.191486027 | 1 |
| WBGene00010726 | K09E9.3 | 0.506534798 | 0.39580565 | 0.159748392 | 0.054 | 0.039 | 0.41589174 | 1 |
| WBGene00011867 | chc-1 | 0.906226776 | 0.795161117 | 0.160233876 | 0.1 | 0.137 | 0.405526579 | 1 |
| WBGene00011051 | pfs-2 | 0.54562567 | 0.433775559 | 0.161365601 | 0.062 | 0.044 | 0.374219641 | 1 |
| WBGene00020031 | suco-1 | 0.608883593 | 0.496742213 | 0.161785813 | 0.062 | 0.06 | 0.865103057 | 1 |
| WBGene00004732 | sax-7 | 0.571666764 | 0.459376239 | 0.162000983 | 0.054 | 0.066 | 0.69446268 | 1 |
| WBGene00006632 | tsp-6 | 0.955468995 | 0.841638234 | 0.164223075 | 0.115 | 0.133 | 0.809311829 | 1 |
| WBGene00009975 | F53C11.5 | 0.437266112 | 0.323216993 | 0.164538097 | 0.054 | 0.043 | 0.553195415 | 1 |
| WBGene00011299 | ikke-1 | 0.918924167 | 0.803754562 | 0.166154619 | 0.1 | 0.124 | 0.610945361 | 1 |
| WBGene00003636 | nhr-46 | 0.563980944 | 0.447272925 | 0.16837408 | 0.046 | 0.064 | 0.501247268 | 1 |
| WBGene00008586 | snrp-40.1 | 0.633859319 | 0.517047054 | 0.168524476 | 0.054 | 0.072 | 0.55119078 | 1 |
| WBGene00011481 | imp-2 | 0.551867705 | 0.434492427 | 0.169336731 | 0.054 | 0.064 | 0.753712417 | 1 |
| WBGene00022635 | ZC581.9 | 1.687964627 | 1.570547481 | 0.169397134 | 0.231 | 0.321 | 0.271793956 | 1 |
| WBGene00021050 | W05H9.3 | 2.57104727 | 2.452795501 | 0.170601241 | 0.454 | 0.528 | 0.983747645 | 1 |
| WBGene00018476 | F45E12.6 | 4.11129118 | 3.992542345 | 0.171318355 | 0.954 | 0.892 | 0.360848908 | 1 |
| WBGene00012602 | Y38E10A.24 | 0.840472162 | 0.72082443 | 0.172615189 | 0.085 | 0.083 | 0.805081821 | 1 |
| WBGene00010964 | ctc-1 | 2.658817438 | 2.53831408 | 0.173849598 | 0.615 | 0.578 | 0.050240075 | 1 |
| WBGene00006708 | ubc-13 | 0.462344119 | 0.340002204 | 0.176502074 | 0.054 | 0.05 | 0.814933494 | 1 |
| WBGene00004136 | pqn-51 | 0.728990887 | 0.606387463 | 0.176879352 | 0.077 | 0.095 | 0.658491206 | 1 |
| WBGene00000840 | cul-5 | 0.596941751 | 0.473071876 | 0.178706455 | 0.062 | 0.054 | 0.678694174 | 1 |
| WBGene00019196 | H14A12.3 | 0.508848431 | 0.384441729 | 0.179480933 | 0.046 | 0.058 | 0.668082847 | 1 |
| WBGene00004698 | rsp-1 | 1.290867121 | 1.166453245 | 0.179491283 | 0.131 | 0.213 | 0.091617552 | 1 |
| WBGene00012005 | 1-Jun | 1.444684376 | 1.320031892 | 0.17983552 | 0.208 | 0.253 | 0.666198999 | 1 |
| WBGene00014569 | bkip-1 | 0.77920652 | 0.654169126 | 0.180390828 | 0.069 | 0.108 | 0.253002749 | 1 |
| WBGene00008690 | F11C1.1 | 0.462250281 | 0.336977894 | 0.180729851 | 0.031 | 0.05 | 0.37911595 | 1 |
| WBGene00006731 | uev-2 | 0.750381917 | 0.624893394 | 0.181041669 | 0.085 | 0.101 | 0.752905638 | 1 |
| WBGene00007490 | mks-5 | 0.570777892 | 0.444420034 | 0.182295854 | 0.054 | 0.072 | 0.552798997 | 1 |
| WBGene00004014 | phb-1 | 0.564309576 | 0.437571368 | 0.182844584 | 0.054 | 0.041 | 0.472516263 | 1 |
| WBGene00012543 | nkcc-1 | 0.584599202 | 0.457752154 | 0.183001608 | 0.054 | 0.058 | 0.935542817 | 1 |
| WBGene00004804 | skn-1 | 1.013840366 | 0.886906218 | 0.183127266 | 0.115 | 0.128 | 0.966499129 | 1 |
| WBGene00018204 | F39F10.4 | 0.954230239 | 0.825180669 | 0.186179174 | 0.115 | 0.124 | 0.965028794 | 1 |
| WBGene00003568 | ncx-3 | 0.802882291 | 0.673601778 | 0.186512355 | 0.085 | 0.103 | 0.679677521 | 1 |
| WBGene00014772 | F56D5.4 | 1.166511334 | 1.037162627 | 0.186610739 | 0.162 | 0.17 | 0.827907265 | 1 |
| WBGene00006707 | ubc-12 | 0.496978442 | 0.366882414 | 0.187688893 | 0.038 | 0.054 | 0.521121837 | 1 |
| WBGene00003082 | lsm-8 | 0.581168595 | 0.449825977 | 0.189487344 | 0.062 | 0.05 | 0.54894032 | 1 |
| WBGene00008034 | prcc-1 | 0.812636591 | 0.678839615 | 0.193028234 | 0.085 | 0.093 | 0.917537881 | 1 |
| WBGene00008549 | din-1 | 0.913540114 | 0.779442299 | 0.193462251 | 0.123 | 0.101 | 0.361927648 | 1 |
| WBGene00005726 | srv-15 | 0.724588316 | 0.58939813 | 0.19503821 | 0.069 | 0.097 | 0.437909151 | 1 |
| WBGene00020710 | T23B3.2 | 1.866259837 | 1.730545602 | 0.195794255 | 0.308 | 0.391 | 0.712447426 | 1 |
| WBGene00195043 | Y57G11C.1135 | 0.604564867 | 0.468549253 | 0.196229051 | 0.069 | 0.074 | 0.980456183 | 1 |
| WBGene00009012 | F21D5.7 | 0.584235335 | 0.446702293 | 0.198418238 | 0.062 | 0.054 | 0.678694174 | 1 |
| WBGene00001172 | egl-3 | 1.819653117 | 1.68211136 | 0.198430811 | 0.3 | 0.385 | 0.735215374 | 1 |
| WBGene00017156 | F01E11.3 | 0.595016035 | 0.457368601 | 0.198583271 | 0.054 | 0.062 | 0.790272057 | 1 |
| WBGene00006537 | tbb-2 | 1.26714668 | 1.129395008 | 0.198733653 | 0.162 | 0.174 | 0.930903668 | 1 |
| WBGene00011287 | glb-24 | 1.065234803 | 0.927234141 | 0.199092871 | 0.131 | 0.174 | 0.470758125 | 1 |
| WBGene00015811 | C16A3.6 | 0.564963362 | 0.426604843 | 0.199609149 | 0.054 | 0.046 | 0.677719601 | 1 |
| WBGene00019995 | R10F2.4 | 2.280736031 | 2.141765361 | 0.200492297 | 0.431 | 0.485 | 0.692326437 | 1 |

| | | | | | | | | |
|---|---|---|---|---|---|---|---|---|
| WBGene00002129 | inx-7 | 0.82266806 | 0.682307301 | 0.202497771 | 0.092 | 0.11 | 0.716602961 | 1 |
| WBGene00002101 | ins-18 | 2.646335553 | 2.505004214 | 0.203898022 | 0.492 | 0.522 | 0.655043051 | 1 |
| WBGene00016977 | akap-1 | 1.081790521 | 0.940186208 | 0.20429184 | 0.123 | 0.147 | 0.742895899 | 1 |
| WBGene00000207 | asb-2 | 0.734183752 | 0.592076467 | 0.205017475 | 0.092 | 0.095 | 0.921233281 | 1 |
| WBGene00022812 | uvs-1 | 0.470585703 | 0.328069789 | 0.205607004 | 0.038 | 0.05 | 0.62963585 | 1 |
| WBGene00003964 | pdi-3 | 1.112874933 | 0.970177283 | 0.205869191 | 0.146 | 0.147 | 0.76569914 | 1 |
| WBGene00018968 | F56D2.8 | 0.614384009 | 0.471215158 | 0.206548991 | 0.077 | 0.07 | 0.669307728 | 1 |
| WBGene00015418 | pac-1 | 1.282252097 | 1.138586683 | 0.207265381 | 0.169 | 0.191 | 0.87004465 | 1 |
| WBGene00012037 | T26E3.4 | 0.552860724 | 0.408939763 | 0.207634058 | 0.054 | 0.05 | 0.814933494 | 1 |
| WBGene00019552 | K09C4.7 | 0.65822913 | 0.513906971 | 0.208212864 | 0.077 | 0.074 | 0.757744204 | 1 |
| WBGene00044077 | pacs-1 | 0.67033196 | 0.525761773 | 0.208570692 | 0.069 | 0.079 | 0.807870812 | 1 |
| WBGene00011629 | T08G11.1 | 0.530044726 | 0.385251093 | 0.208893056 | 0.054 | 0.058 | 0.935542817 | 1 |
| WBGene00011289 | R102.2 | 7.235796502 | 7.089558181 | 0.210977301 | 1 | 1 | 3.58E-06 | 0.02286638 |
| WBGene00020037 | mrps-6 | 0.571137068 | 0.424878634 | 0.211006318 | 0.062 | 0.066 | 0.964216034 | 1 |
| WBGene00003080 | lsm-6 | 0.529185456 | 0.382355217 | 0.211831258 | 0.046 | 0.058 | 0.671933972 | 1 |
| WBGene00004920 | snr-7 | 1.223080229 | 1.076182434 | 0.211928721 | 0.177 | 0.18 | 0.740163138 | 1 |
| WBGene00006528 | tba-1 | 1.087254897 | 0.938416094 | 0.214729003 | 0.138 | 0.161 | 0.886346133 | 1 |
| WBGene00007979 | imp-1 | 0.6262891 | 0.477230796 | 0.215045676 | 0.062 | 0.031 | 0.088801749 | 1 |
| WBGene00007646 | nkb-1 | 0.808317597 | 0.659026967 | 0.215380851 | 0.085 | 0.079 | 0.733742238 | 1 |
| WBGene00007886 | ethe-1 | 0.694289854 | 0.544798238 | 0.215670812 | 0.077 | 0.062 | 0.464948061 | 1 |
| WBGene00019185 | H10E21.5 | 1.299124004 | 1.149591414 | 0.215729926 | 0.169 | 0.226 | 0.449007666 | 1 |
| WBGene00010603 | nhr-198 | 0.912685583 | 0.762802833 | 0.2162351 | 0.108 | 0.103 | 0.723790145 | 1 |
| WBGene00017150 | EGAP5.1 | 2.059809899 | 1.908260153 | 0.218640067 | 0.369 | 0.412 | 0.807022223 | 1 |
| WBGene00020846 | T27A10.6 | 0.576642414 | 0.424959584 | 0.218832067 | 0.069 | 0.066 | 0.786776881 | 1 |
| WBGene00003901 | paa-1 | 0.881749659 | 0.729575895 | 0.219540334 | 0.077 | 0.106 | 0.402485727 | 1 |
| WBGene00021693 | Y48G8AR.2 | 0.645826398 | 0.492005915 | 0.221916048 | 0.069 | 0.072 | 0.981670261 | 1 |
| WBGene00010269 | F58G1.7 | 0.658761104 | 0.504401369 | 0.222694024 | 0.054 | 0.068 | 0.64088736 | 1 |
| WBGene00020092 | pcf-11 | 0.692834741 | 0.538149335 | 0.223163868 | 0.077 | 0.087 | 0.836217793 | 1 |
| WBGene00001008 | dlk-1 | 0.864648094 | 0.708918474 | 0.224670351 | 0.115 | 0.097 | 0.397415453 | 1 |
| WBGene00014159 | arrd-4 | 1.85598099 | 1.698523041 | 0.227163802 | 0.308 | 0.387 | 0.683327413 | 1 |
| WBGene00010756 | K10G9.2 | 0.770628633 | 0.612314438 | 0.228399105 | 0.085 | 0.106 | 0.617643961 | 1 |
| WBGene00007534 | fubl-1 | 1.398668864 | 1.239733436 | 0.229295354 | 0.208 | 0.209 | 0.622044314 | 1 |
| WBGene00019069 | lgc-30 | 0.617560536 | 0.458489529 | 0.229490952 | 0.062 | 0.074 | 0.744596187 | 1 |
| WBGene00003674 | nhr-84 | 1.473166103 | 1.313803413 | 0.229911763 | 0.215 | 0.273 | 0.570916807 | 1 |
| WBGene00015729 | C13A10.2 | 0.668458005 | 0.508471377 | 0.230811915 | 0.069 | 0.066 | 0.812134077 | 1 |
| WBGene00001505 | fut-1 | 0.527926364 | 0.367935253 | 0.230818382 | 0.054 | 0.052 | 0.878617554 | 1 |
| WBGene00001439 | fkh-7 | 1.490237838 | 1.330121229 | 0.230999438 | 0.215 | 0.261 | 0.714954792 | 1 |
| WBGene00010037 | F54C8.4 | 1.661002058 | 1.500197045 | 0.231992595 | 0.292 | 0.309 | 0.568261157 | 1 |
| WBGene00022696 | ift-139 | 0.887544639 | 0.72636005 | 0.232540207 | 0.085 | 0.122 | 0.345804593 | 1 |
| WBGene00006698 | uaf-2 | 0.793757328 | 0.632106176 | 0.233213315 | 0.085 | 0.103 | 0.706223737 | 1 |
| WBGene00003892 | osm-12 | 1.177842686 | 1.015852133 | 0.233702968 | 0.146 | 0.199 | 0.415991507 | 1 |
| WBGene00008444 | tbc-14 | 0.56074869 | 0.39872665 | 0.233748394 | 0.062 | 0.06 | 0.862101696 | 1 |
| WBGene00044742 | Y55F3BR.10 | 0.598169036 | 0.435907868 | 0.234093382 | 0.046 | 0.058 | 0.664240463 | 1 |
| WBGene00004739 | scd-1 | 0.614927001 | 0.450658849 | 0.236988847 | 0.054 | 0.056 | 0.988930299 | 1 |
| WBGene00015926 | C17H11.6 | 0.505989205 | 0.341638123 | 0.237108491 | 0.038 | 0.054 | 0.519334947 | 1 |
| WBGene00019128 | F59G1.4 | 0.890375383 | 0.725999483 | 0.237144296 | 0.1 | 0.095 | 0.727899589 | 1 |
| WBGene00010333 | copz-1 | 0.70976329 | 0.54529262 | 0.23728102 | 0.077 | 0.093 | 0.713188035 | 1 |
| WBGene00005015 | spt-5 | 0.554035185 | 0.389297938 | 0.237665609 | 0.054 | 0.041 | 0.474353818 | 1 |
| WBGene00010993 | vha-20 | 0.475841494 | 0.310270913 | 0.238867856 | 0.054 | 0.044 | 0.59999877 | 1 |
| WBGene00011723 | T12A7.2 | 0.882510221 | 0.716279127 | 0.239820775 | 0.1 | 0.112 | 0.873177866 | 1 |
| WBGene00007684 | C18E9.4 | 0.509191039 | 0.342423444 | 0.240594783 | 0.062 | 0.054 | 0.649910541 | 1 |
| WBGene00077761 | zip-9 | 0.517500319 | 0.349916107 | 0.241772913 | 0.062 | 0.037 | 0.191568524 | 1 |
| WBGene00015367 | srt-64 | 1.993435348 | 1.8257631 | 0.241899921 | 0.362 | 0.395 | 0.475653152 | 1 |
| WBGene00022347 | Y82E9BR.14 | 0.871032278 | 0.702572098 | 0.243036666 | 0.123 | 0.12 | 0.701812351 | 1 |
| WBGene00001122 | dyf-6 | 1.873549548 | 1.704708458 | 0.243586202 | 0.315 | 0.4 | 0.659016597 | 1 |
| WBGene00009584 | drap-1 | 2.09018881 | 1.921056459 | 0.244006404 | 0.385 | 0.435 | 0.631003404 | 1 |
| WBGene00016458 | C35E7.6 | 0.717119343 | 0.54771803 | 0.244394434 | 0.085 | 0.101 | 0.771181606 | 1 |
| WBGene00010096 | rskd-1 | 0.484347356 | 0.314064797 | 0.245665804 | 0.054 | 0.027 | 0.112962734 | 1 |
| WBGene00015501 | C06A5.3 | 0.759324091 | 0.588795411 | 0.246020881 | 0.085 | 0.072 | 0.524809081 | 1 |
| WBGene00004503 | rpt-3 | 0.625904125 | 0.455013541 | 0.246542998 | 0.038 | 0.056 | 0.466421403 | 1 |
| WBGene00011270 | R31.2 | 0.90139162 | 0.730381005 | 0.246716166 | 0.092 | 0.104 | 0.846660775 | 1 |
| WBGene00019322 | ahcy-1 | 0.944257622 | 0.771533783 | 0.249187827 | 0.123 | 0.112 | 0.574861938 | 1 |
| WBGene00022712 | ZK355.2 | 0.540728392 | 0.367890332 | 0.249352613 | 0.069 | 0.058 | 0.551481996 | 1 |
| WBGene00004980 | spk-1 | 0.580583002 | 0.407734433 | 0.249367773 | 0.054 | 0.068 | 0.655912846 | 1 |

| | | | | | | | | |
|---|---|---|---|---|---|---|---|---|
| WBGene00008587 | F08G12.3 | 0.799371763 | 0.625587735 | 0.250717356 | 0.077 | 0.081 | 0.991560493 | 1 |
| WBGene00019840 | R02F11.2 | 3.190621765 | 3.016626688 | 0.251021833 | 0.785 | 0.814 | 0.058400227 | 1 |
| WBGene00012625 | Y38H6C.13 | 0.654447687 | 0.480318032 | 0.25121599 | 0.069 | 0.048 | 0.291827614 | 1 |
| WBGene00012130 | slc-17.4 | 0.601708765 | 0.42641626 | 0.252893629 | 0.062 | 0.072 | 0.792051989 | 1 |
| WBGene00000910 | daf-14 | 1.323756364 | 1.148325671 | 0.25309299 | 0.208 | 0.199 | 0.497271725 | 1 |
| WBGene00000876 | cyl-1 | 0.687159141 | 0.509837718 | 0.255820738 | 0.069 | 0.07 | 0.923192847 | 1 |
| WBGene00008989 | F20G2.6 | 3.074649793 | 2.89671806 | 0.256701229 | 0.723 | 0.77 | 0.138812802 | 1 |
| WBGene00021960 | tmem-258 | 0.988048972 | 0.810012283 | 0.256852649 | 0.1 | 0.126 | 0.576955391 | 1 |
| WBGene00001017 | dnc-1 | 0.515968004 | 0.337686842 | 0.257205349 | 0.054 | 0.043 | 0.525142174 | 1 |
| WBGene00004800 | sir-2.1 | 0.526084525 | 0.347165335 | 0.258125828 | 0.038 | 0.052 | 0.581523943 | 1 |
| WBGene00009888 | F49E2.5 | 0.747791311 | 0.568700315 | 0.258373692 | 0.092 | 0.081 | 0.570057274 | 1 |
| WBGene00004751 | sea-2 | 1.408428288 | 1.228361948 | 0.259780815 | 0.215 | 0.242 | 0.909393637 | 1 |
| WBGene00004106 | pqn-15 | 1.157807693 | 0.977624542 | 0.259949338 | 0.123 | 0.166 | 0.413325646 | 1 |
| WBGene00015508 | mvb-12 | 0.409107911 | 0.228506176 | 0.260553227 | 0.054 | 0.031 | 0.19838098 | 1 |
| WBGene00016294 | C31H2.4 | 0.712283453 | 0.53127577 | 0.261138887 | 0.046 | 0.085 | 0.174237893 | 1 |
| WBGene00010427 | hpo-11 | 0.64228393 | 0.461000789 | 0.261536288 | 0.069 | 0.052 | 0.387858525 | 1 |
| WBGene00012897 | pisy-1 | 0.774389107 | 0.592670007 | 0.262165245 | 0.108 | 0.064 | 0.07150155 | 1 |
| WBGene00021097 | cdc-37 | 0.801030372 | 0.618869017 | 0.262803284 | 0.092 | 0.087 | 0.731397716 | 1 |
| WBGene00000567 | cnx-1 | 1.015801083 | 0.833154235 | 0.263503701 | 0.115 | 0.139 | 0.714985829 | 1 |
| WBGene00018519 | F46H5.3 | 0.938279653 | 0.755562456 | 0.263605194 | 0.123 | 0.106 | 0.440976689 | 1 |
| WBGene00010864 | xbx-3 | 3.368909382 | 3.18612731 | 0.26369879 | 0.869 | 0.814 | 0.085336961 | 1 |
| WBGene00016415 | ampd-1 | 0.682057823 | 0.499245845 | 0.263741934 | 0.077 | 0.072 | 0.739102246 | 1 |
| WBGene00001915 | his-41 | 3.219287644 | 3.035743813 | 0.264797775 | 0.769 | 0.743 | 0.052682206 | 1 |
| WBGene00022817 | upp-1 | 1.601937825 | 1.418080962 | 0.265249384 | 0.277 | 0.277 | 0.409781843 | 1 |
| WBGene00009774 | F46B6.9 | 0.681499742 | 0.497623262 | 0.265277685 | 0.069 | 0.06 | 0.605710064 | 1 |
| WBGene00010560 | eif-2beta | 0.974536513 | 0.790483483 | 0.265532395 | 0.138 | 0.095 | 0.090448969 | 1 |
| WBGene00003884 | osm-3 | 1.011840928 | 0.827522178 | 0.265915747 | 0.138 | 0.128 | 0.553064789 | 1 |
| WBGene00269432 | ZC132.11 | 1.394244649 | 1.209761493 | 0.266152934 | 0.162 | 0.242 | 0.173563753 | 1 |
| WBGene00018042 | mks-3 | 0.859801862 | 0.674001536 | 0.268053209 | 0.1 | 0.108 | 0.948441189 | 1 |
| WBGene00007645 | C17E4.6 | 0.796883121 | 0.610826728 | 0.268422635 | 0.108 | 0.095 | 0.510413849 | 1 |
| WBGene00020629 | T20F5.7 | 0.522081074 | 0.335280481 | 0.26949629 | 0.054 | 0.054 | 0.921067205 | 1 |
| WBGene00007752 | wdfy-3 | 0.657923792 | 0.470351597 | 0.270609476 | 0.062 | 0.077 | 0.622222934 | 1 |
| WBGene00018564 | F47D12.9 | 0.953459198 | 0.765398448 | 0.271314312 | 0.138 | 0.118 | 0.421420312 | 1 |
| WBGene00019900 | vdac-1 | 1.751526534 | 1.562331418 | 0.272950854 | 0.3 | 0.338 | 0.775089869 | 1 |
| WBGene00004862 | sma-9 | 1.196841676 | 1.00739556 | 0.273312972 | 0.162 | 0.184 | 0.895087137 | 1 |
| WBGene00007761 | rad-26 | 0.758183501 | 0.568679164 | 0.273396966 | 0.077 | 0.075 | 0.83547357 | 1 |
| WBGene00013219 | Y54G11A.11 | 0.96898324 | 0.778171132 | 0.275283681 | 0.123 | 0.124 | 0.777025614 | 1 |
| WBGene00002191 | kin-3 | 0.821368415 | 0.630426335 | 0.275471191 | 0.077 | 0.085 | 0.888599408 | 1 |
| WBGene00001910 | C50F4.6 | 1.635548614 | 1.44421767 | 0.276032204 | 0.277 | 0.236 | 0.127186807 | 1 |
| WBGene00000175 | aqp-7 | 0.667284727 | 0.475768604 | 0.276299361 | 0.069 | 0.044 | 0.220738285 | 1 |
| WBGene00008254 | C51E3.9 | 0.513544578 | 0.32168861 | 0.276789653 | 0.038 | 0.052 | 0.57483854 | 1 |
| WBGene00009931 | F52D10.4 | 0.970104114 | 0.77667653 | 0.279057016 | 0.1 | 0.114 | 0.783337446 | 1 |
| WBGene00016292 | tbc-7 | 0.626930187 | 0.433485388 | 0.279081853 | 0.062 | 0.056 | 0.751760808 | 1 |
| WBGene00219912 | F22H10.10 | 0.507273113 | 0.313746107 | 0.279200452 | 0.062 | 0.037 | 0.188750119 | 1 |
| WBGene00010260 | ddx-17 | 1.152670798 | 0.958681001 | 0.279868118 | 0.138 | 0.162 | 0.759135995 | 1 |
| WBGene00000415 | ced-1 | 0.619334439 | 0.424417907 | 0.281205114 | 0.046 | 0.07 | 0.391265307 | 1 |
| WBGene00002999 | lin-10 | 0.798217832 | 0.603184485 | 0.281373641 | 0.062 | 0.081 | 0.503940448 | 1 |
| WBGene00013111 | sta-1 | 1.330044964 | 1.134997444 | 0.28139409 | 0.208 | 0.201 | 0.44472316 | 1 |
| WBGene00006889 | pfd-3 | 0.566621128 | 0.37135474 | 0.281709849 | 0.062 | 0.043 | 0.334061089 | 1 |
| WBGene00014213 | ZK1073.1 | 1.386621301 | 1.191114478 | 0.282056724 | 0.192 | 0.224 | 0.857822135 | 1 |
| WBGene00044442 | T08A9.13 | 2.066464135 | 1.870884077 | 0.28216238 | 0.354 | 0.429 | 0.827534246 | 1 |
| WBGene00010074 | irld-34 | 1.983764245 | 1.787875366 | 0.282607913 | 0.377 | 0.358 | 0.118095332 | 1 |
| WBGene00000242 | bbs-2 | 1.079448013 | 0.88261501 | 0.283969997 | 0.123 | 0.137 | 0.931569869 | 1 |
| WBGene00004468 | rpn-12 | 0.510841888 | 0.31239765 | 0.286294517 | 0.054 | 0.043 | 0.533769875 | 1 |
| WBGene00006773 | unc-37 | 0.541131164 | 0.342660092 | 0.286333231 | 0.062 | 0.056 | 0.738926159 | 1 |
| WBGene00017914 | F29A7.4 | 0.607221792 | 0.40739504 | 0.288289064 | 0.077 | 0.05 | 0.202036462 | 1 |
| WBGene00001129 | dyf-13 | 1.344124362 | 1.144016398 | 0.288694768 | 0.177 | 0.217 | 0.657449028 | 1 |
| WBGene00045483 | oxy-5 | 0.942449073 | 0.740275935 | 0.291674183 | 0.115 | 0.12 | 0.89639648 | 1 |
| WBGene00016496 | C37C3.2 | 1.007664807 | 0.805356924 | 0.291868579 | 0.123 | 0.11 | 0.50874239 | 1 |
| WBGene00017973 | ift-81 | 1.89359255 | 1.690221694 | 0.293402127 | 0.338 | 0.34 | 0.28105028 | 1 |
| WBGene00021355 | Y37E3.17 | 0.808394988 | 0.604515215 | 0.294136337 | 0.092 | 0.097 | 0.964337935 | 1 |
| WBGene00044143 | K09E9.4 | 0.805325545 | 0.600938649 | 0.294867962 | 0.108 | 0.091 | 0.435290205 | 1 |
| WBGene00004095 | pqe-1 | 0.597191134 | 0.392226278 | 0.295701781 | 0.069 | 0.054 | 0.445417179 | 1 |
| WBGene00019323 | teg-4 | 0.58905775 | 0.383841389 | 0.296064626 | 0.054 | 0.037 | 0.344714921 | 1 |

| | | | | | | | | |
|---|---|---|---|---|---|---|---|---|
| WBGene00001336 | qars-1 | 1.470529987 | 1.265258489 | 0.296144173 | 0.208 | 0.261 | 0.643633119 | 1 |
| WBGene00016378 | imp-3 | 0.627610736 | 0.422275652 | 0.296235909 | 0.062 | 0.072 | 0.811376146 | 1 |
| WBGene00003826 | ntl-3 | 0.547383684 | 0.340993334 | 0.297758334 | 0.054 | 0.054 | 0.924257344 | 1 |
| WBGene00007227 | C01G6.5 | 0.887888534 | 0.681315941 | 0.298021255 | 0.1 | 0.089 | 0.563894502 | 1 |
| WBGene00235102 | Y54G2A.75 | 1.022696453 | 0.815926737 | 0.298305644 | 0.115 | 0.133 | 0.79541236 | 1 |
| WBGene00018152 | acs-4 | 0.686237993 | 0.479117251 | 0.298812067 | 0.069 | 0.046 | 0.244090182 | 1 |
| WBGene00014021 | svop-1 | 0.640357925 | 0.430541353 | 0.302701328 | 0.077 | 0.066 | 0.554409489 | 1 |
| WBGene00022069 | eel-1 | 1.032872463 | 0.822341927 | 0.303731361 | 0.1 | 0.126 | 0.589377816 | 1 |
| WBGene00016115 | mdt-26 | 2.50634702 | 2.295659371 | 0.303958027 | 0.562 | 0.526 | 0.055864668 | 1 |
| WBGene00003085 | ccar-1 | 0.917114173 | 0.705720449 | 0.304976678 | 0.108 | 0.112 | 0.936778928 | 1 |
| WBGene00000102 | akt-1 | 1.196049991 | 0.984572234 | 0.305097911 | 0.169 | 0.197 | 0.847290396 | 1 |
| WBGene00015205 | moa-2 | 0.703416747 | 0.490727118 | 0.306846272 | 0.085 | 0.06 | 0.262560035 | 1 |
| WBGene00006890 | vem-1 | 2.593513433 | 2.380390229 | 0.30747179 | 0.569 | 0.58 | 0.091950222 | 1 |
| WBGene00271808 | F21C10.19 | 0.864795846 | 0.650616556 | 0.308995399 | 0.077 | 0.099 | 0.578559504 | 1 |
| WBGene00010966 | nduo-3 | 0.857268435 | 0.642377627 | 0.310021903 | 0.092 | 0.097 | 0.92915549 | 1 |
| WBGene00044418 | C08G5.7 | 1.045437032 | 0.830208712 | 0.31050883 | 0.123 | 0.133 | 0.984661949 | 1 |
| WBGene00001070 | dpy-8 | 0.703712224 | 0.48706841 | 0.312550956 | 0.085 | 0.052 | 0.141449451 | 1 |
| WBGene00021931 | Y55F3AM.14 | 0.557336433 | 0.340281836 | 0.313143591 | 0.062 | 0.044 | 0.368150091 | 1 |
| WBGene00016558 | pks-1 | 0.658965594 | 0.441304533 | 0.314018532 | 0.069 | 0.054 | 0.430015214 | 1 |
| WBGene00004743 | scm-1 | 0.764206331 | 0.545963427 | 0.314857955 | 0.085 | 0.087 | 0.939941245 | 1 |
| WBGene00009804 | F47B8.3 | 1.310668242 | 1.091954901 | 0.315536652 | 0.177 | 0.219 | 0.724918295 | 1 |
| WBGene00020932 | dhod-1 | 0.480937548 | 0.262058985 | 0.315775018 | 0.069 | 0.039 | 0.113480558 | 1 |
| WBGene00019032 | srsx-17 | 1.05170589 | 0.8325879 | 0.316120437 | 0.169 | 0.135 | 0.207745063 | 1 |
| WBGene00019200 | H14E04.3 | 0.571610399 | 0.352026496 | 0.316792609 | 0.046 | 0.056 | 0.734271881 | 1 |
| WBGene00001137 | eat-6 | 1.426385777 | 1.205808235 | 0.318226126 | 0.215 | 0.222 | 0.699223799 | 1 |
| WBGene00009580 | xbx-6 | 1.910305013 | 1.689608561 | 0.318397678 | 0.308 | 0.344 | 0.711526405 | 1 |
| WBGene00017769 | hmgs-1 | 0.458913207 | 0.237751805 | 0.319068457 | 0.054 | 0.039 | 0.381201098 | 1 |
| WBGene00018164 | sup-36 | 0.951143972 | 0.729718589 | 0.319449301 | 0.123 | 0.128 | 0.848391236 | 1 |
| WBGene00017811 | F26A1.13 | 0.808997525 | 0.58747848 | 0.319584427 | 0.085 | 0.095 | 0.867855084 | 1 |
| WBGene00008348 | C56G7.3 | 0.501609083 | 0.279792071 | 0.320014304 | 0.054 | 0.031 | 0.195541252 | 1 |
| WBGene00019698 | riok-1 | 0.541091316 | 0.318627558 | 0.32094736 | 0.062 | 0.044 | 0.368150091 | 1 |
| WBGene00008643 | tmem-138 | 0.554586509 | 0.330775653 | 0.322890812 | 0.054 | 0.054 | 0.912566497 | 1 |
| WBGene00006059 | stc-1 | 0.535981957 | 0.312157396 | 0.322910585 | 0.054 | 0.025 | 0.081980084 | 1 |
| WBGene00013094 | Y51H1A.3 | 0.907738209 | 0.683845713 | 0.323008593 | 0.1 | 0.087 | 0.542458993 | 1 |
| WBGene00020852 | nhr-228 | 0.58161333 | 0.357491263 | 0.323339795 | 0.062 | 0.052 | 0.601511236 | 1 |
| WBGene00011288 | R102.1 | 4.83313741 | 4.608625046 | 0.323902875 | 0.877 | 0.778 | 0.009309723 | 1 |
| WBGene00022748 | oaz-1 | 2.326186747 | 2.10130673 | 0.324433285 | 0.531 | 0.507 | 0.041879815 | 1 |
| WBGene00009137 | bath-39 | 0.792479617 | 0.567541899 | 0.32451653 | 0.062 | 0.091 | 0.359422851 | 1 |
| WBGene00013020 | Y48G10A.3 | 0.698526321 | 0.47229674 | 0.326380295 | 0.077 | 0.064 | 0.521331805 | 1 |
| WBGene00003603 | nhr-4 | 1.019997156 | 0.792742834 | 0.327858683 | 0.123 | 0.124 | 0.826697907 | 1 |
| WBGene00023491 | tsp-21 | 1.935090483 | 1.707588261 | 0.328216327 | 0.385 | 0.381 | 0.261087005 | 1 |
| WBGene00304815 | Y40A1A.6 | 1.824283924 | 1.596757826 | 0.328250773 | 0.054 | 0.079 | 0.364791944 | 1 |
| WBGene00014058 | ZK673.2 | 0.611229502 | 0.383105202 | 0.329113796 | 0.062 | 0.046 | 0.419763649 | 1 |
| WBGene00011849 | T19H5.4 | 0.57190897 | 0.343575556 | 0.329415484 | 0.069 | 0.043 | 0.171534207 | 1 |
| WBGene00017982 | hpo-18 | 0.917037704 | 0.68748795 | 0.331170292 | 0.108 | 0.116 | 0.979917864 | 1 |
| WBGene00021926 | Y55F3AM.9 | 0.609423226 | 0.378485875 | 0.333172172 | 0.069 | 0.064 | 0.700899791 | 1 |
| WBGene00019466 | tos-1 | 3.956834855 | 3.723968332 | 0.335955378 | 0.923 | 0.845 | 0.003880036 | 1 |
| WBGene00020231 | T05A8.5 | 0.497486474 | 0.264214843 | 0.336539826 | 0.062 | 0.039 | 0.22184503 | 1 |
| WBGene00022750 | tres-1 | 0.668134939 | 0.434578149 | 0.336951223 | 0.062 | 0.041 | 0.273117083 | 1 |
| WBGene00006750 | unc-10 | 0.612751963 | 0.378826262 | 0.337483449 | 0.069 | 0.05 | 0.334317889 | 1 |
| WBGene00000063 | act-1 | 0.662878792 | 0.426226876 | 0.341416546 | 0.046 | 0.05 | 0.901596934 | 1 |
| WBGene00013703 | rbm-12 | 0.501894857 | 0.264675479 | 0.34223522 | 0.054 | 0.039 | 0.389494139 | 1 |
| WBGene00006434 | prdx-2 | 1.651230052 | 1.413847941 | 0.342469994 | 0.285 | 0.277 | 0.224083534 | 1 |
| WBGene00003904 | pabp-2 | 0.669383058 | 0.431065419 | 0.343819676 | 0.069 | 0.068 | 0.825318376 | 1 |
| WBGene00010550 | K04B12.2 | 0.440527885 | 0.202086764 | 0.343997823 | 0.054 | 0.027 | 0.114918293 | 1 |
| WBGene00021469 | Y39G10AR.11 | 0.699119097 | 0.460574359 | 0.344147311 | 0.069 | 0.058 | 0.548085819 | 1 |
| WBGene00000508 | cit-1.2 | 0.631051702 | 0.392506381 | 0.344148152 | 0.054 | 0.058 | 0.939693143 | 1 |
| WBGene00016293 | C31H2.3 | 1.933905316 | 1.695240111 | 0.344321108 | 0.308 | 0.319 | 0.618044419 | 1 |
| WBGene00013298 | Y57G11B.6 | 0.975819249 | 0.736687813 | 0.344993736 | 0.123 | 0.124 | 0.781603206 | 1 |
| WBGene00002980 | lgg-1 | 3.53093268 | 3.290772279 | 0.34647822 | 0.9 | 0.847 | 0.000702201 | 1 |
| WBGene00001561 | gei-4 | 1.096402203 | 0.85565063 | 0.347331101 | 0.138 | 0.133 | 0.613220764 | 1 |
| WBGene00004220 | ptr-5 | 0.999210259 | 0.75788969 | 0.348151987 | 0.131 | 0.122 | 0.547043802 | 1 |
| WBGene00003406 | mrg-1 | 0.715873088 | 0.474368569 | 0.348417373 | 0.077 | 0.066 | 0.56006422 | 1 |
| WBGene00002113 | ins-30 | 1.520159185 | 1.278429874 | 0.348741679 | 0.215 | 0.162 | 0.055248892 | 1 |

| | | | | | | | | |
|---|---|---|---|---|---|---|---|---|
| WBGene00013340 | tcc-1 | 0.583660212 | 0.341647251 | 0.349150898 | 0.069 | 0.027 | 0.017893687 | 1 |
| WBGene00007980 | C36E8.1 | 0.738858477 | 0.496609344 | 0.349491623 | 0.077 | 0.06 | 0.405280881 | 1 |
| WBGene00000829 | ctb-1 | 2.013244525 | 1.769812718 | 0.351197861 | 0.354 | 0.364 | 0.271086724 | 1 |
| WBGene00045171 | K09E9.5 | 0.484363938 | 0.240828413 | 0.351347494 | 0.054 | 0.031 | 0.188297514 | 1 |
| WBGene00019402 | syf-2 | 0.62302065 | 0.377484038 | 0.354234451 | 0.069 | 0.052 | 0.387858558 | 1 |
| WBGene00006753 | unc-14 | 1.240106186 | 0.993558946 | 0.355692481 | 0.154 | 0.153 | 0.654832791 | 1 |
| WBGene00019870 | R04E5.7 | 1.033341832 | 0.786025547 | 0.356801978 | 0.131 | 0.124 | 0.595468687 | 1 |
| WBGene00017536 | F17A9.4 | 1.020902233 | 0.773576652 | 0.356815389 | 0.138 | 0.128 | 0.491740957 | 1 |
| WBGene00004783 | seu-1 | 0.792096097 | 0.544760121 | 0.356830386 | 0.092 | 0.083 | 0.607356852 | 1 |
| WBGene00006798 | unc-64 | 1.114145215 | 0.866226915 | 0.357670502 | 0.146 | 0.128 | 0.379904923 | 1 |
| WBGene00000424 | ced-10 | 0.570060201 | 0.321709848 | 0.358293823 | 0.054 | 0.025 | 0.08865298 | 1 |
| WBGene00006003 | srx-112 | 0.415335973 | 0.16647155 | 0.359035469 | 0.054 | 0.023 | 0.057818588 | 1 |
| WBGene00050939 | C05G5.7 | 2.13361499 | 1.884030536 | 0.360074254 | 0.331 | 0.29 | 0.160780231 | 1 |
| WBGene00011302 | R166.2 | 0.719210409 | 0.468627843 | 0.361514226 | 0.085 | 0.068 | 0.435755475 | 1 |
| WBGene00009850 | F48F7.5 | 0.709204305 | 0.458293534 | 0.361987724 | 0.069 | 0.075 | 0.93141549 | 1 |
| WBGene00014079 | ZK792.7 | 0.836555768 | 0.584624312 | 0.363460263 | 0.123 | 0.087 | 0.140930849 | 1 |
| WBGene00000834 | cua-1 | 0.67367849 | 0.421272423 | 0.364144982 | 0.069 | 0.029 | 0.028646925 | 1 |
| WBGene00006713 | ubc-18 | 0.915862377 | 0.663415312 | 0.364204128 | 0.108 | 0.106 | 0.754254377 | 1 |
| WBGene00017676 | F21F3.7 | 0.878218987 | 0.625593965 | 0.364460867 | 0.1 | 0.104 | 0.902032937 | 1 |
| WBGene00219609 | linc-7 | 4.467611626 | 4.214604205 | 0.365012551 | 0.631 | 0.615 | 0.250038343 | 1 |
| WBGene00001262 | emb-8 | 1.499591354 | 1.246559621 | 0.365047627 | 0.223 | 0.219 | 0.453808672 | 1 |
| WBGene00001127 | dyf-11 | 0.859017403 | 0.605107899 | 0.366313983 | 0.1 | 0.089 | 0.566738116 | 1 |
| WBGene00012434 | Y11D7A.10 | 0.640818531 | 0.38689626 | 0.3663324 | 0.077 | 0.041 | 0.069775954 | 1 |
| WBGene00022465 | ift-20 | 2.084535041 | 1.830401424 | 0.36663731 | 0.431 | 0.453 | 0.183263312 | 1 |
| WBGene00018946 | F56C3.9 | 0.593819396 | 0.339654835 | 0.366681952 | 0.069 | 0.046 | 0.263562344 | 1 |
| WBGene00009563 | F39H2.3 | 0.480262538 | 0.225752494 | 0.367180378 | 0.054 | 0.029 | 0.150261691 | 1 |
| WBGene00009294 | F31D4.9 | 0.526991163 | 0.272312494 | 0.367423653 | 0.054 | 0.044 | 0.578637132 | 1 |
| WBGene00044771 | tmem-218 | 0.603566443 | 0.348668319 | 0.36774026 | 0.062 | 0.056 | 0.714450216 | 1 |
| WBGene00004321 | rcan-1 | 1.190262701 | 0.935307291 | 0.367822905 | 0.177 | 0.135 | 0.12052649 | 1 |
| WBGene00012412 | Y7A5A.6 | 0.675022849 | 0.419396293 | 0.368791166 | 0.069 | 0.07 | 0.887361784 | 1 |
| WBGene00004947 | sos-1 | 0.498056785 | 0.241246583 | 0.370498804 | 0.054 | 0.031 | 0.187200848 | 1 |
| WBGene00010236 | F58B4.3 | 3.928262604 | 3.670754935 | 0.371505037 | 0.954 | 0.901 | 4.86E-06 | 0.030985415 |
| WBGene00019808 | R01B10.6 | 0.782622914 | 0.524008446 | 0.37310181 | 0.092 | 0.087 | 0.702680786 | 1 |
| WBGene00014036 | ZK643.5 | 0.939069598 | 0.67881869 | 0.375462695 | 0.123 | 0.104 | 0.358349006 | 1 |
| WBGene00011983 | fbxa-81 | 0.510576366 | 0.248231196 | 0.378484076 | 0.054 | 0.027 | 0.116104351 | 1 |
| WBGene00018586 | ubxn-3 | 0.567340214 | 0.303144322 | 0.381154104 | 0.062 | 0.046 | 0.415749882 | 1 |
| WBGene00021920 | mrps-25 | 1.40581083 | 1.14142468 | 0.381428588 | 0.192 | 0.203 | 0.793213528 | 1 |
| WBGene00011763 | T14B1.1 | 1.76149188 | 1.496988011 | 0.38159842 | 0.285 | 0.267 | 0.178217195 | 1 |
| WBGene00001684 | gpd-2 | 1.560448778 | 1.295375832 | 0.382419425 | 0.231 | 0.238 | 0.59666433 | 1 |
| WBGene00008000 | slc-17.2 | 1.044118095 | 0.778657575 | 0.382978576 | 0.138 | 0.11 | 0.25008612 | 1 |
| WBGene00016397 | maph-9 | 1.023889667 | 0.757836781 | 0.383833179 | 0.131 | 0.116 | 0.447362329 | 1 |
| WBGene00003660 | nhr-70 | 0.509669676 | 0.24264631 | 0.385233286 | 0.054 | 0.031 | 0.185564821 | 1 |
| WBGene00011735 | hip-1 | 0.722672041 | 0.454074173 | 0.387504812 | 0.069 | 0.07 | 0.889242546 | 1 |
| WBGene00011500 | T05G5.5 | 2.000082731 | 1.731334496 | 0.387721746 | 0.408 | 0.387 | 0.110257159 | 1 |
| WBGene00000249 | bir-1 | 0.566469943 | 0.297422184 | 0.388153868 | 0.054 | 0.044 | 0.582584234 | 1 |
| WBGene00004109 | stam-1 | 0.692816762 | 0.423406478 | 0.388676881 | 0.054 | 0.062 | 0.813885242 | 1 |
| WBGene00003885 | osm-5 | 1.432647448 | 1.162158825 | 0.390232594 | 0.223 | 0.221 | 0.461809139 | 1 |
| WBGene00022019 | sut-2 | 0.603257763 | 0.331834954 | 0.39158034 | 0.054 | 0.048 | 0.72112046 | 1 |
| WBGene00000490 | che-11 | 1.348759827 | 1.075773855 | 0.393835509 | 0.192 | 0.176 | 0.339439931 | 1 |
| WBGene00003218 | mep-1 | 1.080520255 | 0.806534637 | 0.395277692 | 0.138 | 0.128 | 0.486378364 | 1 |
| WBGene00000509 | cka-1 | 0.925285614 | 0.650820907 | 0.395968872 | 0.123 | 0.106 | 0.405149678 | 1 |
| WBGene00001145 | eat-16 | 1.44803152 | 1.173111942 | 0.396625111 | 0.215 | 0.221 | 0.55301477 | 1 |
| WBGene00001089 | dre-1 | 0.407986157 | 0.132897643 | 0.396868836 | 0.054 | 0.015 | 0.009008982 | 1 |
| WBGene00003954 | pcm-1 | 0.9881905 | 0.711757555 | 0.398808439 | 0.115 | 0.112 | 0.702992331 | 1 |
| WBGene00007883 | C33A12.4 | 5.197146631 | 4.920412688 | 0.399242687 | 0.992 | 0.985 | 6.58E-06 | 0.041975633 |
| WBGene00012351 | W09C5.1 | 0.609126269 | 0.329023294 | 0.404103172 | 0.062 | 0.043 | 0.314268951 | 1 |
| WBGene00004776 | ser-1 | 0.56936115 | 0.287817984 | 0.406180929 | 0.054 | 0.025 | 0.086381667 | 1 |
| WBGene00235287 | Y55F3C.17 | 0.6453043 | 0.363384904 | 0.406723715 | 0.069 | 0.054 | 0.440763875 | 1 |
| WBGene00020447 | mob-3 | 0.856069072 | 0.5738649 | 0.40713456 | 0.1 | 0.089 | 0.545580581 | 1 |
| WBGene00009973 | F53C11.3 | 1.132985387 | 0.850728034 | 0.407211284 | 0.154 | 0.112 | 0.10660454 | 1 |
| WBGene00000170 | aqp-2 | 1.794633466 | 1.512162778 | 0.407519062 | 0.338 | 0.308 | 0.066363896 | 1 |
| WBGene00012764 | Y41E3.6 | 2.829726348 | 2.546742555 | 0.408259313 | 0.385 | 0.325 | 0.103930529 | 1 |
| WBGene00013538 | ari-1.4 | 0.648284062 | 0.364299897 | 0.409702547 | 0.062 | 0.037 | 0.187225733 | 1 |
| WBGene00013272 | aqp-12 | 1.234069203 | 0.949672451 | 0.410297784 | 0.138 | 0.162 | 0.837709519 | 1 |

| | | | | | | | | |
|---|---|---|---|---|---|---|---|---|
| WBGene00012760 | Y41C4A.17 | 4.644270366 | 4.359398578 | 0.410983115 | 0.885 | 0.778 | 0.000980047 | 1 |
| WBGene00012413 | Y7A5A.7 | 1.893550107 | 1.607890093 | 0.412120286 | 0.346 | 0.286 | 0.007922642 | 1 |
| WBGene00005182 | srg-25 | 0.567749631 | 0.281053257 | 0.413615437 | 0.054 | 0.037 | 0.333029189 | 1 |
| WBGene00001494 | frm-8 | 1.121393662 | 0.834492147 | 0.413911393 | 0.138 | 0.157 | 0.904285825 | 1 |
| WBGene00001946 | his-72 | 2.378633637 | 2.090994954 | 0.414974901 | 0.423 | 0.491 | 0.627703503 | 1 |
| WBGene00016025 | xbx-4 | 0.810380363 | 0.522675089 | 0.415070973 | 0.092 | 0.075 | 0.40929451 | 1 |
| WBGene00012332 | rpi-1 | 1.164528438 | 0.875779398 | 0.416576808 | 0.154 | 0.168 | 0.895113761 | 1 |
| WBGene00006438 | nrfl-1 | 0.723913124 | 0.434829035 | 0.417060182 | 0.085 | 0.066 | 0.358461422 | 1 |
| WBGene00004378 | rme-8 | 0.630475294 | 0.340710933 | 0.418041607 | 0.062 | 0.052 | 0.588986287 | 1 |
| WBGene00012290 | W05H12.2 | 0.775845989 | 0.485817785 | 0.418422252 | 0.085 | 0.074 | 0.537388625 | 1 |
| WBGene00009403 | F35C11.4 | 0.864719993 | 0.574494246 | 0.418707245 | 0.1 | 0.075 | 0.248472869 | 1 |
| WBGene00021461 | nekl-1 | 0.801207288 | 0.510724527 | 0.419078039 | 0.085 | 0.072 | 0.492949956 | 1 |
| WBGene00021906 | Y55B1AR.2 | 0.555275058 | 0.264745646 | 0.419145342 | 0.054 | 0.027 | 0.111033496 | 1 |
| WBGene00006864 | npr-16 | 0.893986549 | 0.603033757 | 0.419756149 | 0.1 | 0.074 | 0.255119629 | 1 |
| WBGene00023414 | R01H2.7 | 0.51512879 | 0.223340392 | 0.420961675 | 0.054 | 0.037 | 0.327664407 | 1 |
| WBGene00006590 | toc-1 | 0.561433296 | 0.268644117 | 0.422405497 | 0.054 | 0.039 | 0.392841854 | 1 |
| WBGene00004437 | rpl-24.2 | 0.625066587 | 0.332213588 | 0.422497569 | 0.054 | 0.043 | 0.531846445 | 1 |
| WBGene00006802 | unc-69 | 1.268502341 | 0.975451488 | 0.422783012 | 0.177 | 0.161 | 0.355487049 | 1 |
| WBGene00001604 | gln-3 | 2.218682983 | 1.925038438 | 0.42363953 | 0.438 | 0.453 | 0.281376698 | 1 |
| WBGene00021312 | szy-2 | 0.621428757 | 0.326022154 | 0.42618164 | 0.069 | 0.043 | 0.170647136 | 1 |
| WBGene00008208 | nhr-167 | 1.938314619 | 1.642612028 | 0.426608661 | 0.338 | 0.36 | 0.488599147 | 1 |
| WBGene00016037 | dop-6 | 1.232019662 | 0.935549492 | 0.427716044 | 0.177 | 0.166 | 0.425168861 | 1 |
| WBGene00022500 | lfi-1 | 1.433949772 | 1.137057883 | 0.428324455 | 0.269 | 0.182 | 0.007203152 | 1 |
| WBGene00304812 | ZK380.6 | 4.488367997 | 4.187756218 | 0.433691122 | 0.692 | 0.609 | 0.039342505 | 1 |
| WBGene00011725 | lips-5 | 1.009256098 | 0.706785487 | 0.436372851 | 0.1 | 0.091 | 0.619879377 | 1 |
| WBGene00019166 | tat-2 | 1.339279176 | 1.03612782 | 0.437354958 | 0.138 | 0.153 | 0.981456551 | 1 |
| WBGene00010974 | bbs-5 | 1.734152189 | 1.430609039 | 0.437920197 | 0.292 | 0.302 | 0.33078615 | 1 |
| WBGene00017290 | F09E5.12 | 0.871619261 | 0.567251319 | 0.43911012 | 0.123 | 0.072 | 0.03803341 | 1 |
| WBGene00003129 | map-1 | 0.886597885 | 0.58212539 | 0.439260959 | 0.092 | 0.077 | 0.505164596 | 1 |
| WBGene00003635 | nhr-45 | 0.856367872 | 0.551705074 | 0.439535509 | 0.1 | 0.062 | 0.095398786 | 1 |
| WBGene00007495 | C09G9.5 | 1.174647604 | 0.867974213 | 0.442436181 | 0.138 | 0.137 | 0.68771917 | 1 |
| WBGene00004459 | rpn-2 | 0.646919642 | 0.340182269 | 0.442528487 | 0.069 | 0.05 | 0.329646337 | 1 |
| WBGene00021474 | dot-1.1 | 0.604970528 | 0.298000243 | 0.442864507 | 0.062 | 0.05 | 0.518600108 | 1 |
| WBGene00003148 | mbf-1 | 1.309969407 | 1.000990912 | 0.445761743 | 0.185 | 0.193 | 0.725191426 | 1 |
| WBGene00011232 | pck-2 | 0.65549424 | 0.345232659 | 0.447612845 | 0.077 | 0.05 | 0.185722324 | 1 |
| WBGene00022568 | ZC239.5 | 1.157596402 | 0.846328297 | 0.449064952 | 0.131 | 0.132 | 0.806455179 | 1 |
| WBGene00271642 | R106.8 | 1.770643376 | 1.457509967 | 0.451756016 | 0.308 | 0.317 | 0.3385094 | 1 |
| WBGene00016162 | crh-2 | 1.322460284 | 1.00810932 | 0.453512577 | 0.208 | 0.191 | 0.318566402 | 1 |
| WBGene00010632 | K07C5.9 | 2.803952089 | 2.489599434 | 0.453515016 | 0.723 | 0.598 | 1.00E-04 | 0.637941026 |
| WBGene00010840 | M03C11.3 | 0.652408699 | 0.336446998 | 0.455836379 | 0.069 | 0.033 | 0.051090789 | 1 |
| WBGene00001452 | flp-9 | 0.593237251 | 0.277084473 | 0.456112045 | 0.062 | 0.014 | 0.001216683 | 1 |
| WBGene00010375 | H09F14.1 | 0.523461045 | 0.207136035 | 0.456360523 | 0.062 | 0.015 | 0.002499206 | 1 |
| WBGene00003391 | mog-3 | 0.882741476 | 0.565669138 | 0.45743869 | 0.1 | 0.077 | 0.316778733 | 1 |
| WBGene00015527 | C06E2.1 | 0.932418261 | 0.612258672 | 0.461892651 | 0.123 | 0.089 | 0.161233751 | 1 |
| WBGene00007630 | har-1 | 1.158452514 | 0.837758221 | 0.462664067 | 0.131 | 0.135 | 0.792611384 | 1 |
| WBGene00019759 | M03F4.6 | 0.845514451 | 0.5241523 | 0.463627582 | 0.108 | 0.064 | 0.057628037 | 1 |
| WBGene00022419 | Y102A11A.9 | 1.3845632 | 1.063088795 | 0.46378953 | 0.169 | 0.168 | 0.579349463 | 1 |
| WBGene00004367 | ric-8 | 0.852140032 | 0.530518242 | 0.464002161 | 0.092 | 0.068 | 0.259308023 | 1 |
| WBGene00044313 | M02D8.7 | 0.490929886 | 0.168819113 | 0.464707615 | 0.054 | 0.023 | 0.057582147 | 1 |
| WBGene00006720 | ubc-25 | 0.848038681 | 0.525849236 | 0.464821115 | 0.1 | 0.066 | 0.134332641 | 1 |
| WBGene00013295 | Y57G11B.2 | 0.612430824 | 0.290208722 | 0.464868229 | 0.069 | 0.043 | 0.171979034 | 1 |
| WBGene00018696 | pho-6 | 1.438540894 | 1.116255743 | 0.464959189 | 0.138 | 0.132 | 0.654190114 | 1 |
| WBGene00020854 | T27C4.1 | 2.149211288 | 1.826213006 | 0.465988021 | 0.423 | 0.402 | 0.046598189 | 1 |
| WBGene00045495 | F11C1.7 | 0.799313009 | 0.476239531 | 0.466096504 | 0.092 | 0.066 | 0.233821148 | 1 |
| WBGene00018511 | F46F11.8 | 1.027228491 | 0.70413404 | 0.466126763 | 0.115 | 0.099 | 0.417365631 | 1 |
| WBGene00019365 | rei-2 | 1.26215061 | 0.938054741 | 0.467571502 | 0.192 | 0.141 | 0.044336701 | 1 |
| WBGene00000221 | atf-4 | 1.662530426 | 1.338122262 | 0.46802205 | 0.262 | 0.292 | 0.672590745 | 1 |
| WBGene00012359 | W09D10.1 | 0.725978436 | 0.400886367 | 0.469008716 | 0.077 | 0.062 | 0.432927949 | 1 |
| WBGene00001119 | dyf-3 | 1.381604186 | 1.056260222 | 0.469372124 | 0.169 | 0.191 | 0.975682325 | 1 |
| WBGene00003653 | nhr-63 | 0.657707354 | 0.330650857 | 0.471842786 | 0.062 | 0.056 | 0.702803524 | 1 |
| WBGene00012193 | vps-25 | 2.137225982 | 1.808273266 | 0.474578452 | 0.4 | 0.441 | 0.386704044 | 1 |
| WBGene00003640 | nhr-50 | 1.065749411 | 0.734810124 | 0.477444467 | 0.108 | 0.112 | 0.909292597 | 1 |
| WBGene00021328 | npr-23 | 0.64605645 | 0.314809362 | 0.477888531 | 0.069 | 0.027 | 0.017041522 | 1 |
| WBGene00010579 | K05C4.2 | 0.675568712 | 0.343948314 | 0.478427104 | 0.077 | 0.05 | 0.191497375 | 1 |

| | | | | | | | | |
|---|---|---|---|---|---|---|---|---|
| WBGene00007009 | wwp-1 | 0.879288521 | 0.547587845 | 0.478542921 | 0.108 | 0.077 | 0.192635315 | 1 |
| WBGene00018756 | plpp-1.1 | 0.773739777 | 0.440816298 | 0.480307053 | 0.085 | 0.054 | 0.158552762 | 1 |
| WBGene00009558 | F39B2.8 | 0.594580396 | 0.260710903 | 0.481671861 | 0.054 | 0.029 | 0.147906199 | 1 |
| WBGene00010153 | srxa-2 | 1.646596179 | 1.308482372 | 0.487795114 | 0.277 | 0.209 | 0.025517305 | 1 |
| WBGene00016931 | C54G6.2 | 0.522481366 | 0.183930487 | 0.488425673 | 0.062 | 0.027 | 0.045463363 | 1 |
| WBGene00000552 | cmd-1 | 2.598593089 | 2.256659065 | 0.493306521 | 0.585 | 0.571 | 0.00893025 | 1 |
| WBGene00015208 | B0495.9 | 0.962717694 | 0.620464148 | 0.493767494 | 0.092 | 0.103 | 0.913062239 | 1 |
| WBGene00007129 | B0272.3 | 0.885919163 | 0.543308038 | 0.494283371 | 0.108 | 0.081 | 0.237063695 | 1 |
| WBGene00013228 | Y56A3A.7 | 0.774634236 | 0.43134278 | 0.495264881 | 0.077 | 0.072 | 0.703299801 | 1 |
| WBGene00007848 | cytb-5.1 | 1.352832493 | 1.007808355 | 0.497764612 | 0.185 | 0.157 | 0.243788026 | 1 |
| WBGene00044007 | ZK1320.13 | 0.860387647 | 0.515146761 | 0.498077315 | 0.1 | 0.066 | 0.146736682 | 1 |
| WBGene00011511 | gpcp-2 | 0.892838939 | 0.547597557 | 0.498078029 | 0.108 | 0.085 | 0.315528641 | 1 |
| WBGene00011279 | asd-1 | 1.405961231 | 1.060001976 | 0.499113701 | 0.223 | 0.186 | 0.126586384 | 1 |
| WBGene00014202 | mmcm-1 | 0.516528327 | 0.170531463 | 0.49916796 | 0.054 | 0.012 | 0.002042513 | 1 |
| WBGene00000369 | ccf-1 | 0.554869583 | 0.206347373 | 0.502811264 | 0.054 | 0.025 | 0.082597513 | 1 |
| WBGene00008023 | C39B10.3 | 0.921525054 | 0.5722579 | 0.503885991 | 0.131 | 0.081 | 0.05220685 | 1 |
| WBGene00017016 | snap-1 | 1.565422785 | 1.215390066 | 0.504990468 | 0.246 | 0.226 | 0.217574709 | 1 |
| WBGene00009164 | hrdl-1 | 1.113058581 | 0.762480335 | 0.505777496 | 0.146 | 0.11 | 0.137991057 | 1 |
| WBGene00003393 | mog-5 | 0.57760207 | 0.225879127 | 0.507428945 | 0.054 | 0.029 | 0.146506509 | 1 |
| WBGene00019743 | nhr-203 | 0.615554255 | 0.26318734 | 0.508358001 | 0.069 | 0.041 | 0.139954149 | 1 |
| WBGene00017472 | cst-1 | 0.690921309 | 0.335273585 | 0.513091207 | 0.077 | 0.043 | 0.088422903 | 1 |
| WBGene00018266 | nhr-183 | 0.641309474 | 0.285571835 | 0.513220928 | 0.069 | 0.041 | 0.133497072 | 1 |
| WBGene00011990 | T24D5.2 | 2.009760741 | 1.652749298 | 0.515058638 | 0.338 | 0.369 | 0.466985435 | 1 |
| WBGene00016622 | mafr-1 | 0.54208451 | 0.184315055 | 0.516152218 | 0.069 | 0.025 | 0.011396139 | 1 |
| WBGene00013920 | pssy-1 | 0.534347536 | 0.176118327 | 0.516815503 | 0.054 | 0.023 | 0.058772404 | 1 |
| WBGene00008265 | kcnl-4 | 0.738572412 | 0.380312784 | 0.516859388 | 0.092 | 0.05 | 0.053095149 | 1 |
| WBGene00021956 | Y57E12AL.1 | 0.882191264 | 0.523162816 | 0.517968561 | 0.1 | 0.075 | 0.267441933 | 1 |
| WBGene00011624 | T08G5.3 | 0.639486808 | 0.279760221 | 0.518975763 | 0.077 | 0.033 | 0.021416068 | 1 |
| WBGene00044026 | cisd-1 | 0.900845869 | 0.539609247 | 0.521154283 | 0.108 | 0.093 | 0.439868516 | 1 |
| WBGene00018252 | F40H3.6 | 0.798981523 | 0.436602946 | 0.522801777 | 0.092 | 0.062 | 0.163857009 | 1 |
| WBGene00009562 | flp-22 | 2.779238873 | 2.415590743 | 0.524633353 | 0.615 | 0.602 | 0.019133557 | 1 |
| WBGene00007107 | pfd-4 | 0.942509835 | 0.577966617 | 0.525924693 | 0.092 | 0.077 | 0.446640969 | 1 |
| WBGene00022733 | tmem-17 | 0.90835113 | 0.542968817 | 0.527135251 | 0.092 | 0.075 | 0.428035831 | 1 |
| WBGene00018886 | npr-28 | 1.059374721 | 0.693832391 | 0.527366106 | 0.131 | 0.114 | 0.405266904 | 1 |
| WBGene00022280 | Y74C10AL.2 | 0.557088066 | 0.191154956 | 0.527929884 | 0.054 | 0.029 | 0.146971942 | 1 |
| WBGene00009534 | F38B7.3 | 3.09354396 | 2.72717945 | 0.528552263 | 0.731 | 0.692 | 0.000102062 | 0.651255424 |
| WBGene00019431 | K06A5.3 | 0.692180965 | 0.325283878 | 0.529320608 | 0.077 | 0.05 | 0.197863171 | 1 |
| WBGene00001909 | his-35 | 2.212768941 | 1.845783561 | 0.529447988 | 0.438 | 0.418 | 0.075603198 | 1 |
| WBGene00021876 | Y54G2A.11 | 0.689049483 | 0.321565866 | 0.530166792 | 0.085 | 0.046 | 0.066382219 | 1 |
| WBGene00007476 | C09E9.1 | 1.237532171 | 0.868701654 | 0.532109958 | 0.192 | 0.153 | 0.100287128 | 1 |
| WBGene00016626 | clhm-1 | 0.910245877 | 0.539958971 | 0.534211083 | 0.1 | 0.074 | 0.231971662 | 1 |
| WBGene00022243 | Y73B6BL.22 | 1.602504337 | 1.232162053 | 0.534290977 | 0.246 | 0.234 | 0.297645861 | 1 |
| WBGene00021922 | atg-3 | 0.703667163 | 0.333041172 | 0.534700279 | 0.062 | 0.052 | 0.588063319 | 1 |
| WBGene00010505 | K02D3.2 | 0.955264938 | 0.583988262 | 0.535639018 | 0.1 | 0.083 | 0.426685599 | 1 |
| WBGene00007068 | cTel55X.1 | 0.62824389 | 0.256798151 | 0.535882926 | 0.077 | 0.033 | 0.020408693 | 1 |
| WBGene00008454 | E02A10.4 | 0.855020496 | 0.482132422 | 0.537963775 | 0.1 | 0.077 | 0.292858346 | 1 |
| WBGene00016246 | daf-37 | 0.589191327 | 0.216155906 | 0.538176351 | 0.069 | 0.033 | 0.050542449 | 1 |
| WBGene00009925 | F52B11.2 | 0.662807692 | 0.289100163 | 0.539145998 | 0.054 | 0.033 | 0.22829074 | 1 |
| WBGene00004902 | snf-3 | 0.47444788 | 0.099762588 | 0.540556612 | 0.054 | 0.015 | 0.008392015 | 1 |
| WBGene00010471 | cdr-6 | 0.553037345 | 0.177821888 | 0.541321478 | 0.069 | 0.027 | 0.016591883 | 1 |
| WBGene00008576 | F08G2.4 | 0.896715772 | 0.52123203 | 0.541708532 | 0.092 | 0.07 | 0.287608681 | 1 |
| WBGene00022434 | Y108G3AL.2 | 0.546509869 | 0.170897783 | 0.541893694 | 0.062 | 0.023 | 0.02088616 | 1 |
| WBGene00016964 | C56C10.7 | 0.768051483 | 0.391790572 | 0.54282975 | 0.062 | 0.068 | 0.927833194 | 1 |
| WBGene00003372 | mlc-4 | 0.653731083 | 0.277031874 | 0.543462081 | 0.077 | 0.039 | 0.049992822 | 1 |
| WBGene00021043 | pck-1 | 0.616341363 | 0.238472138 | 0.545150057 | 0.077 | 0.037 | 0.038355715 | 1 |
| WBGene00010305 | F59A2.5 | 0.560422407 | 0.181457718 | 0.546730478 | 0.062 | 0.029 | 0.059861456 | 1 |
| WBGene00018327 | F42A6.5 | 0.848578333 | 0.468947857 | 0.547691005 | 0.1 | 0.056 | 0.050807383 | 1 |
| WBGene00016303 | fbxc-32 | 0.446971893 | 0.06656264 | 0.548814543 | 0.054 | 0.01 | 0.000757134 | 1 |
| WBGene00012132 | ifta-2 | 1.879119539 | 1.498372604 | 0.549301716 | 0.346 | 0.282 | 0.02529178 | 1 |
| WBGene00020997 | gpa-18 | 1.866173073 | 1.485232045 | 0.549581731 | 0.323 | 0.309 | 0.198842349 | 1 |
| WBGene00016304 | fbxc-41 | 1.483241216 | 1.102242031 | 0.549665634 | 0.223 | 0.213 | 0.38531422 | 1 |
| WBGene00021883 | Y54G2A.18 | 0.615521066 | 0.234392425 | 0.5498524 | 0.069 | 0.025 | 0.011234249 | 1 |
| WBGene00004350 | rgs-7 | 1.496471158 | 1.113868539 | 0.551978901 | 0.177 | 0.18 | 0.684014155 | 1 |
| WBGene00001496 | frm-10 | 1.932343315 | 1.549421025 | 0.55244009 | 0.354 | 0.294 | 0.015148486 | 1 |

| | | | | | | | | |
|---|---|---|---|---|---|---|---|---|
| WBGene00014166 | ZK945.4 | 0.802366768 | 0.418184557 | 0.55425777 | 0.069 | 0.07 | 0.889242552 | 1 |
| WBGene00000899 | daf-3 | 1.032658699 | 0.647287652 | 0.555972898 | 0.1 | 0.106 | 0.944572717 | 1 |
| WBGene00016126 | nhr-258 | 0.982161316 | 0.596494334 | 0.556399842 | 0.115 | 0.104 | 0.490348976 | 1 |
| WBGene00000514 | ckb-4 | 0.508576862 | 0.121197178 | 0.558870748 | 0.054 | 0.012 | 0.001983989 | 1 |
| WBGene00002855 | let-721 | 0.668830474 | 0.279629648 | 0.561498101 | 0.069 | 0.043 | 0.164964772 | 1 |
| WBGene00007200 | vamp-8 | 1.217421694 | 0.827093195 | 0.56312499 | 0.162 | 0.13 | 0.170326091 | 1 |
| WBGene00021804 | Y53G8AM.4 | 1.022679369 | 0.631989188 | 0.563646786 | 0.123 | 0.081 | 0.087416852 | 1 |
| WBGene00219313 | C17E4.20 | 0.791266861 | 0.400191414 | 0.564202609 | 0.085 | 0.064 | 0.305189315 | 1 |
| WBGene00010737 | K10C8.4 | 0.610937236 | 0.219016139 | 0.565422623 | 0.062 | 0.027 | 0.046372954 | 1 |
| WBGene00017180 | F02E11.2 | 3.108753517 | 2.716644662 | 0.565693501 | 0.762 | 0.665 | 1.55E-05 | 0.098948875 |
| WBGene00020294 | T06F4.1 | 0.608847795 | 0.216392678 | 0.566193052 | 0.069 | 0.029 | 0.025377698 | 1 |
| WBGene00018847 | 6-Mar | 1.158446643 | 0.762456782 | 0.57129261 | 0.162 | 0.103 | 0.026718862 | 1 |
| WBGene00015791 | C15C7.5 | 0.617959833 | 0.221209988 | 0.572389035 | 0.054 | 0.031 | 0.187748614 | 1 |
| WBGene00020008 | R11F4.2 | 1.1160801 | 0.718971741 | 0.572906261 | 0.146 | 0.112 | 0.150576911 | 1 |
| WBGene00010732 | K10C3.5 | 0.781326762 | 0.382229117 | 0.575776193 | 0.092 | 0.058 | 0.116077002 | 1 |
| WBGene00020482 | T13C2.7 | 1.330092413 | 0.929935405 | 0.57730453 | 0.192 | 0.162 | 0.16758635 | 1 |
| WBGene00007260 | olrn-1 | 0.756374129 | 0.354001242 | 0.580501368 | 0.085 | 0.048 | 0.083809665 | 1 |
| WBGene00017735 | did-2 | 1.055297586 | 0.652892615 | 0.580547655 | 0.108 | 0.081 | 0.248347192 | 1 |
| WBGene00019872 | R04E5.9 | 1.059356712 | 0.653135876 | 0.586052786 | 0.1 | 0.095 | 0.670081556 | 1 |
| WBGene00021349 | arl-13 | 1.243198142 | 0.832820885 | 0.592049235 | 0.169 | 0.153 | 0.330719263 | 1 |
| WBGene00000970 | dhs-6 | 0.576375001 | 0.165412328 | 0.59289381 | 0.062 | 0.023 | 0.020029637 | 1 |
| WBGene00015295 | acl-12 | 0.557739422 | 0.146430638 | 0.593393143 | 0.054 | 0.021 | 0.037087262 | 1 |
| WBGene00017298 | toca-1 | 0.587809222 | 0.175709943 | 0.594533587 | 0.062 | 0.023 | 0.020693111 | 1 |
| WBGene00006704 | ubc-7 | 1.457628875 | 1.045418648 | 0.59469365 | 0.231 | 0.211 | 0.198393231 | 1 |
| WBGene00008530 | F02E9.5 | 1.139162773 | 0.7266775 | 0.595090458 | 0.146 | 0.122 | 0.268460386 | 1 |
| WBGene00011681 | T10B10.4 | 0.987243144 | 0.574174976 | 0.595931398 | 0.123 | 0.095 | 0.217709468 | 1 |
| WBGene00001094 | dars-1 | 0.592114064 | 0.177915698 | 0.597561929 | 0.077 | 0.023 | 0.002248614 | 1 |
| WBGene00012136 | T28F4.1 | 0.782590172 | 0.368301259 | 0.59769256 | 0.085 | 0.05 | 0.1038233 | 1 |
| WBGene00018509 | xrep-4 | 1.075785601 | 0.661424971 | 0.597796026 | 0.146 | 0.116 | 0.184556224 | 1 |
| WBGene00011434 | T04D3.5 | 0.965878116 | 0.550040193 | 0.599927309 | 0.138 | 0.077 | 0.019978187 | 1 |
| WBGene00018145 | F37C4.5 | 1.792572586 | 1.375787261 | 0.601294123 | 0.338 | 0.29 | 0.02433792 | 1 |
| WBGene00012471 | Y17G7B.20 | 2.009990823 | 1.59168558 | 0.603486899 | 0.377 | 0.321 | 0.009289136 | 1 |
| WBGene00013734 | Y111B2A.10 | 1.131835086 | 0.711641027 | 0.606211885 | 0.146 | 0.118 | 0.227022817 | 1 |
| WBGene00012928 | aakb-2 | 0.663731887 | 0.242262245 | 0.608052162 | 0.062 | 0.033 | 0.108664763 | 1 |
| WBGene00020865 | T27E4.7 | 0.808654796 | 0.386051864 | 0.609687154 | 0.085 | 0.041 | 0.035374214 | 1 |
| WBGene00001178 | egl-9 | 1.269874377 | 0.847214922 | 0.6097687 | 0.215 | 0.128 | 0.003265021 | 1 |
| WBGene00017084 | E01A2.1 | 1.068059578 | 0.645279645 | 0.609942513 | 0.138 | 0.104 | 0.151300732 | 1 |
| WBGene00015002 | B0034.1 | 0.618309764 | 0.193695695 | 0.612588611 | 0.054 | 0.021 | 0.038792892 | 1 |
| WBGene00045190 | R01H2.8 | 1.921406422 | 1.496346914 | 0.613231244 | 0.362 | 0.319 | 0.031970569 | 1 |
| WBGene00008973 | F20D1.1 | 0.713531566 | 0.288043372 | 0.613849706 | 0.077 | 0.041 | 0.06572261 | 1 |
| WBGene00009116 | snx-27 | 1.010141021 | 0.584101981 | 0.61464441 | 0.108 | 0.072 | 0.124753571 | 1 |
| WBGene00016961 | vps-32.1 | 1.589939513 | 1.163544176 | 0.615158439 | 0.269 | 0.211 | 0.034409021 | 1 |
| WBGene00016005 | ift-74 | 1.529119956 | 1.101463642 | 0.616977642 | 0.231 | 0.188 | 0.078476325 | 1 |
| WBGene00007703 | gbf-1 | 0.799906862 | 0.372119208 | 0.617167128 | 0.077 | 0.062 | 0.417585324 | 1 |
| WBGene00010882 | atg-7 | 0.886487763 | 0.458431036 | 0.617555318 | 0.092 | 0.072 | 0.328746087 | 1 |
| WBGene00006373 | syx-5 | 0.65310058 | 0.224748571 | 0.61798132 | 0.062 | 0.027 | 0.045644068 | 1 |
| WBGene00009112 | tag-353 | 0.639380806 | 0.210758329 | 0.618371523 | 0.062 | 0.025 | 0.031681513 | 1 |
| WBGene00021333 | Y34D9A.8 | 1.077099058 | 0.647814894 | 0.619326134 | 0.131 | 0.099 | 0.181955671 | 1 |
| WBGene00012929 | rsks-1 | 1.041576275 | 0.611852799 | 0.619959928 | 0.131 | 0.081 | 0.049466465 | 1 |
| WBGene00019301 | K02D10.1 | 0.999670629 | 0.569138908 | 0.621125979 | 0.131 | 0.074 | 0.023059737 | 1 |
| WBGene00011280 | pelo-1 | 0.667373968 | 0.23535231 | 0.623275505 | 0.077 | 0.029 | 0.009213894 | 1 |
| WBGene00016633 | C44B11.1 | 0.751959517 | 0.317736451 | 0.626451464 | 0.092 | 0.043 | 0.016359768 | 1 |
| WBGene00011263 | R13H4.5 | 0.564139139 | 0.128193141 | 0.62893713 | 0.062 | 0.014 | 0.001078627 | 1 |
| WBGene00017504 | F16B4.2 | 0.706688679 | 0.27022051 | 0.629690463 | 0.077 | 0.029 | 0.009513995 | 1 |
| WBGene00000390 | cdc-42 | 0.921330586 | 0.48435284 | 0.630425627 | 0.1 | 0.072 | 0.201971958 | 1 |
| WBGene00010262 | droe-4 | 3.515015976 | 3.077671353 | 0.630954919 | 0.392 | 0.321 | 0.039400042 | 1 |
| WBGene00019781 | M60.6 | 0.667707396 | 0.229963779 | 0.631530545 | 0.069 | 0.023 | 0.006866568 | 1 |
| WBGene00009069 | F23A7.4 | 1.77342892 | 1.335444593 | 0.631877817 | 0.038 | 0.099 | 0.0358767 | 1 |
| WBGene00022503 | ZC13.2 | 0.935396806 | 0.497154735 | 0.632249662 | 0.115 | 0.058 | 0.016200597 | 1 |
| WBGene00007969 | acs-19 | 0.87915991 | 0.440772161 | 0.632459831 | 0.1 | 0.041 | 0.005111807 | 1 |
| WBGene00020347 | ech-1.2 | 0.650608902 | 0.212037044 | 0.632725445 | 0.062 | 0.031 | 0.082666624 | 1 |
| WBGene00021416 | mksr-2 | 1.643863519 | 1.204640323 | 0.633665127 | 0.308 | 0.226 | 0.004546482 | 1 |
| WBGene00014066 | rev-1 | 0.639492363 | 0.197891219 | 0.63709578 | 0.062 | 0.025 | 0.031681513 | 1 |
| WBGene00077771 | C41G7.9 | 0.821851669 | 0.379120822 | 0.638725598 | 0.085 | 0.06 | 0.23581778 | 1 |

| | | | | | | | | |
|---|---|---|---|---|---|---|---|---|
| WBGene00004990 | spp-5 | 0.730897651 | 0.285511255 | 0.642556745 | 0.077 | 0.039 | 0.05274843 | 1 |
| WBGene00022254 | nlp-71 | 3.073245748 | 2.625987949 | 0.645256609 | 0.592 | 0.542 | 0.021161642 | 1 |
| WBGene00015935 | C17H12.10 | 0.635519925 | 0.188101924 | 0.64548773 | 0.054 | 0.014 | 0.004426881 | 1 |
| WBGene00077526 | C25A1.16 | 0.91188815 | 0.458937735 | 0.653469318 | 0.092 | 0.07 | 0.275734143 | 1 |
| WBGene00007653 | cysl-1 | 2.627557097 | 2.173964358 | 0.654395995 | 0.569 | 0.507 | 0.000247874 | 1 |
| WBGene00017563 | F18C5.10 | 0.562347932 | 0.10852826 | 0.65472339 | 0.069 | 0.014 | 0.000227912 | 1 |
| WBGene00003048 | lit-1 | 2.186091513 | 1.730991347 | 0.656570752 | 0.438 | 0.36 | 0.004109931 | 1 |
| WBGene00015161 | pho-5 | 0.729955514 | 0.274777645 | 0.656682854 | 0.062 | 0.031 | 0.086422643 | 1 |
| WBGene00021352 | pcyt-2.1 | 0.796253522 | 0.338922308 | 0.659789474 | 0.077 | 0.043 | 0.083199322 | 1 |
| WBGene00002001 | hars-1 | 0.632231418 | 0.174312507 | 0.660637341 | 0.062 | 0.017 | 0.004501045 | 1 |
| WBGene00011298 | R107.2 | 0.610499013 | 0.152055859 | 0.661393664 | 0.077 | 0.025 | 0.003355173 | 1 |
| WBGene00007191 | lgc-20 | 1.97899517 | 1.520095008 | 0.662052988 | 0.392 | 0.323 | 0.006353394 | 1 |
| WBGene00015206 | B0495.7 | 0.973755173 | 0.512885663 | 0.664894157 | 0.092 | 0.068 | 0.255144053 | 1 |
| WBGene00007786 | rmd-2 | 0.897279901 | 0.433244516 | 0.66946155 | 0.108 | 0.072 | 0.10512459 | 1 |
| WBGene00008214 | gem-1 | 0.750581188 | 0.286345152 | 0.669751028 | 0.069 | 0.035 | 0.066326273 | 1 |
| WBGene00006363 | syd-1 | 0.67773564 | 0.210453696 | 0.674145343 | 0.062 | 0.027 | 0.04510376 | 1 |
| WBGene00011382 | dop-5 | 1.077205508 | 0.608133712 | 0.676727554 | 0.131 | 0.097 | 0.148915625 | 1 |
| WBGene00000545 | clp-4 | 0.586044654 | 0.114893706 | 0.679727137 | 0.054 | 0.015 | 0.008292944 | 1 |
| WBGene00044004 | C24A8.6 | 1.552918933 | 1.080788054 | 0.681140878 | 0.254 | 0.222 | 0.097283596 | 1 |
| WBGene00007363 | stdh-1 | 0.602842138 | 0.129242235 | 0.683260232 | 0.069 | 0.014 | 0.000227041 | 1 |
| WBGene00000896 | dad-1 | 1.192317542 | 0.718635827 | 0.683378261 | 0.169 | 0.122 | 0.065477615 | 1 |
| WBGene00004108 | pqn-18 | 1.308856421 | 0.834786293 | 0.683938622 | 0.208 | 0.137 | 0.015833308 | 1 |
| WBGene00020812 | acdh-7 | 0.716259361 | 0.24111785 | 0.685484301 | 0.085 | 0.029 | 0.003049408 | 1 |
| WBGene00003903 | pab-2 | 1.876541051 | 1.399873877 | 0.687685368 | 0.331 | 0.253 | 0.009766109 | 1 |
| WBGene00020857 | srt-62 | 0.686673148 | 0.205753643 | 0.693820185 | 0.069 | 0.025 | 0.010968968 | 1 |
| WBGene00011309 | R186.8 | 0.571151153 | 0.089836261 | 0.694390609 | 0.062 | 0.014 | 0.000990061 | 1 |
| WBGene00007484 | C09G1.2 | 2.402381967 | 1.921029872 | 0.69444428 | 0.485 | 0.418 | 0.003344014 | 1 |
| WBGene00009672 | F43G9.12 | 0.72041357 | 0.239051259 | 0.694459018 | 0.077 | 0.031 | 0.013723214 | 1 |
| WBGene00008419 | wdr-23 | 1.373233276 | 0.891563855 | 0.694902084 | 0.185 | 0.143 | 0.098197897 | 1 |
| WBGene00007364 | stdh-3 | 0.721073782 | 0.238861033 | 0.695685942 | 0.085 | 0.029 | 0.003221795 | 1 |
| WBGene00012648 | Y39A1A.9 | 1.163524787 | 0.680724602 | 0.696533433 | 0.131 | 0.101 | 0.195234655 | 1 |
| WBGene00000004 | aat-3 | 0.91083049 | 0.426130779 | 0.699273868 | 0.131 | 0.07 | 0.012075746 | 1 |
| WBGene00077536 | F38B7.10 | 1.396983868 | 0.910567116 | 0.701751036 | 0.215 | 0.159 | 0.03064011 | 1 |
| WBGene00012472 | Y17G7B.21 | 1.071327912 | 0.584042383 | 0.703004416 | 0.123 | 0.083 | 0.098353054 | 1 |
| WBGene00010470 | cdr-4 | 0.588232589 | 0.099035062 | 0.705762845 | 0.069 | 0.015 | 0.000494172 | 1 |
| WBGene00003962 | pdi-1 | 0.86187543 | 0.372380803 | 0.70619147 | 0.077 | 0.058 | 0.340175155 | 1 |
| WBGene00016634 | C44B11.4 | 3.054766214 | 2.564143647 | 0.707818744 | 0.762 | 0.615 | 4.93E-07 | 0.00314605 |
| WBGene00020807 | T25F10.4 | 0.626821067 | 0.13530785 | 0.70910368 | 0.062 | 0.021 | 0.01219693 | 1 |
| WBGene00008227 | C50B8.5 | 0.766391864 | 0.274721612 | 0.709330234 | 0.062 | 0.039 | 0.211871135 | 1 |
| WBGene00015517 | C06A8.8 | 1.115157916 | 0.618440885 | 0.716611197 | 0.154 | 0.081 | 0.006016527 | 1 |
| WBGene00016305 | fbxc-57 | 1.488236101 | 0.989064416 | 0.720152514 | 0.231 | 0.195 | 0.100559381 | 1 |
| WBGene00006523 | tam-1 | 1.204634473 | 0.705406735 | 0.720233382 | 0.162 | 0.116 | 0.074339147 | 1 |
| WBGene00000293 | cap-2 | 1.040048392 | 0.540551146 | 0.7206222 | 0.146 | 0.083 | 0.014789134 | 1 |
| WBGene00021237 | crtc-1 | 0.945296671 | 0.438369538 | 0.73134126 | 0.1 | 0.064 | 0.107138002 | 1 |
| WBGene00009512 | bigr-1 | 0.721692652 | 0.214665725 | 0.731485232 | 0.085 | 0.021 | 0.00029612 | 1 |
| WBGene00016475 | srab-9 | 0.507622649 | 0 | 0.732344678 | 0.062 | 0 | 1.43E-08 | 9.12E-05 |
| WBGene00017618 | F20A1.6 | 0.59573681 | 0.088011877 | 0.732492243 | 0.054 | 0.012 | 0.001955308 | 1 |
| WBGene00019776 | M6.4 | 1.136970384 | 0.627701255 | 0.734720048 | 0.115 | 0.099 | 0.401198617 | 1 |
| WBGene00003915 | pan-1 | 1.643161569 | 1.132459646 | 0.736787132 | 0.254 | 0.184 | 0.016355924 | 1 |
| WBGene00019767 | rpa-2 | 1.367594822 | 0.856229617 | 0.737744045 | 0.177 | 0.128 | 0.062597073 | 1 |
| WBGene00000224 | atgp-1 | 1.020406456 | 0.507104532 | 0.74053814 | 0.138 | 0.083 | 0.025594196 | 1 |
| WBGene00000370 | ccg-1 | 0.524743882 | 0.011156995 | 0.740949255 | 0.054 | 0.002 | 1.65E-06 | 0.010523291 |
| WBGene00016316 | C32D5.8 | 2.083968202 | 1.566211271 | 0.746965357 | 0.446 | 0.329 | 6.58E-05 | 0.419889664 |
| WBGene00000843 | cup-2 | 0.937532697 | 0.418448761 | 0.748879821 | 0.108 | 0.064 | 0.053173095 | 1 |
| WBGene00001034 | dnj-16 | 0.815528067 | 0.296353799 | 0.749010142 | 0.1 | 0.046 | 0.011862808 | 1 |
| WBGene00000684 | col-110 | 0.543861898 | 0.024403485 | 0.749420077 | 0.054 | 0.004 | 1.29E-05 | 0.082027911 |
| WBGene00016192 | bcl-7 | 0.586529199 | 0.064724993 | 0.75280434 | 0.062 | 0.012 | 0.000407156 | 1 |
| WBGene00008975 | F20D1.3 | 0.644464605 | 0.122610476 | 0.752876364 | 0.085 | 0.019 | 0.000127242 | 0.811928315 |
| WBGene00010318 | F59B10.2 | 1.051613854 | 0.528702496 | 0.754401624 | 0.138 | 0.091 | 0.050806784 | 1 |
| WBGene00019339 | K02F6.5 | 2.200066645 | 1.673567898 | 0.759577132 | 0.446 | 0.385 | 0.001617119 | 1 |
| WBGene00003602 | nhr-3 | 1.135429932 | 0.602855514 | 0.768342472 | 0.154 | 0.081 | 0.006689208 | 1 |
| WBGene00020896 | T28D9.4 | 0.702412525 | 0.169610142 | 0.768671355 | 0.062 | 0.023 | 0.020029637 | 1 |
| WBGene00007048 | nfx-1 | 0.659418526 | 0.125859403 | 0.769763101 | 0.069 | 0.015 | 0.000534075 | 1 |
| WBGene00011474 | aldo-1 | 2.242901142 | 1.704385029 | 0.776914525 | 0.508 | 0.385 | 1.03E-05 | 0.065888251 |

| | | | | | | | | |
|---|---|---|---|---|---|---|---|---|
| WBGene00018413 | cyp-33C4 | 0.82248575 | 0.280068463 | 0.782542731 | 0.092 | 0.039 | 0.007925942 | 1 |
| WBGene00021345 | Y37B11A.3 | 0.841455609 | 0.296397478 | 0.786352662 | 0.092 | 0.048 | 0.035893043 | 1 |
| WBGene00007929 | dmd-10 | 0.881427546 | 0.33628261 | 0.786477896 | 0.108 | 0.044 | 0.003605255 | 1 |
| WBGene00000279 | cah-1 | 0.906483574 | 0.359449078 | 0.789203954 | 0.108 | 0.041 | 0.001800425 | 1 |
| WBGene00007030 | epc-1 | 1.776181414 | 1.227470803 | 0.791622077 | 0.292 | 0.236 | 0.023337882 | 1 |
| WBGene00020366 | acdh-10 | 0.800468229 | 0.248786993 | 0.795907783 | 0.1 | 0.035 | 0.00131555 | 1 |
| WBGene00000066 | act-4 | 2.654807639 | 2.102824868 | 0.796342806 | 0.608 | 0.499 | 2.93E-06 | 0.018664725 |
| WBGene00044437 | Y47G6A.32 | 0.846693024 | 0.291082664 | 0.801576311 | 0.092 | 0.039 | 0.008294512 | 1 |
| WBGene00012339 | W07G4.5 | 1.3404804 | 0.782948529 | 0.804348465 | 0.185 | 0.141 | 0.079106786 | 1 |
| WBGene00194713 | F19B10.13 | 0.813901129 | 0.254496543 | 0.807050222 | 0.077 | 0.031 | 0.013143591 | 1 |
| WBGene00017385 | F11G11.5 | 0.629078171 | 0.068409498 | 0.808873915 | 0.054 | 0.012 | 0.001885262 | 1 |
| WBGene00249817 | F42G8.19 | 1.613393822 | 1.047742448 | 0.816062432 | 0.238 | 0.184 | 0.027791231 | 1 |
| WBGene00007485 | C09G1.3 | 2.34200108 | 1.77624205 | 0.816217746 | 0.508 | 0.369 | 3.11E-05 | 0.198219636 |
| WBGene00001757 | gst-9 | 0.581396589 | 0.013444803 | 0.819381224 | 0.054 | 0.002 | 1.67E-06 | 0.010668123 |
| WBGene00014255 | ZK1320.5 | 0.867421243 | 0.298258018 | 0.821128962 | 0.062 | 0.017 | 0.004215502 | 1 |
| WBGene00001812 | haf-2 | 0.652005151 | 0.081974151 | 0.822380896 | 0.069 | 0.012 | 8.06E-05 | 0.514026895 |
| WBGene00000446 | ceh-23 | 1.67397938 | 1.099750584 | 0.828437036 | 0.285 | 0.176 | 0.000830922 | 1 |
| WBGene00020775 | T24E12.6 | 0.669233264 | 0.092222726 | 0.832450242 | 0.062 | 0.012 | 0.000397633 | 1 |
| WBGene00003610 | nhr-11 | 1.210403149 | 0.632353697 | 0.833949077 | 0.177 | 0.079 | 0.000326234 | 1 |
| WBGene00008310 | C54E10.6 | 1.029021989 | 0.450594156 | 0.834494967 | 0.1 | 0.068 | 0.1455087 | 1 |
| WBGene00002219 | klp-7 | 0.770242027 | 0.190761334 | 0.836013922 | 0.085 | 0.019 | 0.000141181 | 0.900876755 |
| WBGene00006139 | str-79 | 2.171594015 | 1.591685446 | 0.836631216 | 0.469 | 0.352 | 2.24E-05 | 0.142739556 |
| WBGene00021059 | atat-2 | 1.088848187 | 0.508856165 | 0.836751614 | 0.123 | 0.079 | 0.071249195 | 1 |
| WBGene00008579 | F08G2.8 | 0.884196761 | 0.303081056 | 0.838372745 | 0.085 | 0.039 | 0.022481825 | 1 |
| WBGene00012009 | T25B9.3 | 1.372841532 | 0.788768791 | 0.842638846 | 0.231 | 0.126 | 0.000562631 | 1 |
| WBGene00004057 | pmk-3 | 0.899291202 | 0.315180364 | 0.842693809 | 0.108 | 0.046 | 0.004607447 | 1 |
| WBGene00017004 | nrd-1 | 0.863659266 | 0.278163306 | 0.844692117 | 0.108 | 0.035 | 0.000429247 | 1 |
| WBGene00007882 | C33A12.3 | 1.980342177 | 1.393799608 | 0.846202055 | 0.338 | 0.279 | 0.009976988 | 1 |
| WBGene00002267 | lec-4 | 0.601066764 | 0.010795813 | 0.851580973 | 0.069 | 0.002 | 2.61E-08 | 0.000166775 |
| WBGene00016321 | tppp-1 | 2.29738388 | 1.705724781 | 0.853583648 | 0.446 | 0.35 | 0.000386547 | 1 |
| WBGene00004859 | sma-5 | 0.994464766 | 0.401732613 | 0.855131739 | 0.115 | 0.056 | 0.009393249 | 1 |
| WBGene00015678 | mcp-1 | 1.305088299 | 0.711116143 | 0.856920684 | 0.138 | 0.087 | 0.046547184 | 1 |
| WBGene00008206 | set-6 | 1.031452241 | 0.433155766 | 0.863159357 | 0.123 | 0.056 | 0.004353884 | 1 |
| WBGene00002783 | let-607 | 0.991486344 | 0.392469897 | 0.864198058 | 0.123 | 0.039 | 0.000133785 | 0.853680824 |
| WBGene00011292 | allo-1 | 2.169847428 | 1.56614848 | 0.870953479 | 0.438 | 0.337 | 0.000278424 | 1 |
| WBGene00005647 | srp-6 | 0.885785257 | 0.272381916 | 0.884953958 | 0.092 | 0.031 | 0.001498033 | 1 |
| WBGene00016395 | spcs-1 | 1.076641798 | 0.463037767 | 0.885243493 | 0.131 | 0.075 | 0.022214427 | 1 |
| WBGene00021556 | Y45G5AM.3 | 0.664091507 | 0.049326719 | 0.886918111 | 0.069 | 0.006 | 1.56E-06 | 0.009961047 |
| WBGene00020736 | T23F2.3 | 3.197866668 | 2.580193585 | 0.891113894 | 0.792 | 0.642 | 3.92E-09 | 2.50E-05 |
| WBGene00010018 | fbxa-69 | 0.728528106 | 0.1102976 | 0.891918085 | 0.062 | 0.017 | 0.004018266 | 1 |
| WBGene00013692 | Y105E8B.7 | 0.911397244 | 0.288665019 | 0.898412693 | 0.108 | 0.044 | 0.003205565 | 1 |
| WBGene00006498 | ten-1 | 1.210184602 | 0.586371452 | 0.899972138 | 0.146 | 0.074 | 0.004733804 | 1 |
| WBGene00018795 | F54C4.4 | 0.793439403 | 0.167822389 | 0.902574563 | 0.085 | 0.023 | 0.000551615 | 1 |
| WBGene00016150 | hint-3 | 0.680572726 | 0.054936945 | 0.902601638 | 0.062 | 0.004 | 1.91E-06 | 0.012218965 |
| WBGene00003387 | mod-5 | 1.19392864 | 0.56496029 | 0.90740952 | 0.169 | 0.079 | 0.00074461 | 1 |
| WBGene00019664 | K11H12.8 | 0.829019091 | 0.196938893 | 0.911898967 | 0.092 | 0.021 | 7.62E-05 | 0.486539299 |
| WBGene00003040 | lin-59 | 1.299432665 | 0.667151546 | 0.912188835 | 0.2 | 0.097 | 0.000307095 | 1 |
| WBGene00022824 | ZK813.5 | 1.863397729 | 1.229255818 | 0.91487339 | 0.362 | 0.248 | 0.000142628 | 0.910107234 |
| WBGene00021610 | nhr-237 | 1.469914291 | 0.834837144 | 0.916222651 | 0.238 | 0.143 | 0.001368902 | 1 |
| WBGene00020481 | T13C2.6 | 1.065350385 | 0.430219198 | 0.916300615 | 0.138 | 0.056 | 0.000635101 | 1 |
| WBGene00021011 | W03G1.5 | 1.142486274 | 0.504720372 | 0.920101704 | 0.146 | 0.062 | 0.000779191 | 1 |
| WBGene00043054 | T21B4.15 | 1.21693063 | 0.579123233 | 0.920161569 | 0.185 | 0.079 | 0.000108615 | 0.693071099 |
| WBGene00007913 | cyp-36A1 | 4.571820217 | 3.927307503 | 0.929835296 | 0.977 | 0.814 | 5.71E-18 | 3.64E-14 |
| WBGene00015626 | C09B9.1 | 2.310454604 | 1.660471359 | 0.937727605 | 0.423 | 0.369 | 0.003414832 | 1 |
| WBGene00021915 | Y55D5A.1 | 0.844195083 | 0.192535207 | 0.940146471 | 0.085 | 0.029 | 0.002715842 | 1 |
| WBGene00018758 | F53E10.1 | 0.709910314 | 0.057583749 | 0.9411083 | 0.092 | 0.008 | 2.52E-08 | 0.000160925 |
| WBGene00000092 | ags-3 | 2.491094912 | 1.832408403 | 0.950283759 | 0.531 | 0.395 | 1.30E-05 | 0.082767667 |
| WBGene00005087 | srd-9 | 0.782011694 | 0.118430387 | 0.957345462 | 0.062 | 0.017 | 0.004115812 | 1 |
| WBGene00000973 | dhs-9 | 0.899161459 | 0.23239283 | 0.961943795 | 0.108 | 0.035 | 0.000362299 | 1 |
| WBGene00007024 | plin-1 | 0.883545901 | 0.214704682 | 0.96493391 | 0.077 | 0.027 | 0.005532649 | 1 |
| WBGene00017934 | F30B5.4 | 1.005245892 | 0.335039088 | 0.966904033 | 0.115 | 0.05 | 0.00350097 | 1 |
| WBGene00021854 | Y54F10AR.1 | 1.068901561 | 0.398536431 | 0.967132448 | 0.169 | 0.05 | 1.86E-06 | 0.011862119 |
| WBGene00010827 | M02B1.2 | 0.937831 | 0.264287938 | 0.971717236 | 0.123 | 0.043 | 0.000262289 | 1 |
| WBGene00004208 | ptc-1 | 2.313613791 | 1.639326841 | 0.97279044 | 0.454 | 0.329 | 2.80E-05 | 0.178565275 |

| | | | | | | | | |
|---|---|---|---|---|---|---|---|---|
| WBGene00003707 | nhr-117 | 1.163399956 | 0.487141367 | 0.975634912 | 0.169 | 0.077 | 0.00049239 | 1 |
| WBGene00003019 | lin-33 | 1.434963259 | 0.757524906 | 0.977336953 | 0.2 | 0.116 | 0.002866257 | 1 |
| WBGene00013647 | Y105C5B.5 | 1.189993005 | 0.512441326 | 0.977500447 | 0.1 | 0.056 | 0.054327243 | 1 |
| WBGene00020737 | T23F2.4 | 0.937691806 | 0.259198676 | 0.978858674 | 0.1 | 0.023 | 3.58E-05 | 0.228694216 |
| WBGene00004134 | myrf-1 | 1.542120053 | 0.861181191 | 0.98238712 | 0.238 | 0.133 | 0.000858502 | 1 |
| WBGene00002007 | hsp-3 | 1.737910328 | 1.056567719 | 0.982969602 | 0.254 | 0.147 | 0.000534418 | 1 |
| WBGene00011935 | scrm-1 | 1.694037406 | 1.011143823 | 0.985207186 | 0.246 | 0.205 | 0.076348747 | 1 |
| WBGene00008021 | C39B10.1 | 0.907373826 | 0.217682657 | 0.995014029 | 0.115 | 0.025 | 4.11E-06 | 0.026207039 |
| WBGene00019305 | snap-29 | 1.014823929 | 0.322374877 | 0.998992813 | 0.069 | 0.046 | 0.246327114 | 1 |
| WBGene00019994 | cdh-1 | 0.831357737 | 0.133070517 | 1.007415509 | 0.092 | 0.019 | 2.87E-05 | 0.182969366 |
| WBGene00019342 | K02F6.8 | 2.103610443 | 1.402287738 | 1.01179479 | 0.431 | 0.304 | 1.70E-05 | 0.108172923 |
| WBGene00002263 | lea-1 | 0.882780171 | 0.172778205 | 1.024316314 | 0.108 | 0.017 | 5.27E-07 | 0.003362012 |
| WBGene00015947 | C18A11.1 | 0.735031002 | 0.024841493 | 1.024586884 | 0.069 | 0.002 | 2.80E-08 | 0.000178934 |
| WBGene00002232 | kpc-1 | 1.356329667 | 0.646083382 | 1.024668793 | 0.185 | 0.087 | 0.000478756 | 1 |
| WBGene00001425 | fis-2 | 0.960455045 | 0.246718286 | 1.029704482 | 0.123 | 0.029 | 4.13E-06 | 0.026360199 |
| WBGene00009726 | tbc-13 | 1.042347357 | 0.326603552 | 1.032600038 | 0.115 | 0.05 | 0.003459305 | 1 |
| WBGene00021039 | W05F2.7 | 1.838722459 | 1.120120337 | 1.036723718 | 0.315 | 0.199 | 0.000242248 | 1 |
| WBGene00020694 | dhhc-11 | 2.355002824 | 1.635145667 | 1.038534351 | 0.446 | 0.354 | 0.000194419 | 1 |
| WBGene00020751 | srbc-69 | 0.850795089 | 0.130791379 | 1.038745782 | 0.123 | 0.017 | 1.63E-08 | 0.000103941 |
| WBGene00004131 | pqn-44 | 2.883540553 | 2.162222676 | 1.040641724 | 0.646 | 0.474 | 4.23E-08 | 0.000269695 |
| WBGene00011425 | T04B2.7 | 0.733966333 | 0.010301341 | 1.044027896 | 0.085 | 0.002 | 4.05E-10 | 2.59E-06 |
| WBGene00007553 | C13G3.1 | 1.67515695 | 0.949144175 | 1.04741503 | 0.192 | 0.05 | 3.72E-08 | 0.000237545 |
| WBGene00012140 | T28F4.5 | 3.16602187 | 2.438133068 | 1.050121565 | 0.677 | 0.53 | 5.55E-09 | 3.54E-05 |
| WBGene00004405 | rop-1 | 1.296982034 | 0.567324411 | 1.052673434 | 0.162 | 0.07 | 0.000344986 | 1 |
| WBGene00010329 | pcdr-1 | 1.608304914 | 0.87631999 | 1.056031019 | 0.269 | 0.145 | 5.45E-05 | 0.347721105 |
| WBGene00002107 | ins-24 | 2.895554176 | 2.160816077 | 1.060003011 | 0.608 | 0.509 | 1.88E-06 | 0.012020575 |
| WBGene00005096 | srd-18 | 0.961325589 | 0.219608635 | 1.070071371 | 0.115 | 0.027 | 9.47E-06 | 0.060398839 |
| WBGene00021979 | Y58A7A.5 | 0.74182496 | 0 | 1.070227191 | 0.062 | 0 | 1.43E-08 | 9.12E-05 |
| WBGene00015955 | C18B2.4 | 1.494393488 | 0.752343241 | 1.070552211 | 0.231 | 0.13 | 0.000438083 | 1 |
| WBGene00000996 | dif-1 | 1.728150258 | 0.98365724 | 1.074076385 | 0.277 | 0.166 | 0.000406442 | 1 |
| WBGene00019465 | acl-14 | 1.516072491 | 0.769754089 | 1.076709857 | 0.208 | 0.116 | 0.001352372 | 1 |
| WBGene00018748 | F53C3.4 | 0.883788862 | 0.134529059 | 1.080953401 | 0.115 | 0.017 | 9.92E-08 | 0.000633296 |
| WBGene00021811 | ral-1 | 1.329174826 | 0.579902236 | 1.08097185 | 0.185 | 0.066 | 8.82E-06 | 0.056266069 |
| WBGene00004443 | rpl-29 | 2.442747865 | 1.687900272 | 1.08901488 | 0.538 | 0.395 | 2.88E-07 | 0.001840461 |
| WBGene00016017 | C23G10.11 | 1.293697283 | 0.538460048 | 1.089577015 | 0.123 | 0.062 | 0.011019908 | 1 |
| WBGene00009165 | gpx-1 | 2.296346712 | 1.54006794 | 1.091079633 | 0.377 | 0.337 | 0.013969521 | 1 |
| WBGene00012907 | cpt-1 | 1.850035367 | 1.089372452 | 1.097404615 | 0.277 | 0.207 | 0.011347363 | 1 |
| WBGene00010828 | M02B1.3 | 1.071970026 | 0.309755243 | 1.099643489 | 0.146 | 0.033 | 3.15E-07 | 0.002009251 |
| WBGene00020596 | oga-1 | 1.411755581 | 0.64780409 | 1.102149027 | 0.208 | 0.095 | 8.27E-05 | 0.527760069 |
| WBGene00010918 | M117.1 | 1.759642042 | 0.9935755 | 1.105200402 | 0.285 | 0.174 | 0.00044159 | 1 |
| WBGene00012444 | Y15E3A.5 | 2.546416761 | 1.75738768 | 1.138328343 | 0.462 | 0.267 | 2.30E-07 | 0.001466159 |
| WBGene00019686 | K12H6.6 | 1.41281098 | 0.621521021 | 1.141590101 | 0.223 | 0.079 | 6.68E-07 | 0.004265579 |
| WBGene00018634 | pigx-1 | 1.057035436 | 0.263176116 | 1.145296905 | 0.138 | 0.037 | 3.77E-06 | 0.024080542 |
| WBGene00017907 | F28F9.3 | 2.500153885 | 1.701864507 | 1.151688127 | 0.477 | 0.348 | 8.73E-06 | 0.055715196 |
| WBGene00000534 | cpi-2 | 1.790793783 | 0.988253929 | 1.157820268 | 0.308 | 0.182 | 5.12E-05 | 0.326390466 |
| WBGene00011631 | T08G11.4 | 1.421298587 | 0.618402054 | 1.158334847 | 0.223 | 0.093 | 7.95E-06 | 0.05073328 |
| WBGene00018259 | F41B4.3 | 0.886865859 | 0.082004426 | 1.161169598 | 0.1 | 0.012 | 6.85E-08 | 0.000437065 |
| WBGene00004793 | shw-3 | 1.045615962 | 0.239690424 | 1.162704777 | 0.123 | 0.029 | 4.73E-06 | 0.030197938 |
| WBGene00007981 | glb-11 | 2.011785945 | 1.198976139 | 1.172636677 | 0.362 | 0.209 | 2.44E-06 | 0.015557312 |
| WBGene00010879 | taap-1 | 1.301932984 | 0.478912292 | 1.18736787 | 0.185 | 0.081 | 0.000120489 | 0.768839724 |
| WBGene00000298 | cat-4 | 2.976272377 | 2.153138885 | 1.187530607 | 0.646 | 0.547 | 1.08E-06 | 0.006884625 |
| WBGene00016506 | abhd-5.1 | 1.189012576 | 0.36563812 | 1.187878244 | 0.154 | 0.06 | 0.000112093 | 0.715266301 |
| WBGene00008745 | F13E6.1 | 1.507874228 | 0.683552461 | 1.189244926 | 0.223 | 0.106 | 5.61E-05 | 0.35797785 |
| WBGene00021611 | nhr-238 | 1.451108254 | 0.623371426 | 1.194171817 | 0.223 | 0.083 | 9.31E-07 | 0.005938633 |
| WBGene00007217 | tads-1 | 1.22820482 | 0.400149821 | 1.194630841 | 0.162 | 0.058 | 2.93E-05 | 0.18679856 |
| WBGene00000802 | crt-1 | 2.020979338 | 1.191793896 | 1.196261725 | 0.292 | 0.211 | 0.00258719 | 1 |
| WBGene00012002 | T24H10.4 | 1.43659185 | 0.606930264 | 1.196948656 | 0.192 | 0.072 | 7.69E-06 | 0.049070326 |
| WBGene00014473 | MTCE.36 | 1.181729438 | 0.350433381 | 1.199306699 | 0.138 | 0.039 | 9.34E-06 | 0.059615133 |
| WBGene00001749 | gst-1 | 1.180877014 | 0.348016451 | 1.201563804 | 0.092 | 0.052 | 0.058543465 | 1 |
| WBGene00043308 | tmem-107 | 1.183666084 | 0.342295678 | 1.213840912 | 0.162 | 0.043 | 4.17E-07 | 0.002660522 |
| WBGene00019304 | K02D10.4 | 1.51090798 | 0.658264083 | 1.230105122 | 0.231 | 0.101 | 9.32E-06 | 0.059474533 |
| WBGene00006840 | unc-116 | 1.42368769 | 0.570421218 | 1.231003307 | 0.192 | 0.087 | 0.000112709 | 0.719198135 |
| WBGene00003989 | pfn-1 | 1.119279846 | 0.264189736 | 1.233634261 | 0.131 | 0.033 | 4.28E-06 | 0.027314695 |
| WBGene00008681 | scrm-4 | 2.36719767 | 1.508583487 | 1.238718425 | 0.492 | 0.288 | 2.81E-09 | 1.79E-05 |

| | | | | | | | | |
|---|---|---|---|---|---|---|---|---|
| WBGene00010724 | K09E9.1 | 1.403091003 | 0.541780118 | 1.242608943 | 0.192 | 0.077 | 2.83E-05 | 0.180803767 |
| WBGene00000442 | ceh-19 | 2.799052077 | 1.93691048 | 1.243807406 | 0.662 | 0.41 | 7.65E-13 | 4.88E-09 |
| WBGene00008343 | C56A3.4 | 1.070923613 | 0.207051149 | 1.246304519 | 0.138 | 0.033 | 8.07E-07 | 0.005148058 |
| WBGene00011772 | T14G8.4 | 2.022230803 | 1.155979113 | 1.249737017 | 0.377 | 0.191 | 6.59E-08 | 0.000420782 |
| WBGene00003408 | mrp-2 | 2.137470293 | 1.267186419 | 1.255554228 | 0.385 | 0.224 | 1.65E-06 | 0.010517707 |
| WBGene00007010 | alx-1 | 1.147816973 | 0.271840062 | 1.263767547 | 0.138 | 0.041 | 1.10E-05 | 0.070381645 |
| WBGene00013878 | atfs-1 | 1.09391629 | 0.204329442 | 1.283402534 | 0.138 | 0.031 | 3.87E-07 | 0.002471053 |
| WBGene00007574 | C14B1.3 | 2.196577328 | 1.306922314 | 1.283500877 | 0.446 | 0.273 | 4.54E-08 | 0.000289403 |
| WBGene00007343 | C05E7.1 | 0.923280162 | 0.009870287 | 1.317771897 | 0.123 | 0.002 | 1.11E-14 | 7.05E-11 |
| WBGene00017675 | F21F3.6 | 2.112807012 | 1.198518152 | 1.319040005 | 0.408 | 0.232 | 1.05E-07 | 0.000670485 |
| WBGene00003930 | pat-3 | 0.964930476 | 0.050243992 | 1.319613655 | 0.115 | 0.008 | 5.93E-11 | 3.78E-07 |
| WBGene00008476 | E03H4.8 | 0.967435575 | 0.052233846 | 1.320356996 | 0.085 | 0.004 | 4.53E-09 | 2.89E-05 |
| WBGene00006923 | vhp-1 | 2.420268705 | 1.49754299 | 1.331211814 | 0.546 | 0.302 | 3.55E-12 | 2.27E-08 |
| WBGene00194982 | Y17D7B.10 | 2.457950974 | 1.534389551 | 1.332417485 | 0.331 | 0.244 | 0.003540875 | 1 |
| WBGene00016783 | irg-2 | 0.968692729 | 0.010485629 | 1.382400631 | 0.062 | 0.002 | 2.08E-07 | 0.001328557 |
| WBGene00001527 | gcs-1 | 1.332325286 | 0.360923416 | 1.40143666 | 0.131 | 0.037 | 1.77E-05 | 0.112769728 |
| WBGene00189933 | F38B7.11 | 1.659727514 | 0.687899624 | 1.402051277 | 0.254 | 0.112 | 3.40E-06 | 0.021702043 |
| WBGene00001779 | gst-31 | 1.115977164 | 0.140106461 | 1.407883823 | 0.131 | 0.017 | 2.43E-09 | 1.55E-05 |
| WBGene00015561 | C07A12.7 | 1.538370842 | 0.553786071 | 1.420455567 | 0.223 | 0.072 | 4.80E-08 | 0.000306167 |
| WBGene00017082 | DC2.5 | 1.349162592 | 0.363590456 | 1.421880033 | 0.138 | 0.056 | 0.000505189 | 1 |
| WBGene00020738 | T23F2.5 | 2.585287785 | 1.59726224 | 1.425419553 | 0.577 | 0.36 | 6.08E-12 | 3.88E-08 |
| WBGene00018290 | vps-60 | 1.52355441 | 0.532597482 | 1.429648646 | 0.192 | 0.077 | 1.84E-05 | 0.117366014 |
| WBGene00003647 | nhr-57 | 1.148383624 | 0.15205057 | 1.437404756 | 0.123 | 0.014 | 1.14E-09 | 7.29E-06 |
| WBGene00018750 | F53C3.6 | 1.269070637 | 0.271565733 | 1.439095378 | 0.185 | 0.041 | 1.82E-09 | 1.16E-05 |
| WBGene00007346 | frpr-2 | 1.373358559 | 0.374293287 | 1.441346513 | 0.223 | 0.05 | 7.34E-11 | 4.69E-07 |
| WBGene00017467 | F14F9.4 | 1.366390267 | 0.36427995 | 1.445739584 | 0.192 | 0.041 | 6.17E-10 | 3.93E-06 |
| WBGene00011480 | enpl-1 | 1.570855454 | 0.560538041 | 1.457579922 | 0.208 | 0.089 | 1.83E-05 | 0.116904658 |
| WBGene00016175 | C27H5.6 | 2.374915227 | 1.363883029 | 1.458611138 | 0.446 | 0.23 | 9.29E-10 | 5.93E-06 |
| WBGene00022252 | Y73B6BL.31 | 1.788689791 | 0.773396451 | 1.464758666 | 0.308 | 0.118 | 2.75E-09 | 1.76E-05 |
| WBGene00010837 | M03B6.5 | 1.276848743 | 0.256086209 | 1.472649046 | 0.169 | 0.033 | 2.11E-09 | 1.35E-05 |
| WBGene00007216 | C01A2.4 | 1.290858325 | 0.269302723 | 1.473793201 | 0.154 | 0.025 | 1.56E-09 | 9.94E-06 |
| WBGene00006372 | syx-2 | 1.333980999 | 0.311109999 | 1.47569092 | 0.192 | 0.048 | 7.78E-09 | 4.96E-05 |
| WBGene00010700 | nipi-3 | 2.093578887 | 1.063399803 | 1.486234256 | 0.385 | 0.195 | 8.45E-09 | 5.39E-05 |
| WBGene00194710 | mks-2 | 1.972785986 | 0.940904181 | 1.488690763 | 0.338 | 0.153 | 1.56E-08 | 9.93E-05 |
| WBGene00002086 | ins-3 | 4.893616146 | 3.846229164 | 1.511060006 | 0.969 | 0.77 | 3.39E-29 | 2.16E-25 |
| WBGene00008075 | C43F9.7 | 2.677985761 | 1.627289238 | 1.515834664 | 0.562 | 0.319 | 1.91E-12 | 1.22E-08 |
| WBGene00010504 | K02D3.1 | 3.210221676 | 2.149396673 | 1.53044697 | 0.623 | 0.455 | 2.74E-10 | 1.75E-06 |
| WBGene00007366 | C06B3.7 | 1.340829152 | 0.269942918 | 1.544962258 | 0.192 | 0.039 | 1.67E-10 | 1.06E-06 |
| WBGene00016488 | C36C9.5 | 1.984613406 | 0.90615025 | 1.555893447 | 0.331 | 0.151 | 2.82E-08 | 0.000179771 |
| WBGene00003963 | pdi-2 | 1.488754269 | 0.405008797 | 1.563514218 | 0.215 | 0.041 | 5.94E-12 | 3.79E-08 |
| WBGene00000097 | aip-1 | 2.190240723 | 1.098991049 | 1.574340493 | 0.346 | 0.226 | 2.41E-05 | 0.153853161 |
| WBGene00008726 | F13A7.11 | 1.137757168 | 0.036247934 | 1.58914191 | 0.138 | 0.002 | 1.96E-16 | 1.25E-12 |
| WBGene00021977 | Y58A7A.3 | 1.297836833 | 0.18429498 | 1.606501309 | 0.169 | 0.014 | 1.32E-14 | 8.44E-11 |
| WBGene00015167 | B0403.3 | 1.127658374 | 0.012152383 | 1.609334961 | 0.131 | 0.002 | 1.33E-15 | 8.48E-12 |
| WBGene00000993 | dhs-30 | 1.147353964 | 0.024528353 | 1.619894942 | 0.146 | 0.004 | 2.30E-16 | 1.47E-12 |
| WBGene00011423 | ipla-7 | 1.343706248 | 0.206530202 | 1.640598242 | 0.177 | 0.033 | 2.72E-10 | 1.74E-06 |
| WBGene00019006 | spg-20 | 1.671797584 | 0.517678996 | 1.665041162 | 0.254 | 0.079 | 1.42E-09 | 9.06E-06 |
| WBGene00022648 | ZK54.3 | 1.849038225 | 0.694190869 | 1.666092553 | 0.238 | 0.093 | 7.77E-07 | 0.004959954 |
| WBGene00044443 | ZC21.10 | 2.100840142 | 0.936228159 | 1.680179933 | 0.3 | 0.12 | 3.05E-08 | 0.000194662 |
| WBGene00006470 | tag-120 | 1.558494247 | 0.380077873 | 1.700095459 | 0.238 | 0.054 | 7.32E-12 | 4.67E-08 |
| WBGene00016739 | pitr-1 | 1.794924519 | 0.601873347 | 1.721209009 | 0.3 | 0.074 | 3.14E-14 | 2.00E-10 |
| WBGene00016507 | abhd-5.2 | 2.895740338 | 1.702067669 | 1.722105641 | 0.715 | 0.377 | 4.32E-23 | 2.76E-19 |
| WBGene00004373 | rme-1 | 1.981682943 | 0.781596338 | 1.731358994 | 0.415 | 0.133 | 1.22E-16 | 7.81E-13 |
| WBGene00019770 | M04F3.4 | 1.528754121 | 0.315051521 | 1.751002723 | 0.254 | 0.039 | 2.22E-16 | 1.42E-12 |
| WBGene00020363 | fbxa-3 | 1.276432701 | 0.050437982 | 1.768736501 | 0.185 | 0.006 | 6.46E-20 | 4.12E-16 |
| WBGene00010742 | K10D6.2 | 2.025253738 | 0.742502571 | 1.850618747 | 0.338 | 0.12 | 1.39E-11 | 8.86E-08 |
| WBGene00010736 | istr-1 | 1.626964509 | 0.340239385 | 1.856351956 | 0.192 | 0.05 | 1.37E-08 | 8.77E-05 |
| WBGene00008865 | F15G9.1 | 1.314656941 | 0.013596014 | 1.877034147 | 0.185 | 0.002 | 4.10E-22 | 2.62E-18 |
| WBGene00017080 | lec-12 | 1.650109136 | 0.334974171 | 1.897338692 | 0.246 | 0.044 | 1.48E-14 | 9.44E-11 |
| WBGene00013266 | Y57A10A.26 | 2.513909736 | 1.192467812 | 1.90643771 | 0.508 | 0.244 | 1.72E-15 | 1.10E-11 |
| WBGene00011045 | gpx-2 | 1.758139198 | 0.433309757 | 1.911324865 | 0.277 | 0.064 | 6.64E-14 | 4.24E-10 |
| WBGene00015168 | pdi-6 | 1.842556083 | 0.51449544 | 1.915986505 | 0.3 | 0.081 | 3.07E-13 | 1.96E-09 |
| WBGene00015162 | algn-11 | 1.782220713 | 0.448021409 | 1.924842719 | 0.277 | 0.07 | 7.14E-13 | 4.56E-09 |
| WBGene00009803 | F47B8.2 | 3.237512468 | 1.890264266 | 1.9436683 | 0.7 | 0.364 | 2.19E-22 | 1.40E-18 |

| | | | | | | | | |
|---|---|---|---|---|---|---|---|---|
| WBGene00018869 | rfip-1 | 3.547946358 | 2.171843953 | 1.985296114 | 0.846 | 0.487 | 5.09E-33 | 3.25E-29 |
| WBGene00022667 | ZK154.5 | 1.897591442 | 0.503348289 | 2.011467682 | 0.292 | 0.072 | 8.91E-14 | 5.69E-10 |
| WBGene00005655 | srr-4 | 2.872536226 | 1.474149912 | 2.017445001 | 0.708 | 0.308 | 3.41E-28 | 2.18E-24 |
| WBGene00019975 | flp-34 | 1.935663811 | 0.533250343 | 2.023254956 | 0.292 | 0.079 | 6.98E-13 | 4.46E-09 |
| WBGene00013845 | fbxa-37 | 1.458705452 | 0.050829869 | 2.03113512 | 0.192 | 0.008 | 5.49E-20 | 3.51E-16 |
| WBGene00010019 | F54B8.4 | 1.630487444 | 0.217277321 | 2.038831237 | 0.208 | 0.029 | 3.85E-14 | 2.46E-10 |
| WBGene00017597 | F19C7.8 | 3.10432436 | 1.680376358 | 2.054322721 | 0.662 | 0.338 | 7.25E-21 | 4.63E-17 |
| WBGene00002008 | hsp-4 | 1.839222381 | 0.408476724 | 2.064129666 | 0.215 | 0.037 | 9.20E-13 | 5.87E-09 |
| WBGene00008777 | F13H10.6 | 3.613243423 | 2.164952912 | 2.089441539 | 0.877 | 0.462 | 1.39E-38 | 8.86E-35 |
| WBGene00006959 | xbp-1 | 4.608774274 | 3.157209607 | 2.094165147 | 0.985 | 0.708 | 7.80E-49 | 4.98E-45 |
| WBGene00009901 | F49E12.8 | 1.961859518 | 0.4961422 | 2.114583105 | 0.354 | 0.07 | 4.09E-20 | 2.61E-16 |
| WBGene00007365 | C06B3.6 | 3.094740831 | 1.525757209 | 2.263564891 | 0.669 | 0.193 | 2.51E-31 | 1.60E-27 |
| WBGene00235164 | K09F6.13 | 1.654256503 | 0.051517483 | 2.312263636 | 0.115 | 0.006 | 9.10E-12 | 5.81E-08 |
| WBGene00010833 | M03B6.1 | 3.478214656 | 1.865772612 | 2.326262141 | 0.769 | 0.424 | 4.98E-28 | 3.18E-24 |
| WBGene00001787 | gst-39 | 2.445204498 | 0.830039437 | 2.330190623 | 0.392 | 0.143 | 4.16E-14 | 2.66E-10 |
| WBGene00019967 | cyp-33C8 | 2.660538472 | 0.979356875 | 2.425432353 | 0.569 | 0.132 | 8.48E-32 | 5.41E-28 |
| WBGene00015164 | ykt-6 | 2.721655197 | 0.996939666 | 2.488238543 | 0.608 | 0.174 | 1.13E-31 | 7.19E-28 |
| WBGene00022668 | ZK154.6 | 2.737048154 | 0.987178027 | 2.524528954 | 0.546 | 0.143 | 1.91E-28 | 1.22E-24 |
| WBGene00019327 | zip-2 | 3.322611872 | 1.559565913 | 2.543537661 | 0.8 | 0.323 | 7.67E-40 | 4.89E-36 |
| WBGene00005097 | srd-19 | 2.125785066 | 0.306078473 | 2.625281677 | 0.362 | 0.046 | 7.91E-27 | 5.05E-23 |
| WBGene00021487 | comt-3 | 2.2651543 | 0.428313528 | 2.650001073 | 0.323 | 0.041 | 1.22E-23 | 7.75E-20 |
| WBGene00016997 | cebp-1 | 2.459385744 | 0.600478742 | 2.681835913 | 0.592 | 0.099 | 2.15E-42 | 1.37E-38 |
| WBGene00001755 | gst-7 | 2.948226195 | 1.077313764 | 2.699156087 | 0.408 | 0.147 | 2.10E-14 | 1.34E-10 |
| WBGene00008922 | F17C11.11 | 3.71940996 | 1.752740517 | 2.837304252 | 0.754 | 0.302 | 7.25E-36 | 4.62E-32 |
| WBGene00012128 | nra-1 | 2.3497975 | 0.336841215 | 2.904082049 | 0.354 | 0.041 | 3.52E-27 | 2.24E-23 |
| WBGene00003991 | pfn-3 | 2.293903437 | 0.175007701 | 3.056920369 | 0.392 | 0.021 | 7.29E-39 | 4.65E-35 |
| WBGene00010291 | F58H10.1 | 2.584278822 | 0.349170738 | 3.224579348 | 0.415 | 0.043 | 3.28E-34 | 2.09E-30 |
| WBGene00018702 | F52E4.5 | 2.499272261 | 0.036116082 | 3.553583204 | 0.477 | 0.006 | 8.71E-58 | 5.56E-54 |
| WBGene00016788 | C49G7.10 | 2.578095595 | 0 | 3.719405729 | 0.162 | 0 | 1.68E-20 | 1.07E-16 |
| WBGene00194689 | Y38H6C.24 | 3.103681306 | 0.393796656 | 3.909537146 | 0.469 | 0.046 | 2.21E-40 | 1.41E-36 |
| WBGene00044644 | B0205.13 | 3.454820966 | 0.645738051 | 4.052649992 | 0.377 | 0.023 | 2.11E-35 | 1.35E-31 |
| WBGene00001753 | gst-5 | 3.656847145 | 0.672639034 | 4.305302243 | 0.523 | 0.075 | 5.88E-39 | 3.76E-35 |
